# Interspecies regulatory landscapes and elements revealed by novel joint systematic integration of human and mouse blood cell epigenomes

**DOI:** 10.1101/2023.04.02.535219

**Authors:** Guanjue Xiang, Xi He, Belinda M. Giardine, Kathryn J. Isaac, Dylan J. Taylor, Rajiv C. McCoy, Camden Jansen, Cheryl A. Keller, Alexander Q. Wixom, April Cockburn, Amber Miller, Qian Qi, Yanghua He, Yichao Li, Jens Lichtenberg, Elisabeth F. Heuston, Stacie M. Anderson, Jing Luan, Marit W. Vermunt, Feng Yue, Michael E.G. Sauria, Michael C. Schatz, James Taylor, Berthold Göttgens, Jim R. Hughes, Douglas R. Higgs, Mitchell J. Weiss, Yong Cheng, Gerd A. Blobel, David M. Bodine, Yu Zhang, Qunhua Li, Shaun Mahony, Ross C. Hardison

## Abstract

Knowledge of locations and activities of *cis*-regulatory elements (CREs) is needed to decipher basic mechanisms of gene regulation and to understand the impact of genetic variants on complex traits. Previous studies identified candidate CREs (cCREs) using epigenetic features in one species, making comparisons difficult between species. In contrast, we conducted an interspecies study defining epigenetic states and identifying cCREs in blood cell types to generate regulatory maps that are comparable between species, using integrative modeling of eight epigenetic features jointly in human and mouse in our **<u>V</u>**al**i**dated **<u>S</u>**ystematic **I**ntegrati**<u>on</u>** (VISION) Project. The resulting catalogs of cCREs are useful resources for further studies of gene regulation in blood cells, indicated by high overlap with known functional elements and strong enrichment for human genetic variants associated with blood cell phenotypes. The contribution of each epigenetic state in cCREs to gene regulation, inferred from a multivariate regression, was used to estimate epigenetic state Regulatory Potential (esRP) scores for each cCRE in each cell type, which were used to categorize dynamic changes in cCREs. Groups of cCREs displaying similar patterns of regulatory activity in human and mouse cell types, obtained by joint clustering on esRP scores, harbored distinctive transcription factor binding motifs that were similar between species. An interspecies comparison of cCREs revealed both conserved and species-specific patterns of epigenetic evolution. Finally, we showed that comparisons of the epigenetic landscape between species can reveal elements with similar roles in regulation, even in the absence of genomic sequence alignment.

## Introduction

The morphology and functions of different cell types are determined by the expression of distinctive sets of genes in each. This differential gene expression is regulated by the interplay of transcription factors (TFs) binding to *cis*-regulatory elements (CREs) in the genomic DNA, such as promoters and enhancers, forging interactions among the CREs and components of transcriptional apparatus and ultimately leading to patterns of gene activation and repression characteristic of each cell type (Maston et al. 2006; Hamamoto and Fukaya 2022). Epigenetic features such as accessibility of DNA and modifications of histone tails in chromatin have pronounced impacts on the ability of TFs to bind to CREs, and furthermore, they serve as a molecular memory of transcription and repression (Strahl and Allis 2000; Ringrose and Paro 2004). Frequently co-occurring sets of chromatin features define epigenetic states, which are associated with gene regulation and expression (Ernst and Kellis 2010; Hoffman et al. 2013; Zhang et al. 2016). Genome-wide assignment of DNA intervals to epigenetic states (annotation) provides a view of the regulatory landscape that can be compared across cell types, which in turn leads to insights into the processes regulating gene expression (Libbrecht et al. 2021).

Comprehensive mapping of CREs within the context of the regulatory landscape in different cell types is needed to achieve a broad understanding of differential gene expression. Maps of candidate CREs (cCREs) provide guidance in understanding how changes in cCREs, including single nucleotide variants and indels, can lead to altered expression (Hardison 2012), and they can inform approaches for activation or repression of specific genes in potential strategies for therapies (Bauer et al. 2013). Indeed, most human genetic variants associated with common traits and diseases are localized in or near cCREs (Hindorff et al. 2009; Maurano et al. 2012; The ENCODE Project Consortium 2012). Thus, knowledge of the activity and epigenetic state of cCREs in each cell type can facilitate understanding the impact of trait-associated genetic variants on specific phenotypes. Furthermore, genome editing approaches in somatic cells have recently been demonstrated to have promise as therapeutic modalities (Frangoul et al. 2021), and a full set of cCREs annotated by activity and state can help advance similar applications.

The different types of blood cells in humans and mice are particularly tractable systems for studying many aspects of gene regulation during differentiation. The striking differences among mature cell types result from progressive differentiation starting from a common hematopoietic stem cell (HSC) (Kondo et al. 2003). While single cell analyses reveal a pattern of ostensibly continuous expression change along each hematopoietic lineage (Laurenti and Göttgens 2018), intermediate populations of multi-lineage progenitor cells with decreasing differentiation potential have been defined, which provide an overall summary and nomenclature for major stages in differentiation. These stem, progenitor, and mature cell populations can be isolated using characteristic cell surface markers (Spangrude et al. 1988; Payne and Crooks 2002), albeit with many fewer cells in progenitor populations. In addition to the primary blood cells, several immortalized cell lines provide amenable systems for intensive study of various aspects of gene regulation during differentiation and maturation of blood cells (Weiss et al. 1997).

The VISION project aims to produce a **<u>V</u>**al**i**dated **<u>S</u>**ystematic **I**ntegrati**<u>on</u>** of hematopoietic epigenomes, harvesting extensive epigenetic and transcriptomic datasets from many investigators and large consortia into concise, systematically integrated summaries of regulatory landscapes and cCREs (Hardison et al. 2020). We previously published the results of these analyses for progenitor and mature blood cell types from mouse (Xiang et al. 2020). In the current study, we generated additional epigenetic datasets and compiled data from human blood cells to expand the integrative analyses to include data from both human and mouse. The systematic integrative analysis of epigenetic features across blood cell types was conducted jointly in both species to learn epigenetic states, generate concise views of epigenetic landscapes, and predict regulatory elements that are comparable in both species. This joint modeling enabled further comparisons using approaches that were not dependent on DNA sequence alignments between species, including a demonstration of the role of orthologous transcription factors in cell type-specific regulation in both species. An exploration of comparisons of epigenetic landscapes between species showed that they were informative for inferring regulatory roles of elements in lineage-specific (i.e., non-aligning) DNA. Together, this work provides valuable community resources that enable researchers to leverage the extensive existing epigenomic data into further mechanistic regulatory studies of both individual loci and genome-wide trends in human and mouse blood cells.

## Results

### Extracting and annotating epigenetic states by modeling epigenomic information jointly in human and mouse

A large number of data sets of epigenetic features related to gene regulation and expression (404 data sets, 216 in human and 188 in mouse; Fig. 1, Supplemental Material “Data generation and collection”, Supplemental Tables S1 and S2) served as the input for our joint integrative analysis of human and mouse regulatory landscapes across progenitor and mature blood cell types. The features included chromatin accessibility, which is a hallmark of almost all regulatory elements, occupancy by the structural protein CTCF, and histone modifications associated with gene activation or repression. After normalizing and denoising these diverse data sets (Supplemental Fig. S1), we conducted an iterative joint modeling to discover epigenetic states, i.e., sets of epigenetic features commonly found together, in a consistent manner for both human and mouse blood cells (Fig. 2). The joint modeling took advantage of the Bayesian framework of the Integrative and Discriminative Epigenomic Annotation System, or IDEAS (Zhang et al. 2016; Zhang and Hardison 2017), to iteratively learn states in both species. The joint modeling proceeded in four steps: initial training on randomly selected regions in both species, retaining the 27 epigenetic states that exhibit similar combinatorial patterns of features in both human and mouse, using these 27 states as prior information to sequentially run the IDEAS genome segmentation on the human and mouse data sets, and removal of two heterogenous states (Fig. 2A and Supplemental Figs. S2, S3, S4, and S5). This procedure ensured that the same set of epigenetic states was learned and applied for both species. Previously, the segmentation and genome annotation (Libbrecht et al. 2021) method ChromHMM (Ernst and Kellis 2012) was used to combine data between species by concatenating the datasets for both human and mouse cell types (Yue et al. 2014). This earlier approach produced common states between species, but it did not benefit from the positional information and automated approach to handling missing data that are embedded in IDEAS.

**Figure 1.**
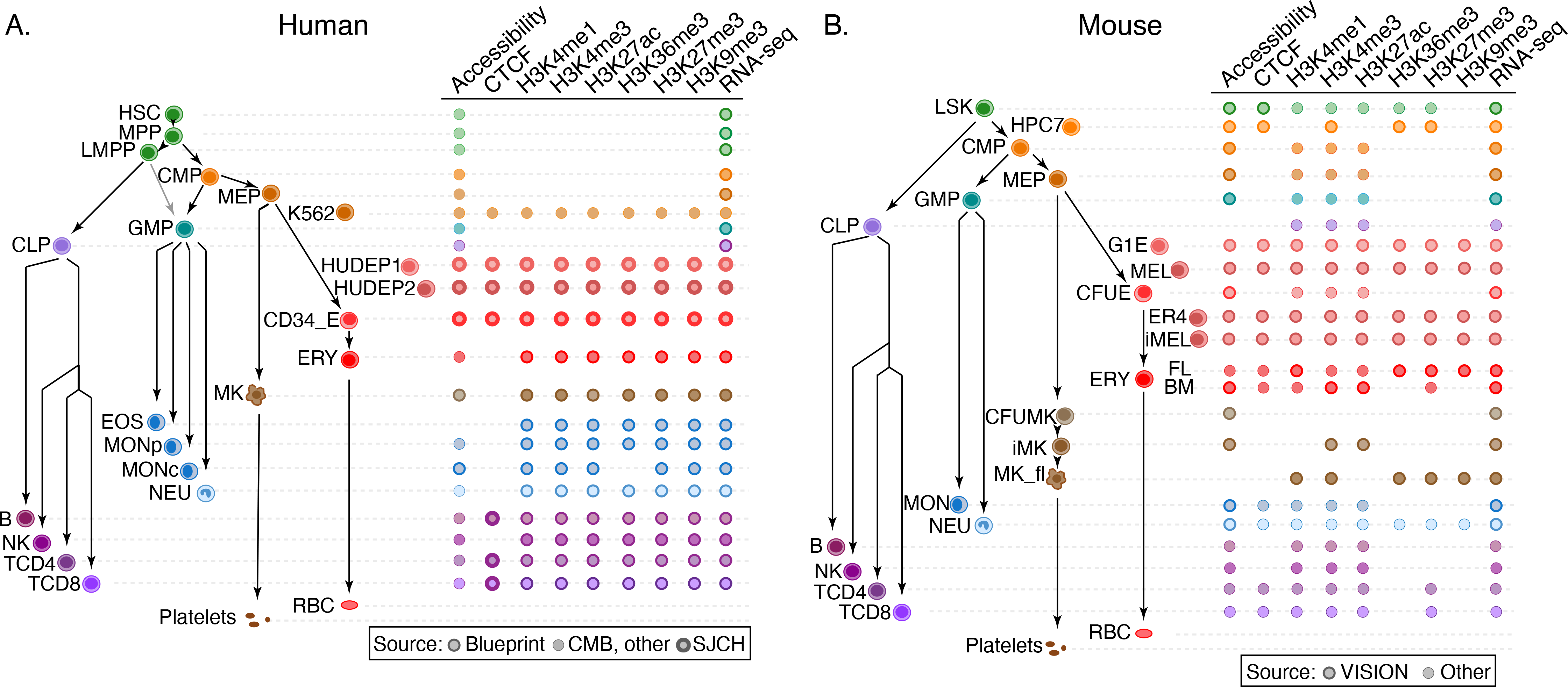
Cell types and data sets used for systematic integration of epigenetic features of blood cells. **(A)** The tree on the left shows the populations of stem, progenitor, and mature blood cells and cell lines in human. The diagram on the right indicates the epigenetic features and transcriptomes for which genome-wide data sets were generated or collected, with distinctive icons for the major sources of data, specifically the Blueprint project (Martens and Stunnenberg 2013; Stunnenberg et al. 2016), Corces et al. (2016), abbreviated CMB, and St. Jude Children’s Research Hospital (SJCRH, Cheng et al. 2021; Qi et al. 2021). (B) Cell types and epigenetic data sets in mouse, diagrammed as for panel A. Sources were described in Xiang et al. (2020) and Supplemental Table S1. Abbreviations for blood cells and lines are: HSC = hematopoietic stem cell, MPP = multipotent progenitor cell, LMPP = lymphoid-myeloid primed progenitor cell, CMP = common myeloid progenitor cell, MEP = megakaryocyte-erythrocyte progenitor cell, K562 = a human cancer cell line with some features of early megakaryocytic and erythroid cells, HUDEP = immortalized human umbilical cord blood-derived erythroid progenitor cell lines expressing fetal globin genes (HUDEP1) or adult globin genes (HUDEP2), CD34_E = human erythroid cells generated by differentiation from CD34+ blood cells, ERY = erythroblast, RBC = mature red blood cell, MK = megakaryocyte, GMP = granulocyte monocyte progenitor cell, EOS = eosinophil, MON = monocyte, MONp = primary monocyte, MONc = classical monocyte, NEU = neutrophil, CLP = common lymphoid progenitor cell, B = B cell, NK = natural killer cell, TCD4 = CD4+ T cell, TCD8 = CD8+ T cell, LSK = Lin-Sca1+Kit+ cells from mouse bone marrow containing hematopoietic stem and progenitor cells, HPC7 = immortalized mouse cell line capable of differentiation in vitro into more mature myeloid cells, G1E = immortalized mouse cell line blocked in erythroid maturation by a knockout of the *Gata1* gene and its subline ER4 that will further differentiate after restoration of *Gata1* function in an estrogen inducible manner (Weiss et al. 1997), MEL = murine erythroleukemia cell line that can undergo further maturation upon induction (designated iMEL), CFUE = colony forming unit erythroid, FL = designates ERY derived from fetal liver, BM = designates ERY derived from adult bone marrow, CFUMK = colony forming unit megakaryocyte, iMK = immature megakaryocyte, MK_fl = megakaryocyte derived from fetal liver.

**Figure. 2.**
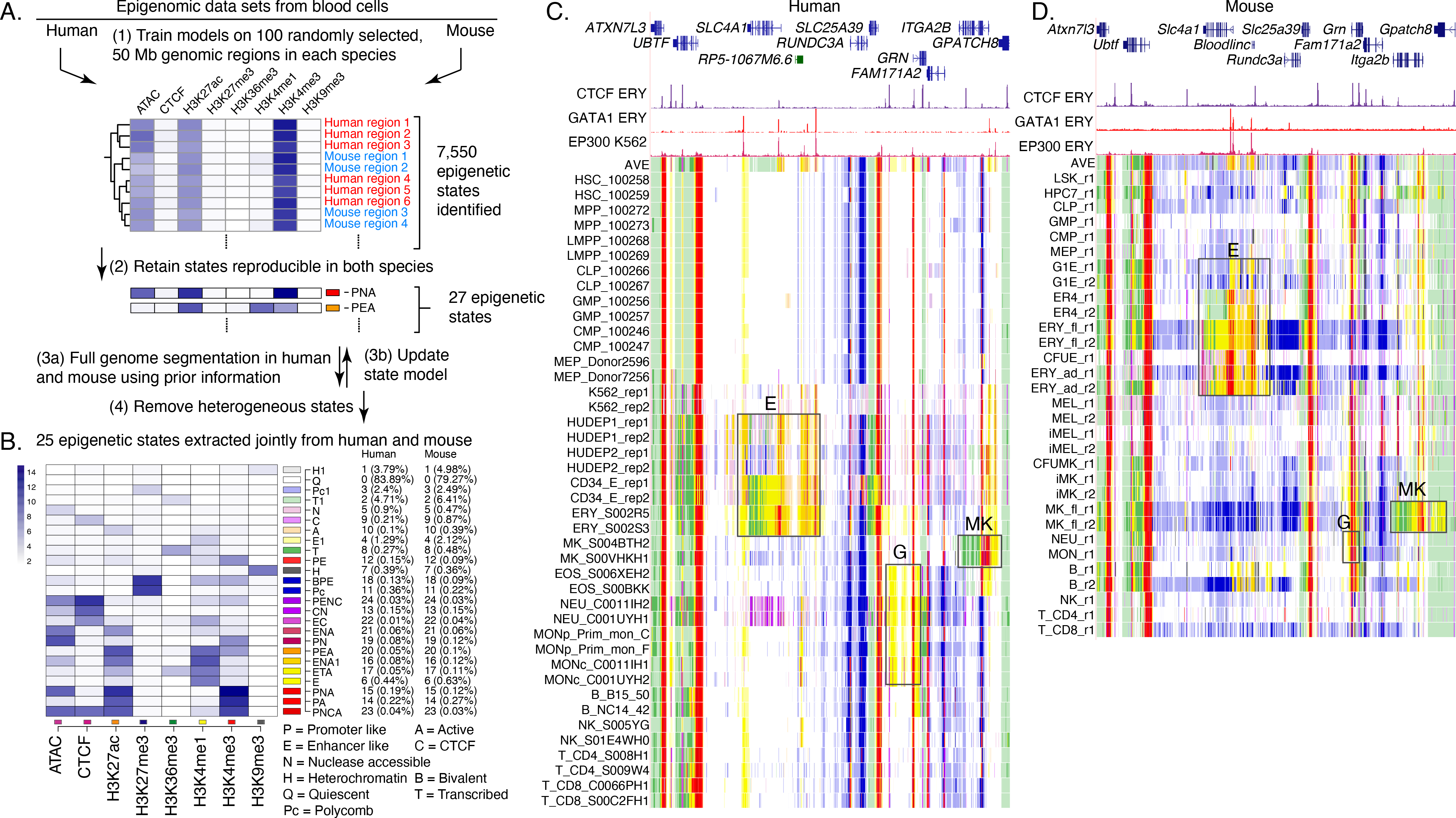
Genome segmentation and annotation jointly between human and mouse using IDEAS. **(A)** Workflow for joint modeling. (1) Initial epigenetic states from 100 randomly selected regions separately in human and mouse hematopoietic cell types were identified in IDEAS runs. (2) States that were reproducible and shared in both species were retained. (3a and 3b) The profile of epigenetic feature contribution to each of the reproducible states was sequentially refined by applying IDEAS across the full genomes of human and of mouse, updating the state model after each IDEAS run. (4) Two heterogeneous states were removed to generate the final joint epigenetic states in the two species. **(B)** The 25 joint epigenetic states for human and mouse hematopoietic cell types. The average signal of the epigenetic features for each state are shown in the heatmap. The corresponding state colors, the state labels based on the function, and the average proportions of the genome covered by each state across cell types are listed on the right-side of the heatmap. **(C)** Annotation of epigenetic states in a large genomic interval containing *SLC4A1* and surrounding genes across human blood cell types. The genomic interval is 210kb, GRCh38 Chr17:44,192,001-44,402,000, with gene annotations from GENCODE V38. Binding patterns for selected transcription factors are from the VISION project ChIP-seq tracks (CTCF and GATA1 in adult erythroblasts, signal tracks from MACS, track heights 100 and 80, respectively) or from the ENCODE data portal (EP300 in K562 cells, experiment ENCSR000EGE, signal track is fold change over background, track height is 50). The epigenetic state assigned to each genomic bin in the different cell types is designated by the color coding shown in panel (B). The replicates in each cell type examined in Blueprint are labeled by the id for the donor of biosamples. Genes and regulatory regions active primarily in erythroid (E), granulocytes (G), and megakaryocytes (MK) are marked by gray rectangles. **(D)** Annotation of epigenetic states in a large genomic interval containing *Slc4a1* and surrounding genes across mouse blood cell types. The genomic interval is 198kb, mm10 Chr11:102,290,001-102,488,000, with gene annotations from GENCODE VM23. Binding patterns for selected transcription factors are from the VISION project ChIP-seq tracks (CTCF in adult erythroblasts, GATA1 and EP300 from the highly erythroid fetal liver, signal tracks from MACS, track heights 200, 200, and 150, respectively; the EP300 track was made by re-mapping reads from ENCODE experiment ENCSR982LJQ). The tracks of epigenetic states and highlighted regions are indicated as in panel (C).

The resulting model with 25 epigenetic states (Fig. 2B) was similar to that obtained from mouse blood cell data (Xiang et al. 2020). The states captured combinations of epigenetic features characteristic of regulatory elements such as promoters and enhancers, transcribed regions, repressed regions marked by either Polycomb (H3K27me3) or heterochromatin (H3K9me3), including states that differ quantitatively in the contribution of specific features to each state. For example, H3K4me1 is the predominant component of states E1 and E, but E1 has a lower contribution of that histone modification. Similar proportions of the genomes of human and mouse were covered by each state (Fig. 2B).

Assigning all genomic bins in human and mouse to one of the 25 states in each hematopoietic cell type produced an annotation of blood cell epigenomes that gave a concise view of the epigenetic landscape and how it changes across cell types, using labels and color conventions consistently for human and mouse. The value of this concise view can be illustrated in orthologous genomic intervals containing genes expressed preferentially in different cell lineages as well as genes that are uniformly expressed (Fig. 2C, D). For example, the gene *SLC4A1*/*Slc4a1*, encoding the anion transporter in the erythrocyte plasma membrane, is expressed in the later stages of erythroid maturation (Dore and Crispino 2011). The epigenetic state assignments across cell types matched the differential expression pattern, with genomic intervals in the gene and its flanking regions, including a non-coding gene located upstream (to its right, *Bloodlinc* in mouse), assigned to states indicative of enhancers (yellow and orange) and promoters (red) only in erythroid cell types, with indications of stronger activation in the more mature erythroblasts (region boxed and labeled E in Fig. 2 C, D). A similar pattern was obtained in both human and mouse. Those genomic intervals assigned to the enhancer- or promoter-like states contain candidates for regulatory elements, an inference that was supported by chromatin binding data including occupancy by the transcription factor GATA1 (Xu et al. 2012; Pimkin et al. 2014) and the co-activator EP300 (ENCODE datasets ENCSR000EGE and ENCSR982LJQ) in erythroid cells. Similarly, the gene and flanking regions for *GRN*/*Grn*, encoding the granulin precursor protein that is produced at high levels in granulocytes and monocytes (Jian et al. 2013), and *ITGA2B*/*Itga2b*, encoding the alpha 2b subunit of integrin that is abundant in mature megakaryocytes (van Pampus et al. 1992; Pimkin et al. 2014), were assigned to epigenetic states indicative of enhancers and promoters in the expressing cell types (boxed regions labeled G and MK, respectively). In contrast, genes expressed in all the blood cell types, such as *UBTF*/*Ubtf*, were assigned to active promoter states and transcribed states across the cell types. We conclude that these concise summaries of the epigenetic landscapes across cell types showed the chromatin signatures for differential or uniform gene expression and revealed discrete intervals as potential regulatory elements, with the consistent state assignments often revealing similar epigenetic landscapes of orthologous genes in human and mouse.

While these resources are useful, some limitations should be kept in mind. For example, IDEAS used data from similar cell types to improve state assignments in cell types with missing data, but the effectiveness of this approach may be impacted by the pattern of missing data. In particular, the epigenetic data on human stem and progenitor cell types were largely limited to ATAC-seq data, whereas histone modification data and CTCF occupancy were available for the analogous cell types in mouse (Fig. 1). Thus, the state assignments for epigenomes in human stem and progenitor cells may be less robust compared to those for similar cell types in mouse. Another limitation is the broad range of quality in the data sets that cannot be completely adjusted by normalization, which leads to over- or under-representation of some epigenetic signals in specific cell types (Supplemental Fig. S5). Despite these limitations, the annotation of blood cell epigenomes after normalization and joint modeling of epigenetic states produced a highly informative painting of the activity and regulatory landscapes across the genomes of human and mouse blood cells.

### Candidate *cis*-regulatory elements in human and mouse

We define a candidate *cis*-regulatory element, or cCRE, as a DNA interval with a high signal for chromatin accessibility in any cell type (Xiang et al. 2020). We utilized a version of the IDEAS methodology to combine peaks of accessibility across different cell types, running it in the signal intensity state (IS) mode only on chromatin accessibility signals (Xiang et al. 2021), which helps counteract excessive expansion of peak calls when combining them (Supplemental Fig. S6).

Employing the same peak-calling procedure to data from human and mouse resulted in 200,342 peaks of chromatin accessibility for human and 96,084 peaks for mouse blood cell types (Supplemental Table S3). Applying the peak caller MACS3 (Zhang et al. 2008) on the same human ATAC-seq data generated a larger number of peaks, but those additional peaks tended to have low signal and less enrichment for overlap with other function-related genomic datasets (Supplemental Fig. S7).

The ENCODE Project released regulatory element predictions in a broad spectrum of cell types in the Index of DHSs (Meuleman et al. 2020) and the SCREEN cCRE catalog (The ENCODE Project Consortium et al. 2020), using data that were largely different from those utilized for the VISION analyses. Almost all the VISION cCRE calls in human blood cells were included in the regulatory element predictions from ENCODE (Supplemental Fig. S8A), supporting the quality of the VISION cCRE calls. Furthermore, as expected from its focus on blood cell types, the VISION cCRE catalog shows stronger enrichment for regulatory elements active in blood cells (Supplemental Fig. S8B, Supplemental Table S4).

### Enrichment of the cCRE catalog for function-related elements and trait-associated genetic variants

Having generated catalogs of cCREs along with an assignment of their epigenetic states in each cell type, we characterized the human cCREs further by connecting them to orthogonal (not included in VISION predictions) datasets of DNA elements implicated in gene regulation or in chromatin structure and architecture (termed structure-related) (Fig. 3A, Supplemental Fig. S9, Supplemental Table S5). About two-thirds (136,664 or 68%) of the VISION human cCREs overlapped with elements in the broad groups of CRE-related (97,361 cCREs overlapped) and structure-related (83,327 cCREs overlapped) elements, with 44,024 cCREs overlapping elements in both categories (Fig. 3A, B). In contrast, ten sets of randomly chosen DNA intervals, matched in length and GC-content with the human cCRE list, showed much less overlap with the orthogonal sets of elements (Fig. 3B). Of the CRE-related superset, the enhancer-related group of datasets contributed the most overlap with VISION cCREs, followed by SuRE peaks, which measure promoter activity in a massively parallel reporter assay (van Arensbergen et al. 2017), and CpG islands (Fig. 3C). Compared to overlaps with the random matched intervals, the VISION cCREs were highly enriched for overlap with each group of CRE-related datasets (Fig. 3C). Of the structure-related superset, the set of CTCF occupied segments (OSs) contributed the most overlap, followed by chromatin loop anchors, again with high enrichment relative to overlaps with random matched sets (Fig. 3D). Considering the VISION cCREs that intersected with both structure- and CRE-related elements, major contributors were the cCREs that overlap with enhancers and CTCF OSs or loop anchors (Supplemental Fig. S10). Furthermore, the VISION cCREs captured known blood cell CREs (Supplemental Table S4) and CREs demonstrated to impact a specific target gene in a high throughput analysis (Gasperini et al. 2019) (Fig. 3E). We conclude that the intersections with orthogonal, function- or structure-related elements lent strong support for the biological significance of the VISION cCRE calls and added to the annotation of potential functions for each cCRE.

**Figure. 3.**
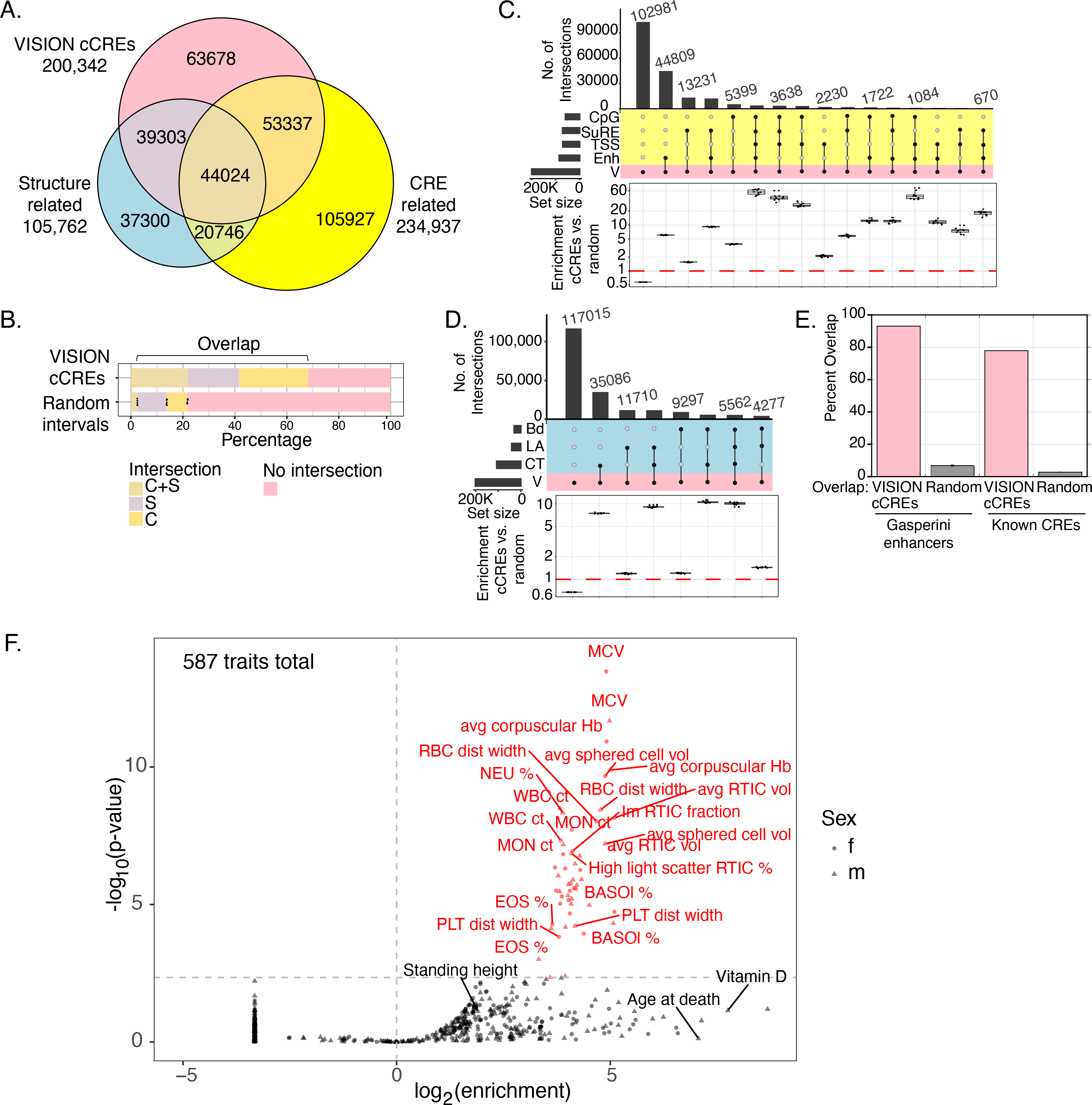
Overlaps of VISION cCREs with other catalogs and enrichment for variants associated with blood cell traits. **(A)** Venn diagram showing intersections of human VISION cCREs with a combined superset of elements associated with nuclear structure (CTCF OSs, loop anchors, and TAD boundaries) and with a combined superset of DNA intervals associated with *cis*-regulatory elements (CREs), including TSSs, CpG islands, peaks from a massively parallel promoter and enhancer assay, and enhancers predicted from enhancer RNAs, peaks of binding by EP300, and histone modifications in erythroblasts (see Supplemental Material, Supplemental Fig. S9, and Supplemental Table S5). **(B)** The proportions of cCREs and randomly selected, matched sets of intervals in the overlap categories are compared in the bar graph. For the random sets, the bar shows the mean, and the dots show the values for each of ten random sets. **(C)** The UpSet plot provides a higher resolution view of intersections of VISION cCREs with the four groups of CRE-related elements, specifically enhancer-related (Enh), transcription start sites (TSS), Survey of Regulatory Elements (SuRE), and CpG islands (CpG). The enrichment for the cCRE overlaps compared to those in randomly selected, matched sets of intervals are shown in the boxplots below each overlap subset, with dots for the enrichment relative to individual random sets. **(D)** Overlaps and enrichments of VISION cCREs for three sets of structure-related elements, specifically CTCF OSs (CT), loop anchors (LA), and TAD boundary elements. **(E)** Overlaps of VISION cCREs with two sets of experimentally determined blood cell cCREs. **(F)** Enrichment of SNPs associated with blood cell traits from UK Biobank in VISION cCREs. Results of the sLDSC analysis of all cCREs are plotted with enrichment of the cCRE annotation in heritability of each trait on the x-axis, and the significance of the enrichment on the y-axis. The analysis covers 292 unique traits with GWAS results from both males and females and 3 traits with results only from males. The vertical dotted line indicates an enrichment of 1, and the horizontal dotted line delineates the 5% FDR significance threshold. Points and labels in red represent traits for which there was significant enrichment of SNPs associated with the VISION cCREs. Traits with a negative enrichment were assigned an arbitrary enrichment of 0.1 for plotting and appear as the column of points at the bottom left of the plot. The shape of the point indicates the sex in which the GWAS analysis was performed for each trait.

The catalog of VISION human blood cell cCREs showed a remarkable enrichment for genetic variants associated with blood cell traits, further supporting the utility of the catalog. We initially observed a strong enrichment by overlap with variants from the NHGRI-EBI GWAS Catalog (Buniello et al. 2019) associated with blood cell traits (Supplemental Fig. S11). We then analyzed the enrichments while considering the haplotype structure of human genomes, whereby association signals measured at assayed genetic markers likely reflect an indirect effect driven by linkage disequilibrium (LD) with a causal variant (that may or may not have been genotyped). We employed stratified linkage disequilibrium score regression (sLDSC, Finucane et al. 2015) to account for LD structure and estimate the proportion of heritability of each trait explained by a given genomic annotation, quantifying the enrichment of heritability in 587 traits from the UK Biobank (UKBB) GWAS (Ge et al. 2017 and http://www.nealelab.is/uk-biobank/) within the VISION cCREs relative to the rest of the genome (Supplemental Material section “Stratified linkage disequilibrium score regression”). These traits encompassed 54 “blood count” traits that measure properties including size and counts of specific blood cell types, 60 “blood biochemistry” traits that measure lipid, enzyme, and other molecular concentrations within whole blood samples, and 473 non-blood-related traits, allowing us to assess the specific relevance of the cCREs to regulation of blood-related versus other phenotypes. At a 5% FDR threshold, we discovered 53 traits for which cCREs were significantly enriched in heritability (Fig. 3F). Of these traits, 52 (98%) were blood-related and 50 were blood count traits, representing 93% of all UKBB blood count traits included in our analysis. The remaining 2 significant traits pertained to blood biochemistry, specifically, the male and female glycated hemoglobin concentrations. These metrics and observations together lend support to the VISION cCRE annotation being composed of informative genomic regions associated with regulation of genes involved in development of blood cell traits.

### Estimates of regulatory impact of cCREs during differentiation

The epigenetic states assigned to cCREs can reveal those that show changes in apparent activity during differentiation. Inferences about the activity of a cCRE in one or more cell types are based on whether the cCRE was actuated, i.e., was found in a peak of chromatin accessibility, and which epigenetic state was assigned to the actuated cCRE. Those states can be associated with activation (e.g., enhancer-like or promoter-like) or repression (e.g., associated with polycomb or heterochromatin). In addition to these categorical state assignments, quantitative estimates of the impact of epigenetic states on expression of target genes are useful, e.g., to provide an estimate of differences in inferred activity when the states change. Previous work used signals from single or multiple individual features such as chromatin accessibility or histone modifications in regression modeling to explain gene expression (e.g., Karlić et al. 2010; Dong et al. 2012), and we applied a similar regression modeling using epigenetic states as predictor variables to infer estimates of regulatory impact of each state on gene expression (Xiang et al. 2020).

We used state assignments of cCREs across cell types in a multivariate regression model to estimate the impact of each state on the expression of local genes (Supplemental Material, “Estimation of the impact of epigenetic states and cCREs on gene expression”). That impact was captured as β coefficients, which showed the expected strong positive impact for promoter and enhancer associated states and negative impacts from heterochromatin and polycomb states (Fig. 4A). The β coefficients were then used in further analysis, such as estimating the change in regulatory impact as a cCRE shifts between states during differentiation (difference matrix to the left of the β coefficient values in Fig. 4A). The β coefficient values also were used to generate an epigenetic state Regulatory Potential (esRP) score for each cCRE in each cell type, calculated as the β coefficient values for the epigenetic states assigned to the cCRE weighted by the coverage of the cCRE by each state (Fig. 4B). These esRP scores were the basis for visualizing the collection of cCREs and how their regulatory impact changed across differentiation (Supplemental Fig. S12 and Supplemental movie S1). Comparison of the integrative esRP scores with signal intensities for single features (ATAC-seq and H3K27ac) showed all were informative for visualizations, and esRP performed slightly better than the single features in differentiating cCREs based on locations within gene bodies (Supplemental Fig. S13).

**Figure 4.**
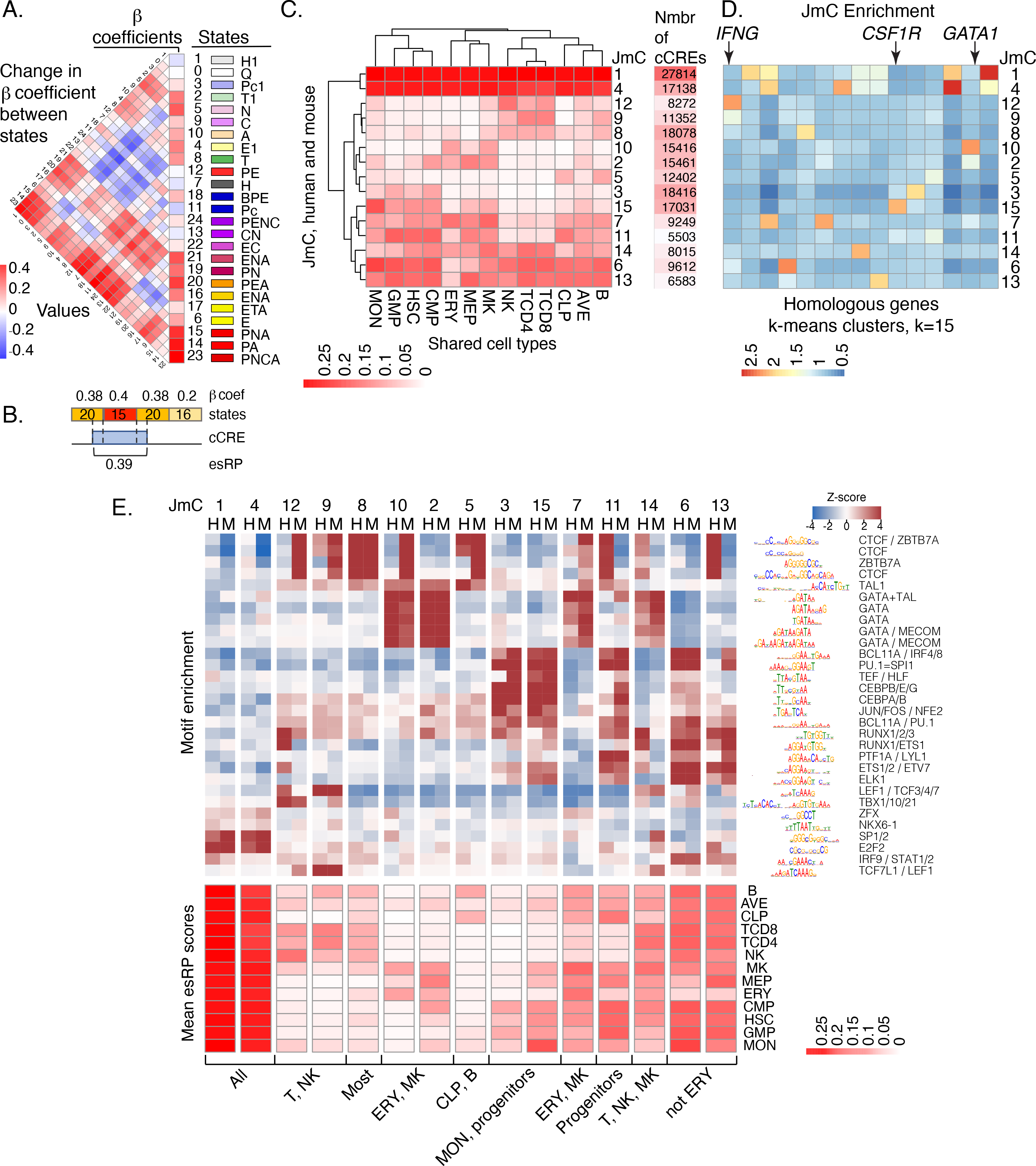
Beta coefficients of states, esRP scores of cCREs, joint human-mouse metaclusters of cCREs based on esRP scores, and enrichment for TFBS motifs. **(A)** Beta coefficients and the difference of beta coefficients of the 25 epigenetic states. The vertical columns on the right show the beta coefficients along with the ID, color, and labels for the 25 joint epigenetic states. The triangular heatmap shows the difference of the beta coefficients between two states in the right columns. Each value in the triangle heatmap shows the difference in beta coefficients between the state on top and the state below based on the order of states in the right columns. **(B)** An example of calculating esRP score for a cCRE in a cell type based on the beta coefficients of states. For a cCRE covering more than one 200bp bin, the esRP equals the weighted sum of beta coefficients of states that covers the cCRE, where the weights are the region covered by different states. **(C)** The average esRP score of all cCREs in JmCs across blood cell types shared by human and mouse. The right column shows the number of human cCREs in each JmC. **(D)** The average enrichment of JmCs in 15 homologous gene clusters. The genes are clustered based on the JmCs’ enrichments by *k*-means. **(E)** Motifs enriched in joint metaclusters. The top heatmap shows the enrichment of motifs in the cCREs in each JmC in human (H) and mouse (M) as a *Z*-score. The logo for each motif is given to the right of the heat map, labeled by the family of transcription factors that recognize that motif. The heatmap below is aligned with the motif enrichment heatmap, showing the mean esRP score for the cCREs in each JmC for all the common cell types examined between human and mouse. A summary description of the cell types in which the cCREs in each JmC are more active is given at the bottom.

In addition, we explored the utility of the esRP scores for clustering the cCREs into groups with similar activity profiles across blood cell types in both human and mouse. Focusing on the esRP scores in 12 cell types shared between human and mouse along with the average across cell types, we identified clusters jointly in both species. The clustering proceeded in three steps, specifically finding robust *k*-means clusters for the combined human and mouse cCREs, identifying the clusters shared by cCREs in both species, and then further grouping those shared *k*-means clusters hierarchically to define fifteen joint metaclusters (JmCs) (Supplemental Fig. S14). Each cCRE in both mouse and human was assigned to one of the fifteen JmCs, and each JmC was populated with cCREs from both mouse and human.

These JmCs established discrete categories for the cCREs based on the cell type distribution of their inferred regulatory impact (Fig. 4C). The clusters of cCREs with high esRP scores across cell types were highly enriched for promoter elements (Supplemental Fig. S15A). The cell type-restricted clusters of cCREs showed enrichment both for selected enhancer catalogs and for functional terms associated with those cell types (Supplemental Fig. S15A and B). Furthermore, clustering of human genes by the JmC assignments of cCREs in a 100kb interval centered on their TSS (Supplemental Material section “Enrichment of JmCs assigned to cCREs in gene loci”) revealed a strong enrichment for JmCs with high activity in the cell type(s) in which the genes are expressed (Fig. 4D). Examples include *IFNG* showing enrichment for JmC 12, which has high esRP scores in T and NK cells, *CSF1R* showing enrichment for JmC 15, which has high scores in monocytes, and *GATA1* showing enrichment for JmC 10, which has high scores in erythroid cells and megakaryocytes. Moreover, running sLDSC on cCREs in individual JmCs showed enrichment for heritability of blood cell-related traits in some specific JmCs (Supplemental Fig. S16).

As expected from previous work (e.g., Heintzman et al. 2009; Meuleman et al. 2020), similar metaclusters of cCREs were generated based on single signals from the histone modification H3K27ac or chromatin accessibility across cell types (Supplemental Fig. S17). Clustering based any of the three features better resolved individual cell types when larger numbers of clusters were considered, prior to collapsing the shared robust clusters into JmCs (Supplemental Fig. S18).

In summary, we show that the β coefficients and esRP scores provide valuable estimates of regulatory impacts of states and cCREs, respectively. The esRP-driven joint metaclusters provide refined subsets of cCREs that should be informative for investigating cell type-specific and general functions of cCREs. We also built self-organizing maps as a complementary approach to systematic integration of epigenetic features and RNA data across cell types (Supplementary Figure S10, Jansen et al. 2019).

### Motif enrichment in joint metaclusters of human and mouse cCREs

We examined the sets of cCREs in each JmC to ascertain enrichment for transcription factor binding site (TFBS) motifs because these enriched motifs suggest the families of transcription factors that play a major role in regulation by each category of cCREs. Furthermore, having sets of cCREs determined and clustered for comparable blood cell types in human and mouse provided the opportunity to discover which TFBS motifs were shared between species and whether any were predominant in only one species.

To find TFBS motifs associated with each JmC, we calculated enrichment for all non-redundant motifs in the Cis-BP database (Weirauch et al. 2014) using Maelstrom from GimmeMotifs (Bruse and van Heeringen 2018) (Supplemental Material “Enrichment for transcription factor binding site motifs in joint metaclusters of cCREs”). The results confirmed previously established roles of specific TFs in cell lineages and showed little evidence for novel motifs (Fig. 4E). For example, TFBS motifs for the GATA family of transcription factors were enriched in JmCs 2 and 10, which have high esRP scores in progenitor and mature cells in the erythroid and megakaryocytic lineages, as expected for the known roles of GATA1 and GATA2 in this lineage (Blobel and Weiss 2009; Fujiwara et al. 2009). The GATA motif was also enriched in JmC 14, as expected for the role of GATA3 in natural killer (NK) and T cells (Rothenberg and Taghon 2005). Furthermore, motifs for the known lymphoid transcription factors TBX21, TCF7L1, and LEF1 (Chi et al. 2009) were enriched in cCREs with high esRP scores in NK and T cells (JmCs 9 and 12), and motifs for myeloid-determining transcription factors CEBPA and CEBPB (Graf and Enver 2009) and the myeloid transcription factor PU.1 (Tenen et al. 1997) were enriched in cCREs that are active in progenitor cells and monocytes (JmCs 3 and 15). TFBS motifs for promoter-associated transcription factors such as E2F2 and SP1 (Dynan and Tjian 1983; Kaczynski et al. 2003) were enriched in broadly active cCREs (JmCs 1 and 4). These patterns of motif enrichments in the JmCs fit well with the expectations from previous studies of transcription factor activity across lineages of blood cells, and thus, they lend further credence to the value of the cCRE calls and the JmC groupings for further studies of regulation in the blood cell types.

The genome-wide collection of cCREs across many blood cell types in human and mouse provided an opportunity for an unbiased and large-scale search for indications of transcription factors that may be active specifically in one species for a shared cell type. Prior studies of transcription factors have shown homologous transcription factors used in analogous cell types across species (e.g., Carroll 2008; Noyes et al. 2008; Schmidt et al. 2010; Cheng et al. 2014; Villar et al. 2014), but it is not clear if there are significant exceptions. In our study, we found that for the most part, the motif enrichments were quite similar between the human and mouse cCREs in each JmC. Note that these similarities were not forced by requiring sequence matches between species; the cCREs were grouped into JmCs based on their pattern of activity, as reflected in the esRP scores, across cell types, not by requiring homologous sequences. This similarity between species indicates that the same transcription factors tend to be active in similar groups of cell types in both mouse and human. An intriguing potential exception to the sharing of motifs between species was the enrichment of TFBS motifs for CTCF and ZBTB7A in some JmCs, suggestive of some species selectivity in their binding in the context of other TFs (Supplemental Figs. S20 and S21). These indications of conditional, preferential usage of these TFs in human or mouse could serve as the basis for more detailed studies in the future.

In summary, after grouping the cCREs in both human and mouse by their inferred regulatory impact across blood cell in a manner agnostic to DNA sequence or occupancy by TFs, the enrichment for TFBS motifs within those groups recapitulated known activities of TFs both broadly and in specific cell lineages. The results also showed considerable sharing of inferred TF activity in both human and mouse.

### Evolution of sequence and inferred function of cCREs

The human and mouse cCREs from blood cells were assigned to three distinct evolutionary categories (Fig. 5A). About one-third of the cCREs were present only in the reference species (39% for human, 28% for mouse), as inferred from the failure to find a matching orthologous sequence in whole-genome alignments with the other species. We refer to these as nonconserved (N) cCREs. Of the two-thirds of cCREs with an orthologous sequence in the second species, slightly over 30,000 were also identified as cCREs in the second species. The latter cCREs comprise the set of cCREs conserved in both sequence and inferred function, which we call SF conserved (SF) cCREs. Almost the same number of cCREs in both species fall into the SF category; the small difference resulted from interval splits during the search for orthologous sequences (Supplemental Fig. S22). The degree of chromatin accessibility in orthologous SF cCREs was positively correlated between the two species (Supplemental Fig. S23). The remaining cCREs (91,000 in human and 36,000 in mouse) were conserved in sequence but not in an inferred function as a regulatory element, and we call them S conserved (S) cCREs. The latter group could result from turnover of regulatory motifs or acquisition of different functions in the second species.

**Figure. 5.**
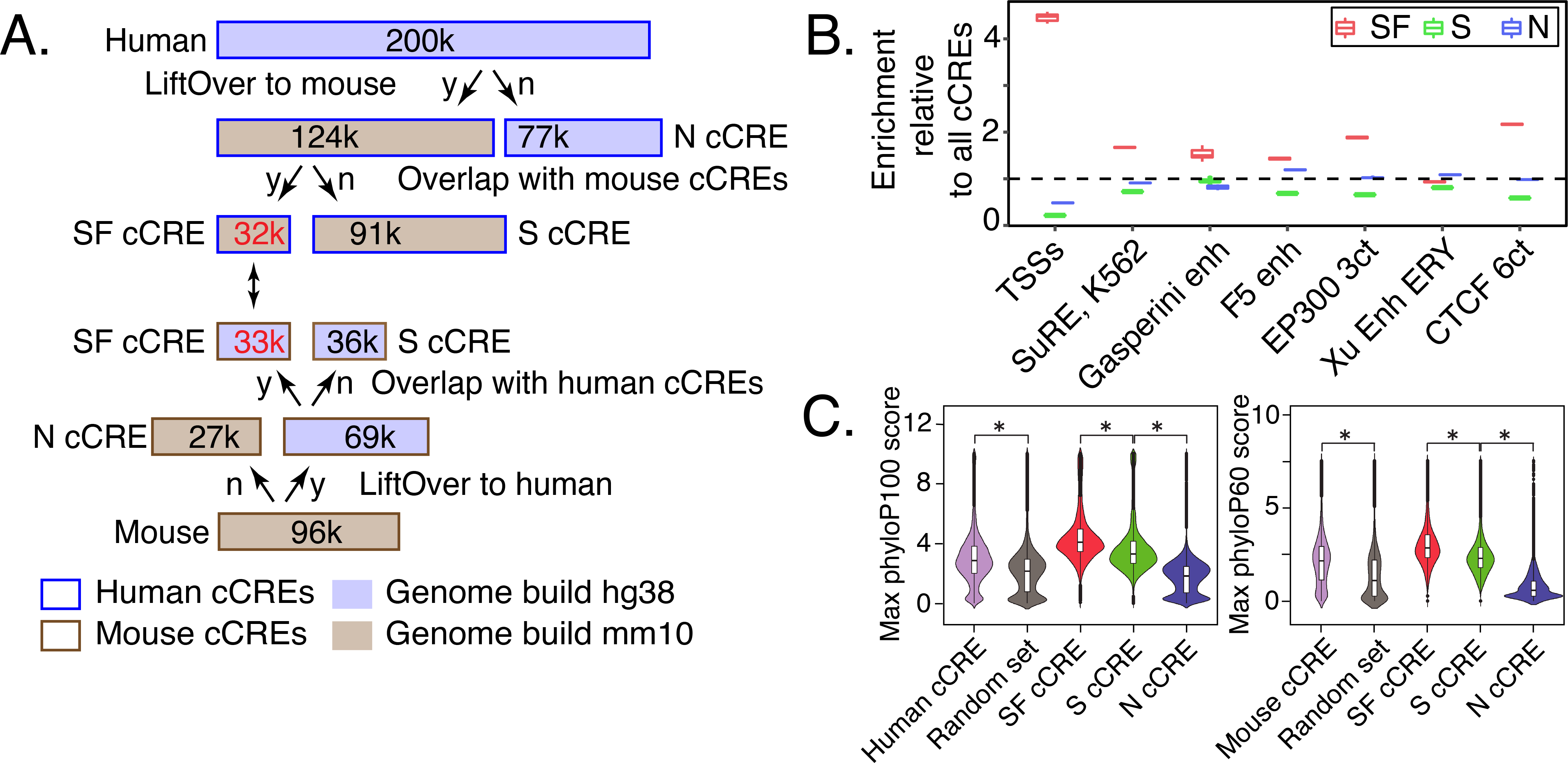
Evolutionary and epigenetic comparisons of cCREs. **(A)** Workflow to partition blood cell cCREs in human and mouse into three evolutionary categories. N=nonconserved, S=conserved in sequence but not inferred function, SF=conserved in both sequence and inferred function as a cCRE, y=yes, n=no. **(B)** Enrichment of SF-conserved human cCREs for TSSs. The number of elements in seven sets of function-related DNA intervals that overlap with the 32,422 SF human cCREs was determined, along with the number that overlap with three subsets (32,422 each) randomly selected from the full set of 200,342 human cCREs. The ratio of the number of function-related elements overlapping SF-cCREs to the number overlapping a randomly chosen subset of all cCREs gave the estimate of enrichment plotted in the graph. The mean for the three determinations of enrichment is indicated by the horizontal line for each set. Results are also shown for a similar analysis for the S and N cCREs. **(C)** Distribution of phyloP scores for three evolutionary categories of cCREs in human and mouse. The maximum phyloP score for each genomic interval was used to represent the score for each cCRE, using genome sequence alignments of 100 species with human as the reference (phyloP100) and alignments of 60 species with mouse as the reference (phyloP60). The distribution of phyloP scores for each group are displayed as a violin plot. All ten random sets had distributions similar to the one shown. The asterisk (*) over brackets indicates comparison for which the P values for Welch’s *t*-test is less than 2.2×10^−16^. **(D)** Proportion of human genomic elements active in a massively parallel reporter assay (MPRA) that align with mouse or are in a state reflecting dynamic chromatin. A set of 57,061 genomic elements found to be active in a lentivirus MPRA that tested a close to comprehensive set of predicted regulatory elements in K562 cells (Agarwal et al. 2023) were assessed for their ability to align with the mouse genome (blue bar) or whether the IDEAS epigenetic state assigned in K562 cells was not quiescent or was in a set of states associated with gene activation (magenta bars). The results are plotted as percentages of the total number of MPRA-active elements.

The distributions of epigenetic states assigned to the blood cell cCREs in each of the three evolutionary categories were similar between human and mouse, but those distributions differed among evolutionary categories, with significantly more SF cCREs assigned to promoter-like states than were S or N cCREs (Supplemental Fig. S24). Indeed, the SF cCREs tended to be close to or encompass the TSSs of genes, showing a substantial enrichment in overlap with TSSs compared to the overlap observed for all cCREs (Fig. 5B). Many of the S and N cCREs were assigned to enhancer-like states (Supplemental Fig. S24D), giving a level of enrichment for overlap with enhancer datasets comparable to that observed for the full set of cCREs (Fig. 5B).

For both human and mouse, the level of sequence conservation, estimated by the maximum phyloP score (Pollard et al. 2010), was higher in the collection of cCREs than in sets of randomly chosen genomic intervals matching the cCREs in length and G+C content (Fig. 5C). Among the evolutionary categories of cCREs, the distribution of phyloP scores for SF cCREs was significantly higher than the distribution for S cCREs, which in turn was higher than that for N cCREs, for both species (Fig. 5C). The whole genome alignments underlying the phyloP scores are influenced by proximity to the highly conserved coding exons (King et al. 2007), and the high phyloP scores of the promoter-enriched SF cCREs could reflect both this effect as well as strong constraint on conserved function (Supplemental Fig. S25). In all three evolutionary categories, the distribution of phyloP scores was higher for promoter-proximal cCREs than for distal ones, but the relative levels of inferred conservation were the same for both, i.e., SF>S>N (Supplemental Fig. S26).

In summary, this partitioning of the cCRE catalogs by conservation of sequence and inferred function revealed informative categories that differed both in evolutionary trajectories and in types of functional enrichment.

Conservation of non-coding genomic DNA sequences among species has been used extensively to predict regulatory elements (Gumucio et al. 1992; Hardison 2000; Pennacchio and Rubin 2001), but the observation that predicted regulatory elements fall into distinct evolutionary categories (SF, S, and N) raised the question of whether inter-species DNA sequence alignments or annotation of epigenetic states would be more effective in finding elements that were experimentally determined to be active in gene regulation. Recent advances in massively parallel reporter assays have enabled the testing of large sets of candidate elements, approaching comprehensive assessment of the predicted elements (Agarwal et al. 2023). We used the set of over 57,000 human genomic elements shown to be active in K562 cells to address this question (Supplemental Material), and we found that requiring alignment to the mouse genome would miss about 40% of the active elements, whereas requiring presence in a non-quiescent epigenetic state or one associated with gene activation would cover 87% or 82.5%, respectively, of the active elements (Fig. 5D). Thus, the epigenetic state annotation can enable a more comprehensive prediction and examination or gene regulatory elements. This realization motivated a comparison of epigenetic states between human and mouse, as described in the next section.

### Comparison of epigenetic states around orthologous genes in human and mouse

The consistent state assignments from the joint modeling facilitated epigenetic comparisons between species. Such comparisons are particularly informative for orthologous genes with similar expression patterns but some differences in their regulatory landscapes. For example, the orthologous genes *GATA1* in human and *Gata1* in mouse each encode a transcription factor with a major role in regulating gene expression in erythroid cells, megakaryocytes, and eosinophils (Ferreira et al. 2005), with a similar pattern of gene expression across blood cell types in both species (Supplemental Fig. S27). The human and mouse genomic DNA sequences aligned around these orthologous genes, including their promoters and proximal enhancers; the alignments continued through the genes downstream of *GATA1*/*Gata1* (Fig. 6A). An additional, distal regulatory element located upstream of the mouse *Gata1* gene, which was bound by GATA1 and EP300 (Fig. 6A), was found only in mouse (Valverde-Garduno et al. 2004). The DNA sequences of the upstream interval harboring the mouse regulatory element did not align between mouse and human except in portions of the *GLOD5*/*Glod5* genes (Fig. 6A). Thus, the interspecies sequence alignments provide limited information about this distal regulatory element.

**Figure. 6.**
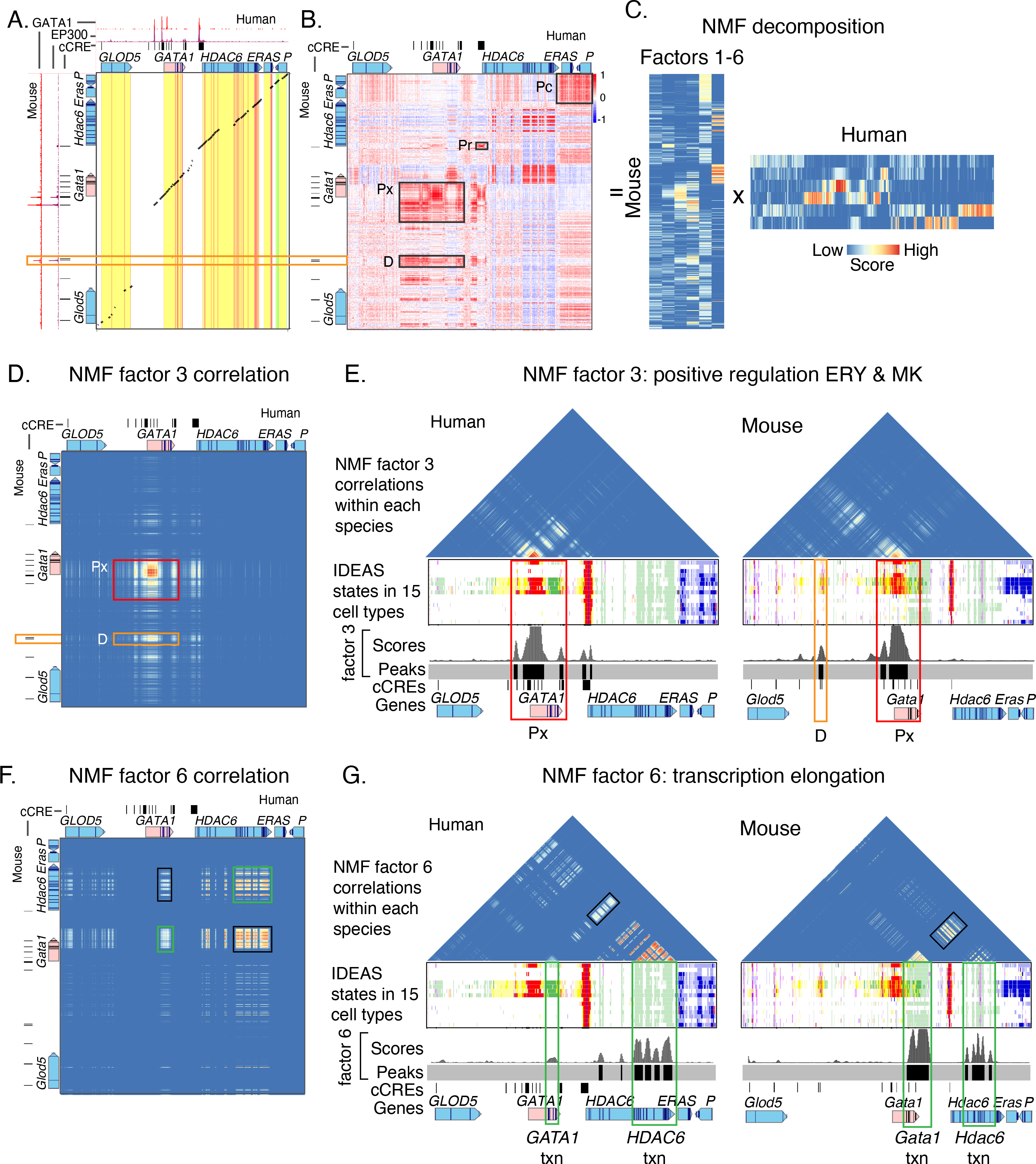
Epigenetic comparisons of regulatory landscapes and cCREs. **(A and B)** DNA sequence alignments and correlations of epigenetic states in human *GATA1* and mouse *Gata1* genes and flanking genes. **(A)** Dot-plot view of chained blastZ alignments by PipMaker (Schwartz et al. 2000) between genomic intervals encompassing and surrounding the human *GATA1* (GRCh38 ChrX:48,760,001-48,836,000; 76kb) and mouse *Gata1* (mm10 ChrX:7,919,401-8,020,800; 101.4kb, reverse complement of reference genome) genes. The axes are annotated with gene locations (GENCODE), predicted *cis*-regulatory elements (cCREs), and binding patterns for GATA1 and EP300 in erythroid cells. **(B)** Matrix of Pearson’s correlation values between epigenetic states (quantitative contributions of each epigenetic feature to the assigned state) across 15 cell types analogous for human and mouse. The correlation is shown for each 200bp bin in one species with all the bins in the other species, using a red-blue heat map to indicate the value of the correlation. Axes are annotated with genes and cCREs in each species. **(C)** Decomposition of the correlation matrix (panel **B**) into six component parts or factors using nonnegative matrix factorization. **(D-G)** Correlation matrices for genomic intervals encompassing *GATA1*/*Gata1* and flanking genes, reconstructed using values from NMF factors. **(D and E)** Correlation matrices using values of NMF factor 3 between human and mouse (panel **D**) or within human and within mouse (panel **E**). The red rectangles highlight the positive regulatory patterns in the *GATA1*/*Gata1* genes (labeled Px), which exhibit conservation of both DNA sequence and epigenetic state pattern. The orange rectangles denote the distal positive regulatory region present only in mouse (labeled D), which shows conservation of epigenetic state pattern without corresponding sequence conservation. Beneath the correlation matrices in panel **E** are maps of IDEAS epigenetic states across 15 cell types, followed by a graph of the score and peak calls for NMF factor 3 and annotation of cCREs (thin black rectangles) and genes. **(F and G)** Correlation matrices using values of NMF factor 6 between human and mouse (panel **F**) or within human and within mouse (panel **G**). The green rectangles highlight the correlation of epigenetic state patterns within the same gene, both across the two species and within each species individually, while the black rectangles highlight the high correlation observed between the two genes *GATA1* and *HDAC6*.

This limitation to sequence alignments led us to explore whether comparisons of epigenetic information would be more informative, utilizing the consistent assignment of epigenetic states in both human and mouse, which do not rely on DNA sequence alignment. In the large genomic regions (76kb and 101kb in the two species) encompassing the orthologous human *GATA1* and mouse *Gata1* genes and surrounding genes, we computed the correlation for each genomic bin between the epigenetic state assignments across cell types in one species and that in the other species for all the bins (Supplemental Fig. S28). This local, all-versus-all comparison of the two loci yielded a matrix of correlation values showing similarities and differences in profiles of epigenetic states in the two species (Fig. 6B). The conserved promoter and proximal enhancers of the *GATA1*/*Gata1* genes were highly correlated in epigenetic states across cell types between the two species, in a region of the matrix that encompassed the aligning DNA sequences (labeled Px in Fig. 6B). In contrast, whereas the mouse-specific distal regulatory element did not align with the human DNA sequence, the epigenetic states annotating it presented high correlations with active epigenetic states in the human *GATA1* locus (labeled D in Fig. 6B).

The complexity of the correlation matrix (Fig. 6B) indicated that multiple epigenetic trends could be contributing to the patterns. To systematically reduce the high dimensionality of the matrix to a set of simpler matrices, we employed nonnegative matrix factorization (NMF) because of its interpretability (Stein-O’Brien et al. 2018; Lee and Roy 2021). The decomposed matrices from NMF revealed a set of factors, each of which (represented by each column in the mouse matrix and each row in the human matrix in Fig. 6C) captures a group of highly correlated elements in the original matrix that show a pattern distinct from the rest of the elements. The complex correlation matrix was decomposed into six distinct factors, as determined by the number of factors at which an “elbow” was found in the BIC score (Supplemental Fig. S29). Each factor encapsulated a specific epigenetic regulatory machinery or process exhibiting consistent cross-cell type patterns in both humans and mice (Supplemental Fig. S30). For example, the correlation matrices reconstructed by using signals from factor 3 exclusively highlighted the cell type-specific positive regulators for the *GATA1*/*Gata1* gene loci; these regulatory elements were evident in reconstructed correlation matrices between species (Fig. 6D) and within individual species (Fig. 6E). By applying a *Z*-score approach to identify peak regions in the factor 3 signal vector (with FDR < 0.1; Supplemental Material), we pinpointed regions in both species showing an epigenetic regulatory machinery exhibiting positive regulatory dynamics for the orthologous *GATA1*/*Gata1* gene loci, particularly in the ERY and MK cell types. In contrast, the correlation matrices reconstructed from the signals for factor 6 (Fig. 6F and G) highlighted regions marked by the transcription elongation modification H3K36me3 (epigenetic states colored green, Fig. 6G). The correlations in the factor 6 elongation signature were observed, as expected, between the human/mouse orthologous gene pairs *GATA1* and *Gata1* as well as between human *HDAC6* and mouse *Hdac6* (green rectangles in Fig. 6F). The factor 6 correlations were also observed between the *GATA1*/*Gata1* and *HDAC6*/*Hdac6* genes (black rectangles in Fig. 6F and G), showing a common process, specifically transcriptional elongation, at both loci. A similar analysis for other factors revealed distinct regulatory processes or elements, such as active promoters (factor 2), exhibiting unique cross-cell type patterns (Supplemental Fig. 30). The genomic bins with high scores for a given NMF factor in human showed high correlation with bins with high scores for that same factor in mouse, indicating that the NMF factors capture a similar set of epigenetic state patterns in each species (Supplemental Fig. S31). The patterns captured by NMF factors 3 and 6 were robust to the choice of *k* in the NMF (Supplemental Fig. S32). Overall, these results underscore this method’s capability to objectively highlight regulatory regions with analogous epigenetic patterns across cell types in both species. This method could aid in extracting additional information about similar epigenetic patterns between human and model organisms such as mice, for which only a portion of their genome aligns with human.

Because some of the NMF factors reflected processes in gene expression and regulation that occur in many genes, some of the highly correlated regions across species could reflect false positives. Thus, it is prudent to restrict the current approach to genomic intervals around orthologous genes to reduce the impact of false discovery. We examined patterns of epigenetic state correlations across cell types between the human *GATA1* gene locus and three non-orthologous loci in mouse to investigate the scope of this issue (Supplemental Material). While genomic bins of high epigenetic state correlation were observed between non-orthologous loci, the discovery of bins implicated in a cell type-specific process, such as erythroid or megakaryocytic regulation, could be enhanced by utilizing a broader background model for computing peaks of NMF signal (Supplemental Fig. S33). With this refined approach to peak identification, the false discovery rate estimated for epigenetic state comparison between the human *GATA1* locus and the mouse *Cd4* locus was reduced to 0.1 or less (Supplemental Fig. S33R). Furthermore, the epigenetic state comparisons between the human *GATA1* locus and the mouse *Rps19* locus revealed a previously unreported region with hallmarks of erythroid regulatory elements (Supplemental Fig. S34). These initial results suggest that the genomic scale of the epigenetic state correlations could be expanded in future work with judicious attention to reducing false discovery, e.g., by linking the discovered elements to evidence of conserved synteny between species.

Examination of human genomic elements shown to be active in a lentiMPRA assay (Agarwal et al. 2023) at 30 loci (Supplemental Table S6) revealed that the active elements were enriched in genomic bins with high cross cell-type epigenetic state correlation between species (Supplemental Fig. S35). The enrichment for active elements was further increased in bins with both high epigenetic state correlation and interspecies sequence conservation, while enrichment was reduced or comparable (depending on approaches used for false discovery thresholds) in bins with only sequence conservation. These results further support the value of the cross cell-type epigenetic state correlation between species in predicting and interpreting cCREs (Supplemental Fig. S36).

The comparison of epigenetic state profiles across cell types also provided a means to categorize cCREs between species that did not require a match in the underlying genomic DNA sequence (Supplemental Figs. S37 and S38). Results from that approach indicated that certain cCREs were potentially involved in regulation of orthologous genes, even for cCREs with DNA sequences that did not align between species.

In summary, the IDEAS joint modeling on the input data compiled here and the consistent state assignments in both mouse and human confirmed and extended previous observations on known regulatory elements, and they revealed both shared and distinctive candidate regulatory elements and states between species. Correlations of state profiles between species provided a comparison of chromatin landscapes even in regions with DNA sequences that were not conserved between species. Our initial results reported here support continuing the development of this approach of comparing cross cell-type epigenetic state profiles between species for functional prediction and interpretation of cCREs.

## Discussion

In this paper, the VISION consortium introduces a set of resources describing the regulatory landscapes of both human and mouse blood cell epigenomes. A key, novel aspect of our work is that the systematic integrative modeling that generated these resources was conducted jointly across the data from both species, which enabled robust comparisons between species without being limited by sequence alignments, allowing comparisons in non-conserved and lineage-specific genomic regions.

One major resource is the annotation of the epigenetic states across the genomes of progenitor and mature blood cells of both species. These state maps show the epigenetic landscape in a compact form, capturing information from the input data on multiple histone modifications, CTCF occupancy, and chromatin accessibility, and they use a common set of epigenetic states to reveal the patterns of epigenetic activity associated with gene expression and regulation both across cell types and between species. A second major resource is a catalog of cCREs actuated in one or more of the blood cell types in each species. The cCREs are predictions of discrete DNA segments likely involved in gene regulation, based on the patterns of chromatin accessibility across cell types, and the epigenetic state annotations suggest the type of activity for each cCRE in each cell type, such as serving as a promoter or enhancer, participating in repression, or inactivity. A third major resource is a quantitative estimate of the regulatory impact of human and mouse cCREs on gene expression in each cell type, i.e., an esRP score, derived from multivariate regression modeling of the epigenetic states in cCREs as predictors of gene expression. The esRP scores are a continuous variable capturing not only the integration of the input epigenetic data, but also the inferred impacts on gene expression. Those impacts may be manifested as activation or repression during regulation or as transcriptional elongation. They are useful for many downstream analyses, such as determining informative groups of cCREs by clustering analysis. These resources along with browsers for visualization and tools for analysis are provided at our project website, http://usevision.org. Among these tools is cCRE_db, which records the several dimensions of annotation of the cCREs and provides a query interface to support custom queries from users.

Our human blood cell cCRE catalog should be valuable for mechanistic interpretations of trait-related human genetic variants. Human genetic variants associated with traits intrinsic to blood cells were significantly enriched in the VISION cCRE catalog, whereas variants associated with a broad diversity of other traits were not enriched. We expect that the extensive annotations in our cCRE catalog combined with information about TFBS motifs and TF occupancy should lead to specific, refined hypotheses for mechanisms by which a variant impacts expression, such as alterations in TF binding, which can be tested experimentally in further work.

The jointly learned state maps and cCRE predictions allowed us to extend previous work on the evolution of regulatory elements between mouse and human. Several previous studies focused on transcription factor (TF) occupancy, e.g. examining key TFs in one tissue across multiple species (Schmidt et al. 2010; Ballester et al. 2014; Villar et al. 2014) or a diverse set of TFs in multiple cell types and in mouse and human (Cheng et al. 2014; Yue et al. 2014; Denas et al. 2015). Other studies focused on discrete regions of high chromatin accessibility in multiple cell types and tissues between mouse and human (Stergachis et al. 2014; Vierstra et al. 2014). These previous studies revealed that only a small fraction of elements was conserved both in genomic sequence and in inferred function. A notable fraction of elements changed considerably during mammalian diversification, including turnover of TF binding site motifs and repurposing of elements (Schmidt et al. 2010; Cheng et al. 2014; Stergachis et al. 2014; Denas et al. 2015). These prior studies focused primarily on regions of the genome with sequences that aligned between human and mouse, with the non-aligning regions used to infer that some elements were lineage-specific and that many were derived from transposable elements and endogenous retroviruses (Bourque 2009; Rebollo et al. 2012; Jacques et al. 2013; Sundaram et al. 2014). Our evolutionary analyses confirmed the previous observations, e.g., finding about one-third of cCREs are conserved in both sequence and inferred function between human and mouse, and further showing that this evolutionary category was highly enriched for proximal regulatory elements.

Going beyond the prior comparative epigenetic studies, our jointly learned epigenetic state maps generated a representation of multiple epigenetic features, not just TF occupancy or chromatin accessibility, and they are continuous in bins across genomes of both species. Using the same set of epigenetic states for annotation of both the human and mouse genomes gave a common “alphabet” (set of states) for both species, which enabled comparisons of the epigenetic profiles between species. In the current work, we explored the utility of these epigenetic comparisons in several ways. For example, the joint clusterings of cCREs between species by esRP scores (derived from the epigenetic state annotations) enabled an analysis that was agnostic to DNA sequence or occupancy by TFs to show considerable sharing of inferred TF activity in both human and mouse. Furthermore, the common alphabet of states allowed us to compare the cross-cell type epigenetic state patterns in large genomic intervals of both species containing orthologous genes, again in a manner agnostic to underlying DNA sequence similarities or differences. These epigenetic comparisons were a strong complement to genomic sequence alignments, revealing regulatory elements with similar epigenetic profiles even in genomic regions in which the DNA sequence does not align between species. Our detection, even in segments of DNA that do not align between species, of epigenetic similarity indicative of a common role in gene regulation suggests that processes or structures, such as chromatin interactions, chromatin complexes, or molecular condensates, may be maintained between species in a manner that is not fully revealed by comparisons of genome sequences. Hence, further studies of this apparent epigenetic dimension of regulatory conservation may be productive. For example, the complex interspecies epigenetic state correlation matrices were decomposed into NMF factors that represented major types of regulatory mechanisms, some that were common across cell types and others that were specific to certain cell types. Further investigation indicated the potential for judicious use of the cell type specific NMF factors in a context of conserved synteny for expanding the scale of the state correlation analysis in future studies.

Previous work compared epigenetic profiles across species, such as the phylo-HMGP method to find different evolutionary states in multi-species epigenomic data (Yang et al. 2018) and the LECIF scores to find evidence of conservation from functional genomic data (Kwon and Ernst 2021). These approaches are powerful but limited to the genomic regions with DNA sequences that align between the species, and thus they will miss the approximately 40% of experimentally demonstrated CREs that are not in aligning regions (Fig. 5D). In contrast, our approach of correlating epigenetic states included both DNA segments that align between human and mouse and those that do not, and it captures more of the experimentally verified cCREs. For comparisons between species, both genomic sequence alignment and epigenetic state annotation across cell types provide important sources of information. Combining both types of data into joint models for predicting CREs could be a productive avenue for future work, not only for improved accuracy but also to allow the contributions of each type of information to determined systematically.

Several innovations were developed to produce the resources introduced here. A major innovation was to extend the IDEAS framework (Zhang et al. 2016) to jointly learn epigenetic states and assign them to annotate the epigenomes in human and mouse blood cells. The IDEAS method employs a Bayesian approach to the modeling to learn the states, which we utilized to bring in states learned from the data in one species as priors for learning states in the data from the second species. Another extension of the IDEAS framework was to learn states based on one feature, specifically ATAC-seq data, defining discrete signal intensity states. This approach was used for calling cCREs, implemented as the IDEAS-IS method (Xiang et al. 2021). The approach is relatively simple and benefits from joint modeling across the input datasets. Other methods for predicting cCREs based on chromatin accessibility across many cell types prevented excessive expansion of the summary calls for overlapping peaks by employing a centroid determination for the DNase hypersensitive sites (DHS) index (Meuleman et al. 2020) or by choosing the highest signal peak for the ENCODE cCRE catalog (The ENCODE Project Consortium et al. 2020). The ENCODE cCRE catalog paired DHS peaks with individual chromatin modifications or CTCF occupancy, which led to complications when data on diagnostic features were missing from some cell types. The IDEAS framework used for the VISION cCRE sets leveraged data in related cell types to ameliorate the impact of missing data.

While the resources introduced here are valuable for many applications, it is prudent to acknowledge their limitations. First, the quality of the products of integrated analyses are limited by the quality and completeness of the input, raw data. We endeavored to reduce the impact of variances in the input data by normalization. The S3V2 procedure (Xiang et al. 2021) systematically normalized the input data to adjust for differences in signal-to-noise and variance in signal across the datasets. Some epigenetic features were not determined in some cell types, and we used the IDEAS method in part because it is able to assign an epigenetic state even in the context of missing data by learning patterns from local similarities in cell types for which the data are present (Zhang and Mahony 2019). However, these approaches cannot completely overcome all issues with variance in input data, and further developments in these directions (such as Shahraki et al. 2023; Xiang et al. 2024) may help to improve integrative resources. Second, the resolution of both the epigenetic state assignments and the cCRE inference is limited to 200 bp, which is the window size we utilized in the IDEAS analyses. Other resources, such as DHS calls (Meuleman et al. 2020), DNase footprints (Vierstra et al. 2020), and motif instances (Weirauch et al. 2014), achieve a higher resolution. Indeed, one can use these higher resolution datasets to derive further information about cCREs, such as families of TFs that are likely to be binding to them. Regarding esRP scores, a third limitation is that we do not make explicit assignments for target genes of cCREs. Predictions of a large number of target gene-cCRE pairs were made in our prior work (Xiang et al. 2020); these assignments cover large genomic intervals around each gene and are most useful when used with further filtering, such as restricting cCREs and target genes to the same topologically associated domains. On-going work is examining other models and approaches for assigning likely target genes to cCREs. A fourth limitation is that our inference of repression-related cCREs applies only to those with stable histone modifications. Elements that had been involved in initiation of repression but eventually were packaged into quiescent chromatin, e.g., via a hit-and-run mechanism (Shah et al. 2019), would not be detected. A fifth limitation concerns the scale of the studies of epigenetic conservation by correlations of epigenetic states. Our current approach is limited to individual examination of specific genetic loci since we used orthologous genes as the initial anchors. Exploring ways to expand the scale of the analytical approach is a goal of future research. Finally, the work presented here was restricted to blood cell types. In future work, extension of the approaches developed in this study to a broader spectrum of cell types would expand the utility of the resulting resources.

In conclusion, we present several important new resources to enable further and more detailed studies of gene regulation in human and mouse blood cells both during normal differentiation and in pathological contexts. The patterns of epigenetic states in cCREs across cell types show value in developing an understanding of how genetic variants impact blood cell traits and diseases. Furthermore, the joint modeling between species opens avenues for further exploration of comparisons of epigenetic landscapes in addition to sequence alignments for insights into evolution and function of regulatory elements between species.

## Methods

### Data generation, collation, normalization, and integration

The data sets used as input, including the ones generated for the work reported here (with methods), are described in Supplemental Material section “Data generation and collection” and Supplemental Tables S1 and S2. The S3V2 approach (Xiang et al. 2021) was used for normalization and denoising the data sets prior to integration. The data sets were integrated to find and assign epigenetic states using IDEAS (Zhang et al. 2016; Zhang and Hardison 2017); the extension of this approach to joint learning and annotation between species is described in Supplemental Material sections “Data normalization” and “Joint systematic integration of human and mouse blood cell epigenomes by IDEAS”.

### Prediction, annotation, and estimation of regulatory impact of cCREs

The identification of cCREs as peaks of chromatin accessibility employed IDEAS in the signal intensity state (IS) mode (Xiang et al. 2021). This approach and comparisons with MACS peaks (Zhang et al. 2008) are described in Supplemental Material section “Prediction of VISION cCREs using IDEAS-IS”. The cCREs are provided in Supplemental Table S3. Annotation of potential cCRE functions used intersections with orthogonal data sets of elements implicated in regulation or chromatin structure (Supplemental Table S5). Enrichment of genetic variants associated with blood cell traits used stratified linkage disequilibrium score regression (sLDSC, Finucane et al. 2015). The impact of epigenetic states in cCREs on regulation of gene expression used a multivariate linear regression approach like one described previously (Xiang et al. 2020). Methods and supplementary results on these analyses are presented in detail in the Supplemental Material.

### Identification of clusters of cCREs based on epigenetic regulatory potential scores

The sets of human and mouse cCREs were placed jointly into groups based on their epigenetic regulatory potential (esRP) scores using a series of *k*-means clustering steps, as described in detail in Supplemental Material and Supplementary Fig. S14. Methods and results for enrichment of the resulting joint meta-clusters (JmCs) for orthogonal sets of regulatory elements and SNPs associated with blood cell traits, along with comparisons of clusters based on chromatin accessibility and H3K27ac signal, are described in Supplemental Material and Supplementary Figs. S15 - S18. Motifs that were differentially enriched across JmCs were identified using the Maelstrom tool in the GimmeMotifs suite (v0.17.1) (Bruse and van Heeringen 2018) and SeqUnwinder (Kakumanu et al. 2017), as described in detail in Supplemental Material and Supplementary Fig. S21.

### Partitioning cCREs to evolutionary categories based on DNA sequence alignments and cCRE calls between species

The human and mouse cCREs were assigned to three evolutionary categories using the following procedure. The set of human cCREs was mapped to mouse genome assembly mm10 using the liftOver tool at the UCSC Genome Browser (Hinrichs et al. 2006). Human cCREs that failed to map to mm10 were grouped as N cCREs. Matches to mouse cCREs for the human cCREs that could be mapped by liftOver to mm10 were determined using the intersect tool in BEDTools (Quinlan and Hall 2010). Human cCREs that overlapped with mouse cCREs were labeled as SF cCREs, while human cCREs that mapped to mm10 but did not match mouse cCREs were labeled as S cCREs. A similar process was performed on the set of mouse cCREs using liftOver to map to human genome build GRCh38

### Calculation of pairwise correlation coefficients for epigenetic landscapes between human and mouse

A bin-to-bin pairwise correlation analysis was used to quantify the similarity of epigenetic landscapes between two DNA regions in human and mouse. For each 200bp bin in one cell type in one species, the assigned epigenetic state was replaced by a vector of mean signals of 8 epigenetic features in the IDEAS state model. After replacing the states in all 15 matched cell types (14 analogous cell types and one pseudo-cell type with average values for all cell types) in the two species, the original two categorical state vectors with 15 elements were converted into two numeric vectors with 120 numbers (Supplemental Fig. S28). The similarity of cross-cell type epigenetic landscape between two bins in the two species was defined as the correlation coefficient between each pair of numeric vectors with 120 numbers. When calculating the correlation coefficients, we added random noise (mean=0, sd=0.2) to the raw values to avoid high correlation coefficients created between regions with states that have low signals. The complex correlation matrix was decomposed into distinctive factors using Nonnegative Matrix Factorization (Lee and Seung 1999). Methods and supplementary results on these analyses are presented in detail in the Supplemental Material.

## Data access

All raw and processed sequencing data generated in this study have been submitted to the NCBI Gene Expression Omnibus (GEO; https://www.ncbi.nlm.nih.gov/geo/) under accession number GSE229101 and the NCBI BioProject database (https://www.ncbi.nlm.nih.gov/bioproject/) under accession number PRJNA952902. Resources developed in the VISION project are available at the website https://usevision.org; the data can be viewed via a track hub at the UCSC Genome Browser or any compatible browser by using this URL: https://usevision.org/data/trackHub/hub.txt or by clicking the track hubs link at usevision.org. The database cCRE db supports flexible user queries on extensive annotation of the cCREs, including epigenetic states and esRP scores across cell types, chromatin accessibility scores across cell types, membership in JmCs, and evolutionary categories. Code developed for this study is in the Supplemental Material and at these GitHub repositories: https://github.com/guanjue/Joint_Human_Mouse_IDEAS_State for the joint human-mouse IDEAS pipeline and https://github.com/usevision/cre_heritability for the sLDSC analysis.

## Supporting information

Scripts for analyzing heritability in cCREs

Scripts for IDEAS modeling of epigenetic states jointly in human and mouse

## Acknowledgments

This work was supported by grants from the National Institutes of Health: R24DK106766 to RCH, GAB, MJW, YZ, FY, JT, MS, DB, DH, JRH, BG; R01DK054937 to GAB; R01GM121613 to YZ and SM; R01GM109453 to QL; R35GM133747 to RCM; F31HG012900 to DJT; R01HG011139; National Science Foundation DBI CAREER 2045500 to SM, and intramural funds from the National Human Genome Research Institute. We dedicate this paper to the memory of JT.

## References

Agarwal V, Inoue F, Schubach M, Martin BK, Dash PM, Zhang Z, Sohota A, Noble WS, Yardimci GG, Kircher M et al. 2023. Massively parallel characterization of transcriptional regulatory elements in three diverse human cell types. bioRxiv doi:10.1101/2023.03.05.531189.

Ballester B, Medina-Rivera A, Schmidt D, Gonzalez-Porta M, Carlucci M, Chen X, Chessman K, Faure AJ, Funnell AP, Goncalves A et al. 2014. Multi-species, multi-transcription factor binding highlights conserved control of tissue-specific biological pathways. eLife 3: e02626.

Bauer DE, Kamran SC, Lessard S, Xu J, Fujiwara Y, Lin C, Shao Z, Canver MC, Smith EC, Pinello L et al. 2013. An erythroid enhancer of BCL11A subject to genetic variation determines fetal hemoglobin level. Science 342: 253–257.

Blobel GA, Weiss MJ. 2009. Nuclear Factors that Regulate Erythropoiesis. In Disorders of Hemoglobin: Genetics, Pathophysiology, and Clinical Management, (ed. MH Steinberg, et al.), pp. 62–85. Cambridge University Press, Cambridge.

Bourque G. 2009. Transposable elements in gene regulation and in the evolution of vertebrate genomes. Curr Opin Genet Dev 19: 607–612.

Bruse N, van Heeringen SJ. 2018. GimmeMotifs: an analysis framework for transcription factor motif analysis. bioRxiv 10.1101/474403.

Buniello A, MacArthur JAL, Cerezo M, Harris LW, Hayhurst J, Malangone C, McMahon A, Morales J, Mountjoy E, Sollis E et al. 2019. The NHGRI-EBI GWAS Catalog of published genome-wide association studies, targeted arrays and summary statistics 2019. Nucleic Acids Res 47: D1005–D1012.

Carroll SB. 2008. Evo-devo and an expanding evolutionary synthesis: a genetic theory of morphological evolution. Cell 134: 25–36.

Cheng L, Li Y, Qi Q, Xu P, Feng R, Palmer L, Chen J, Wu R, Yee T, Zhang J et al. 2021. Single-nucleotide-level mapping of DNA regulatory elements that control fetal hemoglobin expression. Nat Genet 53: 869–880.

Cheng Y, Ma Z, Kim BH, Wu W, Cayting P, Boyle AP, Sundaram V, Xing X, Dogan N, Li J et al. 2014. Principles of regulatory information conservation between mouse and human. Nature 515: 371–375.

Chi AW, Bell JJ, Zlotoff DA, Bhandoola A. 2009. Untangling the T branch of the hematopoiesis tree. Curr Opin Immunol 21: 121–126.

Corces MR, Buenrostro JD, Wu B, Greenside PG, Chan SM, Koenig JL, Snyder MP, Pritchard JK, Kundaje A, Greenleaf WJ et al. 2016. Lineage-specific and single-cell chromatin accessibility charts human hematopoiesis and leukemia evolution. Nat Genet 48: 1193–1203.

Denas O, Sandstrom R, Cheng Y, Beal K, Herrero J, Hardison RC, Taylor J. 2015. Genome-wide comparative analysis reveals human-mouse regulatory landscape and evolution. BMC Genomics 16: 87.

Dong X, Greven MC, Kundaje A, Djebali S, Brown JB, Cheng C, Gingeras TR, Gerstein M, Guigo R, Birney E et al. 2012. Modeling gene expression using chromatin features in various cellular contexts. Genome Biol 13: R53.

Dore LC, Crispino JD. 2011. Transcription factor networks in erythroid cell and megakaryocyte development. Blood 118: 231–239.

Dynan WS, Tjian R. 1983. The promoter-specific transcription factor Sp1 binds to upstream sequences in the SV40 early promoter. Cell 35: 79–87.

Ernst J, Kellis M. 2010. Discovery and characterization of chromatin states for systematic annotation of the human genome. Nat Biotechnol 28: 817–825.

Ernst J, Kellis M. 2012. ChromHMM: automating chromatin-state discovery and characterization. Nat Methods 9: 215–216.

Ferreira R, Ohneda K, Yamamoto M, Philipsen S. 2005. GATA1 function, a paradigm for transcription factors in hematopoiesis. Mol Cell Biol 25: 1215–1227.

Finucane HK, Bulik-Sullivan B, Gusev A, Trynka G, Reshef Y, Loh PR, Anttila V, Xu H, Zang C, Farh K et al. 2015. Partitioning heritability by functional annotation using genome-wide association summary statistics. Nat Genet 47: 1228–1235.

Frangoul H, Altshuler D, Cappellini MD, Chen YS, Domm J, Eustace BK, Foell J, de la Fuente J, Grupp S, Handgretinger R et al. 2021. CRISPR-Cas9 Gene Editing for Sickle Cell Disease and beta-Thalassemia. The New England journal of medicine 384: 252–260.

Fujiwara T, O’Geen H, Keles S, Blahnik K, Linnemann AK, Kang YA, Choi K, Farnham PJ, Bresnick EH. 2009. Discovering hematopoietic mechanisms through genome-wide analysis of GATA factor chromatin occupancy. Mol Cell 36: 667–681.

Gasperini M, Hill AJ, McFaline-Figueroa JL, Martin B, Kim S, Zhang MD, Jackson D, Leith A, Schreiber J, Noble WS et al. 2019. A Genome-wide Framework for Mapping Gene Regulation via Cellular Genetic Screens. Cell 176: 377–390 e319.

Ge T, Chen CY, Neale BM, Sabuncu MR, Smoller JW. 2017. Phenome-wide heritability analysis of the UK Biobank. PLoS Genet 13: e1006711.

Graf T, Enver T. 2009. Forcing cells to change lineages. Nature 462: 587–594.

Gumucio DL, Heilstedt-Williamson H, Gray TA, Tarle SA, Shelton DA, Tagle D, Slightom J, Goodman M, Collins FS. 1992. Phylogenetic footprinting reveals a nuclear protein which binds to silencer sequences in the human g and e globin genes. Mol Cell Biol 12: 4919–4929.

Hamamoto K, Fukaya T. 2022. Molecular architecture of enhancer-promoter interaction. Curr Opin Cell Biol 74: 62–70.

Hardison RC. 2000. Conserved noncoding sequences are reliable guides to regulatory elements. Trends in Genetics 16: 369–372.

Hardison RC. 2012. Genome-wide epigenetic data facilitate understanding of disease susceptibility association studies. J Biol Chem 287: 30932–30940.

Hardison RC, Zhang Y, Keller CA, Xiang G, Heuston EF, An L, Lichtenberg J, Giardine BM, Bodine D, Mahony S et al. 2020. Systematic integration of GATA transcription factors and epigenomes via IDEAS paints the regulatory landscape of hematopoietic cells. IUBMB Life 72: 27–38.

Heintzman ND, Hon GC, Hawkins RD, Kheradpour P, Stark A, Harp LF, Ye Z, Lee LK, Stuart RK, Ching CW et al. 2009. Histone modifications at human enhancers reflect global cell-type-specific gene expression. Nature 459: 108–112.

Hindorff LA, Sethupathy P, Junkins HA, Ramos EM, Mehta JP, Collins FS, Manolio TA. 2009. Potential etiologic and functional implications of genome-wide association loci for human diseases and traits. Proc Natl Acad Sci U S A 106: 9362–9367.

Hinrichs AS, Karolchik D, Baertsch R, Barber GP, Bejerano G, Clawson H, Diekhans M, Furey TS, Harte RA, Hsu F et al. 2006. The UCSC Genome Browser Database: update 2006. Nucleic Acids Res 34: D590–598.

Hoffman MM, Ernst J, Wilder SP, Kundaje A, Harris RS, Libbrecht M, Giardine B, Ellenbogen PM, Bilmes JA, Birney E et al. 2013. Integrative annotation of chromatin elements from ENCODE data. Nucleic Acids Res 41: 827–841.

Jacques PE, Jeyakani J, Bourque G. 2013. The majority of primate-specific regulatory sequences are derived from transposable elements. PLoS Genet 9: e1003504.

Jansen C, Ramirez RN, El-Ali NC, Gomez-Cabrero D, Tegner J, Merkenschlager M, Conesa A, Mortazavi A. 2019. Building gene regulatory networks from scATAC-seq and scRNA-seq using Linked Self Organizing Maps. PLoS Comput Biol 15: e1006555.

Jian J, Konopka J, Liu C. 2013. Insights into the role of progranulin in immunity, infection, and inflammation. J Leukoc Biol 93: 199–208.

Kaczynski J, Cook T, Urrutia R. 2003. Sp1- and Kruppel-like transcription factors. Genome Biol 4: 206.

Kakumanu A, Velasco S, Mazzoni E, Mahony S. 2017. Deconvolving sequence features that discriminate between overlapping regulatory annotations. PLoS Comput Biol 13: e1005795.

Karlić R, Chung HR, Lasserre J, Vlahovicek K, Vingron M. 2010. Histone modification levels are predictive for gene expression. Proc Natl Acad Sci U S A 107: 2926–2931.

King DC, Taylor J, Zhang Y, Cheng Y, Lawson HA, Martin J, ENCODE groups for Transcriptional Regulation and Multispecies Sequence Analysis, Chiaromonte F, Miller W, Hardison RC. 2007. Finding cis-regulatory elements using comparative genomics: some lessons from ENCODE data. Genome Res 17: 775–786.

Kondo M, Wagers AJ, Manz MG, Prohaska SS, Scherer DC, Beilhack GF, Shizuru JA, Weissman IL. 2003. Biology of hematopoietic stem cells and progenitors: implications for clinical application. Annu Rev Immunol 21: 759–806.

Kwon SB, Ernst J. 2021. Learning a genome-wide score of human-mouse conservation at the functional genomics level. Nature communications 12: 2495.

Laurenti E, Göttgens B. 2018. From haematopoietic stem cells to complex differentiation landscapes. Nature 553: 418–426.

Lee DD, Seung HS. 1999. Learning the parts of objects by non-negative matrix factorization. Nature 401: 788–791.

Lee DI, Roy S. 2021. GRiNCH: simultaneous smoothing and detection of topological units of genome organization from sparse chromatin contact count matrices with matrix factorization. Genome Biol 22: 164.

Libbrecht MW, Chan RCW, Hoffman MM. 2021. Segmentation and genome annotation algorithms for identifying chromatin state and other genomic patterns. PLoS Comput Biol 17: e1009423.

Martens JH, Stunnenberg HG. 2013. BLUEPRINT: mapping human blood cell epigenomes. Haematologica 98: 1487–1489.

Maston GA, Evans SK, Green MR. 2006. Transcriptional Regulatory Elements in the Human Genome. Annu Rev Genomics Hum Genet 7: 29–59.

Maurano MT, Humbert R, Rynes E, Thurman RE, Haugen E, Wang H, Reynolds AP, Sandstrom R, Qu H, Brody J et al. 2012. Systematic localization of common disease-associated variation in regulatory DNA. Science 337: 1190–1195.

Meuleman W, Muratov A, Rynes E, Halow J, Lee K, Bates D, Diegel M, Dunn D, Neri F, Teodosiadis A et al. 2020. Index and biological spectrum of human DNase I hypersensitive sites. Nature 584: 244–251.

Noyes MB, Christensen RG, Wakabayashi A, Stormo GD, Brodsky MH, Wolfe SA. 2008. Analysis of homeodomain specificities allows the family-wide prediction of preferred recognition sites. Cell 133: 1277–1289.

Payne KJ, Crooks GM. 2002. Human hematopoietic lineage commitment. Immunol Rev 187: 48–64.

Pennacchio LA, Rubin EM. 2001. Genomic strategies to identify mammalian regulatory sequences. Nat Rev Genet 2: 100–109.

Pimkin M, Kossenkov AV, Mishra T, Morrissey CS, Wu W, Keller CA, Blobel GA, Lee D, Beer MA, Hardison RC et al. 2014. Divergent functions of hematopoietic transcription factors in lineage priming and differentiation during erythro-megakaryopoiesis. Genome Res 24: 1932–1944.

Pollard KS, Hubisz MJ, Rosenbloom KR, Siepel A. 2010. Detection of nonneutral substitution rates on mammalian phylogenies. Genome Res 20: 110–121.

Qi Q, Cheng L, Tang X, He Y, Li Y, Yee T, Shrestha D, Feng R, Xu P, Zhou X et al. 2021. Dynamic CTCF binding directly mediates interactions among cis-regulatory elements essential for hematopoiesis. Blood 137: 1327–1339.

Quinlan AR, Hall IM. 2010. BEDTools: a flexible suite of utilities for comparing genomic features. Bioinformatics 26: 841–842.

Rebollo R, Romanish MT, Mager DL. 2012. Transposable elements: an abundant and natural source of regulatory sequences for host genes. Annu Rev Genet 46: 21–42.

Ringrose L, Paro R. 2004. Epigenetic regulation of cellular memory by the Polycomb and Trithorax group proteins. Annu Rev Genet 38: 413–443.

Rothenberg EV, Taghon T. 2005. Molecular genetics of T cell development. Annu Rev Immunol 23: 601–649.

Schmidt D, Wilson MD, Ballester B, Schwalie PC, Brown GD, Marshall A, Kutter C, Watt S, Martinez-Jimenez CP, Mackay S et al. 2010. Five-vertebrate ChIP-seq reveals the evolutionary dynamics of transcription factor binding. Science 328: 1036–1040.

Schwartz S, Zhang Z, Frazer KA, Smit A, Riemer C, Bouck J, Gibbs R, Hardison R, Miller W. 2000. PipMaker-A web server for aligning two genomic DNA sequences. Genome Res 10: 577–586.

Shah M, Funnell APW, Quinlan KGR, Crossley M. 2019. Hit and Run Transcriptional Repressors Are Difficult to Catch in the Act. Bioessays 41: e1900041.

Shahraki MF, Farahbod M, Libbrecht MW. 2023. Robust chromatin state annotation. bioRxiv 10.1101/2023.07.15.549175.

Spangrude GJ, Heimfeld S, Weissman IL. 1988. Purification and characterization of mouse hematopoietic stem cells. Science 241: 58–62.

Stein-O’Brien GL, Arora R, Culhane AC, Favorov AV, Garmire LX, Greene CS, Goff LA, Li Y, Ngom A, Ochs MF et al. 2018. Enter the Matrix: Factorization Uncovers Knowledge from Omics. Trends Genet 34: 790–805.

Stergachis AB, Neph S, Sandstrom R, Haugen E, Reynolds AP, Zhang M, Byron R, Canfield T, Stelhing-Sun S, Lee K et al. 2014. Conservation of trans-acting circuitry during mammalian regulatory evolution. Nature 515: 365–370.

Strahl BD, Allis CD. 2000. The language of covalent histone modifications. Nature 403: 41–45.

Stunnenberg HG, International Human Epigenome C, Hirst M. 2016. The International Human Epigenome Consortium: A Blueprint for Scientific Collaboration and Discovery. Cell 167: 1145–1149.

Sundaram V, Cheng Y, Ma Z, Li D, Xing X, Edge P, Snyder MP, Wang T. 2014. Widespread contribution of transposable elements to the innovation of gene regulatory networks. Genome Res 24: 1963–1976.

Tenen DG, Hromas R, Licht JD, Zhang DE. 1997. Transcription factors, normal myeloid development, and leukemia. Blood 90: 489–519.

The ENCODE Project Consortium. 2012. An integrated encyclopedia of DNA elements in the human genome. Nature 489: 57–74.

The ENCODE Project Consortium, Moore JE, Purcaro MJ, Pratt HE, Epstein CB, Shoresh N, Adrian J, Kawli T, Davis CA, Dobin A et al. 2020. Expanded encyclopaedias of DNA elements in the human and mouse genomes. Nature 583: 699–710.

Valverde-Garduno V, Guyot B, Anguita E, Hamlett I, Porcher C, Vyas P. 2004. Differences in the chromatin structure and cis-element organization of the human and mouse GATA1 loci: implications for cis-element identification. Blood 104: 3106–3116.

van Arensbergen J, FitzPatrick VD, de Haas M, Pagie L, Sluimer J, Bussemaker HJ, van Steensel B. 2017. Genome-wide mapping of autonomous promoter activity in human cells. Nat Biotechnol 35: 145–153.

van Pampus EC, Denkers IA, van Geel BJ, Huijgens PC, Zevenbergen A, Ossenkoppele GJ, Langenhuijsen MM. 1992. Expression of adhesion antigens of human bone marrow megakaryocytes, circulating megakaryocytes and blood platelets. Eur J Haematol 49: 122–127.

Vierstra J, Lazar J, Sandstrom R, Halow J, Lee K, Bates D, Diegel M, Dunn D, Neri F, Haugen E et al. 2020. Global reference mapping of human transcription factor footprints. Nature 583: 729–736.

Vierstra J, Rynes E, Sandstrom R, Zhang M, Canfield T, Hansen RS, Stehling-Sun S, Sabo PJ, Byron R, Humbert R et al. 2014. Mouse regulatory DNA landscapes reveal global principles of cis-regulatory evolution. Science 346: 1007–1012.

Villar D, Flicek P, Odom DT. 2014. Evolution of transcription factor binding in metazoans - mechanisms and functional implications. Nat Rev Genet 15: 221–233.

Weirauch MT, Yang A, Albu M, Cote AG, Montenegro-Montero A, Drewe P, Najafabadi HS, Lambert SA, Mann I, Cook K et al. 2014. Determination and inference of eukaryotic transcription factor sequence specificity. Cell 158: 1431–1443.

Weiss MJ, Yu C, Orkin SH. 1997. Erythroid-cell-specific properties of transcription factor GATA-1 revealed by phenotypic rescue of a gene-targeted cell line. Mol Cell Biol 17: 1642–1651.

Xiang G, Giardine BM, Mahony S, Zhang Y, Hardison RC. 2021. S3V2-IDEAS: a package for normalizing, denoising and integrating epigenomic datasets across different cell types. Bioinformatics 37: 3011–3013.

Xiang G, Guo Y, Bumcrot D, Sigova A. 2024. JMnorm: a novel joint multi-feature normalization method for integrative and comparative epigenomics. Nucleic Acids Res 52: e11.

Xiang G, Keller CA, Heuston E, Giardine BM, An L, Wixom AQ, Miller A, Cockburn A, Sauria MEG, Weaver K et al. 2020. An integrative view of the regulatory and transcriptional landscapes in mouse hematopoiesis. Genome Res 30: 472–484.

Xu J, Shao Z, Glass K, Bauer DE, Pinello L, Van Handel B, Hou S, Stamatoyannopoulos JA, Mikkola HK, Yuan GC et al. 2012. Combinatorial assembly of developmental stage-specific enhancers controls gene expression programs during human erythropoiesis. Dev Cell 23: 796–811.

Yang Y, Gu Q, Zhang Y, Sasaki T, Crivello J, O’Neill RJ, Gilbert DM, Ma J. 2018. Continuous-Trait Probabilistic Model for Comparing Multi-species Functional Genomic Data. Cell Syst 7: 208–218 e211.

Yue F, Cheng Y, Breschi A, Vierstra J, Wu W, Ryba T, Sandstrom R, Ma Z, Davis C, Pope BD et al. 2014. A comparative encyclopedia of DNA elements in the mouse genome. Nature 515: 355–364.

Zhang Y, An L, Yue F, Hardison RC. 2016. Jointly characterizing epigenetic dynamics across multiple human cell types. Nucleic Acids Res 44: 6721–6731.

Zhang Y, Hardison RC. 2017. Accurate and reproducible functional maps in 127 human cell types via 2D genome segmentation. Nucleic Acids Res 45: 9823–9836.

Zhang Y, Liu T, Meyer CA, Eeckhoute J, Johnson DS, Bernstein BE, Nussbaum C, Myers RM, Brown M, Li W et al. 2008. Model-based analysis of ChIP-Seq (MACS). Genome Biol 9: R137.

Zhang Y, Mahony S. 2019. Direct prediction of regulatory elements from partial data without imputation. PLoS Comput Biol 15: e1007399.

