## Supplemental Material, Text and Figures for "Interspecies regulatory landscapes and elements revealed by novel joint systematic integration of human and mouse blood cell epigenomes"

For

**Table of Contents**

| Topic | Suppl. Fig. or Table. | Pages |
| --- | --- | --- |
| Data generation and collection | Tables S1, S2 | 4-7 |
| Data normalization | Fig. S1 | 7-9 |
| Joint systematic integration of human and mouse blood cell epigenomes by IDEAS | Figs. S2, S3, S4, S5 | 9-18 |
| Prediction of VISION cCREs using IDEAS-IS | Fig. S6; Table S3 | 18-21 |
| Comparison of ATAC-seq peak calls between IDEAS-IS and MACS3 | Fig. S7 | 22-23 |
| Annotation of VISION cCREs using other datasets of elements | Figs. S8, S9, S10; Tables S4, S5 | 24-28 |
| Enrichment of genetic variants for blood cell related traits in the VISION human cCRE collection | Fig. S11 | 29-33 |
| Estimation of the impact of epigenetic states and cCREs on gene expression |  | 33-40 |
| Visualization of cCRE esRP scores across cell types using UMAPs | Fig. S12 | 40-41 |
| Comparisons of esRP, chromatin accessibility, and H3K27ac in cCREs | Figs. S12, S13 | 42-44 |
| Clustering of cCREs based on esRP scores | Fig. S14 | 45-48 |
| Enrichment of specific JmCs for promoter or enhancer annotations | Fig. S15 | 49-50 |
| Enrichment of Joint MetaClusters (JmCs) assigned to cCREs in gene loci |  | 50 |
| Enrichment of trait-associated SNPs in the joint metaclusters | Fig. S16 | 51-52 |
| Comparisons of cCREs clusters based on esRP, chromatin accessibility, and H3K27ac for distinguishing among cell types | Figs. S17, S18 | 52-55 |
| Self-organizing maps | Fig. S19 | 55 |
| Enrichment for transcription factor binding site motifs in joint metaclusters of cCREs |  | 56 |
| Binding of CTCF to cCREs in JmCs: species similarities and differences | Fig. S20, S21 | 57-60 |
| Small differences in the number of cCREs in the structure and function conserved category (SF cCREs) | Fig. S22 | 61 |
| Comparison of magnitude of chromatin accessibility in orthologous cCREs in human and mouse blood cells | Fig. S23 | 62 |
| Profiles of epigenetic states in the three evolutionary categories of cCREs in human and mouse blood cells | Fig. S24 | 63-64 |
| Impact of cCRE location relative to the transcription start site (TSS) of genes on conservation scores | Fig. S25, S26 | 65-67 |
| Correlation between human and mouse of gene expression levels in blood cell types | Fig. S27 | 67 |
| Comparison of epigenetic state landscapes between human and mouse blood cells | Fig. S28 | 68 |
| Decomposition of the correlation matrix of epigenetic states using nonnegative matrix factorization (NMF) | Fig. S29, S30, S31, S32 | 69-74 |
| False discovery rate in finding related elements using NMF of correlation matrices of epigenetic states | Fig. S33, S34 | 75-84 |
| Effectiveness of interspecies sequence alignment and epigenetic state correlation for regulatory element discovery | Fig. S35, S36 | 85-97 |
| Enrichment of cCREs in the three evolutionary categories for specific JmCs in the vicinity of orthologous genes | Fig. S37, S38 | 97-100 |
| Code Access |  | 101 |
| References |  | 101-109 |

### **Data generation and collection**

To conduct our joint integrative analysis of human and mouse regulatory landscapes across progenitor and mature blood cell types, we collected 404 data sets of epigenetic features related to gene regulation and expression, 216 in human and 188 in mouse. These datasets (illustrated in Figure 1 of the main text) are listed in Supplemental Table S1 along with metadata such as source laboratory (consortia and individual laboratories), biosamples, the molecule ascertained, and file IDs. The 404 data sets included 44 new ones that were generated for this work; these are listed in Supplemental Table S2 along with information about the biosamples and molecules ascertained. Many of the new data sets were for epigenetic features and RNA in the HUDEP1 cell line, which is Human Umbilical Cord Blood-Derived Erythroid Progenitor cell line 1 that expresses gamma-globin upon induced differentiation.

Most data collected were from primary cell populations, and data from commonly used cell lines were also included. Much of the epigenetic data for mature blood cell types in humans was collected as mapped reads (human genome build GRCh38) in bigWig format from the data portal for the BLUEPRINT Project (Martens and Stunnenberg 2013; Stunnenberg et al. 2016). Additional data, including ATAC-seq for human progenitor cells (Corces et al. 2016), and the full set of features in HUDEP1 (generated for this paper), HUDEP2 (Cheng et al. 2021; Qi et al. 2021), and K562 (The ENCODE Project Consortium et al. 2020) cell lines were collected as sequencing reads and processed through the mapping pipelines described in a previous VISION paper (Xiang et al. 2020b), mapping the reads to human genome build GRCh38. Replicate data were obtained for most but not all features across the cell types, especially for the human blood cell types (Supplemental Table S1), and integrative analysis was conducted keeping the replicate sets separate for each cell type. The sequencing reads in the data for mouse hematopoietic cells, both new and described previously, were mapped to mouse genome build mm10 (Xiang et al. 2020b).

The epigenetic features studied here were associated with several processes in gene expression and regulation. Chromatin accessibility is a general feature of almost all regulatory elements, and it was measured by the Assay for Transposase Accessible Chromatin with high throughput sequencing (ATAC-seq, Buenrostro et al. 2013; Corces et al. 2016) or by DNase-seq (Thurman et al. 2012) for almost all cell types in both species. Available ChIP-seq data for up to six histone modifications provided information related to different elements or processes in gene expression, specifically H3K4me3 for promoters and H3K4me1 for enhancers (Birney et al. 2007; Heintzman et al. 2007), H3K27ac for activation (Roh et al. 2005; Smith and Shilatifard 2014), H3K36me3 for transcriptional elongation (Li et al. 2002), H3K27me3 for repression by the Polycomb repressor complex (Muller et al. 2002; Schwartz et al. 2006), and H3K9me3 for heterochromatin (Padeken et al. 2022). ChIP-seq data on occupancy by the structural protein CTCF associated with insulation (West et al. 2002) were available in many cell types. Bulk RNA-seq data were collected for all cell types.

Different data sets on epigenetic features in K562 cells were used for the integrative analysis and for the compilation of orthogonal annotations. Specifically, the CTCF ChIP-seq data from K562 cells used for the integration were ENCODE files ENCFF000YLW and ENCFF000YLY. For the orthogonal annotations, we used the CTCF ChIP-seq peaks reported by an independent laboratory (Pugacheva et al. 2015). Other data sets from K562 cells used in the compilation of orthogonal annotations (see later section) were not used in the IDEAS integrative analysis.

The newly generated data utilized in this paper came from purified populations of mouse blood cells and from human and mouse blood cell lines. Populations of primary blood cells were purified from mouse bone marrow by cell sorting using distinctive panels of surface markers listed in Supplemental Table S1. Cell lines included Human Umbilical Cord Blood-Derived Erythroid Progenitor cell line 1, abbreviated HUDEP1, a mouse model of *Gata1*-deficient erythroid progenitor cells (G1E), and a subclone of G1E cells conditionally rescued with a hybrid GATA1-estrogen receptor protein (G1E-ER4). The latter cell line was induced for erythroid maturation by treatment with 1x10^-8^ M estradiol for 24 hr (G1E-ER4+E2 cells).

The protocol for ChIP-seq followed previously published procedures (Cheng et al. 2009). In general, approximately 2.5 × 10^7^ cells were used for each immunoprecipitation. Cells were cross-linked with 1% formaldehyde for 10 min at room temperature with rotation, and the reaction was quenched with glycine at a final concentration of 125 mM. Cross-linked cells were then lysed and resuspended in 2 mL of RIPA buffer and sonicated for 12 cycles with a Branson 250 sonifier (10 s on/90 s off for a total of 2 min of pulses with 20% output from the micro-tip) or equivalent to obtain fragments of chromatin approximately 200–300 bp in size. Supernatants were precleared by incubation with 200 µL of protein A/G agarose bead slurry (Thermo Fisher Scientific, cat. #15918014) overnight at 4°C with rotation. Meanwhile, 12.5 µg of IP antibody was incubated with 50 µL of protein A/G agarose bead slurry in 1 mL of PBS overnight at 4°C with rotation. Saved precleared chromatin (20 µL) was used as the input sample. Precleared chromatin was incubated with the antibody–bead complex for 7 hr at 4°C with rotation. Cross-linking of DNA was reversed by incubation with RNase A (1 µg/µL), proteinase K (0.2 mg/mL), and 0.25 M NaCl overnight at 65°C. Immunoprecipitated DNA was purified using the Qiagen PCR Extraction Kit and eluted with 20 µL of EB elution buffer. Sequencing libraries were prepared using the NEBNext Ultra II DNA Library Prep Kit (NEB, cat. #E7645) with TruSeq adaptors. Libraries were sequenced on an Illumina system, either HiSeq 4000, HiSeq 2000, or NextSeq 2000.

The protocol for RNA-seq followed previously published procedures (Cheng et al. 2021; Qi et al. 2021). RNA was extracted from 1 million cells using the RNeasy Mini Kit (Qiagen). The TruSeq Stranded mRNA Library Prep Kit (Illumina) was used to enrich for polyA+ RNA and to create libraries for HiSeq2000 sequencing (Illumina).

**Data normalization**

The epigenomic data from multiple sources differed in many properties, including sequencing depth, fraction of reads on target, and signal-to-noise ratio (Xiang et al. 2020a). To reduce the impact of these technical differences, we used an improved version of the S3norm method, called S3V2, to normalize and denoise all data sets. This normalization method was designed to match the ranges of both signal intensities and variances across epigenetic datasets (Xiang et al. 2021). In this procedure, we first generated a reference signal track for each epigenetic feature by computing the mean signal of all data sets for that feature at each genomic location (200 bp bins). Then, the peak means and the background non-zero means of the reference signal tracks for the different epigenetic features were equalized by the S3norm method (Xiang et al. 2020a). We then used these mean-adjusted references as the new reference signal tracks for each epigenetic feature. For all datasets of the same epigenetic feature, we normalized their signal against the reference signal track using the recently developed S3V2 method (Xiang et al. 2021). The S3V2 version of the method was designed to adjust both the non-zero means and the standard deviations of the background regions, so that it can better reduce background noise in some data sets with higher variance at the background regions.

This adjustment produced a stronger and more consistent correlation by feature across cell types, indicating that the denoising and normalization were effective (Supplemental Fig. S1).

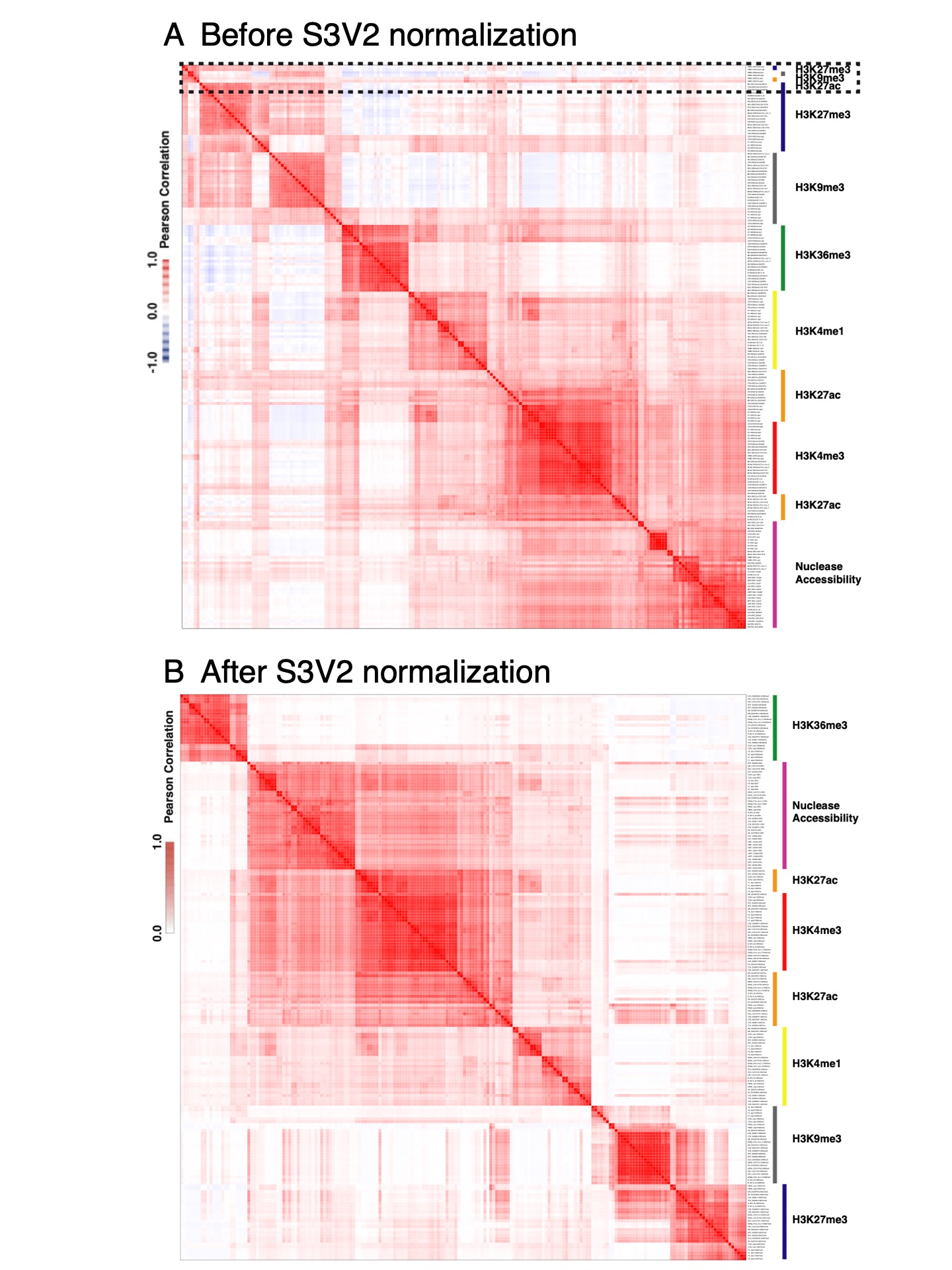

**Supplemental Figure S1.** *Improved consistency of epigenetic signals after normalization and denoising.* The Pearson’s correlation coefficient for the signal intensity values for each epigenetic feature in each human blood cell type (in 200 bp bins) across the human genome (GRCh38) was computed, organized by hierarchical clustering, and displayed as these heatmaps, with a stronger intensity of red meaning a higher positive correlation and blue indicating a negative correlation. These correlations were computed before (A) and after (B) normalization using the S3V2 method. The epigenetic features are labeled by different color bars on the right side the heatmap. Most datasets clustered by epigenetic features across cell types after normalization, whereas the groupings by feature were not as apparent prior to normalization. For example, the box with dotted lines in panel A highlights datasets of epigenetic features associated with different functions (H3K27me3, H3K9me3 and H3K27ac), which form an unexpected group prior to normalization and then are grouped with other datasets for those individual features after normalization and denoising. The clear separation between active features and the repressive features after normalization and denoising by S3V2 indicated that this procedure can reduce false positive signals, which may increase the correlation between sets of different features.

**Joint systematic integration of human and mouse blood cell epigenomes by IDEAS**

*Strategy for the joint systematic integration*

A powerful class of methods for integrative analysis of epigenomes involves statistical modeling to discover frequently occurring combinations of epigenetic features, comprising epigenetic states, and then assigning DNA intervals (often of 200 bp) to those states to produce regulatory annotations across the genome. These segmentation and genome annotation (SAGA) methods (Libbrecht et al. 2021) include ChromHMM (Ernst and Kellis 2012), Segway (Hoffman et al. 2012), and IDEAS (Zhang et al. 2016; Zhang and Hardison 2017). We employed IDEAS because its simultaneous two-dimensional modeling along chromosomes and across cell types provides a consistent and well-resolved annotation while leveraging epigenetic information from locally related cell types when assigning states in cell types with missing data (Zhang and Mahony 2019). Moreover, its Bayesian statistical framework allows the incorporation of epigenetic models from different studies and even from different species. This latter feature was critical to the joint modeling between species described here.

We conducted an iterative, joint training on the epigenomic data of both human and mouse blood cells to ensure that the same set of epigenetic states was learned and applied for both species. Previous studies showed that the epigenetic states uncovered by SAGA methods such as ChromHMM (Ernst and Kellis 2012) were similar in both mouse and human (Yue et al. 2014; Roadmap Epigenomics Consortium et al. 2015; Gorkin et al. 2020). Indeed, when the epigenomic data from mouse or human were used separately as input to IDEAS, most of the resulting states were shared between the species (Supplemental Fig. S2). The states specific to human or mouse were often similar to the shared states but with small variations in one or more epigenetic features; no clear evidence for a state specific to either species was found.

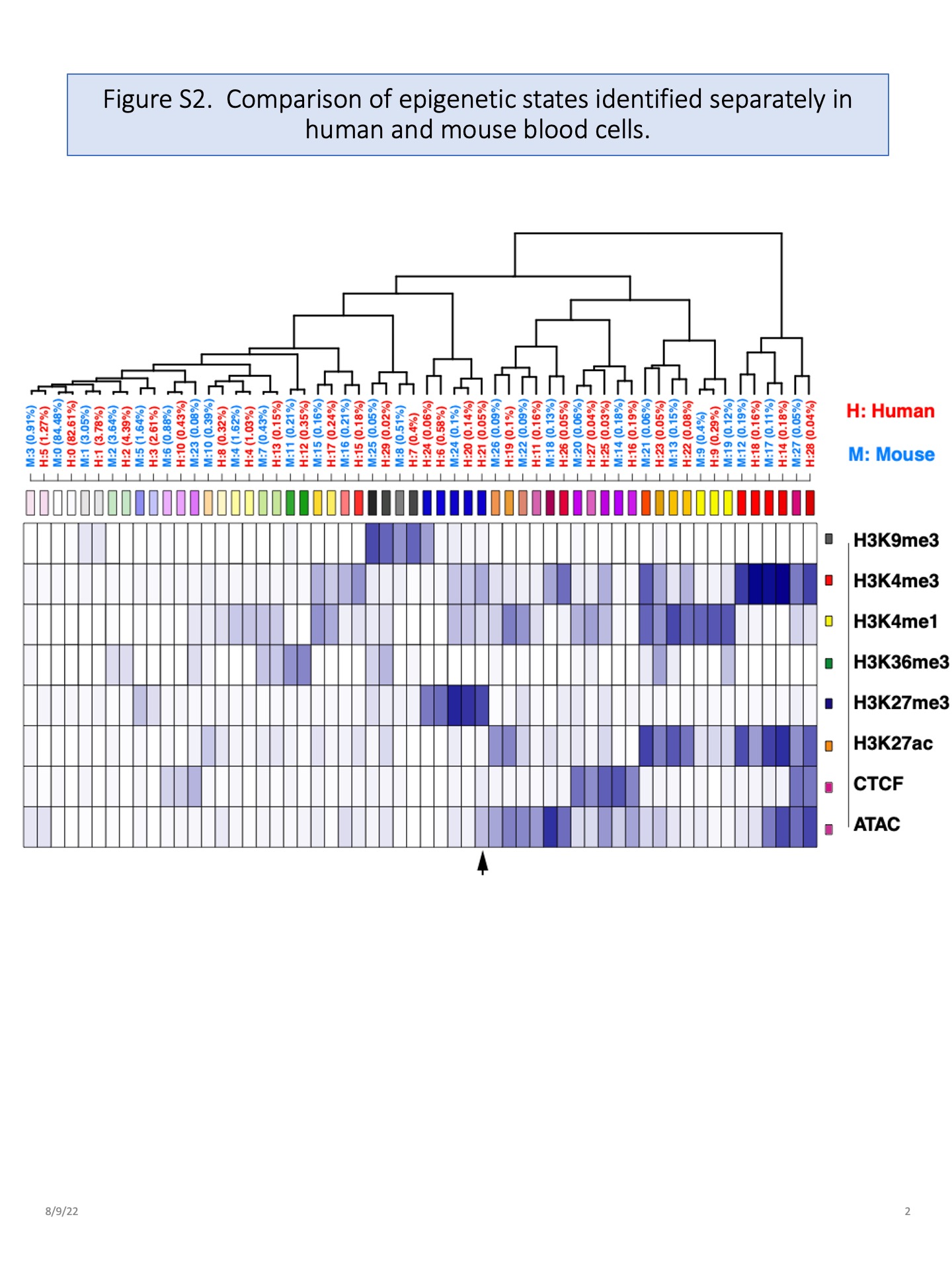

**Supplemental Figure S2.** *Similar epigenetic states learned by modeling data separately in mouse and human blood cells.* The profiles of epigenetic feature emissions from the epigenetic states identified separately in human and mouse blood cells were compared and presented as a heatmap organized by hierarchical clustering of the columns, with each column representing one state learned in the modeling. The arrow at the bottom points to an example of a state found only in one species; note that it is very similar to other states found in both species.

*Methods for the joint systematic integration*

The joint modeling began with a search for epigenetic states that exhibit similar combinatorial patterns across different epigenetic features in both human and mouse, which we defined as reproducible epigenetic states (illustrated in Figure 2A in main text). Initially, 200 sets of epigenetic states were identified by IDEAS at 100 randomly selected, 50 MB regions from each species, using the S3V2-IDEAS pipeline (Xiang et al. 2021) on both species. The IDEAS method learns the number of states in each run. In practice, the number of states within each of the 200 initial epigenetic state sets can be influenced by a fixed parameter, “mycut”, at line 472 of the “S3V2_IDEAS_ESMP/bin/IDEAS_2018/bin/ideaspipe.R” script in the IDEAS package. The "mycut" parameter can range from 0 to 1, such that a larger "mycut" setting can decrease the number of states within each set, and vice versa. For this study, the "mycut" value is set to 0.55. The 200 sets of epigenetic states contained 7,550 states, meaning that the average number of states learned in each run was 37.75. The reproducible IDEAS states were selected by an internal *combineState* function in the IDEAS pipeline. Intuitively, this function clusters the 200 sets of epigenetic states from the initial IDEAS runs into groups based on the Euclidean distances between their average epigenetic features’ signal vectors using hierarchical clustering. If an output cluster contains epigenetic states generated by more than a certain proportion of the 200 sets, these states are merged to define a reproducible epigenetic state. Here, we wanted to include as reproducible states not only those found at high frequency in all the runs on data from both species but also those that were either highly reproducible in one species and also found in the other species or those that were moderately reproducible in both species. By requiring that a state appear in at least 52% of the 200 runs, we ensured that the collected states included the latter two categories. This search led to the retention of 27 reproducible states (steps 1 and 2).

Then, to analyze the full epigenomic information in each species, we used these 27 reproducible states as the priors for the distribution parameters in the two rounds of IDEAS runs across the whole genomes of both species. The two rounds of IDEAS runs for the two species were performed sequentially, alternating between human and mouse. After each round of IDEAS run, the frequency, mean, and variance parameters for each epigenetic state were updated, so that the information of the species at the current round was then integrated into the next IDEAS run (steps 3a and 3b). We found that two rounds of whole genome IDEAS runs were sufficient, since the mean squared error of the mutual nearest neighbors of epigenetic states was very small when comparing human and mouse after the second round (Supplemental Fig. S3).

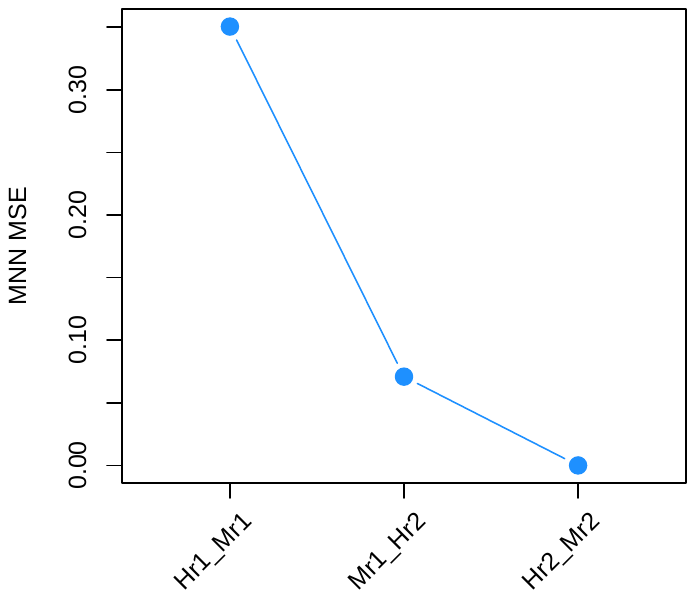

**Supplemental Figure S3.** *The Mean Squared Errors (MSEs) of the Mutual Nearest Neighbor epigenetic states observed in the following IDEAS runs pairs:* Human_round1_VS_ Mouse_round1 (Hr1_Mr1), Mouse_round1_VS_ Human_round2 (Mr1_Hr2), and Human_round2_VS_ Mouse_round2 (Hr2_Mr2).

Two heterogenous states, identified by their coefficient of variance (Supplemental Fig. S4), were removed, because such states previously had been observed to be composites of low frequency states (Xiang et al. 2020b).

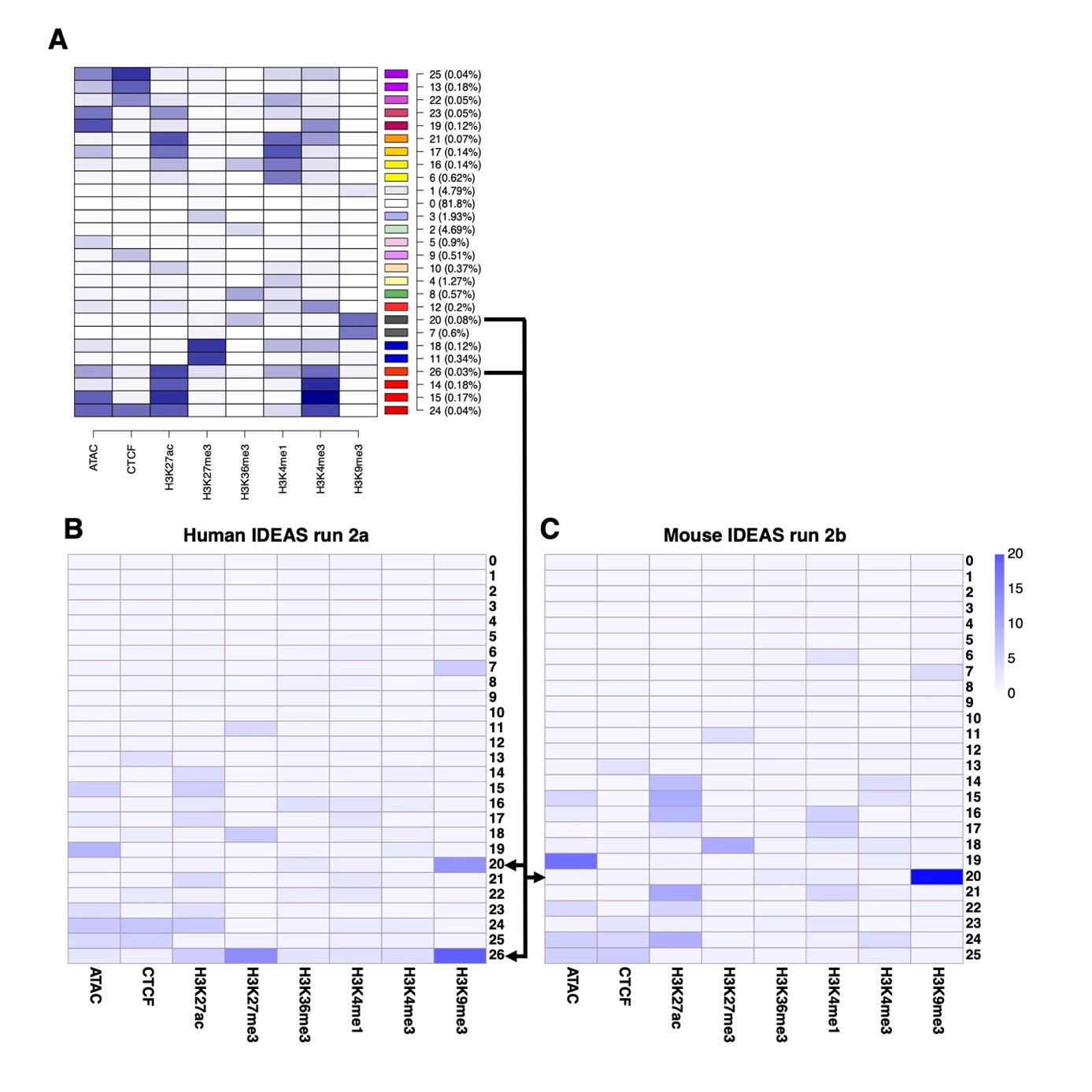

**Supplemental Figure S4**. *Variance in epigenetic signals in the states learned in intermediate stages of the IDEAS models.* (**A**) The emission profiles of an intermediate, 27-state model are represented by different intensities of blue, with darker blue indicating a stronger signal. The two states subsequently removed because of high variance, indicative of heterogeneous states, are indicated by horizontal lines that are the beginnings of arrows. (**B**) and (**C** ) These heatmaps show the coefficient of variance in signal levels of each epigenetic feature in each state across all the genomic bins assigned to that state. Observing a low variance for signal strength in the features for a given state indicates good consistency in the features characteristic of that state; this is the case for almost all the states shown. In contrast, states such as state 20 in both species and state 26 in human, with high variance for one or multiple features, reflect heterogeneity in the epigenetic profiles of the genomic intervals contained within them (Xiang et al. 2020b) and were removed (indicated by the ends of arrows). State 26 was removed after the IDEAS run on human data, and thus it is not present in the heatmap of variance for the mouse data.

After the two rounds of IDEAS runs for the two species, a set of 25 epigenetic states (main Figure 2B) were used as the final joint epigenetic states for both species. To assign these final joint epigenetic states to each genomic location in each cell type in both species, another two rounds of IDEAS runs for the two species were performed in parallel. The proportions of the genomes covered by each state were similar in human and mouse (main Figure 2B). The largest portions of the genomes were in the quiescent state 0, characterized by no significant detectable contribution from any feature. Some of the input ATAC-seq data sets had particularly high signal with low noise (e.g., for HUDEP cells and K562 cells) or relatively high noise (neutrophils), based both on inspection of signal tracks and the calling of higher numbers of peaks in those cell types compared to others (Supplemental Fig. S5). These high ATAC-seq signals may have resulted in the frequent assignment of genomic intervals to state 5 (low ATAC-seq signal alone, light mauve color, main Figure 2B and Supplemental Fig. S5) in HUDEP cells, K562 cells, and human neutrophils (example locus shown in main Figure 2C).

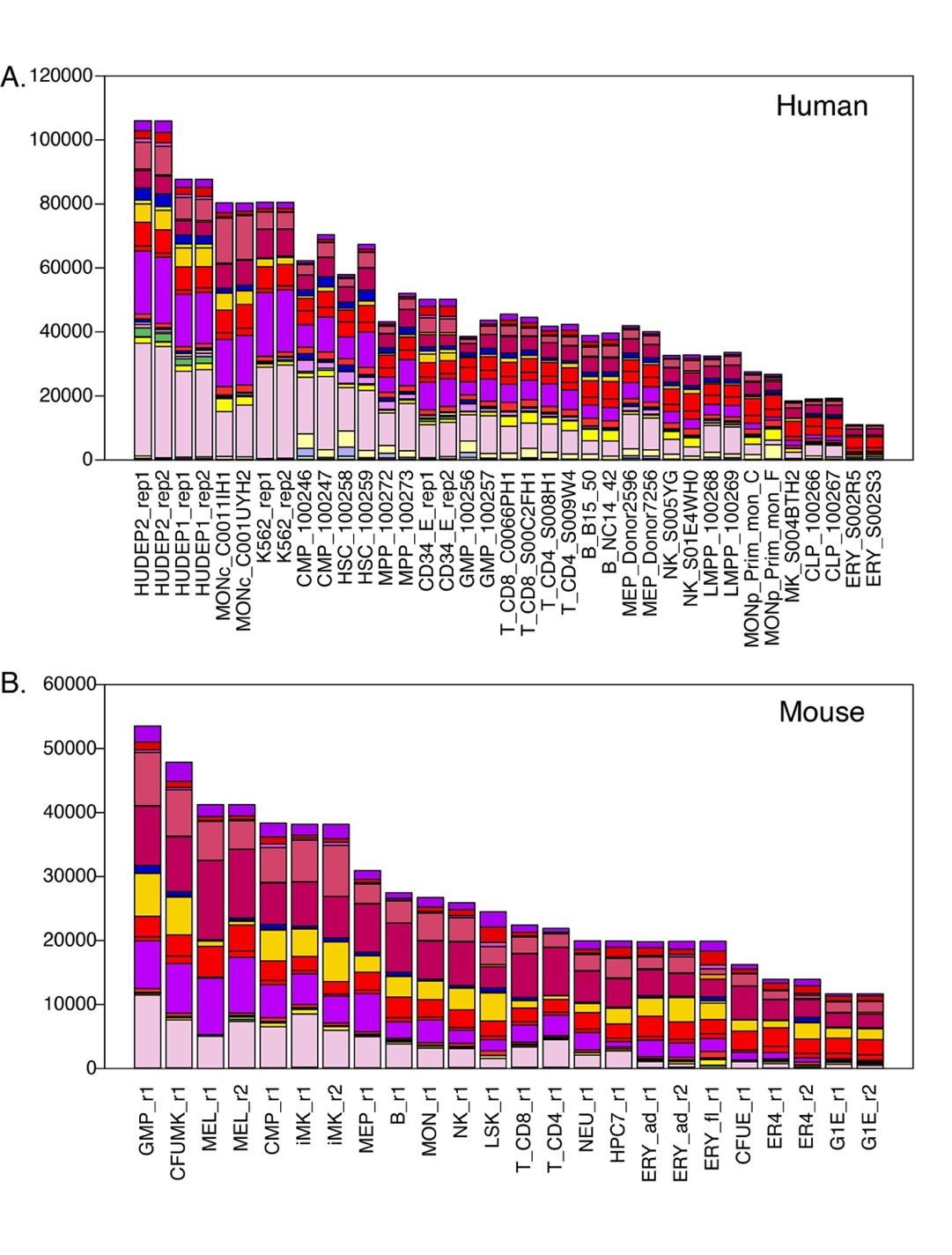

**Supplemental Figure S5**. *Numbers of cCREs assigned to each epigenetic state across cell types.* (**A**) The numbers of cCREs (y-axis) assigned to each epigenetic state in each human blood cell type and replicate are indicated by the color distinctive for each state. The neutrophil cCREs were not included because the very high noise in the ATAC-seq data led to many apparently false positives in the peak calls. **(B)** Same analysis for mouse blood cell types.

*Heterogeneous states and epigenetic states localized to specific regions of the genome*

Some epigenetic states are concentrated primarily in specific locations of the genome. A notable example is the combination of H3K36me3 and H3K9me3 found in chromatin containing genes for zinc finger proteins. SAGA methods, including IDEAS, are designed to find combinations of epigenetic features that consistently found across the genome, and thus uncovering such localized states is challenging. In the joint human-mouse IDEAS modeling described here, no state with an exclusive combination of H3K36me3 and H3K9me3 was found (Fig. 2B). However, our earlier work using IDEAS to integrate epigenetic data in mouse blood cells did discover this state (Xiang et al. 2020b). Also, in our current work the IDEAS method found a state comprised of H3K36me3 and H3K9me3 in both human and mouse. Examination of an intermediate, 27-state model (Supplemental Fig. S4A) shows that state 20 has the H3K36me3 + H3K9me3 profile. However, the H3K9me3 signal across bins assigned to this state had high variance (Supplemental Fig. S4B), which led to its elimination as a heterogeneous state, as described above. A similar state (also state 20) was learned in the mouse data, but again, the high variance in the H3K9me3 signal led to its elimination (Suppl. Fig. S4C).

We have observed that the high variance in signal for one or more features in the heterogeneous states arises from the combining of sets of genomic bins with related but distinct profiles of epigenetic features. While IDEAS and other SAGA methods are successful in learning the abundant, dominant combinations of epigenetic features in their state models, many other, less frequent combinations of features are in the epigenome and the modeling eventually groups them into a heterogeneous state. These high variance, heterogeneous states can be subdivided into component states using *ad hoc* procedures, but we have not developed a systematic way to accomplish this task. Thus, the IDEAS method (and other SAGA methods) can retain an epigenetic state that is limited to a subset of the genome as reproducible, but the ability to do so is at the margins of the ability to distinguish consistent states from heterogeneous ones. We suggest that the ability to systematically handle this challenge is an open issue for SAGA methods.

*Limited impact of number of datasets on the portion of the genome assigned to each state in each cell type*

The analysis in Supplemental Fig. S5 gives some insight into the question of whether the number of available datasets had an impact on the amount of the genome in each cell type assigned to each state. The IDEAS method learns the distribution of epigenetic signals locally in related cell types and uses that information in the inference of epigenetic states, even for cell types with missing data (Zhang and Mahony 2019). Thus, we expected the missing data to not have a large impact. The pattern of availability for epigenetic data in human cells provided some leverage for examining this question further. Specifically, only ATAC-seq data were available for most stem and progenitor cell types in human, as shown in main Fig. 1. If the missing data had a strong impact on the assignment of chromatin states from IDEAS, then the stem and progenitor cells should show a different distribution of states from those in mature cells, for which the feature data was complete or almost complete. We specifically examined the distribution of states in the cCREs. The results show that while the number of cCREs does vary considerably across cell types, the cCREs in each cell type, including the stem and progenitor cells, are assigned to all the epigenetic states (Supplemental Fig. S5). For example, human HSC, MPP, CMP, and GMP cells have roughly average numbers of actuated cCREs (from the ATAC-seq data), and in each cell type those cCREs are assigned to a similar distribution of states to those in many mature cell types, even though no histone modification or CTCF binding data were included in the input to IDEAS. The broad range of state assignments for cCREs in mouse CFUMK (Supplemental Fig. S5B), which also has only chromatin accessibility data (main Fig. 1), also supports the limited impact of missing data on state assignments by IDEAS. Despite this ability to handle missing data, we do point out that the state assignments for epigenomes in human stem and progenitor cells may be less robust compared to those for similar cell types in mouse.

*Prior use of BluePrint Consortium data for integrative segmentation*

An earlier publication used the BluePrint Consortium data on 91 blood cell types as input for a segmentation using ChromHMM (Carrillo-de-Santa-Pau et al. 2017). The results are available at <http://ftp.ebi.ac.uk/pub/databases/blueprint/releases/current_release/homo_sapiens/secondary_analysis/Segmentation_of_ChIP-Seq_data>. Our IDEAS-based segmentation serves as a useful complement to this earlier segmentation. The input data for the two SAGA methods had some notable differences. The histone modifications were the same for the BluePrint data in both SAGAs, but the IDEAS analysis also included ATAC-seq and CTCF ChIP-seq data. The input data for the IDEAS analysis included replicates of datasets in selected blood cell types from healthy donors to the BluePrint project, whereas the ChromHMM segmentation used data from more cell types obtained from both healthy and diseased donors. The ChromHMM SAGA modeled 12 states. The IDEAS SAGA learned more states, and many of them have similar emission profiles to those learned by ChromHMM (for the epigenetic features used as input in common), but the states can include more features and can have states distinguishing strong from weak signals. The current IDEAS SAGA was conducted jointly in both human and mouse, whereas the ChromHMM analysis was only human blood cells.

**Prediction of VISION cCREs using IDEAS-IS**

We define a candidate *cis*-regulatory element, or cCRE, as a DNA interval with a high signal for chromatin accessibility in any cell type (Xiang et al. 2020b). When peaks of accessibility are called independently on different cell types and then combined across cell types, the genomic intervals inferred as peaks can enlarge excessively unless special procedures are employed to prevent expansion (Meuleman et al. 2020; The ENCODE Project Consortium et al. 2020). We reasoned that this expansion could be avoided by using both a combination of all the chromatin accessibility signals and the original data for each cell type as input for modeling across all these datasets to call peaks. We utilized a version of the IDEAS methodology for this purpose, running it in the signal intensity state (IS) mode on ATAC-seq and DNase-seq signals only (Xiang et al. 2021), in contrast to the epigenetic state mode used for integrating data on multiple epigenetic features.

The input data for the peak calling were the ATAC-seq signals for each replicate of each cell type plus a track of combined average ATAC-seq signal. This combined average was computed by averaging the normalized ATAC-seq signal in 200 bp bins for each cell type, and then averaging these values per bin for all cell types (Supplemental Fig. S6). IDEAS in the IS mode learned four signal intensity states, with state 0 being no detectable signal and state 3 being the highest signal state, which were then used to annotate the genomes of all cell types plus the combined cell data. The peaks were called using a hierarchical process designed to find genomic DNA intervals in the high signal intensity states, compared to the local background, both in many cell types and in restricted sets of cell types (Supplemental Fig. S6). Specifically, in the first step (1), the DNA bins in the higher signal states, compared to the local background, in the average track were collected as peaks. If a contiguous series of bins was in higher signal states, indicating a longer accessible region, only the bin(s) in the highest signal state were called as peaks. In the second step (2), bins in a high signal state in individual cell types were included in the set of peaks. The next two steps added bins in a lower signal state, but still above the local background, as peaks, with step (3) adding such bins from the average signal track and step (4) adding such bins from signal tracks from individual cells. Juxtaposed peak calls were combined into a single peak. If replicate determinations were available for chromatin accessibility in a given cell type, the peak call had to be replicated. This procedure resulted in a collection of peak calls that included both peaks present in many cell types as well as those in a single cell type. Furthermore, the preference given to the peaks in the average signal track helped prevent excessive lengthening of the peak calls after combining them.

We employed the same peak-calling procedure for the blood cell epigenomes of human as well as mouse, resulting in 200,342 peaks of chromatin accessibility for human blood cell types and 96,084 peaks for mouse blood cell types; these cCREs are listed in Supplemental Table S3. Unlike the set of accessibility peaks used in earlier work (Xiang et al. 2020b), which were called using the HOMER program (Heinz et al. 2010), all of the IDEAS-IS peaks were in a non-quiescent state in at least one cell type. Thus, the sets of IDEAS-IS peaks comprised the sets of VISION cCREs. The larger number of cCREs called in human than in mouse resulted at least in part from the very high signal in chromatin accessibility data from some human cell lines (HUDEP1, HUDEP2, and K562) and cell types (e.g., monocytes; Supplemental Fig. S5). We found that the ATAC-seq data sets for human neutrophils had an excessive level of noise, such that some extremely long genomic regions were called as peaks. To prevent the noise from this apparent false discovery from compromising further analysis, ATAC-seq peaks that were found only in the human neutrophil data were removed from the set of human blood cell cCREs. Thus, while the human neutrophil ATAC-seq data were included in the joint IDEAS modeling, the peak calls exclusive to human neutrophils were excluded from the 200,342 human blood cell cCREs.

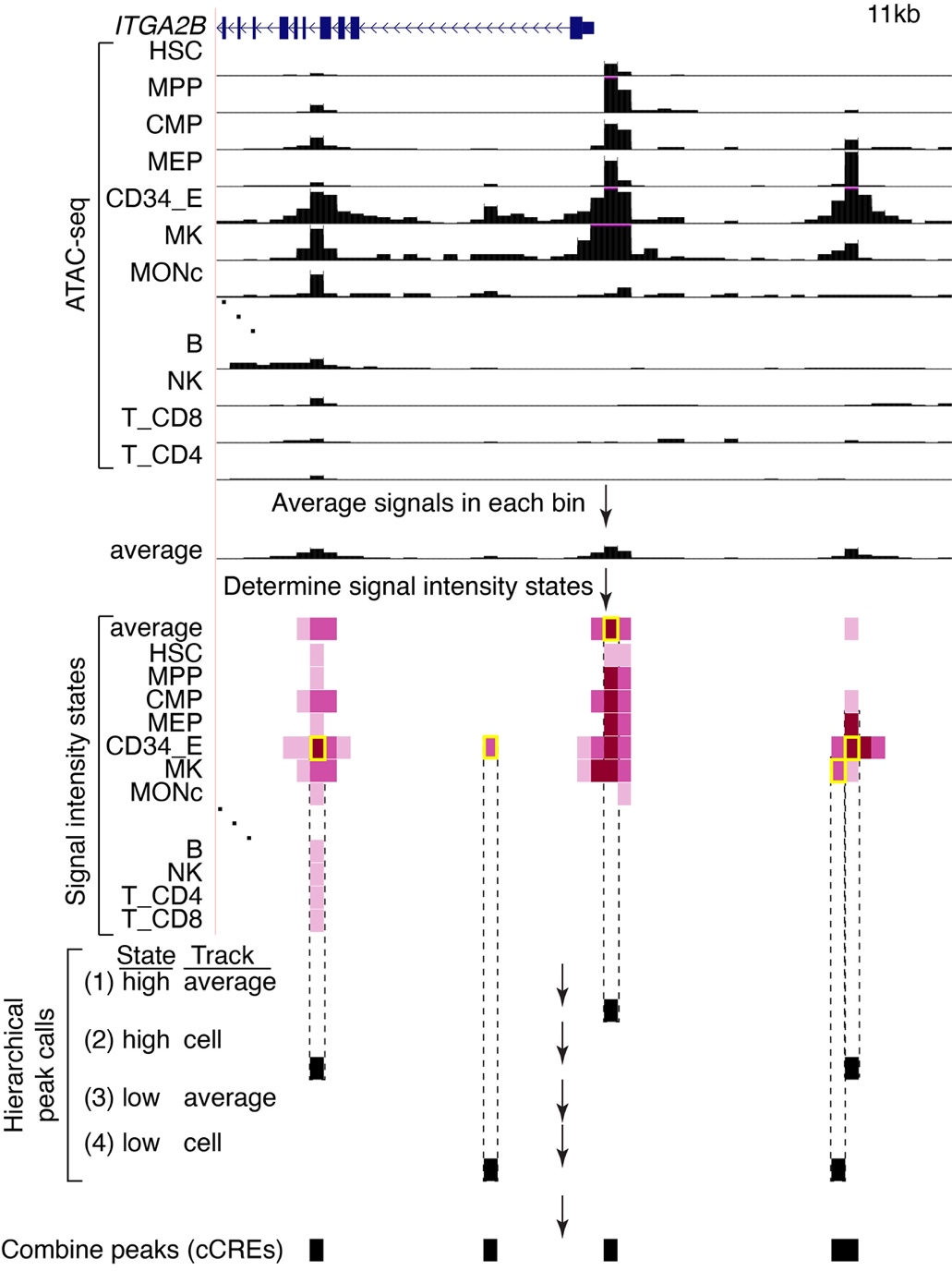

**Supplemental Figure S6.** *Method for calling cCREs using S3V2-IDEAS in the IS mode.* The normalized ATAC-seq signals (expressed as the negative log10 p-value for fitting a negative binomial distribution, signal range 0-10, 200bp bins) are shown for a selected subset of the 39 human biosamples plus the average signal track in an 11kb genomic interval around the transcription start site (TSS) of the *ITGB2B* gene is shown (GRCh38 Chr17:44,384,001-44,395,000). The signal intensity states learned by IDEAS in the IS mode are shown as shades of violet (state 0 is white, darker shades represent higher signal states). Genomic intervals in high signal states were called as peaks (yellow rectangles) in a four-step hierarchical process designed to limit the peak calls to local maxima while also finding cell type-specific peaks (see Methods). Peaks in this genomic region illustrate calls at steps 1, 2, and 4 of the hierarchical process.

**Comparison of ATAC-seq peak calls between IDEAS-IS and MACS3**

We compared the peak calls between IDEAS-IS and MACS3 (Zhang et al. 2008) on ATAC-seq data from 36 experiments in human blood cells that were available in a form that could be analyzed by both tools. Using these ATAC-seq data as input, the IDEAS pipeline operating in the signal intensity state (IS) mode called fewer peaks (200,342 peaks) than did the MACS3 peak caller (277,064, Supplemental Fig. S7A). A majority of both peak sets overlapped, but the MACS3 peak set has almost three times more peaks distinctive to it than did the IDEAS_IS set. The distribution of ATAC-seq signal strengths for the peaks called by both pipelines were similar, but the MACS-only peaks tended to have lower signal strength (Supplemental Fig. S7B). We compared the sets of peaks for their ability to capture orthogonal, function-associated genomic intervals, such as sets of active and predicted enhancers and DNA segments bound by co-activators and CTCF (described in the section “Annotation of VISION cCREs using orthogonal datasets of elements”), and found that the peaks called by IDEAS_IS were significantly more effective than those called by MACS3 (p-value = 2.384e-07; Paired Wilcoxon test; Supplemental Figure S7C).

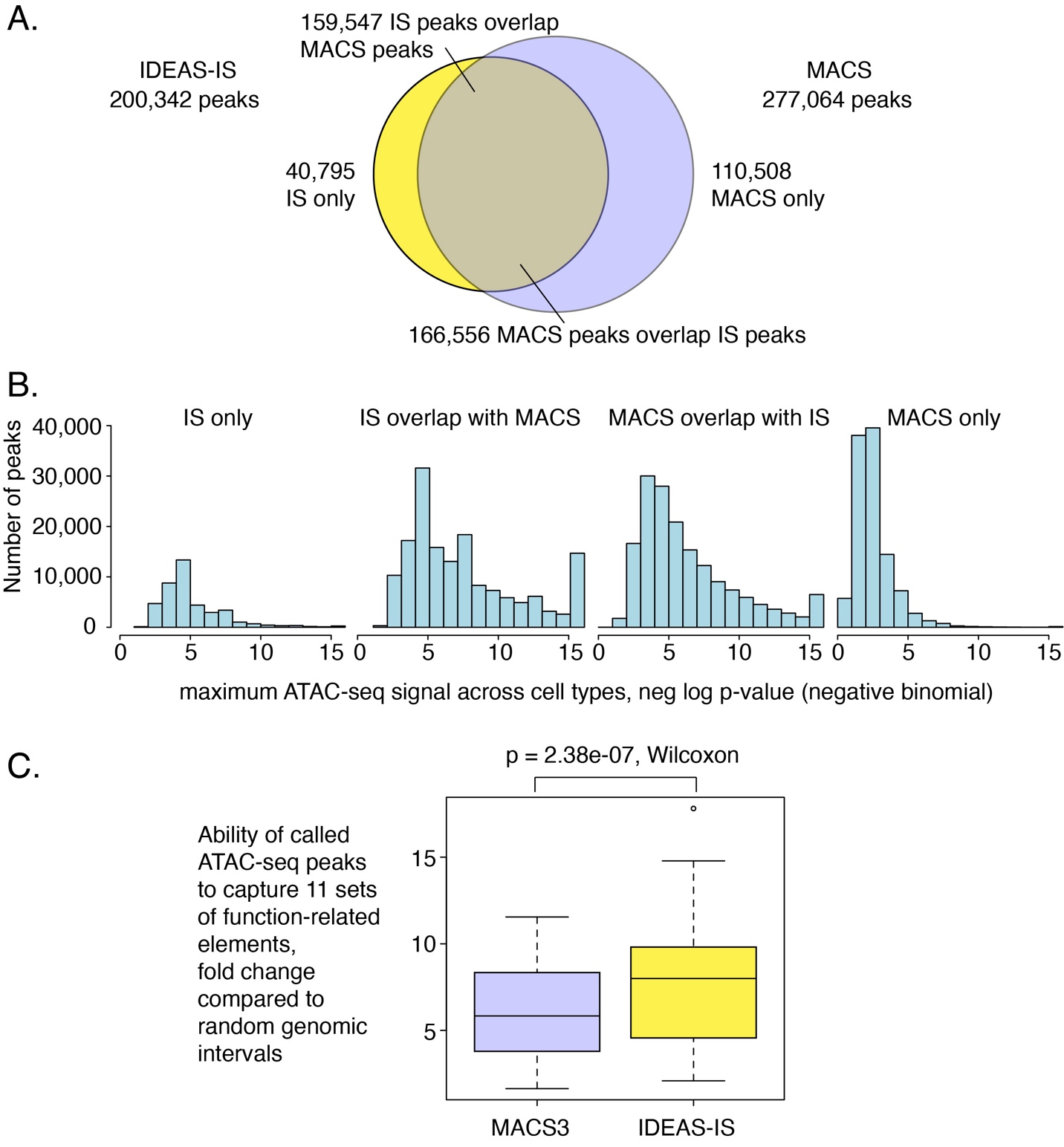

**Supplemental Figure S7.** *Comparison of ATAC-seq peak calls from IDEAS-IS and MACS3.* (A) Overlap in peaks called by each method. (B) Comparison of ATAC-seq signal strength for shared peaks shared and those returned by only one method. (C) Differences in ability of each ATAC-seq peak set to capture elements in 11 orthogonal function-related sets. The fold change (y-axis) is the ratio of number of peaks from the designated peak caller that overlaps with a set of function-associated genomic intervals (11 different sets) to the number of randomly selected matched genomic intervals that overlap with those function-associated sets. Those 11 values for fold change are summarized as boxplots.

**Annotation of VISION cCREs using other datasets of elements**

*Comparison with other large catalogs of cCREs*

The VISION human blood cell cCREs were found to be a sub-set of two large collections of cCREs predicted from ENCODE data on a much larger number of cell types (Supplemental Fig. S8). These large collections were the 926,535 ENCODE cCREs provided in Supplemental Table 10 of reference (The ENCODE Project Consortium et al. 2020) and the 3,591,898 elements in the Index of DNase hypersensitive sites from Meuleman et al. (2020).

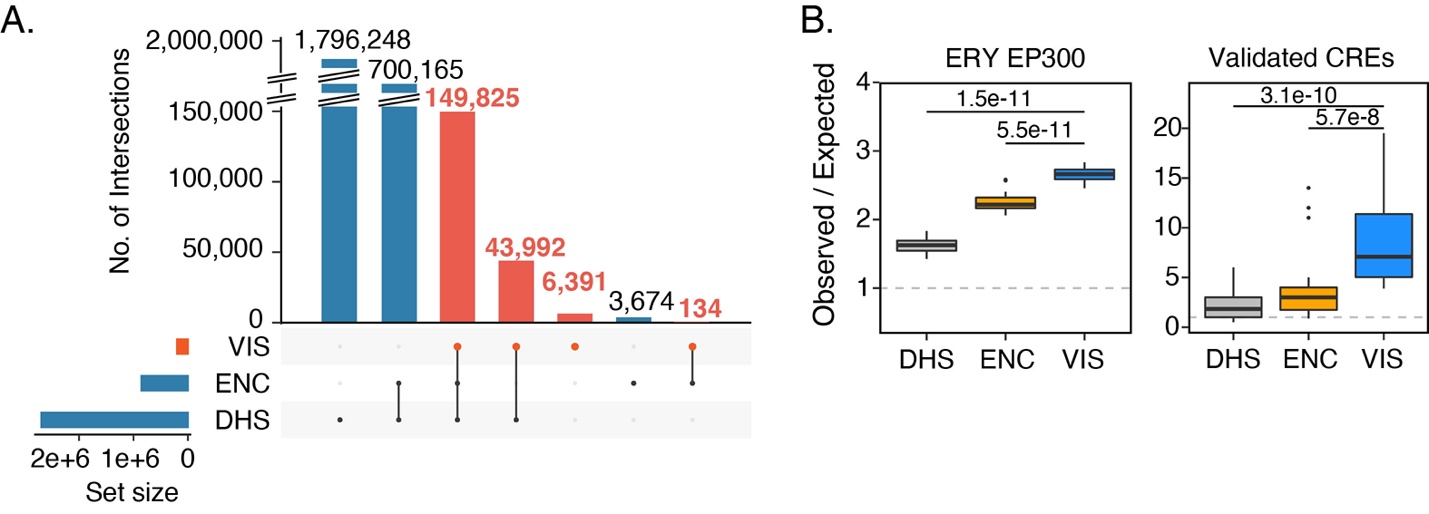

**Supplemental Figure S8**. *VISION cCREs are supported by elements from ENCODE.* (**A**) Overlap of the human VISION cCREs with the Index of consensus DHSs (Meuleman et al. 2020) and CRE catalog from ENCODE phase 3 (The ENCODE Project Consortium et al. 2020), displayed as an UpSet plot. (**B**) Enrichments in the three large cCRE catalogs for overlap with orthogonal sets of elements indicative of regulatory function. Abbreviations are DHS = Index of consensus DHSs, ENC = CRE catalog from ENCODE phase 3, VIS= VISION cCREs, ERY EP300 = peaks for EP300 ChIP-seq in three erythroid cell types or lines (Supplemental Table S5), Validated CREs = a curated set of human erythroid regulatory elements gathered from the literature (Supplemental Table S4).

*Annotation of VISION cCREs by overlap with elements defined by orthogonal data*

Additional, orthogonal data sets that were not included in the prediction and epigenetic state annotation of VISION cCREs, but which annotate potential roles of human genomic intervals in transcriptional regulation (CREs) or in chromatin structure (chromatin architecture), were curated from the literature and associated databases. Details of these orthogonal data sets, including references and data sources, are presented in Supplemental Table S5. The organization of the data sets into supersets of elements related to CREs and related to structure and genome architecture is illustrated in Supplemental Fig. S9A.

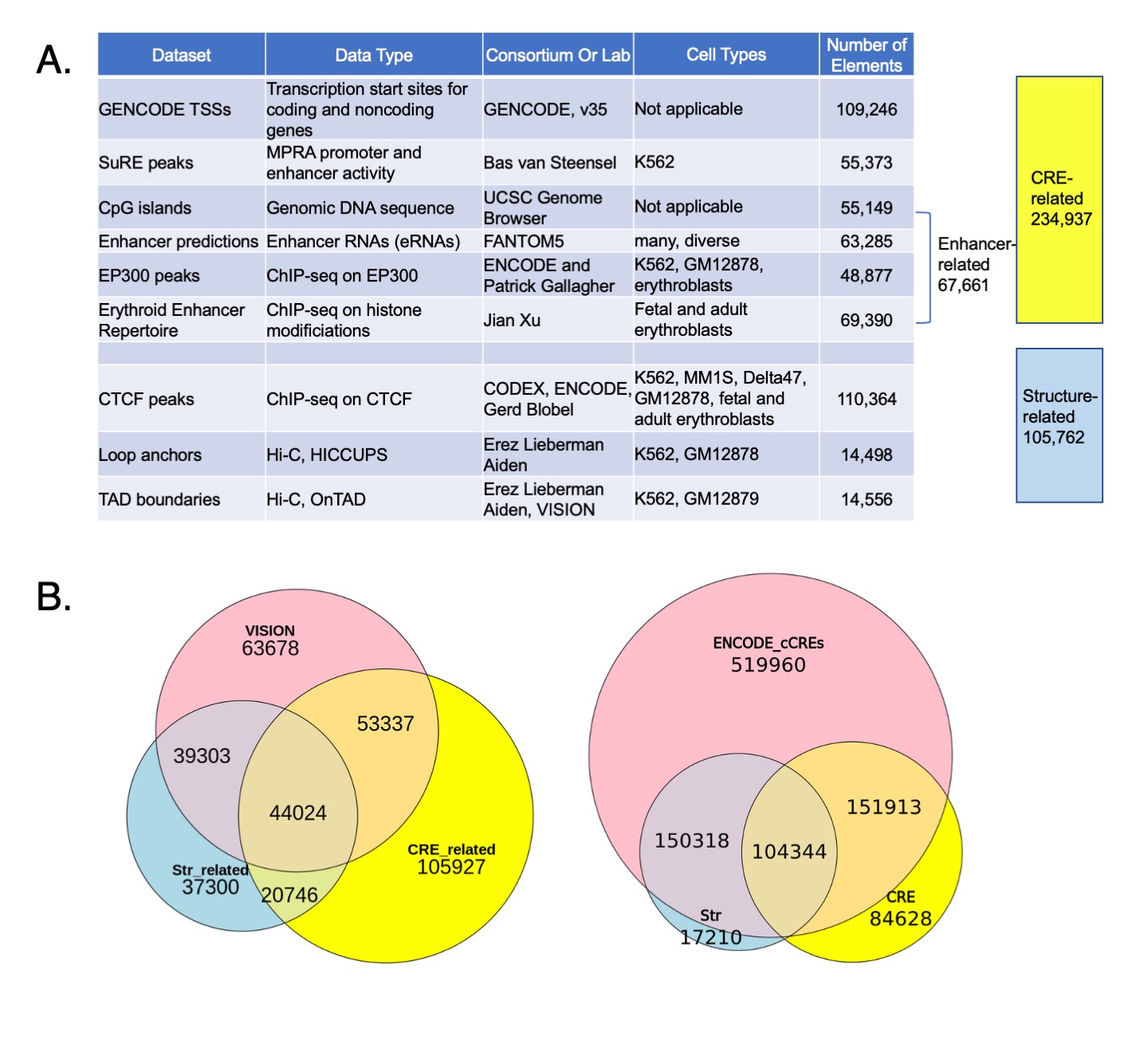

**Supplemental Figure S9.** *Relationships among orthogonal datasets of likely functional elements, VISION cCREs, and ENCODE cCREs.*

**A.** Orthogonal sets of regulatory and structural genomic elements. Organization into supersets of elements is indicated on the right, along with the numbers of elements in each superset. Additional information is in Supplemental Table S5. **B.** Venn diagram showing the overlaps of the ENCODE cCRE catalog with the orthogonal data sets of elements related to cCRE function or structure in chromatin (right) along with a comparison with Venn diagram using the VISION cCREs (left, which is Fig. 3A).

The orthogonal sets of CREs included TSSs from the GENCODE basic gene set (Frankish et al. 2021), peaks from the Survey of Regulatory Elements (SuRE), which is a massively parallel reporter assay that reveals both promoter and enhancer activity, in K562 cells (van Arensbergen et al. 2017), unmasked CpG islands downloaded from UCSC genome browser (Nassar et al. 2023), and a group of enhancer-related elements. The latter group of enhancer-related elements were a combination of three sets: (1) enhancers predicted from eRNAs in hundreds of human cell types (Andersson et al. 2014; https://fantom.gsc.riken.jp/5/datafiles/reprocessed/hg38_latest/extra/enhancer/), (2) a combined set of EP300 ChIP-seq peaks from K562 and GM12878 cell lines (The ENCODE Project Consortium et al. 2020) and erythroblasts (Su et al. 2013), and (3) the erythroid Enhancer Repertoire deduced from histone modification data (Huang et al. 2016). While the Survey of Regulatory Elements (SuRE) is a massively parallel reporter assay designed to detect promoter activity (van Arensbergen et al. 2017), many enhancers also generate a peak in this assay, albeit weaker than those for promoters. The chromatin structure category included CTCF-occupied DNA segments (CTCF OSs), chromatin loop anchors, and TAD boundaries in primary blood cells and related cell lines. The set of CTCF OSs was generated by combining peaks from ChIP-seq experiments in human fetal and adult erythroblasts (Huang et al. 2017), and from K562, MM1S, Delta47, and GM12878 cell lines (Sánchez-Castillo et al. 2015; The ENCODE Project Consortium et al. 2020). Loop anchors determined by HICCUPS from Hi-C data in K562 and GM12878 cell lines (Rao et al. 2014) were downloaded from GEO [accession GSE63525] and combined. TAD boundaries were called by OnTAD (An et al. 2019) using Hi-C data at 10kb resolution in K562 and GM12878 cell lines (Rao et al. 2014). Even though CTCF ChIP-seq data were used both in the IDEAS epigenetic state modeling and in the structure-related datasets, no CTCF ChIP-seq dataset was used in both. Data for K562 CTCF ChIP-seq is in both collections, but the orthogonal data came from Pugacheva et al. (2015) while the data used in the IDEAS modeling came from the ENCODE consortium.

Pairwise overlaps between these orthogonal sets of elements and the VISION cCREs were computed using the bedtool intersect tool (-u option) (Quinlan and Hall 2010). Intersections among multiple datasets were computed using the intervene tool, version 0.6.5 (Khan and Mathelier 2017), and displayed in UpSet plots using the UpSetR package (Conway et al. 2017) and 3-way Venn diagrams using the eulerr package (https://CRAN.R-project.org/package=eulerr).

Ten different randomly chosen sets of human genomic DNA intervals, matched with the cCREs in length and G+C content, were generated by the script in the following link <https://github.com/YenLab/Tn5InsertPrefer/blob/main/StandaloneScripts/Negetive_sequence_matched_length>. Enrichments of overlaps between the VISION cCREs relative to those in the random set of intervals were determined by computing overlaps (using tools described above) between the sets of function- and structure-related elements and each of the ten sets of random intervals, dividing the number of overlapped cCREs by the number of overlapped random intervals (ten times), and using the average of the ten quotients as the enrichment. All ten results were shown in boxplots.

The catalog of cCREs released by ENCODE (The ENCODE Project Consortium et al. 2020) contained about 920,000 cCREs in the human genome. This catalog was determined from epigenetic data on a much larger and diverse set of cell types and cell lines compared to the blood cell types examined in the VISION project. Thus, the size of the cCRE catalog was much larger for ENCODE than for VISION. When that larger number of ENCODE cCREs was intersected with the orthogonal sets of cCRE-related and structure-related cCREs, we find that the ENCODE set overlapped with larger numbers of the orthogonally defined elements compared to the overlaps by the VISION catalog, but a substantial number of the ENCODE cCREs (almost 520,000) did not overlap (Supplemental Fig. S9B). These results likely reflect the larger number of cell types examined in ENCODE, most of which are distinct from the blood cells that were the focus of the orthogonal sets.

Considering the VISION cCREs that intersected with both structure- and CRE-related elements, the largest group are those that overlap with enhancers and CTCF OSs, followed by enhancers and loop anchors, and then promoter-like elements and CTCF OSs (Supplemental Fig. S10).

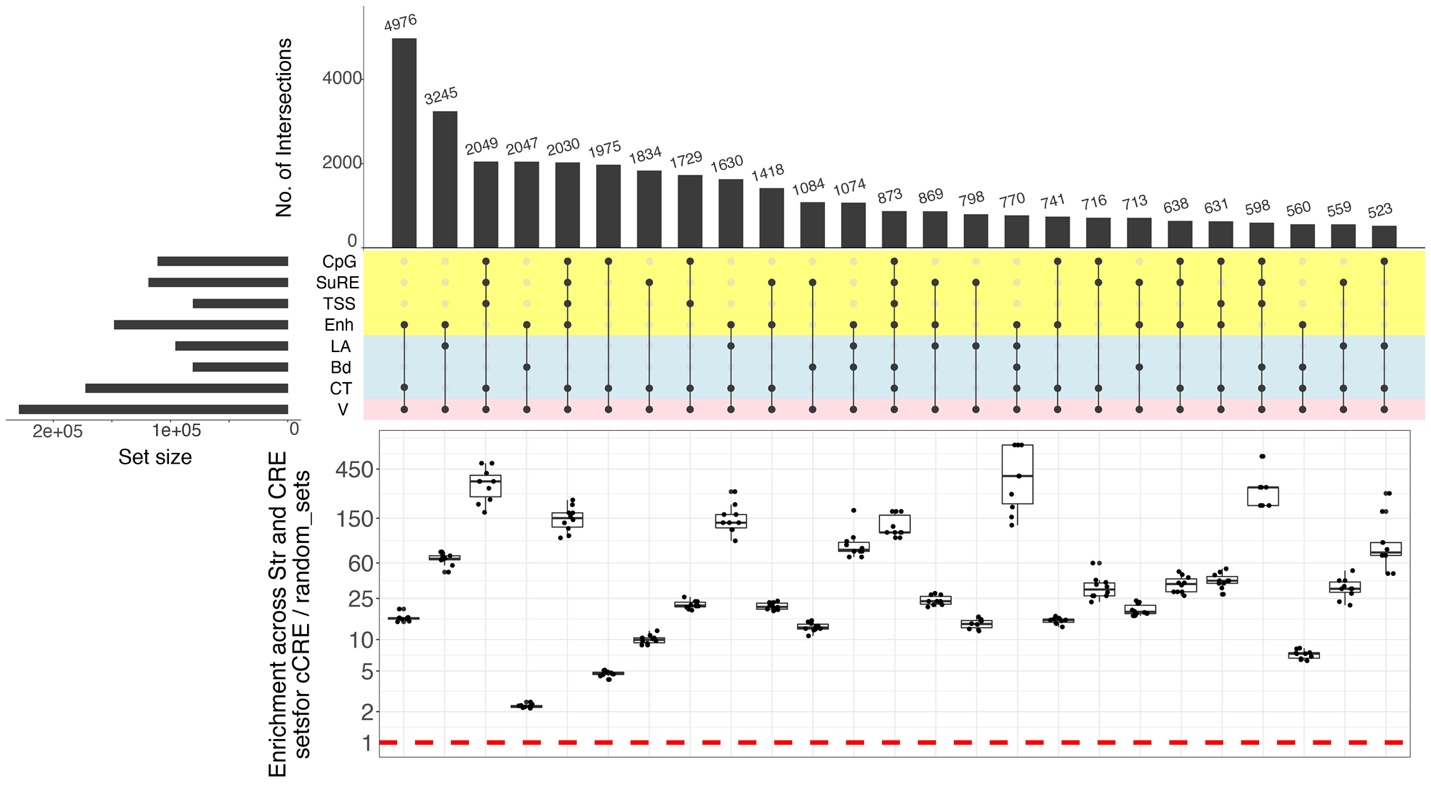

**Supplemental Figure S10**. *Numbers and enrichment of VISION cCREs that overlap with both structure and CRE-related elements*

The human VISION human cCREs were also compared to two sets of known CREs (main Fig. 3E). One was a compilation of 109 demonstrated regulatory elements in blood cells from the literature (Supplemental Table S4). The other set was 664 enhancers with target genes, determined by a high throughput mutagenesis and eQTL analysis in K562 cells (Gasperini et al. 2019).

**Enrichment of genetic variants for blood cell related traits in the VISION human cCRE collection**

*Overlaps of cCREs with trait-associated SNPs*

Most genetic variants associated with complex phenotypes occur in noncoding regions of the genome (Hindorff et al. 2009), and candidate regulatory elements are strongly enriched for such variants (Maurano et al. 2012; The ENCODE Project Consortium 2012). Thus, collections of high quality predictions of CREs could provide insights into the functional mechanisms by which noncoding variants mediate phenotypic variation (Hardison 2012). We reasoned that our collection of cCREs would be informative for interpreting non-coding variants influencing blood cell traits and blood diseases. Therefore, we computed the overlap of the human VISION cCREs with several sets of phenotype-associated variants from the NHGRI-EBI GWAS Catalog (Buniello et al. 2019), specifically those associated with red blood cell traits, immune diseases, and brain measurements (as a negative control). The number of single nucleotide polymorphisms (SNPs) associated with red cell traits overlapping with VISION cCREs was much higher than the number overlapping a randomly selected set of genomic intervals matched for GC content and length (Supplemental Fig. S11A). Refining the analysis to cCREs that are actuated in each cell type, we found that the cCREs actuated in erythroid and related cell types showed high overlap with SNPs associated with red cell traits (Supplemental Fig. S11B). Conversely, SNPs associated with immune disease showed high overlap with cCREs actuated in lymphoid cells (Supplemental Fig. S11C), whereas SNPs associated with brain measurements showed infrequent overlap with cCREs actuated in any blood cell type (Supplemental Fig. S11B).

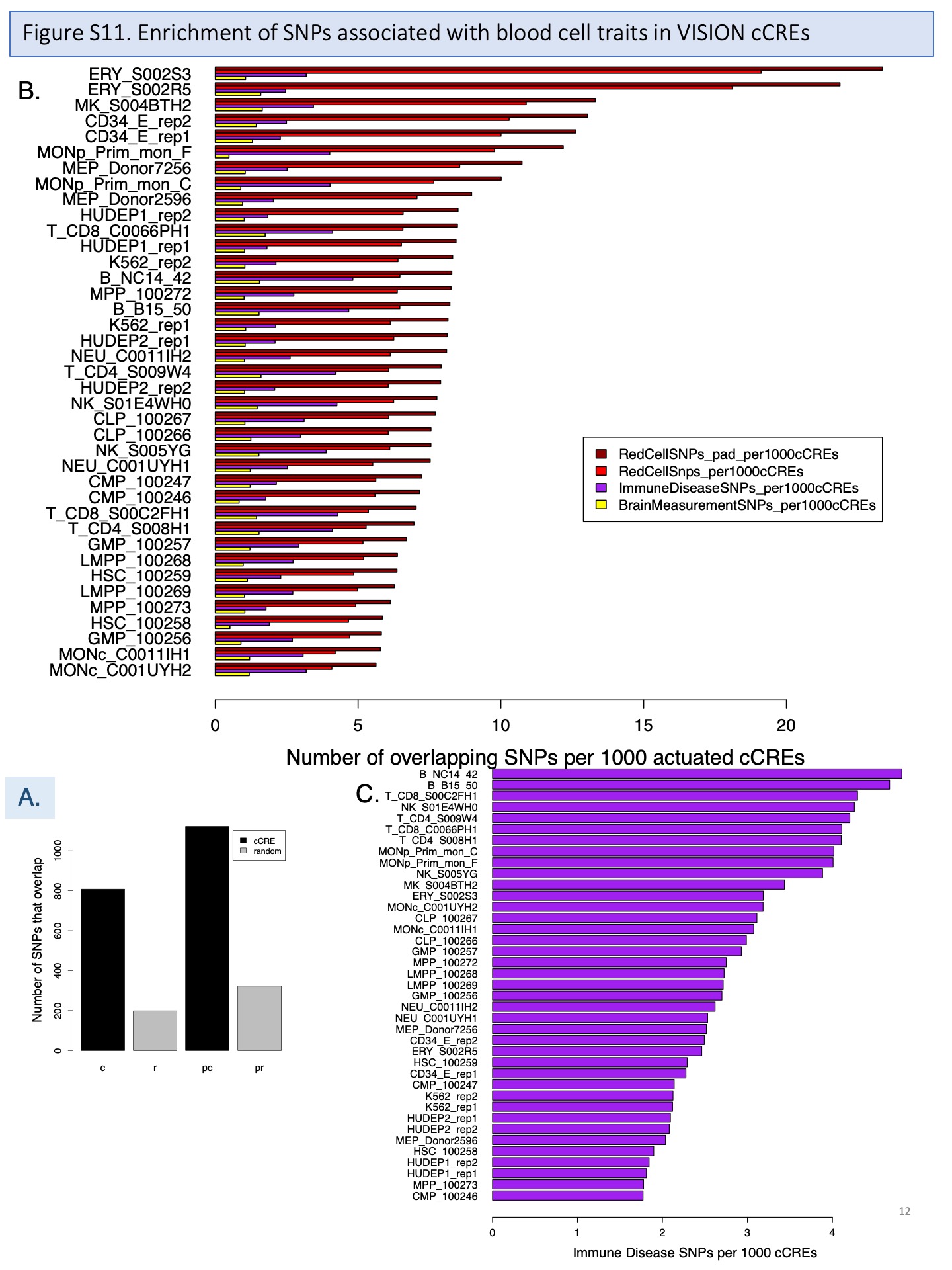

**Supplemental Figure S11.** *Overlap of SNPs associated with blood cell phenotypes with VISION cCREs***.** (**A.**) Overlaps with all VISION cCREs (c) compared with matched random DNA intervals (r), without or with (pc and pr) padding of each SNP to a 200bp interval. The higher numbers of SNPs captured by cCREs is significantly different from the numbers captured by random sets (chi-squared test; p<0.0001 for both cCREs alone and for padded cCREs).

(**B.**) Overlaps of SNPs with cCREs actuated in specific cell types. (**C.**) Overlaps of SNPs associated with immune disease with cCREs actuated in specific cell types.

*Stratified linkage disequilibrium score regression (sLDSC)*

When measuring enrichments from GWAS data, it is important to consider the haplotype structure of human genomes, whereby association signals measured at assayed genetic markers likely reflect an indirect effect driven by linkage disequilibrium (LD) with a causal variant (that may or may not have been genotyped). Stratified linkage disequilibrium score regression (sLDSC, Finucane et al. 2015) offers one principled approach to account for LD structure and estimate the proportion of heritability of each trait explained by a given genomic annotation. We applied sLDSC to quantify the enrichment of heritability in many traits from the UK Biobank (UKBB) GWAS (Ge et al. 2017 and http://www.nealelab.is/uk-biobank/) within the VISION cCREs relative to the rest of the genome. The UKBB (Ge et al. 2017 and http://www.nealelab.is/uk-biobank/) is a database comprising genotypic data as well as data for several medical traits from over 400,000 individuals, and GWAS summary statistic results are publicly available for a number of these traits, stratified by sex. We used all 587 sex-stratified traits for which inverse rank-normalized data was available (representing 295 unique traits) in our analysis, including 54 traits labeled “blood count”-related traits by UKBB, 60 traits labeled “blood biochemistry”-related, and 473 traits that are not blood-related. Blood count-related traits reflect cell morphology and number while blood biochemistry-related traits reflect the concentrations of certain proteins and metabolic products. (Note: hemoglobin concentration is an exception and considered a blood count trait).

For each of these 587 traits, we used sLDSC (Finucane et al. 2015) to quantify the extent to which our cCRE annotation is enriched in the heritability of the trait. Using SNPs within some window of the annotation, this approach regresses the GWAS chi square summary statistic (for the focal trait) of these SNPs onto the LD scores of the SNPs with respect to the annotation. The LD score of a SNP reflects the extent to which that SNP is in linkage disequilibrium with the annotation. If the annotation is associated with the focal trait, we expect a linear relationship between the LD scores of the tested SNPs with the annotation and the chi square values of those SNPs. The slope of the regression line is an estimate of the SNP heritability of the trait with respect to the annotation. By dividing this estimate by the overall SNP heritability of the trait, we obtain an estimate of the proportion of heritability explained by the annotation. Finally, dividing this by the portion of SNPs falling within the annotation provides an estimate of the enrichment of that annotation in heritability of the focal trait.

The sLDSC tool recommends using a set of SNPs from HapMap 3 for analysis, and because these SNPs are reported on GRCh37, we first lifted over (Hinrichs et al. 2006) our cCRE annotations from GRCh38 to GRCh37. A total of 826 cCREs (0.4% of all cCREs) failed to liftOver and were excluded from analysis. Using LDSC v1.0.1, and with these lifted over annotations, we first computed LD scores for HapMap 3 SNPs within 1 cM of cCRE annotations. Then, for each of the 587 UKBB traits, we performed sLDSC, regressing the trait summary statistics onto the LD scores. From this, we obtain an estimate of the enrichment of the cCRE annotation in the heritability of each trait.

The results of the sLDSC analysis for the human VISION catalog of cCREs were reported in main text (main Fig. 3F), emphasizing the remarkable enrichment in heritability for blood count traits. A set of 58 blood biochemistry traits were not enriched for heritability in the human VISION cCREs. We note that those blood biochemistry traits were from screenings of metabolites, proteins, and enzymes that were not produced in blood cells, but rather by the liver (e.g., albumin, alkaline phosphatase, alanine aminotransferase, apolipoproteins, aspartate aminotransferase, bilirubin, urea, cholesterol), kidney (e.g., creatinine), or other organs. While these traits were labeled as blood-related traits by the UKBB, they are largely controlled by organs, tissues, and cell types that we did not assay when developing the VISION CRE annotation.

**Estimation of the impact of epigenetic states and cCREs on gene expression**

*Summary of the calculation of β coefficients for epigenetic states and esRPs for cCREs*

In order to use the categorical state assignments to estimate the impact of each cCRE in each cell type on gene expression, we applied a modified version of the iterative multivariate linear regression model (MVLR) developed previously (Xiang et al. 2020b) to quantify the biological functions of each epigenetic state in terms of regulating gene expression. In this model, we introduced two measurements: β coefficients for each epigenetic state and an esRP (epigenetic state Regulatory Potential) score for each cCRE in each cell type or sample. The biological interpretation of the two measurements are as follows. The β coefficients measure the contribution of each epigenetic state to the expression of local genes; they are calculated in a multivariate regression evaluating how changes in the coverage of cCREs and promoters by each epigenetic state across cell types impact expression levels. The esRP score measures the contribution of individual cCREs on regulating its target gene’s expression level; it is calculated from the overall epigenetic state coverage of the cCRE in each cell type (Figure 4B). In contrast to our previous modeling (Xiang et al. 2020b), our current model does not aim to identify the likely target gene(s) for each cCRE; that will be the subject of a subsequent report. In brief, for the current regression model, the epigenetic state coverage was computed on all cCREs and promoter regions within 50 kb on both sides of the TSS of each gene (an interval of 100kb). We first calculated the β coefficients of the promoter intervals and distal cCREs as separate terms in the regression model. For further analyses and visualization, including computation of the esRP scores, the β coefficients of each state were merged into a single value that was the average of the β coefficients for promoters and for distal cCREs. A more detailed presentation on the calculations of the β coefficients and the esRPs is in the following subsection.

*Detailed methods for the calculation of β coefficients and esRPs*

We identified 14 cell types matched between human and mouse with RNA-seq datasets in the VISION project. For each gene, the correlation coefficient was calculated between the two vectors of 14 values for log2(TPM+1), generating one correlation coefficient value per cell type per gene. When calculating the correlation coefficients, we added random noise (mean=0, sd=1) to the raw values to avoid high correlation coefficients created between vectors with low signals.

In the MVLR model, we assumed the contribution of epigenetic states on gene regulation are different at the promoter regions and the cCRE regions. For the independent variables in the model, we thus separated the information of two classes of regions as follows: one class is the epigenetic state coverage at the cCRE regions (cCREs within TSS +/- 50 kb window) and the other class is the epigenetic state coverage at the promoter regions (TSS +/- 1 kb window). The promoter region was the entire 2Kb window centered on the TSS, regardless of whether the DNA was called as a cCRE. In contrast, the cCRE regions were only the cCREs, not any intervening DNA, within the 100Kb region centered on the TSS. The cCRE regions included cCREs within the promoter region.

For a ${cCRE}_{i}$, the state coverage ($L_{i,j,c}$) for ${Epigenetic\_State}_{j}$ in ${Cell\_Type}_{c}$ is defined as the number of base-pairs covered by ${Epigenetic\_State}_{j}$ in ${cCRE}_{i}$ in ${Cell\_Type}_{c}$. For each ${Gene}_{g}$, the overall coverage of ${Epigenetic\_State}_{j}$ at the cCRE regions (${CC}_{g,j,c}$) is defined as the sum of ${Epigenetic\_State}_{j}$*’*s coverages of all individual cCREs within the TSS +/- 50 kb window:

$${CC}_{g,j,c}=\sum_{i} L_{i,j,c}$$

For the promoter regions, the state coverage (${CP}_{g,j,c}$) for ${Epigenetic\_State}_{j}$ at ${Gene}_{g}$ in ${Cell\_Type}_{c}$ is directly defined as the number of base-pairs covered by ${Epigenetic\_State}_{j}$ within the TSS +/- 1 kb window.

Previously, we directly used the ${CP}_{g,j,c}$ and the ${CC}_{g,j,c}$ as the input features for our iterative regression model (Xiang et al. 2020b). However, the state coverage of epigenetic states with related functions are likely collinear (i.e., are positively or negatively correlated), which can lead to unstable estimation for the β coefficients in the regression model. To avoid this problem, we first transformed the original input features matrix into orthogonal principal components (PCs), and then use the PCs that can explain 95% of the variance in the original data matrix (${CP}_{g,j,c}$ and ${CC}_{g,j,c}$) as the input feature matrix for the regression model.

We first matched the mean and standard deviation of the ${CC}_{g,j,c}$ matrix to the ${CP}_{g,j,c}$ matrix to generate a Standardized cCRE state Coverage, or *SCC*, computed by the following equation.

$${SCC}_{g,j,c}={(CC}_{g,j,c}-mean(CC))\times\frac{sd(CP)}{sd(CC)}+mean(CP)$$

We next combined the $SCC$ matrix and the $CP$ matrix as $C$ matrix, and then used PCA to transform the $C$ matrix into a $PC$ matrix that can explain 95% of the total variance in the $C$ matrix.

$$C=cbind(CP, SCC)$$

$$PC=C\otimes R$$

where *cbind* refers to the operation of combining the matrices, and $R$ represents the rotation matrix in the PCA analysis.

For the multivariate linear regression model, the values in the $PC$ matrix were used as the independent variables. The quantile normalized TPM values of RNA-seq data for protein coding genes were used as the dependent variable ($Y$). The signal of both RNA-seq and the state coverage were transformed into a natural log scale for this analysis.

$$Y={\beta_{0}PC+\alpha}$$

$$\beta_{ES}=R\otimes\beta_{0}$$

where the $\beta_{0}$ represents the coefficient vector for all PCs learnt by the MVLR model, and the $\beta_{ES}$ represents the coefficient vector for all epigenetic states transformed by the rotation matrix ($R$) in the PCA analysis. The biological interpretation of the $\beta_{ES\_j}$ is the contribution of one unit increase of ${Epigenetic\_State}_{j}$’s coverage on regulating the target gene’s expression. In this part, we used only the 24 non-quiescent epigenetic states in the model, so the $\beta_{ES\_0}$ for the quiescent state (state 0) naturally becomes 0, which means that we infer no contribution on gene expression regulation. Since we used the information of the promoter regions and the cCRE regions as two separate terms in the MVLR model, $\beta_{ES}$ is a vector with 50 elements, which can be further separated into 25 $\beta_{ES}$ values for promoter regions ($\beta_{ES\_P}$) and 25 $\beta_{ES}$ values for all cCRE regions ($\beta_{ES\_c}$). The MVLR model was trained multiple times by a leave-one-cell-type-out strategy. For downstream analysis, we combined the $\beta_{ES}$ values for proximal cCREs and distal cCREs to generate an average $\beta_{ES}$ coefficient. These are shown in panel A of Figure 4.

After we obtained the $\beta_{ES}$ coefficient vector, we further used the $\beta_{ES}$ coefficient vector to define the epigenetic-state gene expression regulatory potential (${esRP}_{i,c}$) for each ${cCRE}_{i}$ in ${Cell\_Type}_{c}$.

For ${cCRE}_{i}$ within the promoter regions, the ${esRP}_{i,c}$ is defined as follows:

$${esRP}_{i,c}=\sum_{j} \beta_{ES\_C\_j}\times L_{i,j,c}+\sum_{j} \beta_{ES\_P\_j}\times L_{i,j,c}$$

For ${cCRE}_{i}$ NOT at the promoter regions, the ${esRP}_{i,c}$ is defined as follows:

$${esRP}_{i,c}=\sum_{j} \beta_{ES\_C\_j}\times L_{i,j,c}$$

The biological interpretation of ${esRP}_{i,c}$ is the contribution of ${cCRE}_{i}$ on regulating its target gene’s expression level based on its overall epigenetic state coverage in ${Cell\_Type}_{c}$.

Since it is unlikely that every cCRE within the TSS +/- 50 kb window would be used to regulate the target gene, it is reasonable to filter out some cCREs that are less likely to be relevant. We hypothesized the esRP pattern of the relevant cCREs should be more correlated with the target gene’s expression pattern across all cell-types. Therefore, we filtered the cCREs by the correlation between the cCRE’s esRP vector and the target gene’s expression vector across all cell-types, requiring a correlation coefficient *r* < 0.2 in order to be retained (Xiang et al. 2020b). After the filtering step, we recalculated the $CC$ matrix and re-trained the regression model. We iterated this filtering and re-training step two times and updated the $\beta_{ES\_C}$and $\beta_{ES\_P}$ vectors for both human and mouse. To simplify the final output, we first took the average of the two species’ $\beta_{ES\_C}$and $\beta_{ES\_P}$ vector. We then defined the final contribution ($\beta_{ES}$) of the epigenetic states on gene regulation as the sum of their contributions at both promoter regions ($\beta_{ES\_P}$) and cCRE regions ($\beta_{ES\_C}$):

$$\beta_{ES}=\beta_{ES\_C}+\beta_{ES\_P}\times\frac{max(\beta_{ES\_C})}{max(\beta_{ES\_P})}$$

where the $\frac{max(\beta_{ES\_C})}{max(\beta_{ES\_P})}$ is a factor to scale the contributions at the promoter regions ($\beta_{ES\_P}$) and the cCRE regions ($\beta_{ES\_C}$) to a similar level.

The $\beta_{ES}$ vector, and the corresponding esRP matrix for all cCREs across all cell-types’ samples in both species, were used as the final outputs for downstream analysis.

*Potential advantage of esRP to consideration of each feature marginally*

This procedure of using epigenetic states from integrative modeling of features to estimate beta-coefficients and esRP scores raises the question of whether there is an advantage to the joint estimate described here compared to directly considering each feature marginally, since the latter approach would avoid the issue of co-linearity. In the series of steps to estimate the beta-coefficients for epigenetic states and the corresponding esRP scores of cCREs based on state coverage at proximal and distal regions, we reduced the dimensionality of the input matrix at the regression step by replacing the state coverage with principal component (PC) values to reduce the impact of collinearity, as described above. While the alternative approach of using the marginal effects of each feature may have merit for future development, we offer three points about the potential advantages of utilizing principal components of the esRP matrix.

(i) The esRP scores (and PCs) allow annotation of the direction of transcriptional regulation (activation versus repression) and magnitudes for all epigenetic states, thereby adding more functional meaning to the categorical epigenetic state labels.

(ii) Instead of examining each feature marginally, the esRP scores may capture (at least in part) the interactions and joint effects of all features in each state. The esRP scores may better handle some complexities of transcription regulation because the multiple epigenetic features often work in concert.

(iii) Another advantage of esRP is the reduced impact of missing data on this score. The computation of esRP scores uses IDEAS epigenetic state tracks as the input, and the modeling and assignment of epigenetic states in IDEASs utilize data from locally similar cell types to reduce the impact of missing data in an individual cell type. As a result, the esRP score can be computed, and it is comparable for all cell types, even for the ones with incomplete data. In contrast, alternative methods, like estimating coefficients marginally for each epigenetic feature, make it challenging to compare between cell types with missing data and those with complete data sets.

*Impact of removing cCREs with a low correlation between states and gene expression across cell types*

In the estimation of esRP scores, we removed cCREs with low correlation values; specifically, we filtered the cCREs by the correlation between the cCRE’s esRP vector and the target gene’s expression vector across all cell-types, requiring a correlation coefficient *r* < 0.2 in order to be retained. This step raised the question of what the impact may be, especially if a state is primarily associated with constitutive gene activity. Indeed, if a state were exclusively associated with high level expression, with little change in the expression level, in all cell types, then the impact of the state on expression would be under-estimated or missed. However, in practice almost every epigenetic state is associated with at least some genes with sufficient variation in expression levels across cell types to be included in the regression modeling. Examination of the beta coefficients computed for the epigenetic states (Fig. 4A in the main text) shows that almost all states have a positive or negative score that is not widely divergent from expectation based on the established associations of the epigenetic marks with expression. We infer that the states associated with constitutive activity (for some genes) are also associated with cell type-specific activity (for other genes), and thus we still see a signal in the regression. Two of the states do have very low scores (close to zero), the quiescent state 0 and the promoter-enhancer-associated state 12. Perhaps there is an association of cCREs in these states with low variance genes that leads to an underestimate of the beta-coefficient. Nevertheless, the current esRP scores do show substantial utility, as explored in this manuscript. Also, our previous work (Xiang et al. 2020b) showed that the eRP scores (computed using a similar regression modeling) had explanatory power even for the consistently expressed genes, albeit less so than for differentially expressed genes (Fig. 6C in that paper). Thus, it is likely that inclusion of all genes, as we did in our current procedure, will provide the opportunity for the impact on expression of all or almost all the states to be estimated in the modeling, even with the potential for under-estimation for states primarily associated with constitutively expressed genes.

**Visualization of cCRE esRP scores across cell types using UMAPs**

The esRP scores were used to visualize the collection of VISION cCREs and follow how their regulatory impact changed across differentiation. Starting with the large matrix of about 200,000 human cCREs with esRP scores for each cell type and replicate, we employed the dimensional reduction visualization method UMAP (McInnes et al. 2018) to visualize the esRP matrix of all cCREs across all available cell types. This method for visualizing high dimensional data in lower dimensional space has been widely used in single cell data analysis where each cell’s transcriptomic or epigenomic profile is projected onto a 2-dimensional UMAP space. Similarly, we projected each cCRE’s esRP scores onto a 2-dimensional UMAP space. Thus, each point on our UMAP represents a cCRE rather than a cell, and cCREs with similar vectors of esRP scores across cell types are placed in similar locations on the UMAP plane. We used the umap library’s umap function in R with default settings to transform the data and generate the UMAP images.

The resulting image (Supplemental Fig. S12A) showed multiple clusters populated with cCREs (dots) that are colored to visualize their activity in individual cell types, with the color determined by the esRP score in that cell type. The darker red dots indicate cCREs more strongly implicated in gene activation in that cell type, as illustrated for erythroblasts (ERY) and monocytes (MON). UMAPs annotated by esRP scores for all cell types in both human and mouse along with movies showing the changes in estimated impact of cCREs across human hematopoietic differentiation are provided in the Supplemental Materials (Supplemental_Movie_S1.mp4) and on our VISION website (<http://usevision.org>). These annotated UMAP projections revealed both cCREs active in all cell types, such as the long arc of red cCREs in the upper right of the graphs, as well as shifts in cCRE activation as cells differentiate. The cCREs in the UMAPs were also shaded by their membership in esRP-based metaclusters distinctive for particular cell types or lineages (see next section on “Clustering of cCREs based on esRP scores”). This visualization revealed well-defined clusters of cCREs active in specific cell types (Supplemental Fig. S12B).

| 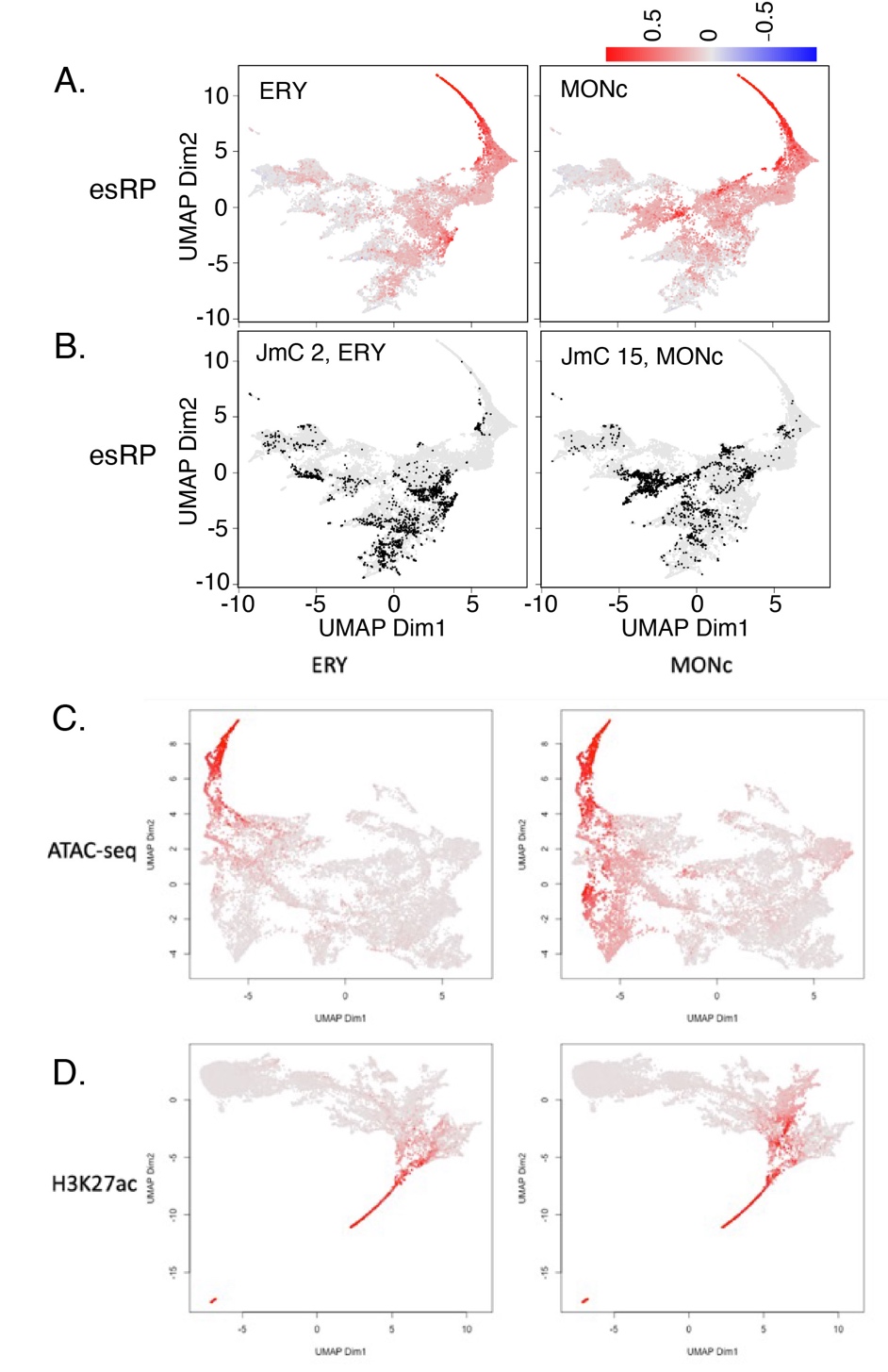 | **Supplemental Figure S12.** *Reduced dimensionality representation of cCREs based on esRP scores, chromatin accessibility, and H3K27ac signal.* **(A)** UMAP of cCREs based on their esRP scores across all cell-types. The points are colored by the esRP scores in the designated cell type. **(B)** UMAP of cCREs based on their esRP scores across all cell-types, with the points colored by the binary label indicating whether a cCRE belongs to the specified joint metacluster (JmC; see next section). **(C)** and **(D)** show UMAPs for the VISION cCREs based on their ATAC-seq signal or their H3K27ac signal, respectively, with the colors indicating the scores in ERY (left) or MONc (right). |
| --- | --- |

**Comparisons of esRP, chromatin accessibility, and H3K27ac in cCREs**

The esRP-based UMAPs used a score that reflected an integration of eight epigenetic features followed by an association with gene expression via multivariate linear regression, and our exploration of these integrative scores for visualization showed that they did reveal informative patterns. We also compare the results of using the integrative esRP scores versus simply using single epigenetic features for visualization. We generated UMAPs of the cCREs based on their signal intensity for ATAC-seq or H3K27ac across cell types. The UMAP based on ATAC-seq signal spread the cCREs over much of the projection plane, and it showed a long arc of cCREs active in all cell types examined along with groups of cCREs active preferentially in ERY or MONc (Supplemental Fig. S12C). The UMAP based on H3K27ac signal intensity revealed a shape on the projection plane that is similar to that for the esRP UMAP, but inverted, along with a prominent long arc of broadly active cCREs (Supplemental Fig. S12D). The comparisons of the UMAPs do not support any of the three metrics as being superior for visualization, but they do indicate that any of the three, including the integrative esRP scores, can reveal informative patterns when used as input for visualizations by dimensional reduction methods.

The three signals showed a positive association in pairwise comparisons (Supplemental Fig. S13A). The H3K27ac signal intensity was strongly and consistently associated with the beta-coefficients in each state, as expected from the well-established association of this histone modification with activation of CREs. Some IDEAS states had high ATAC-seq signal but low values for beta-coefficients, as expected for CREs serving predominantly structural roles, but the overall trend of the comparison was positive.

We also compared the three signals in a discriminatory task, specifically the ability to distinguish cCREs overlapping with transcription start sites (TSS) *versus* those that overlap with “transcription end sites” (TES, poly-A addition sites) or cCREs located elsewhere with respect to gene bodies. These annotations were chosen for the discriminatory task because they could be determined for all cCREs and are invariant across cell types. The UMAP projections based on each of the three signals in cCREs showed some clusters that appeared enriched for TSS and others that were somewhat enriched for TES (Supplemental Fig. S13B). The discriminatory ability of each signal was quantified using the Local Inverse Simpson’s Index, or LISI (Korsunsky et al. 2019). The distributions of LISI scores for distinguishing TSS or TES were high for all three signals, but the distributions were higher for esRP compared to the signals for the two individual features, indicating a somewhat better performance for esRP (Supplemental Fig. S13C).

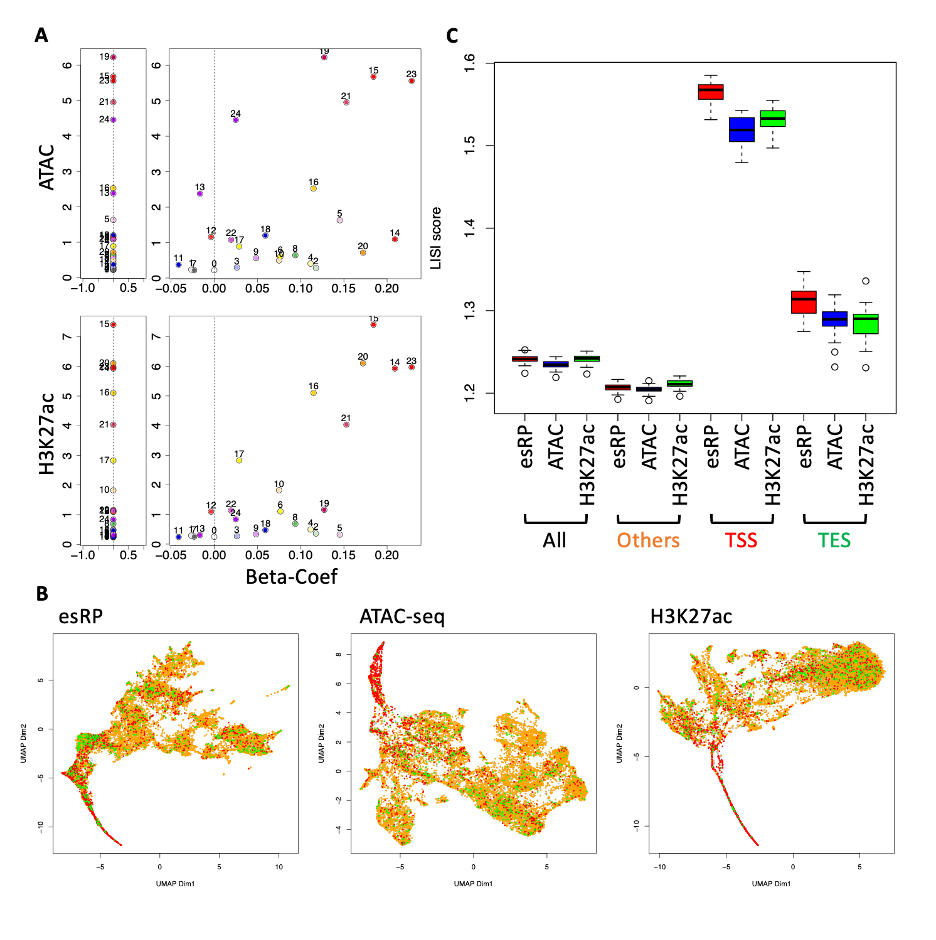

**Supplemental Figure S13.** *Relationships among esRP, ATAC-seq and H3K27ac signals and their ability to distinguish cCREs based on position within gene bodies.* (**A**) This panel depicts the relationship between ATAC-seq or H3K27ac signal and the beta-coefficient for each IDEAS state. The left part of each graph shows the mean ATAC-seq (top) or H3K27ac (bottom) signals in cCREs in each of the 25 IDEAS states, and the right part of each graph is a scatter plot of the means signals for the individual feature versus the beta-coefficient computed for each state in the MVLR. (**B**) UMAPs of cCREs, color-coded according to their intersection with gene structures, specifically red for overlap with transcription start sites (TSS, +/- 2kb), green for overlap with “transcription end sites” (TES, poly-A addition sites, +/- 2kb), and orange for all other cCREs. (**C**) This panel presents the LISI score for the UMAP, demonstrating its ability to differentiate specific gene structure cCREs from the remaining cCREs.

**Clustering of cCREs based on esRP scores**

*Rationale*

To infer the potential biological functions of the cCREs, we clustered them based on their esRP scores across different cell types. The traditional methods for clustering cCREs are based on a pairwise distance matrix of signals measured in different epigenomic datasets across different cell-types. However, these approaches cannot capture the information of the potential biological effects of the epigenetic modifications on gene expression regulation. For example, transitions from an initial state with only H3K4me3 to a second state with only H3K9me3 or to a second state with only H3K27ac have opposite associations with biological processes (repression versus activation, respectively), but the pairwise distances determined from the vectors of epigenetic signals would be very similar. In contrast to the signals measured directly by epigenomic datasets, the esRP score has already integrated all available epigenetic features’ information as a quantitative score that can reflect the overall potential of each cCRE for regulating gene expression. Thus, we hypothesized that clustering the cCREs using a distance based on esRP score would produce groups that better capture the changes of biological effects of the cCRE across different cell types.

*Method for generating esRP-based joint metaclusters of cCREs*

In this study, we conducted a series of clustering steps to generate robust clusters representing the prevalent patterns of inferred regulatory potential (esRP scores) across cell types jointly in both human and mouse species (Supplemental Fig. S14). We first created a combined matrix of esRP scores for all human and mouse cCREs across all shared cell types in both species. This ensured that the identified clusters were based on esRP patterns across the same set of cell types in both species. To mitigate the potential issues of cross-cell type collinearity and overfitting, we performed a principal component analysis and selected the top 10 principal components (PCs), which account for 99% of the variance. This generated a matrix containing 10 PC values corresponding to each cCRE. Although we had data for neutrophils from both species, we excluded the esRP scores for cCREs in neutrophils due to the significant noise in the ATAC-seq, which affected the overall quality of our results.

To identify robust clusters, we employed an iterative *k*-means clustering strategy, finding clusters with high consensus across repeated clustering rounds and then re-assigning all cCREs to these robust clusters. The initial clustering round (Step 1, Supplemental Fig. S14) used *k*-means clustering (K=100) based on the pairwise Euclidean distance of PCs of esRP scores across all cell types. This was performed 100 times, generating 10,000 clusters that could contain both human and mouse cCREs or cCREs from only one species. To identify consensus clusters across the 100 *k*-means runs, we combined the vectors of mean PC values for each of the 10,000 clusters from Step 1 into a matrix and clustered them again using *k*-means (K=100, Step 2, Supplemental Fig. S14). This step partitions the Step 1 clusters with similar characteristics into groups of clusters, where a large group size implies high consensus.

We considered groups with more than 70 Step 1 clusters as high consensus clusters, determined by calculating the Z score for the group size and finding the group size corresponding to the upper tail probability of 0.05. We identified 61 such groups, which we called robust clusters. However, these 61 clusters did not contain all cCREs, so we reassigned all cCREs in both human and mouse to one of the 61 robust clusters based on the cosine distance between each cCRE's esRP score profile and each cluster’s average esRP score profile, which was computed by averaging the mean-esRP-signal-vectors of the Step 1 clusters within each robust cluster.

Since our goal was to find clusters with cCREs in both species, we counted the number of cCREs from each species in each robust cluster and calculated the proportion of cCREs in each species for each robust cluster. We identified 44 clusters in which the proportion of cCREs did not significantly differ between the two species (i.e., the absolute log2 fold changes of cCRE proportion between the species were <= 2.18 (p-value = 0.05)), which we designated as robust clusters shared across species. Using the cosine distance, we reassigned each cCRE in each species to one of these 44 shared robust clusters.

Lastly, we observed that the average esRP score profiles across cell types for some subsets of the 44 clusters were similar, indicating they were not distinct clusters. To better differentiate groups of cCREs shared between species, we created joint metaclusters (JmCs) by clustering the clusters (Step 3, Supplemental Fig. S14). We combined the 44 clusters to generate 15 JmCs using hierarchical clustering (Hclust) and dynamic trimming with dynamicTreeCut (Murtagh and Legendre 2014). We ran Hclust followed by dynamicTreeCut 100 times, adding different noise (uniformly distributed within a range of -0.001 to 0.001) for each run and using the cosine distance matrix. We then created a count matrix to record the frequency of each cluster being found in the same JmC. We used Hclust to cluster the count matrix and applied dynamicTreeCut to cut the Hclust tree into 15 JmCs. As a result, each cCRE in mouse and human was assigned to one of the 15 JmCs. These JmCs provide discrete categories for cCREs based on the cell type distribution of their estimated regulatory impact.

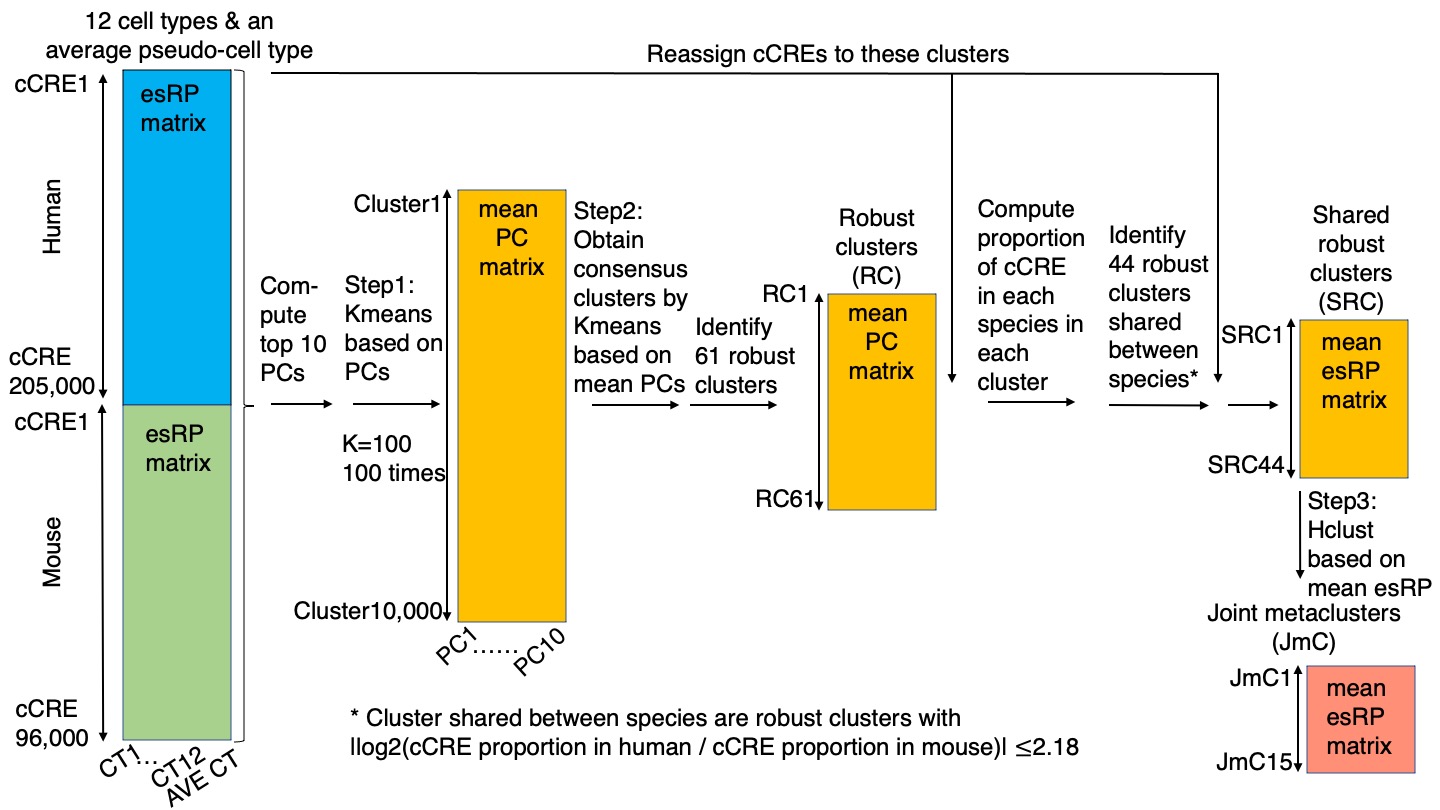

**Supplemental Figure S14.** *Overview of clustering of cCREs based on their esRP scores*, leading to the partitioning of all human and mouse cCREs into one of 15 joint metaclusters (JmCs)*.* The steps in the clustering are described in the Methods. Abbreviations are CT = cell type, esRP = epigenetic state Regulatory Potential, PC = principal component, RC = robust cluster, SRC = shared robust cluster, JmC = joint metacluster.

**Enrichment of specific JmCs for promoter or enhancer annotations**
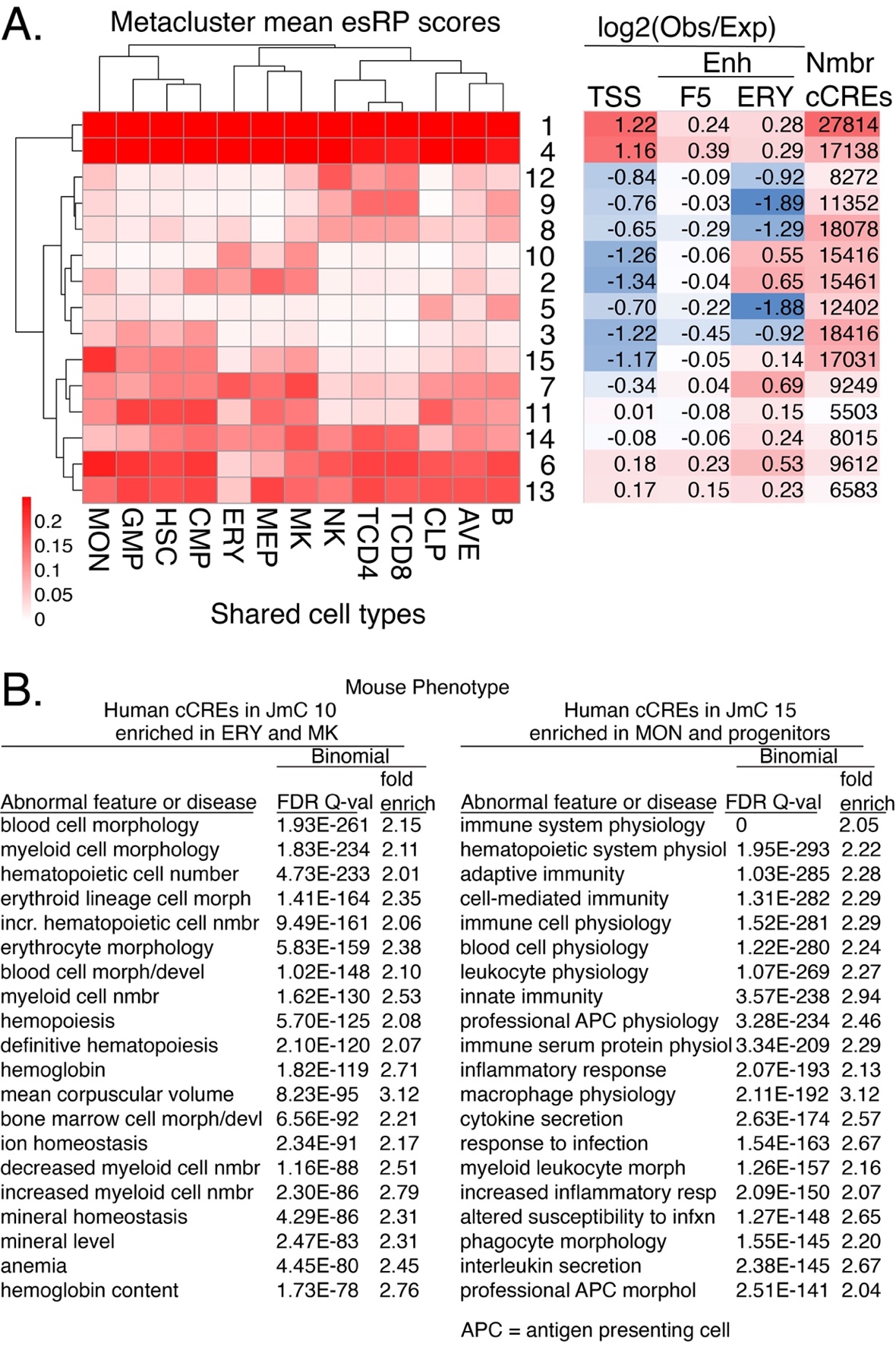

**Supplemental Figure S15.** *Enrichment of orthogonal functional annotations and mouse phenotype terms in the joint metaclusters of cCREs based on their esRP scores across cell types.* **(A)** Enrichment and depletion of cCREs in esRP joint metaclusters (JmCs) for function-associated elements. Enrichment or depletion is the log2(Obs/Exp) for overlaps of the cCREs in each JmC with the designated set of function-associated elements. The expected value was calculated from the fraction of all human VISION cCREs (200,342) that overlap with the sets of elements. TSSs were from GENCODE (extended +/- 500bp for a 1000bp interval). Enhancer sets were from FANTOM5 enhancers (F5) determined from enhancer RNAs in many cell types (Andersson et al. 2014) or the Xu laboratory erythroid (ERY) enhancer repertoire (Huang et al. 2016), determined from histone modification patterns in adult and fetal erythroblasts. **(B)** Enrichment of functional terms from phenotype ontologies for human cCREs in JmC 10 (enriched in erythroid and megakaryocytic cells) and JmC 15 (enriched in monocytes and progenitor cells). Queries on the VISION cCRE_db returned 15,416 human cCREs in JmC 10 and 17,031 human cCREs in JmC 15. These sets of genomic intervals were analyzed for functional term enrichment of linked genes using the GREAT tool, v.4.0.4 (McLean et al. 2010), using the "Basal plus extension" option to associate genomic regions with genes, choosing the default proximal = 5 kb upstream and 1 kb downstream but limiting the extension to 100 kb rather than the default of 1000 kb. The table lists the top 20 Mouse Phenotype terms ordered by the FDR q-value; the fold enrichment is also given. Enriched terms from the Human Phenotype ontology also were associated with the blood cell types in which the JmCs were enriched, but those terms were not as precise as the ones from the Mouse Phenotype ontology.

**Enrichment of JmCs assigned to cCREs in gene loci**

*Related to main Figure 4D*.

We calculated the enrichment of JmC assignments of cCREs at gene loci (TSS +/-50kb) in both species. For each ${Gene}_{i}$, the enrichment of ${JmC}_{j}$ is defined as follows:

$${Enrichment}_{i, j}= \frac{\left( {Gene}_{i}\_{JmC}_{j}\_{cCRE}_{Human}+1 \right)\times\left( {Gene}_{i}\_{JmC}_{j}\_{cCRE}_{Mouse}+1 \right)}{\left( expected\_{Gene}_{i}\_{JmC}_{j}\_{cCRE}_{Human}+1 \right)\times\left( {expected\_Gene}_{i}\_{JmC}_{j}\_{cCRE}_{Mouse}+1 \right)}$$

where the ${Gene}_{i}\_{JmC}_{j}\_{cCRE}_{Human/mouse}$ is the number cCRE assigned to ${JmC}_{j}$ at the ${Gene}_{i}$ locus in human or mouse, the $expected\_{Gene}_{i}\_{JmC}_{j}\_{cCRE}_{Human/Mouse}$is the expected number of cCRE assigned to ${JmC}_{j}$ at the ${Gene}_{i}$ locus in human or mouse by random chance.

**Enrichment of trait-associated SNPs in the joint metaclusters (JmCs)**

We sought to determine the enrichment of trait-associated SNPs in the subsets of VISION cCREs based on activity within groups of cell types, i.e., the joint metaclusters (JmCs). We re-analyzed the blood trait-associated variants by running sLDSC with fifteen separate annotations, each annotation defined by a JmC, to see if the trait heritability was enriched in any JmCs. Running these annotations through the same pipeline described above (see section “Enrichment of genetic variants for blood cell related traits in the VISION human cCRE collection”), we obtained estimates of the enrichment of each JmC in the heritability of each of 587 traits.

We found five JmCs with significant results at a 5% FDR (Supplemental Fig. S16). The cCREs more active in erythroid and megakaryocytic cells, i.e., those in JmCs 10 and 2, were significantly enriched for heritability of several blood traits, including many related to erythroid cells. Several of the enrichments were for cCREs in JmCs 1 and 4, which are active across all cell types examined and are themselves highly enriched for proximal regulatory regions such as promoters (Supplemental Fig. S15A). While this result may suggest that many blood-trait associated variants were in proximal regulatory regions of genes with active epigenetic marks broadly present across blood cells, more study is needed to establish such a relationship. One caveat is that the large number of cCREs in JmCs 1 and 4 makes it more likely for them to overlap with any feature, and thus the large overlap with proximal regulatory regions could be separable, at least in part, from the overlap with trait-associated variants. Many of the JmCs showed no significant enrichment, perhaps reflecting a reduced power for JmCs comprising fewer cCREs.

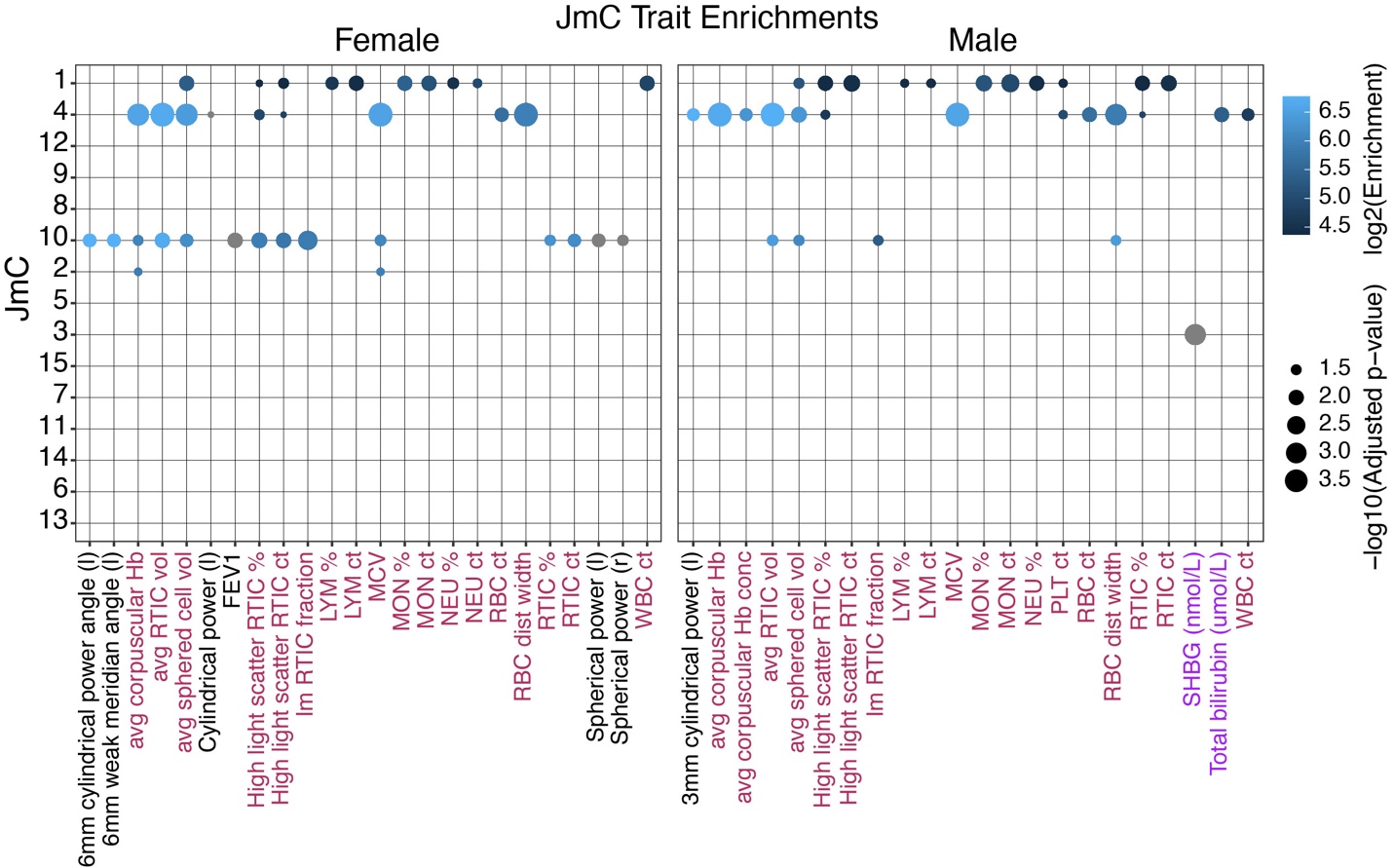

**Supplemental Figure S16. Enrichment of SNPs associated with blood cell traits from UK Biobank in VISION cCREs.** Results of the JmC sLDSC analysis where each set of cCREs within a JmC was considered as a separate annotation. The plot lists a trait on the x-axis if any JmC had a significant enrichment for it. The labels for these traits are maroon for blood count traits, purple for blood biochemistry traits, and black for non-blood related traits. The plot lists the JmC on the y-axis. For a given JmC and trait combination, a dot is plotted if and only if there was an observed significant enrichment for that combination. Size of the dot reflects the significance of the enrichment, and the color of the dot reflects the size of the enrichment itself. Negative enrichments are colored gray. Panels separate the sex in which the GWAS analysis was performed for each trait.

**Comparisons of cCREs clusters based on esRP, chromatin accessibility, and H3K27ac for distinguishing among cell types**

Based on prior work, it was expected that some cCRE clusters would be distinctive for individual cell types. The JmCs based on esRP scores tended to contain cCREs active in groups of cell types in a lineage (e.g., lymphoid cells) rather than in individual cell types. We investigated whether this apparent loss of specificity was a result of the metric used for clustering or whether it resulted from combining previously distinctive clusters.

First, we generated joint metaclusters of the cCREs using the procedure illustrated in Supplemental Fig. S14 but using the signals in each cCRE for ATAC-seq or for H3K27ac in each cell type as the input for clustering. The patterns of the single or integrative (esRP) signals across cell types had broad similarities, with some JmCs composed of cCREs with high signal across all cell types and other JmCs more distinctive to related groups of cell types (Supplemental Fig. S17). None of the three signal sources showed a superior ability to distinguish among individual cell types.

| 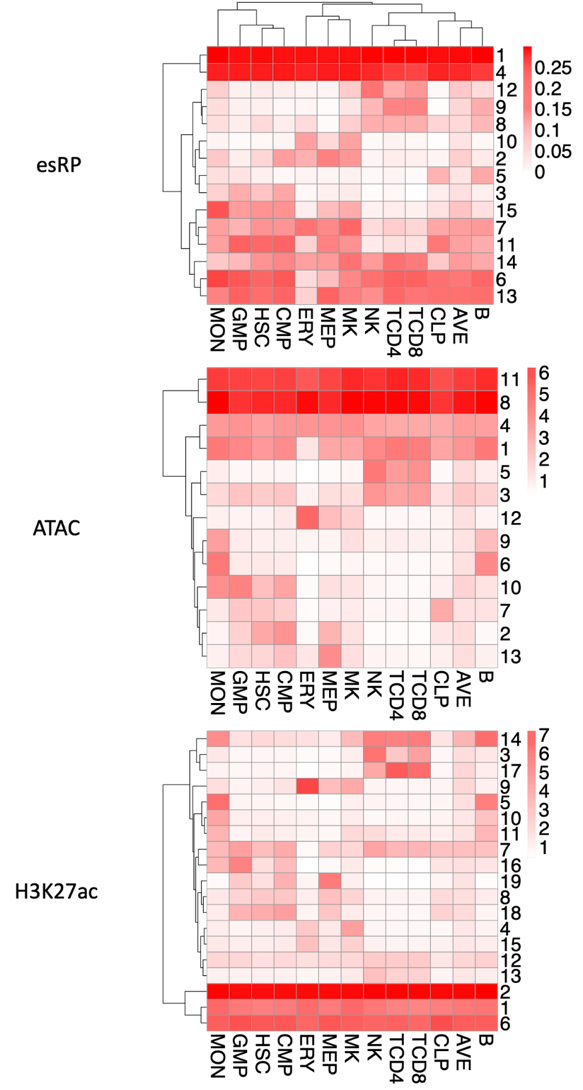 | **Supplemental Figure S17.** *Joint metaclusters of cCREs based on three different input signals.* The metaclusters were combinations of shared robust clusters (see Supplemental Fig. S13) generated by a series of *k*-means clusterings using as input the esRP scores (top), ATAC-seq signal (middle) or H3K27ac signal (bottom) of cCREs in each cell type. The mean signal for the cCREs in each metacluster in each cell type was indicated by the intensity of red color in the heat map. |
| --- | --- |

We then examined the larger number of shared robust clusters (SRCs) generated though this clustering pipeline for each input signal. For all three input signals, the SRCs had clusters of cCREs that were distinctive for individual cell types (Supplemental Fig. S18). For example, esRP-based SRCs 9:37 and 12:54 (JmCid:SRCid) distinguished cCREs active in T cells from those in NK cells, and SRCs 5:5 and 5:52 distinguished cCREs active in B cells from those active in CLPs. The ATAC-seq SRCs 6:80 and 9:36 were distinctive for cCREs active in MON, while SRC 6:93 was distinctive for B cells. Similarly, the H3K27ac SRCs 5:74 and 10:45 were distinctive for cCREs active in MON, while SRCs 5:78 and 10:90 were distinctive for B cells. We conclude that clusters of cCREs based on any of the three signals could be distinctive for individual cell types when a sufficiently large number of clusters were considered.

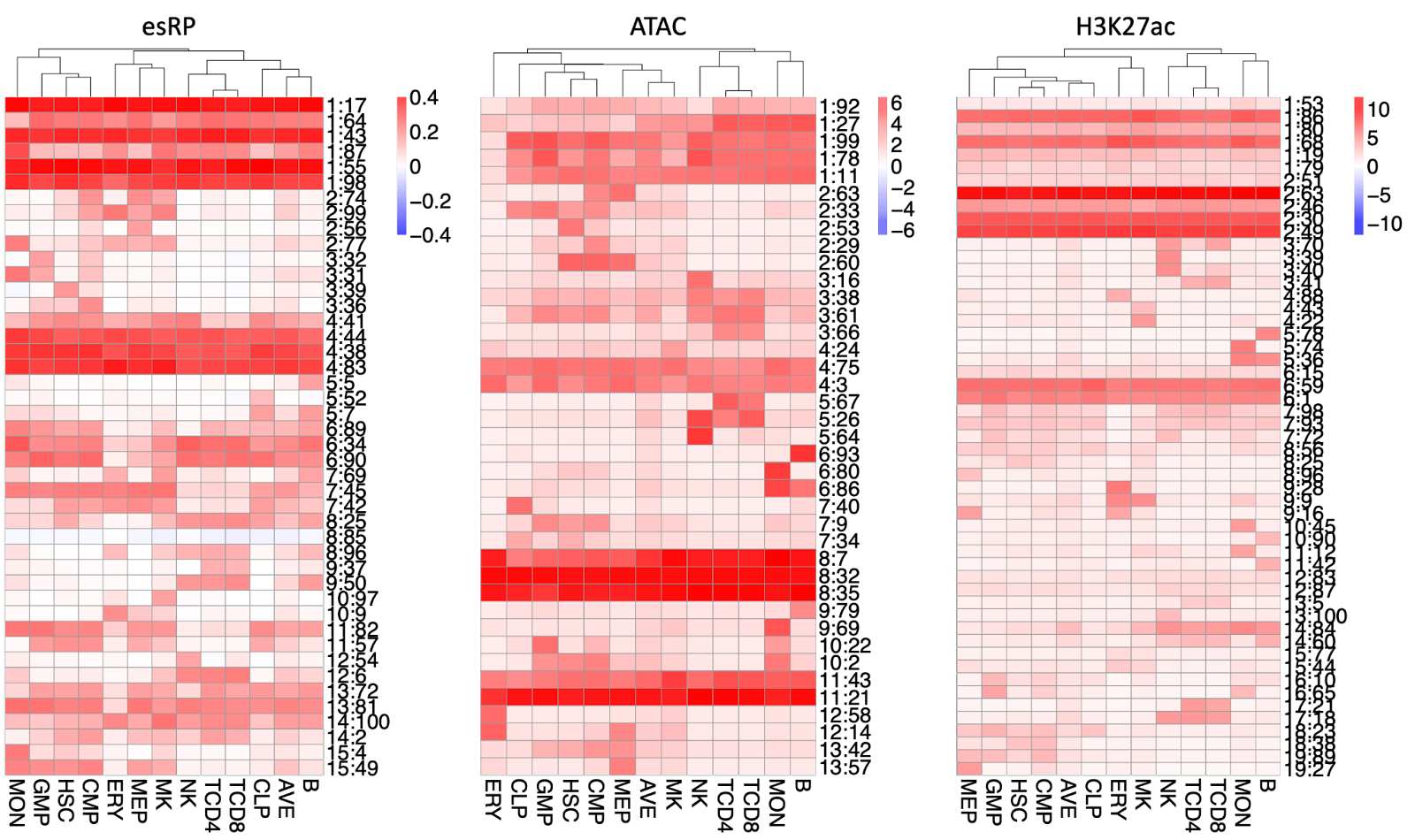

**Supplemental Figure S18.** *Shared robust clusters of cCREs based on three different input signals.* The shared robust clusters (SRCs) were generated by a series of *k*-means clusterings using as input the esRP scores (left), ATAC-seq signal (middle) or H3K27ac signal (right) of cCREs in each cell type (see Supplemental Fig. S13). The SRCs were clusters of cCREs found frequently in multiple rounds of *k*-means clustering that also contained cCREs from both mouse and human. The mean signal for the cCREs in each SRC in each cell type was indicated by the intensity of red color in the heat map. Each row was labeled by the joint metacluster assignment followed by the SRC assignment, i.e., JmCid:SRCid.

**Self-organizing maps**

A complementary approach to systematic integration of epigenetic features or RNA data across cell types is to build self-organizing maps (SOMs) such that each map unit contains elements (cCREs or genes) with similar profiles (epigenetic or expression) across cell types, and units with similar collections of profiles are close to each other in the SOM. We built DNA and RNA SOMs for the human VISION data (Jansen et al. 2019). To build DNA SOMs, the ChIP-seq and ATAC-seq data over all the cCREs were trained on a 60x90 SOM using the SOMatic package (https://github.com/csjansen/SOMatic). This approach yielded 160 different metaclusters, each of which represents a collection of cCREs with a similar signal profile across each cell type and experiment (Supplemental Fig. S19A). An RNA SOM was built by training a 40x60 SOM on the expression profiles of genes across cell types (Supplemental Fig. S19B), which generated 21 metaclusters, each of which contained genes with similar expression profiles across cell types. These SOMs can be viewed interactively at the server at <http://usevision.org>.

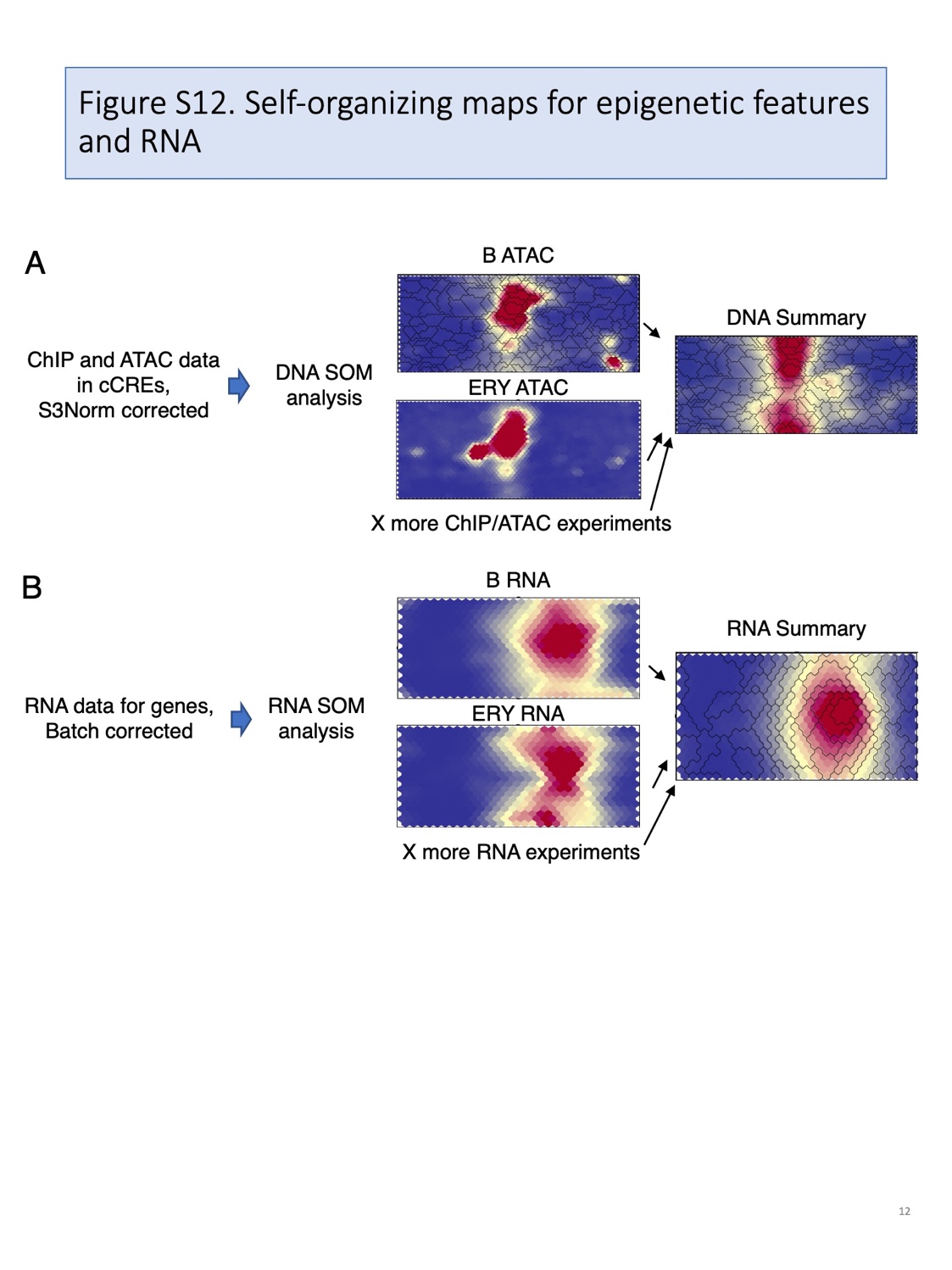

**Supplemental Figure S19.** *Self-organizing maps of DNA-based data and RNA-based data.*

**Enrichment for transcription factor binding site motifs in joint metaclusters of cCREs**

We used the Maelstrom tool in the GimmeMotifs suite (v0.17.1) to identify motifs that are differentially enriched across JmCs (Bruse and van Heeringen 2018). In contrast to previous analyses in which all cCREs active in a specific cell type or those that are distinctive for a cell type were examined (e.g., Neph et al. 2012; Vierstra et al. 2020), our search for motif enrichment in the JmCs examined not only sets of cCREs that are distinctive for a lineage, but also sets of cCREs with a broader distribution of activity. We first labeled all cCREs according to their JmC membership. We then ran separate Maelstrom analyses on human and mouse cCRE sets to find enrichment of motifs in GimmeMotif’s default “gimme.vertebrate.v5.0.pfm” collection of non-redundant clustered vertebrate motifs derived from the Cis-BP database (Weirauch et al. 2014). Maelstrom’s --filter-cutoff parameter was set to the default value of 0.8, which has the effect of filtering redundant motif enrichment results based on scores across the input sets. We filtered out motifs that did not achieve a Maelstrom *Z*-score of at least 4 in any JmC. We then combined results across human and mouse (which required running Maelstrom again for each species using the --no-filter option to fill in *Z*-scores for motifs that were found in one species but not the other). Heatmaps were constructed using the Python seaborn package (Waksom 2021) and motif logos were plotted using WebLogo3 (Crooks et al. 2004). Putative TF names were associated with each motif by examining the identities of Cis-BP motifs that were clustered into the relevant non-redundant motifs within the GimmeMotifs non-redundant set, and by matching enriched motifs against mouse and human motifs from Cis-BP (v.2.0) using STAMP (v.1.0) (Mahony and Benos 2007) with arguments “-cc PCC -align SWU”.

**Binding of CTCF to cCREs in JmCs: species similarities and differences**

The enrichment of TFBS motifs for CTCF and ZBTB7A presented some potential exceptions to the sharing of motifs across species. The cCREs in JmC 8 showed the expected strong enrichment for these motifs in both human and mouse, with little enrichment for binding site motifs of other TFs (main Fig. 4E). These cCREs had modest regulatory impact, as estimated by esRP scores, across most cell types, suggesting the hypothesis that cCREs in JmC 8 may consist of CTCF-bound sites that are not involved in gene activation. Indeed, examination of ChIP-seq results showed that the cCREs in JmC 8 were enriched for CTCF occupancy in both mouse and human and for overlap with loop anchors (Supplemental Fig. S20A and B). In contrast, the cCREs in several JmCs were enriched for CTCF and ZBTB7A motifs only in mouse (e.g., JmCs 12, 7, and 10) or only in human (e.g., JmCs 11 and 13; see main Fig. 4E). In these JmCs, the cCREs were also enriched for binding sites for other TFs, with those other motifs enriched in the cCREs from both species. The frequency of occupancy by CTCF in the cCREs in these latter JmCs corresponded well with the enrichments for the motifs, with JmCs with greater enrichment for CTCF motifs in human also more enriched for CTCF occupancy in human, and *vice versa* for mouse (Supplemental Fig. S20A). These observations lent support to the suggested species-specificity. A parallel analysis of the cCREs in human and mouse JmCs by the multi-label discriminative motif-finder SeqUnwinder (Kakumanu et al. 2017) also uncovered enrichment of some motifs that apparently systematically vary in enrichment between mouse and human cCREs, but the magnitude of enrichment was limited (Supplemental Fig. S21).

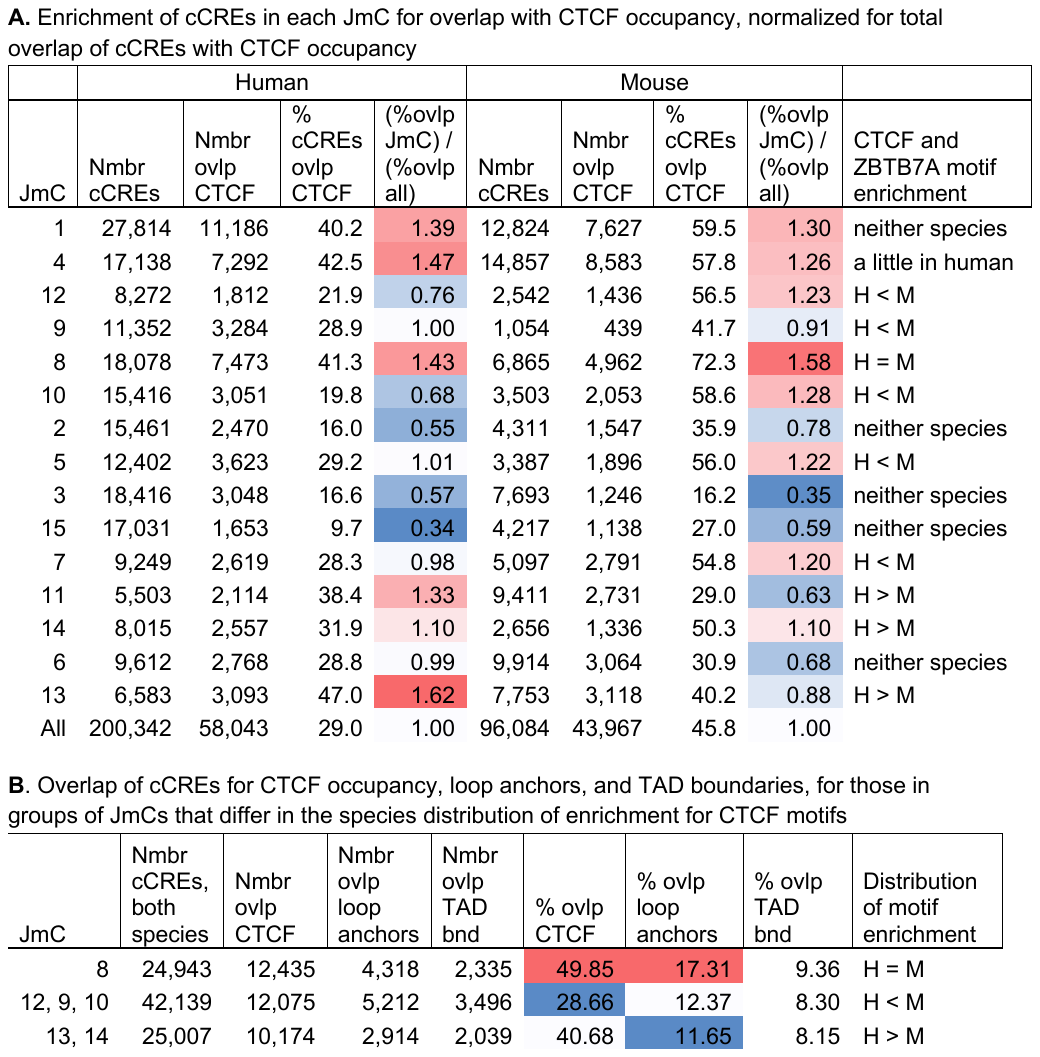

**Supplemental Figure S20.** *Overlap of cCREs in joint metaclusters with DNA segments bound by CTCF and with loop anchors.* **(A)** Enrichment for overlap of cCREs in JmCs for occupancy by CTCF. Extensive data on CTCF occupancy in many blood cell types were recorded in our cCRE db, and we used its query capacity to find the numbers of cCREs in each JmC (human and mouse separately) that were bound by CTCF, according to the ChIP-seq results compiled in the VISION project. These numbers were listed in the indicated column, along with percentage of cCREs that overlap and an estimated enrichment for overlap (colored cells, red for higher enrichment), determined by dividing by the percentage of all cCREs that are bound by CTCF. The final column summarizes the results on motif enrichment from main Fig. 4E. **(B)** Enrichment for overlap of cCREs in JmCs for loop anchors and TAD boundaries. This analysis was conducted on the groups of JmCs that differ in the species distribution of enrichment for CTCF motifs. Abbreviations in both panels: JmC = joint metacluster, Nmbr = number, ovlp = overlap, bnd = boundaries, H = human, M = mouse.

*Method for SeqUnwinder*

We used the SeqUnwinder multi-label discriminative motif finder to discover de novo motifs associated with each JmC and to search for motifs that were potentially constitutively differentially enriched across species. We first gave every cCRE two labels: their JmC membership and their species of origin. We then filtered out sequences that were larger than 1kbp and randomly selected 50,000 sequences from the remaining set. We then provided these doubly-labeled sequences to SeqUnwinder (v.0.1.5) (Kakumanu et al. 2017) and ran analysis using the following options: --threads 8 --win 200 --mink 4 --maxk 5 --r 10 --x 3 --a 200 --hillsthresh 0.1 --memesearchwin 16 --minsubclass 150. Heatmaps were again constructed using the Python seaborn package (Waksom 2021) and motif logos were plotted using WebLogo3 (Crooks et al. 2004). Putative TF names were associated with each SeqUnwinder-discovered motif by matching against mouse and human motifs from Cis-BP (v.2.0) using STAMP (v.1.0) (Mahony and Benos 2007) with arguments “-cc PCC -align SWU”.

| 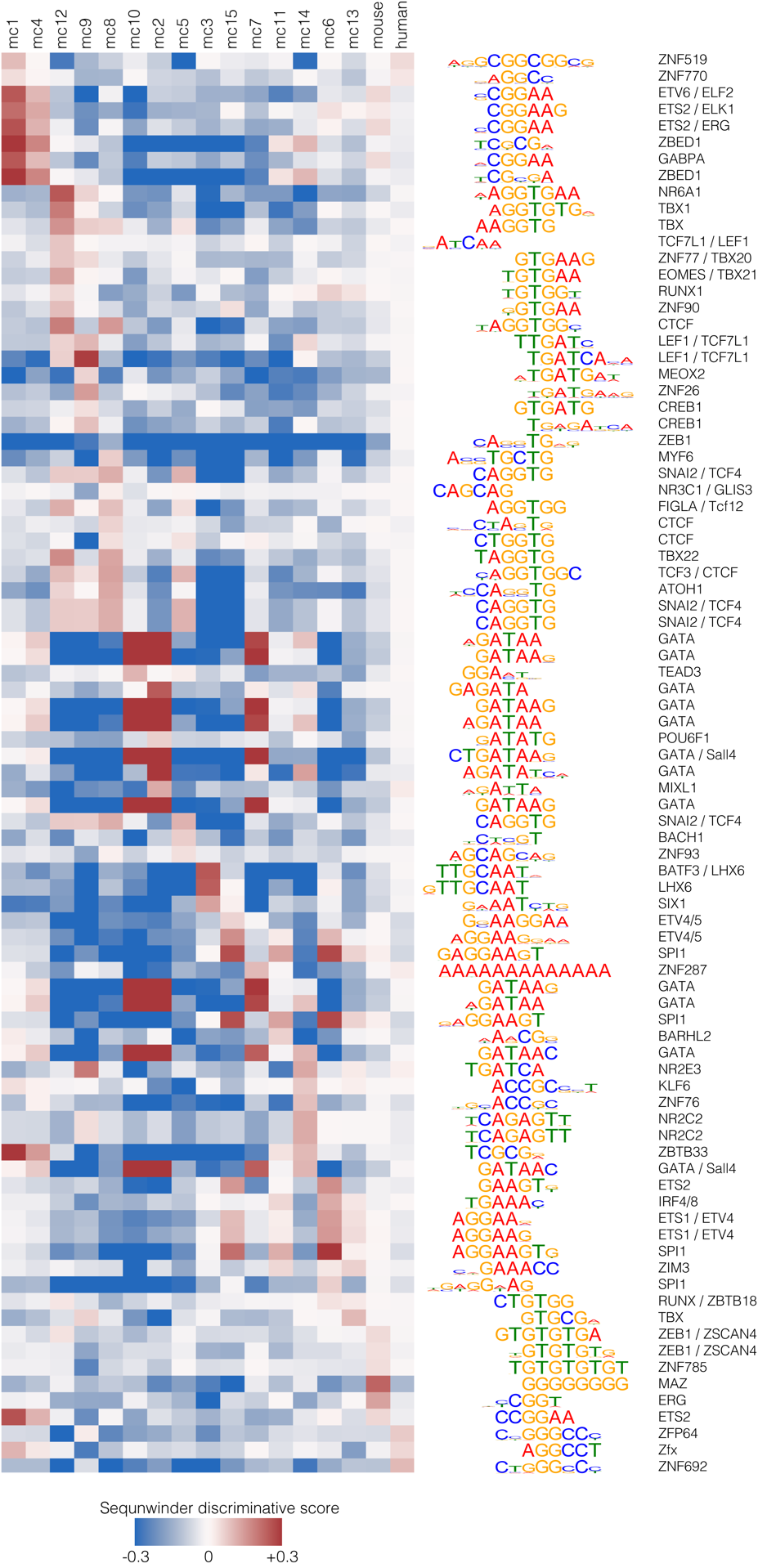 | **Supplemental Figure S21.** *Discriminatory motifs discovered by SeqUnwinder.* This multi-label discriminative motif-finder can deconvolute contributions of the esRP pattern from the contributions of the species in the motif enrichment patterns. The first 15 columns show motif enrichments in each of the joint metaclusters of cCREs, and the last two columns show motifs that were enriched specifically in each of the two species. As shown toward the bottom of the matrix, these species-specific motifs were primarily a GC-rich motif that was systematically enriched in human cCREs and a TGTG-repeat motif enriched in mouse cCREs. The magnitude of enrichment of these motifs was limited. These apparently species-specific motifs can serve as the basis for more detailed studies in the future. |
| --- | --- |

**Small differences in the number of cCREs in the structure and function conserved category (SF cCREs)**

The sets of cCREs conserved both in interspecies sequence alignments and inferred function as regulatory elements (SF cCREs, main Fig. 5A) were expected to have the same numbers of cCREs in both human and mouse. While the numbers were quite similar, they were not identical. We discovered “breakage” of a cCRE in one species into multiple cCREs in the other species was a contributor to this apparent discrepancy (Supplemental Fig. S22). The most common splits were one cCRE being divided into two cCREs in the other species. Such a split could reflect a gap in the interspecies alignment or an evolutionary separation into two elements in the other species.

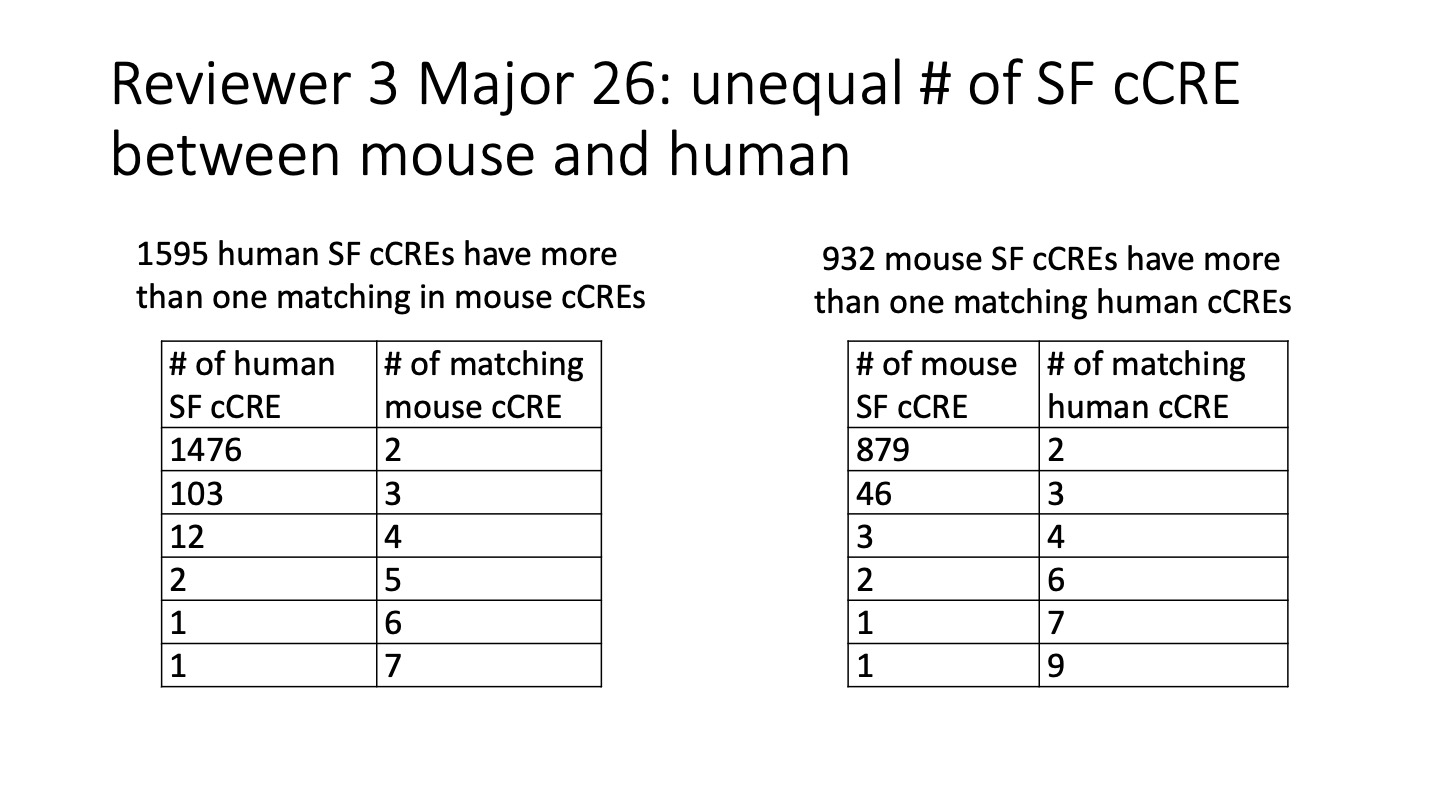

**Supplemental Figure S22.** *Frequencies at which a single cCRE in one species was split into multiple cCREs in the other species during the liftOver procedure.*

**Comparison of magnitude of chromatin accessibility in orthologous cCREs in human and mouse blood cells**

The sets of SF cCREs presented the opportunity to examine whether orthologous, presumptive regulatory elements were involved in similar regulatory processes in both species. We reasoned that the level of chromatin accessibility would reflect the role and frequency of utilization of an element, and we used the accessibility signal as a proxy for regulatory role. We found that SF cCREs in analogous cell types in human and mouse showed a substantial positive correlation in ATAC-seq signal strength (Supplemental Fig. S23), with Pearson’s correlation coefficients ranging from 0.52 to 0.67. This result indicated that the SF cCREs may be playing similar regulatory roles in both species.

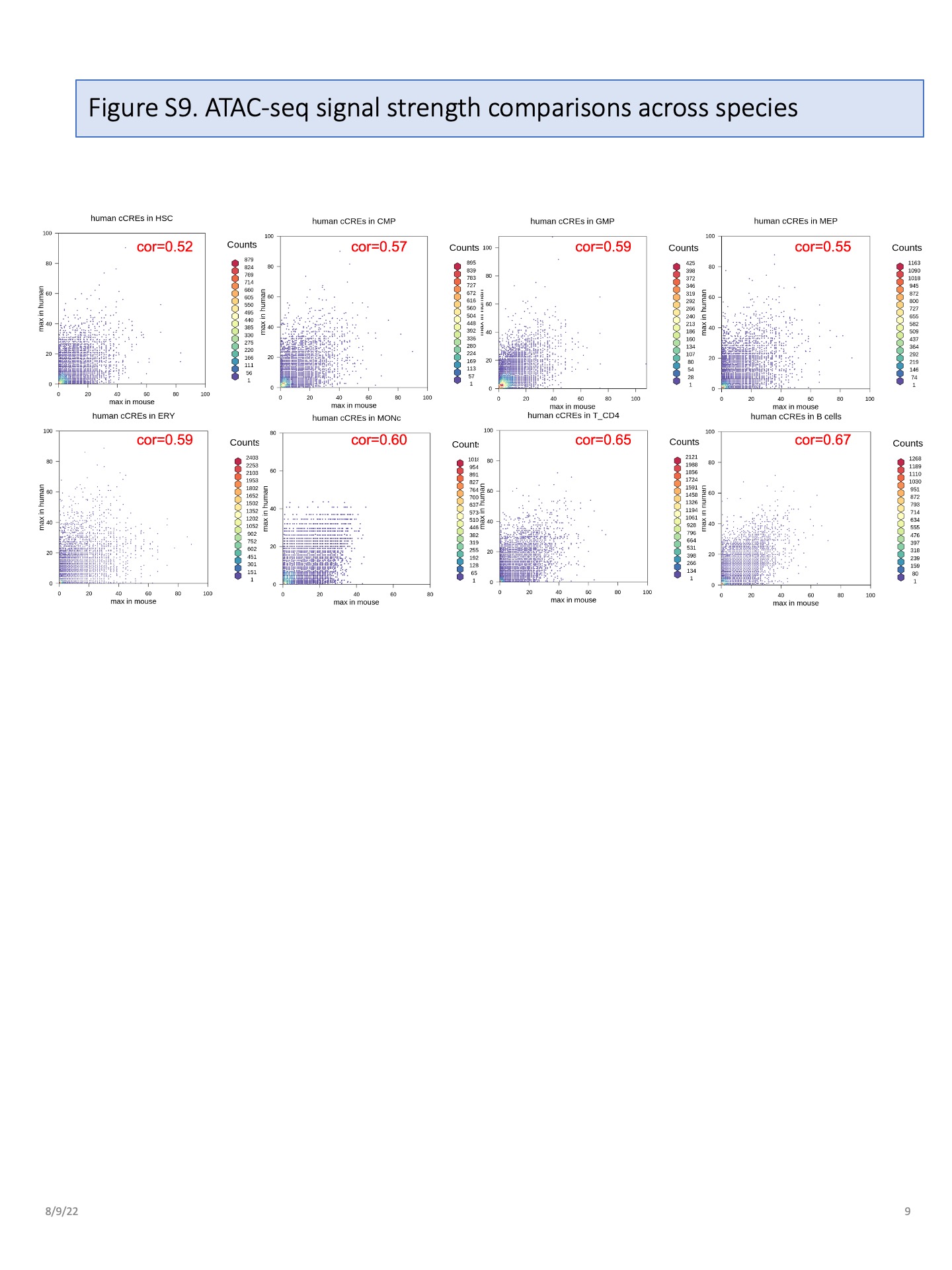

**Supplemental Figure S23.** *ATAC-seq signal strength comparisons across species.* The signal strengths for chromatin accessibility for SF cCREs (conserved both in sequence and inferred function) were represented by the maximum normalized ATAC-seq signal across replicates in each cell type. Two-dimensional density plots were generated for the signal strength for chromatin accessibility of SF cCREs in analogous cell types between mouse and human, and the correlation coefficients were used to estimate the similarities of the signal strength in those cCREs. Quantile normalization was applied to make the distribution of ATAC-seq signals comparable between the two species. In general, the signal strength of chromatin accessibility of SF cCREs were positively correlated in analogous cell types between mouse and human.

**Profiles of epigenetic states in the three evolutionary categories of cCREs in human and mouse blood cells**

To investigate whether any epigenetic states occurred more frequently in one of the three evolutionary categories of cCREs, we determined the fraction of cCREs in each epigenetic state in the various cell types for cCREs in each evolutionary category in human and in mouse. We focused on eight cell types, specifically hematopoietic stem and progenitor cells (human HSC and mouse LSK), CMP, GMP, MEP, erythroblasts (ERY), B cells, CD4+ T cells and CD8+ T cells, that were considered analogous between human and mouse. All had replicated epigenetic datasets on human (for a total of 16 biosamples), and two had replicated datasets in mouse (10 biosamples). The cCREs in each evolutionary category were assigned to a single dominant epigenetic state in each cell type (and replicate), and the distribution of epigenetic states on cCREs was computed. Because the full set of human cCREs was about twice the size of the set of mouse cCREs, a selection of 96,084 human cCREs based on the strongest mean ATAC-seq signal across all cell types was used for comparison (Supplemental Fig. S24A, B). Epigenetic states associated with promoter function were prevalent in the SF-conserved cCREs in both human and mouse, notably states 14 (PA), 15 (PNA), and 19 (PN) (Supplemental Fig. S24C). In contrast, greater proportions of epigenetic states associated with enhancers were assigned to S cCREs and N cCREs in human and mouse, such as states 4 (E1) and 5 (N) (Supplemental Fig. S24D). States associated with transcription were also common in the S and N categories of cCREs. State 13 (CN), associated with CTCF binding and chromatin accessibility, was common in all three categories, but notably higher in SF cCREs (Supplemental Fig. S24E). The distributions of epigenetic states were quite similar between the S-conserved and nonconserved cCREs across cell types and between species. These results indicated that the stringency of conservation of cCREs was related to their inferred function, with the cCREs conserved in sequence and inferred function associated with promoter-like states. This association led to the hypothesis that many SF cCREs were promoters, and this hypothesis was supported by the analyses presented in the main text, Fig. 5B.

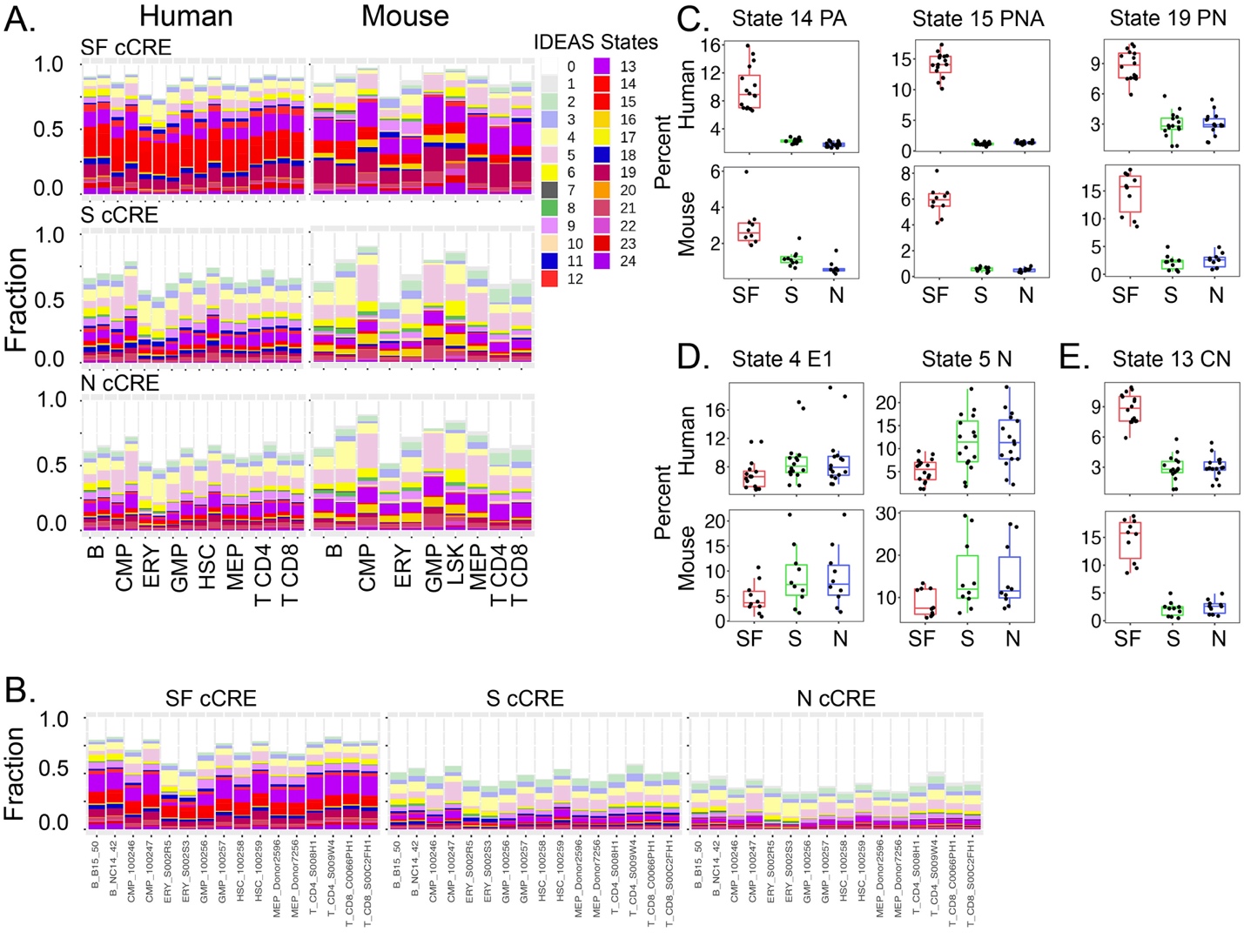

**Supplemental Figure S24.** *Epigenetic state profile comparisons of cCREs across evolutionary categories and between human and mouse.* **(A)** Proportions of 25 epigenetic states in eight analogous cell types between mouse and human for three evolutionary categories. The top 96,084 Human cCREs were selected to match the total number of mouse cCREs based on maximum averaged ATAC-seq signal strength across cell types. The proportions for the full list of human cCREs were shown in **(B)**. Examples of the percentages for promoter-related **(C)**, enhancer-related **(D)**, and CTCF-binding-related **(E)** epigenetic states in three evolutionary categories among eight analogous cell types between mouse and human.

**Impact of cCRE location relative to the transcription start site (TSS) of genes on conservation scores**

The differences in distributions of phyloP scores were interpreted as reflections of differing levels of evolutionary constraint on cCREs in the three evolutionary categories. However, the positions of cCREs relative to strongly conserved elements such as coding exons can influence the phyloP score because the DNA sequence alignments used in computing the phyloP scores are impacted by positions of aligning DNA segments relative to highly conserved sequences (King et al. 2007). Specifically, the local alignments were built by extensions from initial high-scoring matches, and those initial matching segments were frequently in coding exons (Schwartz et al. 2003). Previous work has shown that the fraction of predicted cCREs with sequences that align between mammalian genomes was impacted by masking the coding exons when constructing the whole genome alignment (King et al. 2007). To investigate the potential impact of this effect, we computed the distribution of distances from the transcription start site (TSS) of genes both for cCREs and for randomly chosen matched genomic intervals (Supplemental Fig. S25). The distributions of distances showed considerable overlap, but the distribution for cCREs had a higher peak close to the TSS compared to the random intervals. The distributions were significantly different, and thus we cannot exclude an effect of position relative to the TSS on the distributions of phyloP scores. However, this effect does not rule out an impact of evolutionary constraint on the phyloP scores, which are themselves derived from a statistical model that includes phylogenetic inferences.

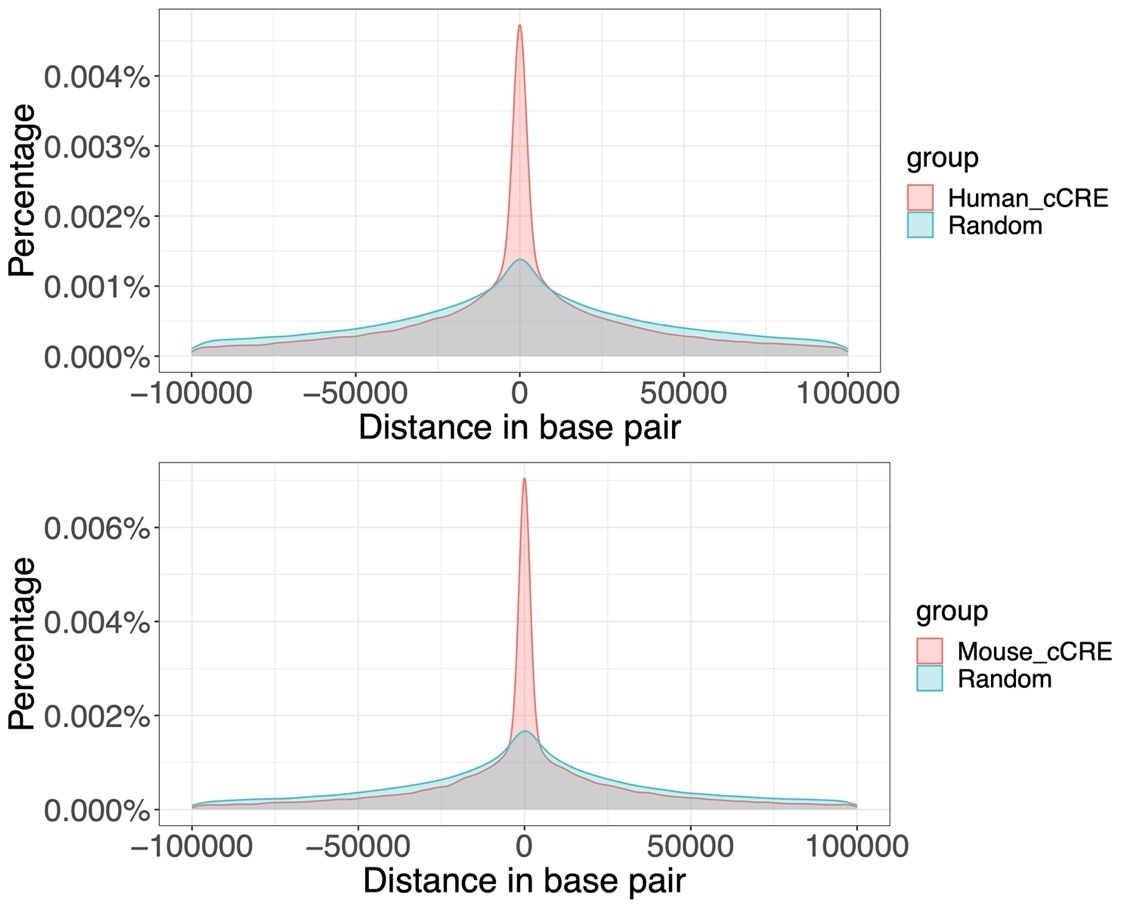

**Supplemental Figure S25.** *Distributions of distances from the TSS of the nearest protein-coding gene for cCREs and for randomly selected matched genomic intervals.* Distributions of distances within a 100 kb interval on each side of the TSS are shown for cCREs and random intervals for (**A)** human and **(B)** mouse. The distributions were significantly different between cCREs and random intervals (p-value < 2.2^-16 by a two-sample Kolmogorov-Smirnov test).

Given that the distance from the TSS could have some impact on the phyloP scores, we computed the distribution of these scores after partitioning the cCREs in each evolutionary category into those that are proximal to the TSS (+/- 1kb) or distal (all other cCREs). For all three evolutionary categories, the distribution of phyloP scores was higher for the proximal than for the distal elements. However, even focusing on the distal elements, the distribution of phyloP scores is still highest for SF cCREs, followed by S cCREs, and lowest for the N cCREs (Supplemental Fig. S26).

| 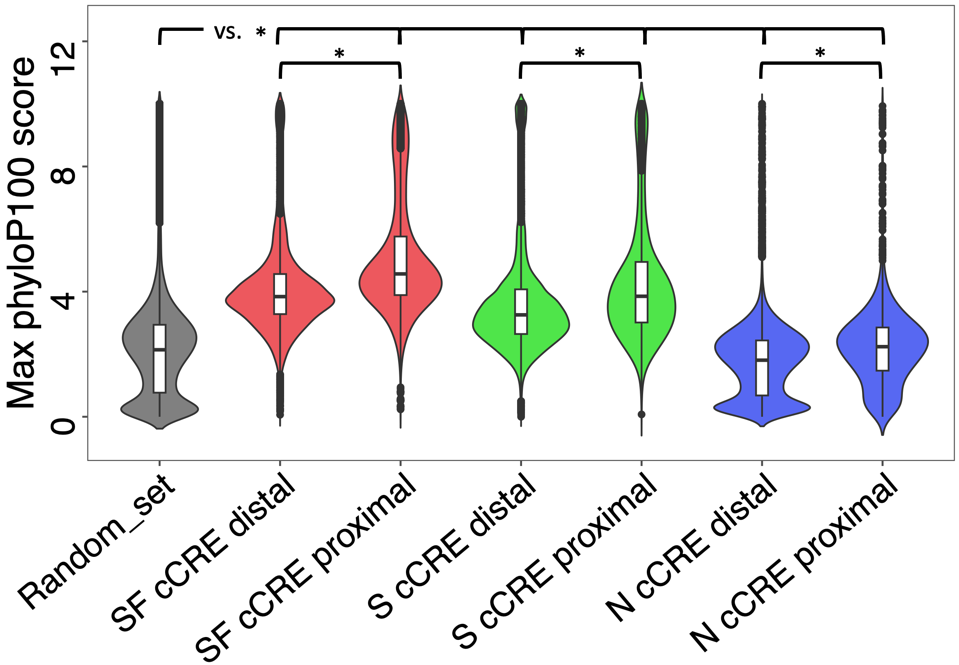 | **Supplemental Figure S26.** *Distributions of maximum phyloP scores for human cCREs in the three evolutionary categories partitioned into proximal or distal groups.* The asterisks (*) mark comparisons that were significant (p-value < 2.2^-16 for all except N proximal vs random, which has p=0.0017) by a Welch two sample *t*-test. The phyloP scores for N distal were significantly less than those for random intervals. |
| --- | --- |

**Correlation between human and mouse of gene expression levels in blood cell types**

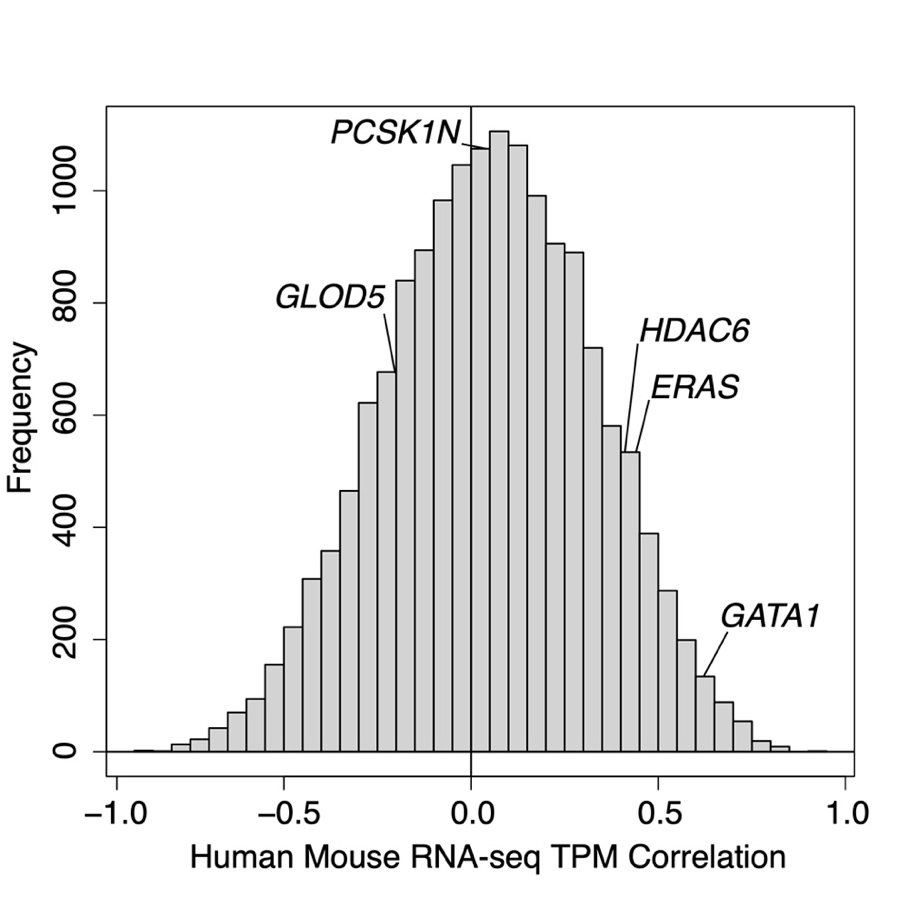

**Supplemental Figure S27.** *Distribution of correlations of RNA levels for protein-coding genes across blood cell types between human and mouse.* The correlations for the *GATA1*/*Gata1* genes and for other closely linked genes are marked with a vertical line.

**Comparison of epigenetic state landscapes between human and mouse blood cells**

To compare the epigenetic state landscapes of blood cells between human and mouse, we computed a correlation based on those state profiles. First, we constructed a vector of values for the epigenetic states across the 15 blood cell types that are analogous for both human and mouse. Specifically, we started with the epigenetic states assigned to each 200bp bin in the human and mouse genomes, based on the IDEAS joint modeling across species. Those states provided a quantitative profile of the eight epigenetic features used in the modeling (Supplemental Fig. S28A). Thus, for each 200bp bin, a vector was constructed, comprised of the eight values for epigenetic features in the state assigned to a cell type, for all 15 cell types. This process produced a vector of 120 (i.e., 8x150) values (Supplemental Fig. S28B). Second, we computed the correlations across those vectors in mouse and human, comparing all 200bp bins across a locus in human with all 200bp bins across the orthologous locus in mouse. This local, all *versus* all comparison gave an estimate of the similarity in epigenetic landscapes across species (Figure 6B). Note that genomic sequence is not included in this comparison.

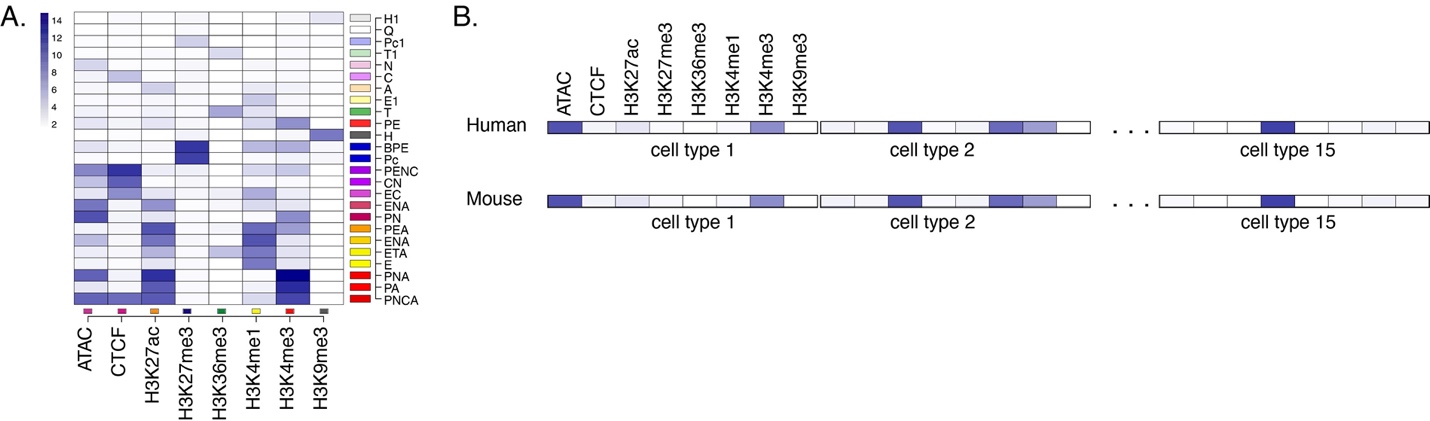

**Supplemental Figure S28.** *Computation of a correlation between epigenetic state landscapes of human and mouse blood cells.*

**Decomposition of the correlation matrix of epigenetic states using nonnegative matrix factorization (NMF)**

*Method for NMF*

To uncover the underlying structure of the correlation matrix of epigenetic states across cell types and to discern relationships between different genomic regions in both human and mouse, we applied Nonnegative Matrix Factorization (NMF) to decompose the correlation matrix. NMF is a dimensionality reduction technique that can efficiently decompose a non-negative matrix into two lower dimensional matrices. Specifically, given an m-by-n correlation matrix, R0, where 'm' represents the number of genomic bins around the target gene in humans and 'n' represents those around the orthologous target gene in mouse, we first converted the correlation matrix R0 to a non-negative correlation matrix R by setting all non-positive values in R0 to a small value of 1x10^-10^. This modified matrix, R, was then decomposed into two matrices - an m-by-k matrix (A) and an n-by-k matrix (B), such that R approximates the product of A and the transpose of B (R ≈ AB^T^). The matrices A and B were learned by using the multiplication update rules presented by Lee and Seung (1999). For the number of factors in NMF, we chose a value of 6 based on the 'elbow' method, which corresponds to the inflection point on the plot of the Bayesian Information Criterion (BIC) versus number of factors K (Supplemental Fig. S29).

| 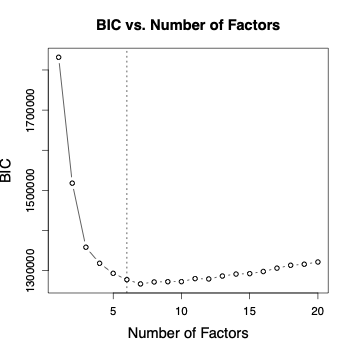 | **Supplemental Figure S29.** *Determination of the number of factors used in NMF*. The x-axis represents the range of numbers of factors tested in various NMF iterations, and the y-axis plots the BIC score for each iteration. The number of factors chosen equals 6 (black dashed line), which corresponds to the "elbow" point where the BIC score curve starts to plateau. |
| --- | --- |

Each of the six NMF factors captured a distinct, non-overlapping set of epigenetic features associated with a particular process in gene expression or regulation (Supplemental Fig. S30).

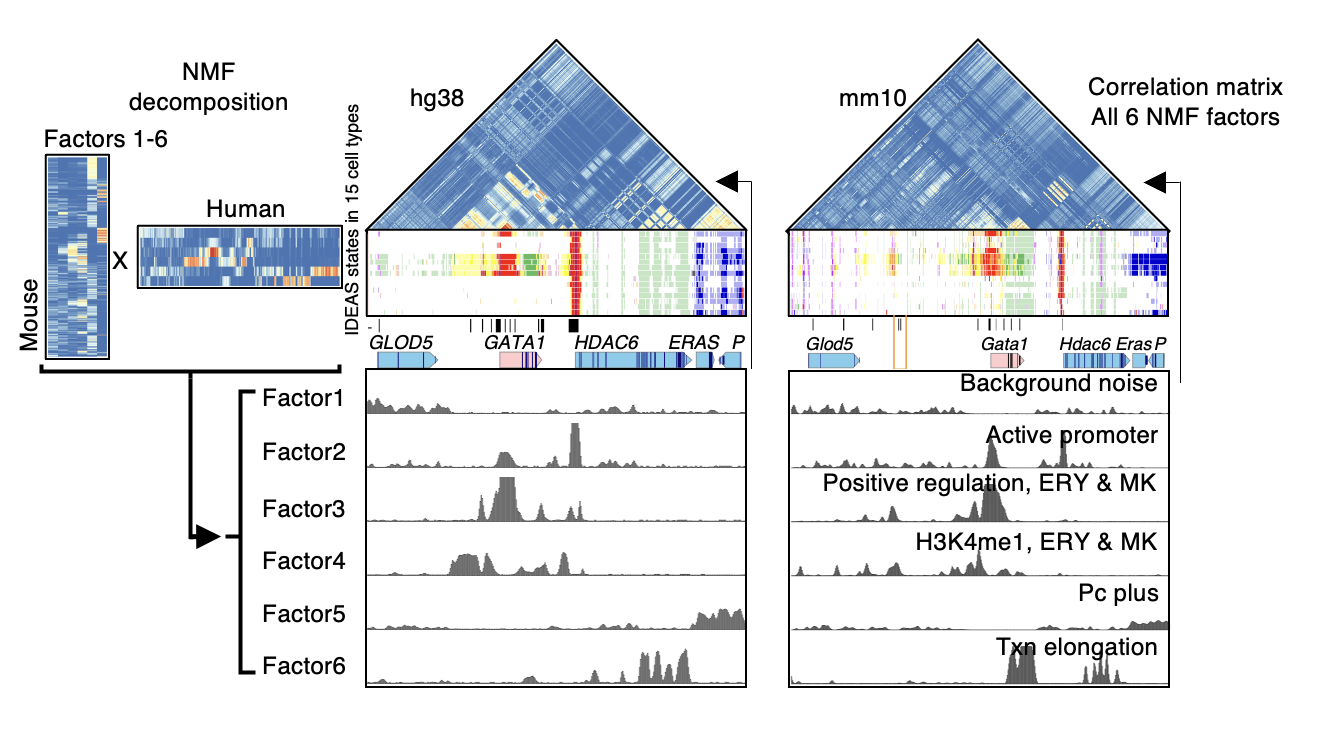

**Supplemental Figure S30.** *Profile of each of the six factors from NMF, highlighting difference processes contributing to each factor*.

To identify genomic regions with high loading to each NMF factor, we employed a *Z*-score based method. One NMF factor k is associated with two NMF loading score vectors: S_k,human_ is the k-th column of the matrix A, which includes the NMF loading scores to factor k for all genomic bins near the target gene in human, and S_k,mouse_ is the k-th column of the matrix B, which includes the NMF loading scores to factor k for all genomic bins near the target gene in mouse. We first smoothed the S_k,human_ and S_k,mouse_ vectors using the loess function in R (with a “span” parameter set at 0.03) to mitigate noise. Subsequently, we implemented two approaches to pinpoint genomic bins with a high loading to this NMF factor:

*FDR adjusted Z-score based approaches* ***without*** *background gene set adjustment*

We first applied a two-round *Z*-score process to pinpoint genomic bins with a high loading to this NMF factor. For the S_k,human_ (or the S_k,mouse_) vector, we first transformed it into its corresponding *Z*-score vector and associated p-values (round 1). Given our assumption that bins contributing to a particular NMF factor should exhibit significantly higher *Z*-scores relative to other bins, we recalculated the *Z*-scores for all bins (round 2). This was achieved by excluding those bins with round 1 *Z*-score p-values < 0.01 during mean and variance calculations. Following this, we further adjusted the round 2 p-values by using the False Discovery Rate (FDR), and those genomic bins with FDR < 0.1 were selected as bins that have significantly higher loading to the NMF factor k. This procedure was independently done in both human and mouse for each NMF factor. As a result, for both human and mouse, we identified a set of genomic bins that have significantly higher loadings to the NMF factor k. We further define the two sets of genomic regions in human and mouse as the peak regions exhibiting common cross cell type epigenetic state patterns (positive regulatory dynamics for *GATA1*/*Gata1* gene loci) between the two species.

*NMF factors in human and mouse capture a similar set of epigenetic features across cell types*

In this analysis, we interpreted the regions within the same Non-negative Matrix Factorization (NMF) factors as genomic regions in the two species that show highly similar cross-cell type epigenetic state patterns. To validate this interpretation, we computed the average correlation between the regions with high loading scores of each pair of the NMF-factors across the two species. Specifically, we first selected genomic bins that exhibit high NMF loading scores (exceeding the 90th percentile) as representative bins for each factor in each of the species. Then, for a pair of the NMF-factors, factor-i and factor-j, the correlation between the two species was computed as the average correlation of epigenetic states across cell types between the representative bins of factor-i in human and the representative bins of factor-j in mouse. As illustrated in Supplemental Fig. S31, each NMF factor in human (each column) exhibits one of the highest average correlations against the same NMF factor in mouse (each row). We conclude that the NMF factors capture a similar set of epigenetic state patterns in each species.

| 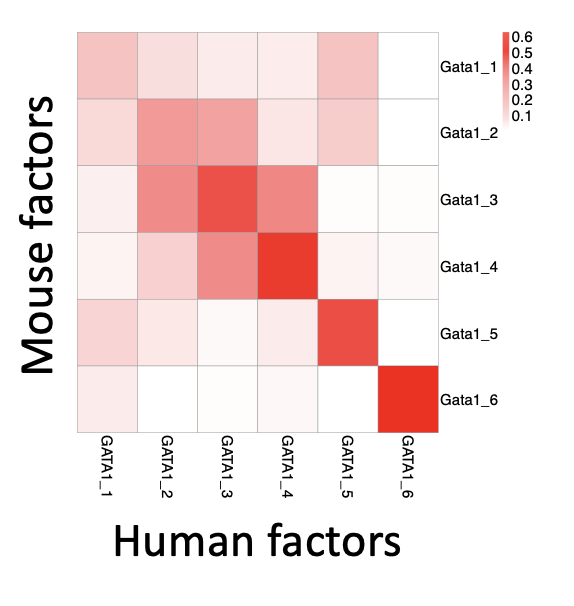 | **Supplemental Figure S31**. The average correlation matrix between human NMF factors and mouse NMF factors. Each entry in the matrix shows the average of Pearson’s correlation coefficients of epigenetic states across cell types between the representative (high scoring) bins of factor-i in human and the representative bins of factor-j in mouse. |
| --- | --- |

*FDR adjusted Z-score based approaches* ***with*** *background gene set adjustment*

Some of the cross-cell-type epigenetic state patterns are more commonly shown across the genome, and thus the previous approach is likely to identify some of these patterns as false positive discoveries that would be found in comparisons of random loci. To reduce the impact of this issue, we refined our approach by incorporating a background gene set adjustment. Specifically, we began with calculating the correlation matrix between human-target gene and mouse-target gene (R_human-tar-vs-mouse-tar_) and decomposing this correlation matrix into two NMF matrices (R_human-tar-vs-mouse-tar_ ≈ AB^T^). Next, instead of directly converting the A matrix's S_k,human_ vectors into *Z*-scores using mean and variance from the A matrix, we revised our method to include an initial selection of 100 random mouse genes. For each of 100 randomly selected mouse genes, we calculated its correlation matrix with the human-target gene, denoting each as one of the background correlation matrices (R_human-tar-vs-mouse-bg-i_). We then decomposed the R_human-tar-vs-mouse-bg-i_ using NMF, keeping the A matrix constant as derived from the original correlation matrix between human-target gene and mouse-target gene decomposition (R_human-tar-vs-mouse-bg-i_ ≈ AB_bg-i_ ^T^). Subsequently, for each S_k,human_ vector, we computed the corresponding *Z*-score vector and associated p-values. The mean and variance for these calculations were derived from all 100 S_k,mouse-bg-i_ vectors. The *Z*-scores were then converted to p-values, which were then adjusted using the FDR. Genomic bins with an FDR < 0.1 were identified as having significantly higher loading scores to the NMF factor k. The same analysis was also done for the B matrix to identify genomic bins with significantly higher loading scores to all NMF factors in mouse. In summary, this approach mitigated the false positive discoveries for the cross-cell-type pattern that are commonly presented across the genome in this analysis. The effectiveness of this bias reduction is demonstrated in Supplemental Fig. S33, as presented in the next section “False discovery rate in finding related elements using NMF of correlation matrices of epigenetic states”.

*Robustness of epigenetic patterns captured by NMF factors*

We found that the identification of the patterns characteristic of factors 3 and 6, discovered using *k*=6 for NMF, was robust to the choice of *k*. The factors identified using *k* ranging from 4 to 9 included factors with patterns similar to those of factors 3 and 6 (from *k*=6), as shown in Supplemental Fig. S32A. Using high values of *k* (15 and 20) decomposed these patterns into multiple factors (Supplemental Fig. S32B) but using these high values for *k* was not supported by the BIC analysis (Supplemental Fig. S29).

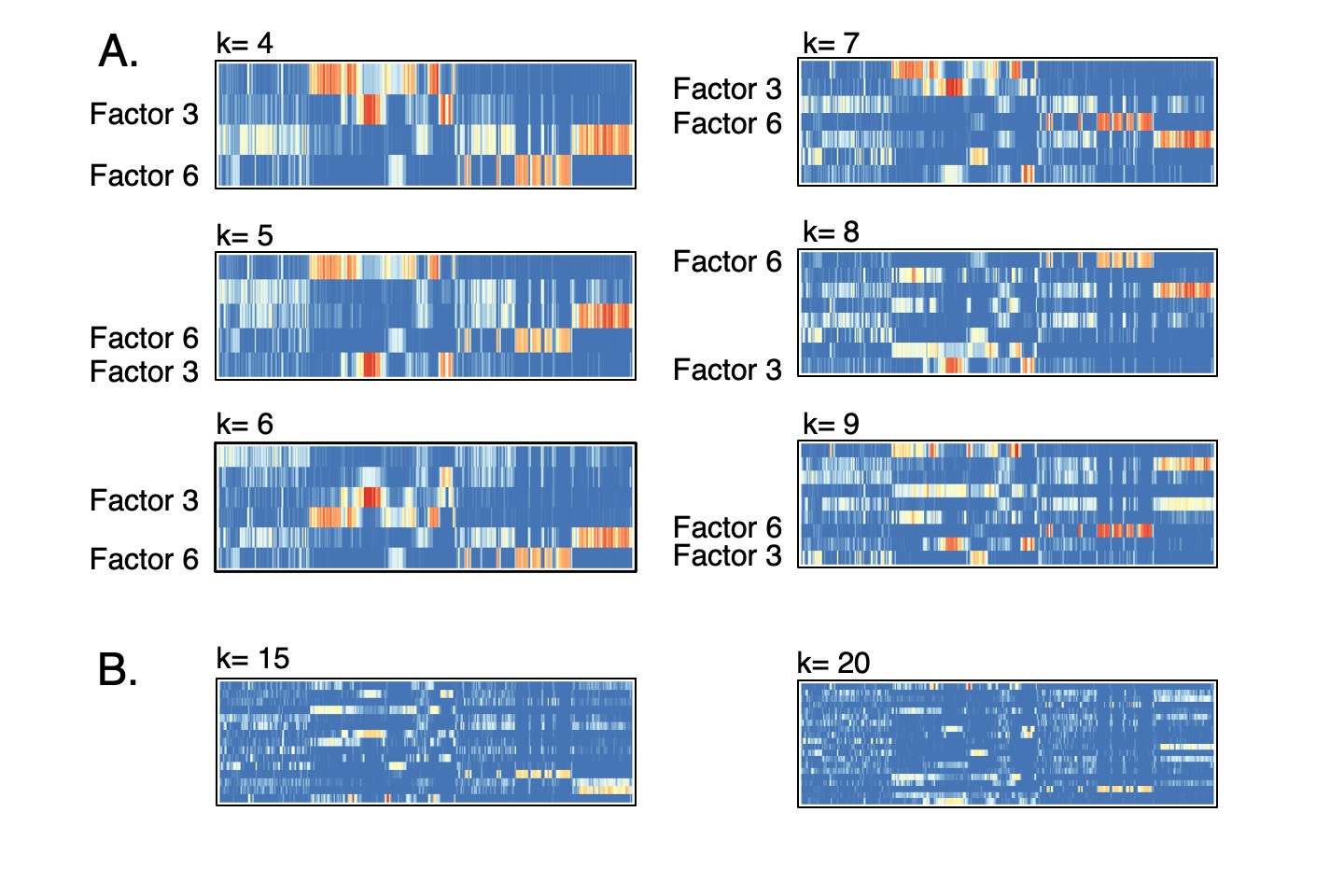

**Supplemental Figure S32.** *The patterns discovered for NMF factors 3 and 6 are robust to the choice of k.* The intensities of signals for the NMF factors from decomposition of the correlation matrix for interspecies comparisons of epigenetic states along the region around the human *GATA1* gene (see main Figure 6) are shown for various values of k used in NMF. The factors corresponding to factor 3 and factor 6 (discovered using *k*=6) are labeled in the results for each value of *k*. (**A**) Results for using *k* from 4 to 9. (**B**) Results for using *k*=15 and *k*=20.

**False discovery rate in finding related elements using NMF of correlation matrices of epigenetic states**

*Statement of the problem*

Having shown that comparisons of the epigenetic landscape between species can reveal elements with similar roles in regulation, even in the absence of genomic sequence alignment, we wanted to estimate the frequency at which these interspecies correlations of epigenetic states across cell types would lead to false discovery. The question of what constitutes a false positive in this context is challenging. A major complication is the well-established association of certain histone modifications with particular processes in gene regulation and expression. Those processes may not show gene specificity, as illustrated by the presentation on NMF factor 6 in Fig. 6 F and G, in which a common function of transcriptional elongation across most cell types is assigned to two different intervals in the same 100kb region, specifically downstream of *GATA1* and the *HDAC6* gene. Of course, one expects to find a high correlation in the epigenetic states across cell types between any genes – orthologous or non-orthologous - in which the H3K36me3 modification is left during elongation in most cell types. Such a case points to a common mechanism of expression or regulation across most cell types. The interesting new discovery in the state correlation and NMF decomposition was that some NMF factors reveal specificity in regulating the same gene or regulating genes with a similar cell type-specific expression profile. In considering the issue of false discovery, the pertinent question is whether the epigenetic comparisons reveal a common function with some specificity. To investigate this issue of potential false discovery, we examined the correlation matrices between cross-cell type epigenetic state profiles of one human locus and those from multiple non-orthologous loci in mouse, under the assumption that elements with high correlations at non-orthologous loci were candidates for false discovery. When evaluating the results, we distinguished between two related goals. The initial goal was (1) to find *tissue-specific regulatory elements that regulate a particular pair of orthologous genes* between species*.* Our examination of epigenetic state correlations across cell types in non-orthologous loci led to the articulation of a second goal, which was (2) to reveal genomic elements that appear to be involved in a common function as *tissue-specific regulatory elements*. The estimate of false discovery rate differs depending on the stated goal.

In the following presentation, we began with a broad assessment in which all discovery between non-orthologous loci was considered a “false positive” (using goal 1 of finding regulatory elements for orthologous genes), which in turn led to an improvement in the method used for calling peaks in the NMF factors. We then assessed some of the signals driving the apparent “false discoveries”, focusing on factors with specificity for regulation in erythroid and megakaryocytic cells. Finally, we show that an apparent “false discovery” (by goal 1) in a non-orthologous locus was actually pointing to a previously unrecognized genomic region with multiple hallmarks of erythroid regulation, and hence it is a candidate for a true discovery by the criterion of goal 2, i.e., finding tissue-specific regulatory elements.

*Analytical approach*

As an approach to systematically evaluating false discoveries, we examined the epigenetic state correlation matrices between orthologous and non-orthologous loci. We compared the epigenetic states across cell types between the human *GATA1* gene locus and four mouse loci, the orthologous *Gata1* gene locus and three non-orthologous loci. For the non-orthologous comparisons, we chose the locus containing *Cd4* because that gene is expressed specifically in CD4+ T-cells, the locus containing *Rps19* because that gene is expressed ubiquitously in all the cell types examined, and the locus containing *Slc4a1* because that gene is specifically expressed in erythroid cells and may reveal some elements with an erythroid regulatory component in common with those identified in comparisons of the human *GATA1* and mouse *Gata1* loci. The distinct expression patterns of genes in these three non-orthologous loci should provide a diverse basis for comparison. We found that certain regions within the locus containing the human *GATA1* gene exhibit ed high state correlation with these non-orthologous mouse loci, with correlation coefficients in a range similar to those for the comparison with the orthologous mouse locus (panels A-D of Supplemental Figure S33). We then applied NMF to decompose the correlation matrices and reveal the NMF factors contributing to the high correlations (panels E-H). In this decomposition, we fixed the human *GATA1* locus factor matrix to be the one obtained from the human *GATA1* locus and mouse *Gata1* locus NMF decomposition and only updated the factor matrix for each of the non-orthologous loci.

*Initial statistical results*

We used FDR-adjusted *Z*-score p-values to identify specific intervals within these mouse loci that correlate highly with each 200 bp bin of the human *GATA1* locus (panels I-L). The state correlations associated with NMF factors 2, 3, 4, and 6 passed the threshold for several genomic bins, and thus these genomic intervals contributed to statistical discovery in this analysis. In contrast, the state correlations associated with NMF factor 5 had relatively low correlation coefficients, and no genomic intervals passed the FDR threshold for highly correlated elements (panels I-L) in any of the loci. NMF factor 1 largely captured the background correlations, none of which passed the threshold for highly correlated.

These results showed that, if one included all the windows in non-orthologous loci with significant state correlations as “false positives” (all the intervals in panels J-L), then many discoveries were “false” by goal 1, approaching the number of “true positive” discoveries in the orthologous comparisons. To estimate the false-discovery rate for *GATA1*-*Gata1* comparisons, we assigned all the genomic intervals in the mouse *Gata1* locus with high correlation (panel I) as “true positives” and those in non-orthologous loci with high correlation as “false positives” (panel J, K, and L). The false discovery rates from this initial estimate were indeed high, even for NMF factors 3 and 4 that are associated with erythroid-megakaryocytic specific activation (panel Q).

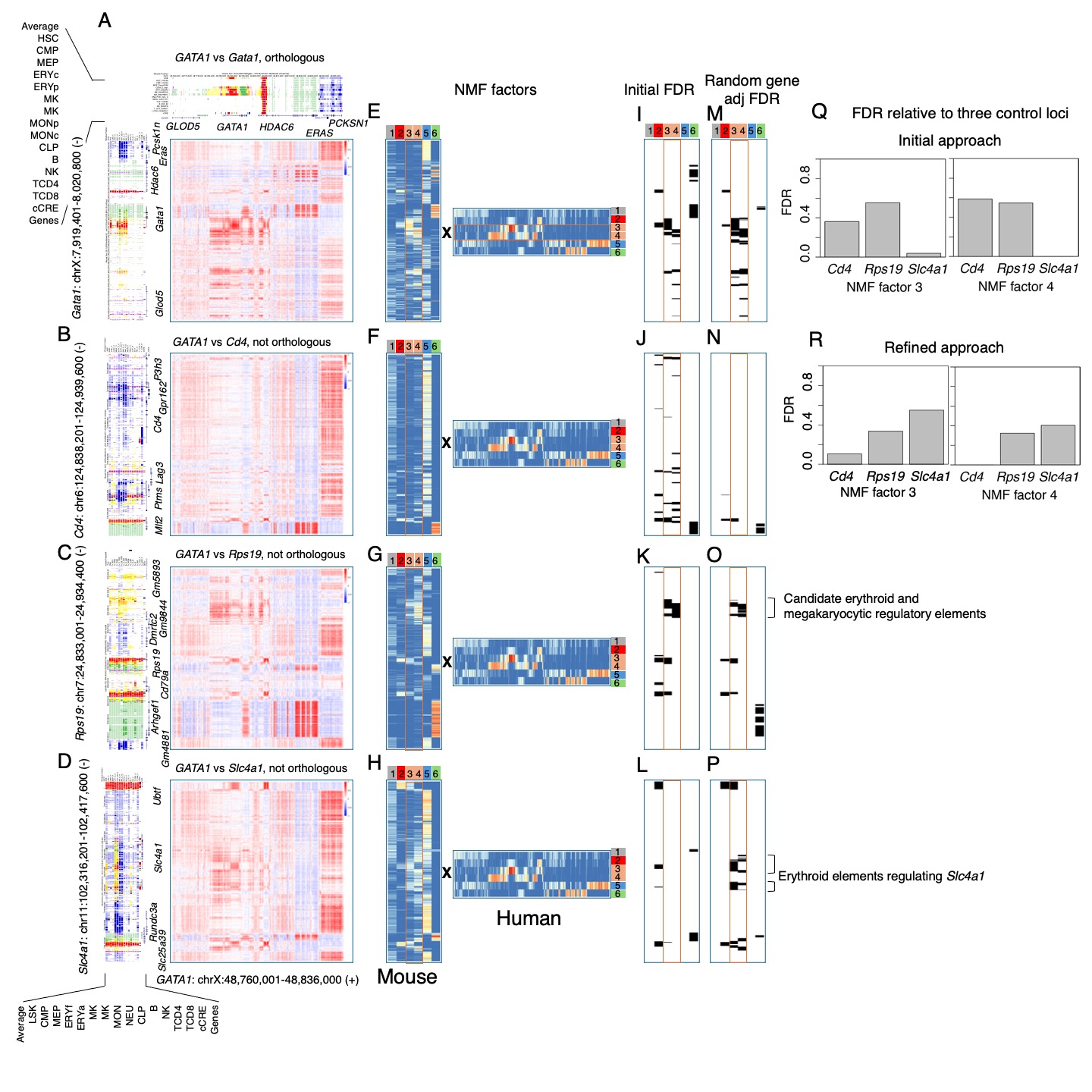
**Supplemental Figure S33.** *Assessment of false and true discovery by epigenetic state comparisons at orthologous and non-orthologous loci*.

(A-D) The state correlation matrices between human *GATA1* locus and four mouse loci, which were approximately 100kb regions containing the genes *Gata1* (orthologous to human *GATA1*, panel A), and three non-orthologous genes *Cd4*, *Rps19*, and *Slc4a1* (panels B-D). The epigenetic state landscapes from the joint human-mouse IDEAS model in 15 cell types are shown along the axes, along with gene annotations. (E-H) The factor matrices of the four correlation matrices after NMF decomposition. (I-P) The mouse genomic intervals identified as having significantly high correlation scores within each of the NMF factors. Panels I-L show the result from the initial approach, which used the mean and standard deviation of the same gene locus to establish the background for the *Z*-score and FDR calculations. Panels M-P show the result based on the mean and standard deviation determined from a diverse set of loci for the background. (Q and R) The FDR of identified mouse genomic intervals with high epigenetic state correlations in the *Gata1* locus based on NMF factors 3 and 4 relative to the three comparison, non-orthologous loci. (Q) The top panel is the result based on the mean and standard deviation of the same gene locus (initial method). (R) The bottom panel is the result based on the mean and standard deviation of the diverse set of loci for the background (refined method).

*Refinement of the criterion for discovery*

These initial results suggested that incorporating epigenetic state correlations between non-orthologous loci could improve our approach to finding highly correlated intervals. Our initial method relied solely on the state correlation matrix between the human *GATA1* locus and *one* specific mouse target locus for the *Z*-score calculation, and we refined the method to incorporate the state correlation matrices between the human *GATA1* locus and *multiple* other mouse loci. The examination of a diverse set of loci was expected to provide a more robust estimate of the background correlations. First, we randomly selected 100 loci of about 100kb surrounding different mouse genes to use as background loci and generated the correlation matrices between the human *GATA1* locus and these mouse background loci. Then, we decomposed the correlation matrices by NMF, again fixing the human *GATA1* locus factor matrix as the one obtained from the human *GATA1* locus and mouse *Gata1* locus NMF decomposition, and only updating the factor matrix for each of the background loci. Next, we used the output of the factor matrices from the 100 background loci to establish a background mean and standard deviation for recomputing the *Z*-score for the factor matrices for the foreground gene loci (*Gata1*, *Cd4*, *Rps19*, *Slc4a1*). In this refined approach, we selected the genomic intervals based on the 10% false discovery rate relative to the 100 background genes.

The impact of this refinement is evident in panels M-P and panels Q vs. R. This refined approach had little impact on the “true discovery” in the orthologous comparisons (compare panels I and M), but it did exclude much of the “false discovery” at the *Cd4* locus (compare panels J and N) while increasing the discovery at the *Slc4a1* locus (compare panels L and P), which was expected to have some regulatory elements similar to those at the *Gata1* locus. We note that the “false discovery” at the *Rps19* locus was not strongly impacted by the refined approach. This recalibration of the background reduced the estimated false discovery rate for comparisons between the human *GATA1* locus and the mouse *Cd4* locus (compare panels Q and R); this estimated rate for NMF factor 3 was reduced from about 0.4 to about 0.1 while the estimated FDR for NMF factor 4 was reduced from about 0.6 to almost zero. The limited impact on the estimated false discovery rate for NMF factors 3 and 4 for comparisons with the *Rps19* locus and the increased rate for comparisons with the *Slc4a1* locus appear to reflect erythroid and megakaryocytic gene regulatory regions, and thus they could be considered “true discovery” by goal 2 (finding tissue-specific regulatory elements), as explained in the next subsection.

*Biochemical interpretations and a candidate for true discovery in a non-orthologous locus*

The statistical analysis using a simple rule for distinguishing “true” from “false” positives gave a low estimate for a false discovery rate for comparison with one non-orthologous locus (panel R), but several discoveries at the non-orthologous loci *Rps19* and *Slc4a1* remained or were enhanced by the refined approach to finding strongly correlated genomic intervals. A deeper examination of the biochemical signatures and epigenetic features driving these correlations improved the interpretation of apparent “false” discoveries (by goal 1) and led us to propose that many appeared to be discoveries of tissue-specific regulatory elements, and hence would be considered “true” discoveries by goal 2.

The state correlations driven by NMF factors 3 and 4 were of particular interest because they were associated with gene activation specifically in erythroid and megakaryocytic cells (Supplemental Fig. S30), just like the *GATA1* gene that was in the center of the human locus used in all the comparisons. The non-orthologous locus containing *Cd4* has no genes expressed specifically in erythroid cells, and the epigenetic state landscape, displayed along the vertical axis of panel B, does not show any regions with an erythroid or megakaryocytic pattern of active states. Most of the genomic intervals with correlations associated with NMF factors 3 and 4 were no longer considered discoveries when the multiple random gene background was adopted for the FDR-adjusted Z score p-value (panel N). In contrast, the gene *Slc4a1* is abundantly expressed specifically during late erythroid maturation, which was reflected in the pattern of active epigenetic states only in erythroid cells (y-axis of panel D). Thus, we thought we might see regulatory regions for *Slc4a1* to have a signal for factors 3 and 4, despite the fact that this gene is not expressed in megakaryocytes, whereas the reference gene (*GATA1*) is expressed in both erythroid and megakaryocytic cells. Indeed, the mouse *Slc4a1* gene and surrounding regions showed state correlations associated with NMF factors 3 and 4 (panel H), but none passed our initial FDR threshold for highly correlated elements (panel L). However, the refined approach revealed several highly correlated elements for NMF factors 3 and 4 (panel P). These elements (indicated on panel P) correspond to regulatory regions demonstrated experimentally to regulate expression of the mouse *Slc4a1* gene (Fox et al. 2020). Thus, while the estimated false discovery rate for NMF factors 3 and 4 are high (about 0.5 and 0.4, respectively) for comparing the human *GATA1* and mouse *Slc4a1* loci, we would argue that most of these discoveries at the non-orthologous locus can be considered “true” discoveries of erythroid regulatory elements (goal 2).

The discovery of elements with high correlations associated with NMF factors 3 and 4 in the comparison between the human *GATA1* and mouse *Rps19* loci were initially puzzling. We had chosen this locus to represent one with genes commonly expressed in all the cell types examined, and indeed the epigenetic state landscape reflects the common expression of genes *Arhgef1* and *Rprs19*, with active promoter states (red) and transcriptional elongation states (green) for all cell types included (y-axis of panel C). The elements with highly correlated epigenetic states and associated with NMF factors 3 and 4 were located in a different part of the locus, between the genes *Gm9844* and *Gm5893* (indicated on panel O). Although these genes are not known to be expressed specifically in erythroid or megakaryocytic cell types, the epigenetic state pattern in this region showed states associated with enhancers (yellow) and promoters (red), with a strong signal in the erythroid and megakaryocytic cells. To obtain further experimental evidence as to whether this region could contain elements with a potential role of regulation in erythroid and/or megakaryocytic cell types, we examined the occupancy profile of transcription factors and co-activators active these two cell types in mouse. The transcription factors GATA1 and TAL1 are major regulators of gene expression in erythroid and megakaryocytic cells, and we found that multiple sites within this region were shown by ChIP-seq to be bound by these transcription factors in erythroid cells, and a different site was bound by them in megakaryocytic cells (Supplemental Fig. S34). These sites bound by GATA1 and TAL1 in erythroid cells were also bound by the co-activator EP300 and the chromatin modulator BRD2. Thus, we infer that the “false” discovery (by the goal 1 criterion of all discovery at non-orthologous loci being false, panel R of Supplemental Fig. S33) actually pointed to a previously unrecognized region with a plausible potential regulatory role specifically in erythroid and megakaryocytic cells, which would be a “true” discovery by goal 2.

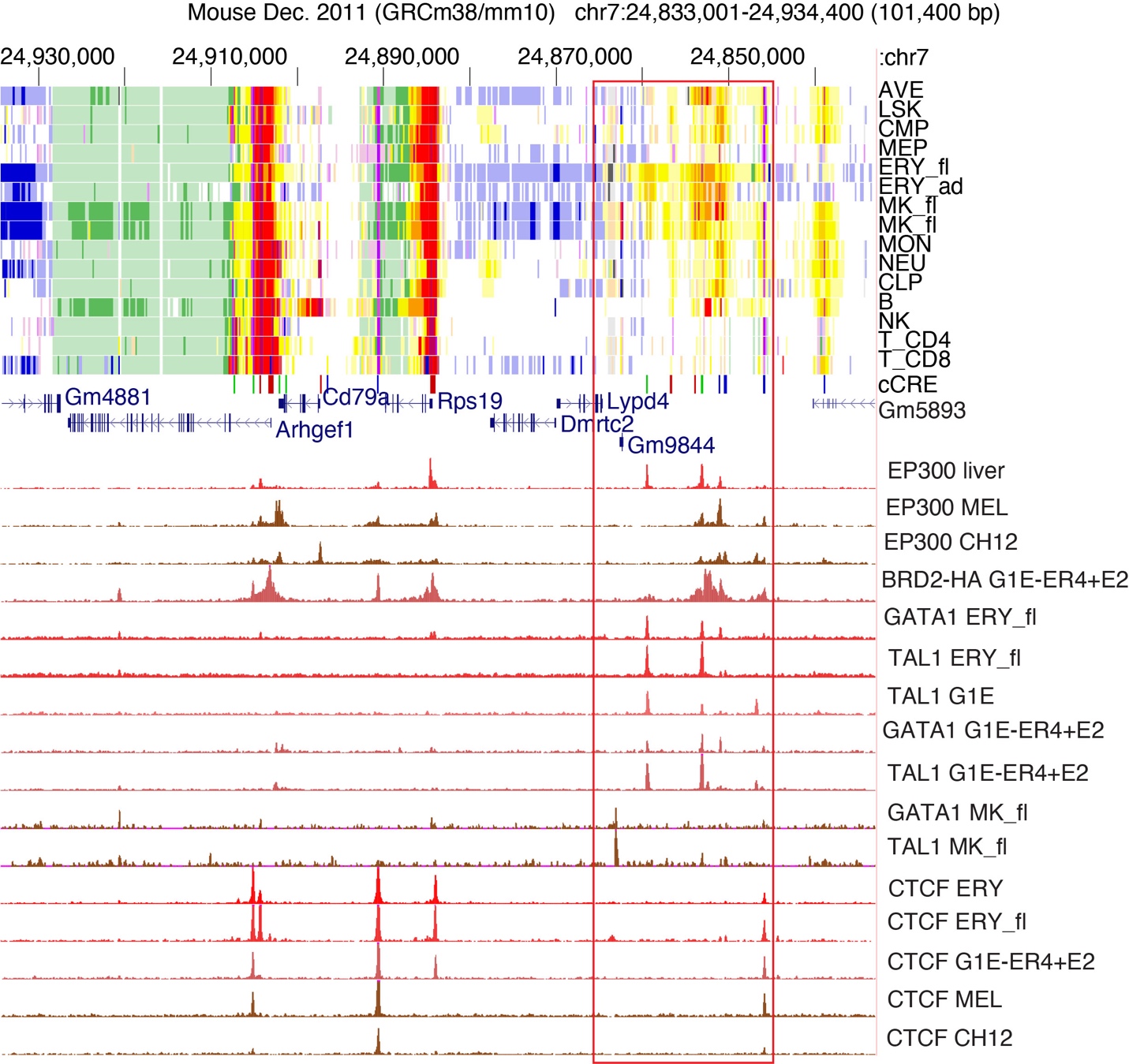

**Supplemental Figure S34.** *A candidate for a discovery of a potential element regulating expression in erythroid and/or megakaryocytic cells based on correlations of epigenetic states at non-orthologous loci.* The 101kb mouse genomic locus containing the *Rps19* gene is shown with (top to bottom) the epigenetic state assignments from the joint human-mouse IDEAS model in 15 blood cell types, the VISION cCREs, the gene annotations, and ChIP-seq profiles for occupancy by the cofactor EP300 in an erythroid tissue (fetal liver) and two cell lines (erythroid MEL and B-cell derived CH12), an HA-tagged form of the chromatin modulator BRD2 in an erythroid cell line model (G1E-ER4+E2), the transcription factors GATA1 and TAL1 in several erythroid or megakaryocytic cell tissues or cell lines, and the structural protein CTCF in several erythroid tissues or cell lines and CH12. Abbreviations include: fl = fetal liver, MK = megakaryocyte, ERY = erythroblast. The red rectangle encompasses the region with chromatin signatures of epigenetic features associated with erythroid and megakaryocytic regulation.

Much of the high state correlation was contributed by epigenetic states around genes expressed or regulated by a shared process *across many cell types*. For example, NMF factor 5, which is dominated by the polycomb mark H3K27me3, accounted for much of the high correlation of states between the *ERAS* and *PCSK1N* genes in the human *GATA1* locus and the non-orthologous mouse loci containing *Cd4* (almost all the genes except *Mlf2*), *Rps19* (the *Dmrtc2* and *Gm4881* genes), and *Slc4a1* (the *Rundc3a* gene and some intergenic intervals) (Supplemental Fig. S33). These intervals with polycomb-modified chromatin across most cell types clearly contributed to the state correlations (panels B-D) and NMF factor 5 (panels F-H)). Another example is NMF factor 6, which is dominated by the elongation mark H3K36me3. Factor 6 accounted for additional correlations of epigenetic states around the *HDAC6* gene in the human *GATA1* locus with specific genes in the non-orthologous loci, such as the *Mlf2* gene in the *Cd4* locus and the *Arhgef1* gene in the *Rps19* locus. Similarly, genomic intervals with states associated with active promoters across all cell types (NMF factor 2) were found in the orthologous and non-orthologous comparisons. We interpret these correlations accounted for by NMF factors 5, 6, and 2 as reflecting the common functions *across cell types* of polycomb-mediated repression of some genes (factor 5) elongation of transcription at expressed genes (factor 6), or active promoters (factor 2). We would argue that these apparent “false” discoveries (by the criterion of all discovery at non-orthologous loci being false) actually indicated a discovery of regulation or expression by these common functions broadly across cell types.

**Effectiveness of interspecies sequence alignment and epigenetic state correlation for regulatory element discovery**

*Statement of the issue*

By utilizing chromatin accessibility and epigenetic state annotation in blood cells in our VISION project, we have predicted cCREs without regard to their sequence conservation between species (human and mouse). The collection of VISION cCREs provided an opportunity to estimate how much true discovery will be missed by focusing only on genomic elements in which the DNA sequence aligns between species. Given that regulatory elements have been discovered in transposable elements that are found only in human (or only in mouse), it is apparent that some regulatory elements would be missed by requiring interspecies sequence alignments. We wanted to use the VISION cCRE collection to assess how large the impact could be.

*Approach to analysis*

Under the assumption that the true discovery of our interspecies analysis lies in identifying specific DNA elements with regulatory functions demonstrated experimentally, we used a large, almost comprehensive collection of short DNA elements shown to be active in a robust assay for elements that regulate reporter gene expression as a proxy for the true discovery. Agarwal et al. (2023) used an enhanced lentivirus massively parallel reporter assay (lentiMPRA) to test over 200,000 DNA segments for activity in boosting or reducing gene expression after integration into three different cell lines. We focused on the results from K652 cells, in which all promoters for protein coding genes and a close to comprehensive set of candidate enhancers (based on DNase-seq peaks from ENCODE), along with many controls, were tested in forward and reverse orientations. The activities of each construct in the lentiMPRA library were reported as mean activity scores, which incorporated results from multiple barcoded constructs for each DNA fragment and running the assay in triplicate. A total of 87,186 lentiMPRA constructs produced scores that were interpreted as reflecting an element active in this assay, i.e., the mean of the activity scores for the three replicates passed a threshold (-0.117) that corresponds to an FDR of 5%. Those active scores were generated from a set of 57,061 elements, reflecting the fact that some elements were active in both orientations.

*A larger fraction of elements active in lentiMPRA are in chromatin with a non-quiescent state than are in DNA segments that align between human and mouse.*

That collection of 57,061 elements shown to be active in at least one orientation in the lentiMPRA in K562 cells can be considered a proxy for a ground truth to ascertain the impact and efficacy on true discovery of (a) requiring that DNA sequences align between human and mouse and of (b) requiring an epigenetic state or esRP score in K562 cells indicative of activity in expression or regulation. For each of the active elements, we determined whether the human DNA sequence aligned with the mouse genome assembly using the liftOver tool. We found that 34,434 (60.8%) had an aligning sequence in the mouse genome, but the remaining 22,627 (39.2%) elements did not align to mouse. These results show that requiring sequence alignment will miss a large portion (about 40%) of elements active in lentiMPRA.

In contrast, about 87% of the active elements were in an epigenetic state in K562 cells indicative of dynamic chromatin modifications and accessibility. We determined the dominant epigenetic state in K562 cells (from our joint IDEAS modeling) for 56,663 elements; this slightly smaller set does not include almost 400 active elements that are located regions of human genome to which large numbers of sequencing reads map (high signal regions or blacklisted regions). The epigenetic states were aggregated into those with histone modifications, CTCF binding, and/or nuclease accessibility, which we termed non-quiescent states, in contrast to the low signal quiescent state. As tabulated below, almost 87% of the active elements were in a non-quiescent state. After aggregating active elements by whether their states are associated with gene activation, repression, or transcriptional elongation, we found that 82.5% of the active elements were in states associated with gene activation.

| Category | Number of elements | Percent |
| --- | --- | --- |
| Non-quiescent state | 49,145 | 86.7 |
| Quiescent state | 7,518 | 13.3 |
| States associated with gene activation | 46,770 | 82.5 |
| States associated with gene repression | 1,713 | 3.0 |
| States associated with transcriptional elongation | 662 | 1.2 |

Since a greater proportion (86.7%) of the active elements are in dynamically active chromatin than are in sequences that align between human and mouse (60.8%), one can make more true discoveries by including sequences that do not align between human and mouse in the predictions of CREs. These results are summarized graphically as panel D of main Figure 5.

*Elements active in lentiMPRA are more highly enriched in DNA intervals with high cross-cell type correlations of epigenetic state with and without DNA sequence alignment to mouse.*

To verify whether our method, which is predicated on correlations derived from cross-cell type epigenetic state patterns, can effectively identify 'true discovery' DNA elements, we applied it to analyze the lentiMPRA results around 30 genes (genome coordinates and names are in Supplemental Table S6). These genes were selected based on their high or specific expression in K562 cells, while including some genes not expressed in K562 cells, such as *Cd4*. We divided a 100kb interval centered on the transcription start site (TSS) into 200bp windows, giving 501 windows (250 windows on each side of the window containing the TSS) for of each of the 30 different genes. Each window was placed into one of four distinct, mutually exclusive groups, which were defined as having: (1) both high cross-cell-type epigenetic state pattern correlation (sufficient to be called as a peak in an NMF factor from the epigenetic state correlation matrix, using as a threshold FDR<=0.1) and sequence conservation between human and mouse, as established by liftOver between the two genome assemblies (S&C); (2) high cross-cell type epigenetic state pattern correlation but not interspecies sequence alignment (C only); or (3) interspecies sequence conservation but not an NMF peak (S only). All remaining windows were placed into the fourth category, Other. For the DNA windows in each of these four categories, we calculated the enrichment of the intersection between the group region set and the collection of 57,061 active DNA elements based on lentiMPRA.

${enrichment}_{i,j}= \frac{N_{obs,i,j,lentiMPRA}+sn}{\left( \frac{N_{i,j}}{N_{i,total}}\times\frac{N_{i,lentiMPRA,}}{N_{i,total}}\times N_{i,total}+sn \right)}$,

where $N_{obs,i,j,lentiMPRA}$ is the number of 200bp windows that intersect with both $N_{lentiMPRA}$ peak regions and group-j regions in the 100kb window round the TSS of gene-i, $N_{i,j}$ is the number of 200bp bins that intersect with group-j regions in the 100kb window around the TSS of gene-i, $N_{i,lentiMPRA}$ is the number of 200bp bins that intersect with lentiMPRA peak regions in the 100kb window around the TSS of gene-i, $N_{i,total}$ is the total number of 200bp bins in the 100kb window around the TSS of gene-i, and $sn$ is an added small number to avoid zeros in the denominator. As shown in Supplemental Fig. S35A, the S&C group and C only group have significantly higher enrichment than the other two groups (pairwise Wilcoxon test p-value < 0.05).

We then delved deeper to understand why the regions that are conserved in sequence but not called as peaks in the epigenetic state correlation (S only) showed lower enrichment. One hypothesis was that these regions might predominantly be located in transcribed regions. To investigate this, we assessed the enrichment of these four sets in DNA regions exhibiting high H3K36me3 signals in K562 cells (IDEAS states 2 and 8). We found that the sequence-only conserved regions (S only) show a notable enrichment in these specific areas (Supplemental Fig. S35B).

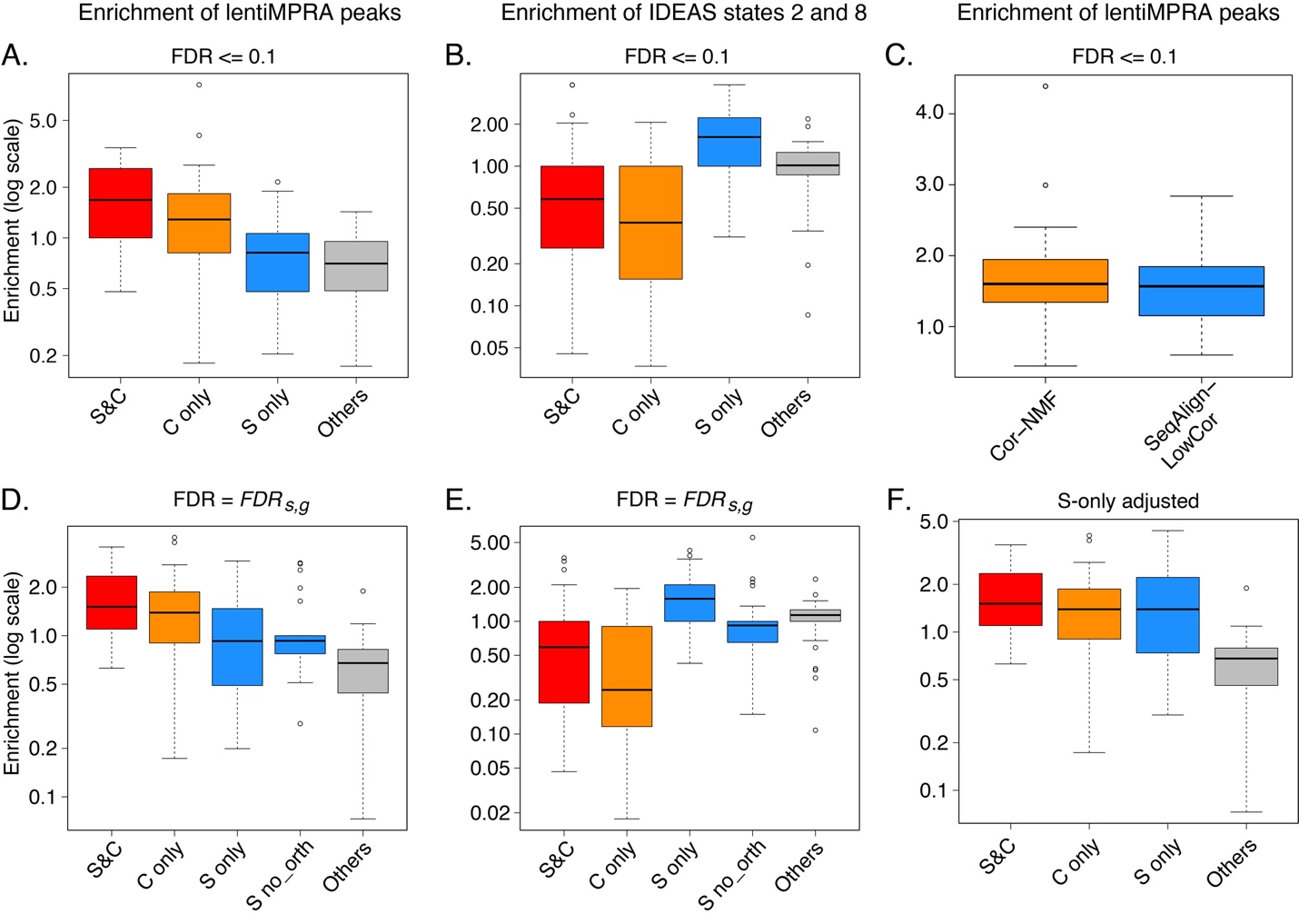

**Supplemental Figure S35**. *The enrichment of regulatory elements and regions of transcriptional elongation in four categories of DNA windows defined by sequence alignment between human and mouse and by high correlation of epigenetic state profiles between human and mouse orthologous genes.* **Panels A, C, D, and F**. The distributions of enrichment values for experimentally determined regulatory elements from lentiMPRA results in K562 cells. **Panels** **B and E**. The distributions of enrichment values for DNA regions labeled as being in IDEAS epigenetic states associated with transcriptional elongation (H3K36me3) in K562 cells. Panels **A,** **B, and C** used an FDR<=0.1 for calling peaks in the NMF signal from the epigenetic state correlation matrix. Panel **C** compares the distribution of enrichment values obtained for DNA bins with a high epigenetic state correlation between species, regardless of interspecies sequence alignments (Cor-NMF; this is the union of S&C and C only categories), versus the distributions found after restricting the analysis to DNA intervals with sequences that aligned between human and mouse, and then lowering the epigenetic state correlation threshold for calling a peak to obtain an FDR <= 0.1 in this context (SeqAlign-LowCor). Panels **D** and **E** present results of similar analyses to those in A and B, respectively, but with categories assigned to DNA intervals after matching the FDR for epigenetic state correlations to the FDR for sequence alignments (*FDR_s,g_*). **F.** The enrichment for MPRA-active elements in four categories of DNA windows (using the FDR for NMF signal from the epigenetic state correlation matrix that equals the *FDRs,g* determined by sequence alignment results), but with the enrichment of S only regions adjusted to accommodate the fact that some MPRA-active elements were in human DNA intervals that did not align with mouse. Each box plot gives a summary of the distribution of enrichment values for DNA bins in the 30 genes for the indicated feature (lentiMPRA active in A, C, D, and F and H3K36me3 state in B and E) and category of DNA windows. The categories are (S&C) both high cross-cell type epigenetic state pattern correlation and sequence alignment between human and mouse, (C only) high cross-cell type epigenetic state pattern correlation only, (S only) interspecies sequence alignment only, and (Others) all remaining windows. In the box plots, the line in the box indicates the median, and the whiskers extend to 1.5 times the interquartile range.

In summary, our findings show that, for the four categories examined, DNA regions exhibiting both sequence conservation and high similarity in cross-cell-type epigenetic state patterns are more likely to function as true regulatory elements. In cases where only one of these criteria is met, the regions with a high similarity in the cross-cell-type epigenetic state patterns tend to be more enriched in regulatory function, as revealed by lentiMPRA assays. The observed low enrichment in the lentiMPRA intersections for regions with only sequence conservation implies that these intervals are less likely to be involved in gene regulation, while their enrichment for the H3K36me3 modification associated with elongation indicates that they are frequently transcribed.

*Using NMF peaks from epigenetic state correlations between species does not inflate the false discovery rate compared to limiting the analysis to sequence-aligning regions.*

We used the set of elements active in lentiMPRA to show that our inclusion of all DNA windows in the epigenetic state correlation analyses does not increase the false discovery rate compared to a method that restricted the analysis to DNA segments that aligned between human and mouse. We used an “aligning positions and lowering the correlation threshold” approach to identify sequence-conserved and epigenetically correlated regions between human and mouse. Our analysis procedure was designed to achieve the same FDR as was used in our epigenetic-state-correlation-NMF method. For each gene locus (100kb DNA intervals centered on the transcription start sites), we first identified human DNA regions that can be aligned to the mouse genome by using the liftOver tool. Here, we adjusted the minMatch threshold in the liftOver step from 0.9 to 0.8 to ensure that a sufficient number of DNA bins were included as aligning so that we can reach the same FDR as used in the epigenetic-state-correlation-NMF method (FDR=0.1). We then calculated the correlation of epigenetic state profiles across cell types between these sequence-aligning human regions and their corresponding mouse counterparts. Subsequently, we ranked the human regions (bins) based on their correlation scores. The top-ranked bins in each locus were selected as predicted positives, using the epigenetic state correlation value that gives an FDR of 0.1 as a threshold. The required correlation threshold was computed for each gene locus. This implementation of an “aligning positions and lowering the correlation threshold” method ensured that we identified the subset of sequence-aligning DNA bins that had an epigenetic state correlation value sufficient to achieve the same FDR as that used in our epigenetic-state-correlation-NMF method (FDR<=0.1).

The enrichment for the elements active in the lentiMPRA assay was then computed. As shown in Supplemental Fig. S35C, using the same FDR, our epigenetic-state-correlation-NMF method, which includes all bins with high epigenetic state correlation regardless of interspecies sequence alignment, shows a distribution of enrichment values for lentiMPRA active elements comparable to that shown after restriction to sequence-aligning DNA bins with a lower epigenetic state correlation (Paired Wilcoxon test p value = 0.14). If the rate of false discovery had increased, then the distribution of enrichment values would have decreased. These similar distributions of enrichment values show that our epigenetic-state-correlation-NMF method does not increase the false discovery-rate compared to the analysis restricting discovery to sequence-aligning regions.

*The strong enrichment for MPRA active elements in DNA sequences with high interspecies epigenetic state correlation is robust to different methods of computing FDR.*

We further investigated the enrichments using an approach that matched the FDR for epigenetic state correlation signals with an estimated FDR for DNA sequence alignments. We devised a procedure to use a property of DNA-sequence aligned regions to compute a sequence alignment based FDR, and then to applied that FDR as the threshold to recompute the epigenetic state correlation-based measures. This procedure varied the FDR thresholds for each gene but ensured the same FDR was used to call conserved regions for both methods in a manner driven by DNA sequence aligned regions.

As is often the case when dealing with a goal for which no clear gold standard of “truth” exists, we needed to find a proxy for false discovery in DNA sequence alignments. In this context of evaluating sequence alignments and epigenetic state correlations in 200bp bins covering 100kb intervals centered on the TSSs of 30 human genes, we found that among the human DNA segments that aligned with the mouse genome, a subset of them actually aligned to mouse regions outside the 100kb interval centered on the TSS of the orthologous gene in mouse. We assigned those human aligning regions that fell outside the orthologous interval in mouse as a false discovery. We acknowledge that the biological relevance of this distinction is not clear; maybe it reflects the level of genomic rearrangement around the orthologous gene. We stress that we are only using this assignment to generate an estimate of false discovery rate in interspecies sequence alignments.

We used this proxy for sequence-alignment based false discovery to compute an *FDR_s,g_* based on the sequence alignment results for each of the 30 human genes.

${FDR}_{s, g}=\frac{N_{aligned\_to\_Mouse\_genome\_but\_ not\_mouse\_orth}}{N_{aligned\_to\_Mouse\_genome}}$,

where $N_{aligned\_to\_Mouse\_genome\_but\_ not\_mouse\_orth}$is the number of 200bp human bins aligned to mouse genome but not aligned to the corresponding mouse orthologous gene locus, and $N_{aligned\_to\_Mouse\_genome}$ is the total number of 200bp human bins aligned to mouse genome. For all 500 bins of 200bp around each of 30 human genes, on average there are about 98 bins that can be mapped to the mouse genome using the liftOver tool (liftover -minMatch=0.9). Among these 98 bins, about 8 bins on average are located outside of the corresponding mouse gene window (TSS+/-50kb). In the case of a gene with these alignment properties, the formula above gives an estimated *FDR_s,g_* = 8/98, which equals 0.079.

An *FDR_s,g_* was computed for each gene. Then the DNA bins in each gene interval were re-assigned based on the NMF signals in the epigenetic state correlation matrices using a new FDR for NMF peak calls that matched the sequence alignment-based *FDR_s,g_* for that gene. We stress that we are varying the thresholds for each gene such that the same FDR is used for both NMF signals in the epigenetic state correlation matrices and for DNA sequence alignments.

The results after these re-assignments presented the same trend as seen in the first analysis, with the enrichment for MPRA-active regions highest in S&C, followed in declining order by C-only, S-only, and Other (Supplemental Fig. S35 panels D and E). However, with the *FDR_s,g_* adjustment for the state correlation FDR, the distribution of enrichment values for C only was not significantly different from that for the S only regions (Paired Wilcoxon test p-value = 0.061), despite having a higher median value. Thus, using this approach to defining FDRs, the results still support the highest enrichment for DNA bins with both high epigenetic state correlation and sequence alignment (Pairwise Wilcoxon test p-value < 0.05), and the use of either feature alone gave a similar enrichment for finding MPRA-active elements.

We also examined the impact of changing the epigenetic state correlation threshold (for calling peaks in the NMF signals) only in the DNA bins with sequences that aligned to mouse. In the analyses summarized in Supplemental Fig. S35, the S only category covered the DNA bins with sequences that aligned to mouse but lacked peak in epigenetic state correlation; thus, the enrichment values for the S only category showed the impact of reducing the epigenetic state correlation thresholds to 0 in the sequence aligned intervals. The S&C category covered the DNA bins with sequences that aligned to mouse **and** with high epigenetic state correlation, using a threshold for peak calling in the NMF signals corresponding to an FDR <=0.1 (panel **A**) or an FDR matching the *FDR_s,g_* for each gene (panel **C**). Thus, the enrichment values for the S&C category showed the impact of increasing the epigenetic state correlation threshold for the sequence aligned intervals. The box-plots for S&C and for S only showed the distributions of enrichment obtained at the two extremes of correlations for epigenetic state profiles. Lowering the correlation threshold to 0 (i.e., S only) did not produce a result that was superior to examining bins with high epigenetic state correlation but lacking interspecies sequence alignment (C only) (Supplemental Fig. S35D).

Furthermore, we took an aggressive approach to finding a metric whereby restricting the target regions to those that aligned between species might show a superior result to that obtained by examining DNA intervals with high correlation in epigenetic state profiles across cell types. Specifically, we used this failure-to-capture rate as an adjustment to the enrichments calculated for the MPRA-active regions. We applied the following formula to the S-only regions:

$${enrichment}_{i,j,adj}= \frac{{aveN}_{C, enh}}{{aveN}_{S, enh}}\times{enrichment}_{i,S-only}$$

where ${aveN}_{C, enh}$ is the average number of 200bp bins identified by our NMF-correlation based approach that intersected with enhancer (MPRA-active) regions across the 30 genes, ${aveN}_{S, enh}$ is the number of 200bp bins identified by the sequence-alignment approach that intersected with enhancer regions across multiple genes, and ${enrichment}_{i,S-only}$ is the original enrichment of S-only regions in MPRA regions for gene-i. We stress that this adjustment was applied solely to the S only DNA bins to accommodate the inability of sequence alignments to find all active elements. This adjustment was an intentional bias in favor of sequence alignment and against the state correlation scores.

With this adjustment, the distribution of enrichments in S-only bins increased, as expected, but it did not significantly exceed the distribution for the C only bins, despite the bias in the evaluation favoring S-only regions (Supplemental Fig. S35, panel F). Even with this approach that aggressively favored a restriction to sequence-aligning regions, we did not find evidence that such a restriction improved the ability to find active elements in the genome over that obtained by examining intervals with high epigenetic state correlations. At best, the category restricted to sequence-aligning regions showed an enrichment for active elements comparable to that for the intervals with high epigenetic state correlations alone (Pairwise Wilcoxon test p-value = 0.5796). Also, even with this adjustment, the S&C group still has a better enrichment than the S only set (Pairwise Wilcoxon test p-value = 0.02349), which confirmed that the value of the correlation based approach for identifying regions with similar regulatory functions in the two species.

We returned to the distinction of interspecies sequence alignments that did or did not align to the orthologous gene region (TSS+/50kb) in the comparison species to compute enrichment for MPRA active elements in DNA intervals that aligned with mouse but outside the orthologous gene region (S no_orth). The DNA intervals in the S no_orth had a lower enrichment for MPRA active elements compared with both S&C and C only (Supplemental Fig. S35, panel D). Given that the full set of DNA intervals identified in the orthologous regions is the union of S&C regions, C only regions, and S only regions, and enrichment of this union set of regions in MPRA active element was also significantly higher than the S no_orth regions (Paired Wilcoxon test p-value = 0.01271), the results indicate that inclusion of DNA intervals that align to a second species but outside the orthologous locus does not improve performance, at least within the scope of this analysis.

*Issues with restricting positive training sets to DNA intervals that align between species.*

To complement this analysis of our method based on epigenetic state correlations, we compared our results to those from another method that examined epigenetic data between species while using interspecies sequence alignments to guide the analysis. The LECIF method (Kwon and Ernst 2021) introduced a conservation score reflecting evidence of conservation at the functional genomics level. We examined the LECIF scores across mouse *Gata1* locus (with surrounding genes), and we found that the promoter for *Hdac6* and the complex proximal regulatory region of *Gata1* were assigned high to moderately high values (Supplemental Fig. S36). However, the distal enhancer for *Gata1* (marked by an orange dashed box) was assigned zero or undetectable LECIF scores. This result was expected given the convention in LECIF to assign positive labels only to DNA sequence-aligned regions in the training data. In contrast, this distal enhancer was assigned high values from our cross cell-type epigenetic state correlation method (Fig. 6 in the main text; NMF factor 3 scores in Supplemental Fig. S36). Missing such regulatory elements could be a critical problem, compromising or confounding the ability to leverage experimental work in a model organism for potential interferences into function in humans.

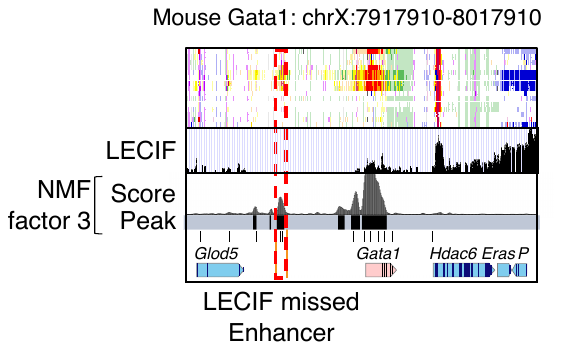

**Supplemental Figure S36.** Comparison of LECIF score tracks with NMF score tracks.

We emphasize that this limitation described for LECIF does *not* directly apply to the basic approach of the LECIF method, but rather we are pointing out that the strategy used in defining the positive set in training the LECIF model was limited to sequence-aligning regions. This restriction introduced a bias that diminished the visibility of regulatory elements in regions that did not align to the comparison species. Given that experimentally validated regulatory elements were found in DNA intervals that failed to align with the comparison species but have high epigenetic state correlations across species, we predict that the output of the LECIF and similar methods could be improved by incorporating a higher or at least equal proportion of S&C regions and C regions compared to S regions in defining their positive set. This adjustment would prevent the model from being overly influenced by the S set, thus offering a more balanced representation of both types of regions.

**Enrichment of cCREs in the three evolutionary categories for specific JmCs in the vicinity of orthologous genes**

The comparison of epigenetic state profiles across cell types also provided a means to categorize cCREs across species that did not require a match in the underlying genomic DNA sequence. We used that information to identify cCREs that may be playing a similar role in regulation in both species despite their lack of conservation in DNA sequence. Specifically, we hypothesized that most cCREs regulating expression of orthologous genes would show a similar epigenetic profile across cell types in both species, regardless of whether the element was conserved in sequence. We leveraged the membership of each cCRE in the joint metaclusters (JmCs) determined in human and mouse because those JmCs reflect the inferred activity (deduced from epigenetic states) of the cCREs across cell types, reasoning that most cCREs regulating a given gene would be in one of the JmCs found frequently in the locus. Orthologous loci in mouse and human were defined as 100 kb genomic DNA intervals centered on the TSS of a gene with an identical name in the two species. Within these orthologous loci, we calculated the enrichment of each joint metacluster (JmC) in the collection of cCREs and assessed whether each individual cCRE was a member of the enriched JmC (Supplemental Fig. S37). Thus, each cCRE was assessed both for its evolutionary history, which relied on DNA sequence alignments, and its regulatory potential deduced from epigenetic state profiles, which did not rely on DNA sequence alignments. The cCREs in these orthologous loci were assigned to a subdivision of the conservation categories; the cCREs in JmCs enriched for a specific orthologous locus were labeled SF+, S+, and N+, whereas those not in enriched JmCs were labeled SF, S, and N cCREs (Supplemental Fig. S37C). Using this approach, one can deduce that even cCREs in non-aligning genomic regions, such as the one upstream of *Gata1*, have epigenetic state profiles suggestive of a role in regulation of the orthologous gene (Supplemental Fig. S37D). Inclusion of the JmC enrichment along with the evolutionary categories increased the correlation between the esRP scores of cCREs and the expression of their inferred target genes (Supplemental Fig. S37E). The increase in correlation was observed for cCREs in all three evolutionary categories, including the species-specific N category, consistent with our hypothesis of common epigenetic profiles across cell types for relevant regulatory elements regardless of evolutionary category. These JmC enrichments provided an opportunity to delve into assessment of potential functions across species even in regions of the genome that no longer align between species.

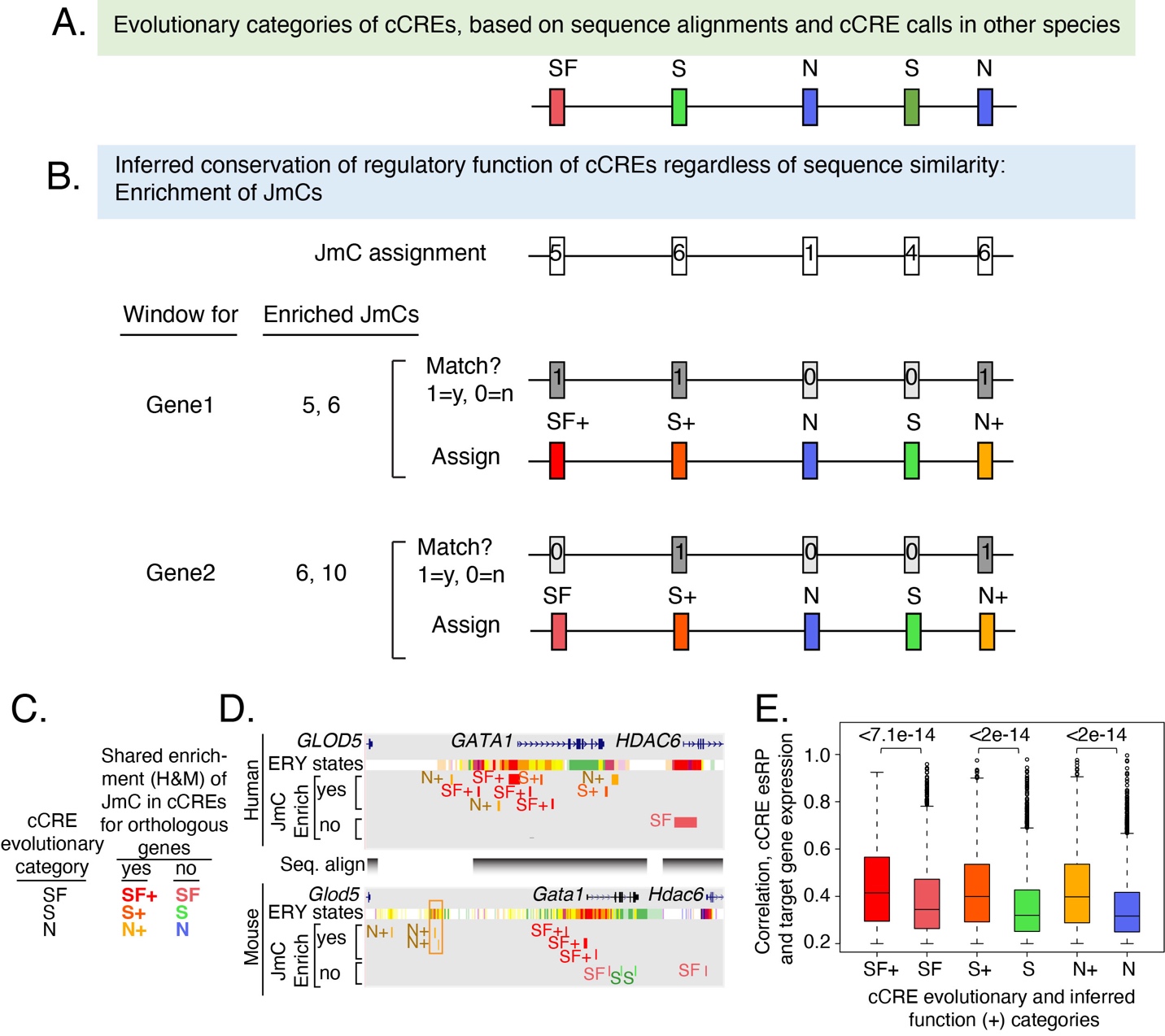

**Supplemental Figure S37.** *Enrichment of epigenetic states in orthologous loci.* **(A)** Categorization of cCREs using liftOver between human and mouse for conservation of sequence (S) and cCRE calls in human and mouse for conservation of inferred function (F). **(B)** A proxy for analogous function is a similar epigenetic pattern across cell types, which was captured as membership in the same joint metaclusters (JmCs) in human and mouse. The JmCs enriched in cCREs in each gene region (TSS+/-50kb) were determined, and then the cCREs were assessed for membership in the enriched JmCs. The JmC assignment for each cCRE does not change as windows are centered on different genes, but enrichment for that JmC can change. **(C and D)** JmC enrichment tracks of cCREs at the human *GATA1* and mouse *Gata1* gene loci. A “+” sign assigned to a cCRE indicates that the JmC to which it belongs was enriched at the GATA1/Gata1 gene locus. **(E)** The correlation between the cCRE’s esRP score and the target gene expression level. The results for each set of cCREs are shown as box plots summarizing the distribution of correlations observed for all loci with orthologous protein-coding genes. The cCREs in each evolutionary category were separated into those that are members of the JmCs enriched for a gene locus (indicated by a +) or those that are not (labeled SF, S, and N).

The enrichment threshold required to consider a JmC enriched in the vicinities of an orthologous gene pair (Supplemental Fig. S37 B-D) was determined based on the distribution of FDR adjusted p-values of the *Z* scores for the computed enrichments of all JmC enrichment for all gene pairs (Supplemental Fig. S38). Those enrichment scores were converted to *Z*-scores, and the FDR-adjusted p-values were computed from the distribution of *Z*-scores. The JmC enrichment of 2.27 was chosen as the threshold based on the FDR adjusted *Z*-score p-value = 0.1.

| 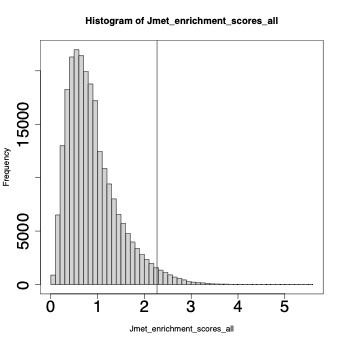 | **Supplemental Figure S38.** *The histogram of JmC gene pair enrichment scores relative to JmCs in all protein coding genes.* The black line represents the enrichment threshold for defining JmC-gene pairs. |
| --- | --- |

**CODE ACCESS**

Scripts and code used in the analyses in this paper are provided as two zipped directories in the Supplemental Material: scripts_for_GR2024_analysis for the joint IDEAS modeling and most other analyses, and cre_heritability for the sLDSC analysis. These scripts and code are also in two GitHub repositories: <https://github.com/guanjue/Joint_Human_Mouse_IDEAS_State> for the joint human-mouse IDEAS pipeline and <https://github.com/usevision/cre_heritability> for the sLDSC analysis.

**REFERENCES**

Agarwal V, Inoue F, Schubach M, Martin BK, Dash PM, Zhang Z, Sohota A, Noble WS, Yardimci GG, Kircher M et al. 2023. Massively parallel characterization of transcriptional regulatory elements in three diverse human cell types. *bioRxiv* doi:10.1101/2023.03.05.531189.

An L, Yang T, Yang J, Nuebler J, Xiang G, Hardison RC, Li Q, Zhang Y. 2019. Hierarchical Domain Structure Reveals the Divergence of Activity among TADs and Boundaries. *Genome Biology* **20**: 282.

Andersson R, Gebhard C, Miguel-Escalada I, Hoof I, Bornholdt J, Boyd M, Chen Y, Zhao X, Schmidl C, Suzuki T et al. 2014. An atlas of active enhancers across human cell types and tissues. *Nature* **507**: 455-461.

Birney E Stamatoyannopoulos JA Dutta A Guigo R Gingeras TR Margulies EH Weng Z Snyder M Dermitzakis ET Thurman RE et al. 2007. Identification and analysis of functional elements in 1% of the human genome by the ENCODE pilot project. *Nature* **447**: 799-816.

Bruse N, van Heeringen SJ. 2018. GimmeMotifs: an analysis framework for transcription factor motif analysis. *bioRxiv* doi:<https://doi.org/10.1101/474403>.

Buenrostro JD, Giresi PG, Zaba LC, Chang HY, Greenleaf WJ. 2013. Transposition of native chromatin for fast and sensitive epigenomic profiling of open chromatin, DNA-binding proteins and nucleosome position. *Nat Methods* **10**: 1213-1218.

Buniello A, MacArthur JAL, Cerezo M, Harris LW, Hayhurst J, Malangone C, McMahon A, Morales J, Mountjoy E, Sollis E et al. 2019. The NHGRI-EBI GWAS Catalog of published genome-wide association studies, targeted arrays and summary statistics 2019. *Nucleic Acids Res* **47**: D1005-D1012.

Carrillo-de-Santa-Pau E, Juan D, Pancaldi V, Were F, Martin-Subero I, Rico D, Valencia A, Consortium B. 2017. Automatic identification of informative regions with epigenomic changes associated to hematopoiesis. *Nucleic Acids Res* **45**: 9244-9259.

Cheng L, Li Y, Qi Q, Xu P, Feng R, Palmer L, Chen J, Wu R, Yee T, Zhang J et al. 2021. Single-nucleotide-level mapping of DNA regulatory elements that control fetal hemoglobin expression. *Nat Genet* **53**: 869-880.

Cheng Y, Wu W, Kumar SA, Yu D, Deng W, Tripic T, King DC, Chen KB, Zhang Y, Drautz D et al. 2009. Erythroid GATA1 function revealed by genome-wide analysis of transcription factor occupancy, histone modifications, and mRNA expression. *Genome Res* **19**: 2172-2184.

Conway JR, Lex A, Gehlenborg N. 2017. UpSetR: an R package for the visualization of intersecting sets and their properties. *Bioinformatics* **33**: 2938-2940.

Corces MR, Buenrostro JD, Wu B, Greenside PG, Chan SM, Koenig JL, Snyder MP, Pritchard JK, Kundaje A, Greenleaf WJ et al. 2016. Lineage-specific and single-cell chromatin accessibility charts human hematopoiesis and leukemia evolution. *Nat Genet* **48**: 1193-1203.

Crooks GE, Hon G, Chandonia JM, Brenner SE. 2004. WebLogo: a sequence logo generator. *Genome Res* **14**: 1188-1190.

Ernst J, Kellis M. 2012. ChromHMM: automating chromatin-state discovery and characterization. *Nat Methods* **9**: 215-216.

Finucane HK, Bulik-Sullivan B, Gusev A, Trynka G, Reshef Y, Loh PR, Anttila V, Xu H, Zang C, Farh K et al. 2015. Partitioning heritability by functional annotation using genome-wide association summary statistics. *Nat Genet* **47**: 1228-1235.

Fox S, Myers JA, Davidson C, Getman M, Kingsley PD, Frankiewicz N, Bulger M. 2020. Hyperacetylated chromatin domains mark cell type-specific genes and suggest distinct modes of enhancer function. *Nature communications* **11**: 4544.

Frankish A, Diekhans M, Jungreis I, Lagarde J, Loveland JE, Mudge JM, Sisu C, Wright JC, Armstrong J, Barnes I et al. 2021. Gencode 2021. *Nucleic Acids Res* **49**: D916-D923.

Gasperini M, Hill AJ, McFaline-Figueroa JL, Martin B, Kim S, Zhang MD, Jackson D, Leith A, Schreiber J, Noble WS et al. 2019. A Genome-wide Framework for Mapping Gene Regulation via Cellular Genetic Screens. *Cell* **176**: 377-390 e319.

Ge T, Chen CY, Neale BM, Sabuncu MR, Smoller JW. 2017. Phenome-wide heritability analysis of the UK Biobank. *PLoS Genet* **13**: e1006711.

Gorkin DU, Barozzi I, Zhao Y, Zhang Y, Huang H, Lee AY, Li B, Chiou J, Wildberg A, Ding B et al. 2020. An atlas of dynamic chromatin landscapes in mouse fetal development. *Nature* **583**: 744-751.

Hardison RC. 2012. Genome-wide epigenetic data facilitate understanding of disease susceptibility association studies. *J Biol Chem* **287**: 30932-30940.

Heintzman ND, Stuart RK, Hon G, Fu Y, Ching CW, Hawkins RD, Barrera LO, Van Calcar S, Qu C, Ching KA et al. 2007. Distinct and predictive chromatin signatures of transcriptional promoters and enhancers in the human genome. *Nat Genet* **39**: 311-318.

Heinz S, Benner C, Spann N, Bertolino E, Lin YC, Laslo P, Cheng JX, Murre C, Singh H, Glass CK. 2010. Simple combinations of lineage-determining transcription factors prime cis-regulatory elements required for macrophage and B cell identities. *Mol Cell* **38**: 576-589.

Hindorff LA, Sethupathy P, Junkins HA, Ramos EM, Mehta JP, Collins FS, Manolio TA. 2009. Potential etiologic and functional implications of genome-wide association loci for human diseases and traits. *Proc Natl Acad Sci U S A* **106**: 9362-9367.

Hinrichs AS, Karolchik D, Baertsch R, Barber GP, Bejerano G, Clawson H, Diekhans M, Furey TS, Harte RA, Hsu F et al. 2006. The UCSC Genome Browser Database: update 2006. *Nucleic Acids Res* **34**: D590-598.

Hoffman MM, Buske OJ, Wang J, Weng Z, Bilmes JA, Noble WS. 2012. Unsupervised pattern discovery in human chromatin structure through genomic segmentation. *Nat Methods* **9**: 473-476.

Huang J, Liu X, Li D, Shao Z, Cao H, Zhang Y, Trompouki E, Bowman TV, Zon LI, Yuan GC et al. 2016. Dynamic Control of Enhancer Repertoires Drives Lineage and Stage-Specific Transcription during Hematopoiesis. *Dev Cell* **36**: 9-23.

Huang P, Keller CA, Giardine B, Grevet JD, Davies JOJ, Hughes JR, Kurita R, Nakamura Y, Hardison RC, Blobel GA. 2017. Comparative analysis of three-dimensional chromosomal architecture identifies a novel fetal hemoglobin regulatory element. *Genes Dev* **31**: 1704-1713.

Jansen C, Ramirez RN, El-Ali NC, Gomez-Cabrero D, Tegner J, Merkenschlager M, Conesa A, Mortazavi A. 2019. Building gene regulatory networks from scATAC-seq and scRNA-seq using Linked Self Organizing Maps. *PLoS Comput Biol* **15**: e1006555.

Kakumanu A, Velasco S, Mazzoni E, Mahony S. 2017. Deconvolving sequence features that discriminate between overlapping regulatory annotations. *PLoS Comput Biol* **13**: e1005795.

Khan A, Mathelier A. 2017. Intervene: a tool for intersection and visualization of multiple gene or genomic region sets. *BMC Bioinformatics* **18**: 287.

King DC, Taylor J, Zhang Y, Cheng Y, Lawson HA, Martin J, ENCODE groups for Transcriptional Regulation and Multispecies Sequence Analysis, Chiaromonte F, Miller W, Hardison RC. 2007. Finding cis-regulatory elements using comparative genomics: some lessons from ENCODE data. *Genome Res* **17**: 775-786.

Korsunsky I, Millard N, Fan J, Slowikowski K, Zhang F, Wei K, Baglaenko Y, Brenner M, Loh PR, Raychaudhuri S. 2019. Fast, sensitive and accurate integration of single-cell data with Harmony. *Nat Methods* **16**: 1289-1296.

Kwon SB, Ernst J. 2021. Learning a genome-wide score of human-mouse conservation at the functional genomics level. *Nature communications* **12**: 2495.

Lee DD, Seung HS. 1999. Learning the parts of objects by non-negative matrix factorization. *Nature* **401**: 788-791.

Li J, Moazed D, Gygi SP. 2002. Association of the histone methyltransferase Set2 with RNA polymerase II plays a role in transcription elongation. *J Biol Chem* **277**: 49383-49388.

Libbrecht MW, Chan RCW, Hoffman MM. 2021. Segmentation and genome annotation algorithms for identifying chromatin state and other genomic patterns. *PLoS Comput Biol* **17**: e1009423.

Mahony S, Benos PV. 2007. STAMP: a web tool for exploring DNA-binding motif similarities. *Nucleic Acids Res* **35**: W253-258.

Martens JH, Stunnenberg HG. 2013. BLUEPRINT: mapping human blood cell epigenomes. *Haematologica* **98**: 1487-1489.

Maurano MT, Humbert R, Rynes E, Thurman RE, Haugen E, Wang H, Reynolds AP, Sandstrom R, Qu H, Brody J et al. 2012. Systematic localization of common disease-associated variation in regulatory DNA. *Science* **337**: 1190-1195.

McInnes L, Healy J, Saul N, Grossberger L. 2018. UMAP: Uniform Manifold Approximation and Projection. *Journal of Open Source Software* **3**: 861.

McLean CY, Bristor D, Hiller M, Clarke SL, Schaar BT, Lowe CB, Wenger AM, Bejerano G. 2010. GREAT improves functional interpretation of cis-regulatory regions. *Nat Biotechnol* **28**: 495-501.

Meuleman W, Muratov A, Rynes E, Halow J, Lee K, Bates D, Diegel M, Dunn D, Neri F, Teodosiadis A et al. 2020. Index and biological spectrum of human DNase I hypersensitive sites. *Nature* **584**: 244-251.

Muller J, Hart CM, Francis NJ, Vargas ML, Sengupta A, Wild B, Miller EL, O'Connor MB, Kingston RE, Simon JA. 2002. Histone methyltransferase activity of a Drosophila Polycomb group repressor complex. *Cell* **111**: 197-208.

Murtagh F, Legendre P. 2014. Ward’s Hierarchical Agglomerative Clustering Method: Which Algorithms Implement Ward’s Criterion? *Journal of Classification* **31**: 274-295.

Nassar LR, Barber GP, Benet-Pages A, Casper J, Clawson H, Diekhans M, Fischer C, Gonzalez JN, Hinrichs AS, Lee BT et al. 2023. The UCSC Genome Browser database: 2023 update. *Nucleic Acids Res* **51**: D1188-D1195.

Neph S, Vierstra J, Stergachis AB, Reynolds AP, Haugen E, Vernot B, Thurman RE, John S, Sandstrom R, Johnson AK et al. 2012. An expansive human regulatory lexicon encoded in transcription factor footprints. *Nature* **489**: 83-90.

Padeken J, Methot SP, Gasser SM. 2022. Establishment of H3K9-methylated heterochromatin and its functions in tissue differentiation and maintenance. *Nat Rev Mol Cell Biol* **23**: 623-640.

Pugacheva EM, Rivero-Hinojosa S, Espinoza CA, Mendez-Catala CF, Kang S, Suzuki T, Kosaka-Suzuki N, Robinson S, Nagarajan V, Ye Z et al. 2015. Comparative analyses of CTCF and BORIS occupancies uncover two distinct classes of CTCF binding genomic regions. *Genome Biol* **16**: 161.

Qi Q, Cheng L, Tang X, He Y, Li Y, Yee T, Shrestha D, Feng R, Xu P, Zhou X et al. 2021. Dynamic CTCF binding directly mediates interactions among cis-regulatory elements essential for hematopoiesis. *Blood* **137**: 1327-1339.

Quinlan AR, Hall IM. 2010. BEDTools: a flexible suite of utilities for comparing genomic features. *Bioinformatics* **26**: 841-842.

Rao SS, Huntley MH, Durand NC, Stamenova EK, Bochkov ID, Robinson JT, Sanborn AL, Machol I, Omer AD, Lander ES et al. 2014. A 3D map of the human genome at kilobase resolution reveals principles of chromatin looping. *Cell* **159**: 1665-1680.

Roadmap Epigenomics Consortium, Kundaje A, Meuleman W, Ernst J, Bilenky M, Yen A, Heravi-Moussavi A, Kheradpour P, Zhang Z, Wang J et al. 2015. Integrative analysis of 111 reference human epigenomes. *Nature* **518**: 317-330.

Roh TY, Cuddapah S, Zhao K. 2005. Active chromatin domains are defined by acetylation islands revealed by genome-wide mapping. *Genes Dev* **19**: 542-552.

Sánchez-Castillo M, Ruau D, Wilkinson AC, Ng FS, Hannah R, Diamanti E, Lombard P, Wilson NK, Göttgens B. 2015. CODEX: a next-generation sequencing experiment database for the haematopoietic and embryonic stem cell communities. *Nucleic Acids Res* **43**: D1117-D1123.

Schwartz S, Kent WJ, Smit A, Zhang Z, Baertsch R, Hardison RC, Haussler D, Miller W. 2003. Human-mouse alignments with *Blastz*. *Genome Res* **13**: 103-105.

Schwartz YB, Kahn TG, Nix DA, Li XY, Bourgon R, Biggin M, Pirrotta V. 2006. Genome-wide analysis of Polycomb targets in Drosophila melanogaster. *Nat Genet* **38**: 700-705.

Smith E, Shilatifard A. 2014. Enhancer biology and enhanceropathies. *Nat Struct Mol Biol* **21**: 210-219.

Stunnenberg HG, International Human Epigenome C, Hirst M. 2016. The International Human Epigenome Consortium: A Blueprint for Scientific Collaboration and Discovery. *Cell* **167**: 1145-1149.

Su MY, Steiner LA, Bogardus H, Mishra T, Schulz VP, Hardison RC, Gallagher PG. 2013. Identification of biologically relevant enhancers in human erythroid cells. *J Biol Chem* **288**: 8433-8444.

The ENCODE Project Consortium. 2012. An integrated encyclopedia of DNA elements in the human genome. *Nature* **489**: 57-74.

The ENCODE Project Consortium, Moore JE, Purcaro MJ, Pratt HE, Epstein CB, Shoresh N, Adrian J, Kawli T, Davis CA, Dobin A et al. 2020. Expanded encyclopaedias of DNA elements in the human and mouse genomes. *Nature* **583**: 699-710.

Thurman RE, Rynes E, Humbert R, Vierstra J, Maurano MT, Haugen E, Sheffield NC, Stergachis AB, Wang H, Vernot B et al. 2012. The accessible chromatin landscape of the human genome. *Nature* **489**: 75-82.

van Arensbergen J, FitzPatrick VD, de Haas M, Pagie L, Sluimer J, Bussemaker HJ, van Steensel B. 2017. Genome-wide mapping of autonomous promoter activity in human cells. *Nat Biotechnol* **35**: 145-153.

Vierstra J, Lazar J, Sandstrom R, Halow J, Lee K, Bates D, Diegel M, Dunn D, Neri F, Haugen E et al. 2020. Global reference mapping of human transcription factor footprints. *Nature* **583**: 729-736.

Waksom ML. 2021. seaborn: statistical data visualization. *Journal of Open Source Software* **6**: 3021.

Weirauch MT, Yang A, Albu M, Cote AG, Montenegro-Montero A, Drewe P, Najafabadi HS, Lambert SA, Mann I, Cook K et al. 2014. Determination and inference of eukaryotic transcription factor sequence specificity. *Cell* **158**: 1431-1443.

West AG, Gaszner M, Felsenfeld G. 2002. Insulators: many functions, many mechanisms. *Genes Dev* **16**: 271-288.

Xiang G, Giardine BM, Mahony S, Zhang Y, Hardison RC. 2021. S3V2-IDEAS: a package for normalizing, denoising and integrating epigenomic datasets across different cell types. *Bioinformatics* **37**: 3011-3013.

Xiang G, Keller CA, Giardine B, An L, Li Q, Zhang Y, Hardison RC. 2020a. S3norm: simultaneous normalization of sequencing depth and signal-to-noise ratio in epigenomic data. *Nucleic Acids Res* **48**: e43.

Xiang G, Keller CA, Heuston E, Giardine BM, An L, Wixom AQ, Miller A, Cockburn A, Sauria MEG, Weaver K et al. 2020b. An integrative view of the regulatory and transcriptional landscapes in mouse hematopoiesis. *Genome Res* **30**: 472-484.

Yue F Cheng Y Breschi A Vierstra J Wu W Ryba T Sandstrom R Ma Z Davis C Pope BD et al. 2014. A comparative encyclopedia of DNA elements in the mouse genome. *Nature* **515**: 355-364.

Zhang Y, An L, Yue F, Hardison RC. 2016. Jointly characterizing epigenetic dynamics across multiple human cell types. *Nucleic Acids Res* **44**: 6721-6731.

Zhang Y, Hardison RC. 2017. Accurate and reproducible functional maps in 127 human cell types via 2D genome segmentation. *Nucleic Acids Res* **45**: 9823-9836.

Zhang Y, Liu T, Meyer CA, Eeckhoute J, Johnson DS, Bernstein BE, Nussbaum C, Myers RM, Brown M, Li W et al. 2008. Model-based analysis of ChIP-Seq (MACS). *Genome Biol* **9**: R137.

Zhang Y, Mahony S. 2019. Direct prediction of regulatory elements from partial data without imputation. *PLoS Comput Biol* **15**: e1007399.
