## Supplementary material for "Interspecies regulatory landscapes and elements revealed by novel joint systematic integration of human and mouse blood cell epigenomes": Scripts for analyzing heritability in cCREs: 01_add_annotations.sh

```
#!/usr/bin/bash

#==========#
# Get args #
#==========#

annotListFile=$1 # This should be a text file with two columns: 1) the annotation name and 2) the path to the bed file for that annotation
baselineBedDir=$2 # Path to directory containing baseline LDSC annotations
outDir=$3 #This should contain the path to the output directory
outPrefix=$4 # Prefix of output annotations


#=========================#
# Create output directory #
#=========================#

if [[ ! -d $outDir ]]; then
	mkdir $outDir
fi


#=============================#
# Create new annotation files #
#=============================#

for i in {1..22}; do
	echo $i

	# Set input file
	raw_annot_file=$baselineBedDir/baselineLD.${i}.annot.gz

	# Set output file
	out_file=$outDir/$outPrefix.baselineLD.${i}.annot.gz

	# Set temp files
	tmpPrefix=$(mktemp)
	raw_annot_bed=$tmpPrefix.${i}.annot.bed
	rsid_annots=$tmpPrefix.${i}.rsid.annot


	# Create bed file of variants in raw annotation
	zcat $raw_annot_file | tail -n +2 | cut -f 1-3 | awk -v OFS="\t" '{print "chr"$1, $2-1, $2, $3}' > $raw_annot_bed

	# Create new annotation file
	while read annotName annotBedFile; do

		bedtools map -a $raw_annot_bed -b $annotBedFile -c 3 -o count | cut -f 5 > $rsid_annots.tmp
		echo -e "$annotName\n$(cat $rsid_annots.tmp)" > $rsid_annots.tmp

		if [[ -f $rsid_annots ]]; then
			paste <(cat $rsid_annots) <(cat $rsid_annots.tmp) > $rsid_annots.tmp2
			mv $rsid_annots.tmp2 $rsid_annots
			rm $rsid_annots.tmp
		else
			mv $rsid_annots.tmp $rsid_annots
		fi
	done < $annotListFile

	paste <(zcat $raw_annot_file) <(cat $rsid_annots) | gzip -c > $out_file
	
	# Clean-up
	rm $tmpPrefix*

done
```
