## Supplementary material for "Interspecies regulatory landscapes and elements revealed by novel joint systematic integration of human and mouse blood cell epigenomes": Scripts for analyzing heritability in cCREs: 02_ldsc_regression.sh

```
#!/usr/bin/bash

#==========#
# Get args #
#==========#

chrom=$1 # chromosome to run LDSC regression on. Should be an integer
kgpPlinkDir=$2 # Path to directory containing LDSC 1KGP plink files
hapmapSNPs=$3 # file with hapmap snps
outDir=$4 #This should contain the path to the output directory (same as 01_add_annotations.sh)
outPrefix=$5 # Prefix of output annotations (same as 01_add_annotations.sh)


#==========#
# Run LDSC #
#==========#

# Set chromosome files
annotFile=$outDir/$outPrefix.baselineLD.$chrom.annot.gz
plinkPrefix=$kgpPlinkDir/1000G.EUR.QC.$chrom

# Format snps
printSNPsFile=$(mktemp)
cat $hapmapSNPs | tail -n +2 | cut -f 1 > $printSNPsFile

# Run LDSC
ldsc.py --l2 \
		--bfile $plinkPrefix \
		--ld-wind-cm 1 \
		--annot $annotFile \
		--out $outDir/$outPrefix.baselineLD.$chrom \
		--print-snps $printSNPsFile

# Clean-up
rm $printSNPsFile
```
