## Supplementary material for "Interspecies regulatory landscapes and elements revealed by novel joint systematic integration of human and mouse blood cell epigenomes": Scripts for analyzing heritability in cCREs: 03_partition_heritability.sh

```
#!/usr/bin/bash

#==========#
# Get args #
#==========#

sumstatsFile=$1 # Path to GWAS sumstats file
ldDir=$2 # This should contain the path to the LDSC output directory (same as first two steps)
ldPrefix=$3 # Prefix of LDSC output annotations (same as first two steps)
kgpFrqDir=$4 # Path to directory containing LDSC 1KGP frq files
kgpWeightsDir=$5 # Path to directory containing LDSC 1KGP snp weights
outDir=$6 # Path to output directory
outPrefix=$7 # Prefix of output files


#=========================#
# Create output directory #
#=========================#

if [[ ! -d $outDir ]]; then
	mkdir $outDir
fi


#========================#
# Partition heritability #
#========================#

ldsc.py --h2 $sumstatsFile \
		--ref-ld-chr $ldDir/$ldPrefix.baselineLD. \
		--frqfile-chr $kgpFrqDir/1000G.EUR.QC. \
		--w-ld-chr $kgpWeightsDir/weights.hm3_noMHC. \
		--overlap-annot \
		--print-coefficients \
		--print-delete-vals \
		--out $outDir/$outPrefix
```
