## Supplementary material for "Interspecies regulatory landscapes and elements revealed by novel joint systematic integration of human and mouse blood cell epigenomes": Scripts for analyzing heritability in cCREs: README.md

# Code to run LDSC

This directory contains the scripts used to run LDSC with the VISION cCREs on 587 traits from UKBB.

Here, we will describe how each script is used. The entire pipeline was run twice: first for the full set of cCREs, and a second time for cCRE annotations stratified by joint metacluster.

These scripts assume you have `bedtools` and `ldsc.py` installed and in your PATH.

## Step 0: Get the raw LDSC data

There are several reference files LDSC needs to run, described below. At time of writing, these files were freely available for download from [here](https://alkesgroup.broadinstitute.org/LDSCORE/). They have since been moved to a requester-pays bucket [here](https://console.cloud.google.com/storage/browser/broad-alkesgroup-public-requester-pays/LDSCORE). The files/directories needed for LDSC analysis are as follows (all files are on GRCh37:
1. LDSC baseline annotations for 1000G Phase3 SNPs
    - [1000G_Phase3_baselineLD_v2.2_ldscores.tgz](https://console.cloud.google.com/storage/browser/_details/broad-alkesgroup-public-requester-pays/LDSCORE/1000G_Phase3_baselineLD_v2.2_ldscores.tgz)
    - This directory should be unzipped after download
2.  List of Hapmap3 SNPs to limit LD scores to
    - [w_hm3.snplist.bz2](https://console.cloud.google.com/storage/browser/_details/broad-alkesgroup-public-requester-pays/LDSCORE/w_hm3.snplist.bz2)
    - This file should be unzipped after download
3. LDSC weights for Hapmap3 SNPs
    - [1000G_Phase3_weights_hm3_no_MHC.tgz](https://console.cloud.google.com/storage/browser/_details/broad-alkesgroup-public-requester-pays/LDSCORE/1000G_Phase3_weights_hm3_no_MHC.tgz)
    - This directory should be unzipped after download
4. 1000G Phase 3 plink files
    - [1000G_Phase3_plinkfiles.tgz](https://console.cloud.google.com/storage/browser/_details/broad-alkesgroup-public-requester-pays/LDSCORE/1000G_Phase3_plinkfiles.tgz)
    - This directory should be unzipped after download
5. 1000 Phase 3 allele frequency files
    - [1000G_Phase3_frq.tgz](https://console.cloud.google.com/storage/browser/_details/broad-alkesgroup-public-requester-pays/LDSCORE/1000G_Phase3_frq.tgz)

## Step 1: Add annotations to baseline annotations

The first step is to add the cCRE annotations to the set of baseline annotations. This is accomplished by the `01_add_annotations.sh` script.

This script takes four inputs:
1. A two-column tab-separated file with the name of each annotation to add in the first column, and the path to the corresponding `.bed` file in the second column.
    - For the analysis of the full set of cCREs, this is just a single line file. For the analysis stratified by metacluster, this is a 15 line file: one line for each metacluster annotation
2. The path to the baseline LDSC annotations (the first file downloaded above)
3. The path of the output directory to write annotations to
4. The prefix of the output files

The script will output a new annotation file for each chromosome, with the cCRE annotations added to the baseline annotations

## Step 2: Run LDSC

Now, we compute LD scores, using our new annotations. This is accomplished by the `02_ldsc_regression.sh` script. This script is designed to run for a single chromosome, so that the full process can be parallelized across chromosomes.

The script takes five inputs:
1. The chromosome to run (no "chr", just the number)
2. The path to the directory containing plink files for 1000G SNPs (the fourth file downloaded above)
3. The path to the file containing the Hapmap3 SNPs to output LD scores for (the second file downloaded above)
4. The path to the output directory from Step 1
5. The prefix of the output files from Step 1

This script will output ldscore files to the same directory where the annotation files from Step 1 exist

## Step 3: Partition heritability

Finally, we use the LD scores from step 2 to quantify the extent to which the cCRE annotations are enriched in heritability of relevant traits. This is accomplished by the `03_partition_heritability.sh` script. This script is designed to run for a single trait (i.e. a single sumstats file), so that the full process can be parallelized across traits.

The script takes seven inputs:
1. The path to the sumstats file for the trait of interest
2. The path to the directory containing the ldscore from step 2
3. The prefix of the ldscore files from step 2
4. The path to the directory containing 1000G SNPs allele frequencies (the fifth file downloaded above)
5. The path to the directory containing Hapmap3 LDSC weights (the third file downloaded above)
6. The path to the directory to write the heritability enrichment results to
7. The prefix of the output heritability enrichment results file
