## Supplementary material for "Interspecies regulatory landscapes and elements revealed by novel joint systematic integration of human and mouse blood cell epigenomes": Scripts for analyzing heritability in cCREs: djt_volcano_plot.py

```
#!/usr/bin/env python3

import pandas as pd
import numpy as np
import sys

enrichments = []
enrichment_ps = []
colors = []

full_enrichment_df = pd.read_csv(sys.argv[1], sep='\t', header=0, index_col=0)

for i in range(full_enrichment_df.shape[0]):
    if full_enrichment_df.iloc[i]['Enrichment'] <= 0:
        enrichments.append(0.1)
    else:
        enrichments.append(full_enrichment_df.iloc[i]['Enrichment'])
    enrichment_ps.append(full_enrichment_df.iloc[i]['Enrichment_p'])

previous_row = None
for trait, row in full_enrichment_df.iterrows():
    if row['Enrichment_p_adj'] > 0.05:
        cutoff = previous_row['Enrichment_p']
        break
    previous_row = row

logged_enrichments = np.log2(enrichments)
logged_enrichments_ps = -1 * np.log10(enrichment_ps)
logged_cutoff = -1 * np.log10(cutoff)
print(logged_cutoff, flush=True)

for p in enrichment_ps:
    if p <= cutoff:
        colors.append('red')
    else:
        colors.append('black')

full_enrichment_df['enrichments_list'] = enrichments
full_enrichment_df['enrichment_ps_list'] = enrichment_ps
full_enrichment_df['colors'] = colors
full_enrichment_df['logged_enrichments'] = logged_enrichments
full_enrichment_df['logged_enrichments_ps'] = logged_enrichments_ps

full_enrichment_df.to_csv("withplotting_raw_cCRE_all_UKBB_traits.enrichment.txt", sep="\t", header=True)
```
