## Supplementary material for "Interspecies regulatory landscapes and elements revealed by novel joint systematic integration of human and mouse blood cell epigenomes": Scripts for analyzing heritability in cCREs: metrics.html

enrichment\_bloodtrait\_metrics


### enrichment\_bloodtrait\_metrics

###### Kate Isaac

#### 2022-11-18

```
blood_traits <- read.delim("~/mccoyLab/VISION/sldsc/all_blood_traits.txt", sep = "\t")
ntotal <- nrow(blood_traits) #114

blood_count_traits <- read.delim("~/mccoyLab/VISION/sldsc/blood_count_traits.txt", sep="\t")
ncount <- nrow(blood_count_traits) #54 

nbiochem <- ntotal - ncount #60

volcano_data <- read.delim("~/mccoyLab/VISION/sldsc/withplotting_raw_cCRE_all_UKBB_traits.enrichment.txt", sep="\t")
sig_traits <- volcano_data[which(volcano_data$colors == "red"),]
not_sig_traits <- volcano_data[which(volcano_data$colors != "red"),]
num_sig_traits <- nrow(sig_traits) #53

ntraitsconsidered <- nrow(volcano_data) #587
nnotblood <- ntraitsconsidered - ntotal #473
num_not_sig <- nrow(not_sig_traits) #534
```

```
num_sig_blood_count <- sum(unlist(lapply(1:nrow(sig_traits), function(x) sig_traits$trait_short[x] %in% blood_count_traits$trait_short))) # 50 significant blood count traits
num_not_sig_butare_bloodcount <- sum(unlist(lapply(1:nrow(not_sig_traits), function(x) not_sig_traits$trait_short[x] %in% blood_count_traits$trait_short))) # 4 not significant blood count trait
num_sig_blood <- sum(unlist(lapply(1:nrow(sig_traits), function(x) sig_traits$trait_short[x] %in% blood_traits$trait_short))) # 52 total significant blood traits (count or biochemistry)
# the one that is significant but not listed as a blood trait according to the UKBB is "Pulse wave Arterial Stiffness index" for males
num_not_sig_butare_blood <- sum(unlist(lapply(1:nrow(not_sig_traits), function(x) not_sig_traits$trait_short[x] %in% blood_traits$trait_short))) #62 not significant blood traits
num_sig_bloodchemistry <- num_sig_blood - num_sig_blood_count # 2 significant biochemistry blood traits
num_not_sig_butare_bloodchem <- num_not_sig_butare_blood - num_not_sig_butare_bloodcount # 58 not significant but are blood chemistry
```

We use sensitivity, precision, and specificity pretty liberally in
this script. We did not use the same language within the manuscript
becuase we weren’t thrilled with suggesting there is some truth we’re
comparing to. But the metrics reported in the manuscript within the last
paragraph of the “Enrichment of the cCRE catalog for function-related
elements and trait-associated genetic variants” section, are from this
analysis script.

```
#overall specificity (TN / (FP + TN)) and sensitivity (TP / (TP + FN)) and precision (TP / (TP + FP))
overall_sensitivity <- num_sig_blood / ntotal
overall_precision <- num_sig_blood / num_sig_traits
overall_specificity <- (num_not_sig - num_not_sig_butare_blood) / ((num_sig_traits - num_sig_blood) + (num_not_sig - num_not_sig_butare_blood))
```

```
#blood count
count_sensitivity <- num_sig_blood_count / ncount
count_precision <- num_sig_blood_count / num_sig_traits
count_specificity <- (num_not_sig - num_not_sig_butare_bloodcount) /((num_sig_traits - num_sig_blood_count) + (num_not_sig - num_not_sig_butare_bloodcount))
```

```
#biochemistry
biochem_sensitivity <- num_sig_bloodchemistry / nbiochem
biochem_precision <- num_sig_bloodchemistry / num_sig_traits
biochem_specificity <- (num_not_sig - num_not_sig_butare_bloodchem) /((num_sig_traits - num_sig_bloodchemistry) + (num_not_sig - num_not_sig_butare_bloodchem))
```

```
df <- data.frame("Metric" = c("Precision", "Sensitivity", "Specificity"), "All Blood Traits" = c(overall_precision, overall_sensitivity, overall_specificity), "Blood Count Traits" = c(count_precision, count_sensitivity, count_specificity), "Blood Chemistry Traits" = c(biochem_precision, biochem_sensitivity, biochem_specificity))
df
```

```
##        Metric All.Blood.Traits Blood.Count.Traits Blood.Chemistry.Traits
## 1   Precision        0.9811321          0.9433962             0.03773585
## 2 Sensitivity        0.4561404          0.9259259             0.03333333
## 3 Specificity        0.9978858          0.9943715             0.90322581
```
