## Supplementary material for "Interspecies regulatory landscapes and elements revealed by novel joint systematic integration of human and mouse blood cell epigenomes": Scripts for analyzing heritability in cCREs: metrics.pdf

### enrichment_bloodtrait_metrics

Kate Isaac

2022-11-18

```r
blood_traits <- read.delim("~/mccoyLab/VISION/sldsc/all_blood_traits.txt", sep = "\t")
ntotal <- nrow(blood_traits) #114

blood_count_traits <- read.delim("~/mccoyLab/VISION/sldsc/blood_count_traits.txt", sep="\t")
ncount <- nrow(blood_count_traits) #54

nbiochem <- ntotal - ncount #60

volcano_data <- read.delim("~/mccoyLab/VISION/sldsc/withplotting_raw_cCRE_all_UKBB_traits.enrichment.txt
sig_traits <- volcano_data[which(volcano_data$colors == "red"),]
not_sig_traits <- volcano_data[which(volcano_data$colors != "red"),]
num_sig_traits <- nrow(sig_traits) #53

ntraitsconsidered <- nrow(volcano_data) #587
nnotblood <- ntraitsconsidered - ntotal #473
num_not_sig <- nrow(not_sig_traits) #534
```

```r
num_sig_blood_count <- sum(unlist(lapply(1:nrow(sig_traits), function(x) sig_traits$trait_short[x] %in%
num_not_sig_butare_bloodcount <- sum(unlist(lapply(1:nrow(not_sig_traits), function(x) not_sig_traits$tr
num_sig_blood <- sum(unlist(lapply(1:nrow(sig_traits), function(x) sig_traits$trait_short[x] %in% blood_
# the one that is significant but not listed as a blood trait according to the UKBB is "Pulse wave Arte
num_not_sig_butare_blood <- sum(unlist(lapply(1:nrow(not_sig_traits), function(x) not_sig_traits$trait_s
num_sig_bloodchemistry <- num_sig_blood - num_sig_blood_count # 2 significant biochemistry blood traits
num_not_sig_butare_bloodchem <- num_not_sig_butare_blood - num_not_sig_butare_bloodcount # 58 not signi
```

We use sensitivity, precision, and specificity pretty liberally in this script. We did not use the same language within the manuscript becuase we weren't thrilled with suggesting there is some truth we're comparing to. But the metrics reported in the manuscript within the last paragraph of the "Enrichment of the cCRE catalog for function-related elements and trait-associated genetic variants" section, are from this analysis script.

```r
#overall specificity (TN / (FP + TN)) and sensitivity (TP / (TP + FN)) and precision (TP / (TP + FP))
overall_sensitivity <- num_sig_blood / ntotal
overall_precision <- num_sig_blood / num_sig_traits
overall_specificity <- (num_not_sig - num_not_sig_butare_blood) / ((num_sig_traits - num_sig_blood) + (n
```

```r
#blood count
count_sensitivity <- num_sig_blood_count / ncount
count_precision <- num_sig_blood_count / num_sig_traits
count_specificity <- (num_not_sig - num_not_sig_butare_bloodcount) /((num_sig_traits - num_sig_blood_cou
```

```r
#biochemistry
biochem_sensitivity <- num_sig_bloodchemistry / nbiochem
biochem_precision <- num_sig_bloodchemistry / num_sig_traits
biochem_specificity <- (num_not_sig - num_not_sig_butare_bloodchem) /((num_sig_traits - num_sig_bloodche
```

```r
df <- data.frame("Metric" = c("Precision", "Sensitivity", "Specificity"), "All Blood Traits" = c(overall
df
```

```
##        Metric All.Blood.Traits Blood.Count.Traits Blood.Chemistry.Traits
## 1   Precision        0.9811321          0.9433962             0.03773585
## 2 Sensitivity        0.4561404          0.9259259             0.03333333
## 3 Specificity        0.9978858          0.9943715             0.90322581
```
