## Supplementary material for "Interspecies regulatory landscapes and elements revealed by novel joint systematic integration of human and mouse blood cell epigenomes": Scripts for analyzing heritability in cCREs: plot_fig3f.html

sldsc\_volcano\_fig3f


### sldsc\_volcano\_fig3f

###### Kate Isaac

#### 2022-09-23

```
library(tidyverse)
```

```
## ── Attaching packages ─────────────────────────────────────── tidyverse 1.3.2 ──
## ✔ ggplot2 3.3.6      ✔ purrr   0.3.4 
## ✔ tibble  3.1.8      ✔ dplyr   1.0.10
## ✔ tidyr   1.2.1      ✔ stringr 1.4.1 
## ✔ readr   2.1.2      ✔ forcats 0.5.2 
## ── Conflicts ────────────────────────────────────────── tidyverse_conflicts() ──
## ✖ dplyr::filter() masks stats::filter()
## ✖ dplyr::lag()    masks stats::lag()
```

```
library(ggplot2)
library(ggthemes)
library(ggrepel)
```

```
full_enrichment_df = read.delim("~/mccoyLab/VISION/sldsc/withplotting_raw_cCRE_all_UKBB_traits.enrichment.txt", sep="\t", header=TRUE, row.names=1)

logged_cutoff <- 2.347514159253684
```

```
full_enrichment_df$label <- full_enrichment_df$description
full_enrichment_df <- full_enrichment_df %>%
  mutate(across("label", str_replace, "White blood cell \\(leukocyte\\)", "WBC")) %>%
  mutate(across("label", str_replace, "Red blood cell \\(erythrocyte\\)", "RBC")) %>%
  mutate(across("label", str_replace, "haemoglobin", "Hb")) %>%
  mutate(across("label", str_replace, "Haemoglobin", "Hb")) %>%
  mutate(across("label", str_replace, "Neutrophill", "NEU")) %>%
  mutate(across("label", str_replace, "Platelet", "PLT")) %>%
  mutate(across("label", str_replace, "platelet \\(thrombocyte\\)", "PLT")) %>%
  mutate(across("label", str_replace, "Monocyte", "MON")) %>%
  mutate(across("label", str_replace, 'reticulocyte', "RTIC")) %>%
  mutate(across("label", str_replace, 'Reticulocyte', "RTIC")) %>%
  mutate(across("label", str_replace, "Lymphocyte", "LYM")) %>%
  mutate(across("label", str_replace, "Basophil", "BASO")) %>%
  mutate(across("label", str_replace, "Eosinophill", "EOS")) %>%
  mutate(across("label", str_replace, "Mean corpuscular volume", "MCV")) %>%
  mutate(across("label", str_replace, "Mean", "avg")) %>%
  mutate(across("label", str_replace, "percentage", "%")) %>%
  mutate(across("label", str_replace, "distribution", "dist")) %>%
  mutate(across("label", str_replace, "volume", "vol")) %>%
  mutate(across("label", str_replace, "concentration", "conc")) %>%
  mutate(across("label", str_replace, "count", "ct")) %>%
  mutate(across("label", str_replace, "Immature", "Im")) %>%
  mutate(across("label", str_replace, "Average", "avg")) %>%
  mutate(across("label", str_replace, "Glycated haemoglobin (mmol/mol)", "Glycated Hb"))
```

add \_f and \_m for male and female to labels

```
full_enrichment_df$Sex <- unlist(lapply(1:nrow(full_enrichment_df), function(x) strsplit(row.names(full_enrichment_df)[x], "\\.")[[1]][2]))
full_enrichment_df <- full_enrichment_df %>%
  mutate(across("Sex", str_replace, "female", "f")) %>%
  mutate(across("Sex", str_replace, "male", "m")) %>%
  unite("full_label", c("label", "Sex"), remove = FALSE)
```

Keep certain labels nonsig: Longest period of depression, Vitamin D,
Standing height, Age at death

Keep certain labels sig: Top 14, BASOl %

> colnames(full\_enrichment\_df) [1] “Enrichment” “Enrichment\_p”
> “Enrichment\_p\_adj” “description”  
> [5] “enrichments\_list” “enrichment\_ps\_list” “colors”
> “logged\_enrichments”  
> [9] “logged\_enrichments\_ps” “full\_label” “label” “Sex”

```
notsig <- which(full_enrichment_df$logged_enrichments_ps < logged_cutoff)
#keep1 <- which(full_enrichment_df$label == "Longest period of depression")
keep2 <- which(full_enrichment_df$label == "Vitamin D")
keep3 <- which(full_enrichment_df$label == "Standing height")
keep4 <- which(full_enrichment_df$label == "Age at death")

sigenrichpanel <- which(full_enrichment_df$logged_enrichments_ps >= logged_cutoff & full_enrichment_df$logged_enrichments == min(full_enrichment_df$logged_enrichments))
```

```
sig <- which(full_enrichment_df$logged_enrichments_ps >= logged_cutoff)
keepsig <- 1:17
keepsigpt2 <- which(full_enrichment_df$label == "BASOl %")
keepsigpt3 <- which(full_enrichment_df$label == "EOS %")
keepsigpt4 <- which(full_enrichment_df$label == "PLT dist width")
keepsigpt5 <- which(full_enrichment_df$label == "Glycated Hb")
```

```
full_enrichment_df$label[setdiff(notsig,c(keep2,keep3,keep4))] <- ""
full_enrichment_df$full_label[setdiff(notsig,c(keep2,keep3,keep4))] <- ""
full_enrichment_df$label[sigenrichpanel] <- ""
full_enrichment_df$full_label[sigenrichpanel] <- ""
full_enrichment_df$label[setdiff(sig, c(keepsig, keepsigpt2, keepsigpt3, keepsigpt4, keepsigpt5))] <- ""
full_enrichment_df$full_label[setdiff(sig, c(keepsig, keepsigpt2, keepsigpt3, keepsigpt4, keepsigpt5))] <- ""
```

```
g <- ggplot(full_enrichment_df, aes(x=logged_enrichments, y=logged_enrichments_ps, color=colors, label=label)) + 
  geom_point(aes(shape = Sex), alpha=0.5, size=1) + 
  geom_hline(yintercept = logged_cutoff, color = "gray", linetype = "dashed") +
  geom_vline(xintercept = 0, color = "gray", linetype = "dashed") +
  geom_text_repel(max.overlaps = 25, nudge_y = 0.75) + 
  theme_bw() + theme(panel.background = element_blank(), panel.grid = element_blank()) + 
  scale_color_manual(values = c("black" = "black", "red" = "red"), labels = c("not sig", "sig"), guide="none") +
  geom_text(data=data.frame(), aes(label = '587 traits total', x = -5, y = 14),
            hjust = 0, vjust = 1, color = "black") +
   xlab('Log2(enrichment)') +
   ylab('Log10(p-value)')
g
```

```
## Warning: ggrepel: 1 unlabeled data points (too many overlaps). Consider
## increasing max.overlaps
```

```
ggsave("~/mccoyLab/VISION/sldsc/enrichment_volcano_fmshape.png", plot=g)
```

```
## Saving 7 x 5 in image
```

```
## Warning: ggrepel: 1 unlabeled data points (too many overlaps). Consider
## increasing max.overlaps
```

```
ggsave("~/mccoyLab/VISION/sldsc/enrichment_volcano_fmshape.pdf", plot=g)
```

```
## Saving 7 x 5 in image
```

```
## Warning: ggrepel: 1 unlabeled data points (too many overlaps). Consider
## increasing max.overlaps
```
