## Supplementary material for "Interspecies regulatory landscapes and elements revealed by novel joint systematic integration of human and mouse blood cell epigenomes": Scripts for analyzing heritability in cCREs: plot_fig3f.pdf

### sldsc_volcano_fig3f

Kate Isaac

2022-09-23

```r
library(tidyverse)
```

```
## -- Attaching packages --------------------------------------- tidyverse 1.3.2 --
## v ggplot2 3.3.6      v purrr   0.3.4
## v tibble  3.1.8      v dplyr   1.0.10
## v tidyr   1.2.1      v stringr 1.4.1
## v readr   2.1.2      v forcats 0.5.2
## -- Conflicts ------------------------------------------ tidyverse_conflicts() --
## x dplyr::filter() masks stats::filter()
## x dplyr::lag()    masks stats::lag()
```

```r
library(ggplot2)
library(ggthemes)
library(ggrepel)
```

```r
full_enrichment_df = read.delim("~/mccoyLab/VISION/sldsc/withplotting_raw_cCRE_all_UKBB_traits.enrichme

logged_cutoff <- 2.347514159253684
```

```r
full_enrichment_df$label <- full_enrichment_df$description
full_enrichment_df <- full_enrichment_df %>%
  mutate(across("label", str_replace, "White blood cell \\(leukocyte\\)", "WBC")) %>%
  mutate(across("label", str_replace, "Red blood cell \\(erythrocyte\\)", "RBC")) %>%
  mutate(across("label", str_replace, "haemoglobin", "Hb")) %>%
  mutate(across("label", str_replace, "Haemoglobin", "Hb")) %>%
  mutate(across("label", str_replace, "Neutrophill", "NEU")) %>%
  mutate(across("label", str_replace, "Platelet", "PLT")) %>%
  mutate(across("label", str_replace, "platelet \\(thrombocyte\\)", "PLT")) %>%
  mutate(across("label", str_replace, "Monocyte", "MON")) %>%
  mutate(across("label", str_replace, 'reticulocyte', "RTIC")) %>%
  mutate(across("label", str_replace, 'Reticulocyte', "RTIC")) %>%
  mutate(across("label", str_replace, "Lymphocyte", "LYM")) %>%
  mutate(across("label", str_replace, "Basophil", "BASO")) %>%
  mutate(across("label", str_replace, "Eosinophill", "EOS")) %>%
  mutate(across("label", str_replace, "Mean corpuscular volume", "MCV")) %>%
  mutate(across("label", str_replace, "Mean", "avg")) %>%
  mutate(across("label", str_replace, "percentage", "%")) %>%
  mutate(across("label", str_replace, "distribution", "dist")) %>%
  mutate(across("label", str_replace, "volume", "vol")) %>%
  mutate(across("label", str_replace, "concentration", "conc")) %>%
  mutate(across("label", str_replace, "count", "ct")) %>%
  mutate(across("label", str_replace, "Immature", "Im")) %>%
  mutate(across("label", str_replace, "Average", "avg")) %>%
```

```
## Warning: ggrepel: 2 unlabeled data points (too many overlaps). Consider
## increasing max.overlaps
```

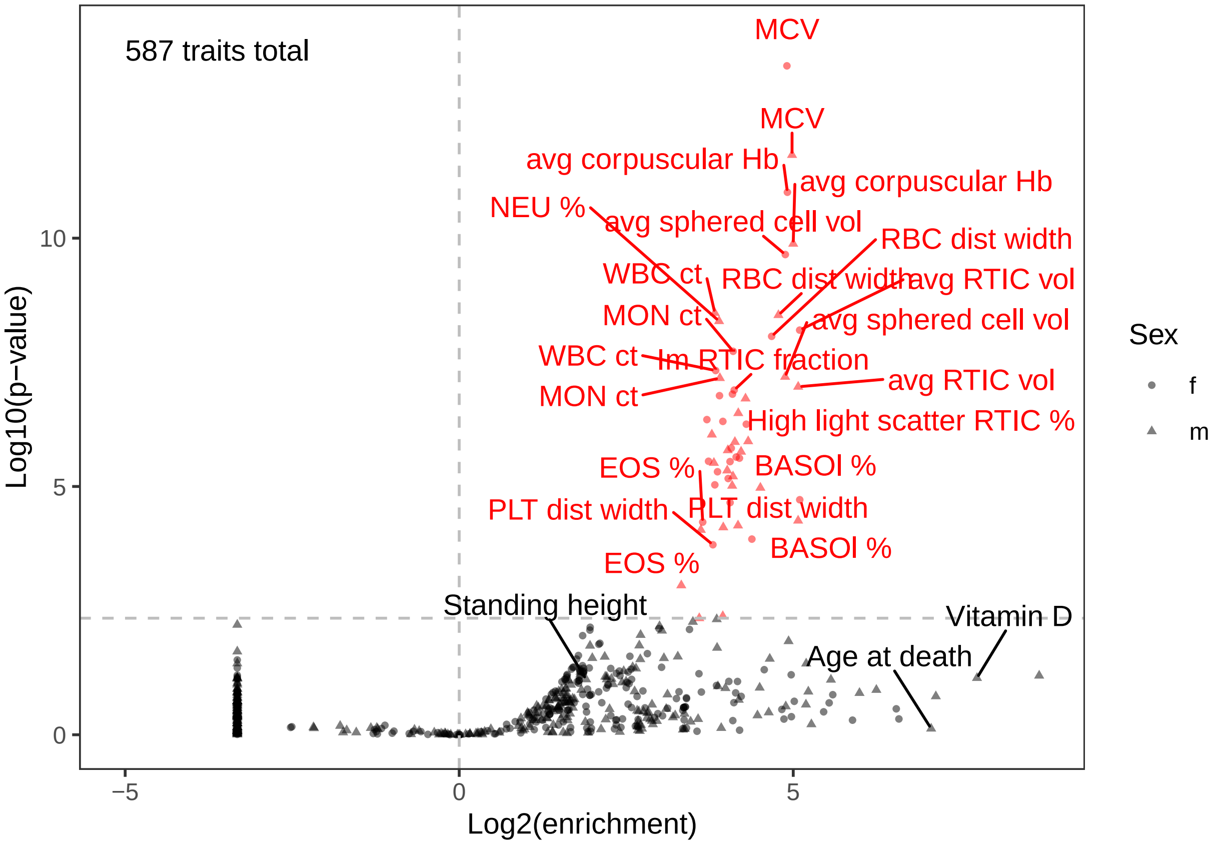


```
ggsave("~/mccoyLab/VISION/sldsc/enrichment_volcano_fmshape.png", plot=g)
```

```
## Saving 6.5 x 4.5 in image
```

```
## Warning: ggrepel: 2 unlabeled data points (too many overlaps). Consider
## increasing max.overlaps
```

```
ggsave("~/mccoyLab/VISION/sldsc/enrichment_volcano_fmshape.pdf", plot=g)
```

```
## Saving 6.5 x 4.5 in image
```

```
## Warning: ggrepel: 2 unlabeled data points (too many overlaps). Consider
## increasing max.overlaps
```
