## Supplementary material for "Interspecies regulatory landscapes and elements revealed by novel joint systematic integration of human and mouse blood cell epigenomes": Scripts for analyzing heritability in cCREs: plot_suppfigS16.html

sldsc\_jmc\_heatmap\_suppfig\_s16


### sldsc\_jmc\_heatmap\_suppfig\_s16

###### Kate Isaac

#### 2022-12-13

Making a heatmap/dotplot hybrid

The desired order of the JMC/metacluster axis

1 4 12 9 8 10 2 5 3 15 7 11 14 6 13 All traits that were found as
significant in at least one metacluster

Enrichment score to be the color of the circle and then we want to
overlay a circle whose size reflects the significance don’t plot a
circle for a trait that is not significant for that given
metacluster

```
library(tidyverse)
```

```
## ── Attaching packages ─────────────────────────────────────── tidyverse 1.3.2 ──
## ✔ ggplot2 3.3.6      ✔ purrr   0.3.4 
## ✔ tibble  3.1.8      ✔ dplyr   1.0.10
## ✔ tidyr   1.2.1      ✔ stringr 1.4.1 
## ✔ readr   2.1.2      ✔ forcats 0.5.2 
## ── Conflicts ────────────────────────────────────────── tidyverse_conflicts() ──
## ✖ dplyr::filter() masks stats::filter()
## ✖ dplyr::lag()    masks stats::lag()
```

```
library(dplyr)
```

read in the metacluster enrichment data read in data that labels the
trait type for all of the traits

```
metacluster_enrichments <- read.delim("~/Downloads/cCRE_15-metaclusters_all_UKBB_traits.enrichment.txt", sep="\t")
trait_type_only <- read.delim("~/mccoyLab/VISION/sldsc/trait_table_description_significance_type.txt", sep="\t") %>% dplyr::select(c(trait_short, type)) %>% `colnames<-`(c("trait", "type"))
```

Add the label for the type of trait to the metacluster enrichment
data using a full join

And set the metacluster number as its own variable, with a factor
level reflecting the order we want it plotted

```
metacluster_enrichments_withtype <- full_join(metacluster_enrichments, trait_type_only, by="trait")
metacluster_enrichments_withtype$metacluster <- factor(unlist(lapply(1:nrow(metacluster_enrichments_withtype), function(x) substr(str_replace(metacluster_enrichments_withtype$Category[x], "metacluster", ""),1,nchar(str_replace(metacluster_enrichments_withtype$Category[x], "metacluster", ""))-4))), levels = rev(c( "1","4","12","9","8","10","2","5","3","15","7","11","14","6","13")))
```

fix our labels to be prettier/match the other volcano figure

```
metacluster_enrichments_withtype$label <- metacluster_enrichments_withtype$description
metacluster_enrichments_withtype <- metacluster_enrichments_withtype %>%
  mutate(across("label", str_replace, "White blood cell \\(leukocyte\\)", "WBC")) %>%
  mutate(across("label", str_replace, "Red blood cell \\(erythrocyte\\)", "RBC")) %>%
  mutate(across("label", str_replace, "haemoglobin", "Hb")) %>%
  mutate(across("label", str_replace, "Haemoglobin", "Hb")) %>%
  mutate(across("label", str_replace, "Neutrophill", "NEU")) %>%
  mutate(across("label", str_replace, "Platelet", "PLT")) %>%
  mutate(across("label", str_replace, "platelet \\(thrombocyte\\)", "PLT")) %>%
  mutate(across("label", str_replace, "Monocyte", "MON")) %>%
  mutate(across("label", str_replace, 'reticulocyte', "RTIC")) %>%
  mutate(across("label", str_replace, 'Reticulocyte', "RTIC")) %>%
  mutate(across("label", str_replace, "Lymphocyte", "LYM")) %>%
  mutate(across("label", str_replace, "Basophil", "BASO")) %>%
  mutate(across("label", str_replace, "Eosinophill", "EOS")) %>%
  mutate(across("label", str_replace, "Mean corpuscular volume", "MCV")) %>%
  mutate(across("label", str_replace, "Mean", "avg")) %>%
  mutate(across("label", str_replace, "percentage", "%")) %>%
  mutate(across("label", str_replace, "distribution", "dist")) %>%
  mutate(across("label", str_replace, "volume", "vol")) %>%
  mutate(across("label", str_replace, "concentration", "conc")) %>%
  mutate(across("label", str_replace, "count", "ct")) %>%
  mutate(across("label", str_replace, "Immature", "Im")) %>%
  mutate(across("label", str_replace, "Average", "avg")) %>%
  mutate(across("label", str_replace, "Glycated haemoglobin (mmol/mol)", "Glycated Hb")) %>%
  mutate(across("label", str_replace, "Forced expiratory vol in 1-second \\(FEV1\\), Best measure", "FEV1")) %>%
  mutate(across("label", str_replace, "\\(left\\)", "\\(l\\)")) %>%
  mutate(across("label", str_replace, "\\(right\\)", "\\(r\\)"))
```

get female only

```
metacluster_enrichments_withtype$sex <- unlist(lapply(1:nrow(metacluster_enrichments_withtype), function(i) unlist(str_split(metacluster_enrichments_withtype$trait[i], "\\."))[2]))

metacluster_enrichments_withtypefem <- metacluster_enrichments_withtype[which(metacluster_enrichments_withtype$sex == "female"),]
```

get male only

```
metacluster_enrichments_withtypemale <- metacluster_enrichments_withtype[which(metacluster_enrichments_withtype$sex == "male"),]
```

#### Focusing on female

Which traits are significant?

```
roi <- which(metacluster_enrichments_withtypefem$Enrichment_p_adj < 0.05)
uniq_sig_traits <- unique(metacluster_enrichments_withtypefem[roi,"trait"])
```

Let’s make some matrices that we can flatten for plotting, but
they’re going to store the size of the point (reflecting significance; 0
if not significant)

and the color of the point (reflecting enrichment)

```
matSize <- matrix(NA, nrow=length(unique(metacluster_enrichments_withtypefem$metacluster)), ncol = length(uniq_sig_traits)) %>% `colnames<-`(uniq_sig_traits) %>% `rownames<-`(names(table(metacluster_enrichments_withtypefem$metacluster)))

matColor <- matrix(NA, nrow=length(unique(metacluster_enrichments_withtypefem$metacluster)), ncol = length(uniq_sig_traits))  %>% `colnames<-`(uniq_sig_traits) %>% `rownames<-`(names(table(metacluster_enrichments_withtypefem$metacluster)))
```

```
for (metacluster_num in names(table(metacluster_enrichments_withtypefem$metacluster))){
  sig_traits <- metacluster_enrichments_withtypefem[which(metacluster_enrichments_withtypefem$metacluster == metacluster_num & metacluster_enrichments_withtypefem$Enrichment_p_adj < 0.05), c("trait", "Enrichment_p_adj", "Enrichment")]
  which_cols_in_mat <- unlist(lapply(1:nrow(sig_traits), function(i) which(colnames(matSize) == sig_traits[i,"trait"])))
  matSize[metacluster_num, which_cols_in_mat] = sig_traits[,"Enrichment_p_adj"]
  matColor[metacluster_num, which_cols_in_mat] = sig_traits[,"Enrichment"]
}
```

#### Focusing on male

Which traits are significant?

```
roi <- which(metacluster_enrichments_withtypemale$Enrichment_p_adj < 0.05)
uniq_sig_traits <- unique(metacluster_enrichments_withtypemale[roi,"trait"])
```

Let’s make some matrices that we can flatten for plotting, but
they’re going to store the size of the point (reflecting significance; 0
if not significant)

and the color of the point (reflecting enrichment)

```
matSizeMale <- matrix(NA, nrow=length(unique(metacluster_enrichments_withtypemale$metacluster)), ncol = length(uniq_sig_traits)) %>% `colnames<-`(uniq_sig_traits) %>% `rownames<-`(names(table(metacluster_enrichments_withtypemale$metacluster)))

matColorMale <- matrix(NA, nrow=length(unique(metacluster_enrichments_withtypemale$metacluster)), ncol = length(uniq_sig_traits))  %>% `colnames<-`(uniq_sig_traits) %>% `rownames<-`(names(table(metacluster_enrichments_withtypemale$metacluster)))
```

```
for (metacluster_num in names(table(metacluster_enrichments_withtypemale$metacluster))){
  sig_traits <- metacluster_enrichments_withtypemale[which(metacluster_enrichments_withtypemale$metacluster == metacluster_num & metacluster_enrichments_withtypemale$Enrichment_p_adj < 0.05), c("trait", "Enrichment_p_adj", "Enrichment")]
  which_cols_in_mat <- unlist(lapply(1:nrow(sig_traits), function(i) which(colnames(matSizeMale) == sig_traits[i,"trait"])))
  matSizeMale[metacluster_num, which_cols_in_mat] = sig_traits[,"Enrichment_p_adj"]
  matColorMale[metacluster_num, which_cols_in_mat] = sig_traits[,"Enrichment"]
}
```

So we need to plot the matrices… we’ll want to collapse the matrices
into a dataframe

```
femMat <- full_join(pivot_longer(cbind(jmc=row.names(matColor), data.frame(matColor)), -c(jmc), names_to = "trait", values_to = "color"), pivot_longer(cbind(jmc=row.names(matSize), data.frame(matSize)), -c(jmc), names_to = "trait", values_to = "size"), by = c("jmc", "trait"))
femMat$trait <- str_replace(femMat$trait, "X", "")
femMatWithLabel <- left_join(femMat, metacluster_enrichments_withtypefem[,c("label", "trait", "type")], by="trait")

maleMat <- full_join(pivot_longer(cbind(jmc=row.names(matColorMale), data.frame(matColorMale)), -c(jmc), names_to = "trait", values_to = "color"), pivot_longer(cbind(jmc=row.names(matSizeMale), data.frame(matSizeMale)), -c(jmc), names_to = "trait", values_to = "size"), by = c("jmc", "trait"))
maleMat$trait <- str_replace(maleMat$trait, "X", "")
maleMatWithLabel <- left_join(maleMat, metacluster_enrichments_withtypemale[,c("label", "trait", "type")], by="trait")
```

set up colors for the tick labels to specify if the trait is a blood
count (maroon/red), blood chemistry (purple), or other (black)
trait.

```
femMatWithLabel <- femMatWithLabel %>%
  mutate(tick_colors = ifelse(type == "count", "maroon", ifelse(type == "chemistry", "purple", "black")))

maleMatWithLabel <- maleMatWithLabel %>%
  mutate(tick_colors = ifelse(type == "count", "maroon", ifelse(type == "chemistry", "purple", "black")))
```

```
tick_colors_fem <- unlist(lapply(1:length(unique(femMatWithLabel$label)), function(x) femMatWithLabel[which(femMatWithLabel$label == unique(femMatWithLabel$label)[order(unique(femMatWithLabel$label))][x])[1], "tick_colors"]))
```

```
gf <- ggplot(femMatWithLabel, aes(y=factor(jmc, levels = rev(c( "1","4","12","9","8","10","2","5","3","15","7","11","14","6","13"))) , x=label)) + geom_tile(aes(fill=factor(0)), show.legend = FALSE) + geom_point(aes(colour = log2(color), size = -log10(size))) + theme_bw() + ylab("JmC") + xlab("") + scale_fill_manual(values = c("transparent")) + theme(axis.text.x = element_text(angle = 90, vjust=0.5,hjust=1, color=tick_colors_fem, size=8)) + labs(size = "-log10(Adjusted p-value)", colour = "log2(Enrichment)") + theme(plot.margin = margin(0.5, 0, 0, 0.25, "cm")) + ggtitle("Female") + theme(panel.grid = element_line(color="black", size=0.1)) + theme(plot.title = element_text(hjust = 0.5))
```

```
## Warning: Vectorized input to `element_text()` is not officially supported.
## Results may be unexpected or may change in future versions of ggplot2.
```

```
gf
```

```
## Warning in FUN(X[[i]], ...): NaNs produced
```

```
## Warning in FUN(X[[i]], ...): NaNs produced
```

```
## Warning: Removed 4875 rows containing missing values (geom_point).
```

```
ggsave("~/mccoyLab/VISION/sldsc/jmc_female_straight.pdf", plot = gf)
```

```
## Saving 7 x 5 in image
```

```
## Warning in FUN(X[[i]], ...): NaNs produced
```

```
## Warning in FUN(X[[i]], ...): NaNs produced
```

```
## Warning: Removed 4875 rows containing missing values (geom_point).
```

```
tick_colors_male <- unlist(lapply(1:length(unique(maleMatWithLabel$label)), function(x) maleMatWithLabel[which(maleMatWithLabel$label == unique(maleMatWithLabel$label)[order(unique(maleMatWithLabel$label))][x])[1], "tick_colors"]))
```

```
gm <- ggplot(maleMatWithLabel, aes(y=factor(jmc, levels = rev(c( "1","4","12","9","8","10","2","5","3","15","7","11","14","6","13"))) , x=label)) + geom_tile(aes(fill=factor(0)), show.legend = FALSE) + geom_point(aes(colour = log2(color), size = -log10(size))) + theme_bw() + ylab("JmC") + xlab("") + scale_fill_manual(values = c("transparent")) + theme(axis.text.x = element_text(angle = 90,vjust=0.5, hjust=1, color=tick_colors_male, size=8)) + labs(size = "-log10(Adjusted p-value)", colour = "log2(Enrichment)") + theme(plot.margin = margin(0.75, 0, 0, 0.25, "cm")) + ggtitle("Male") + theme(panel.grid = element_line(color="black", size=0.1)) + theme(plot.title = element_text(hjust = 0.5))
```

```
## Warning: Vectorized input to `element_text()` is not officially supported.
## Results may be unexpected or may change in future versions of ggplot2.
```

```
gm
```

```
## Warning in FUN(X[[i]], ...): NaNs produced
```

```
## Warning in FUN(X[[i]], ...): NaNs produced
```

```
## Warning: Removed 4515 rows containing missing values (geom_point).
```

```
ggsave("~/mccoyLab/VISION/sldsc/jmc_male_straight.pdf", plot = gm)
```

```
## Saving 7 x 5 in image
```

```
## Warning in FUN(X[[i]], ...): NaNs produced
```

```
## Warning in FUN(X[[i]], ...): NaNs produced
```

```
## Warning: Removed 4515 rows containing missing values (geom_point).
```

```
library(patchwork)

combo <- gf + theme(legend.position = "none") + gm + theme(axis.text.y = element_blank(),
                                                             axis.ticks.y = element_blank(),
                                                             axis.title.y = element_blank() ) + plot_annotation(title = 'JmC Trait Enrichments', theme=theme(plot.title=element_text(hjust=0.5))) + plot_layout(guides = 'collect')
combo
```

```
## Warning in FUN(X[[i]], ...): NaNs produced

## Warning in FUN(X[[i]], ...): NaNs produced
```

```
## Warning: Removed 4875 rows containing missing values (geom_point).
```

```
## Warning in FUN(X[[i]], ...): NaNs produced

## Warning in FUN(X[[i]], ...): NaNs produced
```

```
## Warning: Removed 4515 rows containing missing values (geom_point).
```

```
ggsave("~/mccoyLab/VISION/sldsc/jmc_combo_straight.pdf", plot=combo, width= 12.23, height = 6.15, units = "in")
```

```
## Warning in FUN(X[[i]], ...): NaNs produced
```

```
## Warning in FUN(X[[i]], ...): NaNs produced
```

```
## Warning: Removed 4875 rows containing missing values (geom_point).
```

```
## Warning in FUN(X[[i]], ...): NaNs produced

## Warning in FUN(X[[i]], ...): NaNs produced
```

```
## Warning: Removed 4515 rows containing missing values (geom_point).
```
