## Supplementary material for "Interspecies regulatory landscapes and elements revealed by novel joint systematic integration of human and mouse blood cell epigenomes": Scripts for analyzing heritability in cCREs: plot_suppfigS16.pdf

```r
  mutate(across("label", str_replace, "Lymphocyte", "LYM")) %>%
  mutate(across("label", str_replace, "Basophil", "BASO")) %>%
  mutate(across("label", str_replace, "Eosinophill", "EOS")) %>%
  mutate(across("label", str_replace, "Mean corpuscular volume", "MCV")) %>%
  mutate(across("label", str_replace, "Mean", "avg")) %>%
  mutate(across("label", str_replace, "percentage", "%")) %>%
  mutate(across("label", str_replace, "distribution", "dist")) %>%
  mutate(across("label", str_replace, "volume", "vol")) %>%
  mutate(across("label", str_replace, "concentration", "conc")) %>%
  mutate(across("label", str_replace, "count", "ct")) %>%
  mutate(across("label", str_replace, "Immature", "Im")) %>%
  mutate(across("label", str_replace, "Average", "avg")) %>%
  mutate(across("label", str_replace, "Glycated haemoglobin (mmol/mol)", "Glycated Hb")) %>%
  mutate(across("label", str_replace, "Forced expiratory vol in 1-second \\(FEV1\\), Best measure", "FE
  mutate(across("label", str_replace, "\\(left\\)", "\\(l\\)")) %>%
  mutate(across("label", str_replace, "\\(right\\)", "\\(r\\)"))
```

get female only

```r
metacluster_enrichments_withtype$sex <- unlist(lapply(1:nrow(metacluster_enrichments_withtype), functio

metacluster_enrichments_withtypefem <- metacluster_enrichments_withtype[which(metacluster_enrichments_w
```

get male only

```r
metacluster_enrichments_withtypemale <- metacluster_enrichments_withtype[which(metacluster_enrichments_
```

### Focusing on female

Which traits are significant?

```r
roi <- which(metacluster_enrichments_withtypefem$Enrichment_p_adj < 0.05)
uniq_sig_traits <- unique(metacluster_enrichments_withtypefem[roi,"trait"])
```

Let's make some matrices that we can flatten for plotting, but they're going to store the size of the point (reflecting significance; 0 if not significant)

and the color of the point (reflecting enrichment)

```r
matSize <- matrix(NA, nrow=length(unique(metacluster_enrichments_withtypefem$metacluster)), ncol = leng

matColor <- matrix(NA, nrow=length(unique(metacluster_enrichments_withtypefem$metacluster)), ncol = len
```

```
tick_colors_fem <- unlist(lapply(1:length(unique(femMatWithLabel$label)), function(x) femMatWithLabel[wh
```

```
gf <- ggplot(femMatWithLabel, aes(y=factor(jmc, levels = rev(c( "1","4","12","9","8","10","2","5","3","
```

```
## Warning: Vectorized input to `element_text()` is not officially supported.
## Results may be unexpected or may change in future versions of ggplot2.
```

```
gf
```

```
## Warning in FUN(X[[i]], ...): NaNs produced
```

```
## Warning in FUN(X[[i]], ...): NaNs produced
```

```
## Warning: Removed 4875 rows containing missing values (geom_point).
```

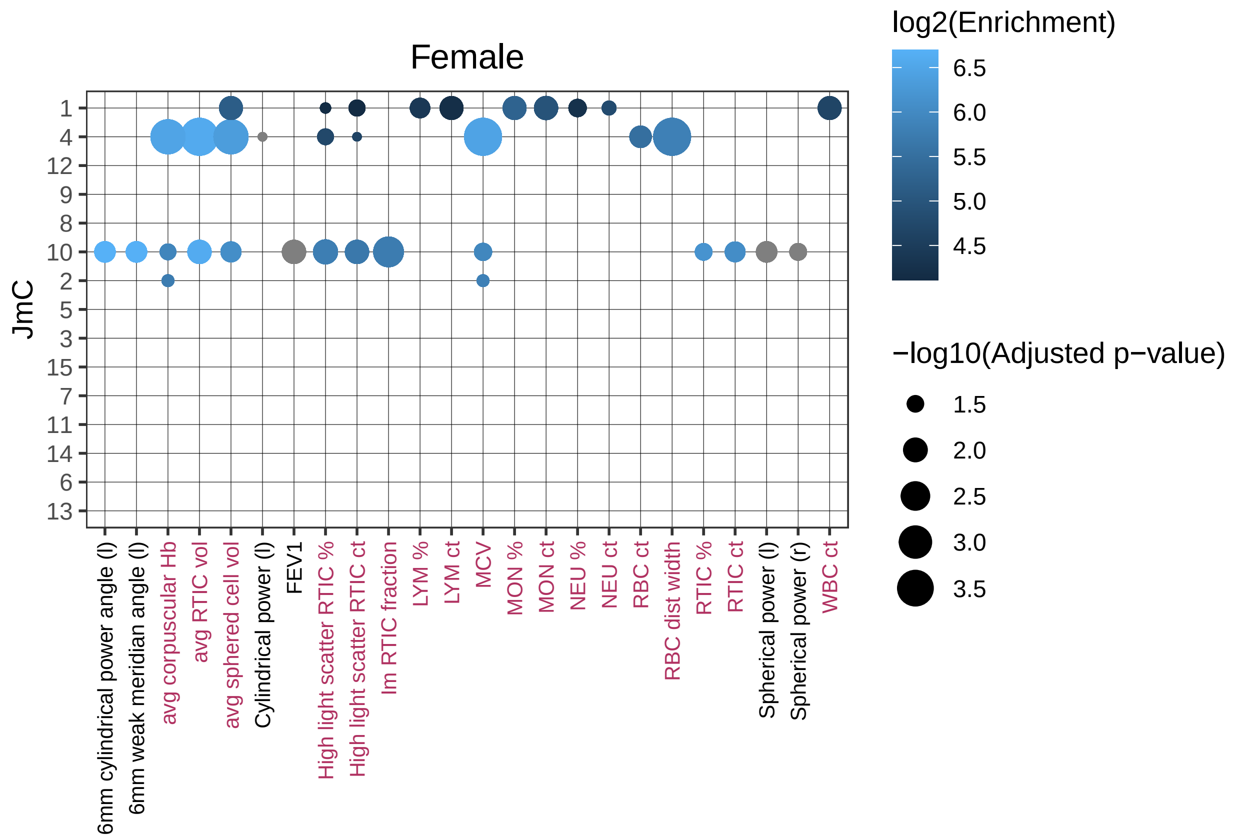


```
ggsave("~/mccoyLab/VISION/sldsc/jmc_female_straight.pdf", plot = gf)
```

```
## Saving 6.5 x 4.5 in image
## Warning in FUN(X[[i]], ...): NaNs produced
## Warning in FUN(X[[i]], ...): NaNs produced
## Warning: Removed 4875 rows containing missing values (geom_point).
```

```
tick_colors_male <- unlist(lapply(1:length(unique(maleMatWithLabel$label)), function(x) maleMatWithLabel
```

```
gm <- ggplot(maleMatWithLabel, aes(y=factor(jmc, levels = rev(c( "1","4","12","9","8","10","2","5","3",
```

```
## Warning: Vectorized input to `element_text()` is not officially supported.
## Results may be unexpected or may change in future versions of ggplot2.
```

```
gm
```

```
## Warning in FUN(X[[i]], ...): NaNs produced
## Warning in FUN(X[[i]], ...): NaNs produced
## Warning: Removed 4515 rows containing missing values (geom_point).
```

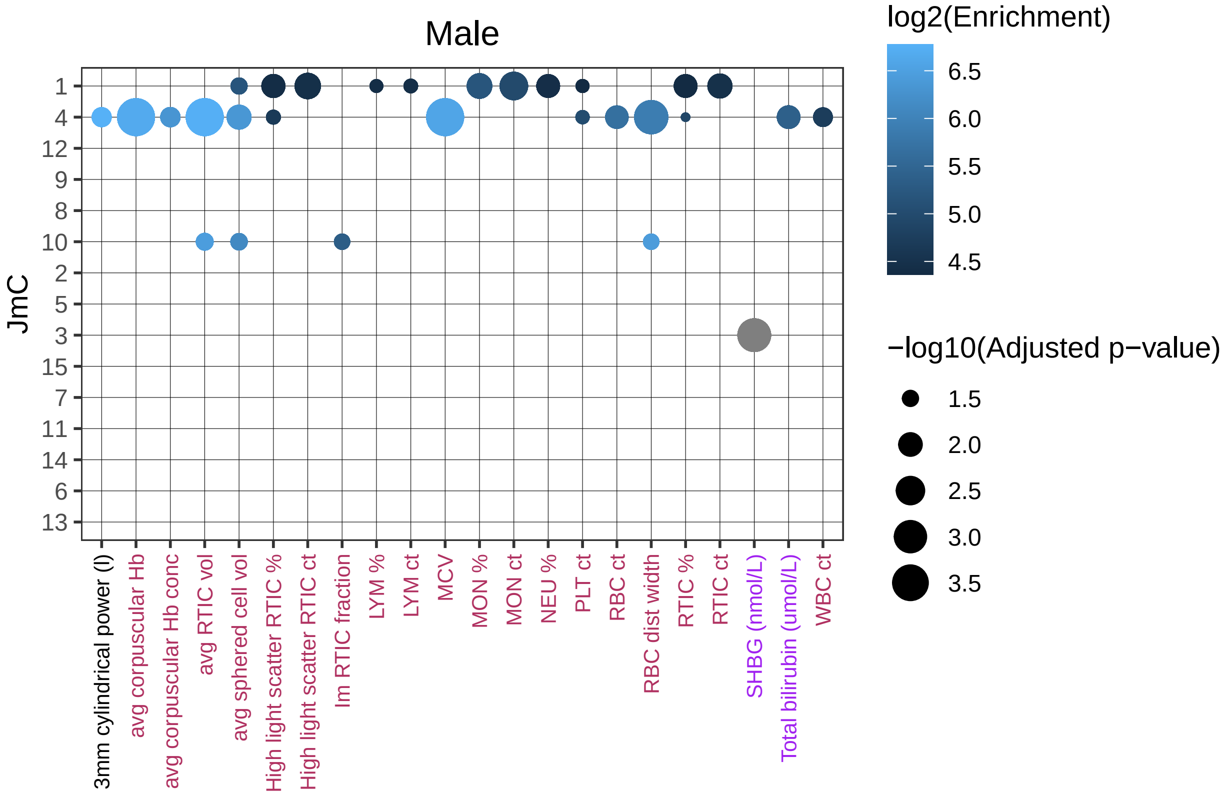


```
ggsave("~/mccoyLab/VISION/sldsc/jmc_male_straight.pdf", plot = gm)
```

```
## Saving 6.5 x 4.5 in image

## Warning in FUN(X[[i]], ...): NaNs produced

## Warning in FUN(X[[i]], ...): NaNs produced

## Warning: Removed 4515 rows containing missing values (geom_point).
```

```
library(patchwork)

combo <- gf + theme(legend.position = "none") + gm + theme(axis.text.y = element_blank(),
                                                          axis.ticks.y = element_blank(),
                                                          axis.title.y = element_blank() ) + plot_an
combo
```

```
## Warning in FUN(X[[i]], ...): NaNs produced

## Warning in FUN(X[[i]], ...): NaNs produced

## Warning: Removed 4875 rows containing missing values (geom_point).

## Warning in FUN(X[[i]], ...): NaNs produced

## Warning in FUN(X[[i]], ...): NaNs produced

## Warning: Removed 4515 rows containing missing values (geom_point).
```

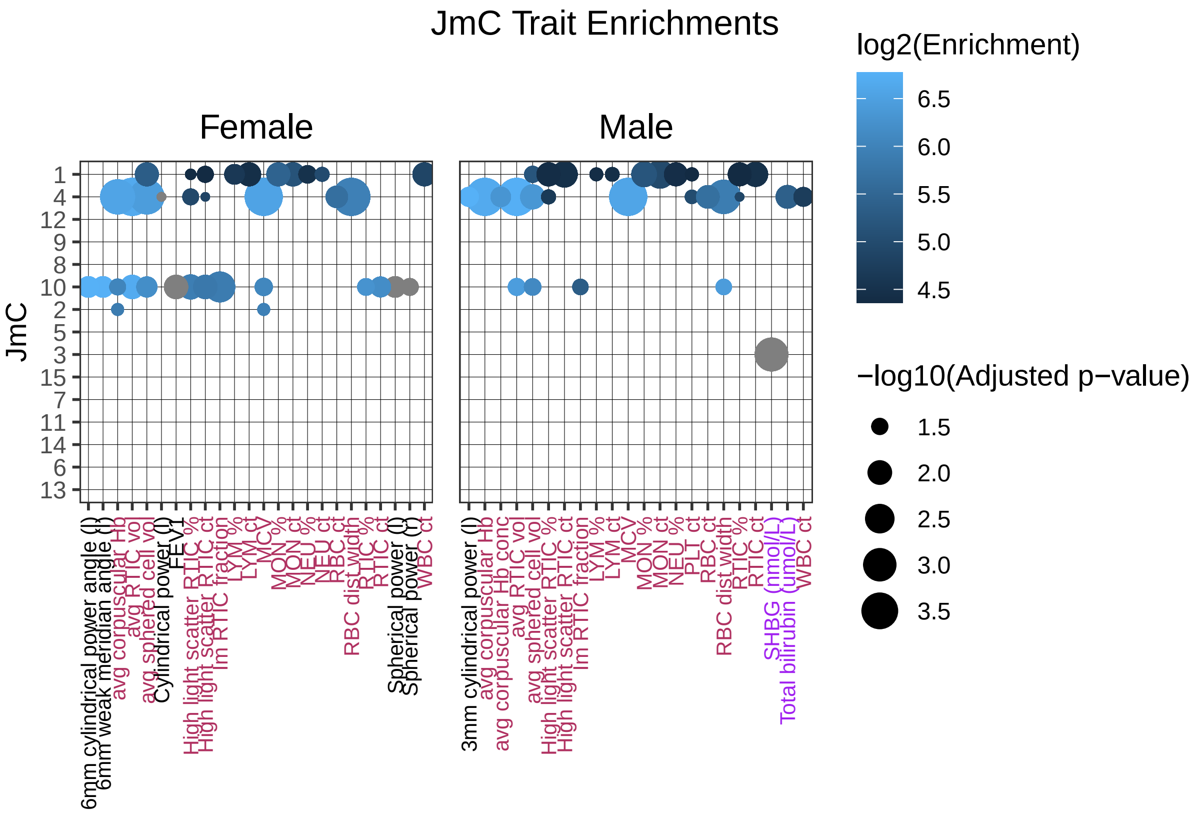


```
ggsave("~/mccoyLab/VISION/sldsc/jmc_combo_straight.pdf", plot=combo, width= 12.23, height = 6.15, units

## Warning in FUN(X[[i]], ...): NaNs produced

## Warning in FUN(X[[i]], ...): NaNs produced

## Warning: Removed 4875 rows containing missing values (geom_point).

## Warning in FUN(X[[i]], ...): NaNs produced

## Warning in FUN(X[[i]], ...): NaNs produced

## Warning: Removed 4515 rows containing missing values (geom_point).
```
