## Supplementary material for "Interspecies regulatory landscapes and elements revealed by novel joint systematic integration of human and mouse blood cell epigenomes": Scripts for analyzing heritability in cCREs: README.md

The files in this repository are used to produce 2 figures ([Figure 3F](../../plots/enrichment_volcano_fmshape.pdf) and [Supplemental Figure S16](../../plots/jmc_combo_straight.pdf)) and some metrics reported in the main text. Each RMarkdown file has a corresponding HTML and PDF file for easier viewing.

Input files to each script are provided in the [data_files directory](../../data_files). The data files themselves are referred to within the script using actual paths that will need to be edited if you intend to run these scripts yourselves to produce the various outputs. 

## Making the [Volcano plot, Figure 3F](../../plots/enrichment_volcano_fmshape.pdf)

  * Step 1: Use [djt_volcano_plot.py](djt_volcano_plot.py)
    * Input to this script is [the output of the LDSC analysis](../../data_files/raw_cCRE_all_UKBB_traits.enrichment.txt)
    * Output of this script is
      * the logged cutoff for significant enrichment (printed line 30)
      * a [further annotation of the input file that provides plotting metrics/values](../../data_files/withplotting_raw_cCRE_all_UKBB_traits.enrichment.txt) -- while this script itself can be used to produce a lovely volcano plot, we wanted a little more control in the aesthetics of the plot and turned to ggplot, but used this original plotting script to provide the values that would be plotted in R.
  * Step 2: Use [plot_fig3f.Rmd](plot_fig3f.Rmd) to produce the volcano plot
    * Input to this script is
      * [the further annotation produced by djt_volcano_plot.py](../../data_files/withplotting_raw_cCRE_all_UKBB_traits.enrichment.txt), specified on line 22
      * the logged cutoff for significant enrichment printed from the first step (specified on line 24)
    * Output of this script is the [Volcano plot, Figure 3F](../../plots/enrichment_volcano_fmshape.pdf) showing the enrichment of heritability of SNPs associated with various traits.

## Finding the metrics reported in the main text associated with the Volcano plot and trait identity

  * Use [metrics.Rmd](metrics.Rmd)
    * Input to this script uses the annotated LDSC output as well as trait annotation which is based on [information provided by the UK Biobank](https://biobank.ndph.ox.ac.uk/ukb/label.cgi?id=100080) on what it considers to be blood traits (specifically blood count and blood biochemistry). A [summary of this](../../data_files/blood_traits_annot.txt) is provided in the data files.
      *  Specifically the [annotated LDSC output produced by djt_volcano_plot.py](../../data_files/withplotting_raw_cCRE_all_UKBB_traits.enrichment.txt), read in on line 19
      *  a list of [all traits considered to be blood traits](../../data_files/all_blood_traits.txt), read in on line 11
      *  a list of [blood traits which are specifically blood count traits](../../data_files/blood_count_traits.txt), read in on line 14
    *  This script provides a lot of metrics which are not reported in the text, but every count or percentage reported in the main text about the identity of the significantly enriched traits can be found using this script.
   
## Making the [heatmap, dotplot hybrid, Supplemental Figure S16](../../plots/jmc_combo_straight.pdf)

  * Use [plot_suppfigS16.Rmd](plot_suppfigS16.Rmd)
    *  Input to this script uses an aggregated output of running LDSC for each individual metacluster/JmC of CREs as its own annotation, and then a file which labels all the traits with the type of trait it is (blood count, blood biochemistry, non-blood)
      * The [aggregated output of running LDSC for each JmC](../../data_files/cCRE_15-metaclusters_all_UKBB_traits.enrichment.txt) is read in on line 48
      * The [file with the trait type annotations](../../data_files/trait_table_description_significance_type.txt) is read in on line 49, selecting only the trait names (`trait_short`) and the `type` columns
    *  The output of this script is the [heatmap, dotplot hybrid for Supplemental Figure S16](../../plots/jmc_combo_straight.pdf) which shows the traits associated with significant enrichment of the heritability of SNPs for each metacluster.    
