## Supplementary material for "Interspecies regulatory landscapes and elements revealed by novel joint systematic integration of human and mouse blood cell epigenomes": Scripts for analyzing heritability in cCREs: 01_download_sumstats.sh

```
#!/usr/bin/bash

#==========#
# Get args #
#==========#

traitDataFile=$1 # Path to traitData.txt file
outDir=$2 # Directory to download sumstats files into


#=========================#
# Create output directory #
#=========================#

if [[ ! -d $outDir ]]; then
	mkdir $outDir
fi


#=========================#
# Download sumstats files #
#=========================#

{
read
while IFS=$'\t' read -r traitID traitDescription traitType downloadLink; do
	wget $downloadLink -P $outDir
done
} < $traitDataFile
```
