## Supplementary material for "Interspecies regulatory landscapes and elements revealed by novel joint systematic integration of human and mouse blood cell epigenomes": Scripts for analyzing heritability in cCREs: 02_download_format_annotations.sh

```
#!/usr/bin/bash

#==========#
# Get args #
#==========#

outDir=$1 # Directory to write files to

annotDir=https://usevision.org/data/ccre/Joint_metaclusters
annotFile=S3V2_IDEAS_hg38_ccre2.cCRE.M.notall0.rmallNEU.withid.clusterID.JclusterID.bed


#==========================#
# Download full annotation #
#==========================#

outFile=$outDir/S3V2_IDEAS_hg38_ccre2.cCRE.M.notall0.rmallNEU.all.bed

wget -P $outDir $annotDir/$annotFile
cat $outDir/$annotFile | tail -n +2 | cut -f 1-3 > $outFile


#==================#
# Extract clusters #
#==================#

for clusterNum in {1..15}; do
	outFile=$outDir/S3V2_IDEAS_hg38_ccre2.cCRE.M.notall0.rmallNEU.JmC${clusterNum}.bed
	cat $outDir/$annotFile | tail -n +2 | awk -v clusterNum=$clusterNum -v OFS="\t" '$4 == clusterNum {print $1, $2, $3}' > $outFile
done


#==========#
# Clean up #
#==========#

rm $outDir/$annotFile
```
