## Supplementary material for "Interspecies regulatory landscapes and elements revealed by novel joint systematic integration of human and mouse blood cell epigenomes": Scripts for analyzing heritability in cCREs: 03_liftover_annotations.sh

```
#!/usr/bin/bash

#==========#
# Get args #
#==========#

annotBedFile=$1 # Input annotation on hg38
outLiftedBedFile=$2 # File to write lifted annotations to (hg19). Should not be gzipped
outFailedFile=$3 # File to write annotations that failed liftover to
chainFile=$4 # Path to chain file (hg38 to hg19)


#======================#
# Liftover annotations #
#======================#

liftOver -minMatch=1 \
		 $annotBedFile \
		 $chainFile \
		 $outLiftedBedFile.unsorted.tmp \
		 $outFailedBedFile


#==================#
# Sort annotations #
#==================#

cat $outLiftedBedFile.unsorted.tmp | sort -t $'\t' -k1,1V -k2,2g -k3,3g > $outLiftedBedFile

# Clean-up
rm $outLiftedBedFile.unsorted.tmp
```
