## Supplementary material for "Interspecies regulatory landscapes and elements revealed by novel joint systematic integration of human and mouse blood cell epigenomes": Scripts for analyzing heritability in cCREs: README.md

# Prepare data for LDSC analysis

This directory contains code to download and format cCRE annotations and UKBB sumstats file for use with LDSC.

The files/scripts in this directory are described here:

## Download UKBB sumstats files

GWAS results (as summary statistics) are available from the [Neale lab](https://www.nealelab.is/uk-biobank).

The 587 traits chosen for this analysis are described in the tab-separated [`traitData.txt`](traitData.txt) file. This file has four fields:
1. The ID of the trait (includes if results are for male or female samples)
2. The long-form description of the trait
3. Whether the trait is a "blood count" trait, a "blood biochemistry" trait, or "non-blood" related
4. The download link to the GWAS summary statistics file

The `01_download_sumstats.sh` script is used to download the summary statistics files for all 587 traits. This script takes two inputs:
1. The path to the `traitData.txt` file
2. The path to the directory to download the sumstats files to

## Download the cCRE annotations

Now, we need to download the human cCRE annotations, and to stratify the full set of cCREs by their joint metacluster. The `02_download_format_annotations.sh` script does both tasks. This script takes just a single input:
1. The path to the directory to write the files to

This script will create a `S3V2_IDEAS_hg38_ccre2.cCRE.M.notall0.rmallNEU.all.bed` file with all cCRE annotations, and 15 `S3V2_IDEAS_hg38_ccre2.cCRE.M.notall0.rmallNEU.<clusterName>.bed` files, one for each joint metacluster

## Liftover the cCRE annotations from hg38 (GRCh38) to hg19 (GRCh37)

While the cCREs are reported on the hg38 reference, the LDSC reference files all use the hg19 reference. As such, we need to liftover the cCRE annotations to hg19. The `03_liftover_annotations.sh` script accomplishes this.

This script assumes you have the UCSC [liftOver](https://genome-store.ucsc.edu/) tool installed and in your PATH. It also requires you to have an hg38 --> hg19 chain file, which can be downloaded from [here](https://hgdownload.soe.ucsc.edu/goldenPath/hg38/liftOver/hg38ToHg19.over.chain.gz).

This script takes four inputs:
1. The path to the cCRE bed file to liftover
    - This script will need to be run 16 times total: once for the full cCRE annotation, and once each for the 15 metacluster annotations
2. The path to the output lifted bed file
3. The path to output file of annotations that fail liftover
4. The path to the hg38 to hg19 chain file

These lifted annotations are used in the LDSC analysis
