## Supplementary material for "Interspecies regulatory landscapes and elements revealed by novel joint systematic integration of human and mouse blood cell epigenomes": Scripts for analyzing heritability in cCREs: README.md

# Code for VISION cCRE heritability enrichment analyses

## Download and prepare data

The [`prep_data`](prep_data) directory contains code to download and prepare the data needed to run the LDSC heritability enrichment analyses

## Run LDSC

The [`ldsc_pipeline`](ldsc_pipeline) directory contains the scripts used to run LDSC

## Plot figures

The [`plot_figures`](plot_figures) directory contains code to plot the figures used in the text
