## Supplementary material for "Interspecies regulatory landscapes and elements revealed by novel joint systematic integration of human and mouse blood cell epigenomes": Scripts for analyzing heritability in cCREs: README.md

# VISION cCRE Trait Heritability Enrichment

This directory contains code and data to replicate the trait heritability analyses done for **Interspecies regulatory landscapes and elements revealed by novel joint systematic integration of human and mouse blood cell epigenomes**. This manuscript was previously posted to BioRxiv as [Cross-species regulatory landscapes and elements revealed by novel joint systematic integration of human and mouse blood cell epigenomes](https://www.biorxiv.org/content/10.1101/2023.04.02.535219v1).
Specifically this directory relates to the following sections and figures of the revised manuscript: 
  * "Enrichment of the cCRE catalog for function-related elements and trait-associated genetic variants" in the main text
    * [Figure 3F](/plots/enrichment_volcano_fmshape.pdf)
  * "Stratified linkage disequilibrium score regression (sLDSC)" subsection of the "Enrichment of genetic variants for blood cell related traits in the VISION human cCRE collection" section in the supplemental material
  * "Enrichment of trait-associated SNPs in the joint metaclusters (JmCs)" in the supplemental material
    * [Supplemental Figure S16](/plots/jmc_combo_straight.pdf)


Specifically, we used [linkage disequilibrium score regression (LDSC)](https://github.com/bulik/ldsc) to quantify whether VISION human blood cell candidate cis-regulatory elements (cCREs) were enriched for heritability of 587 traits from [UKBB](https://www.nealelab.is/uk-biobank).

## Code

The [`code`](/code) directory contains code to download and prepare the data for the analyses, run LDSC, and create the plots used in the paper

## Data

The [`data_files`](/data_files) directory contains results from LDSC as well as supplementary annotation files, used to create the plots found in the paper.

## Plots

The [`plots`](/plots) directory contains the exact plots produced by the scripts in the [`code/plot_figures`](/code/plot_figures) directory.
