## Supplementary figures and images for "Interspecies regulatory landscapes and elements revealed by novel joint systematic integration of human and mouse blood cell epigenomes"

### 01a2_S3V2_IDEAS_hg38_r1_rmh_rerun2.para.modified.para.pdf

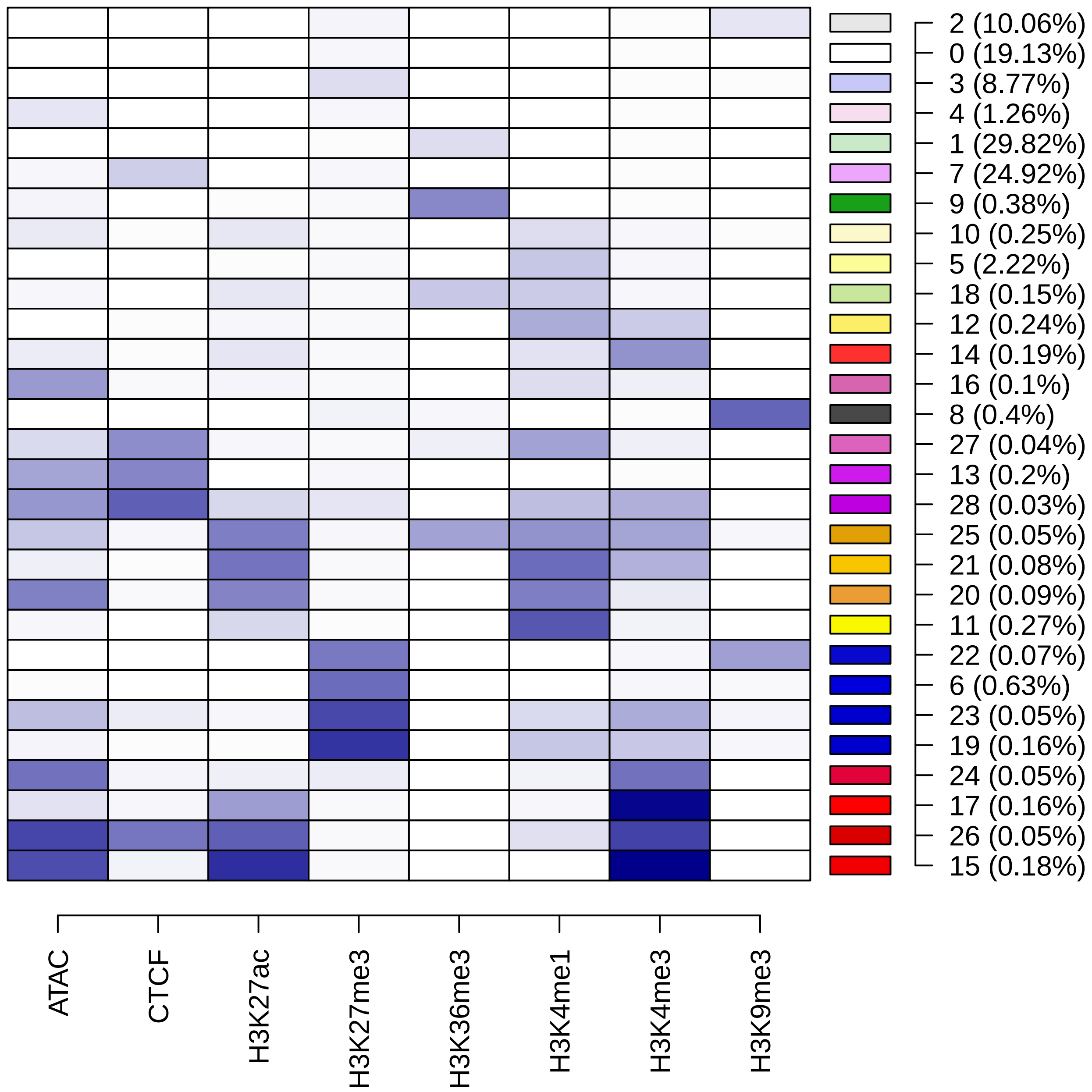

### 01a_S3V2_IDEAS_hg38_r1.all.m_v.pdf

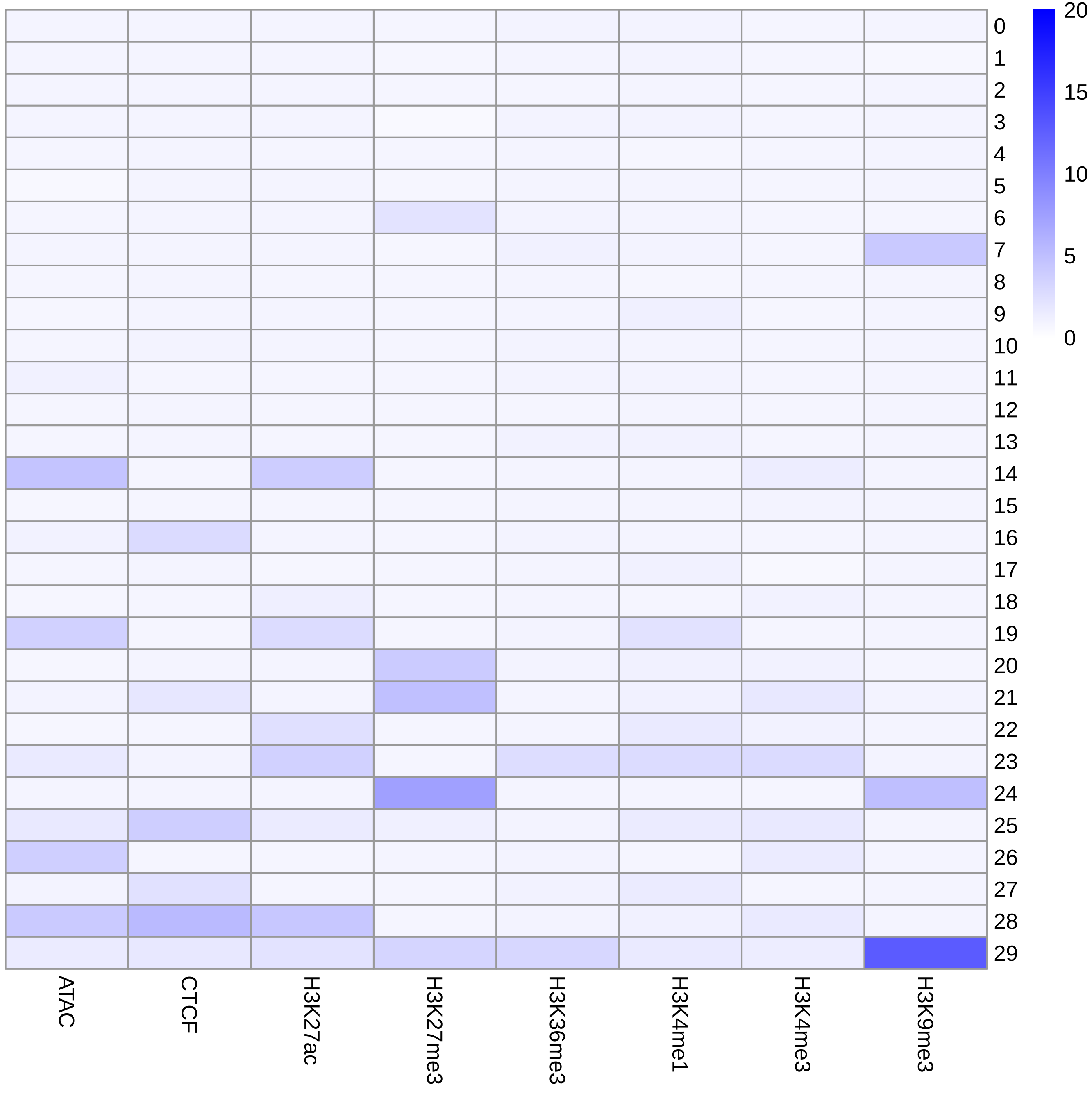

### 01a_S3V2_IDEAS_hg38_r1.pdf

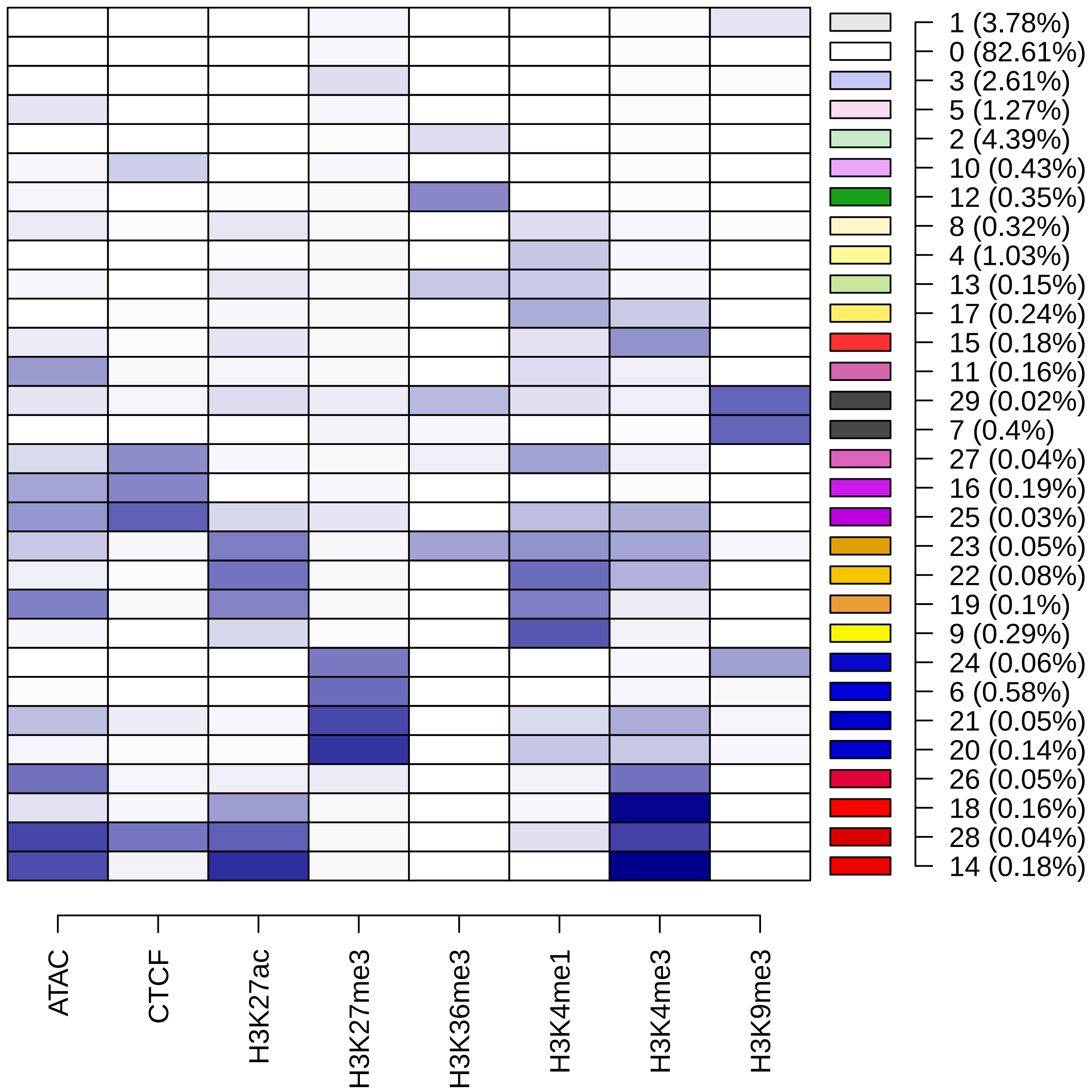

### 01b_S3V2_IDEAS_mm10_r1.pdf

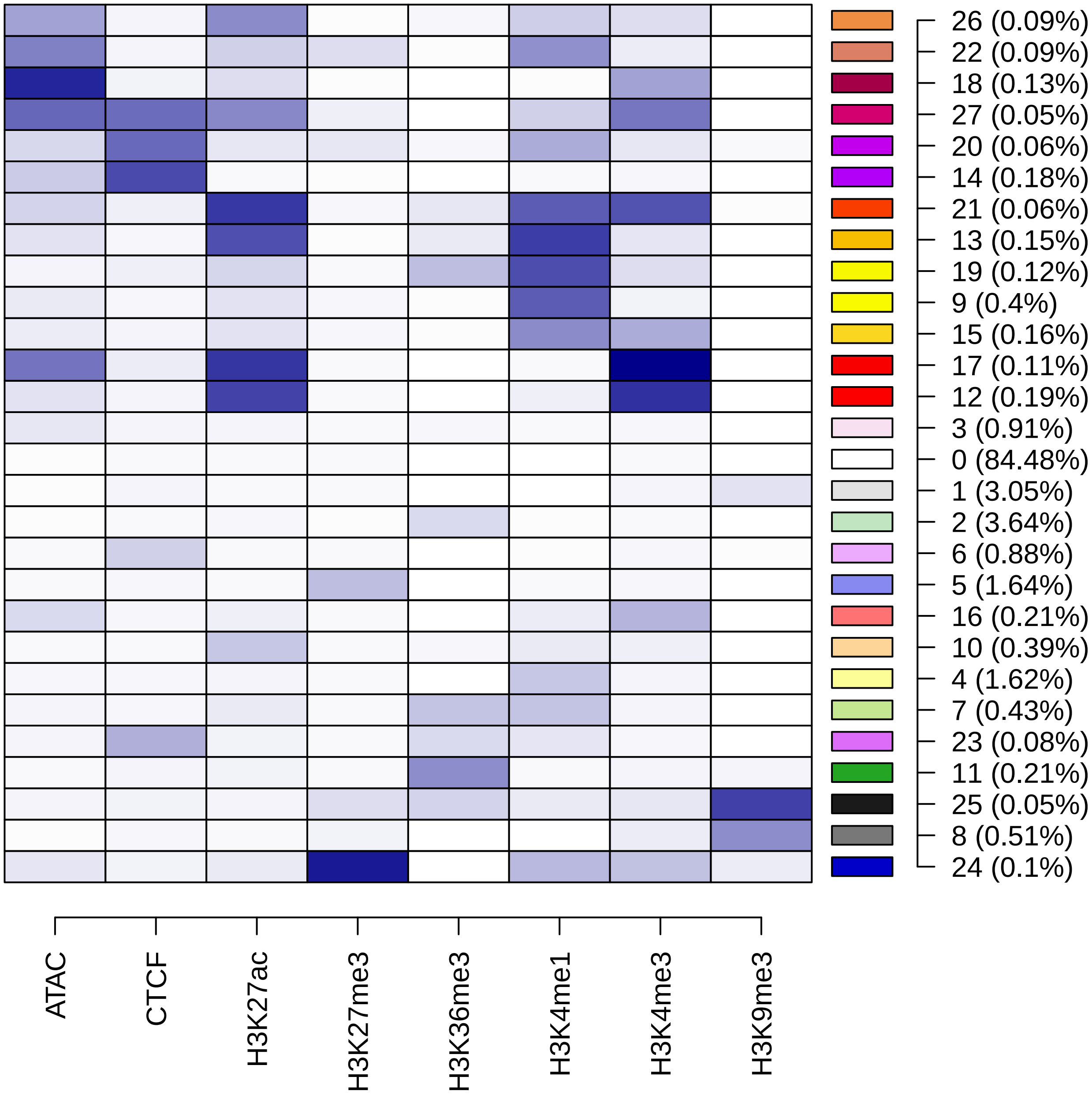

### 02_hg38_mm10_joint_states_8mk.52.para.pdf

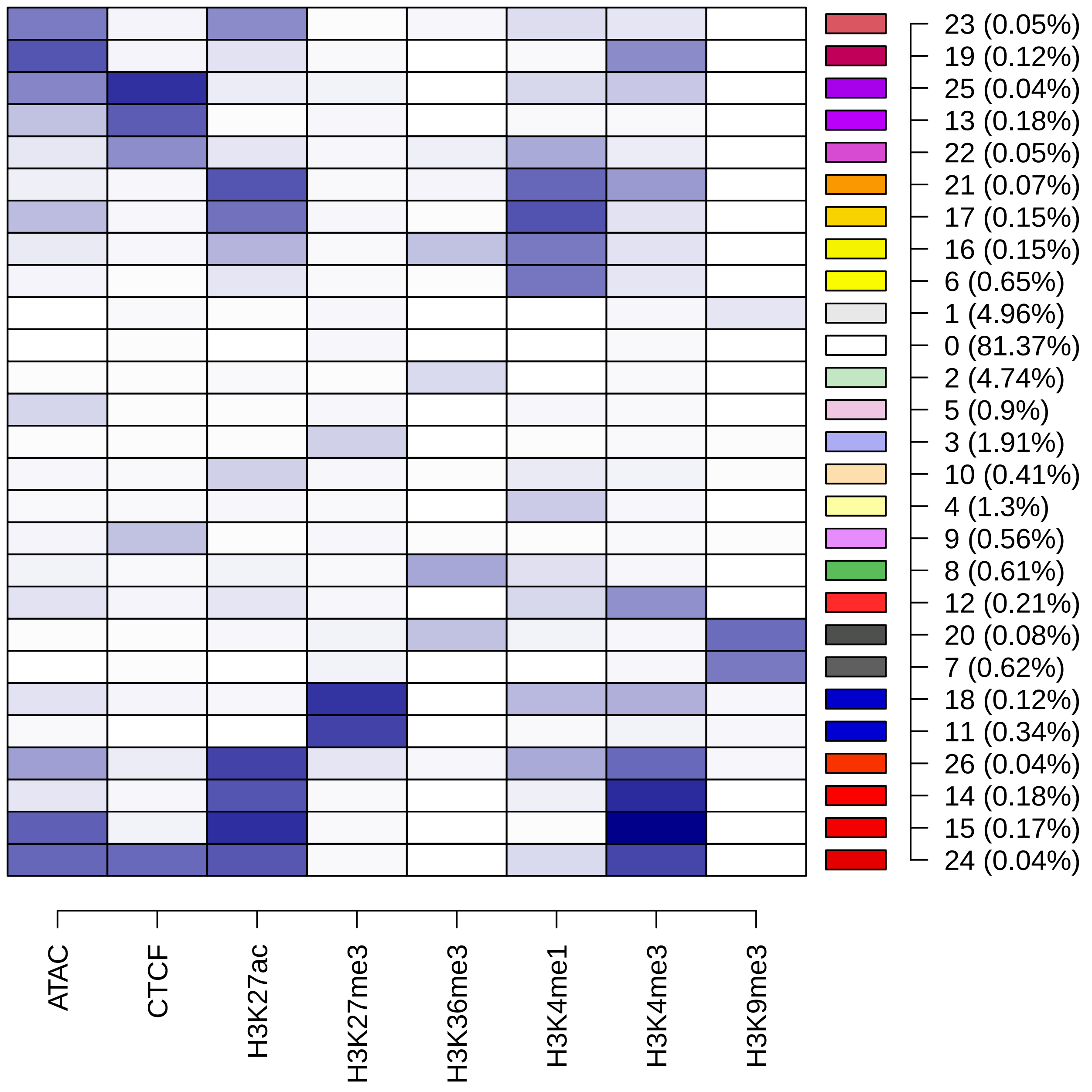

### 02_hg38_mm10_joint_states_IS.52.1.para.pdf

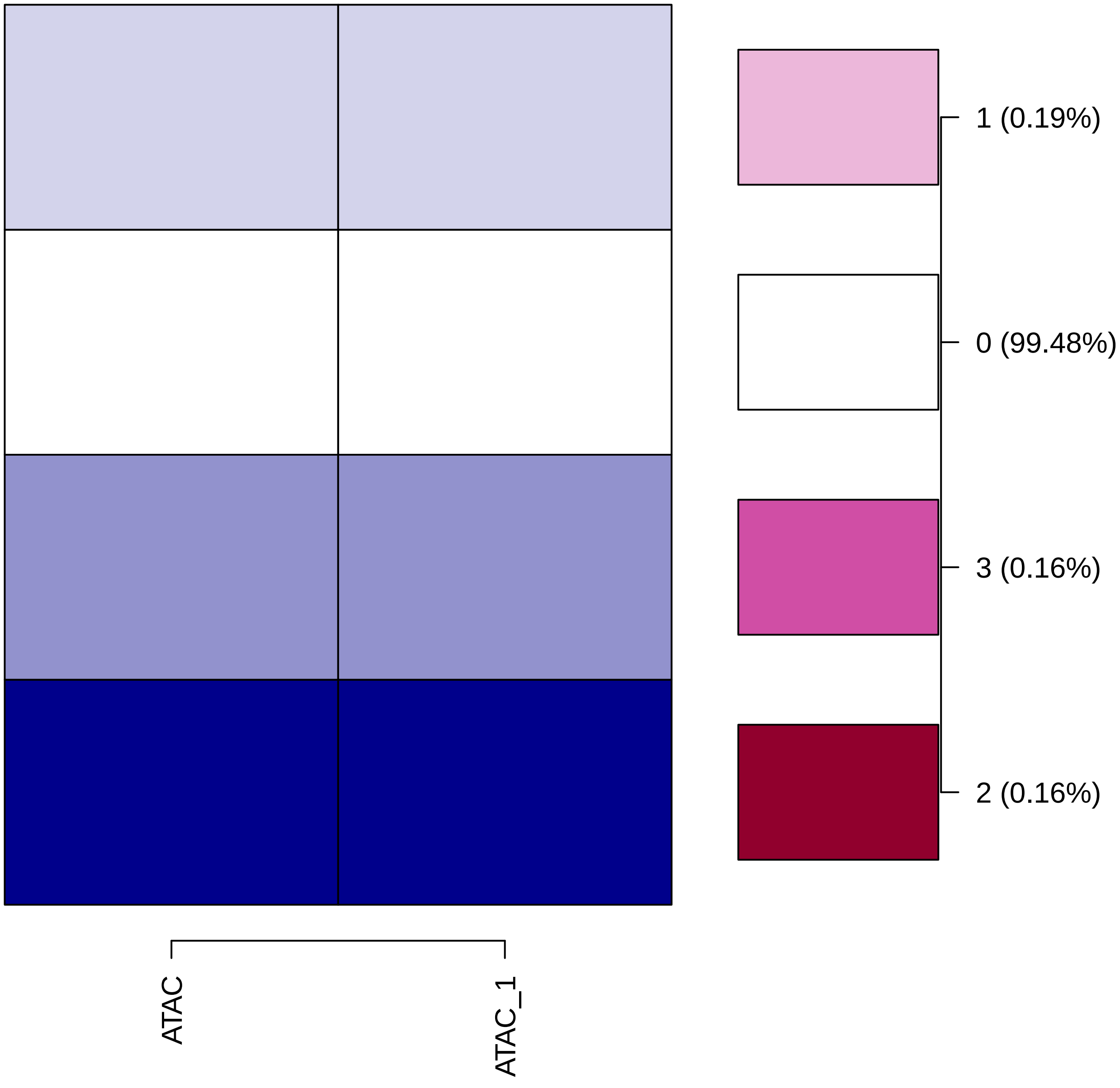

### 03_S3V2_IDEAS_hg38_r2_with_prior_J.all.m_v.pdf

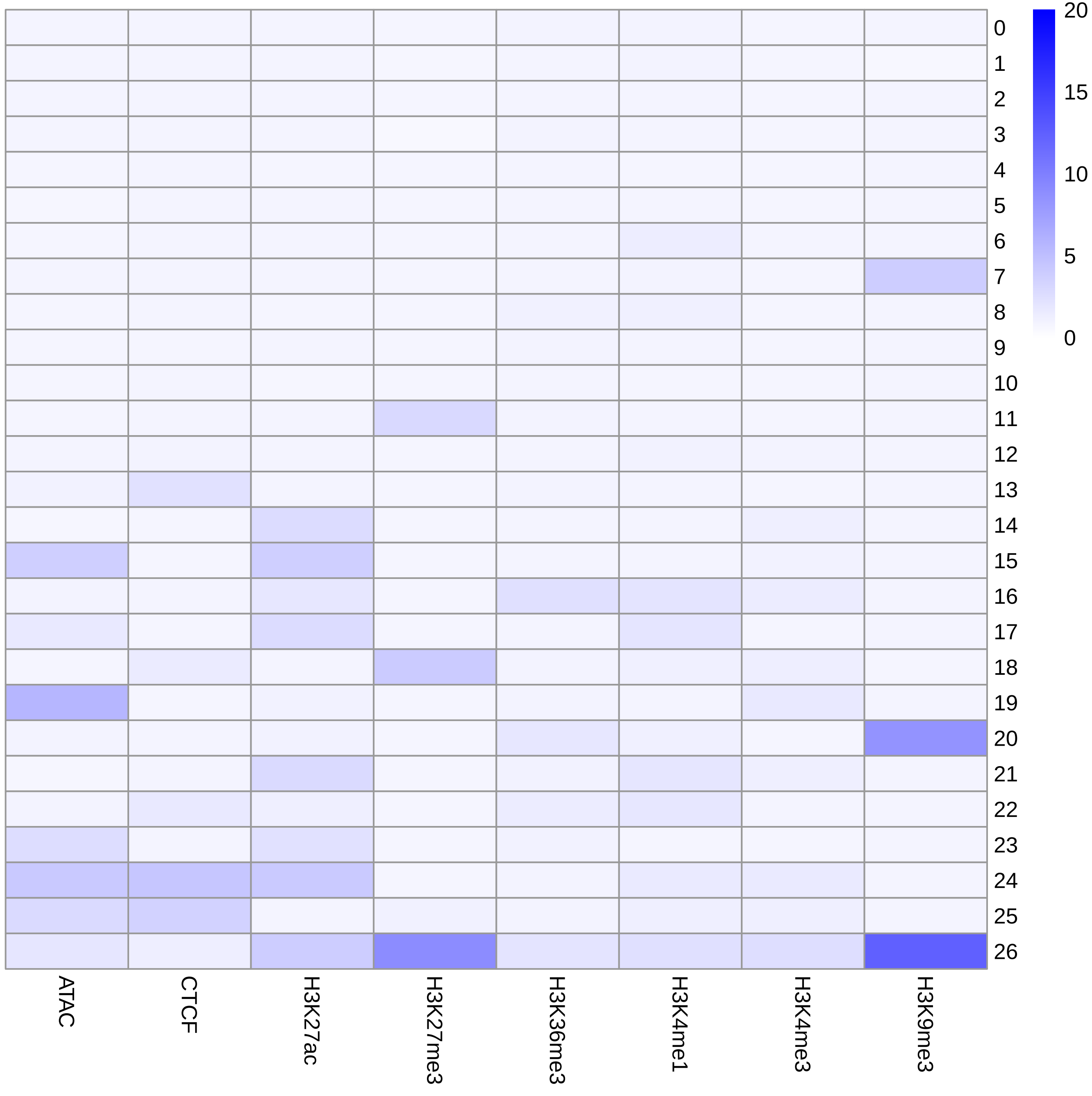

### 03_S3V2_IDEAS_hg38_r2_with_prior_J.pdf

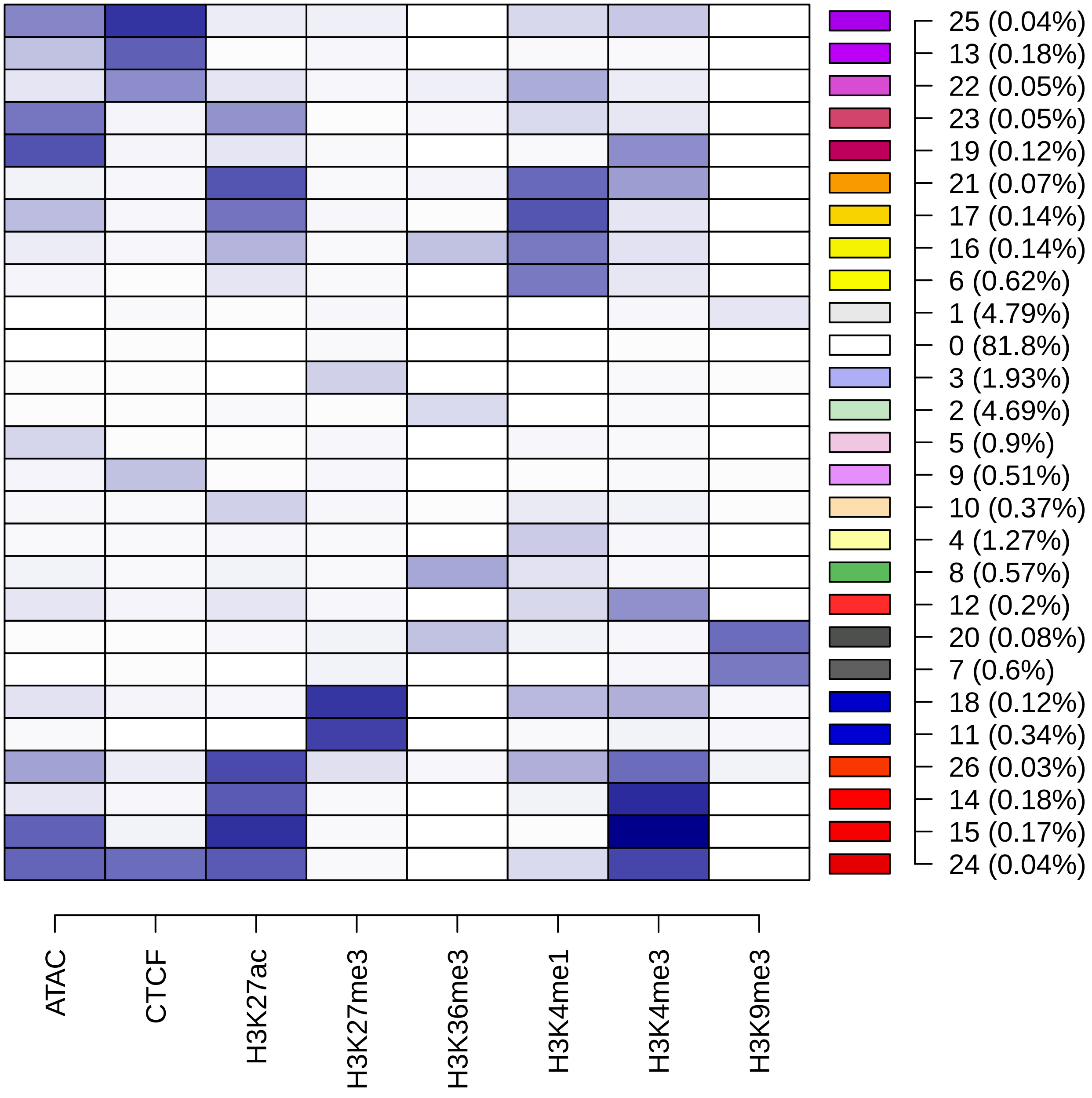

### 03a_S3V2_IDEAS_hg38_ccre2.pdf

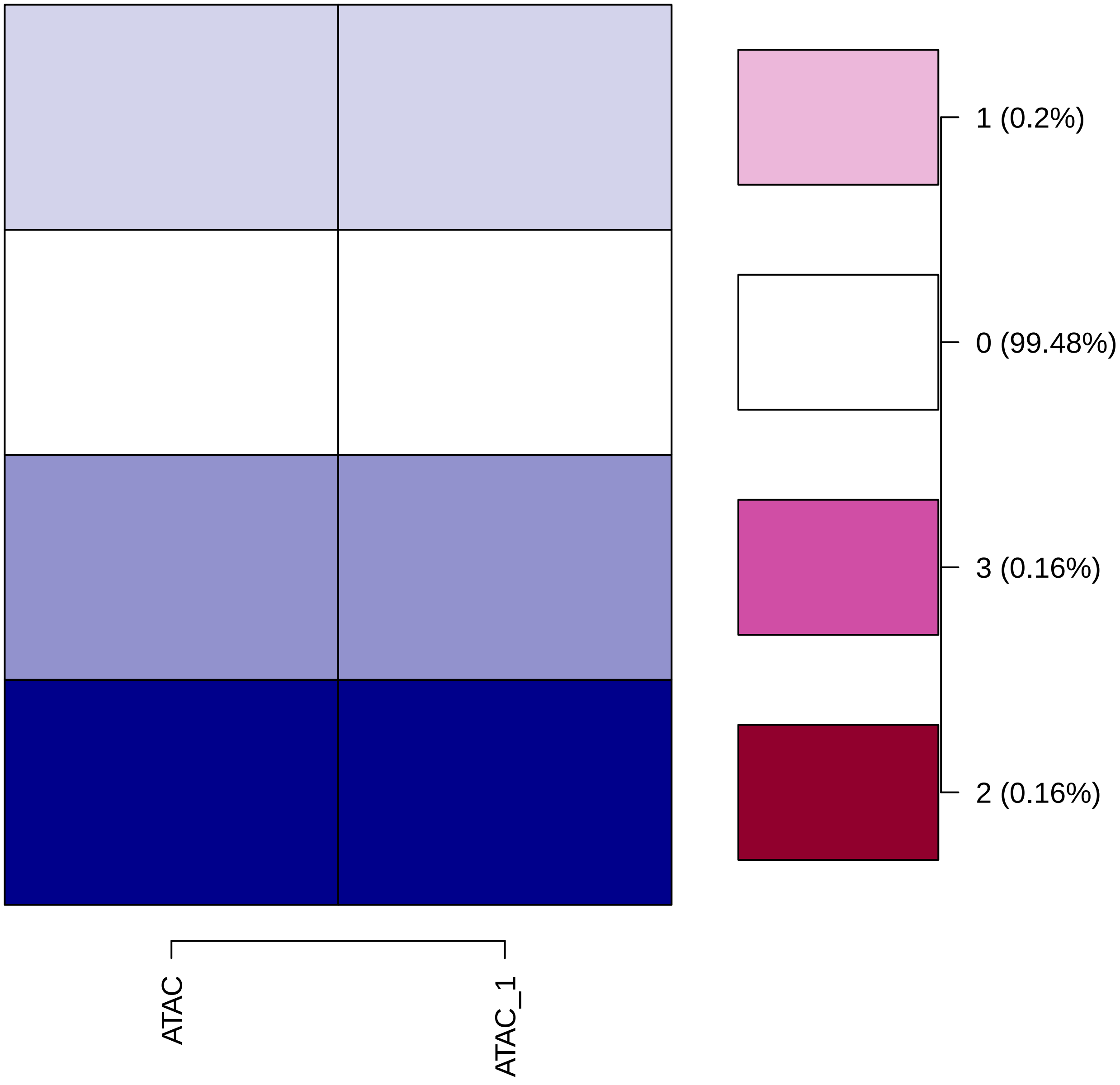

### 03b_S3V2_IDEAS_mm10_ccre2.pdf

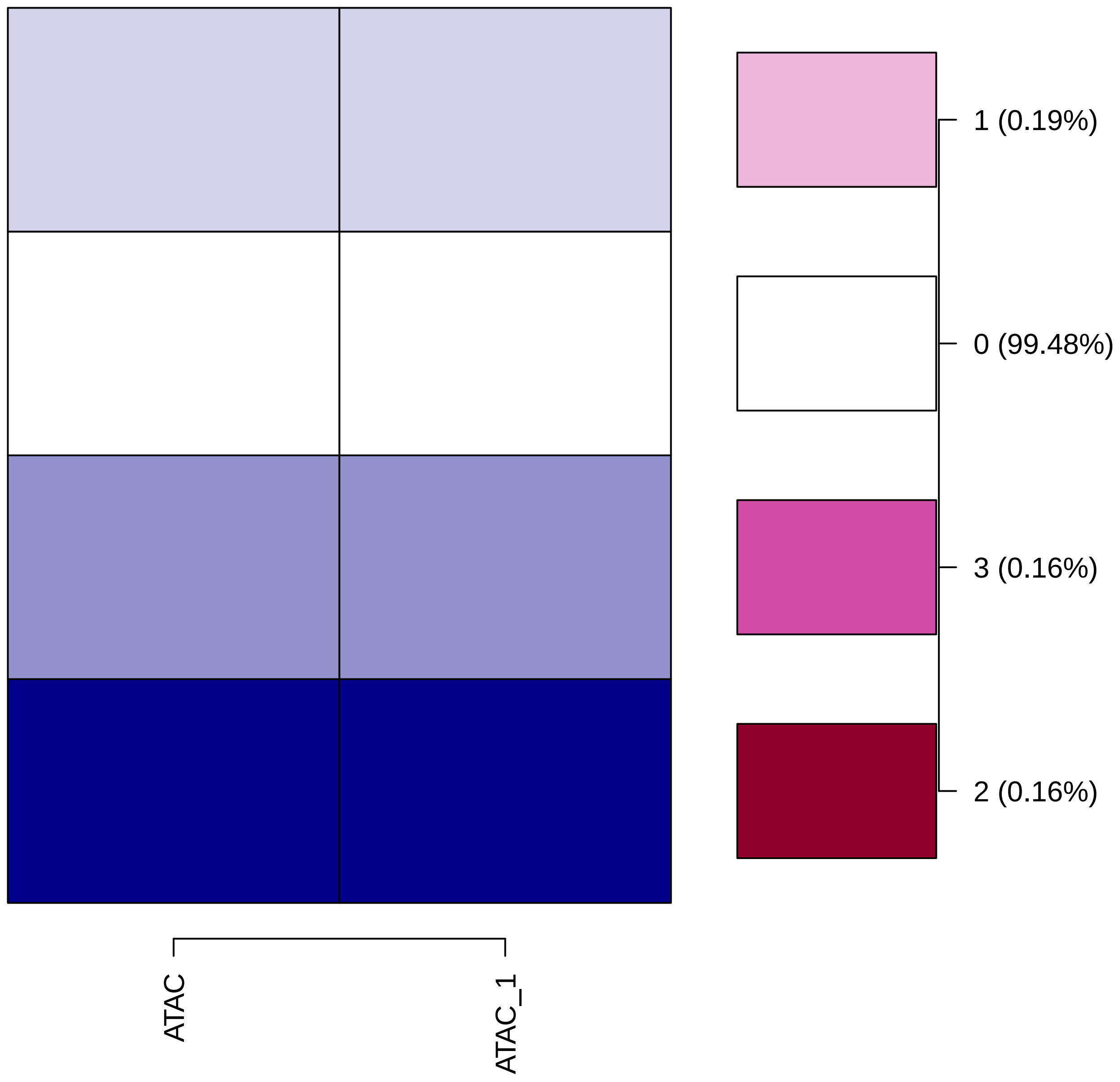

### 04_S3V2_IDEAS_mm10_r2_withhg38prior.all.m_v.r2.pdf

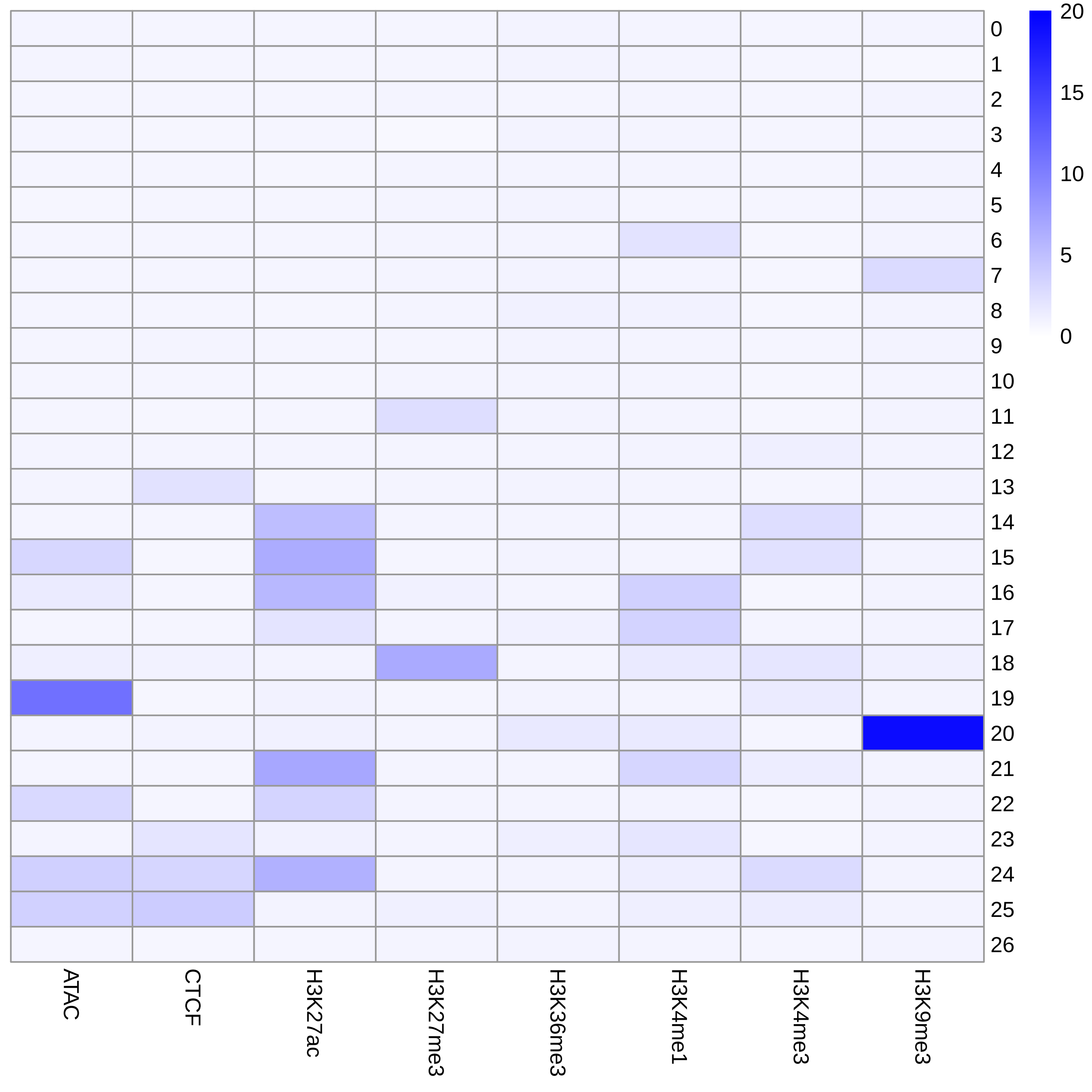

### 04_S3V2_IDEAS_mm10_r2_withhg38prior.pdf

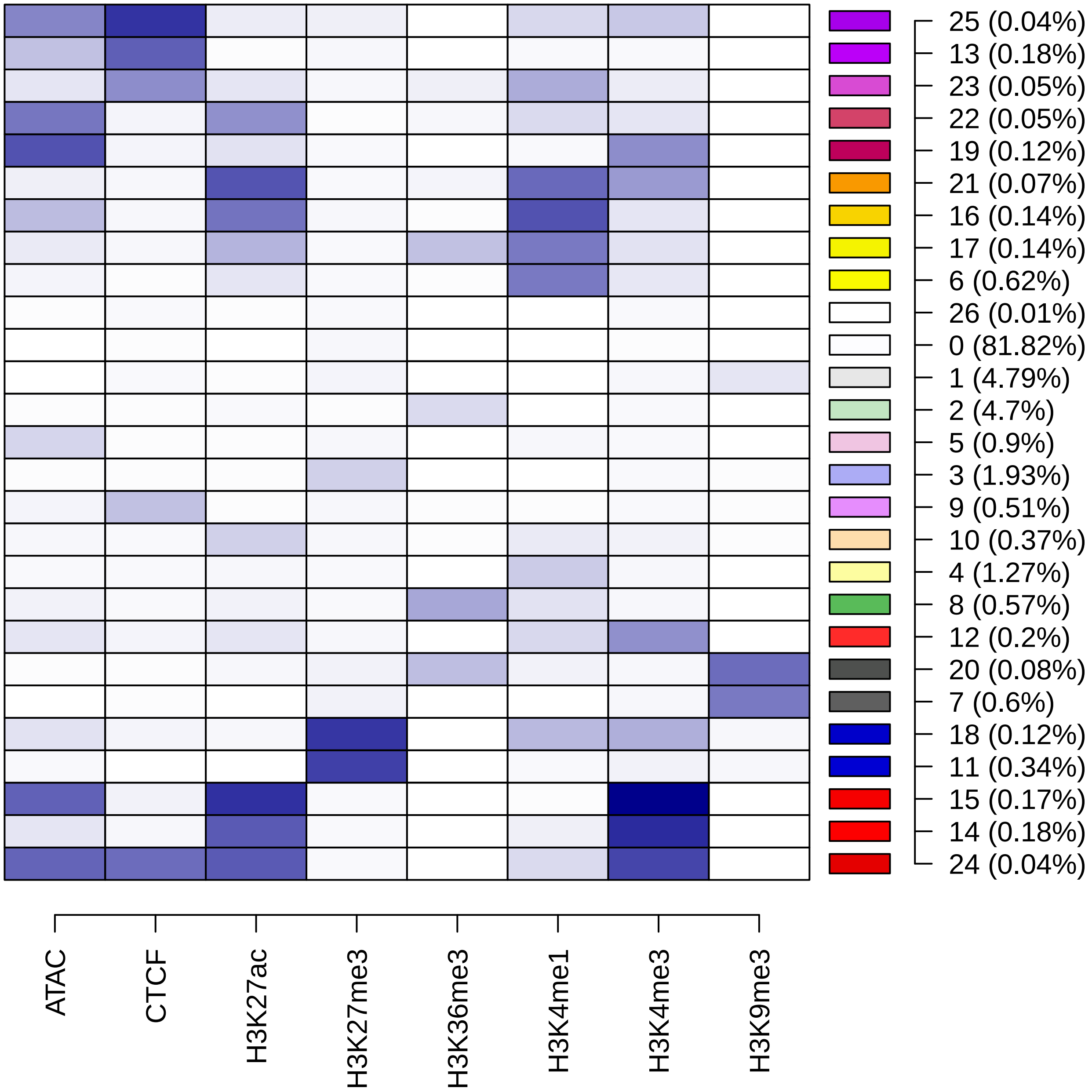

### 05_S3V2_IDEAS_mm10_r2_withhg38prior.rmh.10.para.pdf

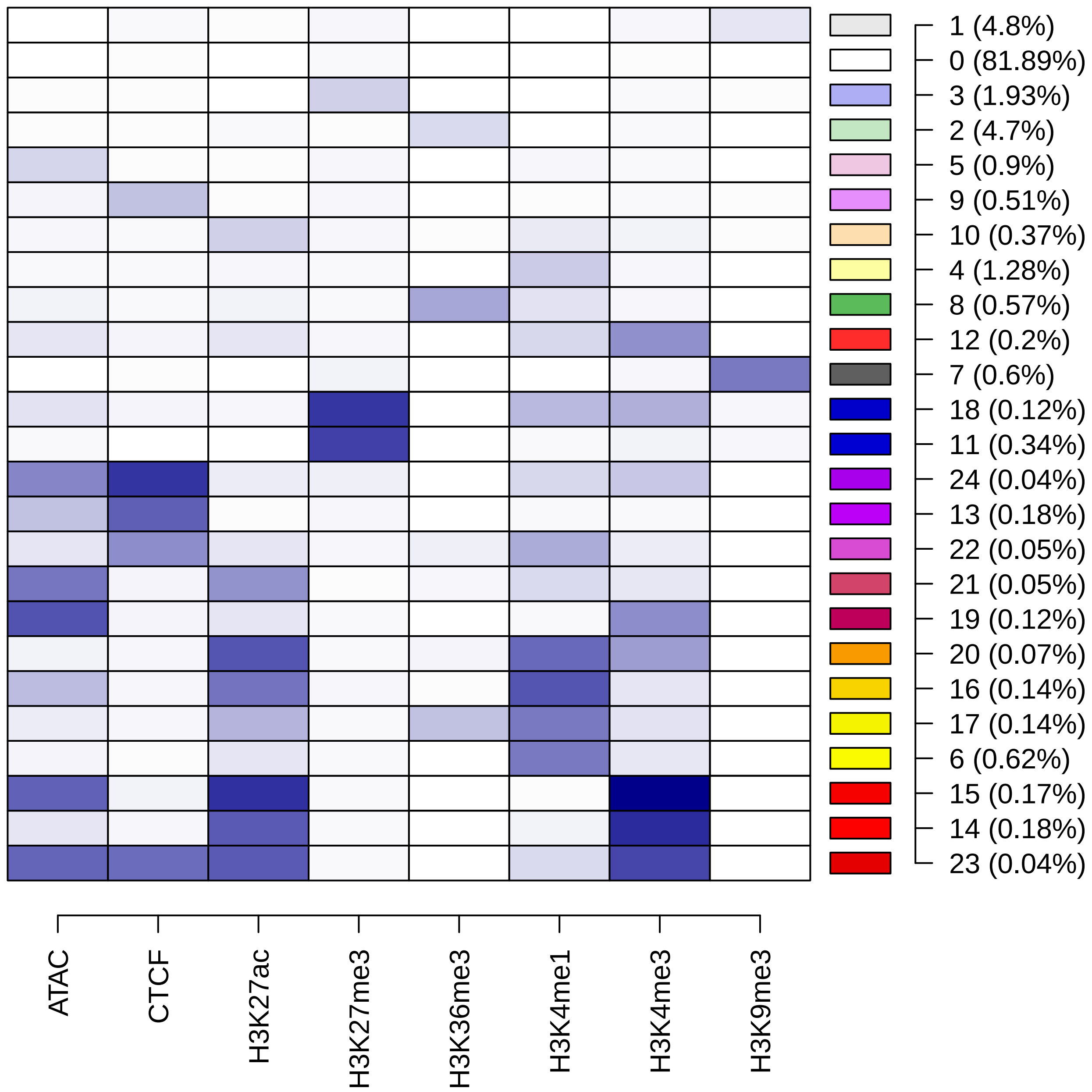

### 06a_S3V2_IDEAS_hg38_r3_withHg38Mm10prior.all.m_v.r3.pdf

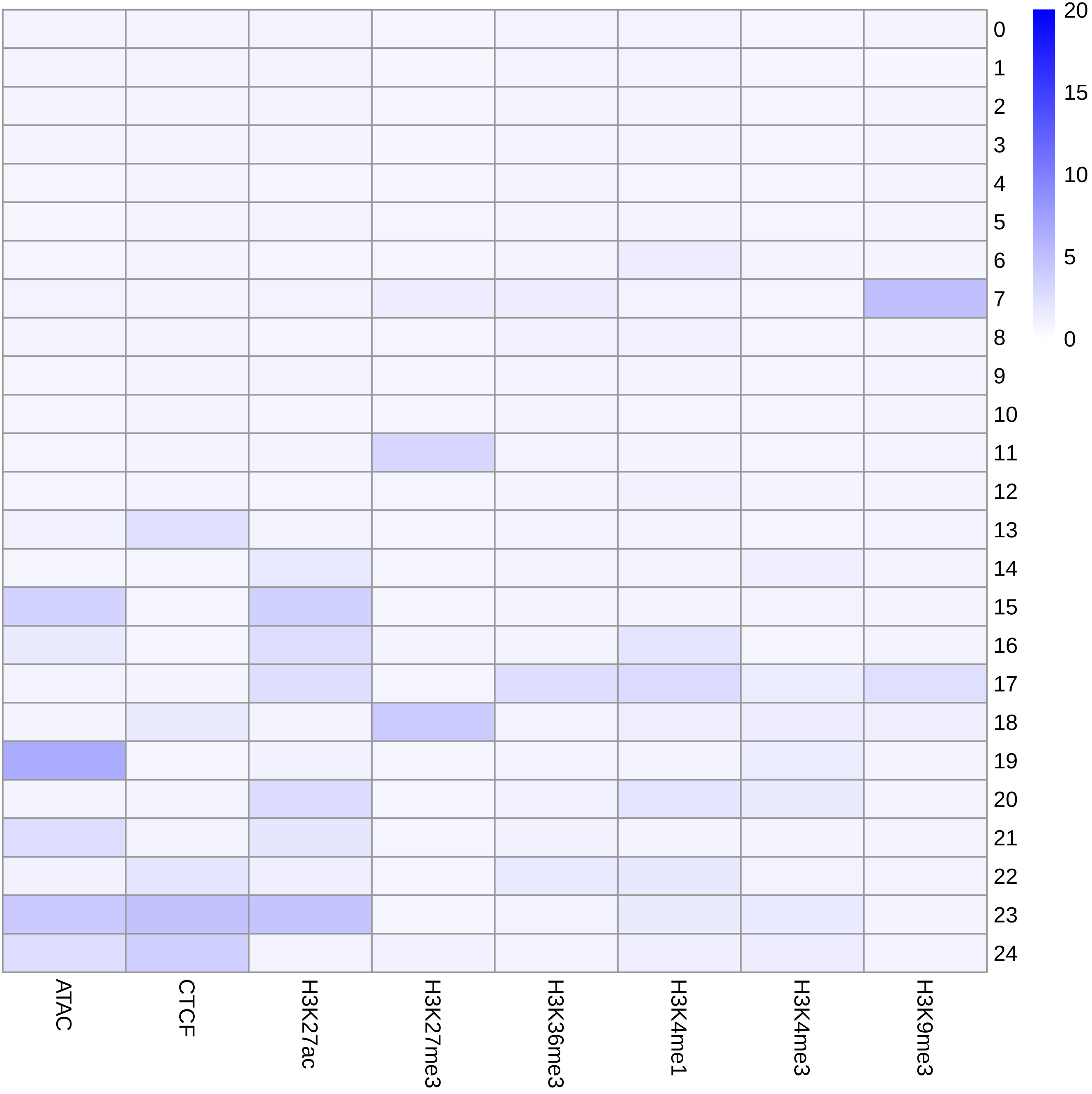

### 06a_S3V2_IDEAS_hg38_r3_withHg38Mm10prior.para.modified.para.pdf

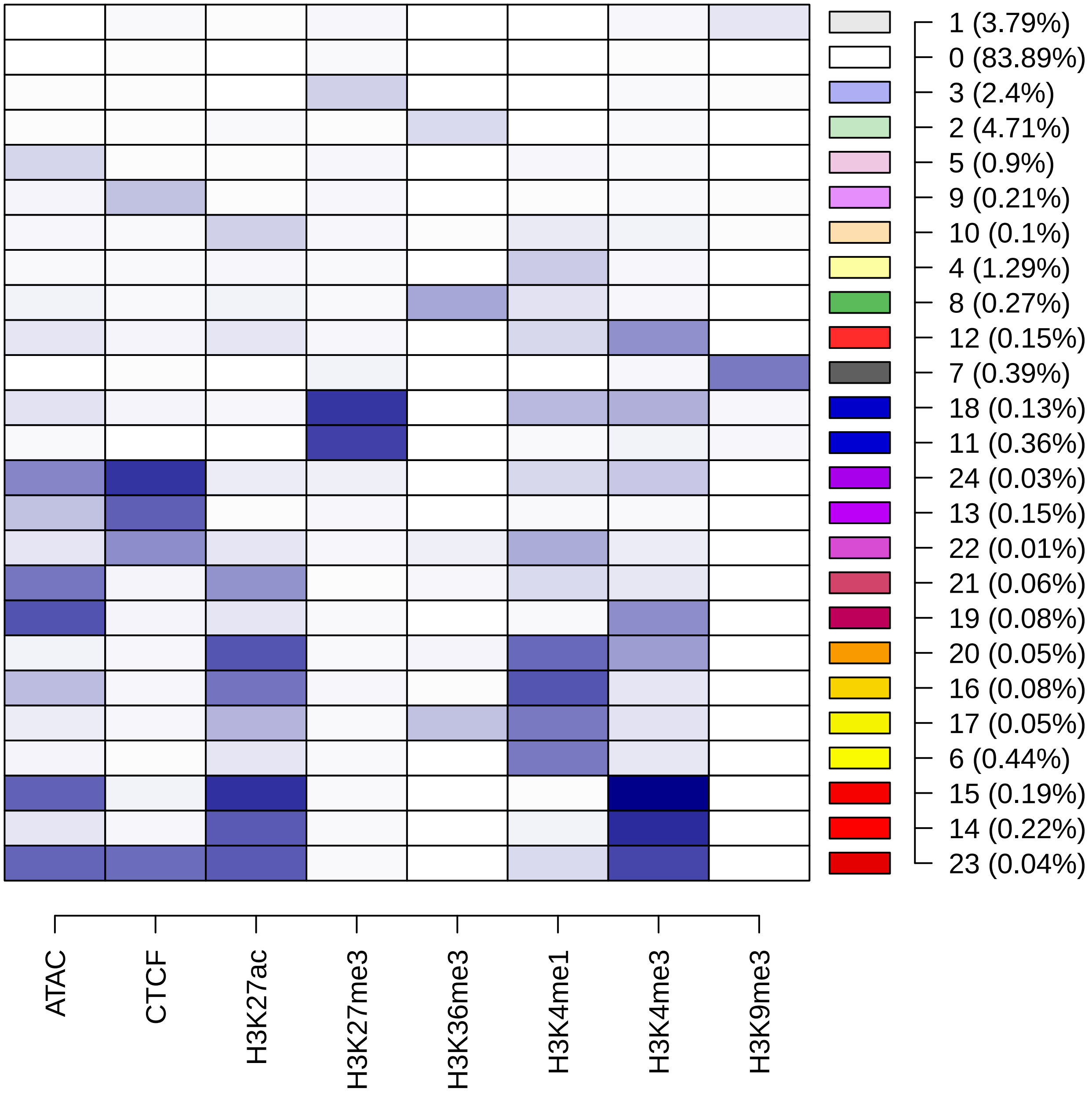

### 06b_S3V2_IDEAS_mm10_r3_withHg38Mm10prior.all.m_v.r3.pdf

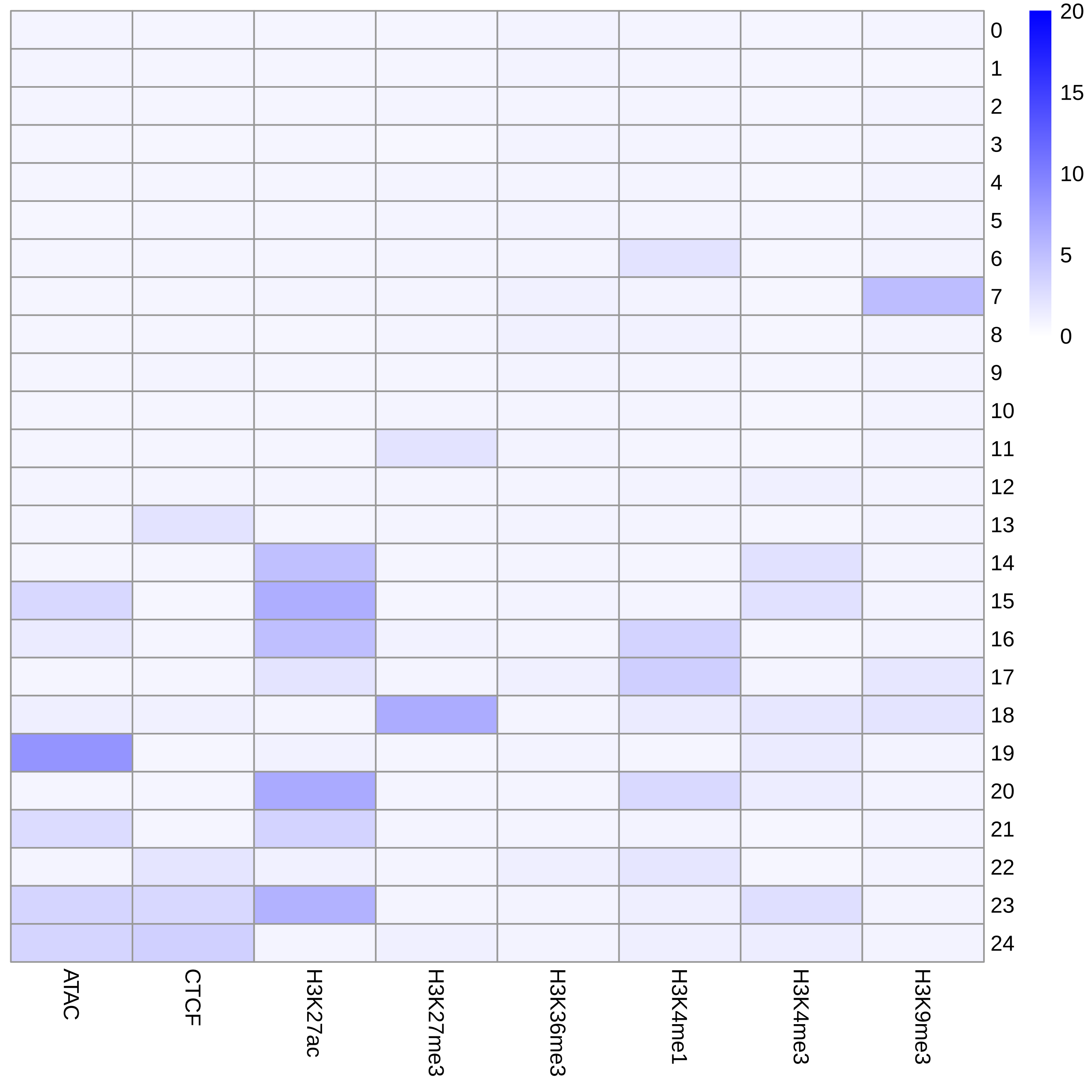

### 06b_S3V2_IDEAS_mm10_r3_withHg38Mm10prior.para.modified.para.pdf

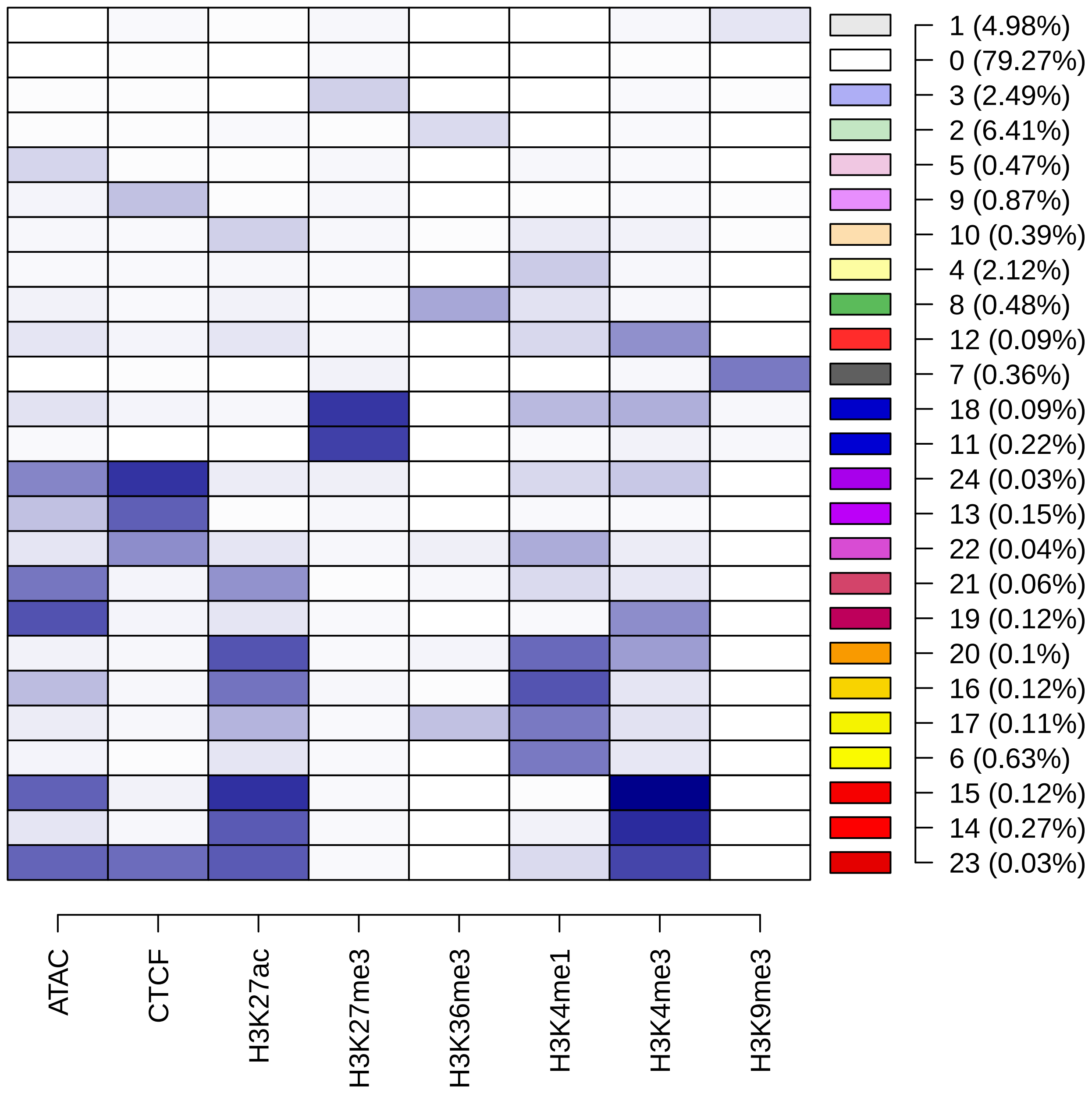

### enrichment_volcano_fmshape.pdf

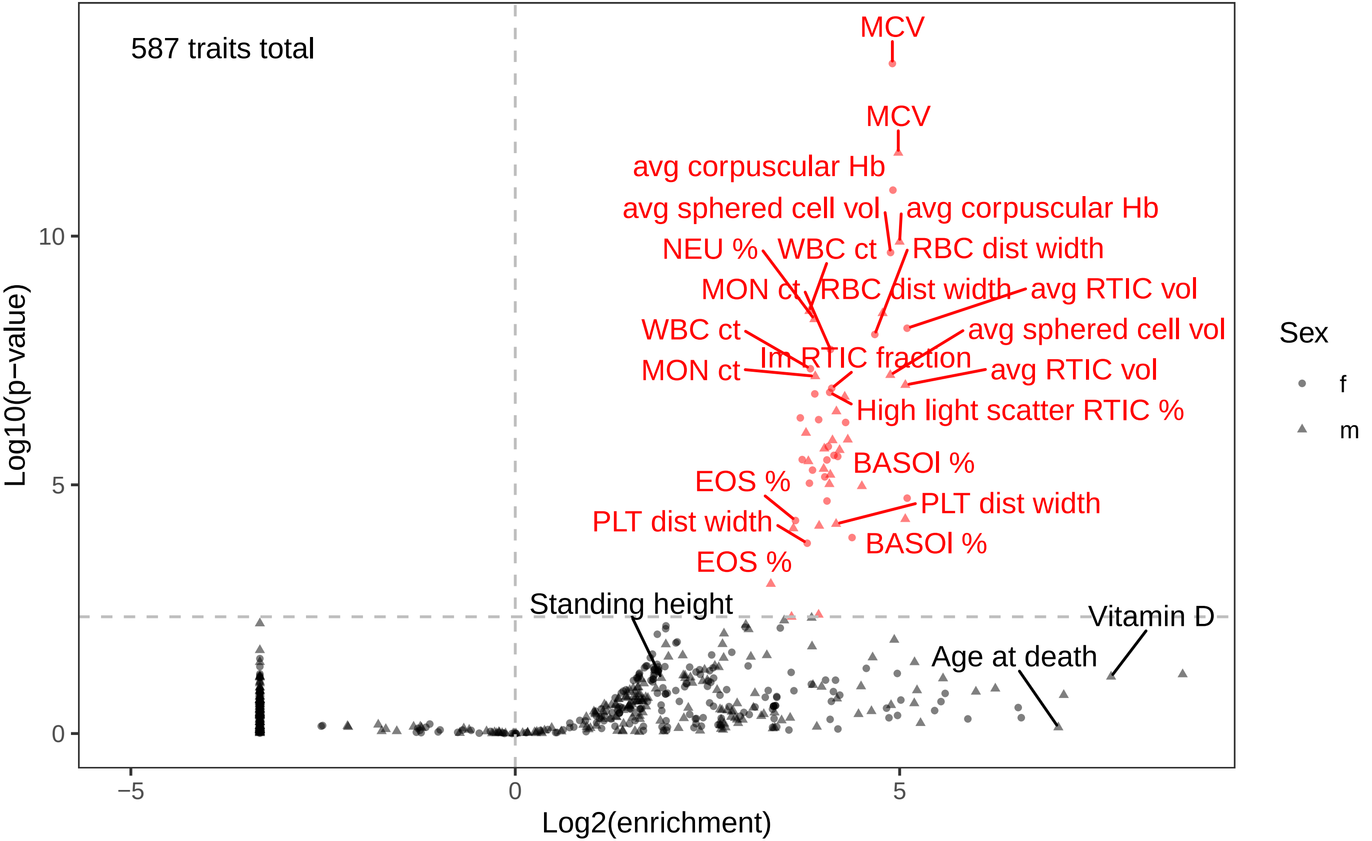

### jmc_combo_straight.pdf

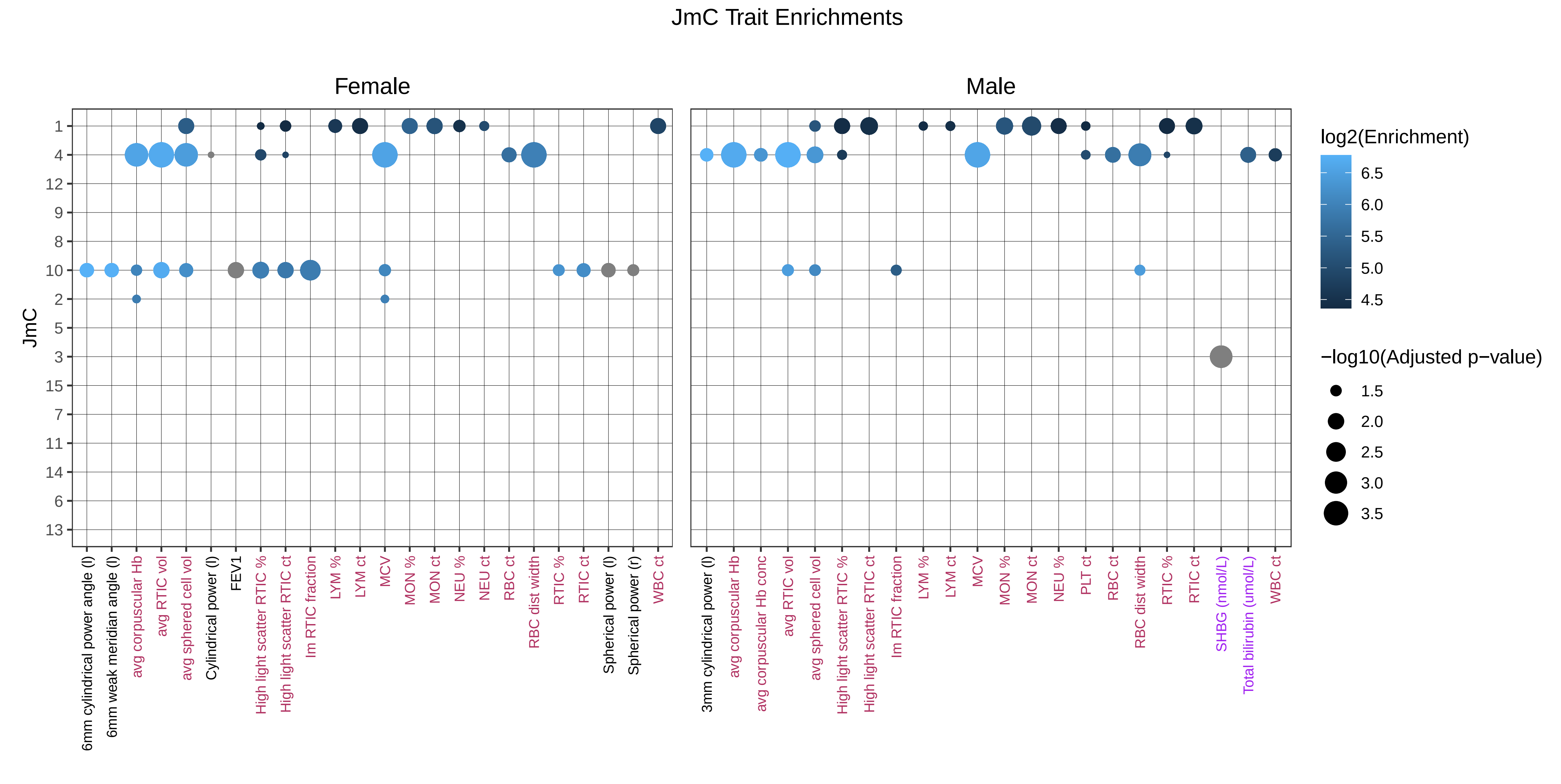

### Joint-VISION-rationale.pdf

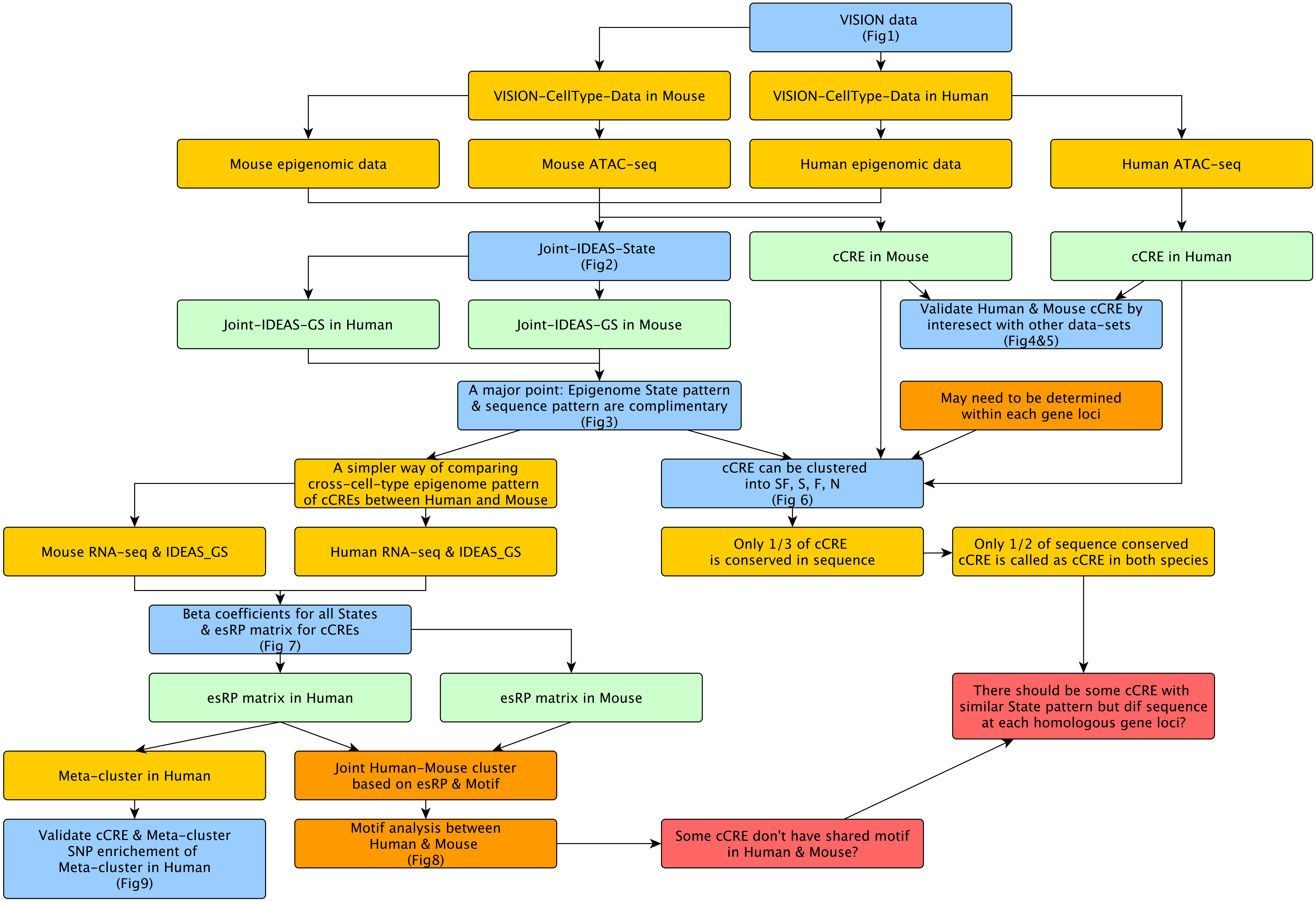

### Joint_ES_mm10_hg38_ES_pipeline.pdf

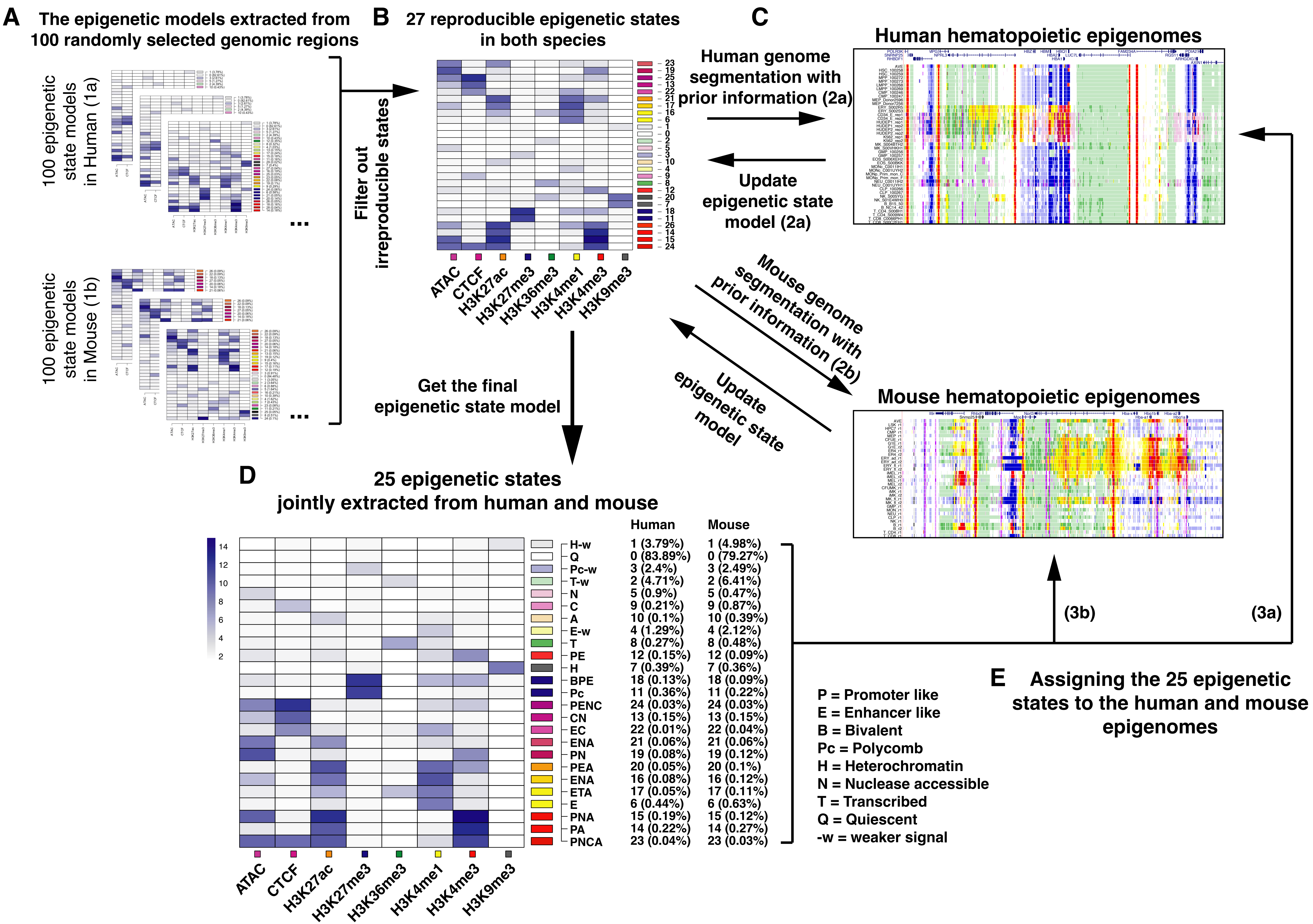

### S3V2_IDEAS_J25_withHg38Mm10prior.pdf

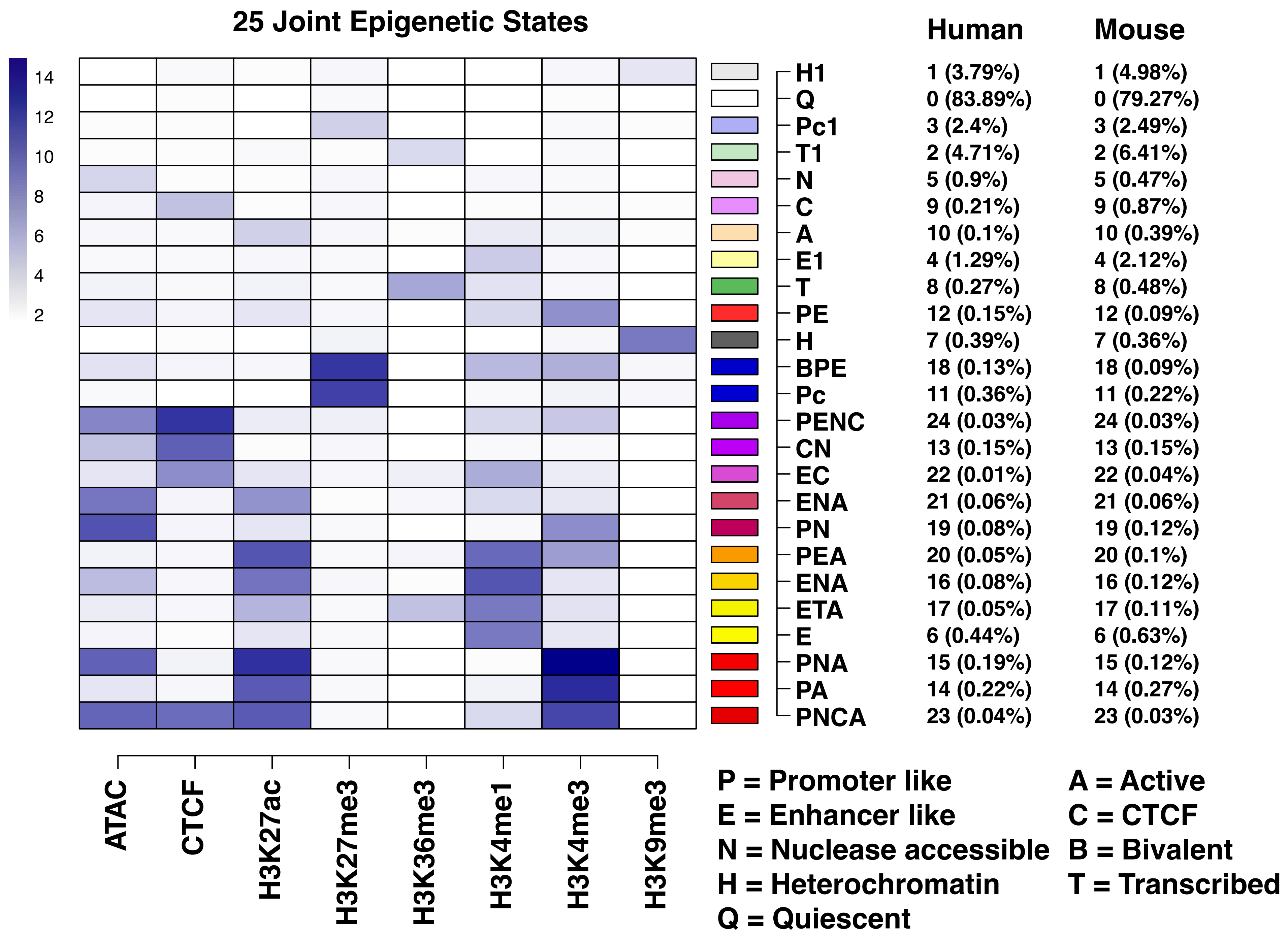

### two_rounds.all.m_v.pdf

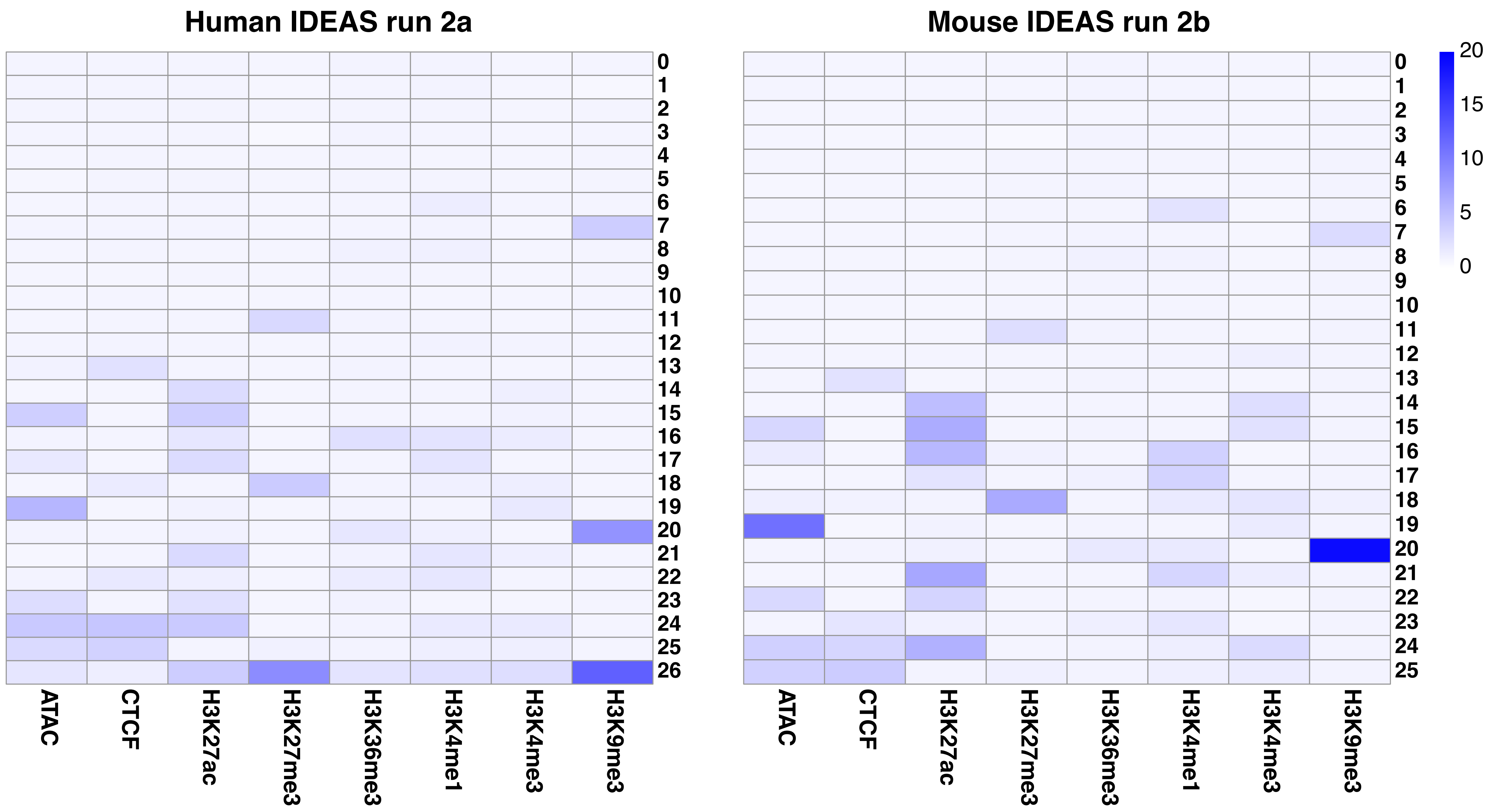
