## Supplementary material for "Interspecies regulatory landscapes and elements revealed by novel joint systematic integration of human and mouse blood cell epigenomes": Scripts for IDEAS modeling of epigenetic states jointly in human and mouse: get_joint_mouse_human_states.R

```
args = commandArgs(trailingOnly=TRUE)

mouse_front = args[1]
human_front = args[2]
output_file = args[3]
n_tmp_file = as.numeric(args[4])
n_used = as.numeric(args[5])
rp = as.numeric(args[6])
#time Rscript get_joint_mouse_human_states.R /absolute_path_to_folder1/hg38bp_forJoint /absolute_path_to_folder1/mm10vis_forJoint output_file_human_mouse_joint_state.para
#human_front = '/storage/home/gzx103/scratch/S3V2norm_compare/human_vision_S3V2/hg38bp0402_forJoint_IDEAS_output_A1_rerun/hg38bp0402_forJoint'
#mouse_front = '/storage/home/gzx103/scratch/S3V2norm_compare/mouse_mm10_for_pipeline_paper_0723_wg/VISION_mouse_ES_S3V2_7mk_NOprior_forJoint_IDEAS_output/VISION_mouse_ES_S3V2_7mk_NOprior_forJoint'
#output = 'mouse_human_joint_state.para'


makepara<-function(myorder,id,mem,mycut,para)
{       rt=NULL;
        j=0;
        for(i in myorder)
        {       t=which(mem==i);
                if(length(unique(id[t]))>mycut)
                {       rt=rbind(rt,cbind(j,array(para[t,],dim=c(length(t),dim(para)[2]))));
                        j=j+1;
                }
        }

        return(rt);
}

getmean<-function(n)
{
        N=NULL;
        for(i in sort(unique(n[,1])))
        {       t=which(n[,1]==i);
                N=rbind(N,apply(array(n[t,],dim=c(length(t),dim(n)[2])),2,sum)[-1]);
        }
        NN=N[,-1]/N[,1];

        return(array(NN,dim=c(length(NN)/(dim(n)[2]-2),dim(n)[2]-2)));
}

compareTwo<-function(n, m, myplot=0)
{       NN=getmean(n);
        MM=getmean(m);

  p=(-3+sqrt(9+8*(dim(n)[2]-2)))/2;
        dd=NULL;
        for(i in 1:dim(NN)[1])
        {       dd[i]=min(apply(array(MM[,1:p],dim=c(dim(MM)[1],p)),1,function(x){sqrt(sum((x-NN[i,1:p])^2))}));
        }
        for(i in 1:dim(MM)[1])
        {       dd[i+dim(NN)[1]]=min(apply(array(NN[,1:p],dim=c(dim(NN)[1],p)),1,function(x){sqrt(sum((x-MM[i,1:p])^2))}));
        }

        if(myplot>0)
        {       M=rbind(NN[,1:myplot],MM[,1:myplot]);
                rownames(M)=c(1:dim(NN)[1],paste("R",1:dim(MM)[1],sep=""));
                source("/storage/home/yzz2/work/IDEAS/ideas_source/heatmap3.R");
                source("/storage/home/yzz2/work/IDEAS/ideas_source/draw.R");
                heatmapwrap(M,cols=c("black","blue","cyan","green","yellow","red","pink"));
        }
        return(dd);
}

combineState<-function(parafiles, method="ward.D", mycut=0.9, pcut=1., fout=NULL)
{ 
#method="ward.D", 
#mycut=0.9, 
#pcut=1., 
rt=c()
nrt=c()
X=K=I=NULL;
  library("data.table");
  myheader=p=NULL;


  ### remove para only has 1 state
  parafiles_no1 = c()
  for(i in 1:length(parafiles))
  {
    x=read.table(parafiles[i], header=T, comment.char='~')
    if (dim(x)[1]>1){
      parafiles_no1 = c(parafiles_no1, parafiles[i])
    }
  }
  parafiles = parafiles_no1


  for(i in 1:length(parafiles))
  { 
  #x=fread('/gpfs/scratch/gzx103/S3V2norm_compare/mouse_mm10_for_pipeline_paper_0723_wg/VISION_mouse_ES_S3V2_7mk_NOprior_IDEAS_output/VISION_mouse_ES_S3V2_7mk_NOprior.modified.10.para')
  x=fread(parafiles[i]);
  t=max(which(is.na(x[1,])==F));
  #x[,1] = lapply(x[,1], as.numeric)
  #x = x*(600338505/sum(x[,1]))
  x=as.matrix(x[,1:t])
    x=as.matrix(rbind(x))
    x=as.matrix(x[which(x[,1]>=sum(x[,1])/1e4 & x[,1]>10),]);
    if(i==1)
    { myheader=colnames(x);
      p=sqrt(9/4-2*(1-length(myheader)))-3/2;
    }
    m=match(myheader[1:p+1],colnames(x)[1:p+1]);
    v=NULL;for(ii in 1:p)for(jj in 1:ii)
    {a=max(m[ii],m[jj]);b=min(m[ii],m[jj]);v=c(v,a*(a+1)/2+b-a);}
    X=rbind(X, array(as.matrix(x[,c(1,1+m,1+p+v)]),dim=c(length(x)/(1+p+length(v)),1+p+length(v))));
    K=c(K,dim(x)[1]);
    I=c(I,rep(i,dim(x)[1]));
    print(K)
    print(dim(x))
  }
  N=length(parafiles);
  
  p = (sqrt(1 + dim(X)[2] * 8) - 3) / 2;
  omycut=mycut;
  mycut=round(length(parafiles)*mycut);

  M=array(X[,1:p+1]/X[,1], dim=c(dim(X)[1],p));
  V=array(0,dim=c(dim(X)[1]*p,p));
  for(i in 1:dim(X)[1])
  { t = (i - 1) * p;
    l = 1;
    for(j in 1:p)
    { for(k in 1:j)
      { V[t + j, k] = V[t + k, j] = X[i,1+p+l] / X[i,1] - M[i,j] * M[i,k];
        l=l+1;
      }
    }
    V[t+1:p,]=t(solve(chol(V[t+1:p,]+diag(1e-1,p))));
#diag(V[t+1:p,])=sqrt(diag(V[t+1:p,]));
  }


  D=array(0,dim=rep(dim(X)[1],2));
  for(i in 2:dim(X)[1])
    for(j in 1:(i-1))
    { D[i,j]=D[j,i]=sqrt((sum((V[(i-1)*p+1:p,]%*%(M[i,]-M[j,]))^2) + sum((V[(j-1)*p+1:p,]%*%(M[i,]-M[j,]))^2)));
      #D[i,j]=D[j,i]=sqrt(sum((M[i,]-M[j,])^2/(diag(V[(i-1)*p+1:p,])^2+diag(V[(j-1)*p+1:p,])^2)));
      #D[i,j]=D[j,i]=sqrt(sum((M[i,]-M[j,])^2));#/(diag(V[(i-1)*p+1:p,])^2+diag(V[(j-1)*p+1:p,])^2)));
    }

  MM = NULL;
  kk=NULL;
  for(i in 1:N)
  { t=1:K[i];
    if(i > 1) t = t + sum(K[1:(i-1)]);
    t=(1:dim(D)[1])[-t];
    h=hclust(as.dist(D[t,t]),method=method);
    k=-1;
    tM = NULL;
    for(j in min(K):(min(length(t),max(K)*2)))
    { m=cutree(h,k=j);
      tt=NULL;
      for(l in 1:j)
      { tt[l] = length(unique(I[t[which(m==l)]]));
      }
      tk=length(which(tt>mycut));
      if(tk > k)
      { k = tk;
        tM = makepara(1:j,I[t],m,mycut,X[t,]);
      } else if(tk < k) { break; }
    }
    kk[i]=k;
    MM = rbind(MM, cbind(i, tM));
  }

  print(kk)
  mysel = median(kk);
  h=hclust(as.dist(D),method=method);
  rt=rep(0,max(K)*2);
  k = -1;
  for(i in min(K):min(dim(D)[1],max(K)*2))
  { m=cutree(h,k=i);
    tt = NULL;
    for(j in 1:i)
    { tt[j] = length(unique(I[which(m==j)]));
    }
    tk=length(which(tt>mycut));
    if(tk==mysel | tk < k) break;
    k = tk;
    rt[i] = length(which(tt>mycut));
  }
  mysel = max(k,tk);
  
  m=cutree(h,k=mysel);
  nn = NULL;
  for(i in 1:mysel)
  { t=which(m==i);
    nn[i] = sum(X[t,1]);
  }
  oo=order(nn,decreasing=T);
  rt = makepara(oo,I,m,mycut,X);
  onstate = max(rt[,1])+1;
        ooo=NULL;
  for(i in oo)
  { t=which(m==i);
          if(length(unique(I[t]))>mycut) ooo=c(ooo,i);
  }

  d=NULL;
  for(i in 1:N)
  { d=rbind(d,compareTwo(rt,MM[MM[,1]==i,-1])[1:onstate]);
  }
  #boxplot(d);
  dd=array(cutree(hclust(dist(c(d))),k=2),dim=dim(d));
  kk=table(c(dd));
  kk=which(as.integer(kk)==max(as.integer(kk)))[1];
  pp=apply(dd,2,function(x){length(which(x!=kk))/length(x)});
  pp0=apply(d,2,function(x){length(which(x>0.5))/length(x)});
  #print(rbind(pp,pp0));
  pp[pp0<pp]=pp0[pp0<pp];
  t=which(pp>pcut);
  if(length(t)>0)
  { j=0;
    nrt = NULL;
    for(i in (1:onstate-1)[-t])
    { nrt = rbind(nrt, cbind(j, rt[rt[,1]==i,-1]));
      j=j+1;
    }
    rt=nrt;
    ooo=ooo[-t];
  }
  message(paste("nstate =", onstate, "-->", max(rt[,1])+1));

  nrt=NULL;
  for(i in 0:max(rt[,1]))
  { t=which(rt[,1]==i);
    nrt=rbind(nrt, apply(array(rt[t,],dim=c(length(t),dim(rt)[2])),2,sum)[-1]);
  }
  rt=nrt;
  colnames(rt)=myheader;

  write.table(rt,fout,quote=F,row.names=F);

}


parafiles = c()

set.seed(2019)
used_id = sample(n_tmp_file, n_used)

for (i in used_id){
  parafiles = c(parafiles, paste(human_front, '.tmp.', i, '.para', sep=''))
  parafiles = c(parafiles, paste(mouse_front, '.tmp.', i, '.para', sep=''))
}


mycut_n = rp
a=combineState(parafiles, method="ward.D", mycut=mycut_n, pcut=1., fout=output_file)
```
