## Supplementary material for "Interspecies regulatory landscapes and elements revealed by novel joint systematic integration of human and mouse blood cell epigenomes": Scripts for IDEAS modeling of epigenetic states jointly in human and mouse: get_cCRE.pipeline.py

```
import os
import glob
import os.path
import numpy as np
import subprocess
from subprocess import call
from multiprocessing import Pool

################################################################################################
### read 2d array
def read1d_array(filename,dtype_used):
	import numpy as np
	data=open(filename,'r')
	data0=[]
	for records in data:
		tmp = [x.strip() for x in records.split('\t')]
		data0.append(tmp[0])
	data.close()
	return(data0)

### get cCRE
def get_cCRE_pipeline(script_dir, id_name, working_dir, bin_size):
	### set working directory
	os.chdir(working_dir)
	### get bins
	b=call('Rscript '+script_dir+'/get_bins_from_IDEAS.n4.R'+' '+id_name, shell=True)


	print('get bed file done')

	### remove problematic NEU chr8 peaks
	b1 = call('for file in $(cat neu_files.txt.tmp); do cat $file | awk -F \'\t\' -v OFS=\'\t\' \'{if ($1!="chr8") print $0}\' > $file.tmp && mv $file.tmp $file; done', shell=True)	
	#b1 = call('cat S3V2_IDEAS_hg38_ccre2.ct.list.txt | awk \'{if ($1!="NEU_C0011IH2" && $1!="NEU_C001UYH1") print $0}\' > tmp1111 && mv tmp1111 S3V2_IDEAS_hg38_ccre2.ct.list.txt', shell=True)

	### define state combinations
	HS=['3','32','321']
	### get bin merged bed
	for st in HS:
		b=call('bedtools merge -i '+id_name+'.AVERAGE.'+st+'.bed'+' > '+id_name+'.AVERAGE.'+st+'.M.bed', shell=True)
		b=call('cp '+id_name+'.AVERAGE.'+st+'.M.bed'+' '+id_name+'.AVERAGE_allct.'+st+'.M.bed', shell=True)

	### read ct list
	ct_list=read1d_array(id_name+'.ct.list.txt', str)
	print(ct_list)

	### merge 3 and 32 because only using 3 may miss some important example cCREs
	for ct in ct_list:
		e = call('cp '+id_name+'.'+ ct +'.32.bed ' +id_name+'.'+ ct +'.3.bed', shell=True)


	AVE_id=ct_list.pop(0)
	for ct in ct_list:
		for st in HS:
			### mergine ct bins
			c=call('bedtools merge -i '+id_name+'.'+ct+'.'+st+'.bed'+' > '+id_name+'.'+ct+'.'+st+'.M.bed', shell=True)
			### intersect
			c=call('bedtools intersect -a '+id_name+'.'+ct+'.'+st+'.M.bed'+' -b '+id_name+'.AVERAGE_allct.'+st+'.M.bed'+' > tmp1.bed', shell=True)
			### ct uniq
			c=call('bedtools intersect -a '+id_name+'.'+ct+'.'+st+'.M.bed'+' -b '+id_name+'.AVERAGE_allct.'+st+'.M.bed -v'+' > tmp2.bed', shell=True)
			### ave uniq
			c=call('bedtools intersect -b '+id_name+'.'+ct+'.'+st+'.M.bed'+' -a '+id_name+'.AVERAGE_allct.'+st+'.M.bed -v'+' > tmp3.bed', shell=True)
			### pool all
			c=call('cat tmp1.bed tmp2.bed tmp3.bed | sort -k1,1 -k2,2n > tmp4.bed && mv tmp4.bed '+id_name+'.AVERAGE_allct.'+st+'.M.bed', shell=True)

	### get ct binary mat
	for st in HS:
		d=call('cp '+id_name+'.AVERAGE_allct.'+st+'.M.bed'+' '+id_name+'.AVERAGE_allct.'+st+'.M.mat.txt', shell=True)

	### get columns
	for ct in ct_list:
		for st in HS:
			e=call('bedtools intersect -a '+id_name+'.AVERAGE_allct.'+st+'.M.bed'+' -b '+id_name+'.'+ct+'.321.bed'+' -c > tmp1.txt', shell=True)
			e=call('cut -f4 tmp1.txt > tmp2.txt', shell=True)
			e=call('paste '+id_name+'.AVERAGE_allct.'+st+'.M.mat.txt'+' '+'tmp2.txt > '+id_name+'.AVERAGE_allct.'+st+'.M.mat.txt.tmp', shell=True)
			e=call('mv '+id_name+'.AVERAGE_allct.'+st+'.M.mat.txt.tmp'+' '+id_name+'.AVERAGE_allct.'+st+'.M.mat.txt', shell=True)


	print(' remove unreproducible ones')
	for st in HS:
		f=call('Rscript '+script_dir+'/get_reproducible_pk.R '+id_name+'.ct.list.txt'+' '+id_name+'.AVERAGE_allct.'+st+'.M.mat.txt'+' '+id_name+'.AVERAGE_allct.'+st+'.M.rep0.bed', shell=True)
		f=call('bedtools merge -i '+id_name+'.AVERAGE_allct.'+st+'.M.rep0.bed'+' > '+id_name+'.AVERAGE_allct.'+st+'.M.rep1.bed', shell=True)
		f=call('bedtools intersect -a '+id_name+'.AVERAGE_allct.'+st+'.M.rep1.bed'+' -b '+id_name+'.AVERAGE.'+st+'.M.bed'+' -v > '+'tmp5.bed', shell=True)
		f=call('cat '+id_name+'.AVERAGE.'+st+'.M.bed'+' tmp5.bed | sort -k1,1 -k2,2n > '+id_name+'.AVERAGE.'+st+'.M.rep.bed', shell=True)

	print(' get final cCRE listi')
	g=call('bedtools intersect -a '+id_name+'.AVERAGE_allct.32.M.rep1.bed'+' -b '+id_name+'.AVERAGE_allct.3.M.rep1.bed -v > '+id_name+'.AVERAGE_allct.32.NOT.3.M.rep.bed', shell=True)
	g=call('bedtools intersect -a '+id_name+'.AVERAGE_allct.321.M.rep1.bed'+' -b '+id_name+'.AVERAGE_allct.32.M.rep1.bed -v > '+id_name+'.AVERAGE_allct.321.NOT.32.M.rep.bed', shell=True)
	g=call('cat '+id_name+'.AVERAGE.3.M.rep.bed'+' '+id_name+'.AVERAGE_allct.32.NOT.3.M.rep.bed'+' '+id_name+'.AVERAGE_allct.321.NOT.32.M.rep.bed'+' | sort -k1,1 -k2,2n > '+id_name+'.cCRE.bed', shell=True)
	g=call('bedtools merge -d '+bin_size+' -i '+id_name+'.cCRE.bed > '+id_name+'.cCRE.M.bed', shell=True)

	### get ct cCREs
	for ct in ct_list:
		h=call('bedtools intersect -a '+id_name+'.'+ct+'.32.M.bed'+' -b '+id_name+'.'+ct+'.3.M.bed -v > '+id_name+'.'+ct+'.32.NOT.3.M.bed', shell=True)
		h=call('bedtools intersect -a '+id_name+'.'+ct+'.321.M.bed'+' -b '+id_name+'.'+ct+'.32.M.bed -v > '+id_name+'.'+ct+'.321.NOT.32.M.bed', shell=True)
		h=call('cat '+id_name+'.'+ct+'.3.M.bed'+' '+id_name+'.'+ct+'.32.NOT.3.M.bed'+' '+id_name+'.'+ct+'.321.NOT.32.M.bed'+' | sort -k1,1 -k2,2n > '+id_name+'.'+ct+'.cCRE.bed', shell=True)

	### clean folder
	#for ct in ct_list:
	#	for filename in glob.glob(id_name+'.'+ct+"*3*"):
	#		os.remove(filename)
	#for filename in glob.glob(id_name+'.'+AVE_id+"*3*"):
	#	os.remove(filename)
	#for filename in glob.glob("tmp*"):
	#	os.remove(filename)

############################################################################
#time python3 get_cCRE.pipeline.py -s $script_dir -i $id_name -w $working_dir -l $bin_size

import getopt
import sys
def main(argv):
	### read user provided parameters
	opts, args = getopt.getopt(argv,"hs:i:w:l:")
	for opt,arg in opts:
		if opt=="-h":
			print('time python3 get_cCRE.pipeline.py -s $script_dir -i $id_name -w $working_dir -l $bin_size')
			return()	
		elif opt=="-s":
			script_dir=str(arg.strip())
		elif opt=="-i":
			id_name=str(arg.strip())
		elif opt=="-w":
			working_dir=str(arg.strip())
		elif opt=="-l":
			bin_size=str(arg.strip())

	############ Default parameters
	###### required parameters
	try:
		print('User provide script_dir: -s '+str(script_dir))
		print('User provide id_name: -i '+str(id_name))
		print('User provide working_dir: -w '+str(working_dir))
		print('User provide bin_size: -l '+str(bin_size))
	except NameError:
		print('Missing required parameter(s):')	
		return()

	######### run get_signal_track
	print('start get_signal_track.......')
	get_cCRE_pipeline(script_dir, id_name, working_dir, bin_size)

if __name__=="__main__":
	main(sys.argv[1:])
```
