## Supplementary material for "Interspecies regulatory landscapes and elements revealed by novel joint systematic integration of human and mouse blood cell epigenomes": Scripts for IDEAS modeling of epigenetic states jointly in human and mouse: S3V2_IDEAS_hg38_ccre2.sh

```
#PBS -l nodes=1:ppn=2
#PBS -l walltime=80:00:00
#PBS -j oe
##PBS -A yzz2_e_g_sc_default
#PBS -A rch8_b_g_bc_default
#PBS -l pmem=40gb

IDEAS_job_name=S3V2_IDEAS_hg38_ccre2
script_dir=/storage/home/gzx103/scratch/joint_human_mouse/S3V2_IDEAS_ESMP/bin//IDEAS_2018/
working_dir=/storage/home/gzx103/scratch/joint_human_mouse/hg38/S3V2_IDEAS_outputs_hg38_CCRE/
output_dir=/storage/home/gzx103/scratch/joint_human_mouse/hg38/S3V2_IDEAS_outputs_hg38_CCRE//S3V2_IDEAS_hg38_ccre2_IDEAS_output/
binfile=/storage/home/gzx103/scratch/joint_human_mouse/hg38/S3V2_IDEAS_outputs_hg38_CCRE//windowsNoBlack.withid.IDEASbins.txt

cd $working_dir
if [ -d $output_dir ]; then rm -r $output_dir; mkdir $output_dir; else mkdir $output_dir; fi
if [ -d bin ]; then rm -r bin; cp -r $script_dir/bin ./ ; else cp -r $script_dir/bin ./ ; fi
if [ -d data ]; then rm -r data; cp -r $script_dir/data/ ./ ; else cp -r $script_dir/data ./ ; fi
time Rscript ./bin/runme.R $IDEAS_job_name'.input' $IDEAS_job_name'.parafile' $output_dir

time python3 /storage/home/gzx103/scratch/joint_human_mouse/S3V2_IDEAS_ESMP/bin/get_cCRE.pipeline.py -s /storage/home/gzx103/scratch/joint_human_mouse/S3V2_IDEAS_ESMP/bin/ -i S3V2_IDEAS_hg38_ccre2 -w /storage/home/gzx103/scratch/joint_human_mouse/hg38/S3V2_IDEAS_outputs_hg38_CCRE/S3V2_IDEAS_hg38_ccre2_IDEAS_output/ -l 200
```
