## Supplementary material for "Interspecies regulatory landscapes and elements revealed by novel joint systematic integration of human and mouse blood cell epigenomes": Scripts for IDEAS modeling of epigenetic states jointly in human and mouse: S3V2_IDEAS_mm10_r3_withHg38Mm10prior.sh

```
#PBS -l nodes=1:ppn=3
#PBS -l walltime=60:00:00
#PBS -j oe
##PBS -A yzz2_e_g_sc_default
#PBS -A rch8_b_g_bc_default
#PBS -l pmem=40gb

R --version

module load r/3.4 
source ~/.bash_profile

IDEAS_job_name=S3V2_IDEAS_mm10_r3_withHg38Mm10prior
script_dir=/storage/home/gzx103/scratch/joint_human_mouse/S3V2_IDEAS_ESMP/bin//IDEAS_2018/
working_dir=/storage/home/gzx103/scratch/joint_human_mouse/mm10/S3V2_IDEAS_outputs_mm10/
output_dir=/storage/home/gzx103/scratch/joint_human_mouse/mm10/S3V2_IDEAS_outputs_mm10//S3V2_IDEAS_mm10_r3_withHg38Mm10prior_IDEAS_output/
binfile=/storage/home/gzx103/scratch/joint_human_mouse/mm10/S3V2_IDEAS_outputs_mm10//windowsNoBlack.withid.IDEASbins.txt

cd $working_dir
if [ -d $output_dir ]; then rm -r $output_dir; mkdir $output_dir; else mkdir $output_dir; fi
if [ -d bin ]; then rm -r bin; cp -r $script_dir/bin ./ ; else cp -r $script_dir/bin ./ ; fi
if [ -d data ]; then rm -r data; cp -r $script_dir/data/ ./ ; else cp -r $script_dir/data ./ ; fi
time Rscript ./bin/runme.R $IDEAS_job_name'.input' $IDEAS_job_name'.parafile' $output_dir
```
