## Supplementary material for "Interspecies regulatory landscapes and elements revealed by novel joint systematic integration of human and mouse blood cell epigenomes": Scripts for IDEAS modeling of epigenetic states jointly in human and mouse: S3V2_IDEAS_mm10_ccre.sh

```
IDEAS_job_name=S3V2_IDEAS_mm10_ccre
script_dir=/storage/home/gzx103/scratch/joint_human_mouse/S3V2_IDEAS_ESMP/bin//IDEAS_2018/
working_dir=/storage/home/gzx103/scratch/joint_human_mouse/mm10/S3V2_IDEAS_outputs_mm10_CCRE/
output_dir=/storage/home/gzx103/scratch/joint_human_mouse/mm10/S3V2_IDEAS_outputs_mm10_CCRE//S3V2_IDEAS_mm10_ccre_IDEAS_output1/
binfile=/storage/home/gzx103/scratch/joint_human_mouse/mm10/S3V2_IDEAS_outputs_mm10_CCRE//windowsNoBlack.withid.IDEASbins.txt

cd $working_dir
if [ -d $output_dir ]; then rm -r $output_dir; mkdir $output_dir; else mkdir $output_dir; fi
if [ -d bin ]; then rm -r bin; cp -r $script_dir/bin ./ ; else cp -r $script_dir/bin ./ ; fi
if [ -d data ]; then rm -r data; cp -r $script_dir/data/ ./ ; else cp -r $script_dir/data ./ ; fi
time Rscript ./bin/runme.R $IDEAS_job_name'.input' $IDEAS_job_name'.parafile' $output_dir
```
