## Supplementary material for "Interspecies regulatory landscapes and elements revealed by novel joint systematic integration of human and mouse blood cell epigenomes": Scripts for IDEAS modeling of epigenetic states jointly in human and mouse: 00.prepare_input_data.sh

```
hg38_chrom_size=/Users/guanjuexiang/Documents/projects/S3V2_IDEAS_ESMP/genomesize/hg38.chrom.1_22XY.sizes
mm10_chrom_size=/Users/guanjuexiang/Documents/projects/S3V2_IDEAS_ESMP/genomesize/mm10.chrom.1_19XY.sizes
script_dir=/Users/guanjuexiang/Documents/projects/Joint_Human_Mouse_IDEAS_State/other_scripts/compare_human_mouse_state_similarity/               
working_dir=/Users/guanjuexiang/Documents/projects/analysis/0813_human_mouse_state_compare_heatmap/

###
cd $working_dir 

### prepare gene set
wget https://ftp.ebi.ac.uk/pub/databases/gencode/Gencode_human/release_39/gencode.v39.basic.annotation.gff3.gz
gunzip gencode.v39.basic.annotation.gff3.gz
cat gencode.v39.basic.annotation.gff3 | awk -F 'gene_type=' '{print $2}' | awk -F ';' '{print $1}' > gene_types.txt
cat gencode.v39.basic.annotation.gff3 | awk -F 'gene_name=' '{print $2}' | awk -F ';' '{print $1}' > gene_names.txt
paste gene_types.txt gene_names.txt gencode.v39.basic.annotation.gff3 | awk -F '\t' -v OFS='\t' '{if ($1=="protein_coding" && $5=="gene") print $3,$6,$7,$9,$2}' > hg38.gene.bed
rm gene_types.txt gene_names.txt

wget https://ftp.ebi.ac.uk/pub/databases/gencode/Gencode_mouse/release_M25/gencode.vM25.basic.annotation.gff3.gz
gunzip gencode.vM25.basic.annotation.gff3.gz
cat gencode.vM25.basic.annotation.gff3 | awk -F 'gene_type=' '{print $2}' | awk -F ';' '{print $1}' > gene_types.txt
cat gencode.vM25.basic.annotation.gff3 | awk -F 'gene_name=' '{print $2}' | awk -F ';' '{print $1}' > gene_names.txt
paste gene_types.txt gene_names.txt gencode.vM25.basic.annotation.gff3 | awk -F '\t' -v OFS='\t' '{if ($1=="protein_coding" && $5=="gene") print $3,$6,$7,$9,$2}' > mm10.gene.bed
rm gene_types.txt gene_names.txt


### prepare matched ct state bed
wget https://usevision.org/data/hg38/IDEASstates/ideasJointMay2021/S3V2_IDEAS_hg38_r3_withHg38Mm10prior.state
wget https://usevision.org/data/mm10/ideasJointMay2021/S3V2_IDEAS_mm10_r3_withHg38Mm10prior.state
wget https://raw.githubusercontent.com/guanjue/Joint_Human_Mouse_IDEAS_State/main/VBSJ_052021_outputs_para_pdf/ES/06a_S3V2_IDEAS_hg38_r3_withHg38Mm10prior.para.modified.para

### hg38
cat S3V2_IDEAS_hg38_r3_withHg38Mm10prior.state | awk -F ' ' -v OFS='\t' '{print $2,$3,$4, $5, $6,$7, $12, $16,$17, $18, $20, $30, $32,$33, $34, $42, $44, $46  }' > S3V2_IDEAS_hg38_r3_withHg38Mm10prior.state.matched_ct.bed
bedtools makewindows -g $hg38_chrom_size -w 200 > hg38.all.200bp.bins.bed
tail -n+2 S3V2_IDEAS_hg38_r3_withHg38Mm10prior.state.matched_ct.bed  > hg38.current.bed
bedtools intersect -a hg38.all.200bp.bins.bed -b hg38.current.bed -v > hg38.current.missed.bed
cat hg38.current.missed.bed | awk -F '\t' -v OFS='\t' '{print $0,0,0,0,0,0,0,0,0,0,0,0,0,0,0,0}' > hg38.current.missed.fill0s.bed
cat S3V2_IDEAS_hg38_r3_withHg38Mm10prior.state.matched_ct.bed hg38.current.missed.fill0s.bed > S3V2_IDEAS_hg38_r3_withHg38Mm10prior.state.matched_ct.full.bed
### mm10
cat S3V2_IDEAS_mm10_r3_withHg38Mm10prior.state | awk -F ' ' -v OFS='\t' '{print $2,$3,$4, $5, $6,$7, $11, $14,$15, $20, $22, $25, $26,$27, $28, $30, $31, $32  }' > S3V2_IDEAS_mm10_r3_withHg38Mm10prior.state.matched_ct.bed
bedtools makewindows -g $mm10_chrom_size -w 200 > mm10.all.200bp.bins.bed
tail -n+2 S3V2_IDEAS_mm10_r3_withHg38Mm10prior.state.matched_ct.bed  > mm10.current.bed
bedtools intersect -a mm10.all.200bp.bins.bed -b mm10.current.bed -v > mm10.current.missed.bed
cat mm10.current.missed.bed | awk -F '\t' -v OFS='\t' '{print $0,0,0,0,0,0,0,0,0,0,0,0,0,0,0,0}' > mm10.current.missed.fill0s.bed
cat S3V2_IDEAS_mm10_r3_withHg38Mm10prior.state.matched_ct.bed mm10.current.missed.fill0s.bed > S3V2_IDEAS_mm10_r3_withHg38Mm10prior.state.matched_ct.full.bed

###
head -1 mm10.gene.bed | awk -F '\t' -v OFS='\t' '{print "chr11",32283511, 32284465, "+", "Hba1"}' >> mm10.gene.bed
head -1 mm10.gene.bed | awk -F '\t' -v OFS='\t' '{print "chr7",103841637, 103843164, "-", "Hbb"}' >> mm10.gene.bed

### get hg38 mm10 shared genes
Rscript $script_dir/get_shared.genes.R hg38.gene.bed mm10.gene.bed hg38.mm10.shared.genes.txt
```
