## Supplementary material for "Interspecies regulatory landscapes and elements revealed by novel joint systematic integration of human and mouse blood cell epigenomes": Scripts for IDEAS modeling of epigenetic states jointly in human and mouse: 01.get_cor_heatmap.sh

```
tail -1 hg38.mm10.shared.genes.txt |while read g_hg38 g_mm10
do
echo $g_hg38 $g_mm10
time bash $script_dir/get_hg38_mm10.state_similarity.sh $g_hg38 $g_mm10 50000 50000 hg38.gene.bed mm10.gene.bed S3V2_IDEAS_hg38_r3_withHg38Mm10prior.state.matched_ct.full.bed S3V2_IDEAS_mm10_r3_withHg38Mm10prior.state.matched_ct.full.bed statep_rna_coe_heatmap.HM.all.ccre.withcorfilter.AVE.txt
done


time bash $script_dir/get_hg38_mm10.state_similarity.sh GATA1 Gata1 26561 49438 58670 42729 hg38.gene.bed mm10.gene.bed S3V2_IDEAS_hg38_r3_withHg38Mm10prior.state.matched_ct.full.bed S3V2_IDEAS_mm10_r3_withHg38Mm10prior.state.matched_ct.full.bed
```
