## Supplementary material for "Interspecies regulatory landscapes and elements revealed by novel joint systematic integration of human and mouse blood cell epigenomes": Scripts for IDEAS modeling of epigenetic states jointly in human and mouse: compress_seq.R

```
args = commandArgs(trailingOnly=TRUE)

input_seq_file = args[1]

output = args[2]

d = read.table(input_seq_file, header=F)


ds = c()
dn = c()

d_i_pre = d[1,1]
d_i_pre_n = 1
ds = d_i_pre

for (i in 2:dim(d)[1]){
d_i = d[i,1]
if (d_i!=d_i_pre){
ds = c(ds, d_i)
dn = c(dn, d_i_pre_n)
d_i_pre_n = 1
d_i_pre = d_i
} else{
d_i_pre_n = d_i_pre_n+1
}
#print(dim(cbind(ds[1:(length(ds)-1)],dn)))

}

dn = c(dn, d_i_pre_n)

write.table(cbind(ds,dn), output, quote=F, sep=' ', col.names=F, row.names=F)
```
