## Supplementary material for "Interspecies regulatory landscapes and elements revealed by novel joint systematic integration of human and mouse blood cell epigenomes": Scripts for IDEAS modeling of epigenetic states jointly in human and mouse: cor.mat.MaxSum.path.R

```
setwd('/Users/guanjuexiang/Documents/projects/analysis/0813_human_mouse_state_compare_heatmap')

# conda activate shiny
library(pheatmap)
library(ComplexHeatmap)
library(InteractiveComplexHeatmap)
library(shiny)
library(circlize)

getMaxSumPath <- function(mat) {
  n <- nrow(mat)
  m <- ncol(mat)

  # Initialize dp (dynamic programming) matrix with input matrix 'mat'
  dp <- mat

  # Initialize a list to store the paths
  path <- vector("list", length=n*m)
  dim(path) <- c(n, m)

  # Boundary conditions
  for(i in 2:m) {
    dp[1,i] <- dp[1,i] + dp[1,i-1]
    path[[1,i]] <- list(c(1, seq(2, i)))
  }

  for(i in 2:n) {
    dp[i,1] <- dp[i,1] + dp[i-1,1]
    path[[i,1]] <- list(c(seq(2, i), 1))
  }

  # Main loop
  for(i in 2:n){
    for(j in 2:m){
      if(dp[i-1,j] > dp[i,j-1]){
        dp[i,j] <- dp[i,j] + dp[i-1,j]
        path[[i,j]] <- c(path[[i-1,j]], list(c(i,j)))
      } else {
        dp[i,j] <- dp[i,j] + dp[i,j-1]
        path[[i,j]] <- c(path[[i,j-1]], list(c(i,j)))
      }
    }
  }

  return(list(max_sum = dp[n,m], path = path[[n,m]]))
}


# output plot folder
hg38_gene = 'GATA1'
mm10_gene = 'Gata1'
output_folder = paste0('NMF_reconstruction_', hg38_gene, '_', mm10_gene)
input_correlation_mat = paste0(hg38_gene, '.', mm10_gene, '.cor.heatmap.png.cor.mat.txt')
hg38_coord = paste0('hg38.gene.', hg38_gene, '.matched_ct.state.bed')
mm10_coord = paste0('mm10.gene.', mm10_gene, '.matched_ct.state.bed')

# read correlation matrix
d_mk_cor = as.matrix(read.table(input_correlation_mat, header=F, sep='\t', comment.char='~'))
d_mk_cor_mat = as.matrix(d_mk_cor)

# 
d_mk_cor_mat1 = d_mk_cor_mat
for (j in 1:dim(d_mk_cor_mat)[2]){
    d_mk_cor_mat1[,j] = d_mk_cor_mat[,dim(d_mk_cor_mat)[2]-j+1]
}
# set negative to 0
d_mk_cor_mat1[d_mk_cor_mat1<0] = 0


# find path
aaa = getMaxSumPath(d_mk_cor_mat1)


d_mk_cor_mat_path = matrix(0, ncol=dim(d_mk_cor_mat1)[2], nrow = dim(d_mk_cor_mat1)[1])


for (i in 1:length(aaa$path)){
    d_mk_cor_mat_path[aaa$path[[i]][1], aaa$path[[i]][2]] = 1
}


library(pheatmap)
d_mk_cor_mat_path1 = d_mk_cor_mat_path
for (j in 1:dim(d_mk_cor_mat_path)[2]){
    d_mk_cor_mat_path1[,j] = d_mk_cor_mat_path[,dim(d_mk_cor_mat_path)[2]-j+1]
}


my_colorbar=colorRampPalette(c('white', 'black'))(n = 2)
png('test_path.DP.png', width = 1000, height = 1000)
pheatmap(d_mk_cor_mat_path1, cluster_rows=F, cluster_cols=F, width = 2, height = 6, legend=F, show_rownames=F, show_colnames=F, color=my_colorbar)
dev.off()


breaksList = seq(-1, 1, by = 0.001)
my_colorbar=colorRampPalette(c('blue', 'white', 'red'))(n = length(breaksList))
png('test_path.cor.png', width = 1000, height = 1000)
pheatmap(d_mk_cor_mat, cluster_rows=F, cluster_cols=F, width = 2, height = 6, legend=F, show_rownames=F, show_colnames=F, color=my_colorbar, breaks=breaksList)
dev.off()
```
