## Supplementary material for "Interspecies regulatory landscapes and elements revealed by novel joint systematic integration of human and mouse blood cell epigenomes": Scripts for IDEAS modeling of epigenetic states jointly in human and mouse: epi_genealign.R

```
cd /homes1/gxiang/softwares/EpiAlign/Ccode/example/VISION_IDEAS_states/

### prepare gene set
wget https://ftp.ebi.ac.uk/pub/databases/gencode/Gencode_human/release_39/gencode.v39.basic.annotation.gff3.gz
gunzip gencode.v39.basic.annotation.gff3.gz
cat gencode.v39.basic.annotation.gff3 | awk -F 'gene_type=' '{print $2}' | awk -F ';' '{print $1}' > gene_types.txt
cat gencode.v39.basic.annotation.gff3 | awk -F 'gene_name=' '{print $2}' | awk -F ';' '{print $1}' > gene_names.txt
paste gene_types.txt gene_names.txt gencode.v39.basic.annotation.gff3 | awk -F '\t' -v OFS='\t' '{if ($1=="protein_coding" && $5=="gene") print $3,$6,$7,$9,$2}' > hg38.gene.bed
rm gene_types.txt gene_names.txt

wget https://ftp.ebi.ac.uk/pub/databases/gencode/Gencode_mouse/release_M25/gencode.vM25.basic.annotation.gff3.gz
gunzip gencode.vM25.basic.annotation.gff3.gz
cat gencode.vM25.basic.annotation.gff3 | awk -F 'gene_type=' '{print $2}' | awk -F ';' '{print $1}' > gene_types.txt
cat gencode.vM25.basic.annotation.gff3 | awk -F 'gene_name=' '{print $2}' | awk -F ';' '{print $1}' > gene_names.txt
paste gene_types.txt gene_names.txt gencode.vM25.basic.annotation.gff3 | awk -F '\t' -v OFS='\t' '{if ($1=="protein_coding" && $5=="gene") print $3,$6,$7,$9,$2}' > mm10.gene.bed
rm gene_types.txt gene_names.txt


### prepare matched ct state bed
cat S3V2_IDEAS_hg38_r3_withHg38Mm10prior.state | awk -F ' ' -v OFS='\t' '{print $2,$3,$4, $5, $6,$7, $12, $16,$17, $18, $20, $32,$33, $34, $42, $44, $46  }' > S3V2_IDEAS_hg38_r3_withHg38Mm10prior.state.matched_ct.bed
cat S3V2_IDEAS_mm10_r3_withHg38Mm10prior.state | awk -F ' ' -v OFS='\t' '{print $2,$3,$4, $5, $6,$7, $11, $14,$15, $20, $22, $26,$27, $28, $30, $31, $32  }' > S3V2_IDEAS_mm10_r3_withHg38Mm10prior.state.matched_ct.bed


hg38_gene=$1
mm10_gene=$2
hg38_gene_set=$3
mm10_gene_set=$4
hg38_state_set=$5
mm10_state_set=$6
hg38_gene_exp_win=$7
mm10_gene_exp_win=$8


hg38_gene='GATA1'
mm10_gene='Gata1'
hg38_gene_set='hg38.gene.bed'
mm10_gene_set='mm10.gene.bed'
hg38_state_set='S3V2_IDEAS_hg38_r3_withHg38Mm10prior.state.matched_ct.bed'
mm10_state_set='S3V2_IDEAS_mm10_r3_withHg38Mm10prior.state.matched_ct.bed'
hg38_gene_exp_win=50000
mm10_gene_exp_win=50000

hg38_gene='HBA1'
mm10_gene='Hba-a1'
hg38_gene_set='hg38.gene.bed'
mm10_gene_set='mm10.gene.bed'
hg38_state_set='S3V2_IDEAS_hg38_r3_withHg38Mm10prior.state.matched_ct.bed'
mm10_state_set='S3V2_IDEAS_mm10_r3_withHg38Mm10prior.state.matched_ct.bed'
hg38_gene_exp_win=100000
mm10_gene_exp_win=100000

cd 
### get query gene
cat $hg38_gene_set | awk -F '\t' -v OFS='\t' -v hg38_gene=$hg38_gene '{if (toupper($5)==toupper(hg38_gene)) print $1,$2,$3,$4,$5}' | awk -F '\t' -v OFS='\t' -v hg38_gene_exp_win=$hg38_gene_exp_win '{if ($4=="+") print $1,$2-hg38_gene_exp_win,$2+hg38_gene_exp_win,$4,$5; else print $1,$3-hg38_gene_exp_win,$3+hg38_gene_exp_win,$4,$5}' | awk -F '\t' -v OFS='\t' '{if ($2<0) print $1,0,$3,$4,$5; else print $0}' > 'hg38.gene.'$hg38_gene'.bed'
cat $mm10_gene_set | awk -F '\t' -v OFS='\t' -v mm10_gene=$mm10_gene '{if (toupper($5)==toupper(mm10_gene)) print $1,$2,$3,$4,$5}' | awk -F '\t' -v OFS='\t' -v mm10_gene_exp_win=$mm10_gene_exp_win '{if ($4=="+") print $1,$2-mm10_gene_exp_win,$2+mm10_gene_exp_win,$4,$5; else print $1,$3-mm10_gene_exp_win,$3+mm10_gene_exp_win,$4,$5}' | awk -F '\t' -v OFS='\t' '{if ($2<0) print $1,0,$3,$4,$5; else print $0}' > 'mm10.gene.'$mm10_gene'.bed'

### intersect state bed
bedtools intersect -a $hg38_state_set -b 'hg38.gene.'$hg38_gene'.bed' -wa -u > 'hg38.gene.'$hg38_gene'.matched_ct.state.bed'
bedtools intersect -a $mm10_state_set -b 'mm10.gene.'$mm10_gene'.bed' -wa -u > 'mm10.gene.'$mm10_gene'.matched_ct.state.bed'

### get state similarity matrix
cat 'hg38.gene.'$hg38_gene'.bed' 'mm10.gene.'$mm10_gene'.bed'
Rscript get_hg38_mm10.cor.heatmap.R 'hg38.gene.'$hg38_gene'.matched_ct.state.bed' 'mm10.gene.'$mm10_gene'.matched_ct.state.bed' $hg38_gene'.cor.heatmap.png' 06a_S3V2_IDEAS_hg38_r3_withHg38Mm10prior.para.modified.para 8 


R

args = commandArgs(trailingOnly=TRUE)

hg38_gene_state = args[1]
mm10_gene_state = args[2]
hg38_gene = args[3]
mm10_gene = args[4]
output_file = args[5]
state_file = args[6]
state_feature_n = as.numeric(args[7])


hg38_gene_state = 'hg38.gene.GATA1.matched_ct.state.bed'
mm10_gene_state = 'mm10.gene.Gata1.matched_ct.state.bed'
state_file = '06a_S3V2_IDEAS_hg38_r3_withHg38Mm10prior.para.modified.para'
state_feature_n = 8
output_file = 'GATA1.cor.heatmap.png'


### read state signal mat
state = read.table(state_file, header=T)
state_mat_od = apply(state[,2:(state_feature_n+1)],2,function(x) x/state[,1])

### state gene assignment
d1 = read.table(hg38_gene_state, header=F, sep='\t', comment.char='~')
d2 = read.table(mm10_gene_state, header=F, sep='\t', comment.char='~')
#d1_gene = read.table(hg38_gene)
#d2_gene = read.table(mm10_gene)
#if (d1[1,5]=='-'){d1 = d1[dim(d1)[1]:1,]}
#if (d2[1,5]=='-'){d2 = d2[dim(d2)[1]:1,]}

### read state coe scores
state_coe = c(0, -1, 0.5, -0.5, 0.6, 0.1, 1, -3, 0.8, 0.3, 0.2, -2, 1, 0.8,
	2, 2.1, 1.3, 1.1, -1.8, 1.8, 1.6, 1.5, 0.8, 3, 0.9)

### get state
d1s = d1[,-c(1:3)]
d2s = d2[,-c(1:3)]

### state to signal matrix
d1s_sigmat = matrix(0, nrow=dim(d1s)[1], ncol=dim(d1s)[2]*dim(state_mat_od)[2])
d2s_sigmat = matrix(0, nrow=dim(d2s)[1], ncol=dim(d2s)[2]*dim(state_mat_od)[2])
###
for (i in 1:dim(d1s)[1]){
	sig_i = c(state_mat_od[unlist(d1s[i,]+1),])
	d1s_sigmat[i,] = as.numeric(state_mat_od[unlist(d1s[i,]+1),])
}
###
for (i in 1:dim(d2s)[1]){
	sig_i = c(state_mat_od[unlist(d2s[i,]+1),])
	d2s_sigmat[i,] = as.numeric(state_mat_od[unlist(d2s[i,]+1),])
}
###
rownames(d1s_sigmat) = paste('H',1:dim(d1s_sigmat)[1], sep=':')
rownames(d2s_sigmat) = paste('M',1:dim(d2s_sigmat)[1], sep=':')

###
d1s_sigmat_rowMax = log(apply(d1s_sigmat,1,max)+1)
d1s_sigmat_rowMax = d1s_sigmat_rowMax/max(d1s_sigmat_rowMax)
d2s_sigmat_rowMax = log(apply(d2s_sigmat,1,max)+1)
d2s_sigmat_rowMax = d2s_sigmat_rowMax/max(d2s_sigmat_rowMax)

### get cor matrix
d12_cor_mat = as.matrix(cor(t(d2s_sigmat), t(d1s_sigmat)))
d12_cor_mat[is.na(d12_cor_mat)] = 0
d12_cor_mat_adj = t(apply(d12_cor_mat,1,function(x) x*(d1s_sigmat_rowMax)))
d12_cor_mat_adj = apply(d12_cor_mat_adj,2,function(x) x*(d2s_sigmat_rowMax))


library(pheatmap)
breaksList = seq(-1, 1, by = 0.001)
my_colorbar=colorRampPalette(c('blue', 'white', 'red'))(n = length(breaksList))
png(output_file, height=800, width=800)
colnames(d12_cor_mat_adj) = NULL
rownames(d12_cor_mat_adj) = NULL
pheatmap(d12_cor_mat_adj, cluster_rows=F, cluster_cols=F, color=my_colorbar, breaks = breaksList)
dev.off()


### state to signal matrix
d1s_sigmat1 = matrix(0, nrow=dim(d1s)[1], ncol=dim(d1s)[2])
d2s_sigmat1 = matrix(0, nrow=dim(d2s)[1], ncol=dim(d2s)[2])
###
for (i in 1:dim(d1s)[1]){
	d1s_sigmat1[i,] = as.numeric(state_coe[unlist(d1s[i,]+1)])
}
###
for (i in 1:dim(d2s)[1]){
	d2s_sigmat1[i,] = as.numeric(state_coe[unlist(d2s[i,]+1)])
}

d12_cor_mat1 = as.matrix(cor(t(d2s_sigmat1), t(d1s_sigmat1)))
d12_cor_mat1[is.na(d12_cor_mat1)] = 0
d12_cor_mat_adj1 = t(apply(d12_cor_mat1,1,function(x) x*d1s_sigmat_rowMax))
d12_cor_mat_adj1 = apply(d12_cor_mat_adj1,2,function(x) x*d2s_sigmat_rowMax)

library(pheatmap)
breaksList = seq(-1, 1, by = 0.001)
my_colorbar=colorRampPalette(c('blue', 'white', 'red'))(n = length(breaksList))
png('d12_cor1_mat.png', height=800, width=800)
colnames(d12_cor_mat_adj1) = NULL
rownames(d12_cor_mat_adj1) = NULL
pheatmap(d12_cor_mat_adj1, cluster_rows=F, cluster_cols=F, color=my_colorbar, breaks = breaksList)
dev.off()


### get ATAC max
ATAC_col_id = which(colnames(state_mat_od)=='ATAC')
d1s_sigmat_atac = c()
for (i in 1:dim(d1s)[2]){
	d1s_sigmat_atac = cbind(d1s_sigmat_atac,d1s_sigmat[,(ATAC_col_id)+8*(i-1)])
}
d1s_sigmat_atac_rowMax = apply(d1s_sigmat_atac,1,max)
d1s_sigmat_atac_rowMax = d1s_sigmat_atac_rowMax/max(d1s_sigmat_atac_rowMax)

d2s_sigmat_atac = c()
for (i in 1:dim(d1s)[2]){
	d2s_sigmat_atac = cbind(d2s_sigmat_atac,d2s_sigmat[,(ATAC_col_id)+8*(i-1)])
}
d2s_sigmat_atac_rowMax = apply(d2s_sigmat_atac,1,max)
d2s_sigmat_atac_rowMax = d2s_sigmat_atac_rowMax/max(d2s_sigmat_atac_rowMax)


### get average mat
d1s_sigmat_ave = d1s_sigmat[,1:8]
for (i in 2:dim(d1s)[2]){
	d1s_sigmat_ave = d1s_sigmat_ave+d1s_sigmat[,(1:8)+8*(i-1)]
}
d1s_sigmat_ave = d1s_sigmat_ave/dim(d1s)[2]

d2s_sigmat_ave = d2s_sigmat[,1:8]
for (i in 2:dim(d2s)[2]){
	d2s_sigmat_ave = d2s_sigmat_ave+d2s_sigmat[,(1:8)+8*(i-1)]
}
d2s_sigmat_ave = d2s_sigmat_ave/dim(d2s)[2]

### get denoised filter
#avefeature = rowMeans(d1s_sigmat_ave)
#avefeature2 = rowMeans(d2s_sigmat_ave)
avefeature = apply(d1s_sigmat_ave,1,max)
avefeature2 = apply(d2s_sigmat_ave,1,max)


### state to signal matrix
d1s_sigmat1 = matrix(0, nrow=dim(d1s)[1], ncol=dim(d1s)[2])
d2s_sigmat1 = matrix(0, nrow=dim(d2s)[1], ncol=dim(d2s)[2])
###
for (i in 1:dim(d1s)[1]){
	d1s_sigmat1[i,] = as.numeric(state_coe[unlist(d1s[i,]+1)])
}
###
for (i in 1:dim(d2s)[1]){
	d2s_sigmat1[i,] = as.numeric(state_coe[unlist(d2s[i,]+1)])
}


avefeature1 = apply(d1s_sigmat1,1,max)
avefeature21 = apply(d2s_sigmat1,1,max)

d12_cor_mat1 = as.matrix(cor(t(d2s_sigmat1), t(d1s_sigmat1)))
d12_cor_mat1[is.na(d12_cor_mat1)] = 0
d12_cor_mat_adj1 = t(apply(d12_cor_mat1,1,function(x) x*avefeature1))
d12_cor_mat_adj1 = apply(d12_cor_mat_adj1,2,function(x) x*avefeature21)

library(pheatmap)
breaksList = seq(-1, 1, by = 0.001)
my_colorbar=colorRampPalette(c('blue', 'white', 'red'))(n = length(breaksList))
png('d12_cor1_mat.png', height=800, width=800)
colnames(d12_cor_mat_adj1) = NULL
rownames(d12_cor_mat_adj1) = NULL
pheatmap(d12_cor_mat_adj1, cluster_rows=F, cluster_cols=F, color=my_colorbar, breaks = breaksList)
dev.off()


d11_cor_mat1 = as.matrix(cor(t(d1s_sigmat1), t(d1s_sigmat1)))
d11_cor_mat1[is.na(d11_cor_mat1)] = 0
d11_cor_mat_adj1 = t(apply(d11_cor_mat1,1,function(x) x*avefeature1))
d11_cor_mat_adj1 = apply(d11_cor_mat_adj1,2,function(x) x*avefeature1)

breaksList = seq(-1, 1, by = 0.001)
my_colorbar=colorRampPalette(c('blue', 'white', 'red'))(n = length(breaksList))
png('d11_cor1_mat.png', height=800, width=800)
colnames(d11_cor_mat_adj1) = NULL
rownames(d11_cor_mat_adj1) = NULL
pheatmap(d11_cor_mat_adj1, cluster_rows=F, cluster_cols=F, color=my_colorbar, breaks = breaksList)
dev.off()


d22_cor_mat1 = as.matrix(cor(t(d2s_sigmat1), t(d2s_sigmat1)))
d22_cor_mat1[is.na(d22_cor_mat1)] = 0
d22_cor_mat_adj1 = t(apply(d22_cor_mat1,1,function(x) x*avefeature21))
d22_cor_mat_adj1 = apply(d22_cor_mat_adj1,2,function(x) x*avefeature21)

breaksList = seq(-1, 1, by = 0.001)
my_colorbar=colorRampPalette(c('blue', 'white', 'red'))(n = length(breaksList))
png('d22_cor1_mat.png', height=800, width=800)
colnames(d22_cor_mat_adj1) = NULL
rownames(d22_cor_mat_adj1) = NULL
pheatmap(d22_cor_mat_adj1, cluster_rows=F, cluster_cols=F, color=my_colorbar, breaks = breaksList)
dev.off()
```
