## Supplementary material for "Interspecies regulatory landscapes and elements revealed by novel joint systematic integration of human and mouse blood cell epigenomes": Scripts for IDEAS modeling of epigenetic states jointly in human and mouse: get_cCRE_state_sig_mat.hg38.R

```
args = commandArgs(trailingOnly=TRUE)

Beta_coe_mat_file = args[1]
cCRE_state_coverage_file_start = args[2]
state_max = as.numeric(args[3])
output_filename_start = args[4]
ct_list_human = args[-c(1:4)]

Beta_coe_mat_file = 'statep_rna_coe_heatmap.human.all.ccre.withcorfilter.AVE.txt'
cCRE_state_coverage_file_start = 'S3V2_IDEAS_hg38_ccre2.cCRE.M.notall0.rmallNEU.withid.S'
state_max = 24
output_filename_start = 'S3V2_IDEAS_hg38_ccre2.cCRE.M.notall0.rmallNEU.withid.coe_mat'
ct_list_human = c('AVE', 'B_B15_50', 'B_NC14_42', 'CD34_E_rep1', 'CD34_E_rep2', 'CLP_100266', 'CLP_100267', 'CMP_100246', 'CMP_100247', 'EOS_S006XEH2', 'EOS_S00BKK', 'ERY_S002R5', 'ERY_S002S3', 'GMP_100256', 'GMP_100257', 'HSC_100258', 'HSC_100259', 'HUDEP1_rep1', 'HUDEP1_rep2', 'HUDEP2_rep1', 'HUDEP2_rep2', 'K562_rep1', 'K562_rep2', 'LMPP_100268', 'LMPP_100269', 'MEP_Donor2596', 'MEP_Donor7256', 'MK_S004BTH2', 'MK_S00VHKH1', 'MONc_C0011IH1', 'MONc_C001UYH2', 'MONp_Prim_mon_C', 'MONp_Prim_mon_F', 'MPP_100272', 'MPP_100273', 'NEU_C0011IH2', 'NEU_C001UYH1', 'NK_S005YG', 'NK_S01E4WH0', 'T_CD4_S008H1', 'T_CD4_S009W4', 'T_CD8_C0066PH1', 'T_CD8_S00C2FH1')

State_sig_mat_file = '/Users/guanjuexiang/Documents/projects/Joint_Human_Mouse_IDEAS_State/VBSJ_052021_outputs_para_pdf/ES/06a_S3V2_IDEAS_hg38_r3_withHg38Mm10prior.para.modified.para'
output_file = 'S3V2_IDEAS_hg38_ccre2.cCRE.M.notall0.rmallNEU.withid.statesigmat.txt'

###################
###### human ######
print('read Beta coefficient mat')
coe = read.table(Beta_coe_mat_file, header=T)
state_sig_mat = read.table(State_sig_mat_file, header=T)
state_sig_mat = state_sig_mat[,2:9]/state_sig_mat[,1]


print('read state count')
Hs = read.table(paste0(cCRE_state_coverage_file_start, '0.mat.txt'), header=F)

print('normalize dist')
HsP0 = as.matrix(Hs[,-c(1:4)])
HsD0 = as.matrix(Hs[,-c(1:4)])

print('get state sig mat')
Hstate_sig = matrix(0, nrow=dim(HsD0)[1], ncol=dim(HsD0)[2]*8)
for (i in 1:dim(HsD0)[2]){
	print(i)
	Hstate_sig[which(HsD0[,i]!=0),(1:8)+(i-1)*8] = as.matrix(state_sig_mat[(HsD0[,i]!=0)*1,])*HsD0[HsD0[,i]!=0,i]
}

print('get colnames')
Hstate_sig_colnames = c()
for (ct_i in ct_list_human){
	for (state_j in colnames(state_sig_mat)){
		Hstate_sig_colnames = c(Hstate_sig_colnames, paste(ct_i, state_j, sep=':'))
	}
}
colnames(Hstate_sig) = Hstate_sig_colnames

Hstate_sig = cbind(Hs[,c(1:4)],Hstate_sig)
colnames(Hstate_sig)[1:4] = c('chr','start','end','id')
write.table(Hstate_sig, output_file,quote=F, sep='\t', col.names=T, row.names=F)
```
