## Supplementary material for "Interspecies regulatory landscapes and elements revealed by novel joint systematic integration of human and mouse blood cell epigenomes": Scripts for IDEAS modeling of epigenetic states jointly in human and mouse: get_hg38.ave.sh

```
#!/bin/bash

#SBATCH --mem=2G
#SBATCH --cpus-per-task=1

source ~/.bashrc

cd /homes1/gxiang/softwares/EpiAlign/Ccode/example/VISION_IDEAS_states
state_bed='S3V2_IDEAS_hg38_r3_withHg38Mm10prior.state.AVE.bed'
while read chr start end st gn
do
echo $st
echo $chr $start $end $st $gn | awk -F ' ' -v OFS='\t' '{print $1,$2,$3,$4,$5}' > 'genes/hg38.gene.tss.exp.'$gn'.'$state_bed'.bed'
bedtools intersect -a $state_bed -b 'genes/hg38.gene.tss.exp.'$gn'.'$state_bed'.bed' -wa -u > 'genes/hg38.gene.tss.exp.'$gn'.'$state_bed'.state.bed'
if [[ $st == "+" ]]
then
cut -f4 'genes/hg38.gene.tss.exp.'$gn'.'$state_bed'.state.bed' > 'genes/hg38.gene.tss.exp.'$gn'.'$state_bed'.state.bed.seq'
fi
if [[ $st == "-" ]]
then
cut -f4 'genes/hg38.gene.tss.exp.'$gn'.'$state_bed'.state.bed' > 'genes/hg38.gene.tss.exp.'$gn'.'$state_bed'.state.bed.r.seq'
Rscript reverse_seq.R 'genes/hg38.gene.tss.exp.'$gn'.'$state_bed'.state.bed.r.seq' 'genes/hg38.gene.tss.exp.'$gn'.'$state_bed'.state.bed.seq'
rm 'genes/hg38.gene.tss.exp.'$gn'.'$state_bed'.state.bed.r.seq'
fi
Rscript compress_seq.R 'genes/hg38.gene.tss.exp.'$gn'.'$state_bed'.state.bed.seq' 'genes/hg38.gene.tss.exp.'$gn'.'$state_bed'.state.bed.Sseq'
rm 'genes/hg38.gene.tss.exp.'$gn'.'$state_bed'.state.bed'
#mv 'genes/hg38.gene.tss.exp.'$gn'.'$state_bed'.state.bed.seq' genes/hg38_seq
#mv 'genes/hg38.gene.tss.exp.'$gn'.'$state_bed'.state.bed.Sseq' genes/hg38_Sseq
done < genes/hg38.gene.tss.exp.bed
```
