## Supplementary material for "Interspecies regulatory landscapes and elements revealed by novel joint systematic integration of human and mouse blood cell epigenomes": Scripts for IDEAS modeling of epigenetic states jointly in human and mouse: get_hg38_mm10.cor.heatmap.coe.pre.R

```
args = commandArgs(trailingOnly=TRUE)

hg38_gene_state = args[1]
mm10_gene_state = args[2]
output_file = args[3]
state_file = args[4]
state_feature_n = as.numeric(args[5])
coe_file = args[6]

#hg38_gene_state = 'hg38.gene.GATA1.matched_ct.state.bed'
#mm10_gene_state = 'mm10.gene.Gata1.matched_ct.state.bed'
#state_file = '06a_S3V2_IDEAS_hg38_r3_withHg38Mm10prior.para.modified.para'
#tate_feature_n = 8
#output_file = 'GATA1.cor.heatmap.png'
#coe_file = 'statep_rna_coe_heatmap.HM.all.ccre.withcorfilter.AVE.txt'

### read state signal mat
state = read.table(state_file, header=T)
state_mat_od = apply(state[,2:(state_feature_n+1)],2,function(x) x/state[,1])

### state gene assignment
d1 = read.table(hg38_gene_state, header=F, sep='\t', comment.char='~')
d2 = read.table(mm10_gene_state, header=F, sep='\t', comment.char='~')

### read state coe scores
coe_mat = read.table(coe_file, header=T)
scale_ratio = max(coe_mat[,2])/max(coe_mat[,1])
### get state coe
state_coe = (coe_mat[,1]*scale_ratio + coe_mat[,2])
print(state_coe)
#state_coe = c(0, -1, 0.5, -0.5, 0.6, 0.1, 1, -3, 0.8, 0.3, 0.2, -2, 1, 0.8, 2, 2.1, 1.3, 1.1, -1.8, 1.8, 1.6, 1.5, 0.8, 3, 0.9)


### get state
d1s = d1[,-c(1:3)]
d2s = d2[,-c(1:3)]

### state to signal matrix
d1s_sigmat = matrix(0, nrow=dim(d1s)[1], ncol=dim(d1s)[2])
d2s_sigmat = matrix(0, nrow=dim(d2s)[1], ncol=dim(d2s)[2])
###
for (i in 1:dim(d1s)[1]){
	d1s_sigmat[i,] = as.numeric(state_coe[unlist(d1s[i,]+1)])
}
###
for (i in 1:dim(d2s)[1]){
	d2s_sigmat[i,] = as.numeric(state_coe[unlist(d2s[i,]+1)])
}

###
rownames(d1s_sigmat) = paste('H',1:dim(d1s_sigmat)[1], sep=':')
rownames(d2s_sigmat) = paste('M',1:dim(d2s_sigmat)[1], sep=':')

###
d1s_sigmat_rowMax = log(apply(d1s_sigmat,1,max)+1)
d1s_sigmat_rowMax = d1s_sigmat_rowMax/max(d1s_sigmat_rowMax)
d2s_sigmat_rowMax = log(apply(d2s_sigmat,1,max)+1)
d2s_sigmat_rowMax = d2s_sigmat_rowMax/max(d2s_sigmat_rowMax)

### get cor matrix
#d12_cor_mat = as.matrix(cor(t(d2s_sigmat), t(d1s_sigmat)))
###d12_cor_mat = as.matrix(cor(t(d2s_sigmat+matrix(rnorm(dim(d2s_sigmat)[1]*dim(d2s_sigmat)[2], sd=0.001), nrow=dim(d2s_sigmat)[1], ncol=dim(d2s_sigmat)[2]) ), t(d1s_sigmat+matrix(rnorm(dim(d1s_sigmat)[1]*dim(d1s_sigmat)[2], sd=0.001), nrow=dim(d1s_sigmat)[1], ncol=dim(d1s_sigmat)[2]) )))
###d12_cor_mat[is.na(d12_cor_mat)] = 0


d12s_sigmat = matrix(0, nrow = dim(d2s_sigmat)[1], ncol=dim(d1s_sigmat)[1])
for (i in 1:dim(d2s_sigmat)[1]){
	for (j in 1:dim(d1s_sigmat)[1]){
		d12s_sigmat[i,j] = sqrt(sum((d2s_sigmat[i,]/2 + d1s_sigmat[j,]/2)^2))
	}
}


d12_cor_mat = matrix(0, nrow = dim(d2s_sigmat)[1], ncol=dim(d1s_sigmat)[1])
for (i in 1:dim(d2s_sigmat)[1]){
	for (j in 1:dim(d1s_sigmat)[1]){
		d12_cor_mat[i,j] = sqrt(sum((d2s_sigmat[i,] - d1s_sigmat[j,])^2)) #/ d12s_sigmat[i,j]+0.01)
	}
}
d12_cor_mat0 = max(d12_cor_mat) - d12_cor_mat

d12_cor_mat = d12_cor_mat0 * (d12s_sigmat+0.1)

#d12_cor_mat_adj = t(apply(d12_cor_mat,1,function(x) x*(d1s_sigmat_rowMax)))
#d12_cor_mat_adj = apply(d12_cor_mat_adj,2,function(x) x*(d2s_sigmat_rowMax))

d12_cor_mat_adj = t(apply(d12_cor_mat,1,function(x) x*1))
d12_cor_mat_adj = apply(d12_cor_mat_adj,2,function(x) x*1)


#library(pheatmap)
breaksList = seq(-1, 1, by = 0.001)
#my_colorbar=colorRampPalette(c('blue', 'white', 'red'))(n = length(breaksList))
#png(output_file, height=800, width=800)
#colnames(d12_cor_mat_adj) = NULL
#rownames(d12_cor_mat_adj) = NULL
#pheatmap(d12_cor_mat_adj, cluster_rows=F, cluster_cols=F, color=my_colorbar, breaks = breaksList)
#dev.off()

library(pheatmap)
my_colorbar=colorRampPalette(c('white', 'red'))(n = length(breaksList))
png(output_file, height=800, width=800)
colnames(d12_cor_mat_adj) = NULL
rownames(d12_cor_mat_adj) = NULL
pheatmap(d12_cor_mat_adj, cluster_rows=F, cluster_cols=F, color=my_colorbar)
dev.off()
```
