## Supplementary material for "Interspecies regulatory landscapes and elements revealed by novel joint systematic integration of human and mouse blood cell epigenomes": Scripts for IDEAS modeling of epigenetic states jointly in human and mouse: get_hg38_mm10.cor.heatmap.R

```
#library(DescTools)
args = commandArgs(trailingOnly=TRUE)

hg38_gene_state = args[1]
mm10_gene_state = args[2]
output_file = args[3]
state_file = args[4]
state_feature_n = as.numeric(args[5])


#hg38_gene_state = 'hg38.gene.GATA1.matched_ct.state.bed'
#mm10_gene_state = 'mm10.gene.Gata1.matched_ct.state.bed'
#state_file = '06a_S3V2_IDEAS_hg38_r3_withHg38Mm10prior.para.modified.para'
#state_feature_n = 8
#output_file = 'GATA1.cor.heatmap.png'


### read state signal mat
state = read.table(state_file, header=T)
state_mat_od = apply(state[,2:(state_feature_n+1)],2,function(x) x/state[,1])

### state gene assignment
d1 = read.table(hg38_gene_state, header=F, sep='\t', comment.char='~')
d2 = read.table(mm10_gene_state, header=F, sep='\t', comment.char='~')

### read state coe scores
state_coe = c(0, -1, 0.5, -0.5, 0.6, 0.1, 1, -3, 0.8, 0.3, 0.2, -2, 1, 0.8,
	2, 2.1, 1.3, 1.1, -1.8, 1.8, 1.6, 1.5, 0.8, 3, 0.9)

### get state
d1s = d1[,-c(1:3)]
d2s = d2[,-c(1:3)]

### state to signal matrix
d1s_sigmat = matrix(0, nrow=dim(d1s)[1], ncol=dim(d1s)[2]*dim(state_mat_od)[2])
d2s_sigmat = matrix(0, nrow=dim(d2s)[1], ncol=dim(d2s)[2]*dim(state_mat_od)[2])
###
for (i in 1:dim(d1s)[1]){
	sig_i = c(state_mat_od[unlist(d1s[i,]+1),])
	d1s_sigmat[i,] = as.numeric(state_mat_od[unlist(d1s[i,]+1),])
}
###
for (i in 1:dim(d2s)[1]){
	sig_i = c(state_mat_od[unlist(d2s[i,]+1),])
	d2s_sigmat[i,] = as.numeric(state_mat_od[unlist(d2s[i,]+1),])
}
###
rownames(d1s_sigmat) = paste('H',1:dim(d1s_sigmat)[1], sep=':')
rownames(d2s_sigmat) = paste('M',1:dim(d2s_sigmat)[1], sep=':')

###
d1s_sigmat_rowMax = log(apply(d1s_sigmat,1,max)+1)
d1s_sigmat_rowMax = d1s_sigmat_rowMax/max(d1s_sigmat_rowMax)
d2s_sigmat_rowMax = log(apply(d2s_sigmat,1,max)+1)
d2s_sigmat_rowMax = d2s_sigmat_rowMax/max(d2s_sigmat_rowMax)

### get cor matrix
#d12_cor_mat = as.matrix(cor(t(d2s_sigmat), t(d1s_sigmat)))
d12_cor_mat = as.matrix(cor(t(d2s_sigmat+matrix(rnorm(dim(d2s_sigmat)[1]*dim(d2s_sigmat)[2], sd=0.2), nrow=dim(d2s_sigmat)[1], ncol=dim(d2s_sigmat)[2]) ), t(d1s_sigmat+matrix(rnorm(dim(d1s_sigmat)[1]*dim(d1s_sigmat)[2], sd=0.2), nrow=dim(d1s_sigmat)[1], ncol=dim(d1s_sigmat)[2]) )))

d12_cor_mat[is.na(d12_cor_mat)] = 0
#d12_cor_mat_adj = t(apply(d12_cor_mat,1,function(x) x*(d1s_sigmat_rowMax)))
#d12_cor_mat_adj = apply(d12_cor_mat_adj,2,function(x) x*(d2s_sigmat_rowMax))

d12_cor_mat_adj = t(apply(d12_cor_mat,1,function(x) x*1))
d12_cor_mat_adj = apply(d12_cor_mat_adj,2,function(x) x*1)


library(pheatmap)
breaksList = seq(-1, 1, by = 0.001)
my_colorbar=colorRampPalette(c('blue', 'white', 'red'))(n = length(breaksList))
png(output_file, height=800, width=800)
colnames(d12_cor_mat_adj) = NULL
rownames(d12_cor_mat_adj) = NULL
pheatmap(d12_cor_mat_adj, cluster_rows=F, cluster_cols=F, color=my_colorbar, breaks = breaksList)
dev.off()

#print(paste('Entropy:', Entropy(d12_cor_mat_adj)))

get_cor_score = function(x){
x = as.numeric(x)
#x[x<0] = 0
#xp = -log10(pnorm(max(x), mean = mean(x), sd = sd(x), lower.tail = F))
#xp = -ppois(max(x), mean(x[x>0]), lower.tail = F, log.p = T)

#x1 = x[xp<0.05]
#x2 = x[xp>=0.05]
#x1 = x[x>quantile(x, 0.99)]
#x2 = x[x<=quantile(x, 0.99)]
#p = t.test(max(x),x, alternative='greater')$p.value
return(mean(x[x>quantile(x, 0.9)]))
}

png(paste(output_file, '.bar.1.png', sep=''), height=100, width=800)
par(mar = c(0, 0, 0, 0))
s1 = apply(d12_cor_mat_adj,2,function(x) get_cor_score(x))
s1logp = -log10(pnorm(s1, mean(s1), sd(s1), lower.tail = F))
barplot(rbind(s1logp), axes = F)
dev.off()

png(paste(output_file, '.bar.2.png', sep=''), height=100, width=800)
par(mar = c(0, 0, 0, 0))
s1 = apply(d12_cor_mat_adj,1,function(x) get_cor_score(x))
s1logp = -log10(pnorm(s1, mean(s1), sd(s1), lower.tail = F))
barplot(rbind(s1logp), axes = F)
dev.off()
```
