## Supplementary material for "Interspecies regulatory landscapes and elements revealed by novel joint systematic integration of human and mouse blood cell epigenomes": Scripts for IDEAS modeling of epigenetic states jointly in human and mouse: get_hg38_mm10.RNA_similarity.sh

```
cd /Users/universe/Documents/2020_BG/compare_genes_RNAseq

wget http://usevision.org/data/hg38/RNA/Oct2021/tpmFeb21_v3.tab
wget https://ftp.ebi.ac.uk/pub/databases/gencode/Gencode_human/release_39/gencode.v39.basic.annotation.gff3.gz
gunzip gencode.v39.basic.annotation.gff3.gz
cat gencode.v39.basic.annotation.gff3 | awk -F 'gene_type=' '{print $2}' | awk -F ';' '{print $1}' > hg38.gene_types.txt
cat gencode.v39.basic.annotation.gff3 | awk -F 'gene_name=' '{print $2}' | awk -F ';' '{print $1}' > hg38.gene_names.txt
cat gencode.v39.basic.annotation.gff3 | awk -F 'ID=' '{print $2}' | awk -F ';' '{print $1}' > hg38.gene_ENS_id.txt
paste hg38.gene_types.txt hg38.gene_names.txt hg38.gene_ENS_id.txt gencode.v39.basic.annotation.gff3 | awk -F '\t' -v OFS='\t' '{if ($1=="protein_coding" && $6=="gene") print $2,$3}' > hg38.gene2id.txt

wget http://usevision.org/data/mm10/rnaTPM_withcoordinates.txt
wget http://usevision.org/data/mm10/gencode.vM4-tRNAs-ERCC.gtf
wget https://ftp.ebi.ac.uk/pub/databases/gencode/Gencode_mouse/release_M25/gencode.vM25.basic.annotation.gff3.gz
gunzip gencode.vM25.basic.annotation.gff3.gz

cat gencode.vM25.basic.annotation.gff3 | awk -F 'gene_type=' '{print $2}' | awk -F ';' '{print $1}' > mm10.gene_types.txt
cat gencode.vM25.basic.annotation.gff3 | awk -F 'gene_name=' '{print $2}' | awk -F ';' '{print $1}' > mm10.gene_names.txt
cat gencode.vM25.basic.annotation.gff3 | awk -F 'ID=' '{print $2}' | awk -F ';' '{print $1}' > mm10.gene_ENS_id.txt
paste mm10.gene_types.txt mm10.gene_names.txt mm10.gene_ENS_id.txt gencode.vM25.basic.annotation.gff3 | awk -F '\t' -v OFS='\t' '{if ($1=="protein_coding" && $6=="gene") print $2,$3}' > mm10.gene2id.txt


cat gencode.vM4-tRNAs-ERCC.gtf | awk -F 'gene_type "' '{print $2}' | awk -F '"' '{print $1}' > mm10.gene_types.txt
cat gencode.vM4-tRNAs-ERCC.gtf | awk -F 'gene_name "' '{print $2}' | awk -F '"' '{print $1}' > mm10.gene_names.txt
cat gencode.vM4-tRNAs-ERCC.gtf | awk -F 'gene_id "' '{print $2}' | awk -F '"' '{print $1}' > mm10.gene_ENS_id.txt
paste mm10.gene_types.txt mm10.gene_names.txt mm10.gene_ENS_id.txt gencode.v39.basic.annotation.gff3 | awk -F '\t' -v OFS='\t' '{if ($1=="protein_coding" && $6=="gene") print $2,$3}' > mm10.gene2id.txt


# HSC, CMP, ERY, MK, GMP, MONO1, NEU
cat tpmFeb21_v3.tab | awk -F '\t' -v OFS='\t' '{print $1, $2,$3, $22, $12,$13, $4,$5, $25,$26, $20,$21, $19, $16,$17 }' > hg38.RNAseq.TMP.matched_ct.txt
cat rnaTPM_withcoordinates.txt | awk -F '\t' -v OFS='\t' '{print $4, $19,$20, $7, $11,$12, $13,$14, $17,$18, $15,$16, $23, $24,$25 }' > mm10.RNAseq.TMP.matched_ct.txt

R

d1 = read.table('hg38.RNAseq.TMP.matched_ct.txt', header=T)
d1[,1] = apply(d1, 1, function(x) unlist(strsplit(x[1],'[.]'))[1])

d2 = read.table('mm10.RNAseq.TMP.matched_ct.txt', header=T)
d2[,1] = apply(d2, 1, function(x) unlist(strsplit(x[1],'[.]'))[1])

d1id2name = read.table('hg38.gene2id.txt', header=F)
d1id2name_id = apply(d1id2name, 1, function(x) unlist(strsplit(x[2],'[.]'))[1])

d2id2name = read.table('mm10.gene2id.txt', header=F)
d2id2name_id = apply(d2id2name, 1, function(x) unlist(strsplit(x[2],'[.]'))[1])

### match gene_id gene_name d1
d1used = d1[is.element(d1[,1], d1id2name_id), ]
d1used_ordered = d1used[order(d1used[,1]),]
d1id_name_ordered = cbind(apply(d1id2name, 1, function(x) toString(x[1])), d1id2name_id)[order(d1id2name_id),]
d1id_name_used_ordered = d1id_name_ordered[is.element(d1id_name_ordered[,2], d1used_ordered[,1]),]
multiple_genes = rownames(table(d1id_name_used_ordered[,2])[table(d1id_name_used_ordered[,2])>=2])
d1id_name_used_ordered_rmmulti = d1id_name_used_ordered[!is.element(d1id_name_used_ordered[,2], multiple_genes),]
d1used_ordered_rmmulti = d1used_ordered[!is.element(d1used_ordered[,1], multiple_genes),]
d1used_final = cbind(d1id_name_used_ordered_rmmulti[,1], d1used_ordered_rmmulti)
### d2
d2used = d2[is.element(d2[,1], d2id2name_id), ]
d2used_ordered = d2used[order(d2used[,1]),]
d2id_name_ordered = cbind(apply(d2id2name, 1, function(x) toString(x[1])), d2id2name_id)[order(d2id2name_id),]
d2id_name_used_ordered = d2id_name_ordered[is.element(d2id_name_ordered[,2], d2used_ordered[,1]),]
multiple_genes2 = rownames(table(d2id_name_used_ordered[,2])[table(d2id_name_used_ordered[,2])>=2])
d2id_name_used_ordered_rmmulti = d2id_name_used_ordered[!is.element(d2id_name_used_ordered[,2], multiple_genes2),]
d2used_ordered_rmmulti = d2used_ordered[!is.element(d2used_ordered[,1], multiple_genes2),]
d2used_final = cbind(d2id_name_used_ordered_rmmulti[,1], d2used_ordered_rmmulti)
d2used_final[,1] = apply(d2used_final,1,function(x) toString(x[1]))
d2used_final[d2used_final[,1]==('Hba-a1'),1] = ('Hba1')

### match gene betw hg38 & mm10
multiple_genes = toupper(c(rownames(table(d1used_final[,1])[table(d1used_final[,1])>=2]), rownames(table(d2used_final[,1])[table(d2used_final[,1])>=2])))
d1used_final_match = d1used_final[is.element(toupper(d1used_final[,1]), toupper(d2used_final[,1])) & (!is.element(toupper(d1used_final[,1]), multiple_genes)),]
d2used_final_match = d2used_final[is.element(toupper(d2used_final[,1]), toupper(d1used_final[,1])) & (!is.element(toupper(d2used_final[,1]), multiple_genes)),]
d1used_final_match_order = d1used_final_match[order(d1used_final_match[,1]), ]
d2used_final_match_order = d2used_final_match[order(d2used_final_match[,1]), ]

### get cor 
set.seed(2019)
log_sd = 1
cor_vec = apply(cbind(log2(d1used_final_match_order[,-c(1,2)]*100+1)+matrix(rnorm(dim(d1used_final_match_order)[1]*(dim(d1used_final_match_order)[2]-2), mean = 0, sd = log_sd), nrow=dim(d1used_final_match_order)[1],ncol=dim(d1used_final_match_order)[2]-2), log(d2used_final_match_order[,-c(1,2)]+1))+matrix(rnorm(dim(d1used_final_match_order)[1]*(dim(d1used_final_match_order)[2]-2), mean = 0, sd = log_sd), nrow=dim(d1used_final_match_order)[1],ncol=dim(d1used_final_match_order)[2]-2), 1, function(x) cor(x[1:(dim(d1)[2]-2)], x[(dim(d1)[2]-2+1):(dim(d1)[2]*2-4)]))

#cor_vec = apply(cbind(log2(d1used_final_match_order[,-c(1,2)]*100+1), log(d2used_final_match_order[,-c(1,2)]+1)), nrow=dim(d1used_final_match_order)[1],ncol=dim(d1used_final_match_order)[2]-2), 1, function(x) cor(x[1:(dim(d1)[2]-2)], x[(dim(d1)[2]-2+1):(dim(d1)[2]*2-4)]))

### write output
gene_cor_mat = cbind(as.data.frame(d1used_final_match_order[,1]), cor_vec, d1used_final_match_order[,-c(1,2)], d2used_final_match_order[,-c(1,2)])[order(-cor_vec),]
gene_cor_mat = gene_cor_mat[!is.na(gene_cor_mat[,2]),]
colnames(gene_cor_mat)[1] = 'genes'
write.table(gene_cor_mat, 'hg38.mm10.TPM.cor.mat.txt', quote=F, sep='\t', col.names=T, row.names=F)

pdf('hg38.mm10.TPM.cor.distribution.pdf')
hist(gene_cor_mat[,2], breaks=50)
abline(v=gene_cor_mat[gene_cor_mat[,1]=='GATA1',2])
abline(v=0, lwd=2)
box()
dev.off()


target_genes = c('GLOD5', 'HDAC6', 'ERAS', 'PCSK1N')
print(gene_cor_mat[is.element(gene_cor_mat[,1], target_genes), ])
```
