## Supplementary material for "Interspecies regulatory landscapes and elements revealed by novel joint systematic integration of human and mouse blood cell epigenomes": Scripts for IDEAS modeling of epigenetic states jointly in human and mouse: get_hg38_mm10.state_similarity.0705_PSU_poster.sh

```
hg38_chrom_size=/Users/guanjuexiang/Documents/projects/S3V2_IDEAS_ESMP/genomesize/hg38.chrom.1_22XY.sizes
mm10_chrom_size=/Users/guanjuexiang/Documents/projects/S3V2_IDEAS_ESMP/genomesize/mm10.chrom.1_19XY.sizes
script_dir=/Users/guanjuexiang/Documents/projects/Joint_Human_Mouse_IDEAS_State/other_scripts/compare_human_mouse_state_similarity/               
working_dir=/Users/guanjuexiang/Documents/projects/analysis/0813_human_mouse_state_compare_heatmap/

### 
cd $working_dir 

### prepare_input_data
#time bash 00.prepare_input_data.sh


### get heatmap
#time bash $script_dir/get_hg38_mm10.state_similarity.sh GATA1 Gata1 50000 50000 hg38.gene.bed mm10.gene.bed S3V2_IDEAS_hg38_r3_withHg38Mm10prior.state.matched_ct.full.bed S3V2_IDEAS_mm10_r3_withHg38Mm10prior.state.matched_ct.full.bed
#time bash $script_dir/get_hg38_mm10.state_similarity.sh Gata1 Gata1 50000 50000 mm10.gene.bed mm10.gene.bed S3V2_IDEAS_mm10_r3_withHg38Mm10prior.state.matched_ct.full.bed S3V2_IDEAS_mm10_r3_withHg38Mm10prior.state.matched_ct.full.bed
#time bash $script_dir/get_hg38_mm10.state_similarity.sh GATA1 GATA1 50000 50000 hg38.gene.bed hg38.gene.bed S3V2_IDEAS_hg38_r3_withHg38Mm10prior.state.matched_ct.full.bed S3V2_IDEAS_hg38_r3_withHg38Mm10prior.state.matched_ct.full.bed
#time bash $script_dir/get_hg38_mm10.state_similarity.sh GATA1 Gata1 50000 50000 hg38.gene.bed mm10.gene.bed S3V2_IDEAS_hg38_r3_withHg38Mm10prior.state.matched_ct.full.bed S3V2_IDEAS_mm10_r3_withHg38Mm10prior.state.matched_ct.full.bed statep_rna_coe_heatmap.HM.all.ccre.withcorfilter.AVE.txt


#chrX	7919401	8020800	-	Gata1
#chrX	48760001	48836000	+	GATA1


hg38_gene=$1
mm10_gene=$2
hg38_gene_exp_win_u=$3
hg38_gene_exp_win_d=$4
mm10_gene_exp_win_u=$5
mm10_gene_exp_win_d=$6
hg38_gene_set=$7
mm10_gene_set=$8
hg38_state_set=$9
mm10_state_set=${10}

# time bash get_hg38_mm10.state_similarity.sh GATA1 Gata1 50000 50000 50000 50000 hg38.gene.bed mm10.gene.bed S3V2_IDEAS_hg38_r3_withHg38Mm10prior.state.matched_ct.full.bed S3V2_IDEAS_mm10_r3_withHg38Mm10prior.state.matched_ct.full.bed
# time bash get_hg38_mm10.state_similarity.sh GATA1 Gata1 26561 49438 58670 42729 hg38.gene.bed mm10.gene.bed S3V2_IDEAS_hg38_r3_withHg38Mm10prior.state.matched_ct.full.bed S3V2_IDEAS_mm10_r3_withHg38Mm10prior.state.matched_ct.full.bed


hg38_gene='GATA1'
mm10_gene='Gata1'
hg38_gene_set='hg38.gene.bed'
mm10_gene_set='mm10.gene.bed'
hg38_state_set='S3V2_IDEAS_hg38_r3_withHg38Mm10prior.state.matched_ct.bed'
mm10_state_set='S3V2_IDEAS_mm10_r3_withHg38Mm10prior.state.matched_ct.bed'
para_file='06a_S3V2_IDEAS_hg38_r3_withHg38Mm10prior.para.modified.para'
hg38_gene_exp_win=50000
mm10_gene_exp_win=50000
hg38_gene_exp_win_u=50000
hg38_gene_exp_win_d=50000
mm10_gene_exp_win_u=50000
mm10_gene_exp_win_d=50000
feature_num=8


para_file='06a_S3V2_IDEAS_hg38_r3_withHg38Mm10prior.para.modified.para'
hg38_gene_exp_win_u=100000
hg38_gene_exp_win_d=100000
mm10_gene_exp_win_u=100000
mm10_gene_exp_win_d=100000
feature_num=8

###
echo $hg38_gene
echo $mm10_gene

### get query gene
#cat $hg38_gene_set | awk -F '\t' -v OFS='\t' -v hg38_gene=$hg38_gene '{if (toupper($5)==toupper(hg38_gene)) print $1,$2,$3,$4,$5}' | awk -F '\t' -v OFS='\t' -v hg38_gene_exp_win_u=$hg38_gene_exp_win_u -v hg38_gene_exp_win_d=$hg38_gene_exp_win_d '{if ($4=="+") print $1,$2-hg38_gene_exp_win_u,$2+hg38_gene_exp_win_d,$4,$5; else print $1,$3-hg38_gene_exp_win_u,$3+hg38_gene_exp_win_d,$4,$5}' | awk -F '\t' -v OFS='\t' '{if ($2<0) print $1,0,$3,$4,$5; else print $0}' > 'hg38.gene.'$hg38_gene'.bed'
#cat $mm10_gene_set | awk -F '\t' -v OFS='\t' -v mm10_gene=$mm10_gene '{if (toupper($5)==toupper(mm10_gene)) print $1,$2,$3,$4,$5}' | awk -F '\t' -v OFS='\t' -v mm10_gene_exp_win_u=$mm10_gene_exp_win_u -v mm10_gene_exp_win_d=$mm10_gene_exp_win_d '{if ($4=="+") print $1,$2-mm10_gene_exp_win_u,$2+mm10_gene_exp_win_d,$4,$5; else print $1,$3-mm10_gene_exp_win_u,$3+mm10_gene_exp_win_d,$4,$5}' | awk -F '\t' -v OFS='\t' '{if ($2<0) print $1,0,$3,$4,$5; else print $0}' > 'mm10.gene.'$mm10_gene'.bed'

### intersect state bed
bedtools intersect -a $hg38_state_set -b 'hg38.gene.'$hg38_gene'.bed' -wa -u > 'hg38.gene.'$hg38_gene'.matched_ct.state.bed'
bedtools intersect -a $mm10_state_set -b 'mm10.gene.'$mm10_gene'.bed' -wa -u > 'mm10.gene.'$mm10_gene'.matched_ct.state.bed'

### get state similarity matrix
cat 'hg38.gene.'$hg38_gene'.bed' 'mm10.gene.'$mm10_gene'.bed'
Rscript $script_dir/get_hg38_mm10.cor.heatmap.R 'hg38.gene.'$hg38_gene'.matched_ct.state.bed' 'mm10.gene.'$mm10_gene'.matched_ct.state.bed' $hg38_gene'.'$mm10_gene'.cor.heatmap.png' $para_file $feature_num 
#Rscript $script_dir/get_hg38_mm10.cor.heatmap.coe.R 'hg38.gene.'$hg38_gene'.matched_ct.state.bed' 'mm10.gene.'$mm10_gene'.matched_ct.state.bed' $hg38_gene'.'$mm10_gene'.cor.coe.heatmap.png' 06a_S3V2_IDEAS_hg38_r3_withHg38Mm10prior.para.modified.para 8 statep_rna_coe_heatmap.HM.all.ccre.withcorfilter.AVE.txt

Rscript $script_dir/get_hg38_mm10.cor.heatmap.R 'hg38.gene.'$hg38_gene'.matched_ct.state.bed' 'hg38.gene.'$hg38_gene'.matched_ct.state.bed' $hg38_gene'.hg38.cor.heatmap.png' $para_file $feature_num 
Rscript $script_dir/get_hg38_mm10.cor.heatmap.R 'mm10.gene.'$mm10_gene'.matched_ct.state.bed' 'mm10.gene.'$mm10_gene'.matched_ct.state.bed' $mm10_gene'.mm10.cor.heatmap.png' $para_file $feature_num 
```
