## Supplementary material for "Interspecies regulatory landscapes and elements revealed by novel joint systematic integration of human and mouse blood cell epigenomes": Scripts for IDEAS modeling of epigenetic states jointly in human and mouse: get_shared.genes.R

```
args = commandArgs(trailingOnly=TRUE)

hg38_gene_file = args[1]
mm10_gene_file = args[2]
output_file = args[3]
#hg38_gene_file = 'hg38.gene.bed'
#mm10_gene_file = 'mm10.gene.bed'
#output_file = 'hg38.mm10.shared.genes'

d1 = read.table(hg38_gene_file, header=F)[,5]
d2 = read.table(mm10_gene_file, header=F)[,5]
d2a = toupper(d2)

chr = read.table(hg38_gene_file, header=F)[,1]

d12 = d2[is.element(d2a, d1[chr!='chrM'])]

d12_mat = cbind(toupper(d12), d12)

write.table(d12_mat, output_file, quote=F, col.names=F, row.names=F)
```
