## Supplementary material for "Interspecies regulatory landscapes and elements revealed by novel joint systematic integration of human and mouse blood cell epigenomes": Scripts for IDEAS modeling of epigenetic states jointly in human and mouse: get_TCD4_enh.sh

```
# get the master enhancers
cd /mnt/ebs/EPIC_input_data/peak_files/master_enh
cat /mnt/ebs/analysis/EPIC/2209.K562-K562.EPIC/K562/K562_EP/K562.enhancers.bed | awk -F '\t' -v OFS='\t' '{if (($2-250)>0) print $1,$2-250,$3+250,$4; else print $1,0,$3+250,$4}' > Master.Enh.1kb.bed


# get the TCD4 enhancers
cd /mnt/ebs/EPIC_input_data/peak_files/TCD4
# 
aws s3 cp s3://maraudertx.rd/processed_data/public/Blood/CD4+TCell_Naive_ATACseq/bwa/merged_replicate/macs2/narrow_peak/CD4+TCell.ATACseq.Naive.20230428.mRp.clN_summits.bed ./
aws s3 cp s3://maraudertx.rd/processed_data/public/Blood/CD4+TCell.ChIPseq.Naive_H3K27ac.rep1.20230428/CD4+TCell.ChIPseq.Naive_H3K27ac.rep1.20230428_peaks.narrowPeak ./
aws s3 cp s3://maraudertx.rd/processed_data/public/Blood/CD4+TCell.ChIPseq.Naive_H3K4me1.rep1.20230428/broad_peaks/bwa/mergedLibrary/macs/broadPeak/CD4+TCell.ChIPseq.Naive_H3K4me1.rep1.20230428_R1_peaks.broadPeak ./
bedtools intersect -a CD4+TCell.ATACseq.Naive.20230428.mRp.clN_summits.bed -b /mnt/ebs/reference_genome/hg38/hg38as.chrom.1_22XY.sizes.canonical.bed > CD4+TCell.ATACseq.Naive.20230428.mRp.clN_summits.hg38.bed
bedtools intersect -a CD4+TCell.ChIPseq.Naive_H3K27ac.rep1.20230428_peaks.narrowPeak -b /mnt/ebs/reference_genome/hg38/hg38as.chrom.1_22XY.sizes.canonical.bed > CD4+TCell.ChIPseq.Naive_H3K27ac.rep1.20230428_peaks.narrowPeak.hg38.bed


# get H3K4me1 H3K27ac union peaks
cut -f1,2,3 CD4+TCell.ChIPseq.Naive_H3K27ac.rep1.20230428_peaks.narrowPeak > CD4+TCell.ChIPseq.Naive.H3K27ac_H3K4me1.union_peaks.bed
cut -f1,2,3 CD4+TCell.ChIPseq.Naive_H3K4me1.rep1.20230428_R1_peaks.broadPeak >> CD4+TCell.ChIPseq.Naive.H3K27ac_H3K4me1.union_peaks.bed

# get the TCD4 enhancers
bedtools intersect -a CD4+TCell.ATACseq.Naive.20230428.mRp.clN_summits.bed -b CD4+TCell.ChIPseq.Naive.H3K27ac_H3K4me1.union_peaks.bed -wa -u > CD4+TCell.ATACseq.Naive.20230428.mRp.clN_summits.H3K27ac_H3K4me1_union.bed
cat CD4+TCell.ATACseq.Naive.20230428.mRp.clN_summits.H3K27ac_H3K4me1_union.bed | awk -F '\t' -v OFS='\t' '{if (($2-250)>0) print $1,$2-250,$3+250, "T"NR; else print $1,0,$3+250, "T"NR}' > CD4+TCell.Naive.20230428.Enh.500bp.bed

# get new master enhancers
cd /mnt/ebs/EPIC_input_data/peak_files/TCD4
mater_enh_od='/mnt/ebs/EPIC_input_data/peak_files/master_enh/HepG2.K562.enhancers.summit_500bp.bed'
bedtools intersect -a $mater_enh_od -b CD4+TCell.Naive.20230428.Enh.500bp.bed -v > Master.Enh.500bp.Not_TCD4.bed
cat Master.Enh.500bp.Not_TCD4.bed CD4+TCell.Naive.20230428.Enh.500bp.bed > Master.Enh.500bp.TCD4adj.bed
bedtools intersect -a Master.Enh.500bp.TCD4adj.bed -b /mnt/ebs/reference_genome/hg38/hg38as.chrom.1_22XY.sizes.canonical.bed > Master.Enh.500bp.TCD4adj.hg38.bed
# get peak center
cat Master.Enh.500bp.TCD4adj.hg38.bed | awk -F '\t' -v OFS='\t' '{print $1,int(($2+$3)/2),int(($2+$3)/2)+1,$4}' > Master.Enh.center.TCD4adj.hg38.bed
bedtools intersect -a CD4+TCell.Naive.20230428.Enh.500bp.bed -b /mnt/ebs/reference_genome/hg38/hg38as.chrom.1_22XY.sizes.canonical.bed > CD4+TCell.Naive.20230428.Enh.500bp.hg38.bed
cat CD4+TCell.Naive.20230428.Enh.500bp.hg38.bed | awk -F '\t' -v OFS='\t' '{print $1,int(($2+$3)/2),int(($2+$3)/2)+1,$4}' > CD4+TCell.Naive.20230428.Enh.center.500bp.hg38.bed


# master enhancer should looks like this in EPIC instance:
# /mnt/ebs/analysis/EPIC/2209.HepG2-K562.EPIC/HepG2_K562_MP/HepG2.K562.enhancers.summit_500bp.bed
chr8	70895	71396	1
chr8	94250	94751	2
chr8	206198	206699	3
chr8	206436	206937	4
chr8	206896	207397	5
chr8	211711	212212	6
chr8	212063	212564	7
chr8	212450	212951	8
chr8	213050	213551	9
chr8	214452	214953	10


# put the Master enhancer in EPIC cell-type folder
mkdir /mnt/ebs_tcd4/analysis/EPIC/2309.K562-K562.EPIC/K562_K562_MP
cp /mnt/ebs/EPIC_input_data/peak_files/TCD4/Master.Enh.500bp.TCD4adj.hg38.bed /mnt/ebs_tcd4/analysis/EPIC/2309.K562-K562.EPIC/K562_K562_MP/
```
