## Supplementary material for "Interspecies regulatory landscapes and elements revealed by novel joint systematic integration of human and mouse blood cell epigenomes": Scripts for IDEAS modeling of epigenetic states jointly in human and mouse: NMF.cor.mat.decompose.R

```
setwd('/Users/guanjuexiang/Documents/projects/analysis/0813_human_mouse_state_compare_heatmap')

# conda activate shiny
library(pheatmap)
library(ComplexHeatmap)
library(InteractiveComplexHeatmap)
library(shiny)
library(circlize)

# define NMF function
nmf <- function(V, k, max_iter = 1000, tol = 1e-4) {
  # Initialize W and H with random values
  n <- dim(V)[1]
  m <- dim(V)[2]
  W <- matrix(runif(n * k), nrow = n)
  H <- matrix(runif(k * m), nrow = k)
  for (i in 1:max_iter) {
    # Update H
    WH <- W %*% H
    H <- H * ((t(W) %*% (V / (WH + 1e-10))) / (t(W) %*% matrix(1, nrow = n, ncol = m)))
    # Update W
    WH <- W %*% H
    W <- W * (((V / (WH + 1e-10)) %*% t(H)) / (matrix(1, nrow = n, ncol = m) %*% t(H)))
    # Normalize W and H
    if (i < round(max_iter*0.5)) {
      W <- W / max(W)
      H <- H / max(H)
    }
    # Check convergence
    WH <- W %*% H
    obj <- sum(V * log(1 / (WH + 1e-10)) + WH)
    if (i > 1 && abs(obj - prev_obj) < tol) {
      break
    }
    prev_obj <- obj
  }  
  output = list(W = W, H = H)
  return(output)
}

# output plot folder
hg38_gene = 'GATA1'
mm10_gene = 'Gata1'
output_folder = paste0('NMF_reconstruction_', hg38_gene, '_', mm10_gene)
input_correlation_mat = paste0(hg38_gene, '.', mm10_gene, '.cor.heatmap.png.cor.mat.txt')
hg38_coord = paste0('hg38.gene.', hg38_gene, '.matched_ct.state.bed')
mm10_coord = paste0('mm10.gene.', mm10_gene, '.matched_ct.state.bed')

# read correlation matrix
d_mk_cor = as.matrix(read.table(input_correlation_mat, header=F, sep='\t', comment.char='~'))
d_mk_cor_mat = as.matrix(d_mk_cor)

# read coordinates
bed_hg38 = read.table(hg38_coord, header=F, sep='\t', comment.char='~')
bed_mm10 = read.table(mm10_coord, header=F, sep='\t', comment.char='~')
# add rownames and colnames
colnames(d_mk_cor_mat) = apply(bed_hg38, 1, function(x) paste0('H_', x[1], '_', x[2], '_', x[3]))
rownames(d_mk_cor_mat) = apply(bed_mm10, 1, function(x) paste0('M_', x[1], '_', x[2], '_', x[3]))

# get positive correlation matrix
d_mk_cor_mat_pos = d_mk_cor_mat
d_mk_cor_mat_pos[d_mk_cor_mat_pos<0] = 0

# decide the number of components based on the BIC
set.seed(2019)
set.seed(2023)
BIC_mat = c()
for (j in 1:1){
    print(j)
    BIC_vec = c()
    for (i in seq(1, 20, by = 1)) {
        print(i)
        sparse_nmf_result <- nmf(d_mk_cor_mat_pos, k = i)
        W <- sparse_nmf_result$W
        H <- sparse_nmf_result$H
        reconstructed_whitened_R <- W %*% H
        # get BIC of the model
        BIC = nrow(d_mk_cor_mat_pos) * ncol(d_mk_cor_mat_pos) * log(sum((reconstructed_whitened_R - d_mk_cor_mat_pos)^2)) + i * (nrow(d_mk_cor_mat_pos) + ncol(d_mk_cor_mat_pos)) * log(nrow(d_mk_cor_mat_pos) * ncol(d_mk_cor_mat_pos))
        print(BIC)
        BIC_vec = c(BIC_vec, BIC)
    }
    BIC_mat = rbind(BIC_mat, BIC_vec)
}

# mkdir
dir.create(output_folder)
# plot the BIC
pdf(paste0(output_folder, '/BIC.1.pdf'), width = 6, height = 6)
plot(seq(1, 20, by = 1), colMeans(BIC_mat), type = 'p', xlab = 'Number of Factors', ylab = 'BIC', main = 'BIC vs. Number of Factors', cex.axis=1.2, cex.lab=1.5, cex.main=1.5)
lines(seq(1, 20, by = 1), colMeans(BIC_mat), type = 'b')
abline(v=6, lty=2)
dev.off()

#  NMF decomposition
set.seed(2019)
n_comp = 6
nmf_result <- nmf(d_mk_cor_mat_pos, k = n_comp)

# Extract the independent components (ICs) and the mixing matrix
nmf_result_W <- nmf_result$W
nmf_result_H <- nmf_result$H

# cluster factor matrix
nmf_W_H = rbind(nmf_result_W, t(nmf_result_H))
nmf_W_H_order_tree = hclust(dist(t(nmf_W_H)), method = 'average')
nmf_W_H_order = nmf_W_H_order_tree$order

# plot NMFs_mm10 heatmap
#my_colorbar=colorRampPalette(c('black', 'white'))(n = 100)
png(paste0(output_folder, '/', hg38_gene, '.', mm10_gene, '.cor.heatmap.png.cor.mat.txt.NMFs_mm10.heatmap.png'), width = 300, height = 1000)
pheatmap(nmf_result_W[,], cluster_rows=F, cluster_cols=F, width = 2, height = 6, legend=F, show_rownames=F, show_colnames=F)
dev.off()
# plot NMFs_hg38 heatmap
png(paste0(output_folder, '/', hg38_gene, '.', mm10_gene, '.cor.heatmap.png.cor.mat.txt.NMFs_hg38.heatmap.png'), width = 300, height = 1000)
pheatmap(t(nmf_result_H)[,], cluster_rows=F, cluster_cols=F, width = 2, height = 6, legend=F, show_rownames=F, show_colnames=F)
dev.off()

# plot NMF i reconstruction
#breaksList = seq(0, 1, by = 0.001)
#my_colorbar=colorRampPalette(c('white', 'red'))(n = length(breaksList))
for (i in 1:n_comp) {
    print(i)
    # hg38 & mm10
    plot_mat = as.matrix(cbind(nmf_result_W[,i]) %*% rbind(nmf_result_H[i,]))
    # high resolution png
    png(paste0(output_folder, '/', hg38_gene, '.', mm10_gene, '.cor.heatmap.png.cor.mat.txt.NMF', i,'.png'), width = 5000, height = 5000)
    pheatmap::pheatmap(plot_mat, cluster_rows=F, cluster_cols=F, legend=F, show_rownames=F, show_colnames=F)
    dev.off()
    # hg38
    plot_mat = as.matrix(cbind(nmf_result_H[i,]) %*% t(nmf_result_H[i,]))
    png(paste0(output_folder, '/', hg38_gene, '.', mm10_gene, '.cor.heatmap.png.cor.mat.txt.NMF.hg38.', i,'.png'), width = 5000, height = 5000)
    pheatmap::pheatmap(plot_mat, cluster_rows=F, cluster_cols=F, legend=F, show_rownames=F, show_colnames=F)
    dev.off()
    # mm10
    plot_mat = as.matrix(cbind(nmf_result_W[,i]) %*% t(nmf_result_W[,i]))
    png(paste0(output_folder, '/', hg38_gene, '.', mm10_gene, '.cor.heatmap.png.cor.mat.txt.NMF.mm10.', i,'.png'), width = 5000, height = 5000)
    pheatmap::pheatmap(plot_mat, cluster_rows=F, cluster_cols=F, legend=F, show_rownames=F, show_colnames=F)
    dev.off()
}

# plot NMF factor i bar plot
for (i in 1:dim(nmf_result_W)[2]){
    # plot hg38
    png(paste0(output_folder, '/', hg38_gene, '.', mm10_gene, '.cor.heatmap.png.cor.mat.txt.NMF.hg38.', i,'.bar.png'), width = 1000, height = 300)
    cor_factor = nmf_result_H[i,]
    span = 0.03
    cor_factor_sm = loess(cor_factor ~ c(1:dim(nmf_result_H)[2]), span=span)$fitted
    plot(1:dim(nmf_result_H)[2], cor_factor_sm, type='h', ylim=c(0,1))
    dev.off()
    # plot mm10
    png(paste0(output_folder, '/', hg38_gene, '.', mm10_gene, '.cor.heatmap.png.cor.mat.txt.NMF.mm10.', i,'.bar.png'), width = 1000, height = 300)
    cor_factor = nmf_result_W[,i]
    cor_factor_sm = loess(cor_factor ~ c(1:dim(nmf_result_W)[1]), span=span)$fitted
    plot(1:dim(nmf_result_W)[1], cor_factor_sm, type='h', ylim=c(0,1))
    dev.off()
}

zp2 = function(x, thresh) {
    # 1st round z score
    z = (x - mean(x)) / sd(x)
    # get z score one-side p value
    zp = 1 - pnorm(z)
    # 2nd round z score
    z2 = (x - mean(x[zp>thresh])) / sd(x[zp>thresh])
    # get z score one-side p value
    zp2 = 1 - pnorm(z2)
    return(zp2)
}

zp = function(x, thresh) {
    # 1st round z score
    z = (x - mean(x)) / sd(x)
    # get z score one-side p value
    zp = 1 - pnorm(z)
    return(zp)
}


# plot NMF factor i z score p-value bar plot
for (i in 2:dim(nmf_result_W)[2]){
    # plot hg38
    png(paste0(output_folder, '/', hg38_gene, '.', mm10_gene, '.cor.heatmap.png.cor.mat.txt.NMF.hg38.', i,'.zp.bar.png'), width = 1000, height = 100)
    par(mar = rep(0.5, 4))
    cor_factor = nmf_result_H[i,]
    span = 0.03
    cor_factor_sm = loess(cor_factor ~ c(1:dim(nmf_result_H)[2]), span=span)$fitted
    cor_factor_sm[cor_factor_sm<=0] = 0.1
    cor_factor_sm_zp = -log10(zp2(cor_factor_sm, 0.01))
    #plot(1:dim(nmf_result_H)[2], cor_factor_sm_zp, type='h', ylim=c(0,16), xlim=c(1,dim(nmf_result_H)[2]), xlab='', ylab='', xaxt='n', yaxt='n')
    barplot(cor_factor_sm_zp, xlab='', ylab='', xaxt='n', yaxt='n', ylim=c(0,8))
    dev.off()
    # fdr pk
    png(paste0(output_folder, '/', hg38_gene, '.', mm10_gene, '.cor.heatmap.png.cor.mat.txt.NMF.hg38.', i,'.zpfdr.bar.png'), width = 1000, height = 50)
    cor_factor = nmf_result_H[i,]
    span = 0.03
    cor_factor_sm = loess(cor_factor ~ c(1:dim(nmf_result_H)[2]), span=span)$fitted
    cor_factor_sm[cor_factor_sm<=0] = 1e-16
    cor_factor_sm_zp_fdr = (p.adjust(zp2(cor_factor_sm, 0.01), method = 'fdr')<0.1)*1
    #plot(1:dim(nmf_result_H)[2], cor_factor_sm_zp_fdr, type='h', ylim=c(0,1))
    pheatmap(rbind(cor_factor_sm_zp_fdr), cluster_rows=F, cluster_cols=F, legend=F, show_rownames=F, show_colnames=F, color=c('gray', 'black'))
    dev.off()
    # plot mm10
    png(paste0(output_folder, '/', hg38_gene, '.', mm10_gene, '.cor.heatmap.png.cor.mat.txt.NMF.mm10.', i,'.zp.bar.png'), width = 1000, height = 100)
    par(mar = rep(0.5, 4))
    cor_factor = nmf_result_W[,i]
    cor_factor_sm = loess(cor_factor ~ c(1:dim(nmf_result_W)[1]), span=span)$fitted
    cor_factor_sm[cor_factor_sm<=0] = 1e-16
    cor_factor_sm_zp = -log10(zp2(cor_factor_sm, 0.01))
    cor_factor_sm_zp[!is.finite(cor_factor_sm_zp)] = 16
    # plot barplot without axis labels
    #plot(1:dim(nmf_result_W)[1], cor_factor_sm_zp, type='h', ylim=c(0,16), xlim=c(1,dim(nmf_result_W)[1]), xlab='', ylab='', xaxt='n', yaxt='n')
    barplot(cor_factor_sm_zp, xlab='', ylab='', xaxt='n', yaxt='n', ylim=c(0,8))
    dev.off()
    # fdr pk
    png(paste0(output_folder, '/', hg38_gene, '.', mm10_gene, '.cor.heatmap.png.cor.mat.txt.NMF.mm10.', i,'.zpfdr.bar.png'), width = 1000, height = 50)
    cor_factor = nmf_result_W[,i]
    cor_factor_sm = loess(cor_factor ~ c(1:dim(nmf_result_W)[1]), span=span)$fitted
    cor_factor_sm[cor_factor_sm<=0] = 1e-16
    cor_factor_sm_zp_fdr = (p.adjust(zp2(cor_factor_sm, 0.01), method = 'fdr')<0.1)*1
    #plot(1:dim(nmf_result_W)[1], cor_factor_sm_zp_fdr, type='h', ylim=c(0,1))
    pheatmap(rbind(cor_factor_sm_zp_fdr), cluster_rows=F, cluster_cols=F, legend=F, show_rownames=F, show_colnames=F, color=c('gray', 'black'))
    dev.off()
}


# reconstructed correlation matrix
reconstructed_d <- nmf_result_W %*% nmf_result_H
reconstructed_d_mm10 <- nmf_result_W %*% t(nmf_result_W)
reconstructed_d_hg38 <- t(nmf_result_H) %*% nmf_result_H

# plot reconstructed correlation matrix
png(paste0(output_folder, '/', hg38_gene, '.', mm10_gene, '.cor.heatmap.png.cor.mat.txt.NMF.hg38_mm10.all.png'), width = 1000, height = 1000)
pheatmap(reconstructed_d , cluster_rows=F, cluster_cols=F, legend=F, show_rownames=F, show_colnames=F)
dev.off()
png(paste0(output_folder, '/', hg38_gene, '.', mm10_gene, '.cor.heatmap.png.cor.mat.txt.NMF.mm10.png'), width = 1000, height = 1000)
pheatmap(reconstructed_d_mm10 , cluster_rows=F, cluster_cols=F, legend=F, show_rownames=F, show_colnames=F)
dev.off()
png(paste0(output_folder, '/', hg38_gene, '.', mm10_gene, '.cor.heatmap.png.cor.mat.txt.NMF.hg38.png'), width = 1000, height = 1000)
pheatmap(reconstructed_d_hg38 , cluster_rows=F, cluster_cols=F, legend=F, show_rownames=F, show_colnames=F)
dev.off()

# plot factor 1 & 5 reconstructed correlation matrix
used_col = c(3,4)
reconstructed_d <- nmf_result_W[,used_col] %*% nmf_result_H[used_col,]
reconstructed_d_mm10 <- nmf_result_W[,used_col] %*% t(nmf_result_W[,used_col])
reconstructed_d_hg38 <- t(nmf_result_H[used_col,]) %*% nmf_result_H[used_col,]
png(paste0(output_folder, '/', hg38_gene, '.', mm10_gene, '.cor.heatmap.png.cor.mat.txt.NMF.3_4.png'), width = 1000, height = 1000)
pheatmap(reconstructed_d, cluster_rows=F, cluster_cols=F, legend=F, show_rownames=F, show_colnames=F)
dev.off()
png(paste0(output_folder, '/', hg38_gene, '.', mm10_gene, '.cor.heatmap.png.cor.mat.txt.NMF.mm10.3_4.png'), width = 1000, height = 1000)
pheatmap(reconstructed_d_mm10 , cluster_rows=F, cluster_cols=F, legend=F, show_rownames=F, show_colnames=F)
dev.off()
png(paste0(output_folder, '/', hg38_gene, '.', mm10_gene, '.cor.heatmap.png.cor.mat.txt.NMF.hg38.3_4.png'), width = 1000, height = 1000)
pheatmap(reconstructed_d_hg38 , cluster_rows=F, cluster_cols=F, legend=F, show_rownames=F, show_colnames=F)
dev.off()

### Generate plots for ShinyApp
# set colorbar
breaks <- seq(-1, 1, length.out = 101)
colors <- colorRamp2(breaks, colorRampPalette(c("blue", "white", "red"))(length(breaks)))

#
cor_mat1 = Heatmap(d_mk_cor_mat_pos, name = "cor_mat1",
    show_row_names = F, show_column_names = F, cluster_columns =F, cluster_rows = F, width=1000, col = colors)
#
cor_mat2 = Heatmap(t(d_mk_cor_mat_pos), name = "cor_mat2",
    show_row_names = F, show_column_names = F, cluster_columns =F, cluster_rows = F, width=1000, col = colors)
#
cor_mat3 = Heatmap(reconstructed_d_mm10, name = "cor_mat_mm10",
    show_row_names = F, show_column_names = F, cluster_columns =F, cluster_rows = F, width=1000, col = colors)
#
cor_mat4 = Heatmap(reconstructed_d_hg38, name = "cor_mat_hg38",
    show_row_names = F, show_column_names = F, cluster_columns =F, cluster_rows = F, width=1000, col = colors)

# plot NMFs_mm10 heatmap
mm10_NMF = Heatmap(nmf_result_W, name = "mm10_NMF", show_row_names = F, show_column_names = F, cluster_columns =F, cluster_rows = F, width=200)
# plot NMFs_hg38 heatmap
hg38_NMF = Heatmap(t(nmf_result_H), name = "hg38_NMF", show_row_names = F, show_column_names = F, cluster_columns =F, cluster_rows = F, width=200)

# add two heatmaps together to get interactive heatmap
ht1 = cor_mat1+mm10_NMF
ht2 = cor_mat2+hg38_NMF
ht3 = cor_mat3+mm10_NMF
ht4 = cor_mat4+hg38_NMF

# plot two interactive heatmaps
ui = fluidPage(
    h3("The first heatmap"),
    InteractiveComplexHeatmapOutput("ht1"),
    hr(),
    h3("The second heatmap"),
    InteractiveComplexHeatmapOutput("ht2"),
    hr(),
    h3("The mm10 heatmap"),
    InteractiveComplexHeatmapOutput("ht3"),
    hr(),
    h3("The hg38 heatmap"),
    InteractiveComplexHeatmapOutput("ht4"),

  titlePanel("Simple 2D Scatterplot"),  
  sidebarLayout(
    sidebarPanel(
      numericInput("n_points", "Number of points:", value = 100, min = 1),
      numericInput("correlation", "Correlation:", value = 0.5, min = -1, max = 1)
    ),
    mainPanel(
      plotOutput("scatterplot")
    )
  )
)

server = function(input, output, session){
    makeInteractiveComplexHeatmap(input, output, session, ht1, "ht1")
    makeInteractiveComplexHeatmap(input, output, session, ht2, "ht2")
    makeInteractiveComplexHeatmap(input, output, session, ht3, "ht3")
    makeInteractiveComplexHeatmap(input, output, session, ht4, "ht4")
    output$scatterplot <- renderPlot({
        # Generate random data
        set.seed(42)
        x <- rnorm(input$n_points)
        y <- rnorm(input$n_points, mean = input$correlation * x)
        data <- data.frame(x = x, y = y)
        
        # Create scatterplot using ggplot2
        ggplot(data, aes(x = x, y = y)) +
        geom_point() +
        theme_minimal()
    })
}

shiny::shinyApp(ui, server)


# Define UI
ui <- fluidPage(
  titlePanel("ENCODE Cross-Feature Correlation Heatmap (K562)"),
  sidebarLayout(
    sidebarPanel(
      selectInput("columns",
                  "Select Columns:",
                  choices = colnames(d_mk_cor_shiny),
                  selected = colnames(d_mk_cor_shiny),
                  multiple = TRUE)
    ),
    mainPanel(
      plotOutput("heatmap", width = "100%", height = "1000px")
    )
  )
)

# Define server
server <- function(input, output) {
  output$heatmap <- renderPlot({
    req(input$columns)
    
    # Subset correlation matrix
    selected_columns <- intersect(colnames(d_mk_cor_shiny), input$columns)
    subset_matrix <- d_mk_cor_shiny[selected_columns, selected_columns]
    
    # Convert matrix to long format for ggplot2
    long_data <- melt(subset_matrix)
    print(long_data)
    
    # Create heatmap
    p <- ggplot(long_data, aes(x = Var1, y = Var2, fill = value)) +
        geom_tile() +
        scale_fill_gradient2(low = "blue", mid = "white", high = "red", midpoint = 0) +
        theme_minimal() +
        theme(axis.text.x = element_text(angle = 45, hjust = 1, size = 18), # Increase x-axis text size
        axis.text.y = element_text(size = 18), # Increase y-axis text size
        axis.title = element_text(size = 18)) + # Increase axis title size
        coord_fixed(ratio = 1) 
    
    return(p)
  })
}

# Run the application
shinyApp(ui = ui, server = server)


library(ComplexHeatmap)
library(InteractiveComplexHeatmap)

# plot correlation matrix
breaksList = seq(-1, 1, by = 0.001)
my_colorbar=colorRampPalette(c('blue', 'white', 'red'))(n = length(breaksList))
ht1 = pheatmap(d_mk_cor, cluster_rows=F, cluster_cols=F, legend=F, show_rownames=F, show_colnames=F, color=my_colorbar, breaks=breaksList) %v%
Heatmap(nmf_result_H, cluster_rows=F, cluster_columns=F, height = unit(10, "cm")) 
ht1 = draw(ht1)

ht2 = pheatmap(d_mk_cor, cluster_rows=F, cluster_cols=F, legend=F, show_rownames=F, show_colnames=F, color=my_colorbar, breaks=breaksList) +
Heatmap(nmf_result_W, cluster_rows=F, cluster_columns=F, width = unit(10, "cm"))
ht2 = draw(ht2)


set.seed(2019)
all_colors <- colors()
# Set the number of colors you want
n <- 14
# Sample n unique random colors
random_colors <- sample(all_colors, n)


df_top = as.data.frame(t(nmf_result_H))
colnames(df_top) = paste0('H',1:dim(df_top)[2])
color_vec = c()
for (i in 1:dim(df_top)[2]){
    col_fun = colorRamp2(c(min(nmf_result_H[i,]), max(nmf_result_H[i,])), c("white", random_colors[i]))
    col_fun1 = col_fun(nmf_result_H[i,])
    color_list[colnames(df_top)[i]] = col_fun1
}

names(color_vec) <- colnames(nmf_result_H)

# NMF matrices
ha = HeatmapAnnotation(df = df_top, 
                       show_annotation_name = TRUE, 
                       show_legend = FALSE, height = unit(2, "mm"), col=color_list)

ha_left = rowAnnotation(df = as.data.frame(nmf_result_W), 
                        show_annotation_name = TRUE, 
                        show_legend = FALSE, width = unit(2, "mm"))

pdf('test.pdf', width=15, height=15)
Heatmap(d_mk_cor, 
        top_annotation = ha, 
        left_annotation = ha_left,
        name = "correlation",
        show_column_names = FALSE,
        show_row_names = FALSE,
        cluster_rows=F, cluster_columns=F)
dev.off()


col_fun = colorRamp2(c(min(nmf_result_H[i,]), max(nmf_result_H[i,])), c("white", random_colors[i]))
bbb = col_fun(nmf_result_H[i,])


# Generate Heatmap
ht0 = Heatmap(d_mk_cor, 
        top_annotation = ha, 
        left_annotation = ha_left,
        name = "correlation",
        show_column_names = FALSE,
        show_row_names = FALSE,
        cluster_rows=F, cluster_columns=F)
#


ha = HeatmapAnnotation(df = as.data.frame(t(nmf_result_H)[,1]), 
                       show_annotation_name = TRUE, 
                       show_legend = FALSE, height = unit(20, "mm"))

ha_left = rowAnnotation(df = as.data.frame(nmf_result_W[,1]), 
                        show_annotation_name = TRUE, 
                        show_legend = FALSE, width = unit(20, "mm"))

pdf('test1.pdf', width=15, height=15)
d_mk_cor1 = cbind(nmf_result_W[,1]) %*% rbind(t(nmf_result_H)[,1])
Heatmap(d_mk_cor1, 
        top_annotation = ha, 
        left_annotation = ha_left,
        name = "correlation",
        show_column_names = FALSE,
        show_row_names = FALSE,
        cluster_rows=F, cluster_columns=F)
dev.off()


# Generate Heatmap
d_mk_cor1 = cbind(nmf_result_W[,1]) %*% rbind(t(nmf_result_H)[,1])
ht1 = Heatmap(d_mk_cor1, 
        top_annotation = ha, 
        left_annotation = ha_left,
        name = "correlation",
        show_column_names = FALSE,
        show_row_names = FALSE,
        cluster_rows=F, cluster_columns=F)
#


ui = fluidPage(
    h3("The first heatmap"),
    InteractiveComplexHeatmapOutput("heatmap_1"),
    hr(),
    h3("The second heatmap"),
    InteractiveComplexHeatmapOutput("heatmap_2")
)
server = function(input, output, session) {
    makeInteractiveComplexHeatmap(input, output, session, ht0, "heatmap_1")
    makeInteractiveComplexHeatmap(input, output, session, ht1, "heatmap_2")
}
shinyApp(ui, server)


ht_list = Heatmap(mat1, name = "mat_a", row_km = 2, column_km = 2,
        top_annotation = HeatmapAnnotation(foo = anno_points(runif(10)))) +
    rowAnnotation(bar = anno_barplot(sample(10, 10))) +
    Heatmap(mat2, name = "mat_b")


htShiny(list(ht1,ht2))


htShiny(ht)


ht_list = Heatmap(mat2, col = colorRamp2(c(-1.5, 0, 1.5), c("blue", "white", "red")), 
    show_row_names = FALSE, name = "scaled_expr", column_title = qq("relative expression for @{nrow(mat)} genes"),
    show_column_names = FALSE, width = unit(8, "cm"),
    heatmap_legend_param = list(title = "Scaled expr")) +
    Heatmap(base_mean, name = "base_expr", show_row_names = FALSE, width = unit(5, "mm"),
        heatmap_legend_param = list(title = "Base expr")) +
    Heatmap(rpl + 0, name = "ribonucleoprotein", col = c("0" = "white", "1" = "purple"), 
        show_heatmap_legend = FALSE, width = unit(5, "mm")) +
    Heatmap(ccl + 0, name = "cell_cycle", col = c("0" = "white", "1" = "red"), 
        show_heatmap_legend = FALSE, width = unit(5, "mm")) +
    rowAnnotation(link = row_anno_link(at = which(ccl & base_mean > quantile(base_mean, 0.25)), 
        labels = cc_gene, labels_gp = gpar(fontsize = 10), padding = 0.5), 
        width = unit(1, "cm") + max_text_width(cc_gene, gp = gpar(fontsize = 8))) +
    Heatmap(cor(t(mat2)), name = "cor", col = colorRamp2(c(-1, 0, 1), c("green", "white", "red")), 
        show_row_names = FALSE, show_column_names = FALSE, row_dend_side = "right", 
        show_column_dend = FALSE, column_title = "pairwise correlation between genes",
        heatmap_legend_param = list(title = "Correlation"))
ht_list = draw(ht_list, main_heatmap = "cor")
decorate_column_dend("scaled_expr", {
    tree = column_dend(ht_list)$scaled_expr
    ind = cutree(as.hclust(tree), k = 2)[order.dendrogram(tree)]

    first_index = function(l) which(l)[1]
    last_index = function(l) { x = which(l); x[length(x)] }
    x1 = c(first_index(ind == 1), first_index(ind == 2)) - 1
    x2 = c(last_index(ind == 1), last_index(ind == 2))
    grid.rect(x = x1/length(ind), width = (x2 - x1)/length(ind), just = "left",
        default.units = "npc", gp = gpar(fill = c("#FF000040", "#00FF0040"), col = NA))
})


# only keep the positive correlation
d = as.matrix(d0)
d[d<0] = 1e-10
dneg = -as.matrix(d0)
dneg[dneg<0] = 1e-10

#####################################################
############  NMF decomposition ################
set.seed(2019)

# define NMF function
nmf <- function(V, k, max_iter = 1000, tol = 1e-4) {
  # Initialize W and H with random values
  n <- dim(V)[1]
  m <- dim(V)[2]
  W <- matrix(runif(n * k), nrow = n)
  H <- matrix(runif(k * m), nrow = k)
  
  for (i in 1:max_iter) {
    # Update H
    WH <- W %*% H
    H <- H * ((t(W) %*% (V / (WH + 1e-10))) / (t(W) %*% matrix(1, nrow = n, ncol = m)))
    
    # Update W
    WH <- W %*% H
    W <- W * (((V / (WH + 1e-10)) %*% t(H)) / (matrix(1, nrow = n, ncol = m) %*% t(H)))
    
    # Normalize W and H
    if (i < 100) {
      W <- W / max(W)
      H <- H / max(H)
    }

    print(c(max(W), max(H)))
    # Check convergence
    WH <- W %*% H
    obj <- sum(V * log(1 / (WH + 1e-10)) + WH)
    if (i > 1 && abs(obj - prev_obj) < tol) {
      break
    }
    prev_obj <- obj
  }
  
  list(W = W, H = H)
}


# define NMF function
nmf <- function(V, k, max_iter = 1000, tol = 1e-4) {
  # Initialize W and H with random values
  n <- dim(V)[1]
  m <- dim(V)[2]
  W <- matrix(runif(n * k), nrow = n)
  H <- matrix(runif(k * m), nrow = k)
  
  for (i in 1:max_iter) {
    # Update H
    WH <- W %*% H
    H <- H * ((t(W) %*% (V / (WH + 1e-10))) / (t(W) %*% matrix(1, nrow = n, ncol = m)))
    
    # Update W
    WH <- W %*% H
    W <- W * (((V / (WH + 1e-10)) %*% t(H)) / (matrix(1, nrow = n, ncol = m) %*% t(H)))
    
    # Normalize W and H
    if (i < round(max_iter*0.5)) {
      W <- W / max(W)
      H <- H / max(H)
    }

    # Check convergence
    WH <- W %*% H
    obj <- sum(V * log(1 / (WH + 1e-10)) + WH)
    if (i > 1 && abs(obj - prev_obj) < tol) {
      break
    }
    prev_obj <- obj
  }
  
  list(W = W, H = H)
}

# decide the number of components based on the BIC
set.seed(2019)
BIC_vec = c()
for (i in seq(2, 20, by = 1)) {
  print(i)
    sparse_nmf_result <- nmf(d, k = i)
    W <- sparse_nmf_result$W
    H <- sparse_nmf_result$H
    reconstructed_whitened_R <- W %*% H
    # get BIC of the model
    BIC = nrow(d) * ncol(d) * log(sum((reconstructed_whitened_R - d)^2)) + i * (nrow(d) + ncol(d)) * log(nrow(d) * ncol(d))
    print(BIC)
    BIC_vec = c(BIC_vec, BIC)
}

#  NMF decomposition
set.seed(2019)
n_comp = 14
nmf_result <- nmf(d, k = n_comp)
nmf_result_neg <- nmf(dneg, k = n_comp)

# Extract the independent components (ICs) and the mixing matrix
nmf_result_W <- nmf_result$W
nmf_result_H <- nmf_result$H
# negative
nmf_result_W_neg <- nmf_result_neg$W
nmf_result_H_neg <- nmf_result_neg$H


# mkdir
dir.create('NMF_reconstruction')

# plot the BIC
pdf('NMF_reconstruction/BIC.pdf', width = 6, height = 6)
plot(seq(2, 20, by = 1), BIC_vec, type = 'p', xlab = 'Number of components', ylab = 'BIC', main = 'BIC vs. Number of components')
lines(seq(2, 20, by = 1), BIC_vec, type = 'b')
dev.off()

# plot correlation matrix
breaksList = seq(-1, 1, by = 0.001)
my_colorbar=colorRampPalette(c('blue', 'white', 'red'))(n = length(breaksList))
pheatmap(d0, cluster_rows=F, cluster_cols=F, legend=F, show_rownames=F, show_colnames=F, color=my_colorbar, breaks=breaksList, filename='NMF_reconstruction/HBA1.Hba-a1.cor.heatmap.png.cor.mat.txt.0.cor.mat.png')

# plot reconstructed correlation matrix
reconstructed_d <- nmf_result_W %*% nmf_result_H
pheatmap(reconstructed_d, cluster_rows=F, cluster_cols=F, legend=F, show_rownames=F, show_colnames=F, color=my_colorbar, breaks=breaksList, filename='NMF_reconstruction/HBA1.Hba-a1.cor.heatmap.png.cor.mat.txt.0.reconstructed_d.heatmap.png')

reconstructed_d_mm10 <- nmf_result_W %*% t(nmf_result_W)
reconstructed_d_mm10_neg <- -nmf_result_W_neg %*% t(nmf_result_W_neg)
pheatmap(reconstructed_d_mm10+reconstructed_d_mm10_neg, cluster_rows=F, cluster_cols=F, legend=F, show_rownames=F, show_colnames=F, filename='NMF_reconstruction/HBA1.Hba-a1.cor.heatmap.png.cor.mat.txt.0.reconstructed_d_mm10.heatmap.png')
reconstructed_d_hg38 <- t(nmf_result_H) %*% nmf_result_H
reconstructed_d_hg38_neg <- -t(nmf_result_H_neg) %*% nmf_result_H_neg
pheatmap(reconstructed_d_hg38+reconstructed_d_hg38_neg, cluster_rows=F, cluster_cols=F, legend=F, show_rownames=F, show_colnames=F,  filename='NMF_reconstruction/HBA1.Hba-a1.cor.heatmap.png.cor.mat.txt.0.reconstructed_d_hg38.heatmap.png')

# cluster factor matrix
nmf_W_H = rbind(nmf_result_W, t(nmf_result_H))
nmf_W_H_order = hclust(dist(t(nmf_W_H)), method = 'average')$order


# plot NMFs_mm10 heatmap
pheatmap(nmf_result_W, cluster_rows=F, cluster_cols=F, width = 2, height = 6, legend=F, show_rownames=F, show_colnames=F, filename='NMF_reconstruction/HBA1.Hba-a1.cor.heatmap.png.cor.mat.txt.NMFs_mm10.heatmap.png')

# plot NMFs_hg38 heatmap
pheatmap(t(nmf_result_H), cluster_rows=F, cluster_cols=F, width = 2, height = 6, legend=F, show_rownames=F, show_colnames=F, filename='NMF_reconstruction/HBA1.Hba-a1.cor.heatmap.png.cor.mat.txt.NMFs_hg38.heatmap.png')


# plot NMF i reconstruction
for (i in 1:n_comp) {
    print(i)
    png(paste0('NMF_reconstruction/', 'HBA1.Hba-a1.cor.heatmap.png.cor.mat.txt.NMF', i,'.png'), units="in", res=500)
    pheatmap(cbind(nmf_result_W[,i]) %*% rbind(nmf_result_H[i,]) , cluster_rows=F, cluster_cols=F, legend=F, show_rownames=F, show_colnames=F)
    dev.off()
}

#####################################################


# NMF matrices hg38 for shiny App
nmf_result_H_2 = matrix(0, nrow = nrow(nmf_result_H), ncol = ncol(nmf_result_H))
nmf_result_H_2[2,] = nmf_result_H[2,]
nmf_result_H_5 = matrix(0, nrow = nrow(nmf_result_H), ncol = ncol(nmf_result_H))
nmf_result_H_5[5,] = nmf_result_H[5,]
nmf_result_H_10 = matrix(0, nrow = nrow(nmf_result_H), ncol = ncol(nmf_result_H))
nmf_result_H_10[10,] = nmf_result_H[10,]

# NMF matrices mm10 for shiny App
nmf_result_W_2 = matrix(0, nrow = nrow(nmf_result_W), ncol = ncol(nmf_result_W))
nmf_result_W_2[,2] = nmf_result_W[,2]
nmf_result_W_5 = matrix(0, nrow = nrow(nmf_result_W), ncol = ncol(nmf_result_W))
nmf_result_W_5[,5] = nmf_result_W[,5]
nmf_result_W_10 = matrix(0, nrow = nrow(nmf_result_W), ncol = ncol(nmf_result_W))
nmf_result_W_10[,10] = nmf_result_W[,10]


library(ComplexHeatmap)
library(InteractiveComplexHeatmap)
library(shiny)
library(circlize)


# set colorbar
breaks <- seq(-1, 1, length.out = 101)
colors <- colorRamp2(breaks, colorRampPalette(c("blue", "white", "red"))(length(breaks)))


#
cor_mat1 = Heatmap(d0, name = "cor_mat1",
    show_row_names = F, show_column_names = F, cluster_columns =F, cluster_rows = F, width=1000, col = colors)
#
cor_mat2 = Heatmap(t(d0), name = "cor_mat2",
    show_row_names = F, show_column_names = F, cluster_columns =F, cluster_rows = F, width=1000, col = colors)
#
cor_mat3 = Heatmap(reconstructed_d_mm10+reconstructed_d_mm10_neg, name = "cor_mat_mm10",
    show_row_names = F, show_column_names = F, cluster_columns =F, cluster_rows = F, width=1000, col = colors)
#
cor_mat4 = Heatmap(reconstructed_d_hg38+reconstructed_d_hg38_neg, name = "cor_mat_hg38",
    show_row_names = F, show_column_names = F, cluster_columns =F, cluster_rows = F, width=1000, col = colors)

# plot NMFs_mm10 heatmap
mm10_NMF = Heatmap(nmf_result_W, name = "mm10_NMF", show_row_names = F, show_column_names = F, cluster_columns =F, cluster_rows = F, width=200)
# plot NMFs_hg38 heatmap
hg38_NMF = Heatmap(t(nmf_result_H), name = "hg38_NMF", show_row_names = F, show_column_names = F, cluster_columns =F, cluster_rows = F, width=200)


# add two heatmaps together to get interactive heatmap
ht1 = cor_mat1+mm10_NMF
ht2 = cor_mat2+hg38_NMF
ht3 = cor_mat3+mm10_NMF
ht4 = cor_mat4+hg38_NMF


# plot two interactive heatmaps
ui = fluidPage(
    h3("The first heatmap"),
    InteractiveComplexHeatmapOutput("ht1"),
    hr(),
    h3("The second heatmap"),
    InteractiveComplexHeatmapOutput("ht2"),
    hr(),
    h3("The mm10 heatmap"),
    InteractiveComplexHeatmapOutput("ht3"),
    hr(),
    h3("The hg38 heatmap"),
    InteractiveComplexHeatmapOutput("ht4"),

  titlePanel("Simple 2D Scatterplot"),  
  sidebarLayout(
    sidebarPanel(
      numericInput("n_points", "Number of points:", value = 100, min = 1),
      numericInput("correlation", "Correlation:", value = 0.5, min = -1, max = 1)
    ),
    mainPanel(
      plotOutput("scatterplot")
    )
  )
)

server = function(input, output, session) {
    makeInteractiveComplexHeatmap(input, output, session, ht1, "ht1")
    makeInteractiveComplexHeatmap(input, output, session, ht2, "ht2")
    makeInteractiveComplexHeatmap(input, output, session, ht3, "ht3")
    makeInteractiveComplexHeatmap(input, output, session, ht4, "ht4")
    output$scatterplot <- renderPlot({
        # Generate random data
        set.seed(42)
        x <- rnorm(input$n_points)
        y <- rnorm(input$n_points, mean = input$correlation * x)
        data <- data.frame(x = x, y = y)
        
        # Create scatterplot using ggplot2
        ggplot(data, aes(x = x, y = y)) +
        geom_point() +
        theme_minimal()
    })
}

shiny::shinyApp(ui, server)


ui <- fluidPage(
    titlePanel("Cross species IDEAS state correlation matrix decomposition:"),

    fluidRow(
        column(1, plotOutput("plot1a", width = "100px", height = "100px")),
        column(3, plotOutput("plot1b", width = "300px", height = "100px")),
        column(1, plotOutput("plot1c", width = "100px", height = "100px")),
        column(3, plotOutput("plot1d", width = "300px", height = "100px")),
        column(1, plotOutput("plot1e", width = "100px", height = "100px")),
        column(3, plotOutput("plot1f", width = "300px", height = "100px"))
    ),
    fluidRow(
        column(1, plotOutput("plot2a", width = "100px", height = "300px")),
        column(3, plotOutput("plot2b", width = "300px", height = "300px")),
        column(1, plotOutput("plot2c", width = "100px", height = "300px")),
        column(3, plotOutput("plot2d", width = "300px", height = "300px")),
        column(1, plotOutput("plot2e", width = "100px", height = "300px")),
        column(3, plotOutput("plot2f", width = "300px", height = "300px"))
    ),
    fluidRow(
        column(12, h3("NMF decomposition:"))
    ),  
    fluidRow(
        column(1, plotOutput("plot3a", width = "100px", height = "100px")),
        column(3, plotOutput("plot3b", width = "300px", height = "100px")),
        column(1, plotOutput("plot3c0", width = "100px", height = "100px")),
        column(3, plotOutput("plot3c", width = "300px", height = "100px")),
        column(1, plotOutput("plot3d0", width = "100px", height = "100px")),
        column(3, plotOutput("plot3d", width = "300px", height = "100px")),
    ),
    fluidRow(
        column(1, plotOutput("plot4a", width = "100px", height = "300px")),
        column(3, plotOutput("plot4b", width = "300px", height = "300px")),
        column(1, plotOutput("plot4c", width = "100px", height = "300px")),
        column(3, plotOutput("plot4d", width = "300px", height = "300px")),
        column(1, plotOutput("plot4e", width = "100px", height = "300px")),
        column(3, plotOutput("plot4f", width = "300px", height = "300px"))
    ),
    fluidRow(
        column(1, plotOutput("plot5a", width = "100px", height = "100px")),
        column(3, plotOutput("plot5b", width = "300px", height = "100px")),
        column(1, plotOutput("plot5c0", width = "100px", height = "100px")),
        column(3, plotOutput("plot5c", width = "300px", height = "100px")),
        column(1, plotOutput("plot5d0", width = "100px", height = "100px")),
        column(3, plotOutput("plot5d", width = "300px", height = "100px")),
    ),
    fluidRow(
        column(1, plotOutput("plot6a", width = "100px", height = "300px")),
        column(3, plotOutput("plot6b", width = "300px", height = "300px")),
        column(1, plotOutput("plot6c", width = "100px", height = "300px")),
        column(3, plotOutput("plot6d", width = "300px", height = "300px")),
        column(1, plotOutput("plot6e", width = "100px", height = "300px")),
        column(3, plotOutput("plot6f", width = "300px", height = "300px"))
    ),
    fluidRow(
        column(1, plotOutput("plot7a", width = "100px", height = "100px")),
        column(3, plotOutput("plot7b", width = "300px", height = "100px")),
        column(1, plotOutput("plot7c0", width = "100px", height = "100px")),
        column(3, plotOutput("plot7c", width = "300px", height = "100px")),
        column(1, plotOutput("plot7d0", width = "100px", height = "100px")),
        column(3, plotOutput("plot7d", width = "300px", height = "100px")),
    ),
    fluidRow(
        column(1, plotOutput("plot8a", width = "100px", height = "300px")),
        column(3, plotOutput("plot8b", width = "300px", height = "300px")),
        column(1, plotOutput("plot8c", width = "100px", height = "300px")),
        column(3, plotOutput("plot8d", width = "300px", height = "300px")),
        column(1, plotOutput("plot8e", width = "100px", height = "300px")),
        column(3, plotOutput("plot8f", width = "300px", height = "300px"))
    ),
    fluidRow(
        column(1, plotOutput("plot9a", width = "100px", height = "100px")),
        column(3, plotOutput("plot9b", width = "300px", height = "100px")),
        column(1, plotOutput("plot9c0", width = "100px", height = "100px")),
        column(3, plotOutput("plot9c", width = "300px", height = "100px")),
        column(1, plotOutput("plot9d0", width = "100px", height = "100px")),
        column(3, plotOutput("plot9d", width = "300px", height = "100px")),
    ),
    fluidRow(
        column(1, plotOutput("plot10a", width = "100px", height = "300px")),
        column(3, plotOutput("plot10b", width = "300px", height = "300px")),
        column(1, plotOutput("plot10c", width = "100px", height = "300px")),
        column(3, plotOutput("plot10d", width = "300px", height = "300px")),
        column(1, plotOutput("plot10e", width = "100px", height = "300px")),
        column(3, plotOutput("plot810f", width = "300px", height = "300px"))
    ),
    fluidRow(
        column(1, plotOutput("plot11a", width = "100px", height = "100px")),
        column(3, plotOutput("plot11b", width = "300px", height = "100px")),
        column(1, plotOutput("plot11c0", width = "100px", height = "100px")),
        column(3, plotOutput("plot11c", width = "300px", height = "100px")),
        column(1, plotOutput("plot11d0", width = "100px", height = "100px")),
        column(3, plotOutput("plot11d", width = "300px", height = "100px")),
    ),
    fluidRow(
        column(1, plotOutput("plot12a", width = "100px", height = "300px")),
        column(3, plotOutput("plot12b", width = "300px", height = "300px")),
        column(1, plotOutput("plot12c", width = "100px", height = "300px")),
        column(3, plotOutput("plot12d", width = "300px", height = "300px")),
        column(1, plotOutput("plot12e", width = "100px", height = "300px")),
        column(3, plotOutput("plot12f", width = "300px", height = "300px"))
    )
)


NMF_i = 5
nmf_result_H_i = matrix(0, nrow = nrow(nmf_result_H), ncol = ncol(nmf_result_H))
nmf_result_H_i[NMF_i,] = nmf_result_H[NMF_i,] 
nmf_result_W_i = matrix(0, nrow = nrow(nmf_result_W), ncol = ncol(nmf_result_W))
nmf_result_W_i[,NMF_i] = nmf_result_W[,NMF_i]


server <- function(input, output) {
    # NMFs mm10
    output$plot2a <- renderPlot({
        pheatmap(nmf_result_W, cluster_rows=F, cluster_cols=F, width = 2, height = 6, legend=F, show_rownames=F, show_colnames=F, main='NMFs_mm10')
    }) 
    output$plot1f <- renderPlot({
        pheatmap(t(nmf_result_W), cluster_rows=F, cluster_cols=F, width = 6, height = 2, legend=F, show_rownames=F, show_colnames=F, main='NMFs_mm10')
    })
    output$plot2e <- renderPlot({
        pheatmap(nmf_result_W, cluster_rows=F, cluster_cols=F, width = 6, height = 2, legend=F, show_rownames=F, show_colnames=F, main='NMFs_mm10')
    })

    # NMFs hg38
    output$plot1b <- renderPlot({
        pheatmap(nmf_result_H, cluster_rows=F, cluster_cols=F, width = 6, height = 2, legend=F, show_rownames=F, show_colnames=F, main='NMFs_hg38')
    })
    output$plot2c <- renderPlot({
        pheatmap(t(nmf_result_H), cluster_rows=F, cluster_cols=F, width = 6, height = 2, legend=F, show_rownames=F, show_colnames=F, main='NMFs_hg38')
    })
    output$plot1d <- renderPlot({
        pheatmap(nmf_result_H, cluster_rows=F, cluster_cols=F, width = 6, height = 2, legend=F, show_rownames=F, show_colnames=F, main='NMFs_hg38')
    })

    output$plot2b <- renderPlot({
        pheatmap(reconstructed_d, cluster_rows=F, cluster_cols=F, legend=F, show_rownames=F, show_colnames=F, color=my_colorbar, breaks=breaksList, main='Reconstructed correlation matrix')
    })
    output$plot2d <- renderPlot({
        pheatmap(reconstructed_d_hg38, cluster_rows=F, cluster_cols=F, legend=F, show_rownames=F, show_colnames=F, main='Reconstructed correlation hg38 matrix')
    })
    output$plot2f <- renderPlot({
        pheatmap(reconstructed_d_mm10, cluster_rows=F, cluster_cols=F, legend=F, show_rownames=F, show_colnames=F, main='Reconstructed correlation mm10 matrix')
    })


    output$plot3b <- renderPlot({
        pheatmap(nmf_result_H_2, cluster_rows=F, cluster_cols=F, width = 6, height = 2, legend=F, show_rownames=F, show_colnames=F, main='NMF_2_hg38')
    })
    output$plot3c <- renderPlot({
        pheatmap(nmf_result_H_2, cluster_rows=F, cluster_cols=F, width = 6, height = 2, legend=F, show_rownames=F, show_colnames=F, main='NMF_2_hg38')
    })
    output$plot3d <- renderPlot({
        pheatmap(t(nmf_result_W_2), cluster_rows=F, cluster_cols=F, width = 6, height = 2, legend=F, show_rownames=F, show_colnames=F, main='NMF_2_mm10')
    })

    output$plot4a <- renderPlot({
        pheatmap(cbind(nmf_result_W_2), cluster_rows=F, cluster_cols=F, width = 2, height = 6, legend=F, show_rownames=F, show_colnames=F, main='NMF_2_mm10')
    }) 
    output$plot4c <- renderPlot({
        pheatmap(cbind(t(nmf_result_H_2)), cluster_rows=F, cluster_cols=F, width = 2, height = 6, legend=F, show_rownames=F, show_colnames=F, main='NMF_2_hg38')
    }) 
    output$plot4e <- renderPlot({
        pheatmap(cbind(nmf_result_W_2), cluster_rows=F, cluster_cols=F, width = 2, height = 6, legend=F, show_rownames=F, show_colnames=F, main='NMF_2_mm10')
    })

    output$plot4b <- renderPlot({
        pheatmap(cbind(nmf_result_W[,2]) %*% rbind(nmf_result_H[2,]) , cluster_rows=F, cluster_cols=F, legend=F, show_rownames=F, show_colnames=F, main=paste0('NMF', 2, ': HBA enhancer'))
    })
    output$plot4d <- renderPlot({
        pheatmap(cbind(nmf_result_H[2,]) %*% rbind(nmf_result_H[2,]) , cluster_rows=F, cluster_cols=F, legend=F, show_rownames=F, show_colnames=F, main=paste0('NMF', 2, ': HBA enhancer hg38'))
    })
    output$plot4f <- renderPlot({
        pheatmap(cbind(nmf_result_W[,2]) %*% rbind(nmf_result_W[,2]) , cluster_rows=F, cluster_cols=F, legend=F, show_rownames=F, show_colnames=F, main=paste0('NMF', 2, ': HBA enhancer mm10'))
    })


    output$plot5b <- renderPlot({
        pheatmap(nmf_result_H_10, cluster_rows=F, cluster_cols=F, width = 6, height = 2, legend=F, show_rownames=F, show_colnames=F, main='NMF_10_hg38')
    })
    output$plot5c <- renderPlot({
        pheatmap(nmf_result_H_10, cluster_rows=F, cluster_cols=F, width = 6, height = 2, legend=F, show_rownames=F, show_colnames=F, main='NMF_10_hg38')
    })
    output$plot5d <- renderPlot({
        pheatmap(t(nmf_result_W_10), cluster_rows=F, cluster_cols=F, width = 6, height = 2, legend=F, show_rownames=F, show_colnames=F, main='NMF_10_mm10')
    })

    output$plot6a <- renderPlot({
        pheatmap(cbind(nmf_result_W_10), cluster_rows=F, cluster_cols=F, width = 2, height = 6, legend=F, show_rownames=F, show_colnames=F, main='NMF_10_mm10')
    }) 
    output$plot6c <- renderPlot({
        pheatmap(cbind(t(nmf_result_H_10)), cluster_rows=F, cluster_cols=F, width = 2, height = 6, legend=F, show_rownames=F, show_colnames=F, main='NMF_10_hg38')
    }) 
    output$plot6e <- renderPlot({
        pheatmap(cbind(nmf_result_W_10), cluster_rows=F, cluster_cols=F, width = 2, height = 6, legend=F, show_rownames=F, show_colnames=F, main='NMF_10_mm10')
    })

    output$plot6b <- renderPlot({
        pheatmap(cbind(nmf_result_W[,10]) %*% rbind(nmf_result_H[10,]) , cluster_rows=F, cluster_cols=F, legend=F, show_rownames=F, show_colnames=F, main=paste0('NMF', 10, ': common promoter'))
    })
    output$plot6d <- renderPlot({
        pheatmap(cbind(nmf_result_H[10,]) %*% rbind(nmf_result_H[10,]) , cluster_rows=F, cluster_cols=F, legend=F, show_rownames=F, show_colnames=F, main=paste0('NMF', 10, ': common promoter hg38'))
    })
    output$plot6f <- renderPlot({
        pheatmap(cbind(nmf_result_W[,10]) %*% rbind(nmf_result_W[,10]) , cluster_rows=F, cluster_cols=F, legend=F, show_rownames=F, show_colnames=F, main=paste0('NMF', 10, ': common promoter mm10'))
    })


    output$plot7b <- renderPlot({
        pheatmap(nmf_result_H_i, cluster_rows=F, cluster_cols=F, width = 6, height = 2, legend=F, show_rownames=F, show_colnames=F, main=paste0('NMF_',NMF_i,'_hg38'))
    })
    output$plot7c <- renderPlot({
        pheatmap(nmf_result_H_i, cluster_rows=F, cluster_cols=F, width = 6, height = 2, legend=F, show_rownames=F, show_colnames=F, main=paste0('NMF_',NMF_i,'_hg38'))
    })
    output$plot7d <- renderPlot({
        pheatmap(t(nmf_result_W_i), cluster_rows=F, cluster_cols=F, width = 6, height = 2, legend=F, show_rownames=F, show_colnames=F, main=paste0('NMF_',NMF_i,'_mm10'))
    })

    output$plot8a <- renderPlot({
        pheatmap(cbind(nmf_result_W_i), cluster_rows=F, cluster_cols=F, width = 2, height = 6, legend=F, show_rownames=F, show_colnames=F, main=paste0('NMF_',NMF_i,'_mm10'))
    }) 
    output$plot8c <- renderPlot({
        pheatmap(cbind(t(nmf_result_H_i)), cluster_rows=F, cluster_cols=F, width = 2, height = 6, legend=F, show_rownames=F, show_colnames=F, main=paste0('NMF_',NMF_i,'_hg38'))
    }) 
    output$plot8e <- renderPlot({
        pheatmap(cbind(nmf_result_W_i), cluster_rows=F, cluster_cols=F, width = 2, height = 6, legend=F, show_rownames=F, show_colnames=F, main=paste0('NMF_',NMF_i,'_mm10'))
    })

    output$plot8b <- renderPlot({
        pheatmap(cbind(nmf_result_W[,NMF_i]) %*% rbind(nmf_result_H[NMF_i,]) , cluster_rows=F, cluster_cols=F, legend=F, show_rownames=F, show_colnames=F, main=paste0('NMF', NMF_i))
    })
    output$plot8d <- renderPlot({
        pheatmap(cbind(nmf_result_H[NMF_i,]) %*% rbind(nmf_result_H[NMF_i,]) , cluster_rows=F, cluster_cols=F, legend=F, show_rownames=F, show_colnames=F, main=paste0('NMF', NMF_i, ' hg38'))
    })
    output$plot8f <- renderPlot({
        pheatmap(cbind(nmf_result_W[,NMF_i]) %*% rbind(nmf_result_W[,NMF_i]) , cluster_rows=F, cluster_cols=F, legend=F, show_rownames=F, show_colnames=F, main=paste0('NMF', NMF_i, ' mm10'))
    })

}


shinyApp(ui = ui, server = server)
```
