## Supplementary material for "Interspecies regulatory landscapes and elements revealed by novel joint systematic integration of human and mouse blood cell epigenomes": Scripts for IDEAS modeling of epigenetic states jointly in human and mouse: reverse_seq.R

```
args = commandArgs(trailingOnly=TRUE)

input_seq_file = args[1]
output = args[2]
#input_seq_file = 'S3V2_IDEAS_mm10_r3_withHg38Mm10prior.state.AVE.Gata1.bed.r.seq'
d = read.table(input_seq_file, header=F)
dr = d[dim(d)[1]:1,]
write.table(dr, output, quote=F, sep='\t', col.names=F, row.names=F)
```
