## Supplementary material for "Interspecies regulatory landscapes and elements revealed by novel joint systematic integration of human and mouse blood cell epigenomes": Scripts for IDEAS modeling of epigenetic states jointly in human and mouse: compare_two_para.R

```
### get parameters
args = commandArgs(trailingOnly=TRUE)

library(pheatmap)

parafile1 = args[1]
parafile2 = args[2]
feature_num = as.numeric(args[3])
outputfile = args[4]
source_createGenomeTracks = args[5]

print(parafile1)
print(parafile2)
print(outputfile)
print(source_createGenomeTracks)
source(source_createGenomeTracks)


pdf(outputfile, height=14, width=7)
statecolor = NULL
markcolor = NULL
show=TRUE
fout=NULL
sortstate=TRUE
cols=c("white","dark blue")
scale=F
print('read parafiles')
	x1=read.table(parafile1,comment="!",header=T);
	x2=read.table(parafile2,comment="!",header=T);


used_col_id1 = 2:(1+feature_num)
x1[,used_col_id1] = x1[,used_col_id1][,order(colnames(x1)[used_col_id1])]
colnames(x1)[used_col_id1] = colnames(x1)[used_col_id1][order(colnames(x1)[used_col_id1])]

used_col_id2 = 2:(1+feature_num)
x2[,used_col_id2] = x2[,used_col_id2][,order(colnames(x2)[used_col_id2])]
colnames(x2)[used_col_id2] = colnames(x2)[used_col_id2][order(colnames(x2)[used_col_id2])]

used_col_id1 = (2+feature_num):dim(x1)[2]
colnames(x1)[used_col_id1] = (2+feature_num):dim(x1)[2]

used_col_id2 = (2+feature_num):dim(x2)[2]
colnames(x2)[used_col_id2] = (2+feature_num):dim(x1)[2]

print(cbind(colnames(x1), colnames(x2)))

print(x1[1:10,1:5])
print(x2[1:10,1:5])
	x = rbind(x1, x2)
print('rbind para files')
	k=dim(x)[2];
	l=dim(x)[1];
	l1=dim(x1)[1];
	l2=dim(x2)[1];

	p=(sqrt(9+8*(k-1))-3)/2;
	m=as.matrix(x[,1+1:p]/x[,1]);
	colnames(m) = colnames(x)[1+1:p];
	#m0 = m[,-6]
	marks=colnames(m);
	rn1 = paste('H:',1:l1-1," (",round(x1[,1]/sum(x1[,1])*10000)/100,"%)",sep="");
	rn2 = paste('M:',1:l2-1," (",round(x2[,1]/sum(x2[,1])*10000)/100,"%)",sep="");
	#rownames(m)=paste(1:l-1," (",round(x[,1]/sum(x[,1])*10000)/100,"%)",sep="");
	rownames(m)=c(rn1, rn2)
if(sortstate)
{
o_tree_dist = dist(m)
o_tree=hclust(o_tree_dist,method="ward.D2");

o=o_tree$order;

### plot tree


m=m[o,];
if(length(statecolor) != 0)
{	statecolor=statecolor[o,];
}
}
type = c(rep(1, length(rn1)), rep(2, length(rn2)))
orn1 = o[type==1]
orn2 = o[type==2]
om=m;
if(scale)
{	m = t((t(m) - apply(m,2,min))/(apply(m,2,max)-apply(m,2,min)+1e-10));
}
	if(length(fout)!=0)
	{	pdf(fout);	}	
	par(mar=c(6,1,1,6));
	rg=range(m);
	colors=0:100/100*(rg[2]-rg[1])+rg[1];
        my_palette=colorRampPalette(cols)(n=100);
	defpalette=palette(my_palette);
if(show)
{
	plot(NA,NA,xlim=c(0,p+0.7),ylim=c(0,l),xaxt="n",yaxt="n",xlab=NA,ylab=NA,frame.plot=F);
	axis(1,at=1:p-0.5,labels=colnames(m),las=2);
	axis(4,at=1:l-0.5,labels=rownames(m),las=2);
	rect(rep(1:p-1,l),rep(1:l-1,each=p),rep(1:p,l),rep(1:l,each=p),col=round((t(m)-rg[1])/(rg[2]-rg[1])*100));#,border=NA);
}
#if(scale)
#{	m = om;
#}
	if(length(statecolor)==0)
	{	if(length(markcolor)==0)
		{	markcolor=t(col2rgb(terrain.colors(ceiling(p))[1:p]));	
			for(i in 1:length(marks))
			{	if(regexpr("h3k4me3",tolower(marks[i]))>0)
				{	markcolor[i,]=c(255,0,0);	}
				if(regexpr("h3k4me2",tolower(marks[i]))>0)
				{	markcolor[i,]=c(250,100,0);	}
				if(regexpr("h3k4me1",tolower(marks[i]))>0)
				{	markcolor[i,]=c(250,250,0);	}
				if(regexpr("h3k36me3",tolower(marks[i]))>0)
				{	markcolor[i,]=c(0,150,0);	}
				if(regexpr("h2a",tolower(marks[i]))>0)
				{	markcolor[i,]=c(0,150,150);	}
				if(regexpr("dnase",tolower(marks[i]))>0)
				{	markcolor[i,]=c(0,200,200);	}
				if(regexpr("atac",tolower(marks[i]))>0)
				{	markcolor[i,]=c(200,50,150);	}
				if(regexpr("dnase",tolower(marks[i]))>0)
				{	markcolor[i,]=c(200,50,150);	}
				if(regexpr("h3k9ac",tolower(marks[i]))>0)
				{	markcolor[i,]=c(250,0,200);	}
				if(regexpr("h3k9me3",tolower(marks[i]))>0)
				{	markcolor[i,]=c(100,100,100);	}
				if(regexpr("h3k27ac",tolower(marks[i]))>0)
				{	markcolor[i,]=c(250,150,0);	}
				if(regexpr("h3k27me3",tolower(marks[i]))>0)
				{	markcolor[i,]=c(0,0,225);	}
				if(regexpr("h3k79me2",tolower(marks[i]))>0)
				{	markcolor[i,]=c(200,0,200);	}
				if(regexpr("h4k20me1",tolower(marks[i]))>0)
				{	markcolor[i,]=c(50,200,50);	}
				if(regexpr("ctcf",tolower(marks[i]))>0)
				{	markcolor[i,]=c(200,0,250);	}
				if(regexpr("wgbs",tolower(marks[i]))>0)
				{	markcolor[i,]=c(30,144,255);	}
			}
		}
		statecolor=array(stateColor(m,markcolor),dim=c(dim(m)[1],2));
	}
	if(show)
	{	rect(rep(p+0.2,l),1:l-0.8,rep(p+0.8,l),1:l-0.2,col=statecolor[,2]);
	}
	if(sortstate)	statecolor[o,]=statecolor;
	palette(defpalette);
	if(length(fout)!=0)
	{	dev.off();	}

dev.off()

pdf(paste(outputfile,'.tree.pdf', sep=''), height=7, width=14)
dhc <- as.dendrogram(o_tree)
plot(as.dendrogram(o_tree))
dev.off()

pdf(paste(outputfile,'.dist.pdf', sep=''), height=14, width=14)
my_colorbar=colorRampPalette(c('red', 'white', 'blue'))(n = 128)
print(c(rn1, rn2))
o_tree_dist_mat = as.matrix(o_tree_dist)
print(dim(o_tree_dist_mat))
colnames(o_tree_dist_mat) = c(rn1, rn2)
rownames(o_tree_dist_mat) = c(rn1, rn2)
pheatmap(o_tree_dist_mat[rev(o),rev(o)], color=my_colorbar, cluster_cols = F,cluster_rows=F, annotation_names_row=F, annotation_names_col=T )
dev.off()


#	return(statecolor);
#}
```
