## Supplementary material for "Interspecies regulatory landscapes and elements revealed by novel joint systematic integration of human and mouse blood cell epigenomes": Scripts for IDEAS modeling of epigenetic states jointly in human and mouse: align_epistate.R

```
cd /homes1/gxiang/softwares/EpiAlign/Ccode/example/VISION_IDEAS_states/

args = commandArgs(trailingOnly=TRUE)

input_seq_file = args[1]
output = args[2]
input_seq_file = 'S3V2_IDEAS_mm10_r3_withHg38Mm10prior.state.AVE.Gata1.bed.r.seq'
d = read.table(input_seq_file, header=F)

ds = c()
dn = c()

d_i_pre = d[1,1]
d_i_pre_n = 1
ds = d_i_pre

for (i in 2:dim(d)[1]){
d_i = d[i,1]
if (d_i!=d_i_pre){
ds = c(ds, d_i)
dn = c(dn, d_i_pre_n)
d_i_pre_n = 1
d_i_pre = d_i
} else{
d_i_pre_n = d_i_pre_n+1
}
print(dim(cbind(ds[1:(length(ds)-1)],dn)))
}

dn = c(dn, d_i_pre_n)
write.table(cbind(ds,dn), output, quote=F, sep=' ', col.names=F, row.names=F)

args = commandArgs(trailingOnly=TRUE)

input_seq_file = args[1]
output = args[2]
input_seq_file = 'S3V2_IDEAS_mm10_r3_withHg38Mm10prior.state.AVE.Gata1.bed.r.seq'
d = read.table(input_seq_file, header=F)
dr = d[dim(d)[1]:1,]
write.table(dr, output, quote=F, sep='\t', col.names=F, row.names=F)


### get all genes
wget https://ftp.ebi.ac.uk/pub/databases/gencode/Gencode_human/release_39/gencode.v39.basic.annotation.gff3.gz
gunzip gencode.v39.basic.annotation.gff3.gz
cat gencode.v39.basic.annotation.gff3 | awk -F 'gene_type=' '{print $2}' | awk -F ';' '{print $1}' > gene_types.txt
cat gencode.v39.basic.annotation.gff3 | awk -F 'gene_name=' '{print $2}' | awk -F ';' '{print $1}' > gene_names.txt
paste gene_types.txt gene_names.txt gencode.v39.basic.annotation.gff3 | awk -F '\t' -v OFS='\t' '{if ($1=="protein_coding" && $5=="gene") print $3,$6,$7,$9,$2}' > hg38.gene.bed
#paste gene_types.txt gene_names.txt gencode.v39.basic.annotation.gff3 | awk -F '\t' -v OFS='\t' '{if ($1=="protein_coding" && $5=="gene") print $0}' > hg38.gene.bed
rm gene_types.txt gene_names.txt

wget https://ftp.ebi.ac.uk/pub/databases/gencode/Gencode_mouse/release_M25/gencode.vM25.basic.annotation.gff3.gz
gunzip gencode.vM25.basic.annotation.gff3.gz
cat gencode.vM25.basic.annotation.gff3 | awk -F 'gene_type=' '{print $2}' | awk -F ';' '{print $1}' > gene_types.txt
cat gencode.vM25.basic.annotation.gff3 | awk -F 'gene_name=' '{print $2}' | awk -F ';' '{print $1}' > gene_names.txt
paste gene_types.txt gene_names.txt gencode.vM25.basic.annotation.gff3 | awk -F '\t' -v OFS='\t' '{if ($1=="protein_coding" && $5=="gene") print $3,$6,$7,$9,$2}' > mm10.gene.bed
#paste gene_types.txt gene_names.txt gencode.vM25.basic.annotation.gff3 | awk -F '\t' -v OFS='\t' '{if ($1=="protein_coding" && $5=="gene") print $0}' > mm10.gene.bed
rm gene_types.txt gene_names.txt


R
setwd('/homes1/gxiang/softwares/EpiAlign/Ccode/example/VISION_IDEAS_states/genes')
d1 = read.table('hg38.gene.bed', header=F)
d2 = read.table('mm10.gene.bed', header=F)
### remove genename multiplied genes
d1 = d1[is.element(as.character(d1[,5]), rownames(table(d1[,5]))[table(d1[,5])==1]),]
d2 = d2[is.element(as.character(d2[,5]), rownames(table(d2[,5]))[table(d2[,5])==1]),]
### get human mouse shared genes
d1s = d1[is.element(d1[,5], toupper(d2[,5])),]
d1s = d1s[order(d1s[,5]),]
d2s = d2[is.element(toupper(d2[,5]), (d1[,5])),]
d2s = d2s[order(d2s[,5]),]
get_tss_exp = function(ds, exp_win){
ds_tss = ds
ds_tss[ds[,4]=='+',3] = ds_tss[ds[,4]=='+',2]
ds_tss[ds[,4]=='-',2] = ds_tss[ds[,4]=='-',3]
ds_tss[,2] = ds_tss[,2]-exp_win
ds_tss[ds_tss[,2]<0,2] = 0 
ds_tss[,3] = ds_tss[,3]+exp_win
return(ds_tss)
}
### get tss exp bed
d1s_tss_exp = get_tss_exp(d1s, 100000)
d2s_tss_exp = get_tss_exp(d2s, 100000)
write.table(d1s_tss_exp, 'hg38.gene.tss.exp.bed', quote=F, col.names=F, row.names=F, sep='\t')
write.table(d2s_tss_exp, 'mm10.gene.tss.exp.bed', quote=F, col.names=F, row.names=F, sep='\t')


### get gene state ave
sbatch get_hg38.ave.sh
sbatch get_mm10.ave.sh
sbatch get_hg38.ERY_S002R5.sh
sbatch get_mm10.ERY_ad_r1.sh


state_bed='S3V2_IDEAS_hg38_r3_withHg38Mm10prior.state.AVE.bed'
while read chr start end st gn
do
echo $st
echo $chr $start $end $st $gn | awk -F ' ' -v OFS='\t' '{print $1,$2,$3,$4,$5}' > 'genes/hg38.gene.tss.exp.'$gn'.'$state_bed'.bed'
bedtools intersect -a $state_bed -b 'genes/hg38.gene.tss.exp.'$gn'.'$state_bed'.bed' -wa -u > 'genes/hg38.gene.tss.exp.'$gn'.'$state_bed'.state.bed'
if [[ $st == "+" ]]
then
cut -f4 'genes/hg38.gene.tss.exp.'$gn'.'$state_bed'.state.bed' > 'genes/hg38.gene.tss.exp.'$gn'.'$state_bed'.state.bed.seq'
fi
if [[ $st == "-" ]]
then
cut -f4 'genes/hg38.gene.tss.exp.'$gn'.'$state_bed'.state.bed' > 'genes/hg38.gene.tss.exp.'$gn'.'$state_bed'.state.bed.r.seq'
Rscript reverse_seq.R 'genes/hg38.gene.tss.exp.'$gn'.'$state_bed'.state.bed.r.seq' 'genes/hg38.gene.tss.exp.'$gn'.'$state_bed'.state.bed.seq'
rm 'genes/hg38.gene.tss.exp.'$gn'.'$state_bed'.state.bed.r.seq'
fi
Rscript compress_seq.R 'genes/hg38.gene.tss.exp.'$gn'.'$state_bed'.state.bed.seq' 'genes/hg38.gene.tss.exp.'$gn'.'$state_bed'.state.bed.Sseq'
rm 'genes/hg38.gene.tss.exp.'$gn'.'$state_bed'.state.bed'
rm 'genes/hg38.gene.tss.exp.'$gn'.'$state_bed'.state.bed.seq'
done < genes/hg38.gene.tss.exp.bed


cd /homes1/gxiang/softwares/EpiAlign/Ccode/example/VISION_IDEAS_states
state_bed='S3V2_IDEAS_hg38_r3_withHg38Mm10prior.state.ERY_S002R5.bed'
while read chr start end st gn
do
echo $st
echo $chr $start $end $st $gn | awk -F ' ' -v OFS='\t' '{print $1,$2,$3,$4,$5}' > 'genes/hg38.gene.tss.exp.'$gn'.'$state_bed'.bed'
bedtools intersect -a $state_bed -b 'genes/hg38.gene.tss.exp.'$gn'.'$state_bed'.bed' -wa -u > 'genes/hg38.gene.tss.exp.'$gn'.'$state_bed'.state.bed'
if [[ $st == "+" ]]
then
cut -f4 'genes/hg38.gene.tss.exp.'$gn'.'$state_bed'.state.bed' > 'genes/hg38.gene.tss.exp.'$gn'.'$state_bed'.state.bed.seq'
fi
if [[ $st == "-" ]]
then
cut -f4 'genes/hg38.gene.tss.exp.'$gn'.'$state_bed'.state.bed' > 'genes/hg38.gene.tss.exp.'$gn'.'$state_bed'.state.bed.r.seq'
Rscript reverse_seq.R 'genes/hg38.gene.tss.exp.'$gn'.'$state_bed'.state.bed.r.seq' 'genes/hg38.gene.tss.exp.'$gn'.'$state_bed'.state.bed.seq'
rm 'genes/hg38.gene.tss.exp.'$gn'.'$state_bed'.state.bed.r.seq'
fi
Rscript compress_seq.R 'genes/hg38.gene.tss.exp.'$gn'.'$state_bed'.state.bed.seq' 'genes/hg38.gene.tss.exp.'$gn'.'$state_bed'.state.bed.Sseq'
rm 'genes/hg38.gene.tss.exp.'$gn'.'$state_bed'.state.bed'
#mv 'genes/hg38.gene.tss.exp.'$gn'.'$state_bed'.state.bed.seq' genes/hg38_seq
#mv 'genes/hg38.gene.tss.exp.'$gn'.'$state_bed'.state.bed.Sseq' genes/hg38_Sseq
done < genes/hg38.gene.tss.exp.GATA1.bed


library(text.alignment)


state = read.table('06a_S3V2_IDEAS_hg38_r3_withHg38Mm10prior.para.modified.para', header=T)
state_mat_od = apply(state[,2:9],2,function(x) x/state[,1])

state_exp_num = (state[,1] %*% t(state[,1]))/sum(state[,1])
add_sm_exp = 1000
scale_mat = 1/(state_exp_num+add_sm_exp)*(mean(state_exp_num+add_sm_exp))
scale_mat[scale_mat<0.1] = 0.1
scale_mat[scale_mat>10] = 10

max(scale_mat)

state_mat_posneg = state_mat_od
state_mat_posneg[c(2,4,8,19,12),] = -state_mat_posneg[c(2,4,8,19,12),]

state_dist_od = as.matrix(dist(state_mat_posneg))

state_dist_adjstatefreq = (max(state_dist_od)-state_dist_od)


state_dist_align0a = (state_dist_adjstatefreq-median(state_dist_adjstatefreq))#*scale_mat

state_dist_align0 = (state_dist_adjstatefreq-median(state_dist_adjstatefreq))*scale_mat
state_dist_align0 = state_dist_align0/max(abs(state_dist_align0))

state_dist_align = state_dist_align0


state_dist_align = (-1+state_dist_adjstatefreq/max(state_dist_adjstatefreq))
diag(state_dist_align) = diag(state_dist_align0)


state_dist_od0 = as.matrix(dist(state_mat_od))

s1_table = s2_table = c(0,dim(state_dist_od0)[1])
for (i in 1:dim(state_dist_od0)[1]){
	s_i = i-1
	s1_table[i] = sum(s1==s_i)
	s2_table[i] = sum(s2==s_i)
}


s1_table = table(s1[,1])
s2_table = table(s2[,1])
s1_table_rowname = rownames(s1_table)
s2_table_rowname = rownames(s2_table)

for (i in 1:dim(state_dist_od0)[1]){
	s_i = toString(i-1)
	s1_table
}


s1 = read.table('genes/hg38.gene.tss.exp.GATA1.S3V2_IDEAS_hg38_r3_withHg38Mm10prior.state.AVE.bed.state.bed.seq')
s2 = read.table('genes/mm10.gene.tss.exp.Gata1.S3V2_IDEAS_mm10_r3_withHg38Mm10prior.state.AVE.bed.state.bed.seq')

s1a = s1
s2a = s2

for (i in 1:25){
	s1a[s1==(i-1)] = LETTERS[i]
	s2a[s2==(i-1)] = LETTERS[i]
}


s1s = paste(s1a[,1], collapse='')
s2s = paste(s2a[,1], collapse='')

chrX	48673045	48904704	GATA1	0	+
chrX	7871432	8066079	GATA1	0	-


state_dist = function(x1,x2){
id1 = which(tolower(LETTERS)==tolower(x1))
id2 = which(tolower(LETTERS)==tolower(x2))
dist_ij = (state_dist_align[id1,id2])*1
return(dist_ij)
}

sw_align1 = smith_waterman(s1s,s2s,similarity=state_dist, gap=0)


cbind(sw_align1$a$alignment$text, sw_align1$b$alignment$text)
sw_align1$sw


m <- text.alignment::smith_waterman_function(a = s1s, b = s2s, score_gap = -1, similarity = state_dist)


###################################################K#K#K#######K#M#########GT#O#O#O##O#O#O#O#O#O#O#O########G#KA###GGAAAAAAA#CAAA####C#C####CCACCACCCAE#K#K###########K######################EAAAAAAAAAAAAAAAAAAAA##############################E#E#
AAAAAAACAAAAAAAAAAAAAAAAAAAAAACAAAAAABAAACAAAAAAACE#K#K#KKKTKKK#K#KKKKKKKKK##K#O#O#OP#O#O#O#O#O#O#O#OOOOOUUU#K##KOO#########K####KKCR#I#IRRR###########I#C#CCCCICCRRCC#CCCCCCCCCCCCCCCCCCCCCA#####################JAAAAAAAAAAAAAADAAAAFAAAAAAAAA#E#E


s1b = read.table('genes/hg38.gene.tss.exp.A1BG.S3V2_IDEAS_hg38_r3_withHg38Mm10prior.state.AVE.bed.state.bed.seq')
s2b = read.table('genes/mm10.gene.tss.exp.A1bg.S3V2_IDEAS_mm10_r3_withHg38Mm10prior.state.AVE.bed.state.bed.seq')

s1bs = paste(s1b[,1], collapse='')
s2bs = paste(s2b[,1], collapse='')

sw_align1bg = smith_waterman(s1bs,s2bs,similarity=state_dist, gap=-100)

cbind(sw_align1bg$a$alignment$text, sw_align1bg$b$alignment$text)
sw_align1bg$sw

02222222222222222222222220000000000000000000####444####614141##91##9##141414140000000022222222
################################################4044###6141####91##9191414141414


dist_0 = function(x, y){
	if (x==y){
		s = 10
	} else{
		s = -100
	}
	return(s)
}

sw_align1bg = smith_waterman('abc','abbc', similarity = function(x, y) ifelse(x == y, 2L, -10))

sw_align1bg2 = smith_waterman('abccc','abbcc', similarity=dist_0, gap=-100)
cbind(sw_align1bg2$a$alignment$text, sw_align1bg2$b$alignment$text)

sw_align1bg2$sw

sw_align1bg = smith_waterman('abc','abc',similarity=state_dist, gap=0)


match = 2L,
       mismatch = -1L,
       gap = -1L

cbind(sw_align1bg$a$alignment$text, sw_align1bg$b$alignment$text)


###
head -5000 genes/mm10.gene.tss.exp.bed > genes/mm10.gene.tss.exp.bed.tmp

while read chr start end st gn
do
gn_upper=${gn^^}
echo $gn_upper
../../bin/TwoRegions.out 'genes/hg38.gene.tss.exp.'$gn_upper'.S3V2_IDEAS_hg38_r3_withHg38Mm10prior.state.AVE.bed.state.bed.Sseq' 'genes/mm10.gene.tss.exp.'$gn'.S3V2_IDEAS_mm10_r3_withHg38Mm10prior.state.AVE.bed.state.bed.Sseq' para > 'genes/'$gn'.AVE.EpiAlign.S.txt'
done < genes/mm10.gene.tss.exp.bed.tmp

while read chr start end st gn
do
gn_upper=${gn^^}
echo $gn_upper
../../bin/TwoRegions.out 'genes/hg38.gene.tss.exp.'$gn_upper'.S3V2_IDEAS_hg38_r3_withHg38Mm10prior.state.ERY_S002R5.bed.state.bed.Sseq' 'genes/mm10.gene.tss.exp.'$gn'.S3V2_IDEAS_mm10_r3_withHg38Mm10prior.state.ERY_ad_r1.bed.state.bed.Sseq' para > 'genes/'$gn'.ERY_S002R5.ERY_ad_r1.EpiAlign.S.txt'
done < genes/mm10.gene.tss.exp.bed.tmp


### all align score
rm -f AVE.EpiAlign.score.txt
while read chr start end st gn
do
gn_upper=${gn^^}
echo $gn_upper
tail -n+2 'genes/'$gn'.AVE.EpiAlign.S.txt' | head -1 | awk -F '=' '{print $2}' >> AVE.EpiAlign.score.txt
done < genes/mm10.gene.tss.exp.bed.tmp

rm -f ERY_S002R5.ERY_ad_r1.EpiAlign.score.txt
while read chr start end st gn
do
gn_upper=${gn^^}
echo $gn_upper
tail -n+2 'genes/'$gn'.ERY_S002R5.ERY_ad_r1.EpiAlign.S.txt' | head -1 | awk -F '=' '{print $2}' >> ERY_S002R5.ERY_ad_r1.EpiAlign.score.txt
done < genes/mm10.gene.tss.exp.bed.tmp


R

d1 = scan('AVE.EpiAlign.score.txt')
d2 = scan('ERY_S002R5.ERY_ad_r1.EpiAlign.score.txt')
genes = read.table('genes/mm10.gene.tss.exp.bed.tmp', header=F)[,5]

plot_lim = c(min(c(d1,d2)), max(c(d1,d2)))
plot_lim_MA = c(-max(abs(log2(d2/d1))), max(abs(log2(d2/d1))))

A = d1/2+d2/2
M = log2((d2+10)/(d1+10))
png('EpiAlign.score.MA.png')
plot(A, M, ylim=plot_lim_MA)
quantil_thresh = 0.99
points(A[abs(M)>quantile(abs(M), quantil_thresh)], M[abs(M)>quantile(abs(M), quantil_thresh)], col='red', pch=16)
text(A[abs(M)>quantile(abs(M), quantil_thresh)], M[abs(M)>quantile(abs(M), quantil_thresh)]+0.2, genes[abs(M)>quantile(abs(M), quantil_thresh)], col='blue')
abline(h=0)
dev.off()

png('EpiAlign.score.png')
plot(d1,d2, xlim=plot_lim, ylim=plot_lim)
points(d1[abs(M)>quantile(abs(M), quantil_thresh)], d2[abs(M)>quantile(abs(M), quantil_thresh)], col='red', pch=16)
text(d1[abs(M)>quantile(abs(M), quantil_thresh)], d2[abs(M)>quantile(abs(M), quantil_thresh)], genes[abs(M)>quantile(abs(M), quantil_thresh)], col='blue')
abline(0,1)
dev.off()


d12 = cbind(d1,d2)
rownames(d12) = genes
colnames(d12) = c('AVE','ERY')

head(d12[order(-abs(d12[,1]-d12[,2])),])


cat S3V2_IDEAS_hg38_r3_withHg38Mm10prior.state | awk -F ' ' -v OFS='\t' '{print $2,$3,$4, $5, $6,$7, $12, $16,$17, $18, $20, $32,$33, $34, $42, $44, $46  }' > S3V2_IDEAS_hg38_r3_withHg38Mm10prior.state.matched_ct.bed
cat S3V2_IDEAS_mm10_r3_withHg38Mm10prior.state | awk -F ' ' -v OFS='\t' '{print $2,$3,$4, $5, $6,$7, $11, $14,$15, $20, $22, $26,$27, $28, $30, $31, $32  }' > S3V2_IDEAS_mm10_r3_withHg38Mm10prior.state.matched_ct.bed


cat S3V2_IDEAS_hg38_r3_withHg38Mm10prior.state | awk -F ' ' -v OFS='\t' '{print $2,$3,$4,$16}' > S3V2_IDEAS_hg38_r3_withHg38Mm10prior.state.ERY_S002R5.bed
cat S3V2_IDEAS_mm10_r3_withHg38Mm10prior.state | awk -F ' ' -v OFS='\t' '{print $2,$3,$4,$14}' > S3V2_IDEAS_mm10_r3_withHg38Mm10prior.state.ERY_ad_r1.bed

bedtools intersect -a S3V2_IDEAS_hg38_r3_withHg38Mm10prior.state.ERY_S002R5.bed -b hg38.GATA1.tss.exp.bed -wa -u > S3V2_IDEAS_hg38_r3_withHg38Mm10prior.state.ERY_S002R5.GATA1.bed
bedtools intersect -a S3V2_IDEAS_mm10_r3_withHg38Mm10prior.state.ERY_ad_r1.bed -b mm10.GATA1.tss.exp.bed -wa -u > S3V2_IDEAS_mm10_r3_withHg38Mm10prior.state.ERY_ad_r1.GATA1.r.bed
Rscript reverse_seq.R S3V2_IDEAS_mm10_r3_withHg38Mm10prior.state.ERY_ad_r1.GATA1.r.bed S3V2_IDEAS_mm10_r3_withHg38Mm10prior.state.ERY_ad_r1.GATA1.bed

###
cut -f4 S3V2_IDEAS_hg38_r3_withHg38Mm10prior.state.ERY_S002R5.GATA1.bed > S3V2_IDEAS_hg38_r3_withHg38Mm10prior.state.ERY_S002R5.GATA1.bed.seq
cut -f4 S3V2_IDEAS_mm10_r3_withHg38Mm10prior.state.ERY_ad_r1.GATA1.bed > S3V2_IDEAS_mm10_r3_withHg38Mm10prior.state.ERY_ad_r1.GATA1.bed.seq

Rscript compress_seq.R S3V2_IDEAS_hg38_r3_withHg38Mm10prior.state.AVE.GATA1.bed.seq S3V2_IDEAS_hg38_r3_withHg38Mm10prior.state.AVE.GATA1.bed.Sseq
Rscript compress_seq.R S3V2_IDEAS_mm10_r3_withHg38Mm10prior.state.AVE.Gata1.bed.r.seq S3V2_IDEAS_mm10_r3_withHg38Mm10prior.state.AVE.Gata1.bed.r.Sseq

../../bin/TwoRegions.out S3V2_IDEAS_hg38_r3_withHg38Mm10prior.state.AVE.GATA1.bed.Sseq S3V2_IDEAS_mm10_r3_withHg38Mm10prior.state.AVE.Gata1.bed.r.Sseq para > GATA1.EpiAlign.S.txt

Rscript compress_seq.R S3V2_IDEAS_hg38_r3_withHg38Mm10prior.state.ERY_S002R5.GATA1.bed.seq S3V2_IDEAS_hg38_r3_withHg38Mm10prior.state.ERY_S002R5.GATA1.bed.Sseq
Rscript compress_seq.R S3V2_IDEAS_mm10_r3_withHg38Mm10prior.state.ERY_ad_r1.GATA1.bed.seq S3V2_IDEAS_mm10_r3_withHg38Mm10prior.state.ERY_ad_r1.GATA1.bed.Sseq

../../bin/TwoRegions.out S3V2_IDEAS_hg38_r3_withHg38Mm10prior.state.ERY_S002R5.GATA1.bed.Sseq S3V2_IDEAS_mm10_r3_withHg38Mm10prior.state.ERY_ad_r1.GATA1.bed.Sseq para > GATA1.ERY.EpiAlign.S.txt

../../bin/TwoRegions.out S3V2_IDEAS_hg38_r3_withHg38Mm10prior.state.ERY_S002R5.GATA1.bed.Sseq S3V2_IDEAS_hg38_r3_withHg38Mm10prior.state.ERY_S002R5.GATA1.bed.Sseq para > GATA1.hg38.EpiAlign.S.txt

../../bin/TwoRegions.out S3V2_IDEAS_mm10_r3_withHg38Mm10prior.state.ERY_ad_r1.GATA1.bed.Sseq S3V2_IDEAS_mm10_r3_withHg38Mm10prior.state.AVE.Gata1.bed.r.Sseq para > GATA1.mm10.EpiAlign.S.txt
```
