## Supplementary material for "Interspecies regulatory landscapes and elements revealed by novel joint systematic integration of human and mouse blood cell epigenomes": Scripts for IDEAS modeling of epigenetic states jointly in human and mouse: get_cor_heatmap.human_vs_mouse.addmultipleIDs.finalused_forJoint_MetaCluster.07202023.onlyATAC.finalused_15clusters.200kbwin.R

```
#conda activate som
library(dynamicTreeCut)
library(pheatmap)
library(lsa)
library(LSD)

noise_mat_fun = function(input_mat, noise_level){
  noise_mat_i = matrix(runif(dim(input_mat)[1]*dim(input_mat)[2], -noise_level, noise_level), dim(input_mat)[1], dim(input_mat)[2])
  return(noise_mat_i)
}

get_smooth_bedID = function(bed_mat_OD, smooth_win, bed_cols, sig_cols){
  #bed_mat_OD = dh
  #smooth_win = 250
  #bed_cols = 1:3
  #sig_cols = 7:dim(dh)[2]
  ###
  bed_mat = bed_mat_OD[,bed_cols]
  sig_mat = bed_mat_OD[,sig_cols]
  ### expand bed file
  bed_mat[,2] = bed_mat[,2]-smooth_win
  bed_mat[bed_mat[,2]<0,2] = 0
  bed_mat[,3] = bed_mat[,3]+smooth_win
  ### get merge bed ID
  write.table(bed_mat, 'tmp.file.bed', quote=F, col.names=F, row.names=F, sep='\t')
  bash1 = 'bedtools merge -i tmp.file.bed > tmp.file.merge.bed'
  system(bash1)
  bash1 = 'cat tmp.file.merge.bed | awk -F \'\t\' -v OFS=\'\t\' \'{print $0, NR}\' > tmp.file.merge.ID.bed'
  system(bash1)
  bash1 = 'bedtools map -a tmp.file.bed -b tmp.file.merge.ID.bed -c 4 -o max > tmp.file.WithmergeID.bed'
  system(bash1)
  ###
  bed_mat_withMergeID = read.table('tmp.file.WithmergeID.bed', header=F, sep='\t')
  merge_IDs = bed_mat_withMergeID[,4]
  ###
  k = 0
  for (id in merge_IDs){
    k = k +1
    if (k%%10000==0){print(k/length(merge_IDs))}
    if (sum(merge_IDs==id)>1){
      colMean_sig = colMeans(sig_mat[merge_IDs==id,])
      sig_mat[merge_IDs==id,] = t(matrix(replicate(sum(merge_IDs==id),colMean_sig),nrow=length(colMean_sig)))
    }
  }
  ### clean tmp files
  bash1 = 'rm tmp.file.bed tmp.file.merge.bed tmp.file.merge.ID.bed tmp.file.WithmergeID.bed'
  system(bash1)
  bed_mat_new = bed_mat_OD
  bed_mat_new[,sig_cols] = sig_mat
  return(bed_mat_new)
}

#################################################
setwd('/Users/guanjuexiang/Documents/projects/analysis/04_05_2022_coe/coe_analysis')

dh_esRP = read.table('S3V2_IDEAS_hg38_ccre2.cCRE.M.notall0.rmallNEU.withid.coe_mat.PDmerged.clusterID.txt', header=T, sep='\t')
dm_esRP = read.table('S3V2_IDEAS_mm10_ccre2.cCRE.M.notall0.withid.coe_mat.PDmerged.clusterID.txt', header=T, sep='\t')

dh0 = read.table('S3V2_IDEAS_hg38_ccre2.cCRE.M.notall0.withid.ATACsig.withid.txt', header=T, sep='\t')
dm0 = read.table('../coe_analysis_mouse/S3V2_IDEAS_mm10_ccre2.cCRE.M.notall0.withid.ATACsig.withid.txt', header=T, sep='\t')

dh0$NEU_C001UYH1 = NULL
dh0$NEU_C0011IH2 = NULL

dh = cbind(dh_esRP[,1:6], dh0[,-c(1:4)])
dm = cbind(dm_esRP[,1:6], dm0[,-c(1:4)])


dh = get_smooth_bedID(dh, 250, 1:3, 7:dim(dh)[2])
dm = get_smooth_bedID(dm, 250, 1:3, 7:dim(dm)[2])

dhs = dh[,-c(1:6)]
rep1_h = c(1, 2, 4, 6, 8, 10, 12, 14, 16, 18, 20, 22, 24, 26, 28, 30, 32, 34, 36, 38, 40)
rep2_h = c(1, 3, 5, 7, 9, 11, 13, 15, 17, 19, 21, 23, 25, 27, 29, 31, 33, 35, 37, 39, 41)
ct_h = c('AVE', 'B', 'CD34', 'CLP', 'CMP', 'EOS', 'ERY', 'GMP', 'HSC', 'HUDEP1', 'HUDEP2', 'K562', 'LMPP', 'MEP', 'MK', 'MON', 'MONp', 'MPP', 'NK', 'TCD4', 'TCD8')
dhs_ctmerge = matrix(0, nrow=dim(dhs)[1], ncol=length(ct_h))
colnames(dhs_ctmerge) = ct_h
for (j in 1:length(ct_h)){
  dhs_ctmerge[,j] = (dhs[,rep1_h[j]] + dhs[,rep2_h[j]])/2
}

dms = dm[,-c(1:6)]
rep1_m = c(1, 2, 4, 5, 6, 7, 8, 10, 12, 14, 16, 17, 18, 19, 21, 22, 24, 25, 26, 27, 28, 29, 31)
rep2_m = c(1, 3, 4, 5, 6, 7, 9, 11, 13, 15, 16, 17, 18, 20, 21, 23, 24, 25, 26, 27, 28, 30, 32)
ct_m = c('AVE', 'B', 'CFUE', 'CFUMK', 'CLP', 'CMP', 'ER4', 'ERY', 'ERYfl', 'G1E', 'GMP', 'HPC7', 'HSC', 'MEL', 'MEP', 'MK', 'MON', 'NEU', 'NK', 'TCD4', 'TCD8', 'iMEL', 'iMK')
dms_ctmerge = matrix(0, nrow=dim(dms)[1], ncol=length(ct_m))
colnames(dms_ctmerge) = ct_m
for (j in 1:length(ct_m)){
  dms_ctmerge[,j] = (dms[,rep1_m[j]] + dms[,rep2_m[j]])/2
}

### shared cts
ct_shared = ct_h[is.element(ct_h, ct_m)]
### human
dhs_ctmerge_shared = dhs_ctmerge[,is.element(colnames(dhs_ctmerge), ct_shared)]
dhs_ctmerge_shared = dhs_ctmerge_shared[,order(colnames(dhs_ctmerge_shared))]
### mouse
dms_ctmerge_shared = dms_ctmerge[,is.element(colnames(dms_ctmerge), ct_shared)]
dms_ctmerge_shared = dms_ctmerge_shared[,order(colnames(dms_ctmerge_shared))]

### ct reorder
ct_shared_reorder = c('AVE', 'HSC', 'CMP', 'CLP', 'MEP', 'ERY', 'MK', 'GMP', 'MON', 'NK', 'B', 'TCD4', 'TCD8')
dhs_ctmerge_shared_reorder = c()
dms_ctmerge_shared_reorder = c()
for (i in 1:length(ct_shared_reorder)){
  dhs_ctmerge_shared_reorder = cbind(dhs_ctmerge_shared_reorder, dhs_ctmerge_shared[,colnames(dhs_ctmerge_shared)==ct_shared_reorder[i]])
  dms_ctmerge_shared_reorder = cbind(dms_ctmerge_shared_reorder, dms_ctmerge_shared[,colnames(dms_ctmerge_shared)==ct_shared_reorder[i]])
}
colnames(dhs_ctmerge_shared_reorder) = ct_shared_reorder
colnames(dms_ctmerge_shared_reorder) = ct_shared_reorder

### recalcualte average
dhs_ctmerge_shared_reorder[,1] = rowMeans(dhs_ctmerge_shared_reorder[,-1])
dms_ctmerge_shared_reorder[,1] = rowMeans(dms_ctmerge_shared_reorder[,-1])
#################################################


#################################################
### merge Human and Mouse matrix
# !!! Using mm10 esRP matrix without scaling or lm batcheffect removal gives better downstream clustering results !!! #
dhs_dms_ctmerge_shared_reorder = rbind(dhs_ctmerge_shared_reorder, dms_ctmerge_shared_reorder)
cCRE_id = c(paste0(rep('H', dim(dhs_ctmerge_shared_reorder)[1])), paste0(rep('M', dim(dms_ctmerge_shared_reorder)[1])))
coordinates = rbind(dh[,c(1:6)], dm[,c(1:6)])
### Human & mouse cCRE PCA
dhs_dms_ctmerge_shared_reorder_pca = prcomp(dhs_dms_ctmerge_shared_reorder, center = F, scale. = F)
dhs_ctmerge_shared_reorder_pca_summary = summary(dhs_dms_ctmerge_shared_reorder_pca)
print(dhs_ctmerge_shared_reorder_pca_summary$importance)
pca_used_num = sum(dhs_ctmerge_shared_reorder_pca_summary$importance[3,]<0.99)+1
set.seed(2019)
used_id = sample(dim(dhs_dms_ctmerge_shared_reorder_pca$x)[1], 10000)
dhs_ctmerge_shared_reorder_pca_x_plot = dhs_dms_ctmerge_shared_reorder_pca$x[used_id,]
cCRE_id_plot = cCRE_id[used_id]
png('ATAC_only/PCA.plot.raw.png')
plot(dhs_ctmerge_shared_reorder_pca_x_plot[,1], dhs_ctmerge_shared_reorder_pca_x_plot[,2])
points(dhs_ctmerge_shared_reorder_pca_x_plot[cCRE_id_plot=='H',1], dhs_ctmerge_shared_reorder_pca_x_plot[cCRE_id_plot=='H',2], col='red')
points(dhs_ctmerge_shared_reorder_pca_x_plot[cCRE_id_plot=='M',1], dhs_ctmerge_shared_reorder_pca_x_plot[cCRE_id_plot=='M',2], col='blue')
points(mean(dhs_ctmerge_shared_reorder_pca_x_plot[cCRE_id_plot=='H',1]), mean(dhs_ctmerge_shared_reorder_pca_x_plot[cCRE_id_plot=='H',2]))
points(mean(dhs_ctmerge_shared_reorder_pca_x_plot[cCRE_id_plot=='M',1]), mean(dhs_ctmerge_shared_reorder_pca_x_plot[cCRE_id_plot=='M',2]))
dev.off()

for (ii in 1:dim(dhs_ctmerge_shared_reorder_pca_x_plot)[2]){
png(paste0('ATAC_only/PCA.plot.raw.1.',ii,'.png'))
pc_i = 1
pc_j = ii
plot(dhs_ctmerge_shared_reorder_pca_x_plot[,pc_i], dhs_ctmerge_shared_reorder_pca_x_plot[,pc_j])
points(dhs_ctmerge_shared_reorder_pca_x_plot[cCRE_id_plot=='H',pc_i], dhs_ctmerge_shared_reorder_pca_x_plot[cCRE_id_plot=='H',pc_j], col='red')
points(dhs_ctmerge_shared_reorder_pca_x_plot[cCRE_id_plot=='M',pc_i], dhs_ctmerge_shared_reorder_pca_x_plot[cCRE_id_plot=='M',pc_j], col='blue')
points(mean(dhs_ctmerge_shared_reorder_pca_x_plot[cCRE_id_plot=='H',pc_i]), mean(dhs_ctmerge_shared_reorder_pca_x_plot[cCRE_id_plot=='H',pc_j]))
points(mean(dhs_ctmerge_shared_reorder_pca_x_plot[cCRE_id_plot=='M',pc_i]), mean(dhs_ctmerge_shared_reorder_pca_x_plot[cCRE_id_plot=='M',pc_j]))
dev.off()
}
#################################################


#################################################
### cluster cCRE by PCA-KM by subsampling and identify reproducible clusters pca 1:5 cover 95% of variance
### 1 example
set.seed(2019)
KMPCA_meansig_mat_1 = c()
start_km_num = 100
km_dhs_ctmerge_shared1 = kmeans(dhs_dms_ctmerge_shared_reorder_pca$x[,1:pca_used_num], centers=start_km_num)
### check KM cluster cCRE Human/Mouse ratio
for (i in 1:start_km_num){
  dhs_dms_ctmerge_shared_reorder_ji = dhs_dms_ctmerge_shared_reorder[km_dhs_ctmerge_shared1$cluster==i,]
  KMPCA_meansig_mat_1 = rbind(KMPCA_meansig_mat_1, colMeans(dhs_dms_ctmerge_shared_reorder_ji))
}
colnames(KMPCA_meansig_mat_1) = colnames(dhs_ctmerge_shared_reorder)
pdf('ATAC_only/KMPCA.Joint.1.cluster.pdf', height=30)
pheatmap(KMPCA_meansig_mat_1, cluster_col=T, cluster_rows=T)
dev.off()
#################################################


#################################################
### Select reproducible State
#################################################
### check initial KM-K
set.seed(2019)
cCRE_esRP_KM_determineK_KM_ratio = c()
for (ini_k in seq(10,200,by=10)){
  print(ini_k)
  used_id_j = sample(dim(dhs_dms_ctmerge_shared_reorder_pca$x)[1], 50000)
  km_dhs_ctmerge_shared_check = kmeans(dhs_dms_ctmerge_shared_reorder_pca$x[used_id_j,1:pca_used_num], centers=ini_k)
  cCRE_esRP_KM_determineK_KM_ratio = c(cCRE_esRP_KM_determineK_KM_ratio, km_dhs_ctmerge_shared_check$tot.withinss/km_dhs_ctmerge_shared_check$betweenss)
}
pdf('ATAC_only/cCRE_esRP_KM.determineK.pdf')
plot(seq(10,200,by=10), cCRE_esRP_KM_determineK_KM_ratio)
lines(seq(10,200,by=10), cCRE_esRP_KM_determineK_KM_ratio)
abline(v=100)
dev.off()

### Initial Kmeans
set.seed(2019)
KMPCA_meansig_mat = c()
KMPCA_iteration = c()
KMPCA_cCRE_num = c()
HM_count = c()
check_reproducible = 100
start_km_num = 100
for (j in 1:check_reproducible){
print(j)
used_id_j = sample(dim(dhs_dms_ctmerge_shared_reorder_pca$x)[1], 50000)
km_dhs_ctmerge_shared1 = kmeans(dhs_dms_ctmerge_shared_reorder_pca$x[used_id_j,1:pca_used_num], centers=start_km_num)
### check KM cluster cCRE Human/Mouse ratio
for (i in 1:start_km_num){
  if (sum(km_dhs_ctmerge_shared1$cluster==i)>100){
  dhs_dms_ctmerge_shared_reorder_ji = dhs_dms_ctmerge_shared_reorder[used_id_j,][km_dhs_ctmerge_shared1$cluster==i,]
  KMPCA_meansig_mat = rbind(KMPCA_meansig_mat, colMeans(dhs_dms_ctmerge_shared_reorder_ji))
  KMPCA_cCRE_num = c(KMPCA_cCRE_num, dim(dhs_dms_ctmerge_shared_reorder_ji)[1])
  cCRE_id_ji = cCRE_id[used_id_j][km_dhs_ctmerge_shared1$cluster==i]
  HM_count = rbind(HM_count, c(sum(cCRE_id_ji=='H'), sum(cCRE_id_ji=='M')))
  }
}
KMPCA_iteration = c(KMPCA_iteration, rep(j, start_km_num))
}
#################################################


#################################################
### identify reproducible state
set.seed(2019)
KMPCA_meansig_mat_pca_x = KMPCA_meansig_mat %*% dhs_dms_ctmerge_shared_reorder_pca$rotation
KMPCA_meansig_mat_pca_joint_KM = kmeans(KMPCA_meansig_mat_pca_x[,1:pca_used_num], centers=start_km_num)

KMPCA_iteration_KM_k = c()
KMPCA_meansig_mat_meansig_mat = c()
HM_count_mean = c()
cCRE_count_KMPCA_iteration_KM_k = c()
cluster_set = min(unique(KMPCA_meansig_mat_pca_joint_KM$cluster)):max(unique(KMPCA_meansig_mat_pca_joint_KM$cluster))
for (KM_k in cluster_set){
  used_KM_k = KMPCA_meansig_mat_pca_joint_KM$cluster==KM_k
  KMPCA_iteration_KM_k = c(KMPCA_iteration_KM_k, length(unique(KMPCA_iteration[used_KM_k])) )
  if (sum(used_KM_k)>1){
    KMPCA_meansig_mat_meansig_mat = rbind(KMPCA_meansig_mat_meansig_mat, colMeans(KMPCA_meansig_mat[used_KM_k,]))
    HM_count_mean = rbind(HM_count_mean, apply(HM_count[used_KM_k,], 2, sum))
    cCRE_count_KMPCA_iteration_KM_k = c(cCRE_count_KMPCA_iteration_KM_k, sum(KMPCA_cCRE_num[used_KM_k]))
  } else{
    KMPCA_meansig_mat_meansig_mat = rbind(KMPCA_meansig_mat_meansig_mat, KMPCA_meansig_mat[used_KM_k,])
    HM_count_mean = rbind(HM_count_mean, HM_count[used_KM_k,])
    cCRE_count_KMPCA_iteration_KM_k = c(cCRE_count_KMPCA_iteration_KM_k, KMPCA_cCRE_num[used_KM_k])
  }
}


HM_count_mean_log2FC = log2((HM_count_mean[,1]+1)/(HM_count_mean[,2]/mean(HM_count_mean[,2])*mean(HM_count_mean[,1])+1))
#select_Kclusters = ((KMPCA_iteration_KM_k>=(check_reproducible*0.9)) * (abs(HM_count_mean_log2FC)<=1000))!=0


### determine check_reproducible threshold
zp_threshold_rmtop = 0.05
zp_threshold = 0.05
iter_n_threshold = 5
check_reproducible_vec = KMPCA_iteration_KM_k/check_reproducible
for (i in 1:iter_n_threshold){
check_reproducible_vec_pre = check_reproducible_vec
zp = pnorm((check_reproducible_vec-mean(check_reproducible_vec))/sd(check_reproducible_vec), lower.tail=T)
if (sum(zp<zp_threshold_rmtop)>0){
print(min(abs(check_reproducible_vec)[zp<zp_threshold_rmtop]))
}
check_reproducible_vec = check_reproducible_vec[zp>=zp_threshold_rmtop]
if (length(check_reproducible_vec)==length(check_reproducible_vec_pre)){
  break
}
}
check_reproducible_vec_new = KMPCA_iteration_KM_k/check_reproducible
zp = pnorm((check_reproducible_vec_new-mean(check_reproducible_vec))/sd(check_reproducible_vec), lower.tail=T)
reproducible_thresh = min(check_reproducible_vec_new[zp>=zp_threshold])
print(reproducible_thresh)

### determine abs_HM_count_mean_log2FC threshold
abs_HM_count_mean_log2FC = abs(HM_count_mean_log2FC)
for (i in 1:iter_n_threshold){
abs_HM_count_mean_log2FC_pre = abs_HM_count_mean_log2FC
zp = pnorm((abs_HM_count_mean_log2FC-mean(abs_HM_count_mean_log2FC))/sd(abs_HM_count_mean_log2FC), lower.tail=F)
if (sum(zp<zp_threshold_rmtop)>0){
print(min(abs(abs_HM_count_mean_log2FC)[zp<zp_threshold_rmtop]))
}
abs_HM_count_mean_log2FC = abs_HM_count_mean_log2FC[zp>=zp_threshold_rmtop]
if (length(abs_HM_count_mean_log2FC)==length(abs_HM_count_mean_log2FC_pre)){
  break
}
}
abs_HM_count_mean_log2FC_new = abs(HM_count_mean_log2FC)
zp = pnorm((abs_HM_count_mean_log2FC_new-mean(abs_HM_count_mean_log2FC))/sd(abs_HM_count_mean_log2FC), lower.tail=F)
HM_count_mean_log2FC_thresh = max(abs_HM_count_mean_log2FC_new[zp>=zp_threshold])
print(HM_count_mean_log2FC_thresh)
2.569671
#HM_count_mean_log2FC_thresh = 2


pdf('ATAC_only/KM.reproducible.p.hist.pdf')
hist(KMPCA_iteration_KM_k/check_reproducible, breaks=30)
#reproducible_thresh = 0.75
abline(v=reproducible_thresh)
box()
dev.off()

pdf('ATAC_only/abs_HM_count_mean_log2FC.hist.pdf')
hist(abs(HM_count_mean_log2FC), breaks=30)
#HM_count_mean_log2FC_thresh = 1.5
abline(v=HM_count_mean_log2FC_thresh)
box()
dev.off()


png('ATAC_only/abs_HM_count_mean_log2FC.reproducible.png')
plot(abs(HM_count_mean_log2FC), KMPCA_iteration_KM_k/check_reproducible)
abline(h=reproducible_thresh)
abline(v=HM_count_mean_log2FC_thresh)
dev.off()


### plot 2 heatmap (1) reproducible and cross species (2) reproducible
select_Kclusters = ((KMPCA_iteration_KM_k>=(check_reproducible*reproducible_thresh)) * (abs(HM_count_mean_log2FC)<=HM_count_mean_log2FC_thresh))!=0
KMPCA_meansig_mat_meansig_mat_reproducible = KMPCA_meansig_mat_meansig_mat[select_Kclusters,]
cCRE_count_KMPCA_iteration_KM_k_reproducible = cCRE_count_KMPCA_iteration_KM_k[select_Kclusters]
colnames(KMPCA_meansig_mat_meansig_mat_reproducible) = colnames(dhs_ctmerge_shared_reorder)
dim(KMPCA_meansig_mat_meansig_mat_reproducible)

pdf('ATAC_only/KMPCA.Joint.reproducible.cross_spec.cluster.pdf', height=10, width=6)
pheatmap(KMPCA_meansig_mat_meansig_mat_reproducible, cluster_col=T, cluster_rows=T, clustering_distance_rows=dist(1-cor(t(KMPCA_meansig_mat_meansig_mat_reproducible))))
dev.off()

dhs_dms_ctmerge_shared_reorder_meansig_ctgroup = c()
#ct_groups = cutreeDynamic(hclust(dist(t(KMPCA_meansig_mat_meansig_mat_reproducible))), minClusterSize=1, deepSplit=4, method='hybrid')
ct_groups = cutree(hclust(dist(t(KMPCA_meansig_mat_meansig_mat_reproducible))), 4)
cbind(ct_groups, colnames(KMPCA_meansig_mat_meansig_mat_reproducible))[order(ct_groups),]

png('ATAC_only/test.tree.png')
plot(hclust(dist(t(KMPCA_meansig_mat_meansig_mat_reproducible))))
dev.off()

###
select_Kclusters1 = ((KMPCA_iteration_KM_k>=(check_reproducible*reproducible_thresh)) )!=0
KMPCA_meansig_mat_meansig_mat_reproducible1 = KMPCA_meansig_mat_meansig_mat[select_Kclusters1,]
cCRE_count_KMPCA_iteration_KM_k_reproducible1 = cCRE_count_KMPCA_iteration_KM_k[select_Kclusters1]
colnames(KMPCA_meansig_mat_meansig_mat_reproducible1) = colnames(dhs_ctmerge_shared_reorder)
dim(KMPCA_meansig_mat_meansig_mat_reproducible1)

pdf('ATAC_only/KMPCA.Joint.reproducible.cluster.pdf', height=12, width=6)
plot_color_lim = 15
breaksList = seq(-plot_color_lim, plot_color_lim, by = 0.001)
my_colorbar=colorRampPalette(c('blue', 'white', 'red'))(n = length(breaksList))
pheatmap(KMPCA_meansig_mat_meansig_mat_reproducible1, cluster_col=T, cluster_rows=T, cex = 1.5, color=my_colorbar, breaks = breaksList, clustering_distance_rows=dist(1-cor(t(KMPCA_meansig_mat_meansig_mat_reproducible1))), clustering_distance_cols=dist(t(KMPCA_meansig_mat_meansig_mat_reproducible)))
dev.off()
#################################################


#################################################
### reproducible cluster ids & prepare Train data for QDA
set.seed(2019)
reproducible_clusters = cluster_set[select_Kclusters]
reproducible_clusters_rows = is.element(KMPCA_meansig_mat_pca_joint_KM$cluster, reproducible_clusters)
train_data_notPCs = KMPCA_meansig_mat[reproducible_clusters_rows,]
train_data_y = KMPCA_meansig_mat_pca_joint_KM$cluster[reproducible_clusters_rows]
QDA_train_data = cbind(train_data_y, train_data_notPCs)
colnames(QDA_train_data)[1] = 'Y'
QDA_cluster_train_data_prior = cCRE_count_KMPCA_iteration_KM_k_reproducible / sum(cCRE_count_KMPCA_iteration_KM_k_reproducible)

### predict QDA id for each cCRE
#all_cCREs_prediction_Y = predict(QDA_model, newdata = as.data.frame(dhs_dms_ctmerge_shared_reorder_pca$x))$class
#################################################
### correlation based clustering
set.seed(2019)
cluster_center_mean = matrix(0, nrow = length(unique(train_data_y)), ncol=dim(train_data_notPCs)[2])
for (i in 1:length(unique(train_data_y))){
  cluster_center_mean[i,] = colMeans(train_data_notPCs[train_data_y==unique(train_data_y)[i],])
}
rownames(cluster_center_mean) = unique(train_data_y)

pdf('ATAC_only/KMnotPCA.Joint.reproducible.cluster.pdf', height=12, width=6)
plot_color_lim = 15
breaksList = seq(-plot_color_lim, plot_color_lim, by = 0.001)
my_colorbar=colorRampPalette(c('blue', 'white', 'red'))(n = length(breaksList))
pheatmap(cluster_center_mean, cluster_col=T, cluster_rows=T, cex = 1.5, clustering_distance_cols=dist(t(KMPCA_meansig_mat_meansig_mat_reproducible)))
dev.off()

noise_level = 0.001
each_cCRE_x = dhs_dms_ctmerge_shared_reorder + noise_mat_fun(dhs_dms_ctmerge_shared_reorder, noise_level)
#each_cCRE_x_cor_mat1 = cor(t(each_cCRE_x), t(cluster_center_mean))
each_cCRE_x_cor_mat = t(apply(each_cCRE_x, 1, function(x) cosine(x, t(cluster_center_mean)) ))
each_cCRE_x_cor_mat_Y_mat = apply(each_cCRE_x_cor_mat, 1, function(x) names(which.max(x)))
print(each_cCRE_x_cor_mat[200342+(94096:94099),])
print(apply(each_cCRE_x_cor_mat[200342+(94092:94102),], 1, which.max))

iter_cCRE_assign_num = 30
for (cor_i in 2:iter_cCRE_assign_num){
  print(cor_i)
each_cCRE_x = dhs_dms_ctmerge_shared_reorder + noise_mat_fun(dhs_dms_ctmerge_shared_reorder, noise_level)
#each_cCRE_x_cor_mat_i = cor(t(each_cCRE_x), t(QDA_model_mean))
each_cCRE_x_cor_mat_i = t(apply(each_cCRE_x, 1, function(x) cosine(x, t(cluster_center_mean)) ))
each_cCRE_x_cor_mat = each_cCRE_x_cor_mat + each_cCRE_x_cor_mat_i
each_cCRE_x_cor_mat_Y_mat = cbind(each_cCRE_x_cor_mat_Y_mat, apply(each_cCRE_x_cor_mat, 1, function(x) names(which.max(x))))
print(each_cCRE_x_cor_mat[200342+(94096:94099),])
}
each_cCRE_x_cor_mat = each_cCRE_x_cor_mat / iter_cCRE_assign_num
### get max.cor as cluster
each_cCRE_x_cor_mat_Y = apply(each_cCRE_x_cor_mat, 1, function(x) names(which.max(x)))
all_cCREs_prediction_Y = each_cCRE_x_cor_mat_Y
###
#################################################


#################################################
### check cCRE counts
cCRE_id_HM_count = c()
dhs_dms_ctmerge_shared_reorder_meansig_H = c()
dhs_dms_ctmerge_shared_reorder_meansig_M = c()
dhs_dms_ctmerge_shared_reorder_meansig =c()
dhs_dms_ctmerge_shared_reorder_meansig_rowN = c()
for (KM_i in unique(all_cCREs_prediction_Y)){
  cCRE_id_KM_i = cCRE_id[all_cCREs_prediction_Y==KM_i]
  cCRE_id_HM_count = rbind(cCRE_id_HM_count, c(sum(cCRE_id_KM_i=='H'), sum(cCRE_id_KM_i=='M')))
  dhs_dms_ctmerge_shared_reorder_meansig = rbind(dhs_dms_ctmerge_shared_reorder_meansig, colMeans(dhs_dms_ctmerge_shared_reorder[all_cCREs_prediction_Y==KM_i,]))
  dhs_dms_ctmerge_shared_reorder_meansig_rowN = c(dhs_dms_ctmerge_shared_reorder_meansig_rowN, sum(all_cCREs_prediction_Y==KM_i))
  ###
  H_rows = cCRE_id=='H'
  dhs_dms_ctmerge_shared_reorder_meansig_H = rbind(dhs_dms_ctmerge_shared_reorder_meansig_H, colMeans(dhs_dms_ctmerge_shared_reorder[H_rows,][all_cCREs_prediction_Y[H_rows]==KM_i,]))
  M_rows = cCRE_id=='M'
  dhs_dms_ctmerge_shared_reorder_meansig_M = rbind(dhs_dms_ctmerge_shared_reorder_meansig_M, colMeans(dhs_dms_ctmerge_shared_reorder[M_rows,][all_cCREs_prediction_Y[M_rows]==KM_i,]))
}
rownames(dhs_dms_ctmerge_shared_reorder_meansig) = unique(all_cCREs_prediction_Y)

### add noise
dhs_dms_ctmerge_shared_reorder_meansig_OD = dhs_dms_ctmerge_shared_reorder_meansig
dhs_dms_ctmerge_shared_reorder_meansig = dhs_dms_ctmerge_shared_reorder_meansig_OD
dhs_dms_ctmerge_shared_reorder_meansig = dhs_dms_ctmerge_shared_reorder_meansig + noise_mat_fun(dhs_dms_ctmerge_shared_reorder_meansig, 0.01)
hclust_cCRE = hclust(dist(1 - cosine(t(dhs_dms_ctmerge_shared_reorder_meansig))))
#################################################


#################################################
### meta-cluster
### Get reproducible Jmeta by hclust with cutreeDynamic with noise mat
set.seed(2019)
pdf('ATAC_only/dhs_dms_ctmerge_shared_reorder_meansig.hist.pdf')
hist(log10(c(abs(dhs_dms_ctmerge_shared_reorder_meansig))), breaks=30, log='')
noise_lim = 0.001
abline(v=log10(noise_lim))
box()
dev.off()
### iteratively cluster QDA clusters
set.seed(2019)
iter_n = 100
### merge ct groups
dhs_dms_ctmerge_shared_reorder_meansig_ctgroup = c()
#ct_groups = cutreeDynamic(hclust(dist(t(KMPCA_meansig_mat_meansig_mat_reproducible))), minClusterSize=1, deepSplit=4, method='hybrid')
ct_groups = cutree(hclust(dist(t(KMPCA_meansig_mat_meansig_mat_reproducible))), 4)
cbind(ct_groups, colnames(KMPCA_meansig_mat_meansig_mat_reproducible))[order(ct_groups),]
for (ctgi in unique(ct_groups)){
  if (sum(ct_groups==ctgi)>1){
    dhs_dms_ctmerge_shared_reorder_meansig_ctgroup = cbind(dhs_dms_ctmerge_shared_reorder_meansig_ctgroup, rowMeans(dhs_dms_ctmerge_shared_reorder_meansig[,ct_groups==ctgi]))
  } else{
    dhs_dms_ctmerge_shared_reorder_meansig_ctgroup = cbind(dhs_dms_ctmerge_shared_reorder_meansig_ctgroup, (dhs_dms_ctmerge_shared_reorder_meansig[,ct_groups==ctgi]))
  }
}
dhs_dms_ctmerge_shared_reorder_meansig_ctgroup[dhs_dms_ctmerge_shared_reorder_meansig_ctgroup<0] = 0
dhs_dms_ctmerge_shared_reorder_meansig_for_cluster = dhs_dms_ctmerge_shared_reorder_meansig
dhs_dms_ctmerge_shared_reorder_meansig_for_cluster[dhs_dms_ctmerge_shared_reorder_meansig_for_cluster<0] = 0

### run hclust 100 times 
KM_by_KM_SameClu_count = matrix(0, dim(dhs_dms_ctmerge_shared_reorder_meansig)[1], dim(dhs_dms_ctmerge_shared_reorder_meansig)[1])
for (iter_i in 1:iter_n){
  print(iter_i)
noise_mat = noise_mat_fun(dhs_dms_ctmerge_shared_reorder_meansig_ctgroup, noise_lim)
dhs_dms_ctmerge_shared_reorder_meansig_noise = dhs_dms_ctmerge_shared_reorder_meansig_ctgroup + noise_mat
hclust_cCRE = hclust(dist(1 - cosine(t( dhs_dms_ctmerge_shared_reorder_meansig_noise ))))
#hclust_cCRE_DTC = cutree(hclust_cCRE, k=15)
hclust_cCRE_DTC = cutreeDynamic(hclust_cCRE, minClusterSize=1, deepSplit=4, dist= as.matrix(dist(1 - cosine(t( dhs_dms_ctmerge_shared_reorder_meansig_noise )))), method='hybrid')
#print(table(hclust_cCRE_DTC))
for (Ki in unique(hclust_cCRE_DTC)){
  used_rows = which(hclust_cCRE_DTC==Ki)
  for (i in 1:length(used_rows)){
    for (j in 1:length(used_rows)){
      KM_by_KM_SameClu_count[used_rows[i],used_rows[j]] = KM_by_KM_SameClu_count[used_rows[i],used_rows[j]]+1
    }
  }
}
}
### count the number runs each row-pair are in the same cluster
colnames(KM_by_KM_SameClu_count) = rownames(dhs_dms_ctmerge_shared_reorder_meansig)
rownames(KM_by_KM_SameClu_count) = rownames(dhs_dms_ctmerge_shared_reorder_meansig)
### plot count in same clusters 
pdf('ATAC_only/KM_by_KM_SameClu_count.reproducible.pdf', height=10, width=10)
pheatmap(KM_by_KM_SameClu_count, cex = 1.5, cluster_col=T, cluster_rows=T, clustering_distance_rows=dist(KM_by_KM_SameClu_count), clustering_distance_cols=dist(KM_by_KM_SameClu_count))#, clustering_distance_rows=dist(dhs_dms_ctmerge_shared_reorder_meansig %*% dhs_dms_ctmerge_shared_reorder_pca$rotation[,]) )
dev.off()
### Use cutreeDynamic to cluster KMs
hclust_bySameClu_count = hclust(dist(KM_by_KM_SameClu_count), method='complete')
hclust_cCRE_DTC = cutreeDynamic(hclust_bySameClu_count, minClusterSize=1, deepSplit=4, dist= as.matrix(dist(KM_by_KM_SameClu_count)), method='hybrid')
table(hclust_cCRE_DTC)
#################################################


#################################################
### get Joint meta-cluster ID
hclust_cCRE_DTC_modified = hclust_cCRE_DTC
all_cCREs_prediction_Y_Jmeta = all_cCREs_prediction_Y
class(all_cCREs_prediction_Y_Jmeta) = 'numeric'
KM_i = 0
for (id_KM in unique(all_cCREs_prediction_Y)){
  KM_i = KM_i+1
  print(as.numeric(id_KM))
  all_cCREs_prediction_Y_Jmeta[all_cCREs_prediction_Y==unique(all_cCREs_prediction_Y)[KM_i]] = hclust_cCRE_DTC_modified[KM_i]
}

row_names_combined = paste(hclust_cCRE_DTC_modified, ':', unique(all_cCREs_prediction_Y), sep='')
rownames(dhs_dms_ctmerge_shared_reorder_meansig) = row_names_combined
rownames(dhs_dms_ctmerge_shared_reorder_meansig_H) = row_names_combined
rownames(dhs_dms_ctmerge_shared_reorder_meansig_M) = row_names_combined

pdf('ATAC_only/KMPCA.Joint.cluster.reproducible.pdf', height=12)
plot_color_lim = max(dhs_dms_ctmerge_shared_reorder_meansig)
breaksList = seq(-plot_color_lim, plot_color_lim, by = 0.001)
my_colorbar=colorRampPalette(c('blue', 'white', 'red'))(n = length(breaksList))
pheatmap(dhs_dms_ctmerge_shared_reorder_meansig[order(hclust_cCRE_DTC_modified),], color=my_colorbar, breaks = breaksList, cex = 1.5, cluster_col=T, cluster_rows=F, clustering_distance_cols = dist(t(dhs_dms_ctmerge_shared_reorder_meansig)), clustering_distance_rows=dist(1 - cosine(t(dhs_dms_ctmerge_shared_reorder_meansig))) )#, clustering_distance_rows=dist(dhs_dms_ctmerge_shared_reorder_meansig %*% dhs_dms_ctmerge_shared_reorder_pca$rotation[,]) )
dev.off()

pdf('ATAC_only/KMPCA.Joint.cluster.reproducible_H.pdf', height=10)
pheatmap(dhs_dms_ctmerge_shared_reorder_meansig_H[order(hclust_cCRE_DTC_modified),], cluster_col=T, cluster_rows=F, cutree_rows=20, clustering_distance_cols = dist(t(dhs_dms_ctmerge_shared_reorder_meansig)), clustering_distance_rows=dist(1 - cosine(t(dhs_dms_ctmerge_shared_reorder_meansig))) )#, clustering_distance_cols = dist(1-cosine(dhs_dms_ctmerge_shared_reorder_meansig)/2 - cor(dhs_dms_ctmerge_shared_reorder_meansig)/2), cclustering_distance_rows=dist(1-cosine(t(dhs_dms_ctmerge_shared_reorder_meansig))/2 - cor(t(dhs_dms_ctmerge_shared_reorder_meansig))/2 ) )
dev.off()

pdf('ATAC_only/KMPCA.Joint.cluster.reproducible_M.pdf', height=10)
pheatmap(dhs_dms_ctmerge_shared_reorder_meansig_M[order(hclust_cCRE_DTC_modified),], cluster_col=T, cluster_rows=F, cutree_rows=20, clustering_distance_cols = dist(t(dhs_dms_ctmerge_shared_reorder_meansig)), clustering_distance_rows=dist(1 - cosine(t(dhs_dms_ctmerge_shared_reorder_meansig))) )#, clustering_distance_cols = dist(1-cosine(dhs_dms_ctmerge_shared_reorder_meansig)/2 - cor(dhs_dms_ctmerge_shared_reorder_meansig)/2), cclustering_distance_rows=dist(1-cosine(t(dhs_dms_ctmerge_shared_reorder_meansig))/2 - cor(t(dhs_dms_ctmerge_shared_reorder_meansig))/2 ) )
dev.off()

png('ATAC_only/cCRE_id_HM_count.png')
plot_lim = c(5, 50000)
plot(cCRE_id_HM_count[,1],cCRE_id_HM_count[,2]/mean(cCRE_id_HM_count[,2])*mean(cCRE_id_HM_count[,1]), log='xy', xlim = plot_lim, ylim=plot_lim, xlab='H', ylab='M')
abline(0,1)
dev.off()

### get Jmet meansignal mat
dhs_dms_ctmerge_shared_reorder_meansig_Jmet_meansig = c()
for (i in unique(hclust_cCRE_DTC_modified)){
if (sum(hclust_cCRE_DTC_modified==i)>1){dhs_dms_ctmerge_shared_reorder_meansig_Jmet_meansig = rbind(dhs_dms_ctmerge_shared_reorder_meansig_Jmet_meansig, colMeans(dhs_dms_ctmerge_shared_reorder_meansig[hclust_cCRE_DTC_modified==i,]))}
if (sum(hclust_cCRE_DTC_modified==i)==1){dhs_dms_ctmerge_shared_reorder_meansig_Jmet_meansig = rbind(dhs_dms_ctmerge_shared_reorder_meansig_Jmet_meansig, dhs_dms_ctmerge_shared_reorder_meansig[hclust_cCRE_DTC_modified==i,])}
}
rownames(dhs_dms_ctmerge_shared_reorder_meansig_Jmet_meansig) = unique(hclust_cCRE_DTC_modified)
###
Jmet_order = as.numeric(rownames(dhs_dms_ctmerge_shared_reorder_meansig_Jmet_meansig)[hclust(dist(dhs_dms_ctmerge_shared_reorder_meansig_Jmet_meansig))$order])
pdf('ATAC_only/KMPCA.Joint.cluster.reproducible.Jmet.pdf', height=6)
dhs_dms_ctmerge_shared_reorder_meansig_Jmet_meansig_plot = dhs_dms_ctmerge_shared_reorder_meansig_Jmet_meansig
dhs_dms_ctmerge_shared_reorder_meansig_Jmet_meansig_plot = dhs_dms_ctmerge_shared_reorder_meansig_Jmet_meansig_plot[,c('MON', 'GMP', 'HSC','CMP','ERY','MEP','MK','NK','TCD4','TCD8','CLP','AVE','B')]
plot_color_lim = max(dhs_dms_ctmerge_shared_reorder_meansig_Jmet_meansig_plot)
breaksList = seq(min(dhs_dms_ctmerge_shared_reorder_meansig_Jmet_meansig_plot), plot_color_lim, by = 0.001)
my_colorbar=colorRampPalette(c('white', 'red'))(n = length(breaksList))
pheatmap(dhs_dms_ctmerge_shared_reorder_meansig_Jmet_meansig_plot, color=my_colorbar, breaks = breaksList, cluster_col=F, cluster_rows=T, clustering_distance_rows = dist((dhs_dms_ctmerge_shared_reorder_meansig_Jmet_meansig)), clustering_distance_cols = dist(t(dhs_dms_ctmerge_shared_reorder_meansig)), cex=1.5)#, clustering_distance_rows=dist(dhs_dms_ctmerge_shared_reorder_meansig %*% dhs_dms_ctmerge_shared_reorder_pca$rotation[,]) )
dev.off()

pdf('ATAC_only/KMPCA.Joint.cluster.reproducible.Jmet.OD.pdf', height=6)
dhs_dms_ctmerge_shared_reorder_meansig_Jmet_meansig_plot = dhs_dms_ctmerge_shared_reorder_meansig_Jmet_meansig
dhs_dms_ctmerge_shared_reorder_meansig_Jmet_meansig_plot = dhs_dms_ctmerge_shared_reorder_meansig_Jmet_meansig_plot[,c('MON', 'GMP', 'HSC','CMP','ERY','MEP','MK','NK','TCD4','TCD8','CLP','AVE','B')]
plot_color_lim = max(dhs_dms_ctmerge_shared_reorder_meansig_Jmet_meansig_plot)
breaksList = seq(min(dhs_dms_ctmerge_shared_reorder_meansig_Jmet_meansig_plot), plot_color_lim, by = 0.001)
my_colorbar=colorRampPalette(c('white', 'red'))(n = length(breaksList))
pheatmap(dhs_dms_ctmerge_shared_reorder_meansig_Jmet_meansig_plot, color=my_colorbar, breaks = breaksList, cluster_col=F, cluster_rows=T, clustering_distance_rows = dist((dhs_dms_ctmerge_shared_reorder_meansig_Jmet_meansig)), clustering_distance_cols = dist(t(dhs_dms_ctmerge_shared_reorder_meansig)), cex=1.5)#, clustering_distance_rows=dist(dhs_dms_ctmerge_shared_reorder_meansig %*% dhs_dms_ctmerge_shared_reorder_pca$rotation[,]) )
dev.off()
#################################################

#################################################
### write table
dh_with_JointClusterID_mat = cbind(dh[,c(1:5)], all_cCREs_prediction_Y[H_rows], all_cCREs_prediction_Y_Jmeta[H_rows], dh[,-c(1:6)])
dm_with_JointClusterID_mat = cbind(dm[,c(1:5)], all_cCREs_prediction_Y[M_rows], all_cCREs_prediction_Y_Jmeta[M_rows], dm[,-c(1:6)])
colnames(dh_with_JointClusterID_mat)[6] = 'SRC_ID'
colnames(dm_with_JointClusterID_mat)[6] = 'SRC_ID'
colnames(dh_with_JointClusterID_mat)[7] = 'JmC_ID'
colnames(dm_with_JointClusterID_mat)[7] = 'JmC_ID'
dm_with_JointClusterID_mat[94092:94102,1:8]
dh_with_JointClusterID_mat[196784:196794,1:8]
write.table(dh_with_JointClusterID_mat[,c(1:3,4,6,7)], 'ATAC_only/S3V2_IDEAS_hg38_ccre2.cCRE.M.notall0.rmallNEU.withid.cCRE_ID.SRC_ID.JmC_ID.bed', quote=F, sep='\t', col.names=T, row.names=F)
write.table(dm_with_JointClusterID_mat[,c(1:3,4,6,7)], 'ATAC_only/S3V2_IDEAS_mm10_ccre2.cCRE.M.notall0.withid.cCRE_ID.SRC_ID.JmC_ID.bed', quote=F, sep='\t', col.names=T, row.names=F)


write.table(dh_with_JointClusterID_mat, 'ATAC_only/S3V2_IDEAS_hg38_ccre2.cCRE.M.notall0.rmallNEU.withid.coe_mat.PDmerged.clusterID.JclusterID.txt', quote=F, sep='\t', col.names=T, row.names=F)
write.table(dm_with_JointClusterID_mat, 'ATAC_only/S3V2_IDEAS_mm10_ccre2.cCRE.M.notall0.withid.coe_mat.PDmerged.clusterID.JclusterID.txt', quote=F, sep='\t', col.names=T, row.names=F)

write.table(dh_with_JointClusterID_mat[,c(1:3,7)], 'ATAC_only/S3V2_IDEAS_hg38_ccre2.cCRE.M.notall0.rmallNEU.withid.clusterID.JclusterID.bed', quote=F, sep='\t', col.names=T, row.names=F)
write.table(dm_with_JointClusterID_mat[,c(1:3,7)], 'ATAC_only/S3V2_IDEAS_mm10_ccre2.cCRE.M.notall0.withid.clusterID.JclusterID.bed', quote=F, sep='\t', col.names=T, row.names=F)

write.table(dh_with_JointClusterID_mat[,c(1:3,4,6,7)], 'ATAC_only/S3V2_IDEAS_hg38_ccre2.cCRE.M.notall0.rmallNEU.withid.cCRE_ID.SRC_ID.JmC_ID.bed', quote=F, sep='\t', col.names=T, row.names=F)
write.table(dm_with_JointClusterID_mat[,c(1:3,4,6,7)], 'ATAC_only/S3V2_IDEAS_mm10_ccre2.cCRE.M.notall0.withid.cCRE_ID.SRC_ID.JmC_ID.bed', quote=F, sep='\t', col.names=T, row.names=F)

###### ct esRP mat for RNA correlation analysis
dh_with_JointClusterID_mat_ct = cbind(apply(cbind(dh[,1], as.character(dh[,2]), as.character(dh[,3])) ,1, function(x) paste(x, collapse='_')), dh[,c(1:6)], all_cCREs_prediction_Y_Jmeta[H_rows], dhs_ctmerge_shared_reorder)
dm_with_JointClusterID_mat_ct = cbind(apply(cbind(dm[,1], as.character(dm[,2]), as.character(dm[,3])) ,1, function(x) paste(x, collapse='_')), dm[,c(1:6)], all_cCREs_prediction_Y_Jmeta[M_rows], dms_ctmerge_shared_reorder)
###### check JMeta number
### hg38
JMeta_vs_KM_H = t(apply(cbind(rownames(dhs_dms_ctmerge_shared_reorder_meansig_H)), 1, function(x) as.numeric(unlist(strsplit(x, ':'))) ))
JMeta_vs_KM_M = t(apply(cbind(rownames(dhs_dms_ctmerge_shared_reorder_meansig_M)), 1, function(x) as.numeric(unlist(strsplit(x, ':'))) ))
JMeta_count_H = c()
JMeta_count_M = c()
for (JMeta_i in unique(JMeta_vs_KM_H[,1])){
  table_vec_names = as.numeric(names(table(dh_with_JointClusterID_mat$J_meta)))
  JMeta_vs_KM_H_i = JMeta_vs_KM_H[is.element(JMeta_vs_KM_H[,1], JMeta_i),1]
  JMeta_vs_KM_M_i = JMeta_vs_KM_M[is.element(JMeta_vs_KM_M[,1], JMeta_i),1]
  print(c(JMeta_i, JMeta_vs_KM_H_i))
  print(c(JMeta_i, JMeta_vs_KM_M_i))
  JMeta_count_H = c(JMeta_count_H, sum(table(dh_with_JointClusterID_mat$J_meta)[is.element(table_vec_names, JMeta_vs_KM_H_i)]))
  JMeta_count_M = c(JMeta_count_M, sum(table(dm_with_JointClusterID_mat$J_meta)[is.element(table_vec_names, JMeta_vs_KM_M_i)]))
}
JMeta_count_H = cbind(unique(JMeta_vs_KM_H[,1]), JMeta_count_H)
JMeta_count_M = cbind(unique(JMeta_vs_KM_M[,1]), JMeta_count_M)
###
JMeta_count_HM = cbind(JMeta_count_H[,2], JMeta_count_M[,2])
JMeta_count_HM[,2] = JMeta_count_HM[,2]/mean(JMeta_count_HM[,2])*mean(JMeta_count_HM[,1])
rownames(JMeta_count_HM) = JMeta_count_H[,1]
colnames(JMeta_count_HM) = c('Human','Mouse')
pdf('ATAC_only/KMPCA.Joint.cluster.reproducible.Jmet.cCRE_count.pdf', height=5,width=3)
pheatmap(log10(JMeta_count_HM), cluster_col=F, cluster_rows=T, cex=1.5, clustering_distance_rows = dist(dhs_dms_ctmerge_shared_reorder_meansig_Jmet_meansig))
dev.off()

cbind(94092:94102, dm_with_JointClusterID_mat[94092:94102,7], as.numeric(all_cCREs_prediction_Y[200342+(94092:94102)]), dms_ctmerge_shared_reorder[94092:94102,])
cbind(196784:196794, dh_with_JointClusterID_mat[196784:196794,7], as.numeric(all_cCREs_prediction_Y[196784:196794]), dhs_ctmerge_shared_reorder[196784:196794,])
#meta_cluster_mat[rownames(meta_cluster_mat)=='GATA1',]
#################################################


#################################################
### gene gene locus
dh_with_JointClusterID = cbind(dh[,c(1:3)], all_cCREs_prediction_Y_Jmeta[H_rows])
dm_with_JointClusterID = cbind(dm[,c(1:3)], all_cCREs_prediction_Y_Jmeta[M_rows])
JointCluster_count_HM = cbind(table(all_cCREs_prediction_Y_Jmeta[H_rows]), table(all_cCREs_prediction_Y_Jmeta[M_rows]))
JointCluster_exp_P_H = JointCluster_count_HM[,1]/sum(JointCluster_count_HM[,1])
JointCluster_exp_P_M = JointCluster_count_HM[,2]/sum(JointCluster_count_HM[,2])
All_Jmet = as.numeric(names(table(all_cCREs_prediction_Y_Jmeta)))

###
get_Function_conserve_cCRE = function(chrH, startH, endH, chrM, startM, endM, Jmet_enrichment_scores_all_zpfdr_thresh, enrich_spe){
#chrH = 'chrX'
#startH = 48760001
#endH = 48836000
#chrM = 'chrX'
#startM = 7919401
#endM = 8020800
add_sm_num = 1
included_rows = ((dh_with_JointClusterID[,1]==chrH) * (dh_with_JointClusterID[,2]>=startH) * (dh_with_JointClusterID[,3]<=endH)) != 0
dh_with_JointClusterID_Gene = dh_with_JointClusterID[included_rows,]
included_rows = ((dm_with_JointClusterID[,1]==chrM) * (dm_with_JointClusterID[,2]>=startM) * (dm_with_JointClusterID[,3]<=endM)) != 0
dm_with_JointClusterID_Gene = dm_with_JointClusterID[included_rows,]
#print(dh_with_JointClusterID_GATA1)
#print(dm_with_JointClusterID_GATA1)
### get share state
### expect state count
JointCluster_count_H_exp = (dim(dh_with_JointClusterID_Gene)[1])*JointCluster_exp_P_H
JointCluster_count_M_exp = (dim(dm_with_JointClusterID_Gene)[1])*JointCluster_exp_P_M
JointCluster_count_HM_exp = (JointCluster_count_H_exp+add_sm_num)*(JointCluster_count_M_exp+add_sm_num)
#print(JointCluster_count_HM_exp)
#print(JointCluster_count_H_exp)
#print(JointCluster_count_M_exp)
#print(dim(dh_with_JointClusterID_Gene)[1])
#print(dim(dm_with_JointClusterID_Gene)[1])
### obs state count
JointCluster_count_H_obs = c()
JointCluster_count_M_obs = c()
for (i in All_Jmet){
JointCluster_count_H_obs = c(JointCluster_count_H_obs, sum(dh_with_JointClusterID_Gene[,4]==i))
JointCluster_count_M_obs = c(JointCluster_count_M_obs, sum(dm_with_JointClusterID_Gene[,4]==i))
}
#print(gene_exp)
Jmet = list()
### get odd ratio
gene_exp = ((JointCluster_count_H_obs+add_sm_num)*(JointCluster_count_M_obs+add_sm_num)) / JointCluster_count_HM_exp
Jmet$Jmet_score = gene_exp
if (enrich_spe){
gene_exp_H = (JointCluster_count_H_obs+add_sm_num) / (JointCluster_count_H_exp+add_sm_num)
gene_exp_M = (JointCluster_count_M_obs+add_sm_num) / (JointCluster_count_M_exp+add_sm_num)
Jmet$Jmet_score_H = gene_exp_H
Jmet$Jmet_score_M = gene_exp_M
}
Jmet$cCRE_H = cbind(dh_with_JointClusterID_Gene, is.element(dh_with_JointClusterID_Gene[,4], as.numeric(names(gene_exp)[gene_exp>=Jmet_enrichment_scores_all_zpfdr_thresh]))*1)
Jmet$cCRE_M = cbind(dm_with_JointClusterID_Gene, is.element(dm_with_JointClusterID_Gene[,4], as.numeric(names(gene_exp)[gene_exp>=Jmet_enrichment_scores_all_zpfdr_thresh]))*1)
return(Jmet)
}
#################################################


#################################################
### read gene pairs
### replace CSF1R gene position 
### from chr5    150053291       150113372       -       CSF1R
### To chr5    150053291       150136554       -       CSF1R
hg38_gene = read.table('/Users/guanjuexiang/Documents/projects/analysis/0813_human_mouse_state_compare_heatmap/hg38.gene.bed', header=F, sep='\t')
mm10_gene = read.table('/Users/guanjuexiang/Documents/projects/analysis/0813_human_mouse_state_compare_heatmap/mm10.gene.bed', header=F, sep='\t')
### mm10
### change Gata1 gene locus from
# chrX  7959260 7978071 - Gata1
# to
# chrX  7959260 7967910 - Gata1 
mm10_gene[,5] = toupper(mm10_gene[,5])
###
shared_genes = hg38_gene[is.element(hg38_gene[,5], mm10_gene[,5]),5]
### get input mat
hg38_gene_shared = hg38_gene[is.element(hg38_gene[,5], shared_genes),]
hg38_gene_shared = hg38_gene_shared[order(hg38_gene_shared[,5]),]
mm10_gene_shared = mm10_gene[is.element(mm10_gene[,5], shared_genes),]
mm10_gene_shared = mm10_gene_shared[order(mm10_gene_shared[,5]),]
### duplicate genes
duplicated_genes = unique(c(names(table(hg38_gene_shared[,5])[table(hg38_gene_shared[,5])>1]), names(table(mm10_gene_shared[,5])[table(mm10_gene_shared[,5])>1])))
### remove duplicate genes
shared_genes = shared_genes[!is.element(shared_genes, duplicated_genes)]
hg38_gene_shared = hg38_gene_shared[!is.element(hg38_gene_shared[,5], duplicated_genes),]
mm10_gene_shared = mm10_gene_shared[!is.element(mm10_gene_shared[,5], duplicated_genes),]
### order genes
shared_genes = shared_genes[order(shared_genes)]
hg38_gene_shared = hg38_gene_shared[order(hg38_gene_shared[,5]),]
mm10_gene_shared = mm10_gene_shared[order(mm10_gene_shared[,5]),]
### expand 50KB
hg38_gene_shared_exp = hg38_gene_shared
# expand 100KB
hg38_gene_shared_exp[hg38_gene_shared_exp[,4]=='+',2] = hg38_gene_shared_exp[hg38_gene_shared_exp[,4]=='+',2] - 100000
# since 2nd column is changed, so the add 200000 should be based on the new 2nd column
hg38_gene_shared_exp[hg38_gene_shared_exp[,4]=='+',3] = hg38_gene_shared_exp[hg38_gene_shared_exp[,4]=='+',2] + 200000
# expand 100KB
hg38_gene_shared_exp[hg38_gene_shared_exp[,4]=='-',2] = hg38_gene_shared_exp[hg38_gene_shared_exp[,4]=='-',3] - 100000
# since 3rd column is NOT changed, so the add 100000 should be based on the original 3rd column
hg38_gene_shared_exp[hg38_gene_shared_exp[,4]=='-',3] = hg38_gene_shared_exp[hg38_gene_shared_exp[,4]=='-',3] + 100000
# remove negative values
hg38_gene_shared_exp[hg38_gene_shared_exp[,2]<0,2] = 0
###
mm10_gene_shared_exp = mm10_gene_shared
# expand 100KB
mm10_gene_shared_exp[mm10_gene_shared_exp[,4]=='+',2] = mm10_gene_shared_exp[mm10_gene_shared_exp[,4]=='+',2] - 100000
# since 2nd column is changed, so the add 200000 should be based on the new 2nd column
mm10_gene_shared_exp[mm10_gene_shared_exp[,4]=='+',3] = mm10_gene_shared_exp[mm10_gene_shared_exp[,4]=='+',2] + 200000
# expand 100KB
mm10_gene_shared_exp[mm10_gene_shared_exp[,4]=='-',2] = mm10_gene_shared_exp[mm10_gene_shared_exp[,4]=='-',3] - 100000
# since 3rd column is NOT changed, so the add 100000 should be based on the original 3rd column
mm10_gene_shared_exp[mm10_gene_shared_exp[,4]=='-',3] = mm10_gene_shared_exp[mm10_gene_shared_exp[,4]=='-',3] + 100000
mm10_gene_shared_exp[mm10_gene_shared_exp[,2]<0,2] = 0
###
#hg38_mm10_gene_shared_exp = cbind(hg38_gene_shared_exp[,1:3], mm10_gene_shared_exp[,1:3])
#################################################


#################################################
### get Gene-by-Jmeta matrix: Round1: Determine enrichment threshold of each Jmet cluster
GATA1_exp = get_Function_conserve_cCRE('chrX', 48760001, 48836000, 'chrX', 7919401,8020800, 2.922063, FALSE)
meta_cluster_mat = as.data.frame(matrix(0, nrow=length(shared_genes), ncol=length(GATA1_exp$Jmet_score)))
cCRE_H_Jmeta = c()
cCRE_M_Jmeta = c()
ptm <- proc.time()
for (i in 1:length(shared_genes)){
  if (i%%1000==0){print(i)}
  shared_genes_i = shared_genes[i]
  bed_H = hg38_gene_shared_exp[i,1:3][1,]
  bed_M = mm10_gene_shared_exp[i,1:3][1,]
  JMeta_i = get_Function_conserve_cCRE(bed_H[1,1], bed_H[1,2], bed_H[1,3], bed_M[1,1], bed_M[1,2], bed_M[1,3], 2.922063, FALSE)
  meta_cluster_mat[i,] = JMeta_i$Jmet_score
  if (dim(JMeta_i$cCRE_H)[1]>0){cCRE_H_Jmeta = rbind(cCRE_H_Jmeta, cbind(JMeta_i$cCRE_H, shared_genes_i))}
  if (dim(JMeta_i$cCRE_M)[1]>0){cCRE_M_Jmeta = rbind(cCRE_M_Jmeta, cbind(JMeta_i$cCRE_M, shared_genes_i))}
}
proc.time() - ptm
###
colnames(meta_cluster_mat) = rownames(JointCluster_count_HM)
rownames(meta_cluster_mat) = shared_genes
### get enrichment threshold for each Jmet
get_thresh = function(x){
  xzp = pnorm((x-mean(x))/sd(x), lower.tail=F)
  thresh0 = 0.05
  thresh1 = 0.01
  xzp = pnorm((x-mean(x[xzp>=thresh1]))/sd(x[xzp>=thresh1]), lower.tail=F)
  return(min(x[xzp<thresh0]))
}
meta_cluster_mat_thresh = apply(meta_cluster_mat,2,get_thresh)
meta_cluster_mat_thresh
#################################################


#################################################
### get Gene-by-Jmeta matrix: Round2
GATA1_exp = get_Function_conserve_cCRE('chrX', 48760001, 48836000, 'chrX', 7919401,8020800, meta_cluster_mat_thresh, TRUE)
meta_cluster_mat = as.data.frame(matrix(0, nrow=length(shared_genes), ncol=length(GATA1_exp$Jmet_score)))
meta_cluster_mat_H = meta_cluster_mat
meta_cluster_mat_M = meta_cluster_mat
cCRE_H_Jmeta = c()
cCRE_M_Jmeta = c()
### change some genes' TSS
hg38_gene_shared_exp[hg38_gene_shared_exp[,5]=='CSF1R',2:3] = c(150036554, 150136554)
###
ptm <- proc.time()
for (i in 1:length(shared_genes)){
  if (i%%1000==0){print(i)}
  shared_genes_i = shared_genes[i]
  bed_H = hg38_gene_shared_exp[i,1:3][1,]
  bed_M = mm10_gene_shared_exp[i,1:3][1,]
  JMeta_i = get_Function_conserve_cCRE(bed_H[1,1], bed_H[1,2], bed_H[1,3], bed_M[1,1], bed_M[1,2], bed_M[1,3], meta_cluster_mat_thresh, TRUE)
  meta_cluster_mat[i,] = JMeta_i$Jmet_score
  meta_cluster_mat_H[i,] = JMeta_i$Jmet_score_H
  meta_cluster_mat_M[i,] = JMeta_i$Jmet_score_M
  if (dim(JMeta_i$cCRE_H)[1]>0){cCRE_H_Jmeta = rbind(cCRE_H_Jmeta, cbind(JMeta_i$cCRE_H, shared_genes_i))}
  if (dim(JMeta_i$cCRE_M)[1]>0){cCRE_M_Jmeta = rbind(cCRE_M_Jmeta, cbind(JMeta_i$cCRE_M, shared_genes_i))}
}
proc.time() - ptm
###
colnames(meta_cluster_mat) = rownames(JointCluster_count_HM)
rownames(meta_cluster_mat) = shared_genes
write.table(cbind(shared_genes, round(meta_cluster_mat,5)), 'ATAC_only/Human_Mouse_shared_genes.Jmeta.enrich.100KB.txt', quote=F, sep='\t', col.names=T, row.names=F)
write.table(cCRE_H_Jmeta, 'ATAC_only/cCRE.Gene100KB.hg38.bed', quote=F, sep='\t', col.names=T, row.names=F)
write.table(cCRE_M_Jmeta, 'ATAC_only/cCRE.Gene100KB.mm10.bed', quote=F, sep='\t', col.names=T, row.names=F)
#################################################
#chrX 48783000 48783200


#################################################
### cluster gene by Jmet Joint-Enrichment
meta_cluster_mat_log2 = log2(meta_cluster_mat+1)
### decide K
set.seed(2019)
meta_cluster_mat_log2_km_ratio = c()
for (k in 2:30){
  meta_cluster_mat_log2_km_test = kmeans(meta_cluster_mat_log2, centers=k)
  meta_cluster_mat_log2_km_ratio = c(meta_cluster_mat_log2_km_ratio, meta_cluster_mat_log2_km_test$tot.withinss/meta_cluster_mat_log2_km_test$betweenss)
}
pdf('ATAC_only/KM_gene_Jmet_enrich.determineK.pdf')
plot(2:30, meta_cluster_mat_log2_km_ratio, cex.axis=2)
lines(2:30, meta_cluster_mat_log2_km_ratio)
abline(v=15)
dev.off()
### KM cluster gene
set.seed(2019)
meta_cluster_mat_log2_km = kmeans(meta_cluster_mat_log2, centers=15)
meta_cluster_mat_log2_km_mean = c()
for (i in as.numeric(names(table(meta_cluster_mat_log2_km$cluster)))){
  meta_cluster_mat_log2_km_mean = rbind(meta_cluster_mat_log2_km_mean, colMeans(meta_cluster_mat_log2[meta_cluster_mat_log2_km$cluster==i,])) 
}
rownames(meta_cluster_mat_log2_km_mean) = as.numeric(names(table(meta_cluster_mat_log2_km$cluster)))
### write gene-by-Jmet-enrichment matrix with Gene KM-ID
meta_cluster_mat_log2_df = cbind(as.data.frame(shared_genes), meta_cluster_mat_log2_km$cluster, meta_cluster_mat_log2)
colnames(meta_cluster_mat_log2_df)[2] = 'GeneKM_ID'
pdf('ATAC_only/meta_cluster_mat_log2_df.hist.1.pdf')
Jmet_enrichment_scores_all = c(as.matrix(meta_cluster_mat_log2_df[,-c(1:2)]))
Jmet_enrichment_scores_all_zpfdr = p.adjust(pnorm((Jmet_enrichment_scores_all - mean(Jmet_enrichment_scores_all)) / sd(Jmet_enrichment_scores_all), lower.tail=F), 'fdr')
Jmet_enrichment_scores_all_zpfdr = p.adjust(pnorm((Jmet_enrichment_scores_all - mean(Jmet_enrichment_scores_all[Jmet_enrichment_scores_all_zpfdr>=0.1])) / sd(Jmet_enrichment_scores_all[Jmet_enrichment_scores_all_zpfdr>=0.1]), lower.tail=F), 'fdr')
Jmet_enrichment_scores_all_zpfdr_thresh = min(Jmet_enrichment_scores_all[Jmet_enrichment_scores_all_zpfdr<0.1])
hist(Jmet_enrichment_scores_all, breaks=50, cex.axis=2)
abline(v=Jmet_enrichment_scores_all_zpfdr_thresh)
box()
dev.off()
Jmet_enrichment_scores_all_zpfdr_thresh
2.270142
write.table(meta_cluster_mat_log2_df, 'ATAC_only/Human_Mouse_shared_genes.Jmeta.enrich.100KB.txt', quote=F, sep='\t', col.names=T, row.names=F)
#################################################


#################################################
### plot heatmap
png('ATAC_only/GeneGroup_by_JMeta.Joint.cluster.png', width = 1000, height = 600)
meta_cluster_mat_GeneGroup_Jmet = c()
used_order = Jmet_order
for (coli in used_order){
  meta_cluster_mat_GeneGroup_Jmet = cbind(meta_cluster_mat_GeneGroup_Jmet, meta_cluster_mat_log2_km_mean[,colnames(meta_cluster_mat)==coli])
}
colnames(meta_cluster_mat_GeneGroup_Jmet) = paste('JmC_',used_order, sep='')
rownames(meta_cluster_mat_GeneGroup_Jmet) = paste('GKM_', rownames(meta_cluster_mat_log2_km_mean), sep='')
pheatmap((t(meta_cluster_mat_GeneGroup_Jmet)), cex=2, cluster_col=F, cluster_rows=F, show_rownames=T, show_colnames=T, clustering_distance_rows = dist(dhs_dms_ctmerge_shared_reorder_meansig_Jmet_meansig), cex=1.5)
dev.off()

pdf('ATAC_only/GeneGroup_by_JMeta.Joint.cluster.pdf', height=9 , width = 12)
meta_cluster_mat_GeneGroup_Jmet = c()
used_order = Jmet_order
for (coli in used_order){
  meta_cluster_mat_GeneGroup_Jmet = cbind(meta_cluster_mat_GeneGroup_Jmet, meta_cluster_mat_log2_km_mean[,colnames(meta_cluster_mat)==coli])
}
colnames(meta_cluster_mat_GeneGroup_Jmet) = paste('JmC_',used_order, sep='')
rownames(meta_cluster_mat_GeneGroup_Jmet) = paste('GKM_', rownames(meta_cluster_mat_log2_km_mean), sep='')
pheatmap((t(meta_cluster_mat_GeneGroup_Jmet)), cex=2, cluster_col=F, cluster_rows=F, show_rownames=T, show_colnames=T, clustering_distance_rows = dist(dhs_dms_ctmerge_shared_reorder_meansig_Jmet_meansig), cex=1.5)
dev.off()


get_enriched_JmC_GKM_pairs = function(meta_cluster_mat_GeneGroup_Jmet){
  meta_cluster_mat_GeneGroup_Jmet_z = (meta_cluster_mat_GeneGroup_Jmet-mean(meta_cluster_mat_GeneGroup_Jmet))/sd(meta_cluster_mat_GeneGroup_Jmet)
  meta_cluster_mat_GeneGroup_Jmet_zp = pnorm(meta_cluster_mat_GeneGroup_Jmet_z, lower.tail=F)
  meta_cluster_mat_GeneGroup_Jmet_zpfdr = p.adjust(meta_cluster_mat_GeneGroup_Jmet_zp, 'fdr')
  min_lim = min(meta_cluster_mat_GeneGroup_Jmet[meta_cluster_mat_GeneGroup_Jmet_zpfdr<0.1])
  ID_pair_list = c()
  meta_cluster_mat_GeneGroup_Jmet_binary = meta_cluster_mat_GeneGroup_Jmet>=min_lim
  for (i in 1:dim(meta_cluster_mat_GeneGroup_Jmet_binary)[1]){
  for (j in 1:dim(meta_cluster_mat_GeneGroup_Jmet_binary)[2]){
  if (meta_cluster_mat_GeneGroup_Jmet_binary[i,j]){
  ID_pair_list = rbind(ID_pair_list, c(colnames(meta_cluster_mat_GeneGroup_Jmet_binary)[j], rownames(meta_cluster_mat_GeneGroup_Jmet_binary)[i]))
  }
  }  
  }
  return(ID_pair_list)
}
meta_cluster_mat_GeneGroup_Jmet_ID_pairs = get_enriched_JmC_GKM_pairs(meta_cluster_mat_GeneGroup_Jmet)
write.table(meta_cluster_mat_GeneGroup_Jmet_ID_pairs, 'ATAC_only/enriched_JmC_ID_KM_ID.100KB.txt', sep='\t', quote=F, col.names=F, row.names=F)


### get binary enrichment based on GeneGroup
meta_cluster_mat_GeneGroup_Jmet_vec = c(meta_cluster_mat_GeneGroup_Jmet)
for (i in 1:10){
  print(length(meta_cluster_mat_GeneGroup_Jmet_vec))
#meta_cluster_mat_GeneGroup_Jmet_vec_zp_fdr = p.adjust(pnorm((meta_cluster_mat_GeneGroup_Jmet_vec - mean(meta_cluster_mat_GeneGroup_Jmet_vec)) / sd(meta_cluster_mat_GeneGroup_Jmet_vec), lower.tail=F), 'fdr')
#meta_cluster_mat_GeneGroup_Jmet_vec = meta_cluster_mat_GeneGroup_Jmet_vec[meta_cluster_mat_GeneGroup_Jmet_vec_zp_fdr>=0.1]
meta_cluster_mat_GeneGroup_Jmet_vec_zp_fdr = pnorm((meta_cluster_mat_GeneGroup_Jmet_vec - mean(meta_cluster_mat_GeneGroup_Jmet_vec)) / sd(meta_cluster_mat_GeneGroup_Jmet_vec), lower.tail=F)
meta_cluster_mat_GeneGroup_Jmet_vec = meta_cluster_mat_GeneGroup_Jmet_vec[meta_cluster_mat_GeneGroup_Jmet_vec_zp_fdr>=0.001]
}
### get Jmet enrichment for each GeneGroup for Downstream analysis
meta_cluster_mat_GeneGroup_Jmet_z = (meta_cluster_mat_GeneGroup_Jmet - mean(meta_cluster_mat_GeneGroup_Jmet_vec)) / sd(meta_cluster_mat_GeneGroup_Jmet_vec)
meta_cluster_mat_GeneGroup_Jmet_zp_01 = t((apply(meta_cluster_mat_GeneGroup_Jmet_z, 2, function(x) pnorm(x, lower.tail=F)) < 0.001) * 1)
meta_cluster_mat_GeneGroup_Jmet_zp_01
### 
#png('Gene_by_JMeta.Joint.cluster.png', width = 1000, height = 500)
pdf('ATAC_only/Gene_by_JMeta.Joint.cluster.pdf', height=5.8, width=9)
meta_cluster_mat_plot = c()
used_order = Jmet_order
for (coli in used_order){
  meta_cluster_mat_plot = cbind(meta_cluster_mat_plot, meta_cluster_mat[,colnames(meta_cluster_mat)==coli])
}
colnames(meta_cluster_mat_plot) = used_order
rownames(meta_cluster_mat_plot) = rownames(meta_cluster_mat)
meta_cluster_mat_plot = meta_cluster_mat_plot[order(meta_cluster_mat_log2_km$cluster),]
pheatmap(log2(t(meta_cluster_mat_plot)+1), cex=1.8, cluster_col=F, cluster_rows=F, show_rownames=T, show_colnames=F, clustering_distance_rows = dist(dhs_dms_ctmerge_shared_reorder_meansig_Jmet_meansig))
dev.off()
pdf('ATAC_only/Gene_by_JMeta.Joint.cluster_H.pdf', height=5, width=9)
meta_cluster_mat_plot_H = c()
used_order = Jmet_order
for (coli in used_order){
  meta_cluster_mat_plot_H = cbind(meta_cluster_mat_plot_H, meta_cluster_mat_H[,colnames(meta_cluster_mat)==coli])
}
colnames(meta_cluster_mat_plot_H) = used_order
rownames(meta_cluster_mat_plot_H) = rownames(meta_cluster_mat)
meta_cluster_mat_plot_H = meta_cluster_mat_plot_H[order(meta_cluster_mat_log2_km$cluster),]
pheatmap(log2(t(meta_cluster_mat_plot_H^2)+1), cex=2, cluster_col=F, cluster_rows=F, show_rownames=T, show_colnames=F, clustering_distance_rows = dist(dhs_dms_ctmerge_shared_reorder_meansig_Jmet_meansig))
dev.off()
pdf('ATAC_only/Gene_by_JMeta.Joint.cluster_M.pdf', height=5, width=9)
meta_cluster_mat_plot_M = c()
used_order = Jmet_order
for (coli in used_order){
  meta_cluster_mat_plot_M = cbind(meta_cluster_mat_plot_M, meta_cluster_mat_M[,colnames(meta_cluster_mat)==coli])
}
colnames(meta_cluster_mat_plot_M) = used_order
rownames(meta_cluster_mat_plot_M) = rownames(meta_cluster_mat)
meta_cluster_mat_plot_M = meta_cluster_mat_plot_M[order(meta_cluster_mat_log2_km$cluster),]
pheatmap(log2(t(meta_cluster_mat_plot_M^2)+1), cex=2, cluster_col=F, cluster_rows=F, show_rownames=T, show_colnames=F, clustering_distance_rows = dist(dhs_dms_ctmerge_shared_reorder_meansig_Jmet_meansig))
dev.off()

get_r2 = function(y1, y2){
  r2 = 1-sum(2*(y1-y2)^2)/sum((mean(y1)-y1)^2+(mean(y2)-y2)^2)
  return(r2)
}

for (i in 1:dim(meta_cluster_mat_plot_H)[2]){
meta_cluster_mat_plot_H_vec = as.numeric(c(meta_cluster_mat_plot_H[,i]))
meta_cluster_mat_plot_M_vec = as.numeric(c(meta_cluster_mat_plot_M[,i]))
#enrich_lm_adj = lm(meta_cluster_mat_plot_H_vec~meta_cluster_mat_plot_M_vec)
#meta_cluster_mat_plot_M_vec_adj = meta_cluster_mat_plot_M_vec * enrich_lm_adj$coefficients[2] + enrich_lm_adj$coefficients[1]
#meta_cluster_mat_plot_M_vec_adj = meta_cluster_mat_plot_M_vec
#M_gene_enrich = log2((meta_cluster_mat_plot_H_vec+0.1)/(meta_cluster_mat_plot_M_vec_adj+0.1))
#A_gene_enrich = log2(meta_cluster_mat_plot_H_vec*meta_cluster_mat_plot_M_vec_adj)
used_id = sample(length(meta_cluster_mat_plot_H_vec), 10000)
png(paste0('ATAC_only/Gene_enrich.heatscatterplot.',colnames(meta_cluster_mat_plot_H)[i],'.100KB.png'))
#heatscatter(A_gene_enrich[used_id], M_gene_enrich[used_id], ylim = c(-max(abs(M_gene_enrich)), max(abs(M_gene_enrich))))
#abline(h=0)
plot_lim_i = c(min(c(meta_cluster_mat_plot_H_vec, meta_cluster_mat_plot_M_vec)), max(c(meta_cluster_mat_plot_H_vec, meta_cluster_mat_plot_M_vec)))+1
print(cor(meta_cluster_mat_plot_H_vec, meta_cluster_mat_plot_M_vec))
heatscatter((meta_cluster_mat_plot_H_vec+1), (meta_cluster_mat_plot_M_vec+1), xlim=plot_lim_i, ylim=plot_lim_i, main=get_r2(meta_cluster_mat_plot_H_vec, meta_cluster_mat_plot_M_vec), log='xy')
abline(0,1)
dev.off()
}
### overall scatterplot & MAplot
used_col_for_plotting_MAplot = !is.element(colnames(meta_cluster_mat_plot_H),c(100))
meta_cluster_mat_plot_H_vec = as.numeric(c(meta_cluster_mat_plot_H[,used_col_for_plotting_MAplot]))
meta_cluster_mat_plot_M_vec = as.numeric(c(meta_cluster_mat_plot_M[,used_col_for_plotting_MAplot]))
#enrich_lm_adj = lm(meta_cluster_mat_plot_H_vec~meta_cluster_mat_plot_M_vec)
#meta_cluster_mat_plot_M_vec_adj = meta_cluster_mat_plot_M_vec * enrich_lm_adj$coefficients[2] + enrich_lm_adj$coefficients[1]
#meta_cluster_mat_plot_M_vec_adj = meta_cluster_mat_plot_M_vec
#M_gene_enrich = log2((meta_cluster_mat_plot_H_vec+0.1)/(meta_cluster_mat_plot_M_vec_adj+0.1))
#A_gene_enrich = log2(meta_cluster_mat_plot_H_vec*meta_cluster_mat_plot_M_vec_adj)
used_id = sample(length(meta_cluster_mat_plot_H_vec), 10000)
png(paste0('ATAC_only/Gene_enrich.heatscatterplot.all.png'))
#heatscatter(A_gene_enrich[used_id], M_gene_enrich[used_id], ylim = c(-max(abs(M_gene_enrich)), max(abs(M_gene_enrich))))
#abline(h=0)
plot_lim_i = c(min(c(meta_cluster_mat_plot_H_vec, meta_cluster_mat_plot_M_vec)), max(c(meta_cluster_mat_plot_H_vec, meta_cluster_mat_plot_M_vec)))+1
print(cor(meta_cluster_mat_plot_H_vec, meta_cluster_mat_plot_M_vec))
#enrich_lm_adj = lm(meta_cluster_mat_plot_H_vec~meta_cluster_mat_plot_M_vec)
#meta_cluster_mat_plot_M_vec_adj = meta_cluster_mat_plot_M_vec * enrich_lm_adj$coefficients[2] + enrich_lm_adj$coefficients[1]
#meta_cluster_mat_plot_M_vec_adj = meta_cluster_mat_plot_M_vec / mean(meta_cluster_mat_plot_M_vec) * mean(meta_cluster_mat_plot_H_vec) 
heatscatter((meta_cluster_mat_plot_H_vec+1), (meta_cluster_mat_plot_M_vec+1), xlim=plot_lim_i, ylim=plot_lim_i, log='xy')
abline(0,1)
dev.off()

png(paste0('ATAC_only/Gene_enrich.blackscatterplot.all.png'))
#heatscatter(A_gene_enrich[used_id], M_gene_enrich[used_id], ylim = c(-max(abs(M_gene_enrich)), max(abs(M_gene_enrich))))
#abline(h=0)
plot_lim_i = c(min(c(meta_cluster_mat_plot_H_vec, meta_cluster_mat_plot_M_vec)), max(c(meta_cluster_mat_plot_H_vec, meta_cluster_mat_plot_M_vec)))+1
print(cor(meta_cluster_mat_plot_H_vec, meta_cluster_mat_plot_M_vec))
#enrich_lm_adj = lm(meta_cluster_mat_plot_H_vec~meta_cluster_mat_plot_M_vec)
#meta_cluster_mat_plot_M_vec_adj = meta_cluster_mat_plot_M_vec * enrich_lm_adj$coefficients[2] + enrich_lm_adj$coefficients[1]
#meta_cluster_mat_plot_M_vec_adj = meta_cluster_mat_plot_M_vec / mean(meta_cluster_mat_plot_M_vec) * mean(meta_cluster_mat_plot_H_vec) 
plot((meta_cluster_mat_plot_H_vec+1), (meta_cluster_mat_plot_M_vec+1), xlim=plot_lim_i, ylim=plot_lim_i, log='xy', pch=16)
abline(0,1, col='red')
dev.off()
###
png('ATAC_only/Gene_enrich.MAplot.png')
meta_cluster_mat_plot_H_vec = as.numeric(c(meta_cluster_mat_plot_H[,used_col_for_plotting_MAplot]))
meta_cluster_mat_plot_M_vec = as.numeric(c(meta_cluster_mat_plot_M[,used_col_for_plotting_MAplot]))
enrich_lm_adj = lm(meta_cluster_mat_plot_H_vec~meta_cluster_mat_plot_M_vec)
#meta_cluster_mat_plot_M_vec_adj = meta_cluster_mat_plot_M_vec * enrich_lm_adj$coefficients[2] + enrich_lm_adj$coefficients[1]
#meta_cluster_mat_plot_M_vec_adj = meta_cluster_mat_plot_M_vec / mean(meta_cluster_mat_plot_M_vec) * mean(meta_cluster_mat_plot_H_vec) 
M_gene_enrich = log2((meta_cluster_mat_plot_H_vec+1)/(meta_cluster_mat_plot_M_vec+1))
A_gene_enrich = log2((meta_cluster_mat_plot_H_vec+1)*(meta_cluster_mat_plot_M_vec+1))
set.seed(2022)
used_id = sample(length(A_gene_enrich), 10000)
heatscatter(A_gene_enrich[used_id], M_gene_enrich[used_id], cex.axis=2, ylim = c(-5, 5)) #, ylim = c(-max(abs(M_gene_enrich)), max(abs(M_gene_enrich)))
abline(h=0)
abline(v=0)
abline(h=2, lty=2)
abline(h=-2, lty=2)
dev.off()

png('ATAC_only/Gene_enrich.MAplot.black.png')
meta_cluster_mat_plot_H_vec = as.numeric(c(meta_cluster_mat_plot_H[,used_col_for_plotting_MAplot]))
meta_cluster_mat_plot_M_vec = as.numeric(c(meta_cluster_mat_plot_M[,used_col_for_plotting_MAplot]))
enrich_lm_adj = lm(meta_cluster_mat_plot_H_vec~meta_cluster_mat_plot_M_vec)
#meta_cluster_mat_plot_M_vec_adj = meta_cluster_mat_plot_M_vec * enrich_lm_adj$coefficients[2] + enrich_lm_adj$coefficients[1]
#meta_cluster_mat_plot_M_vec_adj = meta_cluster_mat_plot_M_vec / mean(meta_cluster_mat_plot_M_vec) * mean(meta_cluster_mat_plot_H_vec) 
M_gene_enrich = log2((meta_cluster_mat_plot_H_vec+1)/(meta_cluster_mat_plot_M_vec+1))
A_gene_enrich = log2((meta_cluster_mat_plot_H_vec+1)*(meta_cluster_mat_plot_M_vec+1))
set.seed(2022)
used_id = sample(length(A_gene_enrich), 10000)
plot(A_gene_enrich[used_id], M_gene_enrich[used_id], cex.axis=2, ylim = c(-5, 5), pch=16) #, ylim = c(-max(abs(M_gene_enrich)), max(abs(M_gene_enrich)))
abline(h=0, col='red')
abline(v=0)
abline(h=2, lty=2)
abline(h=-2, lty=2)
dev.off()
#################################################


#################################################
### prepare SFN files
bash1 = 'mkdir -p /Users/guanjuexiang/Documents/projects/analysis/04_05_2022_coe/coe_analysis/SF_S_N_100KB/'
system(bash1)
### get cCRE with H/MIDs
bash2a = 'cat /Users/guanjuexiang/Documents/projects/analysis/04_05_2022_coe/coe_analysis/SF_S_N_100KB/human_ccre_hg38_sf.bed /Users/guanjuexiang/Documents/projects/analysis/04_05_2022_coe/coe_analysis/SF_S_N_100KB/human_ccre_hg38_s.bed /Users/guanjuexiang/Documents/projects/analysis/04_05_2022_coe/coe_analysis/SF_S_N_100KB/human_ccre_hg38_n.bed | sort -k1,1 -k2,2n > /Users/guanjuexiang/Documents/projects/analysis/04_05_2022_coe/coe_analysis/SF_S_N_100KB/human_ccre_hg38.withHID.bed'
system(bash2a)
bash2b = 'cat /Users/guanjuexiang/Documents/projects/analysis/04_05_2022_coe/coe_analysis/SF_S_N_100KB/mouse_ccre_mm10_sf.bed /Users/guanjuexiang/Documents/projects/analysis/04_05_2022_coe/coe_analysis/SF_S_N_100KB/mouse_ccre_mm10_s.bed /Users/guanjuexiang/Documents/projects/analysis/04_05_2022_coe/coe_analysis/SF_S_N_100KB/mouse_ccre_mm10_n.bed | sort -k1,1 -k2,2n > /Users/guanjuexiang/Documents/projects/analysis/04_05_2022_coe/coe_analysis/SF_S_N_100KB/mouse_ccre_mm10.withMID.bed'
system(bash2b)
### for hg38
bash3a = 'cat /Users/guanjuexiang/Documents/projects/analysis/04_05_2022_coe/coe_analysis/SF_S_N_100KB/mouse_ccre_hg38_sf.bed | awk -F \'\t\' -v OFS=\'\t\' \'{print $2,$3,$4,$1}\' > /Users/guanjuexiang/Documents/projects/analysis/04_05_2022_coe/coe_analysis/SF_S_N_100KB/hg38_sf_s.withMID.bed'
system(bash3a)
bash3b = 'cat /Users/guanjuexiang/Documents/projects/analysis/04_05_2022_coe/coe_analysis/SF_S_N_100KB/human_ccre_hg38_s.bed | awk -F \'\t\' -v OFS=\'\t\' \'{print $1,$2,$3,"X"}\' >> /Users/guanjuexiang/Documents/projects/analysis/04_05_2022_coe/coe_analysis/SF_S_N_100KB/hg38_sf_s.withMID.bed'
system(bash3b)
bash3c = 'sort -k1,1 -k2,2n /Users/guanjuexiang/Documents/projects/analysis/04_05_2022_coe/coe_analysis/SF_S_N_100KB/hg38_sf_s.withMID.bed > /Users/guanjuexiang/Documents/projects/analysis/04_05_2022_coe/coe_analysis/SF_S_N_100KB/hg38_sf_s.withMID.sort.bed'
system(bash3c)
### for mm10
bash5a = 'cat /Users/guanjuexiang/Documents/projects/analysis/04_05_2022_coe/coe_analysis/SF_S_N_100KB/human_ccre_mm10_sf.bed | awk -F \'\t\' -v OFS=\'\t\' \'{print $2,$3,$4,$1}\' > /Users/guanjuexiang/Documents/projects/analysis/04_05_2022_coe/coe_analysis/SF_S_N_100KB/mm10_sf_s.withHID.bed'
system(bash5a)
bash5b = 'cat /Users/guanjuexiang/Documents/projects/analysis/04_05_2022_coe/coe_analysis/SF_S_N_100KB/mouse_ccre_mm10_s.bed | awk -F \'\t\' -v OFS=\'\t\' \'{print $1,$2,$3,"X"}\' >> /Users/guanjuexiang/Documents/projects/analysis/04_05_2022_coe/coe_analysis/SF_S_N_100KB/mm10_sf_s.withHID.bed'
system(bash5b)
bash5c = 'sort -k1,1 -k2,2n /Users/guanjuexiang/Documents/projects/analysis/04_05_2022_coe/coe_analysis/SF_S_N_100KB/mm10_sf_s.withHID.bed > /Users/guanjuexiang/Documents/projects/analysis/04_05_2022_coe/coe_analysis/SF_S_N_100KB/mm10_sf_s.withHID.sort.bed'
system(bash5c)
### add both H & M IDs
bash6 = 'bedtools map -a /Users/guanjuexiang/Documents/projects/analysis/04_05_2022_coe/coe_analysis/SF_S_N_100KB/human_ccre_hg38.withHID.bed -b /Users/guanjuexiang/Documents/projects/analysis/04_05_2022_coe/coe_analysis/SF_S_N_100KB/hg38_sf_s.withMID.sort.bed -c 4 -o concat -null NA > /Users/guanjuexiang/Documents/projects/analysis/04_05_2022_coe/coe_analysis/SF_S_N_100KB/human_ccre_hg38.withHID.with_Mouse_S_MID.bed'
system(bash6)
bash7 = 'bedtools map -a /Users/guanjuexiang/Documents/projects/analysis/04_05_2022_coe/coe_analysis/SF_S_N_100KB/mouse_ccre_mm10.withMID.bed -b /Users/guanjuexiang/Documents/projects/analysis/04_05_2022_coe/coe_analysis/SF_S_N_100KB/mm10_sf_s.withHID.sort.bed -c 4 -o concat -null NA > /Users/guanjuexiang/Documents/projects/analysis/04_05_2022_coe/coe_analysis/SF_S_N_100KB/mouse_ccre_mm10.withMID.with_Human_S_HID.bed'
system(bash7)
#################################################


#################################################
######################## hg38
### Intersect with SF_S cCREs hg38
bedtools1 = 'cat cCRE.Gene100KB.hg38.bed | awk -F \'\t\' \'{if ($1!="NA" && $1!="chr") print $0}\' | sort -k1,1 -k2,2n > cCRE.Gene100KB.hg38.sort.bed'
system(bedtools1)
bedtools2a = 'bedtools map -a cCRE.Gene100KB.hg38.sort.bed -b SF_S_N/human_ccre_hg38.withHID.bed -c 4 -o concat -null NA > cCRE.Gene100KB.hg38.SF_S.HID.bed'
system(bedtools2a)
bedtools2b = 'bedtools map -a cCRE.Gene100KB.hg38.sort.bed -b SF_S_N/human_ccre_hg38.withHID.with_Mouse_S_MID.bed -c 5 -o concat -null NA > cCRE.Gene100KB.hg38.SF_S.MID.bed'
system(bedtools2b)
bedtools2c = 'paste cCRE.Gene100KB.hg38.SF_S.HID.bed cCRE.Gene100KB.hg38.SF_S.MID.bed | cut -f1,2,3,4,5,6,7,14 > cCRE.Gene100KB.hg38.SF_S.bed'
system(bedtools2c)
### Intersect with SF_S cCREs mm10
bedtools3 = 'cat cCRE.Gene100KB.mm10.bed | awk -F \'\t\' \'{if ($1!="NA" && $1!="chr") print $0}\' | sort -k1,1 -k2,2n > cCRE.Gene100KB.mm10.sort.bed '
system(bedtools3)
bedtools4 = 'bedtools map -a cCRE.Gene100KB.mm10.sort.bed -b SF_S_N/mouse_ccre_mm10.withMID.with_Human_S_HID.bed -c 4 -o concat -null NA > cCRE.Gene100KB.mm10.withMID.bed'
system(bedtools4)
bedtools5 = 'cat cCRE.Gene100KB.hg38.SF_S.bed | awk \'{if ($6=="GATA1") print $0}\''
bedtools6 = 'cat cCRE.Gene100KB.mm10.withMID.bed | awk \'{if ($6=="GATA1") print $0}\''
system(bedtools5)
system(bedtools6)
######################## mm10
### Intersect with SF_S cCREs mm10
bedtools1 = 'cat cCRE.Gene100KB.mm10.bed | awk -F \'\t\' \'{if ($1!="NA" && $1!="chr") print $0}\' | sort -k1,1 -k2,2n > cCRE.Gene100KB.mm10.sort.bed'
system(bedtools1)
#bedtools2 = 'bedtools map -a cCRE.Gene100KB.mm10.sort.bed -b SF_S_N/mouse_ccre_mm10.withMID.with_Human_S_HID.bed -c 5 -o concat -null NA > cCRE.Gene100KB.mm10.SF_S.bed'
#system(bedtools2)
bedtools2a = 'bedtools map -a cCRE.Gene100KB.mm10.sort.bed -b SF_S_N/mouse_ccre_mm10.withMID.bed -c 4 -o concat -null NA > cCRE.Gene100KB.mm10.SF_S.MID.bed'
system(bedtools2a)
bedtools2b = 'bedtools map -a cCRE.Gene100KB.mm10.sort.bed -b SF_S_N/mouse_ccre_mm10.withMID.with_Human_S_HID.bed -c 5 -o concat -null NA > cCRE.Gene100KB.mm10.SF_S.HID.bed'
system(bedtools2b)
bedtools2c = 'paste cCRE.Gene100KB.mm10.SF_S.MID.bed cCRE.Gene100KB.mm10.SF_S.HID.bed | cut -f1,2,3,4,5,6,7,14 > cCRE.Gene100KB.mm10.SF_S.bed'
system(bedtools2c)
### Intersect with SF_S cCREs hg38
bedtools3 = 'cat cCRE.Gene100KB.hg38.bed | awk -F \'\t\' \'{if ($1!="NA" && $1!="chr") print $0}\' | sort -k1,1 -k2,2n > cCRE.Gene100KB.hg38.sort.bed '
system(bedtools3)
bedtools4 = 'bedtools map -a cCRE.Gene100KB.hg38.sort.bed -b SF_S_N/human_ccre_hg38.withHID.with_Mouse_S_MID.bed -c 4 -o concat -null NA > cCRE.Gene100KB.hg38.withMID.bed'
system(bedtools4)
bedtools5 = 'cat cCRE.Gene100KB.mm10.SF_S.bed | awk \'{if ($6=="GATA1") print $0}\''
bedtools6 = 'cat cCRE.Gene100KB.hg38.withMID.bed | awk \'{if ($6=="GATA1") print $0}\''
system(bedtools5)
system(bedtools6)
#################################################


#################################################
##################
### read cCRE with SF_S MID hg38
cCRE_H_Jmeta_withMSF_hg38_regions = read.table('cCRE.Gene100KB.hg38.SF_S.bed', header=F)
cCRE_M_Jmeta_withMSF_mm10_regions = read.table('cCRE.Gene100KB.mm10.withMID.bed', header=F)
colnames(cCRE_H_Jmeta_withMSF_hg38_regions) = c('chr','start','end','JmetID','F01','GeneName','HID','MID')
cCRE_H_Jmeta_withMSF_hg38_regions_S = rep(0, dim(cCRE_H_Jmeta_withMSF_hg38_regions)[1])
#all_genes_hg38 = unique(cCRE_H_Jmeta_withMSF_hg38_regions[,6])
#kkk = 0
#for (gene_i in all_genes_hg38){
# if (kkk%%1000==0){print(kkk)}
# kkk = kkk+1
# ### get mm10 cCRE MID in gene_i
# cCRE_M_Jmeta_withMSF_mm10_regions_gene_i = cCRE_M_Jmeta_withMSF_mm10_regions[cCRE_M_Jmeta_withMSF_mm10_regions[,6]==gene_i,7]
# ### get hg38 cCRE MID in gene_i
# cCRE_H_Jmeta_withMSF_hg38_regions_MID_gene_i = cCRE_H_Jmeta_withMSF_hg38_regions[cCRE_H_Jmeta_withMSF_hg38_regions[,6]==gene_i,8]
# ### define the S cCRE 
# cCRE_H_Jmeta_withMSF_hg38_regions_S[cCRE_H_Jmeta_withMSF_hg38_regions[,6]==gene_i] = is.element(cCRE_H_Jmeta_withMSF_hg38_regions_MID_gene_i, cCRE_M_Jmeta_withMSF_mm10_regions_gene_i)*1
#}
###
cCRE_H_Jmeta_withMSF_hg38_regions_S = (!is.na(cCRE_H_Jmeta_withMSF_hg38_regions$MID))*1
### add GeneS01 binary label
cCRE_H_Jmeta_withMSF_hg38_regions_set1 = cbind(cCRE_H_Jmeta_withMSF_hg38_regions, cCRE_H_Jmeta_withMSF_hg38_regions_S)
colnames(cCRE_H_Jmeta_withMSF_hg38_regions_set1) = c(colnames(cCRE_H_Jmeta_withMSF_hg38_regions), 'GeneS01')
write.table(cCRE_H_Jmeta_withMSF_hg38_regions_set1, 'cCRE.Gene100KB.hg38.JmetID.FS01.geneName.HID.MID.S01.bed', quote=F, sep='\t', col.names=F, row.names=F)
#####################
### read cCRE with SF_S MID mm10
cCRE_M_Jmeta_withHSF_mm10_regions = read.table('cCRE.Gene100KB.mm10.SF_S.bed', header=F)
cCRE_H_Jmeta_withHSF_hg38_regions = read.table('cCRE.Gene100KB.hg38.withMID.bed', header=F)
colnames(cCRE_M_Jmeta_withHSF_mm10_regions) = c('chr','start','end','JmetID','F01','GeneName','MID','HID')
cCRE_M_Jmeta_withHSF_mm10_regions_S = rep(0, dim(cCRE_M_Jmeta_withHSF_mm10_regions)[1])
#all_genes_hg38 = unique(cCRE_M_Jmeta_withHSF_mm10_regions[,6])
#kkk = 0
#for (gene_i in all_genes_hg38){
# if (kkk%%1000==0){print(kkk)}
# kkk = kkk+1
# ### get mm10 cCRE MID in gene_i
# cCRE_H_Jmeta_withHSF_hg38_regions_gene_i = cCRE_H_Jmeta_withHSF_hg38_regions[cCRE_H_Jmeta_withHSF_hg38_regions[,6]==gene_i,7]
# ### get hg38 cCRE MID in gene_i
# cCRE_M_Jmeta_withHSF_mm10_regions_HID_gene_i = cCRE_M_Jmeta_withHSF_mm10_regions[cCRE_M_Jmeta_withHSF_mm10_regions[,6]==gene_i,8]
# ### define the S cCRE 
# cCRE_M_Jmeta_withHSF_mm10_regions_S[cCRE_M_Jmeta_withHSF_mm10_regions[,6]==gene_i] = is.element(cCRE_M_Jmeta_withHSF_mm10_regions_HID_gene_i, cCRE_H_Jmeta_withHSF_hg38_regions_gene_i)*1
#}
###
cCRE_M_Jmeta_withHSF_mm10_regions_S = (!is.na(cCRE_M_Jmeta_withHSF_mm10_regions$HID))*1
### add GeneS01 binary label
cCRE_M_Jmeta_withHSF_mm10_regions_set1 = cbind(cCRE_M_Jmeta_withHSF_mm10_regions, cCRE_M_Jmeta_withHSF_mm10_regions_S)
colnames(cCRE_M_Jmeta_withHSF_mm10_regions_set1) = c(colnames(cCRE_M_Jmeta_withHSF_mm10_regions), 'GeneS01')
write.table(cCRE_M_Jmeta_withHSF_mm10_regions_set1, 'cCRE.Gene100KB.mm10.JmetID.FS01.geneName.MID.HID.S01.bed', quote=F, sep='\t', col.names=F, row.names=F)
bedtools5 = 'cat cCRE.Gene100KB.hg38.JmetID.FS01.geneName.HID.MID.S01.bed | awk \'{if ($6=="GATA1") print $0}\''
bedtools6 = 'cat cCRE.Gene100KB.mm10.JmetID.FS01.geneName.MID.HID.S01.bed | awk \'{if ($6=="GATA1") print $0}\''
system(bedtools5)
system(bedtools6)
#################################################


#################################################
### Add protein coding gene TSS label
bedtools1 = 'cat /Users/guanjuexiang/Documents/projects/analysis/0813_human_mouse_state_compare_heatmap/hg38.gene.bed | awk -F \'\t\' -v OFS=\'\t\' -v exp_win=1000 \'{if (($2-exp_win>0) && ($4=="+")) print $1,$2-exp_win, $2+exp_win, $4,$5; if (($3-exp_win>0) && ($4=="-")) print $1,$3-exp_win, $3+exp_win, $4,$5; if (($2-exp_win<0) && ($4=="+")) print $1,0, $2+exp_win, $4,$5; if (($3-exp_win<0) && ($4=="-")) print $1,0, $3+exp_win, $4,$5}\' | sort -k1,1 -k2,2n > /Users/guanjuexiang/Documents/projects/analysis/0813_human_mouse_state_compare_heatmap/hg38.gene.sort.TSS.bed'
bedtools2 = 'bedtools map -a cCRE.Gene100KB.hg38.JmetID.FS01.geneName.HID.MID.S01.bed -b /Users/guanjuexiang/Documents/projects/analysis/0813_human_mouse_state_compare_heatmap/hg38.gene.sort.TSS.bed -c 5 -o concat -null NA > cCRE.Gene100KB.hg38.JmetID.FS01.geneName.HID.MID.S01.TSS.bed'
system(bedtools1)
system(bedtools2)
bedtools1 = 'cat /Users/guanjuexiang/Documents/projects/analysis/0813_human_mouse_state_compare_heatmap/mm10.gene.bed | awk -F \'\t\' -v OFS=\'\t\' -v exp_win=1000 \'{if (($2-exp_win>0) && ($4=="+")) print $1,$2-exp_win, $2+exp_win, $4,$5; if (($3-exp_win>0) && ($4=="-")) print $1,$3-exp_win, $3+exp_win, $4,$5; if (($2-exp_win<0) && ($4=="+")) print $1,0, $2+exp_win, $4,$5; if (($3-exp_win<0) && ($4=="-")) print $1,0, $3+exp_win, $4,$5}\' | sort -k1,1 -k2,2n > /Users/guanjuexiang/Documents/projects/analysis/0813_human_mouse_state_compare_heatmap/mm10.gene.sort.TSS.bed'
bedtools2 = 'bedtools map -a cCRE.Gene100KB.mm10.JmetID.FS01.geneName.MID.HID.S01.bed -b /Users/guanjuexiang/Documents/projects/analysis/0813_human_mouse_state_compare_heatmap/mm10.gene.sort.TSS.bed -c 5 -o concat -null NA > cCRE.Gene100KB.mm10.JmetID.FS01.geneName.MID.HID.S01.TSS.bed'
system(bedtools1)
system(bedtools2)
###
bedtools5 = 'cat cCRE.Gene100KB.hg38.JmetID.FS01.geneName.HID.MID.S01.TSS.bed | awk \'{if ($6=="GATA1") print $0}\''
system(bedtools5)
bedtools5 = 'cat cCRE.Gene100KB.mm10.JmetID.FS01.geneName.MID.HID.S01.TSS.bed | awk \'{if ($6=="GATA1") print $0}\''
system(bedtools5)
#################################################


#################################################
### add Gene Jmet enrichment KM cluster ID
add_Gene_KMID = function(input_mat_file, output_mat_file, meta_cluster_mat_log2_km, cCRE_HM_Jmeta_withMSF_hg38mm10_regions_set1) {
  cCRE_gene_JmetID_FS01_geneName_MID_HID_S01_TSS = read.table(input_mat_file, header=F)
  cCRE_gene_JmetID_FS01_geneName_MID_HID_S01_TSS_GeneKM = rep(0,dim(cCRE_gene_JmetID_FS01_geneName_MID_HID_S01_TSS)[1])
  for (gene_i in unique(cCRE_gene_JmetID_FS01_geneName_MID_HID_S01_TSS[,6])){
  cCRE_gene_JmetID_FS01_geneName_MID_HID_S01_TSS_GeneKM[cCRE_gene_JmetID_FS01_geneName_MID_HID_S01_TSS[,6]==gene_i] = meta_cluster_mat_log2_km$cluster[names(meta_cluster_mat_log2_km$cluster)==gene_i]
  }
  cCRE_gene_JmetID_FS01_geneName_MID_HID_S01_TSS_GeneKM_mat = cbind(cCRE_gene_JmetID_FS01_geneName_MID_HID_S01_TSS, cCRE_gene_JmetID_FS01_geneName_MID_HID_S01_TSS_GeneKM)
  colnames(cCRE_gene_JmetID_FS01_geneName_MID_HID_S01_TSS_GeneKM_mat) = c(colnames(cCRE_HM_Jmeta_withMSF_hg38mm10_regions_set1), 'TSS', 'GeneKMID')
  write.table(cCRE_gene_JmetID_FS01_geneName_MID_HID_S01_TSS_GeneKM_mat, output_mat_file, quote=F, sep='\t', col.names=T, row.names=F)
  return(cCRE_gene_JmetID_FS01_geneName_MID_HID_S01_TSS_GeneKM_mat)
}
cCRE_gene_JmetID_FS01_geneName_MID_HID_S01_TSS_GeneKM_mat_hg38 = add_Gene_KMID('cCRE.Gene100KB.hg38.JmetID.FS01.geneName.HID.MID.S01.TSS.bed', 'cCRE.Gene100KB.hg38.JmetID.FS01.geneName.HID.MID.S01.TSS.GeneKMID.bed', meta_cluster_mat_log2_km, cCRE_H_Jmeta_withMSF_hg38_regions_set1)
cCRE_gene_JmetID_FS01_geneName_MID_HID_S01_TSS_GeneKM_mat_mm10 = add_Gene_KMID('cCRE.Gene100KB.mm10.JmetID.FS01.geneName.MID.HID.S01.TSS.bed', 'cCRE.Gene100KB.mm10.JmetID.FS01.geneName.MID.HID.S01.TSS.GeneKMID.bed', meta_cluster_mat_log2_km, cCRE_M_Jmeta_withHSF_mm10_regions_set1)
### check
cCRE_gene_JmetID_FS01_geneName_MID_HID_S01_TSS_GeneKM_mat_hg38[cCRE_gene_JmetID_FS01_geneName_MID_HID_S01_TSS_GeneKM_mat_hg38[,6]=='GATA1',]
cCRE_gene_JmetID_FS01_geneName_MID_HID_S01_TSS_GeneKM_mat_mm10[cCRE_gene_JmetID_FS01_geneName_MID_HID_S01_TSS_GeneKM_mat_mm10[,6]=='GATA1',]
cCRE_gene_JmetID_FS01_geneName_MID_HID_S01_TSS_GeneKM_mat_hg38[cCRE_gene_JmetID_FS01_geneName_MID_HID_S01_TSS_GeneKM_mat_hg38[,6]=='ALAS2',]
cCRE_gene_JmetID_FS01_geneName_MID_HID_S01_TSS_GeneKM_mat_mm10[cCRE_gene_JmetID_FS01_geneName_MID_HID_S01_TSS_GeneKM_mat_mm10[,6]=='ALAS2',]
cCRE_gene_JmetID_FS01_geneName_MID_HID_S01_TSS_GeneKM_mat_hg38[cCRE_gene_JmetID_FS01_geneName_MID_HID_S01_TSS_GeneKM_mat_hg38[,6]=='CSF1R',]
#################################################


'''
#################################################
### get F01 based on Gene Kmeans clusters
add_F01_KM = function(cCRE_gene_JmetID_FS01_geneName_MID_HID_S01_TSS_GeneKM_mat, meta_cluster_mat_GeneGroup_Jmet_zp_01){
  F01KM = rep(0, dim(cCRE_gene_JmetID_FS01_geneName_MID_HID_S01_TSS_GeneKM_mat)[1])
  for (i in as.numeric(colnames(meta_cluster_mat_GeneGroup_Jmet_zp_01))){
    print(i)
    meta_cluster_mat_GeneGroup_Jmet_zp_01_i = as.numeric(rownames(meta_cluster_mat_GeneGroup_Jmet_zp_01)[meta_cluster_mat_GeneGroup_Jmet_zp_01[,i]==1])
    meta_cluster_mat_GeneGroup_Jmet_zp_01_i_binary = is.element(cCRE_gene_JmetID_FS01_geneName_MID_HID_S01_TSS_GeneKM_mat$JmetID, meta_cluster_mat_GeneGroup_Jmet_zp_01_i) & (cCRE_gene_JmetID_FS01_geneName_MID_HID_S01_TSS_GeneKM_mat$GeneKMID==i)
    F01KM[meta_cluster_mat_GeneGroup_Jmet_zp_01_i_binary] = 1
  }
  return(F01KM)
}
### get F01 based on Gene Kmeans clusters
hg38_F01KM = add_F01_KM(cCRE_gene_JmetID_FS01_geneName_MID_HID_S01_TSS_GeneKM_mat_hg38, meta_cluster_mat_GeneGroup_Jmet_zp_01)
mm10_F01KM = add_F01_KM(cCRE_gene_JmetID_FS01_geneName_MID_HID_S01_TSS_GeneKM_mat_mm10, meta_cluster_mat_GeneGroup_Jmet_zp_01)
### check
cbind(cCRE_gene_JmetID_FS01_geneName_MID_HID_S01_TSS_GeneKM_mat_hg38, hg38_F01KM)[cCRE_gene_JmetID_FS01_geneName_MID_HID_S01_TSS_GeneKM_mat_hg38$GeneName=='GATA1',]
cbind(cCRE_gene_JmetID_FS01_geneName_MID_HID_S01_TSS_GeneKM_mat_hg38, hg38_F01KM)[cCRE_gene_JmetID_FS01_geneName_MID_HID_S01_TSS_GeneKM_mat_hg38$GeneName=='ALAS2',]
### 
hg38_F01_OD = cCRE_gene_JmetID_FS01_geneName_MID_HID_S01_TSS_GeneKM_mat_hg38$F01
mm10_F01_OD = cCRE_gene_JmetID_FS01_geneName_MID_HID_S01_TSS_GeneKM_mat_mm10$F01
### replace original F01
cCRE_gene_JmetID_FS01_geneName_MID_HID_S01_TSS_GeneKM_mat_hg38$F01 = hg38_F01KM
cCRE_gene_JmetID_FS01_geneName_MID_HID_S01_TSS_GeneKM_mat_mm10$F01 = mm10_F01KM
### 
hg38_F01_new = cCRE_gene_JmetID_FS01_geneName_MID_HID_S01_TSS_GeneKM_mat_hg38$F01
mm10_F01_new = cCRE_gene_JmetID_FS01_geneName_MID_HID_S01_TSS_GeneKM_mat_mm10$F01
###
#cCRE_gene_JmetID_FS01_geneName_MID_HID_S01_TSS_GeneKM_mat_hg38$F01 = hg38_F01_OD
#cCRE_gene_JmetID_FS01_geneName_MID_HID_S01_TSS_GeneKM_mat_mm10$F01 = mm10_F01_OD
#################################################
'''


#################################################
### Get Gene-by-Jmet RNA cor mat hg38
RNA_mat = read.table('HumanVISION_RNAseq_hg38_genes_tpm.txt', header=T)
RNA_mat_ct_list = c('HSC', 'HSC', 'ERY', 'ERY', 'TCD4', 'TCD4', 'TCD8', 'TCD8', 'B', 'B', 'CMP', 'CMP', 'MON', 'MON', 'NEU', 'NEU', 'MON', 'MON', 'GMP', 'GMP', 'CFUE', 'NK', 'NK', 'MK', 'MK', 'CLP', 'MPP', 'MPP', 'EOS', 'EOS', 'MEP', 'MEP', 'MK', 'MK', 'CLP', 'ERY', 'ERY', 'ERY', 'HUDEP1', 'HUDEP1', 'HUDEP2', 'HUDEP2', 'CD34', 'CD34')
RNA_mat_sig = RNA_mat[,-c(1:4)]
### ct average
unique_ct = unique(RNA_mat_ct_list)
RNA_mat_ct_ave = matrix(0, nrow=dim(RNA_mat)[1], ncol=length(unique_ct))
k = 0
for (ct in unique_ct){
  k = k+1
  if (sum(RNA_mat_ct_list==ct)>1){
    RNA_mat_ct_ave[,k] = rowMeans(RNA_mat_sig[,RNA_mat_ct_list==ct])
  } else{
    RNA_mat_ct_ave[,k] = RNA_mat_sig[,RNA_mat_ct_list==ct]
  }
}
colnames(RNA_mat_ct_ave) = unique_ct
### shared ct with HM_cts
RNA_mat_ct_ave_shared = c()
for (ct_i in colnames(dhs_dms_ctmerge_shared_reorder_meansig_Jmet_meansig)[-1]){
  RNA_mat_ct_ave_shared = cbind(RNA_mat_ct_ave_shared, RNA_mat_ct_ave[,colnames(RNA_mat_ct_ave)==ct_i])
}
#RNA_mat_ct_ave_shared = RNA_mat_ct_ave[,is.element(colnames(RNA_mat_ct_ave), colnames(dhs_dms_ctmerge_shared_reorder_meansig_Jmet_meansig))]
RNA_mat_ct_ave_shared_ave = rowMeans(RNA_mat_ct_ave_shared)
RNA_mat_ct_ave_shared = cbind(RNA_mat_ct_ave_shared_ave, RNA_mat_ct_ave_shared)
colnames(RNA_mat_ct_ave_shared) = c(colnames(dhs_dms_ctmerge_shared_reorder_meansig_Jmet_meansig))
### log QTnorm
quantile_norm = function(x){
  xm = (x[,1])
  refsig_sort = xm[order(xm)]
  for (i in 1:dim(x)[2]){
    sigtmp = x[,i]
    x[,i] = refsig_sort[rank(sigtmp)]
  }
  return(x)
}
smallnum = 1e-1 
RNA_mat_ct_ave_shared_logqtnorm = quantile_norm(log(RNA_mat_ct_ave_shared+smallnum))
### select used genes
RNA_ENSID2GeneName = read.table('gene.ENS_id2GeneName.hg38.txt', header=F)
colnames(RNA_ENSID2GeneName) = c('gene_id', 'gene_name')
RNA_mat_ENSID_GeneName = merge(RNA_mat[,c(1:4)], RNA_ENSID2GeneName, by='gene_id', all.x=T)
### select used genes
RNA_mat_ENSID_GeneName_used_binary = is.element(RNA_mat_ENSID_GeneName$gene_name, cCRE_gene_JmetID_FS01_geneName_MID_HID_S01_TSS_GeneKM_mat_hg38$GeneName)
RNA_mat_ENSID_GeneName_used = RNA_mat_ENSID_GeneName[RNA_mat_ENSID_GeneName_used_binary,]
RNA_mat_ct_ave_shared_logqtnorm_used = RNA_mat_ct_ave_shared_logqtnorm[RNA_mat_ENSID_GeneName_used_binary,]
rownames(RNA_mat_ct_ave_shared_logqtnorm_used) = RNA_mat_ENSID_GeneName_used[,5]
### gene-by-Jmet RNA-esRP correlation matrix
gene_RNA_Jmet_cor = cor(t(RNA_mat_ct_ave_shared_logqtnorm_used), t(dhs_dms_ctmerge_shared_reorder_meansig_Jmet_meansig))
gene_RNA_Jmet_cor = gene_RNA_Jmet_cor[,order(as.numeric(colnames(gene_RNA_Jmet_cor)))]
png('gene_Jmet_RNA_esRP_cor.png', width = 1000, height = 600)
set.seed(2019)
gene_RNA_Jmet_cor_plot_df = cbind(as.data.frame(rownames(gene_RNA_Jmet_cor)), gene_RNA_Jmet_cor)
colnames(gene_RNA_Jmet_cor_plot_df)[1] = 'shared_genes'

gene_RNA_Jmet_cor_plot_df_merge = merge(gene_RNA_Jmet_cor_plot_df, meta_cluster_mat_log2_df, by='shared_genes', all.x=T)
gene_RNA_Jmet_cor_plot_df_merge_plot = gene_RNA_Jmet_cor_plot_df_merge[order(gene_RNA_Jmet_cor_plot_df_merge$GeneKM_ID),2:(dim(dhs_dms_ctmerge_shared_reorder_meansig_Jmet_meansig)[1]+1)]
#gene_RNA_Jmet_cor_plot_df_merge_plot = gene_RNA_Jmet_cor_plot_df_merge[order(gene_RNA_Jmet_cor_plot_df_merge$GeneKM_ID),11:18]
colnames(gene_RNA_Jmet_cor_plot_df_merge_plot) = 1:dim(dhs_dms_ctmerge_shared_reorder_meansig_Jmet_meansig)[1]

gene_RNA_Jmet_cor_plot_df_merge_plot1 = c()
used_order = Jmet_order
for (coli in used_order){
  gene_RNA_Jmet_cor_plot_df_merge_plot1 = cbind(gene_RNA_Jmet_cor_plot_df_merge_plot1, gene_RNA_Jmet_cor_plot_df_merge_plot[,colnames(gene_RNA_Jmet_cor_plot_df_merge_plot)==coli])
}
colnames(gene_RNA_Jmet_cor_plot_df_merge_plot1) = used_order
pdf('gene_Jmet_RNA_esRP_cor.pdf', height=5, width=9)
pheatmap(t(abs(gene_RNA_Jmet_cor_plot_df_merge_plot1)), cluster_col=F, cluster_rows=F, show_colnames=F, cex=2)
dev.off()
###
pdf('gene_Jmet_RNA_esRP_cor.hist.pdf')
hist(as.numeric(gene_RNA_Jmet_cor), breaks=50, cex=1.5)
box()
dev.off()
print(gene_RNA_Jmet_cor[rownames(gene_RNA_Jmet_cor)=='GATA1',])
#################################################


#################################################
### add Jmet-GeneTPM correlation hg38
#Jmet_gene_pairs = unique(cbind(as.data.frame(cCRE_gene_JmetID_FS01_geneName_MID_HID_S01_TSS_GeneKM_mat_hg38$Jmet), cCRE_gene_JmetID_FS01_geneName_MID_HID_S01_TSS_GeneKM_mat_hg38$GeneName))
#hg38_Jmet_RNA_cor = rep(0, dim(cCRE_gene_JmetID_FS01_geneName_MID_HID_S01_TSS_GeneKM_mat_hg38)[1])
#for (i in 1:dim(Jmet_gene_pairs)[1]){
# if (i%%10000==0){print(i)}
# if (sum(rownames(gene_RNA_Jmet_cor)==Jmet_gene_pairs[i,2])>0){
# used_rows = cCRE_gene_JmetID_FS01_geneName_MID_HID_S01_TSS_GeneKM_mat_hg38$Jmet==Jmet_gene_pairs[i,1] & cCRE_gene_JmetID_FS01_geneName_MID_HID_S01_TSS_GeneKM_mat_hg38$GeneName==Jmet_gene_pairs[i,2]
# hg38_Jmet_RNA_cor[used_rows] = gene_RNA_Jmet_cor[rownames(gene_RNA_Jmet_cor)==Jmet_gene_pairs[i,2],Jmet_gene_pairs[i,1]]
# }
#}
#cCRE_gene_JmetID_FS01_geneName_MID_HID_S01_TSS_GeneKM_mat_hg38_RNAcor = cbind(cCRE_gene_JmetID_FS01_geneName_MID_HID_S01_TSS_GeneKM_mat_hg38, hg38_Jmet_RNA_cor)
### get esRP RNAlogTPM cor
cCRE_hg38_coordinates = cCRE_gene_JmetID_FS01_geneName_MID_HID_S01_TSS_GeneKM_mat_hg38[,1:3]
cCRE_gene_JmetID_FS01_geneName_MID_HID_S01_TSS_GeneKM_mat_hg38_withPKID = cbind(apply(cbind(cCRE_hg38_coordinates[,1], as.character(cCRE_hg38_coordinates[,2]), as.character(cCRE_hg38_coordinates[,3])), 1, function(x) paste(x, collapse='_')), cCRE_gene_JmetID_FS01_geneName_MID_HID_S01_TSS_GeneKM_mat_hg38)
cCRE_gene_PKID = cCRE_gene_JmetID_FS01_geneName_MID_HID_S01_TSS_GeneKM_mat_hg38_withPKID[,1]
cCRE_gene_genes = cCRE_gene_JmetID_FS01_geneName_MID_HID_S01_TSS_GeneKM_mat_hg38_withPKID[,7]
RNA_mat_genes = rownames(RNA_mat_ct_ave_shared_logqtnorm_used)
cCRE_PKID = dh_with_JointClusterID_mat_ct[,1]
cCRE_esRPmat = dh_with_JointClusterID_mat_ct[,-c(1:8)]
set.seed(2019)
cCRE_esRPmat_withNoise = cCRE_esRPmat + matrix(runif(dim(cCRE_esRPmat)[1]*dim(cCRE_esRPmat)[2], -0.0001, 0.0001), dim(cCRE_esRPmat)[1], dim(cCRE_esRPmat)[2])

###
ptm <- proc.time()
cCRE_gene_esRP_RNA_cor = rep(-100,dim(cCRE_gene_JmetID_FS01_geneName_MID_HID_S01_TSS_GeneKM_mat_hg38_withPKID)[1])
for (i in 1:dim(cCRE_gene_JmetID_FS01_geneName_MID_HID_S01_TSS_GeneKM_mat_hg38_withPKID)[1]){
  if (i%%10000==0){print(i)}
  gene_i = cCRE_gene_genes[i]
  cCRE_i = cCRE_gene_PKID[i]
  if (is.element(gene_i, RNA_mat_genes)){
  cCRE_esRP_i = cCRE_esRPmat_withNoise[cCRE_PKID==cCRE_i,]
  RNA_tpm_i = RNA_mat_ct_ave_shared_logqtnorm_used[RNA_mat_genes==gene_i,]
  if (!is.null(dim(RNA_tpm_i))){
      cCRE_gene_esRP_RNA_cor[i] = cor(as.numeric(cCRE_esRP_i), RNA_tpm_i[1,])
    } else{
      cCRE_gene_esRP_RNA_cor[i] = cor(as.numeric(cCRE_esRP_i), RNA_tpm_i)
    }
  }
}
proc.time() - ptm

###
cCRE_gene_JmetID_FS01_geneName_MID_HID_S01_TSS_GeneKM_mat_hg38_RNAcor = cbind(cCRE_gene_JmetID_FS01_geneName_MID_HID_S01_TSS_GeneKM_mat_hg38, cCRE_gene_esRP_RNA_cor)
colnames(cCRE_gene_JmetID_FS01_geneName_MID_HID_S01_TSS_GeneKM_mat_hg38_RNAcor)[dim(cCRE_gene_JmetID_FS01_geneName_MID_HID_S01_TSS_GeneKM_mat_hg38_RNAcor)[2]] = 'RNAcor'
cCRE_gene_JmetID_FS01_geneName_MID_HID_S01_TSS_GeneKM_mat_hg38_RNAcor[cCRE_gene_JmetID_FS01_geneName_MID_HID_S01_TSS_GeneKM_mat_hg38_RNAcor$GeneName=='GATA1',-c(1:3)]
### add SFN label
cCRE_gene_JmetID_FS01_geneName_MID_HID_S01_TSS_GeneKM_mat_hg38_RNAcor_SFN = rep('N', dim(cCRE_gene_JmetID_FS01_geneName_MID_HID_S01_TSS_GeneKM_mat_hg38_RNAcor)[1])
# exclude TSS
exclude_TSS = (cCRE_gene_JmetID_FS01_geneName_MID_HID_S01_TSS_GeneKM_mat_hg38_RNAcor$GeneName!=cCRE_gene_JmetID_FS01_geneName_MID_HID_S01_TSS_GeneKM_mat_hg38_RNAcor$TSS) | is.na(cCRE_gene_JmetID_FS01_geneName_MID_HID_S01_TSS_GeneKM_mat_hg38_RNAcor$TSS)
#exclude_TSS = cCRE_gene_JmetID_FS01_geneName_MID_HID_S01_TSS_GeneKM_mat_hg38_RNAcor$F01==0
#cCRE_gene_JmetID_FS01_geneName_MID_HID_S01_TSS_GeneKM_mat_hg38_RNAcor_SFN[((cCRE_gene_JmetID_FS01_geneName_MID_HID_S01_TSS_GeneKM_mat_hg38_RNAcor$F01==1) | (!exclude_TSS)) & (cCRE_gene_JmetID_FS01_geneName_MID_HID_S01_TSS_GeneKM_mat_hg38_RNAcor$GeneS01==1) ] = 'SF'
#cCRE_gene_JmetID_FS01_geneName_MID_HID_S01_TSS_GeneKM_mat_hg38_RNAcor_SFN[((cCRE_gene_JmetID_FS01_geneName_MID_HID_S01_TSS_GeneKM_mat_hg38_RNAcor$F01==1) | (!exclude_TSS)) & (cCRE_gene_JmetID_FS01_geneName_MID_HID_S01_TSS_GeneKM_mat_hg38_RNAcor$GeneS01==0) ] = 'F'
#cCRE_gene_JmetID_FS01_geneName_MID_HID_S01_TSS_GeneKM_mat_hg38_RNAcor_SFN[((cCRE_gene_JmetID_FS01_geneName_MID_HID_S01_TSS_GeneKM_mat_hg38_RNAcor$F01==0) & (exclude_TSS)) & (cCRE_gene_JmetID_FS01_geneName_MID_HID_S01_TSS_GeneKM_mat_hg38_RNAcor$GeneS01==1) ] = 'S'
cCRE_gene_JmetID_FS01_geneName_MID_HID_S01_TSS_GeneKM_mat_hg38_RNAcor_SFN[(cCRE_gene_JmetID_FS01_geneName_MID_HID_S01_TSS_GeneKM_mat_hg38_RNAcor$F01==1) & (cCRE_gene_JmetID_FS01_geneName_MID_HID_S01_TSS_GeneKM_mat_hg38_RNAcor$GeneS01==1) & (!is.na(cCRE_gene_JmetID_FS01_geneName_MID_HID_S01_TSS_GeneKM_mat_hg38_RNAcor$MID)) & (cCRE_gene_JmetID_FS01_geneName_MID_HID_S01_TSS_GeneKM_mat_hg38_RNAcor$MID!='X') ] = 'SF+'
cCRE_gene_JmetID_FS01_geneName_MID_HID_S01_TSS_GeneKM_mat_hg38_RNAcor_SFN[(cCRE_gene_JmetID_FS01_geneName_MID_HID_S01_TSS_GeneKM_mat_hg38_RNAcor$F01==1) & (cCRE_gene_JmetID_FS01_geneName_MID_HID_S01_TSS_GeneKM_mat_hg38_RNAcor$GeneS01==1) & (cCRE_gene_JmetID_FS01_geneName_MID_HID_S01_TSS_GeneKM_mat_hg38_RNAcor$MID=='X') ] = 'S+'
cCRE_gene_JmetID_FS01_geneName_MID_HID_S01_TSS_GeneKM_mat_hg38_RNAcor_SFN[(cCRE_gene_JmetID_FS01_geneName_MID_HID_S01_TSS_GeneKM_mat_hg38_RNAcor$F01==1) & (cCRE_gene_JmetID_FS01_geneName_MID_HID_S01_TSS_GeneKM_mat_hg38_RNAcor$GeneS01==0) ] = 'J'
cCRE_gene_JmetID_FS01_geneName_MID_HID_S01_TSS_GeneKM_mat_hg38_RNAcor_SFN[(cCRE_gene_JmetID_FS01_geneName_MID_HID_S01_TSS_GeneKM_mat_hg38_RNAcor$F01==0) & (cCRE_gene_JmetID_FS01_geneName_MID_HID_S01_TSS_GeneKM_mat_hg38_RNAcor$GeneS01==1) ] = 'S'
cCRE_gene_JmetID_FS01_geneName_MID_HID_S01_TSS_GeneKM_mat_hg38_RNAcor_SFN[(cCRE_gene_JmetID_FS01_geneName_MID_HID_S01_TSS_GeneKM_mat_hg38_RNAcor$F01==0) & (cCRE_gene_JmetID_FS01_geneName_MID_HID_S01_TSS_GeneKM_mat_hg38_RNAcor$GeneS01==1) & (!is.na(cCRE_gene_JmetID_FS01_geneName_MID_HID_S01_TSS_GeneKM_mat_hg38_RNAcor$MID)) & (cCRE_gene_JmetID_FS01_geneName_MID_HID_S01_TSS_GeneKM_mat_hg38_RNAcor$MID!='X')] = 'SF'
#cCRE_gene_JmetID_FS01_geneName_MID_HID_S01_TSS_GeneKM_mat_hg38_RNAcor_SFN[!exclude_TSS] = 'TSS'
###
cCRE_gene_JmetID_FS01_geneName_MID_HID_S01_TSS_GeneKM_mat_hg38_RNAcor_SFN_mat = cbind(cCRE_gene_JmetID_FS01_geneName_MID_HID_S01_TSS_GeneKM_mat_hg38_RNAcor, cCRE_gene_JmetID_FS01_geneName_MID_HID_S01_TSS_GeneKM_mat_hg38_RNAcor_SFN)
colnames(cCRE_gene_JmetID_FS01_geneName_MID_HID_S01_TSS_GeneKM_mat_hg38_RNAcor_SFN_mat)[dim(cCRE_gene_JmetID_FS01_geneName_MID_HID_S01_TSS_GeneKM_mat_hg38_RNAcor_SFN_mat)[2]] = 'SFNID'
cCRE_gene_JmetID_FS01_geneName_MID_HID_S01_TSS_GeneKM_mat_hg38_RNAcor_SFN_mat[cCRE_gene_JmetID_FS01_geneName_MID_HID_S01_TSS_GeneKM_mat_hg38_RNAcor_SFN_mat$GeneName=='GATA1',]
meta_cluster_mat_log2_df[meta_cluster_mat_log2_df$shared_genes=='GATA1',]
cCRE_gene_JmetID_FS01_geneName_MID_HID_S01_TSS_GeneKM_mat_hg38_RNAcor_SFN_mat[cCRE_gene_JmetID_FS01_geneName_MID_HID_S01_TSS_GeneKM_mat_hg38_RNAcor_SFN_mat$GeneName=='CSF1R',]
meta_cluster_mat_log2_df[meta_cluster_mat_log2_df$shared_genes=='CSF1R',]
write.table(cCRE_gene_JmetID_FS01_geneName_MID_HID_S01_TSS_GeneKM_mat_hg38_RNAcor_SFN_mat, 'cCRE.Gene100KB.hg38.JmetID.FS01.geneName.HID.MID.S01.TSS.GeneKMID.RNAcor.SFNID.bed', quote=F, sep='\t', col.names=T, row.names=F)
### Gene cor of 'SF','F','S','N' of each gene
SFN_ID_list = c('SF+','SF','S+','S','J','N')
score_mat = matrix(0, nrow = length(unique(cCRE_gene_JmetID_FS01_geneName_MID_HID_S01_TSS_GeneKM_mat_hg38_RNAcor_SFN_mat$GeneName)), ncol=length(SFN_ID_list))
k = 0
for (gene_i in unique(cCRE_gene_JmetID_FS01_geneName_MID_HID_S01_TSS_GeneKM_mat_hg38_RNAcor_SFN_mat$GeneName)){
  k = k+1
  if (k%%1000==0){print(k)}
  SFN_mat_gene_i = cCRE_gene_JmetID_FS01_geneName_MID_HID_S01_TSS_GeneKM_mat_hg38_RNAcor_SFN_mat[cCRE_gene_JmetID_FS01_geneName_MID_HID_S01_TSS_GeneKM_mat_hg38_RNAcor_SFN_mat$GeneName==gene_i,]
  SFN_mat_gene_i_dat = unique(SFN_mat_gene_i[,c(4,5,9,12,13,2)])
  score_mat_i = rep(-100, length(SFN_ID_list))
  for (j in 1:length(SFN_ID_list)){
    id = SFN_ID_list[j]
    if (sum(SFN_mat_gene_i_dat$SFNID==id)>0){
      score_mat_i[j] = mean(SFN_mat_gene_i_dat[SFN_mat_gene_i_dat$SFNID==id,4], na.rm=T)
    } 
  }
  score_mat[k,] = score_mat_i
}


### cor boxplot
colnames(score_mat) = SFN_ID_list
score_mat_abs = abs(score_mat)
used_rows = apply(score_mat, 1, sum)>-10
### cor boxplot
score_mat11 = score_mat
score_mat11[score_mat11==-100]=NA
#score_mat11 = abs(score_mat11)
wilcox.test(score_mat11[,1], score_mat11[,2], alternative='greater', paired=T)
wilcox.test(score_mat11[,3], score_mat11[,4], alternative='greater', paired=T)
wilcox.test(score_mat11[,5], score_mat11[,6], alternative='greater', paired=T)
  Wilcoxon signed rank test with continuity correction

data:  score_mat11[, 1] and score_mat11[, 2]
V = 9482405, p-value = 1.615e-12
alternative hypothesis: true location shift is greater than 0


  Wilcoxon signed rank test with continuity correction

data:  score_mat11[, 3] and score_mat11[, 4]
V = 27795990, p-value < 2.2e-16
alternative hypothesis: true location shift is greater than 0


  Wilcoxon signed rank test with continuity correction

data:  score_mat11[, 5] and score_mat11[, 6]
V = 18981879, p-value < 2.2e-16
alternative hypothesis: true location shift is greater than 0


pdf('corabs_SFNJ_mat.box.pdf',width=8)
#boxplot(score_mat[used_rows,]-score_mat[used_rows,4], cex.axis=2)
boxplot(score_mat11, cex.axis=2, ylim=c(-1,1))
points(1:dim(score_mat)[2], colMeans(score_mat11, na.rm=T))
lines(1:dim(score_mat)[2], colMeans(score_mat11, na.rm=T))
abline(h=0)
dev.off()

pdf('corabs_SFNJ_mat.box.cor02.pdf',width=8.5)
score_mat11_02 = score_mat11
plotg_cor_thresh = 0.2
score_mat11_02[score_mat11_02<plotg_cor_thresh] = NA
colnames(score_mat11_02) = c('SF+','SF','S+','S','+','N')
boxplot(score_mat11_02, cex.axis=2, ylim=c(plotg_cor_thresh,1.1), boxwex=0.5, col=c('#FF0000','#ED585E','#FF5500','#4FE54A','#FFAB00','#5868F2'))
abline(h=0)
dev.off()
wilcox.test(score_mat11_02[,1], score_mat11_02[,2], alternative='greater', paired=T)
wilcox.test(score_mat11_02[,3], score_mat11_02[,4], alternative='greater', paired=T)
wilcox.test(score_mat11_02[,5], score_mat11_02[,6], alternative='greater', paired=T)

  Wilcoxon signed rank test with continuity correction

data:  score_mat11_02[, 1] and score_mat11_02[, 2]
V = 197576, p-value = 3.229e-14
alternative hypothesis: true location shift is greater than 0


  Wilcoxon signed rank test with continuity correction

data:  score_mat11_02[, 3] and score_mat11_02[, 4]
V = 544941, p-value < 2.2e-16
alternative hypothesis: true location shift is greater than 0


  Wilcoxon signed rank test with continuity correction

data:  score_mat11_02[, 5] and score_mat11_02[, 6]
V = 326251, p-value < 2.2e-16
alternative hypothesis: true location shift is greater than 0


#pdf('corabs_SFN_mat.box.dif.pdf')
#boxplot(score_mat[used_rows,]-score_mat[used_rows,4], cex.axis=2)
#boxplot(score_mat[used_rows,], cex.axis=2, ylim=c(-1,1))
#abline(h=0)
#dev.off()
########################
### add SFNID to mm10
cCRE_gene_JmetID_FS01_geneName_MID_HID_S01_TSS_GeneKM_mat_mm10_SFN = rep('N', dim(cCRE_gene_JmetID_FS01_geneName_MID_HID_S01_TSS_GeneKM_mat_mm10)[1])
cCRE_gene_JmetID_FS01_geneName_MID_HID_S01_TSS_GeneKM_mat_mm10_SFN[(cCRE_gene_JmetID_FS01_geneName_MID_HID_S01_TSS_GeneKM_mat_mm10$F01==1) & (cCRE_gene_JmetID_FS01_geneName_MID_HID_S01_TSS_GeneKM_mat_mm10$GeneS01==1) & (cCRE_gene_JmetID_FS01_geneName_MID_HID_S01_TSS_GeneKM_mat_mm10$HID!='X') & (!is.na(cCRE_gene_JmetID_FS01_geneName_MID_HID_S01_TSS_GeneKM_mat_mm10$HID)) ] = 'SF+'
cCRE_gene_JmetID_FS01_geneName_MID_HID_S01_TSS_GeneKM_mat_mm10_SFN[(cCRE_gene_JmetID_FS01_geneName_MID_HID_S01_TSS_GeneKM_mat_mm10$F01==1) & (cCRE_gene_JmetID_FS01_geneName_MID_HID_S01_TSS_GeneKM_mat_mm10$GeneS01==1) & (cCRE_gene_JmetID_FS01_geneName_MID_HID_S01_TSS_GeneKM_mat_mm10$HID=='X') ] = 'S+'
cCRE_gene_JmetID_FS01_geneName_MID_HID_S01_TSS_GeneKM_mat_mm10_SFN[(cCRE_gene_JmetID_FS01_geneName_MID_HID_S01_TSS_GeneKM_mat_mm10$F01==1) & (cCRE_gene_JmetID_FS01_geneName_MID_HID_S01_TSS_GeneKM_mat_mm10$GeneS01==0)] = 'J'
cCRE_gene_JmetID_FS01_geneName_MID_HID_S01_TSS_GeneKM_mat_mm10_SFN[(cCRE_gene_JmetID_FS01_geneName_MID_HID_S01_TSS_GeneKM_mat_mm10$F01==0) & (cCRE_gene_JmetID_FS01_geneName_MID_HID_S01_TSS_GeneKM_mat_mm10$GeneS01==1)] = 'S'
cCRE_gene_JmetID_FS01_geneName_MID_HID_S01_TSS_GeneKM_mat_mm10_SFN[(cCRE_gene_JmetID_FS01_geneName_MID_HID_S01_TSS_GeneKM_mat_mm10$F01==0) & (cCRE_gene_JmetID_FS01_geneName_MID_HID_S01_TSS_GeneKM_mat_mm10$GeneS01==1) & (!is.na(cCRE_gene_JmetID_FS01_geneName_MID_HID_S01_TSS_GeneKM_mat_mm10$HID)) & (cCRE_gene_JmetID_FS01_geneName_MID_HID_S01_TSS_GeneKM_mat_mm10$HID!='X')] = 'SF'
cCRE_gene_JmetID_FS01_geneName_MID_HID_S01_TSS_GeneKM_mat_mm10_SFN_mat = cbind(cCRE_gene_JmetID_FS01_geneName_MID_HID_S01_TSS_GeneKM_mat_mm10, cCRE_gene_JmetID_FS01_geneName_MID_HID_S01_TSS_GeneKM_mat_mm10_SFN)
colnames(cCRE_gene_JmetID_FS01_geneName_MID_HID_S01_TSS_GeneKM_mat_mm10_SFN_mat)[dim(cCRE_gene_JmetID_FS01_geneName_MID_HID_S01_TSS_GeneKM_mat_mm10_SFN_mat)[2]] = 'SFNID'
### add P to mm10
cCRE_gene_JmetID_FS01_geneName_MID_HID_S01_TSS_GeneKM_mat_mm10_SFN_mat[,10] = toupper(cCRE_gene_JmetID_FS01_geneName_MID_HID_S01_TSS_GeneKM_mat_mm10_SFN_mat[,10])
exclude_TSS_mm10 = (cCRE_gene_JmetID_FS01_geneName_MID_HID_S01_TSS_GeneKM_mat_mm10_SFN_mat$GeneName!=cCRE_gene_JmetID_FS01_geneName_MID_HID_S01_TSS_GeneKM_mat_mm10_SFN_mat$TSS) | is.na(cCRE_gene_JmetID_FS01_geneName_MID_HID_S01_TSS_GeneKM_mat_mm10_SFN_mat$TSS)
#cCRE_gene_JmetID_FS01_geneName_MID_HID_S01_TSS_GeneKM_mat_mm10_SFN_mat$SFNID[!exclude_TSS_mm10] = 'TSS'
###
write.table(cCRE_gene_JmetID_FS01_geneName_MID_HID_S01_TSS_GeneKM_mat_mm10_SFN_mat, 'cCRE.Gene100KB.mm10.JmetID.FS01.geneName.MID.HID.S01.TSS.GeneKMID.SFNID.bed', quote=F, sep='\t', col.names=T, row.names=F)
write.table(cCRE_gene_JmetID_FS01_geneName_MID_HID_S01_TSS_GeneKM_mat_hg38_RNAcor_SFN_mat[,colnames(cCRE_gene_JmetID_FS01_geneName_MID_HID_S01_TSS_GeneKM_mat_hg38_RNAcor_SFN_mat)!='RNAcor'], 'cCRE.Gene100KB.hg38.JmetID.FS01.geneName.HID.MID.S01.TSS.GeneKMID.SFNID.bed', quote=F, sep='\t', col.names=T, row.names=F)
write.table(cCRE_gene_JmetID_FS01_geneName_MID_HID_S01_TSS_GeneKM_mat_mm10_SFN_mat[,c(1:4,11,6,12)], 'cCRE.Gene100KB.mm10.JmC_ID.geneName.GeneKMID.SFN+_ID.bed', quote=F, sep='\t', col.names=T, row.names=F)
write.table(cCRE_gene_JmetID_FS01_geneName_MID_HID_S01_TSS_GeneKM_mat_hg38_RNAcor_SFN_mat[,c(1:4,11,6,13)], 'cCRE.Gene100KB.hg38.JmC_ID.geneName.GeneKMID.SFN+_ID.bed', quote=F, sep='\t', col.names=T, row.names=F)
### save enriched JmC-GKM pairs
JmC_GKM_pairs0 = read.table('enriched_JmC_ID_KM_ID.100KB.txt', header=F)
JmC_GKM_pairs1 = t(apply(JmC_GKM_pairs0, 1, function(x) c(unlist(strsplit(x[1], '_')), unlist(strsplit(x[2], '_'))) ))
JmC_GKM_pairs2 = apply(JmC_GKM_pairs1, 1, function(x) paste(x[2], x[4], sep='_'))
###
JmC_GKM_pairs_hg38 = read.table('cCRE.Gene100KB.hg38.JmC_ID.geneName.GeneKMID.SFN+_ID.bed', header=T)
JmC_GKM_pairs_hg38_ID = apply(JmC_GKM_pairs_hg38, 1, function(x) paste(as.numeric(x[4]), as.numeric(x[5]), sep='_'))
JmC_GKM_pairs_hg38_enriched = JmC_GKM_pairs_hg38[is.element(JmC_GKM_pairs_hg38_ID, JmC_GKM_pairs2),]
write.table(JmC_GKM_pairs_hg38_enriched, 'cCRE.Gene100KB.hg38.JmC_ID.geneName.GeneKMID.SFN+_ID.enriched.bed', quote=F, sep='\t', col.names=T, row.names=F)
###
JmC_GKM_pairs_mm10 = read.table('cCRE.Gene100KB.mm10.JmC_ID.geneName.GeneKMID.SFN+_ID.bed', header=T)
JmC_GKM_pairs_mm10_ID = apply(JmC_GKM_pairs_mm10, 1, function(x) paste(as.numeric(x[4]), as.numeric(x[5]), sep='_'))
JmC_GKM_pairs_mm10_enriched = JmC_GKM_pairs_mm10[is.element(JmC_GKM_pairs_mm10_ID, JmC_GKM_pairs2),]
write.table(JmC_GKM_pairs_mm10_enriched, 'cCRE.Gene100KB.mm10.JmC_ID.geneName.GeneKMID.SFN+_ID.enriched.bed', quote=F, sep='\t', col.names=T, row.names=F)

#chr start end JmC_ID  GeneName  Gene_KMeans_ID  SFN+_ID
### info
# 118435 unique hg38 cCREs & 66106 unique mm10 cCREs
# 234762 unique hg38 cCREs-protein-coding-gene pairs & 145109 unique mm10 cCREs-protein-coding-gene pairs (TSS +/-50kb)
table(cCRE_gene_JmetID_FS01_geneName_MID_HID_S01_TSS_GeneKM_mat_hg38_RNAcor_SFN_mat$SFNID)
table(cCRE_gene_JmetID_FS01_geneName_MID_HID_S01_TSS_GeneKM_mat_mm10_SFN_mat$SFNID)
# hg38
    J     N     S    S+    SF   SF+ 
26179 52417 64871 37932 33532 19831 
# mm10
    J     N     S    S+    SF   SF+ 
13740 26072 26387 15112 39999 23799
kkk = table(apply(cCRE_gene_JmetID_FS01_geneName_MID_HID_S01_TSS_GeneKM_mat_mm10_SFN_mat,1,function(x) paste(x[11],x[12])))
kkk1 = table(apply(bbbbbb,1, function(x) paste(x[11],x[12])))
png('GeneKMID_SFN+ID.count.Human_Mouse.png')
heatscatter(kkk,kkk1)
heatscatter(as.numeric(kkk),as.numeric(kkk1))
abline(0,1)
dev.off()
#################################################


#################################################
### count number 
SFNP_ID_order = SFN_ID_list
Jmet_i_SFNP_count = c()
Jmet_i_SFNP_enrich = c()
for (Jmet_i in used_order){
  print(Jmet_i)
  Jmet_i_binary = cCRE_gene_JmetID_FS01_geneName_MID_HID_S01_TSS_GeneKM_mat_mm10_SFN_mat$JmetID==Jmet_i
  SFNP_ID_order_count_i = c()
  SFNP_ID_order_enrich_i = c()
  for (SFNP_j in SFNP_ID_order){
    obs_Jmet_i_SFNP_j = sum((Jmet_i_binary * (cCRE_gene_JmetID_FS01_geneName_MID_HID_S01_TSS_GeneKM_mat_mm10_SFN_mat$SFNID==SFNP_j))!=0)
    exp_Jmet_i_SFNP_j = sum(Jmet_i_binary) / dim(cCRE_gene_JmetID_FS01_geneName_MID_HID_S01_TSS_GeneKM_mat_mm10_SFN_mat)[1] * sum(cCRE_gene_JmetID_FS01_geneName_MID_HID_S01_TSS_GeneKM_mat_mm10_SFN_mat$SFNID==SFNP_j) 
    SFNP_ID_order_count_i = c(SFNP_ID_order_count_i, obs_Jmet_i_SFNP_j)
    SFNP_ID_order_enrich_i = c(SFNP_ID_order_enrich_i, (obs_Jmet_i_SFNP_j+100)/(exp_Jmet_i_SFNP_j+100))
  }
  Jmet_i_SFNP_count = rbind(Jmet_i_SFNP_count, SFNP_ID_order_count_i)
  Jmet_i_SFNP_enrich = rbind(Jmet_i_SFNP_enrich, SFNP_ID_order_enrich_i)
}
###
colnames(Jmet_i_SFNP_count) = SFNP_ID_order
rownames(Jmet_i_SFNP_count) = used_order
colnames(Jmet_i_SFNP_enrich) = SFNP_ID_order
rownames(Jmet_i_SFNP_enrich) = used_order
###
pdf('Jmet_i_SFNP_count.pdf', width=4, height=5)
pheatmap(log10(Jmet_i_SFNP_count), cluster_cols=F, cluster_rows=F, cex=1.5)
dev.off()
pdf('Jmet_i_SFNP_enrich.pdf', width=4, height=5)
pheatmap(log10(Jmet_i_SFNP_enrich), cluster_cols=F, cluster_rows=F, cex=1.5)
dev.off()
#################################################


#################################################
### write Gene-cCRE loop file
hg38_gene_locus = read.table('/Users/guanjuexiang/Documents/projects/analysis/0813_human_mouse_state_compare_heatmap/hg38.gene.bed', header=F)
hg38_gene_locus_TSS = hg38_gene_locus
hg38_gene_locus_TSS[hg38_gene_locus_TSS[,4]=='+',3] = hg38_gene_locus_TSS[hg38_gene_locus_TSS[,4]=='+',2]+1
hg38_gene_locus_TSS[hg38_gene_locus_TSS[,4]=='-',2] = hg38_gene_locus_TSS[hg38_gene_locus_TSS[,4]=='-',3]-1
### modify CSF1R gene locus
hg38_gene_locus_TSS[hg38_gene_locus_TSS[,5]=='CSF1R',c(2:3)] = c(150086553, 150086554)
### add TSS position
hg38_SFN_loops = cCRE_gene_JmetID_FS01_geneName_MID_HID_S01_TSS_GeneKM_mat_hg38_RNAcor_SFN_mat
hg38_SFN_loops_TSS = hg38_SFN_loops[,1:3]
k = 0
for (gene_i in unique(hg38_SFN_loops$GeneName)){
  k = k+1
  if (k%%1000==0){print(k)}
  used_n = sum(hg38_SFN_loops$GeneName==gene_i)
  TSS_gene_i = hg38_gene_locus_TSS[hg38_gene_locus_TSS[,5]==gene_i,1:3]
  hg38_SFN_loops_TSS[hg38_SFN_loops$GeneName==gene_i,] = cbind(as.data.frame(rep(TSS_gene_i[1,1], used_n)), rep(TSS_gene_i[1,2], used_n), rep(TSS_gene_i[1,3], used_n))
}
###
hg38_SFN_loops = cbind(hg38_SFN_loops[,1:5], round(hg38_SFN_loops[12]*1000), hg38_SFN_loops[,13], rep('#DEDEDE', dim(hg38_SFN_loops)[1]), hg38_SFN_loops[,c(1:3,7)], rep('.', dim(hg38_SFN_loops)[1]), hg38_SFN_loops_TSS, hg38_SFN_loops$GeneName, rep('.', dim(hg38_SFN_loops)[1]) )
### remove neg
#hg38_SFN_loops[(hg38_SFN_loops[,6]<0) | (is.na(hg38_SFN_loops[,6])),6] = 0
#hg38_SFN_loops[(is.na(hg38_SFN_loops[,6])),6] = 0
### change color
#hg38_SFN_loops[hg38_SFN_loops[,7]=='SFJ',8] = '#FF0000'
#hg38_SFN_loops[hg38_SFN_loops[,7]=='SF',8] = '#DE00FF'
#hg38_SFN_loops[hg38_SFN_loops[,7]=='SJ',8] = '#FF5500'
#hg38_SFN_loops[hg38_SFN_loops[,7]=='S',8] = '#006FFF'
#hg38_SFN_loops[hg38_SFN_loops[,7]=='J',8] = '#FFAB00'
#hg38_SFN_loops[hg38_SFN_loops[,7]=='N',8] = '#DEDEDE'
#hg38_SFN_loops[hg38_SFN_loops[,7]=='TSS',8] = '#000000'

hg38_SFN_loops[hg38_SFN_loops[,7]=='SF+',8] = '#FF0000'
hg38_SFN_loops[hg38_SFN_loops[,7]=='SF',8] = '#ED585E'
hg38_SFN_loops[hg38_SFN_loops[,7]=='S+',8] = '#FF5500'
hg38_SFN_loops[hg38_SFN_loops[,7]=='S',8] = '#4FE54A'
hg38_SFN_loops[hg38_SFN_loops[,7]=='J',8] = '#FFAB00'
hg38_SFN_loops[hg38_SFN_loops[,7]=='N',8] = '#5868F2'
#hg38_SFN_loops[hg38_SFN_loops[,7]=='TSS',8] = '#000000'

hg38_SFN_loops[hg38_SFN_loops[,17]=='GATA1',]
hg38_gene_shared_exp[hg38_gene_shared_exp[,5]=='GATA1',]
hg38_SFN_loops[hg38_SFN_loops[,17]=='CSF1R',]
hg38_gene_shared_exp[hg38_gene_shared_exp[,5]=='CSF1R',]
options("scipen"=100, "digits"=4)
hg38_SFN_loops$HID = apply(cbind(hg38_SFN_loops$JmetID, hg38_SFN_loops$HID), 1, function(x) paste(x[1], x[2], sep=':'))
write.table(hg38_SFN_loops, 'hg38.SFN.loop.interact', quote=F, sep='\t', col.names=F, row.names=F)

### get new loop interact file with hierachy TSS>SFJ>SF; TSS>SJ>S; TSS>J>N
### hg38
ptm = proc.time()
unique_HID = unique(hg38_SFN_loops$HID)
hg38_SFN_loops_new = hg38_SFN_loops
k = 0
for ( cCRE_i in unique_HID){
if (k%%1000==0){print(k)}
  d_cCRE_i = hg38_SFN_loops[hg38_SFN_loops$HID==cCRE_i,]
if (dim(d_cCRE_i)[1]==1){
  k = k+1
  hg38_SFN_loops_new[k,] = d_cCRE_i
} else {
  if (sum(d_cCRE_i[,7]=='TSS')>0){
    d_cCRE_i = d_cCRE_i[is.element(d_cCRE_i[,7], c('TSS')),]
  } else if (sum(d_cCRE_i[,7]=='SF+')>0) {
    d_cCRE_i = d_cCRE_i[is.element(d_cCRE_i[,7], c('SF+')),]
  } else if (sum(d_cCRE_i[,7]=='S+')>0) {
    d_cCRE_i = d_cCRE_i[is.element(d_cCRE_i[,7], c('S+')),]
  } else if (sum(d_cCRE_i[,7]=='J')>0) {
    d_cCRE_i = d_cCRE_i[is.element(d_cCRE_i[,7], c('J')),]
  }
  ###
  if (dim(d_cCRE_i)[1]==1){
    k = k+1
    hg38_SFN_loops_new[k,] = d_cCRE_i
  } else{
    k_vec = (k+1):(k+dim(d_cCRE_i)[1])
    hg38_SFN_loops_new[k_vec,] = d_cCRE_i
    k = k_vec[length(k_vec)]
  }
}
}
hg38_SFN_loops_new1 = hg38_SFN_loops_new[1:k,]
proc.time() - ptm

### get the cCRE pass a certain correlation threshold
cor_pass_P_mat = c()
cor_thresh_vec = seq(0,1000,by=100)
for (cor_thresh in cor_thresh_vec){
#PP = sum(hg38_SFN_loops_new1[hg38_SFN_loops_new1[,7]=='TSS',6]>cor_thresh) / length(hg38_SFN_loops_new1[hg38_SFN_loops_new1[,7]=='TSS',6])
PSAF = sum(hg38_SFN_loops_new1[hg38_SFN_loops_new1[,7]=='SF+',6]>cor_thresh) / length(hg38_SFN_loops_new1[hg38_SFN_loops_new1[,7]=='SF+',6])
PSA = sum(hg38_SFN_loops_new1[hg38_SFN_loops_new1[,7]=='SF',6]>cor_thresh) / length(hg38_SFN_loops_new1[hg38_SFN_loops_new1[,7]=='SF',6])
PSF = sum(hg38_SFN_loops_new1[hg38_SFN_loops_new1[,7]=='S+',6]>cor_thresh) / length(hg38_SFN_loops_new1[hg38_SFN_loops_new1[,7]=='S+',6])
PS = sum(hg38_SFN_loops_new1[hg38_SFN_loops_new1[,7]=='S',6]>cor_thresh) / length(hg38_SFN_loops_new1[hg38_SFN_loops_new1[,7]=='S',6])
PF = sum(hg38_SFN_loops_new1[hg38_SFN_loops_new1[,7]=='J',6]>cor_thresh) / length(hg38_SFN_loops_new1[hg38_SFN_loops_new1[,7]=='J',6])
PN = sum(hg38_SFN_loops_new1[hg38_SFN_loops_new1[,7]=='N',6]>cor_thresh) / length(hg38_SFN_loops_new1[hg38_SFN_loops_new1[,7]=='N',6])
BG = sum(hg38_SFN_loops_new1[,6]>cor_thresh) / length(hg38_SFN_loops_new1[,6])
cor_pass_P_mat = cbind(cor_pass_P_mat, c(PSAF,PSF,PSF,PS,PF,PN,BG))
}
pdf('prop.cCRE.pass.corthresh.pdf', width=5, height=5)
cor_thresh_vec_plot = cor_thresh_vec/1000
plot(cor_thresh_vec_plot, cor_pass_P_mat[1,], type='l', cex.axis=1.5)
lines(cor_thresh_vec_plot, cor_pass_P_mat[1,], col='#FF0000')
lines(cor_thresh_vec_plot, cor_pass_P_mat[2,], col='#ED585E')
lines(cor_thresh_vec_plot, cor_pass_P_mat[3,], col='#FF5500')
lines(cor_thresh_vec_plot, cor_pass_P_mat[4,], col='#4FE54A')
lines(cor_thresh_vec_plot, cor_pass_P_mat[5,], col='#FFAB00')
lines(cor_thresh_vec_plot, cor_pass_P_mat[6,], col='#5868F2')
points(cor_thresh_vec_plot, cor_pass_P_mat[1,], col='#FF0000')
points(cor_thresh_vec_plot, cor_pass_P_mat[2,], col='#ED585E')
points(cor_thresh_vec_plot, cor_pass_P_mat[3,], col='#FF5500')
points(cor_thresh_vec_plot, cor_pass_P_mat[4,], col='#4FE54A')
points(cor_thresh_vec_plot, cor_pass_P_mat[5,], col='#FFAB00')
points(cor_thresh_vec_plot, cor_pass_P_mat[6,], col='#5868F2')
dev.off()

### write interaction passing correlation threshold
library(tidyverse)
library(grDevices)
hg38_SFN_loops_new2 = hg38_SFN_loops_new1
hg38_SFN_loops_new2[,5] = hg38_SFN_loops_new1[,6]
hg38_SFN_loops_new2[,6] = hg38_SFN_loops_new1[,5]
colnames(hg38_SFN_loops_new2)[c(6,5)] = colnames(hg38_SFN_loops_new1)[c(6,5)]
write.table(hg38_SFN_loops_new2, 'hg38.SFNJ.loop.OD.interact', quote=F, sep='\t', col.names=F, row.names=F)
### write interact file with header
bash1 = 'cat loop.header.txt > hg38.SFN.loop.interact.tmp'
bash2 = 'cat hg38.SFNJ.loop.OD.interact | awk -F \'\t\' -v OFS=\'\t\' \'{if ($5>=0 || $7=="TSS") print $0; else print $1,$2,$3,$4,0,$6,$7,$8,$9,$10,$11,$12,$13,$14,$15,$16,$17,$18}\' | sort -u >> hg38.SFN.loop.interact.tmp && mv hg38.SFN.loop.interact.tmp hg38.SFN+.loop.All.interact'
system(bash1)
system(bash2)
bash1 = 'cat loop.header.txt > hg38.SFN.loop.interact.tmp'
bash2 = 'cat hg38.SFNJ.loop.OD.interact | awk -F \'\t\' \'{if ($5>=200 || $7=="TSS") print $0}\' | sort -u >> hg38.SFN.loop.interact.tmp && mv hg38.SFN.loop.interact.tmp hg38.SFNJ.loop.cor02.interact'
system(bash1)
system(bash2)
bash1 = 'cat loop.header.txt > hg38.SFN.loop.interact.tmp'
bash2 = 'cat hg38.SFNJ.loop.OD.interact | awk -F \'\t\' -v OFS=\'\t\'  \'{if ($5>=0) print $0; else print $1,$2,$3,$4,0,$6,$7,$8,$9,$10,$11,$12,$13,$14,$15,$16,$17,$18}\' | awk -F \'\t\' \'{if ($17=="GATA1") print $0}\' >> hg38.SFN.loop.interact.tmp && mv hg38.SFN.loop.interact.tmp hg38.SFN+.loop.GATA1.interact'
system(bash1)
system(bash2)

cor_mat_hg38_SFN_loops_new1 = c()
set.seed(2019)
SFN_ID_list1 = SFN_ID_list
for (SFN_ID in SFN_ID_list1){
  cor_SFN_ID = hg38_SFN_loops[hg38_SFN_loops[,7]==SFN_ID,6]
  #cor_SFN_ID = cor_SFN_ID[abs(cor_SFN_ID)>200]
  cor_mat_hg38_SFN_loops_new1 = cbind(cor_mat_hg38_SFN_loops_new1, cor_SFN_ID[sample(length(cor_SFN_ID), 10000, replace=T)])
}
colnames(cor_mat_hg38_SFN_loops_new1) = SFN_ID_list
pdf('corabs_SFN_mat.box.cor02.cCREeach.pdf',width=8.5)
cor_mat_hg38_SFN_loops_new1_plot = cor_mat_hg38_SFN_loops_new1/1000
cor_mat_hg38_SFN_loops_new1_plot[cor_mat_hg38_SFN_loops_new1_plot<(0.2)] = NA
boxplot((cor_mat_hg38_SFN_loops_new1_plot), cex.axis=2)
points(1:dim(cor_mat_hg38_SFN_loops_new1_plot)[2], colMeans(cor_mat_hg38_SFN_loops_new1_plot, na.rm=T))
lines(1:dim(cor_mat_hg38_SFN_loops_new1_plot)[2], colMeans(cor_mat_hg38_SFN_loops_new1_plot, na.rm=T))
abline(h=0)
dev.off()


wilcox.test(cor_mat_hg38_SFN_loops_new1_plot[,1], cor_mat_hg38_SFN_loops_new1_plot[,2], alternative='greater', paired=F)
wilcox.test(cor_mat_hg38_SFN_loops_new1_plot[,3], cor_mat_hg38_SFN_loops_new1_plot[,4], alternative='greater', paired=F)
wilcox.test(cor_mat_hg38_SFN_loops_new1_plot[,5], cor_mat_hg38_SFN_loops_new1_plot[,6], alternative='greater', paired=F)

  Wilcoxon rank sum test with continuity correction

data:  cor_mat_hg38_SFN_loops_new1_plot[, 1] and cor_mat_hg38_SFN_loops_new1_plot[, 2]
W = 7998060, p-value = 0.0000000000007
alternative hypothesis: true location shift is greater than 0


  Wilcoxon rank sum test with continuity correction

data:  cor_mat_hg38_SFN_loops_new1_plot[, 3] and cor_mat_hg38_SFN_loops_new1_plot[, 4]
W = 7821947, p-value = 0.0000000000000007
alternative hypothesis: true location shift is greater than 0


  Wilcoxon rank sum test with continuity correction

data:  cor_mat_hg38_SFN_loops_new1_plot[, 5] and cor_mat_hg38_SFN_loops_new1_plot[, 6]
W = 7176522, p-value <0.0000000000000002
alternative hypothesis: true location shift is greater than 0


#################################################
### Output cCRE with SFNID and colors file
hg38_SFN_loops_new1_bed = unique(hg38_SFN_loops_new1[,c(1:3,12,7,8)])
hg38_SFN_loops_new1_bed[,4] = apply(hg38_SFN_loops_new1_bed, 1, function(x) unlist(strsplit(x[4], ':'))[2])
hg38_SFN_loops_new1_bed1 = merge(hg38_SFN_loops_new1_bed, cCRE_gene_JmetID_FS01_geneName_MID_HID_S01_TSS_GeneKM_mat_hg38_RNAcor_SFN_mat[,c(7,8)], by='HID')
hg38_SFN_loops_new1_bed1 = unique(hg38_SFN_loops_new1_bed1)
### get cCRE name
hg38_SFN_loops_new1_bed1_cCRE_name = apply(hg38_SFN_loops_new1_bed1, 1, function(x) paste(x[c(5,1,7)], collapse=':'))
### convert rgb to num
hg38_SFN_loops_new1_bed1_rgb = apply(apply(hg38_SFN_loops_new1_bed1, 1, function(x) col2rgb(x[6])), 2, function(x) paste(x, collapse=','))
### reorder columns
hg38_SFN_loops_new1_bed2 = cbind(hg38_SFN_loops_new1_bed1[,2:4], hg38_SFN_loops_new1_bed1_cCRE_name, rep(0,dim(hg38_SFN_loops_new1_bed1)[1]), rep('+',dim(hg38_SFN_loops_new1_bed1)[1]), rep(0,dim(hg38_SFN_loops_new1_bed1)[1]), rep(0,dim(hg38_SFN_loops_new1_bed1)[1]), hg38_SFN_loops_new1_bed1_rgb)
###
write.table(hg38_SFN_loops_new1_bed2, 'hg38.cCRE.SFNJ.colored.bed', quote=F, sep='\t', col.names=F, row.names=F)
###
bash1 = 'cat bed.SFNJ.header.txt > hg38.cCRE.SFNJ.colored.bed.tmp'
bash2 = 'cat hg38.cCRE.SFNJ.colored.bed | awk \'{print $0}\' >> hg38.cCRE.SFNJ.colored.bed.tmp && mv hg38.cCRE.SFNJ.colored.bed.tmp hg38.cCRE.SFN+.colored.bed'
system(bash1)
system(bash2)
table(apply(hg38_SFN_loops_new1_bed2,1,function(x) unlist(strsplit(x[4], ':'))[1]))
    J     N     S    S+    SF   SF+ 
16802 25948 29189 22449 13885 10162
#################################################


#################################################
### get cCRE with SFN label colors
### 
bash0 = 'cat bed.SFN.header.txt > /Users/guanjuexiang/Documents/projects/analysis/04_05_2022_coe/coe_analysis/SF_S_N/human_ccre_hg38.withHID.withSFN_color.bed'
system(bash0)
bash0 = 'cat /Users/guanjuexiang/Documents/projects/analysis/04_05_2022_coe/coe_analysis/SF_S_N/human_ccre_hg38_sf.bed | awk -F \'\t\' -v OFS=\'\t\' \'{print $1,$2,$3,$4,0,"+",0,0,"237,88,94"}\' >> /Users/guanjuexiang/Documents/projects/analysis/04_05_2022_coe/coe_analysis/SF_S_N/human_ccre_hg38.withHID.withSFN_color.bed'
system(bash0)
bash0 = 'cat /Users/guanjuexiang/Documents/projects/analysis/04_05_2022_coe/coe_analysis/SF_S_N/human_ccre_hg38_s.bed | awk -F \'\t\' -v OFS=\'\t\' \'{print $1,$2,$3,$4,0,"+",0,0,"79,229,74"}\' >> /Users/guanjuexiang/Documents/projects/analysis/04_05_2022_coe/coe_analysis/SF_S_N/human_ccre_hg38.withHID.withSFN_color.bed'
system(bash0)
bash0 = 'cat /Users/guanjuexiang/Documents/projects/analysis/04_05_2022_coe/coe_analysis/SF_S_N/human_ccre_hg38_n.bed | awk -F \'\t\' -v OFS=\'\t\' \'{print $1,$2,$3,$4,0,"+",0,0,"88,104,242"}\' >> /Users/guanjuexiang/Documents/projects/analysis/04_05_2022_coe/coe_analysis/SF_S_N/human_ccre_hg38.withHID.withSFN_color.bed'
system(bash0)
#################################################


#################################################
### write Gene-cCRE loop file
mm10_gene_locus = read.table('/Users/guanjuexiang/Documents/projects/analysis/0813_human_mouse_state_compare_heatmap/mm10.gene.bed', header=F)
mm10_gene_locus_TSS = mm10_gene_locus
mm10_gene_locus_TSS[mm10_gene_locus_TSS[,4]=='+',3] = mm10_gene_locus_TSS[mm10_gene_locus_TSS[,4]=='+',2]+1
mm10_gene_locus_TSS[mm10_gene_locus_TSS[,4]=='-',2] = mm10_gene_locus_TSS[mm10_gene_locus_TSS[,4]=='-',3]-1
### add TSS position
mm10_SFN_loops = cCRE_gene_JmetID_FS01_geneName_MID_HID_S01_TSS_GeneKM_mat_mm10_SFN_mat
mm10_SFN_loops_TSS = mm10_SFN_loops[,1:3]
k = 0
for (gene_i in unique(mm10_SFN_loops$GeneName)){
  k = k+1
  if (k%%1000==0){print(k)}
  used_n = sum(mm10_SFN_loops$GeneName==gene_i)
  TSS_gene_i = mm10_gene_locus_TSS[toupper(mm10_gene_locus_TSS[,5])==gene_i,1:3]
  mm10_SFN_loops_TSS[mm10_SFN_loops$GeneName==gene_i,] = cbind(as.data.frame(rep(TSS_gene_i[1,1], used_n)), rep(TSS_gene_i[1,2], used_n), rep(TSS_gene_i[1,3], used_n))
}
###
mm10_SFN_loops = cbind(mm10_SFN_loops[,1:5], round(rep(1, dim(mm10_SFN_loops)[1])*1000), mm10_SFN_loops[,12], rep('#DEDEDE', dim(mm10_SFN_loops)[1]), mm10_SFN_loops[,c(1:3,7)], rep('.', dim(mm10_SFN_loops)[1]), mm10_SFN_loops_TSS, mm10_SFN_loops$GeneName, rep('.', dim(mm10_SFN_loops)[1]) )
### remove neg
mm10_SFN_loops[mm10_SFN_loops[,7]=='SF+',8] = '#FF0000'
mm10_SFN_loops[mm10_SFN_loops[,7]=='SF',8] = '#ED585E'
mm10_SFN_loops[mm10_SFN_loops[,7]=='S+',8] = '#FF5500'
mm10_SFN_loops[mm10_SFN_loops[,7]=='S',8] = '#4FE54A'
mm10_SFN_loops[mm10_SFN_loops[,7]=='J',8] = '#FFAB00'
mm10_SFN_loops[mm10_SFN_loops[,7]=='N',8] = '#5868F2'

mm10_SFN_loops[mm10_SFN_loops[,17]=='GATA1',]
mm10_gene_shared_exp[mm10_gene_shared_exp[,5]=='GATA1',]
mm10_SFN_loops[mm10_SFN_loops[,17]=='CSF1R',]
mm10_gene_shared_exp[mm10_gene_shared_exp[,5]=='CSF1R',]
options("scipen"=100, "digits"=4)
mm10_SFN_loops$HID = apply(cbind(mm10_SFN_loops$JmetID, mm10_SFN_loops$MID), 1, function(x) paste(x[1], x[2], sep=':'))
write.table(mm10_SFN_loops, 'mm10.SFN.loop.interact', quote=F, sep='\t', col.names=F, row.names=F)

### get new loop interact file with hierachy TSS>SFJ>SF; TSS>SJ>S; TSS>J>N
### mm10
ptm = proc.time()
unique_MID = unique(mm10_SFN_loops$HID)
mm10_SFN_loops_new = mm10_SFN_loops
k = 0
for ( cCRE_i in unique_MID){
if (k%%1000==0){print(k)}
  d_cCRE_i = mm10_SFN_loops[mm10_SFN_loops$HID==cCRE_i,]
if (dim(d_cCRE_i)[1]==1){
  k = k+1
  mm10_SFN_loops_new[k,] = d_cCRE_i
} else {
  if (sum(d_cCRE_i[,7]=='TSS')>0){
    d_cCRE_i = d_cCRE_i[is.element(d_cCRE_i[,7], c('TSS')),]
  } else if (sum(d_cCRE_i[,7]=='SF+')>0) {
    d_cCRE_i = d_cCRE_i[is.element(d_cCRE_i[,7], c('SF+')),]
  } else if (sum(d_cCRE_i[,7]=='S+')>0) {
    d_cCRE_i = d_cCRE_i[is.element(d_cCRE_i[,7], c('S+')),]
  } else if (sum(d_cCRE_i[,7]=='J')>0) {
    d_cCRE_i = d_cCRE_i[is.element(d_cCRE_i[,7], c('J')),]
  }
  ###
  if (dim(d_cCRE_i)[1]==1){
    k = k+1
    mm10_SFN_loops_new[k,] = d_cCRE_i
  } else{
    k_vec = (k+1):(k+dim(d_cCRE_i)[1])
    mm10_SFN_loops_new[k_vec,] = d_cCRE_i
    k = k_vec[length(k_vec)]
  }
}
}
mm10_SFN_loops_new1 = mm10_SFN_loops_new[1:k,]
proc.time() - ptm

### write interaction passing correlation threshold
library(tidyverse)
library(grDevices)
mm10_SFN_loops_new2 = mm10_SFN_loops_new1
mm10_SFN_loops_new2[,5] = mm10_SFN_loops_new1[,6]
mm10_SFN_loops_new2[,6] = mm10_SFN_loops_new1[,5]
colnames(mm10_SFN_loops_new2)[c(6,5)] = colnames(mm10_SFN_loops_new1)[c(6,5)]
mm10_SFN_loops_new2[,12] = mm10_SFN_loops_new2[,19]
mm10_SFN_loops_new2 = mm10_SFN_loops_new2[,c(1:18)]
write.table(mm10_SFN_loops_new2, 'mm10.SFNJ.loop.OD.interact', quote=F, sep='\t', col.names=F, row.names=F)
### write interact file with header
bash1 = 'cat loop.mm10.header.txt > mm10.SFN.loop.interact.tmp'
bash2 = 'cat mm10.SFNJ.loop.OD.interact | awk -F \'\t\' -v OFS=\'\t\' \'{if ($5>=0 || $7=="TSS") print $0; else print $1,$2,$3,$4,0,$6,$7,$8,$9,$10,$11,$12,$13,$14,$15,$16,$17,$18}\' | sort -u >> mm10.SFN.loop.interact.tmp && mv mm10.SFN.loop.interact.tmp mm10.SFN+.loop.All.interact'
system(bash1)
system(bash2)
bash1 = 'cat loop.mm10.header.txt > mm10.SFN.loop.interact.tmp'
bash2 = 'cat mm10.SFNJ.loop.OD.interact | awk -F \'\t\' -v OFS=\'\t\' \'{if ($5>=0 || $7=="TSS") print $0; else print $1,$2,$3,$4,0,$6,$7,$8,$9,$10,$11,$12,$13,$14,$15,$16,$17,$18}\' | sort -u | awk -F \'\t\' \'{if ($17=="GATA1") print $0}\' >> mm10.SFN.loop.interact.tmp && mv mm10.SFN.loop.interact.tmp mm10.SFN+.loop.GATA1.interact'
system(bash1)
system(bash2)


#################################################
### Output cCRE with SFNID and colors file
mm10_SFN_loops_new1_bed = unique(mm10_SFN_loops_new1[,c(1:3,12,7,8)])
#mm10_SFN_loops_new1_bed[,4] = apply(mm10_SFN_loops_new1_bed, 1, function(x) unlist(strsplit(x[4], ':'))[2])
mm10_SFN_loops_new1_bed1 = merge(mm10_SFN_loops_new1_bed, cCRE_gene_JmetID_FS01_geneName_MID_HID_S01_TSS_GeneKM_mat_mm10_SFN_mat[,c(7,8)], by='MID')
mm10_SFN_loops_new1_bed1 = unique(mm10_SFN_loops_new1_bed1)
### get cCRE name
mm10_SFN_loops_new1_bed1_cCRE_name = apply(mm10_SFN_loops_new1_bed1, 1, function(x) paste(x[c(5,1,7)], collapse=':'))
### convert rgb to num
mm10_SFN_loops_new1_bed1_rgb = apply(apply(mm10_SFN_loops_new1_bed1, 1, function(x) col2rgb(x[6])), 2, function(x) paste(x, collapse=','))
### reorder columns
mm10_SFN_loops_new1_bed2 = cbind(mm10_SFN_loops_new1_bed1[,2:4], mm10_SFN_loops_new1_bed1_cCRE_name, rep(0,dim(mm10_SFN_loops_new1_bed1)[1]), rep('+',dim(mm10_SFN_loops_new1_bed1)[1]), rep(0,dim(mm10_SFN_loops_new1_bed1)[1]), rep(0,dim(mm10_SFN_loops_new1_bed1)[1]), mm10_SFN_loops_new1_bed1_rgb)
###
write.table(mm10_SFN_loops_new1_bed2, 'mm10.cCRE.SFNJ.colored.bed', quote=F, sep='\t', col.names=F, row.names=F)
###
bash1 = 'cat bed.mm10.SFNJ.header.txt > mm10.cCRE.SFNJ.colored.bed.tmp'
bash2 = 'cat mm10.cCRE.SFNJ.colored.bed | awk \'{print $0}\' >> mm10.cCRE.SFNJ.colored.bed.tmp && mv mm10.cCRE.SFNJ.colored.bed.tmp mm10.cCRE.SFN+.colored.bed'
system(bash1)
system(bash2)
table(apply(mm10_SFN_loops_new1_bed2,1,function(x) unlist(strsplit(x[4], ':'))[1]))
    J     N     S    S+    SF   SF+ 
 8123 10735 11837  9386 14377 11648
#################################################


#################################################
### get cCRE with SFN label colors
### 
bash0 = 'cat bed.mm10.SFN.header.txt > /Users/guanjuexiang/Documents/projects/analysis/04_05_2022_coe/coe_analysis/SF_S_N/mouse_ccre_mm10.withHID.withSFN_color.bed'
system(bash0)
bash0 = 'cat /Users/guanjuexiang/Documents/projects/analysis/04_05_2022_coe/coe_analysis/SF_S_N/mouse_ccre_mm10_sf.bed | awk -F \'\t\' -v OFS=\'\t\' \'{print $1,$2,$3,$4,0,"+",0,0,"237,88,94"}\' >> /Users/guanjuexiang/Documents/projects/analysis/04_05_2022_coe/coe_analysis/SF_S_N/mouse_ccre_mm10.withHID.withSFN_color.bed'
system(bash0)
bash0 = 'cat /Users/guanjuexiang/Documents/projects/analysis/04_05_2022_coe/coe_analysis/SF_S_N/mouse_ccre_mm10_s.bed | awk -F \'\t\' -v OFS=\'\t\' \'{print $1,$2,$3,$4,0,"+",0,0,"79,229,74"}\' >> /Users/guanjuexiang/Documents/projects/analysis/04_05_2022_coe/coe_analysis/SF_S_N/mouse_ccre_mm10.withHID.withSFN_color.bed'
system(bash0)
bash0 = 'cat /Users/guanjuexiang/Documents/projects/analysis/04_05_2022_coe/coe_analysis/SF_S_N/mouse_ccre_mm10_n.bed | awk -F \'\t\' -v OFS=\'\t\' \'{print $1,$2,$3,$4,0,"+",0,0,"88,104,242"}\' >> /Users/guanjuexiang/Documents/projects/analysis/04_05_2022_coe/coe_analysis/SF_S_N/mouse_ccre_mm10.withHID.withSFN_color.bed'
system(bash0)
#################################################


bash1 = 'cat loop.header.txt > hg38.SFN.loop.interact.all.tmp'
bash2 = 'cat hg38.SFN.loop.OD.interact | awk -F \'\t\' \'{print $0}\' >> hg38.SFN.loop.interact.all.tmp && mv hg38.SFN.loop.interact.all.tmp hg38.SFN.loop.all.interact'
system(bash1)
system(bash2)


###
bash1 = 'cat loop.header.txt > hg38.SFN.loop.interact.tmp'
bash2 = 'cat hg38.SFN.loop.interact | awk -F \'\t\' -v OFS=\'\t\' \'{if ($17=="GATA1") print $0}\' >> hg38.SFN.loop.interact.tmp && mv hg38.SFN.loop.interact.tmp hg38.SFN.loop.GATA1.interact'
system(bash1)
system(bash2)
bash1 = 'cat loop.header.txt > hg38.SFN.loop.interact.tmp'
bash2 = 'cat hg38.SFN.loop.interact | awk -F \'\t\' -v OFS=\'\t\' \'{if ($17=="HBA1") print $0}\' >> hg38.SFN.loop.interact.tmp && mv hg38.SFN.loop.interact.tmp hg38.SFN.loop.HBA1.interact'
system(bash1)
system(bash2)
bash1 = 'cat loop.header.txt > hg38.SFN.loop.interact.tmp'
bash2 = 'cat hg38.SFN.loop.interact | awk -F \'\t\' -v OFS=\'\t\' \'{if ($17=="CSF1R") print $0}\' >> hg38.SFN.loop.interact.tmp && mv hg38.SFN.loop.interact.tmp hg38.SFN.loop.CSF1R.interact'
system(bash1)
system(bash2)
bash1 = 'cat loop.header.txt > hg38.SFN.loop.interact.tmp'
bash2 = 'cat hg38.SFN.loop.interact | awk -F \'\t\' -v OFS=\'\t\' \'{if ($17=="IFNG") print $0}\' >> hg38.SFN.loop.interact.tmp && mv hg38.SFN.loop.interact.tmp hg38.SFN.loop.IFNG.interact'
system(bash1)
system(bash2)
bash1 = 'cat loop.header.txt > hg38.SFN.loop.interact.tmp'
bash2 = 'cat hg38.SFN.loop.interact | awk -F \'\t\' -v OFS=\'\t\' \'{if ($17=="H2AC4") print $0}\' >> hg38.SFN.loop.interact.tmp && mv hg38.SFN.loop.interact.tmp hg38.SFN.loop.H2AC4.interact'
system(bash1)
system(bash2)
bash1 = 'cat loop.header.txt > hg38.SFN.loop.interact.tmp'
bash2 = 'cat hg38.SFN.loop.interact | awk -F \'\t\' -v OFS=\'\t\' \'{if ($17=="ALAS2") print $0}\' >> hg38.SFN.loop.interact.tmp && mv hg38.SFN.loop.interact.tmp hg38.SFN.loop.ALAS2.interact'
system(bash1)
system(bash2)
hg38_gene_shared_exp[hg38_gene_shared_exp[,5]=='ALAS2',]
#################################################


#################################################
### write Gene-cCRE loop file mm10
mm10_gene_locus = read.table('/Users/guanjuexiang/Documents/projects/analysis/0813_human_mouse_state_compare_heatmap/mm10.gene.bed', header=F)
mm10_gene_locus_TSS = mm10_gene_locus
mm10_gene_locus_TSS[mm10_gene_locus_TSS[,4]=='+',3] = mm10_gene_locus_TSS[mm10_gene_locus_TSS[,4]=='+',2]+1
mm10_gene_locus_TSS[mm10_gene_locus_TSS[,4]=='-',2] = mm10_gene_locus_TSS[mm10_gene_locus_TSS[,4]=='-',3]-1
mm10_gene_locus_TSS[,5] = toupper(mm10_gene_locus_TSS[,5])
### add TSS position
mm10_SFN_loops = cCRE_gene_JmetID_FS01_geneName_MID_HID_S01_TSS_GeneKM_mat_mm10_SFN_mat
mm10_SFN_loops_TSS = mm10_SFN_loops[,1:3]
k = 0
for (gene_i in unique(mm10_SFN_loops$GeneName)){
  k = k+1
  if (k%%1000==0){print(k)}
  used_n = sum(mm10_SFN_loops$GeneName==gene_i)
  TSS_gene_i = mm10_gene_locus_TSS[mm10_gene_locus_TSS[,5]==gene_i,1:3]
  mm10_SFN_loops_TSS[mm10_SFN_loops$GeneName==gene_i,] = cbind(as.data.frame(rep(TSS_gene_i[1,1], used_n)), rep(TSS_gene_i[1,2], used_n), rep(TSS_gene_i[1,3], used_n))
}
###
mm10_SFN_loops = cbind(mm10_SFN_loops[,1:5], round(rep(1,dim(mm10_SFN_loops)[1])*1000), mm10_SFN_loops[,12], rep('#DEDEDE', dim(mm10_SFN_loops)[1]), mm10_SFN_loops[,c(1:3,7)], rep('.', dim(mm10_SFN_loops)[1]), mm10_SFN_loops_TSS, mm10_SFN_loops$GeneName, rep('.', dim(mm10_SFN_loops)[1]) )
### remove neg
mm10_SFN_loops[(mm10_SFN_loops[,6]<0) | (is.na(mm10_SFN_loops[,6])),6] = 0
### change color
mm10_SFN_loops[mm10_SFN_loops[,7]=='SF',8] = '#FD0606'
mm10_SFN_loops[mm10_SFN_loops[,7]=='F',8] = '#FFAB00'
mm10_SFN_loops[mm10_SFN_loops[,7]=='S',8] = '#696969'
mm10_SFN_loops[mm10_SFN_loops[,7]=='N',8] = '#DEDEDE'
mm10_SFN_loops[mm10_SFN_loops[,7]=='P',8] = '#FF0000'
mm10_SFN_loops[mm10_SFN_loops[,17]=='GATA1',]
mm10_gene_shared_exp[mm10_gene_shared_exp[,5]=='GATA1',]
#mm10_SFN_loops[mm10_SFN_loops[,17]=='CSF1R',]
#mm10_gene_shared_exp[mm10_gene_shared_exp[,5]=='CSF1R',]
options("scipen"=100, "digits"=4)
mm10_SFN_loops$MID = apply(cbind(mm10_SFN_loops$JmetID, mm10_SFN_loops$MID), 1, function(x) paste(x[1], x[2], sep=':'))
write.table(mm10_SFN_loops[mm10_SFN_loops[,6]>=0,], 'mm10.SFN.loop.interact', quote=F, sep='\t', col.names=F, row.names=F)
bash1 = 'cat loop.header.txt > mm10.SFN.loop.interact.tmp'
bash2 = 'cat mm10.SFN.loop.interact >> mm10.SFN.loop.interact.tmp && mv mm10.SFN.loop.interact.tmp mm10.SFN.loop.interact'
system(bash1)
system(bash2)
bash1 = 'cat loop.header.txt > mm10.SFN.loop.interact.tmp'
bash2 = 'cat mm10.SFN.loop.interact | awk -F \'\t\' -v OFS=\'\t\' \'{if ($17=="GATA1") print $0}\' >> mm10.SFN.loop.interact.tmp && mv mm10.SFN.loop.interact.tmp mm10.SFN.loop.GATA1.interact'
system(bash1)
system(bash2)
bash1 = 'cat loop.header.txt > mm10.SFN.loop.interact.tmp'
bash2 = 'cat mm10.SFN.loop.interact | awk -F \'\t\' -v OFS=\'\t\' \'{if ($17=="ALAS2") print $0}\' >> mm10.SFN.loop.interact.tmp && mv mm10.SFN.loop.interact.tmp mm10.SFN.loop.ALAS2.interact'
system(bash1)
system(bash2)
#################################################


### get Gasperini Enh-gene pairs
bash_i = 'wget http://hgdownload.soe.ucsc.edu/admin/exe/macOSX.x86_64/liftOver'
system(bash_i)
bash_i = 'wget wget https://hgdownload.soe.ucsc.edu/goldenPath/hg19/liftOver/hg19ToHg38.over.chain.gz'
system(bash_i)
bash_i = 'chmod 777 liftOver'
system(bash_i)
bash_i = './liftOver -minMatch=0.95 Gasperini.Enh.Gene.pairs.hg19.bed hg19ToHg38.over.chain.gz Gasperini.Enh.Gene.pairs.hg19Tohg38.bed Gasperini.Enh.Gene.pairs.hg19Tohg38.unmapped.bed'
system(bash_i)

###
K562_CRISPRi = read.table('Gasperini.Enh.Gene.pairs.hg19Tohg38.bed', header=F)
#K562_CRISPRi = K562_CRISPRi[K562_CRISPRi[,5],]
###
hg38_SFN_loops_crispr_label = rep(-1, dim(cCRE_gene_JmetID_FS01_geneName_MID_HID_S01_TSS_GeneKM_mat_hg38_RNAcor_SFN_mat)[1])
for (i in 1:dim(K562_CRISPRi)[1]){
  if (i%%100==0) {print(i)}
intersect_Enh_gene_rows = ((cCRE_gene_JmetID_FS01_geneName_MID_HID_S01_TSS_GeneKM_mat_hg38_RNAcor_SFN_mat[,1]==K562_CRISPRi[i,1]) & (cCRE_gene_JmetID_FS01_geneName_MID_HID_S01_TSS_GeneKM_mat_hg38_RNAcor_SFN_mat[,6]==K562_CRISPRi[i,4]) & (cCRE_gene_JmetID_FS01_geneName_MID_HID_S01_TSS_GeneKM_mat_hg38_RNAcor_SFN_mat[,2]<=K562_CRISPRi[i,3]) & (cCRE_gene_JmetID_FS01_geneName_MID_HID_S01_TSS_GeneKM_mat_hg38_RNAcor_SFN_mat[,3]>=K562_CRISPRi[i,2]) )
#intersect_Enh_gene_rows = ((cCRE_gene_JmetID_FS01_geneName_MID_HID_S01_TSS_GeneKM_mat_hg38_RNAcor_SFN_mat[,1]==K562_CRISPRi[i,1]) & (cCRE_gene_JmetID_FS01_geneName_MID_HID_S01_TSS_GeneKM_mat_hg38_RNAcor_SFN_mat[,2]<=K562_CRISPRi[i,3]) & (cCRE_gene_JmetID_FS01_geneName_MID_HID_S01_TSS_GeneKM_mat_hg38_RNAcor_SFN_mat[,3]>=K562_CRISPRi[i,2]) )
if (sum(intersect_Enh_gene_rows)>0){
  hg38_SFN_loops_crispr_label[intersect_Enh_gene_rows] = K562_CRISPRi[i,5]
}
}

hg38_SFN_loops_with_crispri_label = cbind(cCRE_gene_JmetID_FS01_geneName_MID_HID_S01_TSS_GeneKM_mat_hg38_RNAcor_SFN_mat, hg38_SFN_loops_crispr_label)[hg38_SFN_loops_crispr_label!=-1,]

sum(hg38_SFN_loops_with_crispri_label[,14]==1) / sum(hg38_SFN_loops_with_crispri_label[,14]!=-1)

used_type = hg38_SFN_loops_with_crispri_label[,13]=='P'
sum(hg38_SFN_loops_with_crispri_label[used_type,14]==1) / sum(hg38_SFN_loops_with_crispri_label[,14]==1)

used_type = hg38_SFN_loops_with_crispri_label[,13]=='SF'
sum(hg38_SFN_loops_with_crispri_label[used_type,14]==1) / sum(hg38_SFN_loops_with_crispri_label[,14]==1)

used_type = hg38_SFN_loops_with_crispri_label[,13]=='F'
sum(hg38_SFN_loops_with_crispri_label[used_type,14]==1) / sum(hg38_SFN_loops_with_crispri_label[,14]==1)

used_type = hg38_SFN_loops_with_crispri_label[,13]=='S'
sum(hg38_SFN_loops_with_crispri_label[used_type,14]==1) / sum(hg38_SFN_loops_with_crispri_label[,14]==1)

used_type = hg38_SFN_loops_with_crispri_label[,13]=='N'
sum(hg38_SFN_loops_with_crispri_label[used_type,14]==1) / sum(hg38_SFN_loops_with_crispri_label[,14]==1)


table(hg38_SFN_loops_with_crispri_label[,13])


library(PRROC)
roc <- roc.curve(scores.class0 = hg38_SFN_loops_with_crispri_label[hg38_SFN_loops_with_crispri_label[,14]==1,12], scores.class1 = hg38_SFN_loops_with_crispri_label[hg38_SFN_loops_with_crispri_label[,14]==0,12], curve = T)
pdf('ROC.crispri.pdf')
plot(roc)
dev.off()


hg38_SFN_loops_with_crispri_label_SF = hg38_SFN_loops_with_crispri_label
hg38_SFN_loops_with_crispri_label_SF[hg38_SFN_loops_with_crispri_label[,7]=='SF',12] = hg38_SFN_loops_with_crispri_label_SF[hg38_SFN_loops_with_crispri_label[,7]=='SF',12]+10
pr <- roc.curve(scores.class0 = hg38_SFN_loops_with_crispri_label_SF[hg38_SFN_loops_with_crispri_label_SF[,19]==1,12], scores.class1 = hg38_SFN_loops_with_crispri_label_SF[hg38_SFN_loops_with_crispri_label[,19]==0,12], curve = T)
pdf('ROC.crispri.SF.pdf')
plot(pr)
dev.off()


# PR Curve
pr <- pr.curve(scores.class0 = hg38_SFN_loops_with_crispri_label[hg38_SFN_loops_with_crispri_label[,14]==1,12], scores.class1 = hg38_SFN_loops_with_crispri_label[hg38_SFN_loops_with_crispri_label[,14]==0,12], curve = T)
pdf('PRC.crispri.pdf')
plot(pr)
dev.off()


hg38_SFN_loops_with_crispri_label_SF = hg38_SFN_loops_with_crispri_label[is.element(hg38_SFN_loops_with_crispri_label[,7], c('SF','F')),]
pr <- pr.curve(scores.class0 = hg38_SFN_loops_with_crispri_label_SF[hg38_SFN_loops_with_crispri_label_SF[,14]==1,6], scores.class1 = hg38_SFN_loops_with_crispri_label_SF[hg38_SFN_loops_with_crispri_label_SF[,14]==0,6], curve = T)
pdf('PRC.crispri.SF.pdf')
plot(pr)
dev.off()


#################################################
### write cCRE-cCRE loop file
### add TSS position
hg38_SFN_CC_loops_new = hg38_SFN_loops[hg38_SFN_loops[,1]!='chrXasdasd',]
### enriched F set in each gene
hg38_SFN_set_per_gene = apply(hg38_SFN_loops[hg38_SFN_loops[,1]!='chrXasdasd',c(4,17)], 1, function(x) paste(x[1], x[2], sep='_'))
hg38_SFN_CC_loops_Jmet_gene_new = c()
k = 0
for (Jmet_gene_i in unique(hg38_SFN_set_per_gene)){
  k = k+1
  if (k%%100==0){print(k)}
  hg38_SFN_CC_loops_Jmet_gene_i = hg38_SFN_CC_loops_new[hg38_SFN_set_per_gene==Jmet_gene_i,]
  if (hg38_SFN_CC_loops_Jmet_gene_i[1,5]!=0){
    hg38_SFN_CC_loops_Jmet_gene_i_new = c()
    for (i in 1:dim(hg38_SFN_CC_loops_Jmet_gene_i)[1]){
      for (j in 1:dim(hg38_SFN_CC_loops_Jmet_gene_i)[1]){
        if (i<j){
          hg38_SFN_CC_loops_Jmet_gene_i_new_j = hg38_SFN_CC_loops_Jmet_gene_i[i,]
          hg38_SFN_CC_loops_Jmet_gene_i_new_j[14:18] = hg38_SFN_CC_loops_Jmet_gene_i[j,c(9:13)]
          hg38_SFN_CC_loops_Jmet_gene_i_new = rbind(hg38_SFN_CC_loops_Jmet_gene_i_new, hg38_SFN_CC_loops_Jmet_gene_i_new_j)
        }
      }
    }
    hg38_SFN_CC_loops_Jmet_gene_new = rbind(hg38_SFN_CC_loops_Jmet_gene_new, hg38_SFN_CC_loops_Jmet_gene_i_new)
  }
}

### change colors
hg38_SFN_CC_loops_Jmet_gene_new_unique[hg38_SFN_CC_loops_Jmet_gene_new_unique[,7]=='SF',8] = '#FD0606'
hg38_SFN_CC_loops_Jmet_gene_new_unique[hg38_SFN_CC_loops_Jmet_gene_new_unique[,7]=='F',8] = '#FFAB00'
hg38_SFN_CC_loops_Jmet_gene_new_unique[hg38_SFN_CC_loops_Jmet_gene_new_unique[,7]=='S',8] = '#000000'
hg38_SFN_CC_loops_Jmet_gene_new_unique[hg38_SFN_CC_loops_Jmet_gene_new_unique[,7]=='N',8] = '#DEDEDE'
### 
hg38_SFN_CC_loops_Jmet_gene_new_unique = unique(hg38_SFN_CC_loops_Jmet_gene_new)
options("scipen"=100, "digits"=4)
write.table(hg38_SFN_CC_loops_Jmet_gene_new_unique[hg38_SFN_CC_loops_Jmet_gene_new_unique[,6]>0.2,], 'hg38.SFN.loop.CC.interact', quote=F, sep='\t', col.names=F, row.names=F)
bash1 = 'cat loop.header.txt > hg38.SFN.loop.CC.interact.tmp'
bash2 = 'cat hg38.SFN.loop.CC.interact >> hg38.SFN.loop.CC.interact.tmp && mv hg38.SFN.loop.CC.interact.tmp hg38.SFN.loop.CC.interact'
system(bash1)
system(bash2)


hg38_SFN_loops = cbind(hg38_SFN_loops[,1:5], round(hg38_SFN_loops[12]*1000), hg38_SFN_loops[,13], rep('#DEDEDE', dim(hg38_SFN_loops)[1]), hg38_SFN_loops[,c(1:3,7)], rep('.', dim(hg38_SFN_loops)[1]), hg38_SFN_loops_TSS, hg38_SFN_loops$GeneName, rep('.', dim(hg38_SFN_loops)[1]) )
### remove neg
hg38_SFN_loops[(hg38_SFN_loops[,6]<0) | (is.na(hg38_SFN_loops[,6])),6] = 0
### change color
hg38_SFN_loops[hg38_SFN_loops[,7]=='SF',8] = '#FD0606'
hg38_SFN_loops[hg38_SFN_loops[,7]=='F',8] = '#FFAB00'
hg38_SFN_loops[hg38_SFN_loops[,7]=='S',8] = '#000000'
hg38_SFN_loops[hg38_SFN_loops[,7]=='N',8] = '#DEDEDE'
hg38_SFN_loops[hg38_SFN_loops[,7]=='P',8] = '#FF0000'
hg38_SFN_loops[hg38_SFN_loops[,17]=='GATA1',]
hg38_gene_shared_exp[hg38_gene_shared_exp[,5]=='GATA1',]
hg38_SFN_loops[hg38_SFN_loops[,17]=='CSF1R',]
hg38_gene_shared_exp[hg38_gene_shared_exp[,5]=='CSF1R',]
options("scipen"=100, "digits"=4)
write.table(hg38_SFN_loops[hg38_SFN_loops[,6]>0.2,], 'hg38.SFN.loop.interact', quote=F, sep='\t', col.names=F, row.names=F)
bash1 = 'cat loop.header.txt > hg38.SFN.loop.interact.tmp'
bash2 = 'cat hg38.SFN.loop.interact >> hg38.SFN.loop.interact.tmp && mv hg38.SFN.loop.interact.tmp hg38.SFN.loop.interact'
system(bash1)
system(bash2)
#################################################


### mm10
cCRE_M_Jmeta1 = cbind(cCRE_M_Jmeta, cCRE_M_Jmeta[,4])
for (gene_i in unique(cCRE_M_Jmeta[,5])){
cCRE_M_Jmeta1[cCRE_M_Jmeta[,5]==gene_i,6] = meta_cluster_mat_log2_km$cluster[shared_genes==gene_i]
}
colnames(cCRE_M_Jmeta1)[4] = 'JmetID'
colnames(cCRE_M_Jmeta1)[6] = 'Gene_Jmet_KMID'
write.table(cCRE_M_Jmeta1, 'cCRE.Gene50KB.All_Info.mm10.bed', quote=F, sep='\t', col.names=T, row.names=F)


cCRE_H_Jmeta1[cCRE_H_Jmeta1[,5]=='GATA1',]
cCRE_M_Jmeta1[cCRE_M_Jmeta1[,5]=='GATA1',]

hg38_gene_shared_exp[hg38_gene_shared_exp[,5]=='GATA1',]
mm10_gene_shared_exp[mm10_gene_shared_exp[,5]=='GATA1',]

i = which(shared_genes == 'GATA1')
bed_H = hg38_gene_shared_exp[i,1:3][1,]
bed_M = mm10_gene_shared_exp[i,1:3][1,]
JMeta_i = get_Function_conserve_cCRE(bed_H[1,1], bed_H[1,2], bed_H[1,3], bed_M[1,1], bed_M[1,2], bed_M[1,3])

### check Jmet_GeneKM count in H & M
pdf('Jmet.GeneKM.count.pdf')
plot_lim = c(1,10000)
Human_count = as.numeric(table(apply(cCRE_H_Jmeta1,1, function(x) paste(x[4],x[6]))))
Mouse_count = as.numeric(table(apply(cCRE_M_Jmeta1,1, function(x) paste(x[4],x[6])))) 
Mouse_count_adj = Mouse_count / mean(Mouse_count) * mean(Human_count)
plot(Human_count,  Mouse_count_adj, xlim=plot_lim, ylim=plot_lim, log='xy', cex.axis=2, pch=16)
abline(0,1)
dev.off()
#################################################


#################################################
### check
module_per_gene = apply(cCRE_H_Jmeta, 1, function(x) paste(x[4], x[5], sep=':'))
head(table(module_per_gene)[order(-table(module_per_gene))])

meta_cluster_mat_log2_df[meta_cluster_mat_log2_df[,1]=='FIBCD1',]
meta_cluster_mat_log2_df[meta_cluster_mat_log2_df[,1]=='GATA1',]
meta_cluster_mat_log2_df[meta_cluster_mat_log2_df[,1]=='CD8A',]
meta_cluster_mat_log2_df[meta_cluster_mat_log2_df[,1]=='CSF1R',]

GATA1_exp = get_Function_conserve_cCRE('chrX', 48760001, 48836000, 'chrX', 7919401,8020800)
CD8A_exp = get_Function_conserve_cCRE('chr2', 86758396, 86858396, 'chr6', 71323427,71423427,JointCluster_count_HM)
CSF1R_exp = get_Function_conserve_cCRE('chr5', 150063372, 150163372, 'chr18', 61050598,61150598,JointCluster_count_HM)
IFNG_exp = get_Function_conserve_cCRE('chr12', 68109740, 68209740, 'chr10', 118391046,118491046,JointCluster_count_HM)


length(table(apply(cCRE_H_Jmeta1,1, function(x) paste(x[4],x[6]))))
length(table(apply(cCRE_M_Jmeta1,1, function(x) paste(x[4],x[6]))))


### check EP-pairs num
EP_pair_num = rep(0, dim(hg38_gene_shared_exp)[1])
for (i in 1:dim(hg38_gene_shared_exp)[1]){
  if (i%%1000==0){print(i)}
hg38_gene_shared_exp_i = hg38_gene_shared_exp[i,]
included_rows_i = ((dh_with_JointClusterID[,1]==hg38_gene_shared_exp_i[1,1]) * (dh_with_JointClusterID[,2]>=hg38_gene_shared_exp_i[1,2]) * (dh_with_JointClusterID[,3]<=hg38_gene_shared_exp_i[1,3])) != 0
EP_pair_num[i] = sum(included_rows_i)
}

#> summary(EP_pair_num)
#   Min. 1st Qu.  Median    Mean 3rd Qu.    Max. 
#   0.00    9.00   16.00   17.88   24.00   97.00 
#> sum(EP_pair_num)
#[1] 286027


bedtools intersect -a cCRE.Gene50KB.hg38.bed -b human_ccre_hg38_sf.bed  -c > cCRE.Gene50KB.hg38.SF01.bed
bedtools intersect -a cCRE.Gene50KB.mm10.bed -b mouse_ccre_mm10_sf.bed  -c > cCRE.Gene50KB.mm10.SF01.bed

input_file_hg38 = 'cCRE.Gene50KB.hg38.SF01.bed'
input_file_mm10 = 'cCRE.Gene50KB.mm10.SF01.bed'
output_file = 'Jmet.SF.p.pdf'

dd_hg38 = read.table(input_file_hg38, header=F)
dd_mm10 = read.table(input_file_mm10, header=F)
dd = rbind(dd_hg38, dd_mm10)
### check gene group SF count
Gene_SF_count = c()
for (gene_i in unique(dd[,5])){
Gene_SF_count = rbind(Gene_SF_count, c(sum(dd[,5]==gene_i), sum(dd[dd[,5]==gene_i,7])))
}
Gene_SF_count = cbind(as.data.frame(unique(dd[,5])), Gene_SF_count)
colnames(Gene_SF_count) = c('gene', 'F_cCREn', 'SF_cCREn')

### check Jmet SF proportion
Jmet_SF = c()
for (i in unique(dd[,4])){
Jmet_SF = rbind(Jmet_SF, c(sum(dd[dd[,4]==i,7]!=0), sum(dd[,4]==i)))
}
Jmet_SF = cbind(unique(dd[,4]), Jmet_SF)
Jmet_SF = cbind(Jmet_SF,Jmet_SF[,2]/Jmet_SF[,3])
colnames(Jmet_SF) = c('Jmet','SF_nCREn','F_nCREn','ratio')
Jmet_SF_plot = c()
used_order = c(1,11,5,8,2,6,9,10)
for (Jmeti in used_order){
  Jmet_SF_plot = rbind(Jmet_SF_plot, Jmet_SF[Jmet_SF[,1]==Jmeti,4])
}
rownames(Jmet_SF_plot) = used_order
colnames(Jmet_SF_plot) = NULL

pdf(output_file, width=3)
pheatmap(cbind(Jmet_SF_plot,Jmet_SF_plot), cluster_rows=F, cex=1.5, )
dev.off()

#################################################


chr     start   end     cCRE    AVE     B_B15_50        B_NC14_42 CD34_E_rep1     CD34_E_rep2     CLP_100266      CLP_100267      CMP_100246 CMP_100247      EOS_S006XEH2    EOS_S00BKK      ERY_S002R5      ERY_S002S3
GMP_100256      GMP_100257      HSC_100258      HSC_100259
HUDEP1_rep1     HUDEP1_rep2     HUDEP2_rep1     HUDEP2_rep2     K562_rep1
K562_rep2       LMPP_100268     LMPP_100269     MEP_Donor2596
MEP_Donor7256   MK_S004BTH2     MK_S00VHKH1     MONc_C0011IH1
MONc_C001UYH2   MONp_Prim_mon_C MONp_Prim_mon_F MPP_100272      MPP_100273
NEU_C0011IH2    NEU_C001UYH1    NK_S005YG       NK_S01E4WH0
T_CD4_S008H1    T_CD4_S009W4    T_CD8_C0066PH1  T_CD8_S00C2FH1
chr2    60550400        60552800        chr2_60550400_60552800  12      14
14      14      20      2       2       2       14      18      18      14
14      2       12      2       14      15      15      15      15      2
2       12      12      2       12      12      14      14      14      14
14      12      12      12      18      18      18      18      18      12


bedtools5 = 'cat cCRE.Gene50KB.hg38.SF_S.bed | awk \'{if ($6=="GATA1") print $0}\''
bedtools6 = 'cat cCRE.Gene50KB.mm10.withMID.bed | awk \'{if ($6=="GATA1") print $0}\''
system(bedtools5)
system(bedtools6)
gene_RNA_Jmet_cor[rownames(gene_RNA_Jmet_cor)=='GATA1',]
hg38_gene_shared_exp[hg38_gene_shared_exp[,5]=='GATA1',]
mm10_gene_shared_exp[mm10_gene_shared_exp[,5]=='GATA1',]


bedtools5 = 'cat cCRE.Gene50KB.hg38.SF_S.bed | awk \'{if ($6=="GLOD5") print $0}\''
bedtools6 = 'cat cCRE.Gene50KB.mm10.withMID.bed | awk \'{if ($6=="GLOD5") print $0}\''
system(bedtools5)
system(bedtools6)
gene_RNA_Jmet_cor[rownames(gene_RNA_Jmet_cor)=='GLOD5',]
hg38_gene_shared_exp[hg38_gene_shared_exp[,5]=='GLOD5',]
mm10_gene_shared_exp[mm10_gene_shared_exp[,5]=='GLOD5',]

bedtools5 = 'cat cCRE.Gene50KB.hg38.SF_S.bed | awk \'{if ($6=="CSF1R") print $0}\''
bedtools6 = 'cat cCRE.Gene50KB.mm10.withMID.bed | awk \'{if ($6=="CSF1R") print $0}\''
bedtools5 = 'cat cCRE.Gene50KB.hg38.SF_S.bed | awk \'{if ($6=="CSF1R" && ($5==1 || $7!="NA")) print $0}\''
bedtools6 = 'cat cCRE.Gene50KB.mm10.withMID.bed | awk \'{if ($6=="CSF1R" && ($5==1 || $7!="NA")) print $0}\''
system(bedtools5)
system(bedtools6)
gene_RNA_Jmet_cor[rownames(gene_RNA_Jmet_cor)=='CSF1R',]
hg38_gene_shared_exp[hg38_gene_shared_exp[,5]=='CSF1R',]
mm10_gene_shared_exp[mm10_gene_shared_exp[,5]=='CSF1R',]


###
Gene_Jmet_RNA_shared_genes = rownames(meta_cluster_mat)[is.element(rownames(meta_cluster_mat), rownames(gene_RNA_Jmet_cor))]
meta_cluster_mat_check_shared = meta_cluster_mat[is.element(rownames(meta_cluster_mat), Gene_Jmet_RNA_shared_genes),]
gene_RNA_Jmet_cor_check_shared = gene_RNA_Jmet_cor[is.element(rownames(gene_RNA_Jmet_cor), Gene_Jmet_RNA_shared_genes),]
### remove uniq
gene_RNA_Jmet_cor_check_shared_unique = c()
k = 0
for (gene_i in rownames(meta_cluster_mat_check_shared)){
  k =k+1
  if (k%%10000==0){print(k)}
  if (sum(rownames(gene_RNA_Jmet_cor_check_shared)==gene_i)>1){
    gene_RNA_Jmet_cor_check_shared_unique = rbind(gene_RNA_Jmet_cor_check_shared_unique, colMeans(gene_RNA_Jmet_cor_check_shared[rownames(gene_RNA_Jmet_cor_check_shared)==gene_i,]))
  } else{
    gene_RNA_Jmet_cor_check_shared_unique = rbind(gene_RNA_Jmet_cor_check_shared_unique, gene_RNA_Jmet_cor_check_shared[rownames(gene_RNA_Jmet_cor_check_shared)==gene_i,])

  }
}
rownames(gene_RNA_Jmet_cor_check_shared_unique) = rownames(meta_cluster_mat_check_shared)
colnames(gene_RNA_Jmet_cor_check_shared_unique) = colnames(gene_RNA_Jmet_cor_check_shared)


gene_RNA_Jmet_cor_check_shared_unique = gene_RNA_Jmet_cor_check_shared_unique[order(rownames(gene_RNA_Jmet_cor_check_shared_unique)),order(colnames(gene_RNA_Jmet_cor_check_shared_unique))]
meta_cluster_mat_check_shared = meta_cluster_mat_check_shared[order(rownames(meta_cluster_mat_check_shared)),order(colnames(meta_cluster_mat_check_shared))]

gene_RNA_Jmet_cor_check_shared_unique_vec = as.numeric(as.matrix(gene_RNA_Jmet_cor_check_shared_unique))
meta_cluster_mat_check_shared_vec = as.numeric(as.matrix(meta_cluster_mat_check_shared))


png('gene_RNA_Jmet_cor.png')
used_id = sample(length(gene_RNA_Jmet_cor_check_shared_unique_vec), 10000)
heatscatter(gene_RNA_Jmet_cor_check_shared_unique_vec[used_id], meta_cluster_mat_check_shared_vec[used_id], log='y')
dev.off()


cat cCRE.Gene50KB.hg38.JmetID.FS01.geneName.HID.MID.S01.TSS.GeneKMID.SFN+ID.bed | awk -F '\t' -v OFS='\t' '{if ($12=="SF+" && $4==1) print $1,$2,$3,$4}' | sort -u | sort -k1,1 -k2,2n > cCRE.Gene50KB.hg38.JmetID.FS01.geneName.HID.MID.S01.TSS.GeneKMID.SFN+ID.SF+.bed
cat cCRE.Gene50KB.hg38.JmetID.FS01.geneName.HID.MID.S01.TSS.GeneKMID.SFN+ID.bed | awk -F '\t' -v OFS='\t' '{if ($12=="SF" && $4==1) print $1,$2,$3,$4}' | sort -u | sort -k1,1 -k2,2n > cCRE.Gene50KB.hg38.JmetID.FS01.geneName.HID.MID.S01.TSS.GeneKMID.SFN+ID.SF.bed
cat cCRE.Gene50KB.hg38.JmetID.FS01.geneName.HID.MID.S01.TSS.GeneKMID.SFN+ID.bed | awk -F '\t' -v OFS='\t' '{if ($12=="S" && $4==1) print $1,$2,$3,$4}' | sort -u | sort -k1,1 -k2,2n > cCRE.Gene50KB.hg38.JmetID.FS01.geneName.HID.MID.S01.TSS.GeneKMID.SFN+ID.S.bed
cat cCRE.Gene50KB.hg38.JmetID.FS01.geneName.HID.MID.S01.TSS.GeneKMID.SFN+ID.bed | awk -F '\t' -v OFS='\t' '{if ($12=="S+" && $4==1) print $1,$2,$3,$4}' | sort -u | sort -k1,1 -k2,2n > cCRE.Gene50KB.hg38.JmetID.FS01.geneName.HID.MID.S01.TSS.GeneKMID.SFN+ID.S+.bed
cat cCRE.Gene50KB.hg38.JmetID.FS01.geneName.HID.MID.S01.TSS.GeneKMID.SFN+ID.bed | awk -F '\t' -v OFS='\t' '{if ($12=="J" && $4==1) print $1,$2,$3,$4}' | sort -u | sort -k1,1 -k2,2n > cCRE.Gene50KB.hg38.JmetID.FS01.geneName.HID.MID.S01.TSS.GeneKMID.SFN+ID.J.bed
cat cCRE.Gene50KB.hg38.JmetID.FS01.geneName.HID.MID.S01.TSS.GeneKMID.SFN+ID.bed | awk -F '\t' -v OFS='\t' '{if ($12=="N" && $4==1) print $1,$2,$3,$4}' | sort -u | sort -k1,1 -k2,2n > cCRE.Gene50KB.hg38.JmetID.FS01.geneName.HID.MID.S01.TSS.GeneKMID.SFN+ID.N.bed

cat hg38.cCRE.SFN\(+\).colored.bed | awk -F ':' -v OFS='\t' '{print $1}' | awk -F '\t' -v OFS='\t' '{if ($4=="SFJ") print $0}' > cCRE.Gene50KB.hg38.JmetID.FS01.geneName.HID.MID.S01.TSS.GeneKMID.SFN+ID.SF+.bed
cat hg38.cCRE.SFN\(+\).colored.bed | awk -F ':' -v OFS='\t' '{print $1}' | awk -F '\t' -v OFS='\t' '{if ($4=="SF") print $0}' > cCRE.Gene50KB.hg38.JmetID.FS01.geneName.HID.MID.S01.TSS.GeneKMID.SFN+ID.SF.bed
cat hg38.cCRE.SFN\(+\).colored.bed | awk -F ':' -v OFS='\t' '{print $1}' | awk -F '\t' -v OFS='\t' '{if ($4=="SJ") print $0}' > cCRE.Gene50KB.hg38.JmetID.FS01.geneName.HID.MID.S01.TSS.GeneKMID.SFN+ID.S+.bed
cat hg38.cCRE.SFN\(+\).colored.bed | awk -F ':' -v OFS='\t' '{print $1}' | awk -F '\t' -v OFS='\t' '{if ($4=="S") print $0}' > cCRE.Gene50KB.hg38.JmetID.FS01.geneName.HID.MID.S01.TSS.GeneKMID.SFN+ID.S.bed
cat hg38.cCRE.SFN\(+\).colored.bed | awk -F ':' -v OFS='\t' '{print $1}' | awk -F '\t' -v OFS='\t' '{if ($4=="J") print $0}' > cCRE.Gene50KB.hg38.JmetID.FS01.geneName.HID.MID.S01.TSS.GeneKMID.SFN+ID.J.bed
cat hg38.cCRE.SFN\(+\).colored.bed | awk -F ':' -v OFS='\t' '{print $1}' | awk -F '\t' -v OFS='\t' '{if ($4=="N") print $0}' > cCRE.Gene50KB.hg38.JmetID.FS01.geneName.HID.MID.S01.TSS.GeneKMID.SFN+ID.N.bed


ref_bed='human_ccre_hg38_LoopAnchor_K5562_GM12878.bed'
ref_bed='human_ccre_hg38_EP300_3ct.bed'
ref_bed='human_ccre_hg38_GENCODEtss.bed'

ref_bed_sort=$ref_bed'.sort.bed'
sort -k1,1 -k2,2n /Users/guanjuexiang/Downloads/$ref_bed > /Users/guanjuexiang/Downloads/$ref_bed'.sort.bed'

bedtools intersect -a cCRE.Gene50KB.hg38.JmetID.FS01.geneName.HID.MID.S01.TSS.GeneKMID.SFN+ID.SF+.bed -b /Users/guanjuexiang/Downloads/$ref_bed_sort -wa -u > Intersect.SF+.txt
bedtools intersect -a cCRE.Gene50KB.hg38.JmetID.FS01.geneName.HID.MID.S01.TSS.GeneKMID.SFN+ID.SF.bed -b /Users/guanjuexiang/Downloads/$ref_bed_sort -wa -u > Intersect.SF.txt
bedtools intersect -a cCRE.Gene50KB.hg38.JmetID.FS01.geneName.HID.MID.S01.TSS.GeneKMID.SFN+ID.S.bed -b /Users/guanjuexiang/Downloads/$ref_bed_sort -wa -u > Intersect.S.txt
bedtools intersect -a cCRE.Gene50KB.hg38.JmetID.FS01.geneName.HID.MID.S01.TSS.GeneKMID.SFN+ID.S+.bed -b /Users/guanjuexiang/Downloads/$ref_bed_sort -wa -u > Intersect.S+.txt
bedtools intersect -a cCRE.Gene50KB.hg38.JmetID.FS01.geneName.HID.MID.S01.TSS.GeneKMID.SFN+ID.J.bed -b /Users/guanjuexiang/Downloads/$ref_bed_sort -wa -u > Intersect.J.txt
bedtools intersect -a cCRE.Gene50KB.hg38.JmetID.FS01.geneName.HID.MID.S01.TSS.GeneKMID.SFN+ID.N.bed -b /Users/guanjuexiang/Downloads/$ref_bed_sort -wa -u > Intersect.N.txt

wc -l Intersect.*.txt

wc -l cCRE.Gene50KB.hg38.JmetID.FS01.geneName.HID.MID.S01.TSS.GeneKMID.SFN+ID.*.bed

d1 = read.table('JI.SF+.txt', header=T)
d2 = read.table('JI.SF.txt', header=T)
d3 = read.table('JI.S+.txt', header=T)
d4 = read.table('JI.S.txt', header=T)
d5 = read.table('JI.J.txt', header=T)
d6 = read.table('JI.N.txt', header=T)
```
