## Supplementary material for "Interspecies regulatory landscapes and elements revealed by novel joint systematic integration of human and mouse blood cell epigenomes": Scripts for IDEAS modeling of epigenetic states jointly in human and mouse: reviewer1_point5_6_get_cCRE.get_coe_umap.human.R

```
library(pheatmap)
library(umap)
library(LSD)
library(lisi)


d = read.table('S3V2_IDEAS_hg38_ccre2.cCRE.M.notall0.rmallNEU.withid.coe_mat.txt', header=T)

ds = d[,-c(1:4)]

library(pheatmap)
library(umap)
library(LSD)

HsP_cCRE = read.table('../coe_analysis/S3V2_IDEAS_hg38_ccre2.cCRE.M.withid.atProximal.bed', header=F)

coe_mat = read.table('../coe_analysis/statep_rna_coe_heatmap.HM.all.ccre.withcorfilter.AVE.txt', header=T)
scale_ratio = max(coe_mat[,2])/max(coe_mat[,1])

### get
dsP = as.matrix(ds[,1:(dim(ds)[2]/2)])
dsD = as.matrix(ds[,(dim(ds)[2]/2+1):dim(ds)[2]])
#dsPadj = (dsP - mean(dsP))/sd(dsP) * sd(dsD) + mean(dsD)

dsPadj = dsP * scale_ratio

#dsMerge = dsD
#dsMerge[HsP_cCRE[,5]!=0,] = dsPadj[HsP_cCRE[,5]!=0,]/2 + dsD[HsP_cCRE[,5]!=0,]/2
dsMerge = dsD + dsPadj
dsMerge_output = cbind(d[,1:4], dsMerge)
colnames(dsMerge_output)[1:4] = c('chr','start','end','id')
dsMerge_output = dsMerge_output[,!is.element(colnames(dsMerge_output), c('NEU_C0011IH2_D', 'NEU_C001UYH1_D'))]
#write.table(dsMerge_output, '../coe_analysis/S3V2_IDEAS_hg38_ccre2.cCRE.M.notall0.rmallNEU.withid.coe_mat.PDmerged.txt', quote=F, col.names=T, row.names=F, sep='\t')


###### For Proximal & Distal
set.seed(2019)
used_row = sample(dim(ds)[1], 20000)
dssP = as.matrix(ds[used_row,1:(dim(ds)[2]/2)])
dssD = as.matrix(ds[used_row,(dim(ds)[2]/2+1):dim(ds)[2]])
#dssPadj = (dssP - mean(dssP))/sd(dssP) * sd(dssD) + mean(dssD)
dssPadj = dssP * scale_ratio
dssDadj = (dssD - mean(dssD))/sd(dssD) * sd(dssP) + mean(dssP)

rep1 = c(1, 2, 4, 6, 8, 10, 12, 14, 16, 18, 20, 22, 24, 26, 28, 30, 32, 34, 36, 38, 40, 42)
rep2 = c(1, 3, 5, 7, 9, 11, 13, 15, 17, 19, 21, 23, 25, 27, 29, 31, 33, 35, 37, 39, 41, 43)

dssP_ct = c()
dssD_ct = c()
dssPadj_ct = c()
dssDadj_ct = c()
for (i in 1:length(rep1)){
	dssP_ct = cbind(dssP_ct, rowMeans(cbind(dssP[,rep1[i]], dssP[,rep2[i]])))
	dssD_ct = cbind(dssD_ct, rowMeans(cbind(dssD[,rep1[i]], dssD[,rep2[i]])))
	dssPadj_ct = cbind(dssPadj_ct, rowMeans(cbind(dssPadj[,rep1[i]], dssPadj[,rep2[i]])))
	dssDadj_ct = cbind(dssDadj_ct, rowMeans(cbind(dssDadj[,rep1[i]], dssDadj[,rep2[i]])))
}

### get subsample umap
HsP_cCRE_s = HsP_cCRE[used_row,]

#dss = dssD
#dss[HsP_cCRE_s[,5]!=0,] = dssPadj[HsP_cCRE_s[,5]!=0,]/2 + dssD[HsP_cCRE_s[,5]!=0,]/2 #( dssP+dssDadj)[HsP_cCRE_s[,5]!=0,]/2
dss = dssD + dssPadj

#dssPD_ct = dssD_ct
#dssPD_ct[HsP_cCRE_s[,5]!=0,] = dssP_ct[HsP_cCRE_s[,5]!=0,] + dssDadj_ct[HsP_cCRE_s[,5]!=0,]

#dssPD_ct = dssD_ct
#dssPD_ct[HsP_cCRE_s[,5]!=0,] = dssPadj_ct[HsP_cCRE_s[,5]!=0,]/2 + dssD_ct[HsP_cCRE_s[,5]!=0,]/2
dssPD_ct = dssD_ct + dssPadj_ct

colnames(dssPD_ct) = colnames(dss)[rep1]

#dssPD_ct = cbind(dssPadj, dssD_ct)
#dssPD_ct = cbind(dssP_ct, dssDadj_ct)
#dssPD_ct = cbind(dssD_ct)
#dssPD_ct = (dssP_ct+dssDadj_ct)/2

### run umap
set.seed(2019)
dss_umap = umap(round(dssPD_ct[, colnames(dssPD_ct)!='NEU_C0011IH2_D'], 5), preserve.seed=T)

dir.create('coe_umap_PD_esRP')
### plot heatscatter plot
png('coe_umap_PD_esRP/state_coe.human.umap.PD.0702.png')
heatscatter(dss_umap$layout[,1], dss_umap$layout[,2], cex=0.5, xlab='UMAP Dim1', ylab='UMAP Dim2')
dev.off()

# read cCRE annotation label
cCRE_label = read.table('S3V2_IDEAS_hg38_ccre2.cCRE.M.notall0.withid.label.bed', header=F)
# get cCRE label sample
cCRE_label_sample = cCRE_label[used_row,]

# plot umap colored by cCRE label
table(cCRE_label_sample[,4])
uniq_cCRE_label = names(table(cCRE_label_sample[,4]))
color_list = c('orange','red','green')
pdf('coe_umap_PD_esRP/state_coe.human.umap.PD.0702.cCRElabel.pdf')
plot(dss_umap$layout[,1], dss_umap$layout[,2], col='gray', pch=16, main = '', xlab='UMAP Dim1', ylab='UMAP Dim2', cex = 0.5)
#for (i in 1:length(uniq_cCRE_label)){
for (i in c(1,2,3)){
	plot_1 = cCRE_label_sample[,4] == uniq_cCRE_label[i]
	points(dss_umap$layout[plot_1,1], dss_umap$layout[plot_1,2], col=color_list[i], pch=16, cex = 0.5)
}
dev.off()

# lisi score
cCRE_label_sample_1 = cbind(cCRE_label_sample[,4])
colnames(cCRE_label_sample_1) = 'cCRE_label'
lisi_score = compute_lisi(dss_umap$layout, cCRE_label_sample_1, 'cCRE_label')

mean(lisi_score$cCRE_label)
mean(lisi_score$cCRE_label[cCRE_label_sample_1=='Intergenic'])
mean(lisi_score$cCRE_label[cCRE_label_sample_1=='Promoter'])
mean(lisi_score$cCRE_label[cCRE_label_sample_1=='TES'])

lisi_score_mat = c()
for (k in 1:30){
set.seed(k)
print(k)
used_row = sample(dim(ds)[1], 20000)
dssP = as.matrix(ds[used_row,1:(dim(ds)[2])])
# merege replicate
dssP_ct = c()
for (i in 1:length(rep1)){
	dssP_ct = cbind(dssP_ct, rowMeans(cbind(dssP[,rep1[i]], dssP[,rep2[i]])))
}
# define cell type names
colnames(dssP_ct) = ct_list_human
# umap
dss_umap = umap(dssP_ct[, colnames(dssP_ct)!='NEU_C0011IH2_D'], preserve.seed=T)
# get cCRE label sample
cCRE_label_sample = cCRE_label[used_row,]
# lisi score
cCRE_label_sample_1 = cbind(cCRE_label_sample[,4])
colnames(cCRE_label_sample_1) = 'cCRE_label'
lisi_score = compute_lisi(dss_umap$layout, cCRE_label_sample_1, 'cCRE_label')
lisi_score_mat = rbind(lisi_score_mat, c(mean(lisi_score$cCRE_label), mean(lisi_score$cCRE_label[cCRE_label_sample_1=='Intergenic']), mean(lisi_score$cCRE_label[cCRE_label_sample_1=='Promoter']), mean(lisi_score$cCRE_label[cCRE_label_sample_1=='TES'])))
print(lisi_score_mat)
write.table(lisi_score_mat, 'lisi_score_mat.Beta_coef.txt', sep='\t', quote=F, row.names=F, col.names=F)
}

write.table(lisi_score_mat, 'lisi_score_mat.Beta_coef.txt', sep='\t', quote=F, row.names=F, col.names=F)

# lisi
n = 30
lisi_score_mat_coe = read.table('lisi_score_mat.Beta_coef.txt', header=F)[1:n,]
lisi_score_mat_ATAC = read.table('lisi_score_mat.ATAC.txt', header=F)[1:n,]
lisi_score_mat_H3K27ac = read.table('lisi_score_mat.H3K27ac.txt', header=F)[1:n,]


pdf('lisi_score_mat.boxplot.pdf', width=6, height=6)
lisi_mat = c()
for (i in 1:4){
lisi_mat = cbind(lisi_mat, lisi_score_mat_coe[,i], lisi_score_mat_ATAC[,i], lisi_score_mat_H3K27ac[,i])
}
sig_type = c('Beta_coef', 'ATAC', 'H3K27ac')
colnames(lisi_mat) = c(paste0('All.', sig_type), paste0('Intergenic.', sig_type), paste0('Promoter.', sig_type), paste0('TES.', sig_type))
# boxplot with different color
boxplot(lisi_mat, cex.axis=1, las=3, col=c('red','blue','green','red','blue','green','red','blue','green','red','blue','green'), main='', ylab='LISI score', xlab='', ylim=c(min(lisi_mat), max(lisi_mat)))
dev.off()


for (i in 1:dim(dss)[2]){
	ct_i = colnames(dss)[i]
png(paste('coe_umap_PD_esRP/', 'state_coe.human.umap.PD.', ct_i,'.png', sep=''))
rbPal = colorRampPalette(c('blue', 'gray90','red'))
color_sig_range = (dss[,i])
print(summary(color_sig_range))
color_sig_range[color_sig_range>quantile(as.numeric(as.matrix(dss)),0.99)] = quantile(as.numeric(as.matrix(dss)),0.99)
plot_color = rbPal(500)[as.numeric(cut(c(-quantile(as.numeric(as.matrix(dss)),0.99), quantile(as.numeric(as.matrix(dss)),0.99), color_sig_range),breaks = 500))][-c(1:2)]
plot(dss_umap$layout[,1], dss_umap$layout[,2], col='white', pch=16, main = ct_i, xlab='UMAP Dim1', ylab='UMAP Dim2', cex = 0.5)
plot_1 = color_sig_range <= quantile(color_sig_range, 0.1)
points(dss_umap$layout[plot_1,1], dss_umap$layout[plot_1,2], col=plot_color[plot_1], pch=16, cex = 0.5)
for (q in seq(0.1,0.99, by=0.01)){
	plot_1 = ((color_sig_range > quantile(color_sig_range, q)) * (color_sig_range <= quantile(color_sig_range, q+0.01))) != 0
	points(dss_umap$layout[plot_1,1], dss_umap$layout[plot_1,2], col=plot_color[plot_1], pch=16, cex = 0.5)
}
dev.off()
}

pdf('umap.esRP.colorbar', width=3)
breaksList = seq(-quantile(as.numeric(as.matrix(dss)),0.99), quantile(as.numeric(as.matrix(dss)),0.99), by = 0.001)
my_colorbar=colorRampPalette(c('blue', 'gray90','red'))(n = length(breaksList))
col_breaks = c(seq(0, 2000,length=33))
pheatmap(a, color=my_colorbar, breaks = breaksList, cluster_cols = FALSE,cluster_rows=F, clustering_method = 'average',annotation_names_row=TRUE,annotation_names_col=TRUE,show_rownames=TRUE,show_colnames=TRUE)
dev.off()

### get umap with meta-clusterID
cCRE_clusters = read.table('../coe_analysis/S3V2_IDEAS_hg38_ccre2.cCRE.M.notall0.rmallNEU.withid.coe_mat.PDmerged.clusterID.txt', header=T)[used_row,]

dir.create('coe_umap_PD_meta_clusterBinary')
for (i in 1:length(unique(cCRE_clusters[,6]))){
	metaC_i = unique(cCRE_clusters[,6])[i]
png(paste('coe_umap_PD_meta_clusterBinary/', 'state_coe.human.umap.PD.', metaC_i,'.png', sep=''))
plot(dss_umap$layout[,1], dss_umap$layout[,2], col='gray90', pch=16, cex = 0.5, main = paste0('Meta-Cluster:', metaC_i), xlab='UMAP Dim1', ylab='UMAP Dim2')
points(dss_umap$layout[cCRE_clusters[,6]==metaC_i,1], dss_umap$layout[cCRE_clusters[,6]==metaC_i,2], col='black', pch=16, cex = 0.5)
dev.off()
}


### get umap with Jmeta-clusterID
cCRE_clusters0 = read.table('../coe_analysis/S3V2_IDEAS_hg38_ccre2.cCRE.M.notall0.rmallNEU.withid.coe_mat.PDmerged.clusterID.JclusterID.txt', header=T)
cCRE_clusters = cCRE_clusters0[used_row,1:7]
dir.create('coe_umap_PD_meta_clusterBinary')
for (i in 1:length(unique(cCRE_clusters[,7]))){
	metaC_i = unique(cCRE_clusters[,7])[i]
png(paste('coe_umap_PD_meta_clusterBinary/', 'state_coe.human.umap.PD.JmC.', metaC_i,'.png', sep=''))
plot(dss_umap$layout[,1], dss_umap$layout[,2], col='gray90', pch=16, cex = 0.5, main = paste0('Meta-Cluster:', metaC_i), xlab='UMAP Dim1', ylab='UMAP Dim2')
points(dss_umap$layout[cCRE_clusters[,7]==metaC_i,1], dss_umap$layout[cCRE_clusters[,7]==metaC_i,2], col='black', pch=16, cex = 0.5)
dev.off()
}
```
