## Supplementary material for "Interspecies regulatory landscapes and elements revealed by novel joint systematic integration of human and mouse blood cell epigenomes": Scripts for IDEAS modeling of epigenetic states jointly in human and mouse: reviewer1_point5_6_get_cCRE_ATACsig.get_ATACsig_umap.human.R

```
library(pheatmap)
library(umap)
library(LSD)
library(lisi)

# read ATACsig
setwd('/Users/guanjuexiang/Documents/projects/analysis/04_05_2022_coe/coe_analysis/')
d = read.table('S3V2_IDEAS_hg38_ccre2.cCRE.M.notall0.withid.ATACsig.withid.txt', header=T)
ds = d[,-c(1:4)]

###### For Proximal & Distal
set.seed(2019)
used_row = sample(dim(ds)[1], 20000)
dssP = as.matrix(ds[used_row,1:(dim(ds)[2])])


rep1 = c(1, 2, 4, 6, 8, 10, 12, 14, 16, 18, 20, 22, 24, 26, 28, 30, 32, 34, 36, 38, 40, 42)
rep2 = c(1, 3, 5, 7, 9, 11, 13, 15, 17, 19, 21, 23, 25, 27, 29, 31, 33, 35, 37, 39, 41, 43)

dssP_ct = c()
for (i in 1:length(rep1)){
	dssP_ct = cbind(dssP_ct, rowMeans(cbind(dssP[,rep1[i]], dssP[,rep2[i]])))
}

ct_list_human = c('AVE', 'B','CD34','CLP','CMP','EOS','ERY','GMP','HSC','HUDEP1','HUDEP2','K562','LMPP','MEP','MK','MONc','MONp','MPP','NEU','NK','T_CD4','T_CD8')

colnames(dssP_ct) = ct_list_human
dss = dssP_ct


### run umap
set.seed(2019)
dss_umap = umap(dssP_ct[, colnames(dssP_ct)!='NEU_C0011IH2_D'], preserve.seed=T)

# color umap by ATAC signal
dir.create('ATACsig_umap_PD_esRP/')
for (i in 1:dim(dssP)[2]){
	ct_i = colnames(dssP)[i]
png(paste('ATACsig_umap_PD_esRP/', 'state_ATAC.human.umap.PD.', ct_i,'.png', sep=''))
rbPal = colorRampPalette(c('blue', 'gray90','red'))
color_sig_range = (dssP[,i])
print(summary(color_sig_range))
color_sig_range[color_sig_range>quantile(as.numeric(as.matrix(dssP)),0.99)] = quantile(as.numeric(as.matrix(dssP)),0.99)
plot_color = rbPal(500)[as.numeric(cut(c(-quantile(as.numeric(as.matrix(dssP)),0.99), quantile(as.numeric(as.matrix(dssP)),0.99), color_sig_range),breaks = 500))][-c(1:2)]
plot(dss_umap$layout[,1], dss_umap$layout[,2], col='white', pch=16, main = ct_i, xlab='UMAP Dim1', ylab='UMAP Dim2', cex = 0.5)
plot_1 = color_sig_range <= quantile(color_sig_range, 0.1)
points(dss_umap$layout[plot_1,1], dss_umap$layout[plot_1,2], col=plot_color[plot_1], pch=16, cex = 0.5)
for (q in seq(0.1,0.99, by=0.01)){
	plot_1 = ((color_sig_range > quantile(color_sig_range, q)) * (color_sig_range <= quantile(color_sig_range, q+0.01))) != 0
	points(dss_umap$layout[plot_1,1], dss_umap$layout[plot_1,2], col=plot_color[plot_1], pch=16, cex = 0.5)
}
dev.off()
}


dir.create('ATACsig_umap_PD_esRP')
### plot heatscatter plot
png('ATACsig_umap_PD_esRP/state_ATACsig.human.umap.PD.png')
heatscatter(dss_umap$layout[,1], dss_umap$layout[,2], cex=0.5, xlab='UMAP Dim1', ylab='UMAP Dim2')
dev.off()

# read cCRE annotation label
cCRE_label = read.table('S3V2_IDEAS_hg38_ccre2.cCRE.M.notall0.withid.label.bed', header=F)

# get cCRE label sample
cCRE_label_sample = cCRE_label[used_row,]

# plot umap colored by cCRE label
uniq_cCRE_label = names(table(cCRE_label_sample[,4]))
color_list = c('orange','red','green')
pdf('ATACsig_umap_PD_esRP/umap.ATACsig.cCRElabel.pdf')
plot(dss_umap$layout[,1], dss_umap$layout[,2], col='gray', pch=16, main = '', xlab='UMAP Dim1', ylab='UMAP Dim2', cex = 0.5)
#for (i in 1:length(uniq_cCRE_label)){
for (i in c(1,2,3)){
	plot_1 = cCRE_label_sample[,4] == uniq_cCRE_label[i]
	points(dss_umap$layout[plot_1,1], dss_umap$layout[plot_1,2], col=color_list[i], pch=16, cex = 0.5)
}
dev.off()


### plot scatterplot
for (i in 1:dim(dss)[2]){
	ct_i = colnames(dss)[i]
png(paste('ATACsig_umap_PD_esRP/', 'state_ATACsig.human.umap.', ct_i,'.png', sep=''))
rbPal = colorRampPalette(c('blue', 'gray90','red'))
color_sig_range = (dss[,i])
print(summary(color_sig_range))
color_sig_range[color_sig_range>quantile(as.numeric(as.matrix(dss)),0.99)] = quantile(as.numeric(as.matrix(dss)),0.99)
plot_color = rbPal(500)[as.numeric(cut(c(-quantile(as.numeric(as.matrix(dss)),0.99), quantile(as.numeric(as.matrix(dss)),0.99), color_sig_range),breaks = 500))][-c(1:2)]
plot(dss_umap$layout[,1], dss_umap$layout[,2], col='white', pch=16, main = ct_i, xlab='UMAP Dim1', ylab='UMAP Dim2', cex = 0.5)
plot_1 = color_sig_range <= quantile(color_sig_range, 0.1)
points(dss_umap$layout[plot_1,1], dss_umap$layout[plot_1,2], col=plot_color[plot_1], pch=16, cex = 0.5)
for (q in seq(0.1,0.99, by=0.01)){
	plot_1 = ((color_sig_range > quantile(color_sig_range, q)) * (color_sig_range <= quantile(color_sig_range, q+0.01))) != 0
	points(dss_umap$layout[plot_1,1], dss_umap$layout[plot_1,2], col=plot_color[plot_1], pch=16, cex = 0.5)
}
dev.off()
}


# plot colorbar
pdf('ATACsig_umap_PD_esRP/umap.ATACsig.colorbar.pdf', width=3)
breaksList = seq(-quantile(as.numeric(as.matrix(dss)),0.99), quantile(as.numeric(as.matrix(dss)),0.99), by = 0.001)
my_colorbar=colorRampPalette(c('blue', 'gray90','red'))(n = length(breaksList))
col_breaks = c(seq(0, 2000,length=33))
a = cbind(dss[sample(dim(dss)[1], 100),1:2])
pheatmap(a, color=my_colorbar, breaks = breaksList, cluster_cols = FALSE,cluster_rows=F, clustering_method = 'average',annotation_names_row=TRUE,annotation_names_col=TRUE,show_rownames=TRUE,show_colnames=TRUE)
dev.off()

# lisi
library(lisi)
# lisi score
cCRE_label_sample_1 = cbind(cCRE_label_sample[,4])
colnames(cCRE_label_sample_1) = 'cCRE_label'
lisi_score = compute_lisi(dss_umap$layout, cCRE_label_sample_1, 'cCRE_label')

mean(lisi_score$cCRE_label)
mean(lisi_score$cCRE_label[cCRE_label_sample_1=='Intergenic'])
mean(lisi_score$cCRE_label[cCRE_label_sample_1=='Promoter'])
mean(lisi_score$cCRE_label[cCRE_label_sample_1=='TES'])


lisi_score_mat = c()
for (k in 1:30){
set.seed(k)
print(k)
used_row = sample(dim(ds)[1], 20000)
dssP = as.matrix(ds[used_row,1:(dim(ds)[2])])
# merege replicate
dssP_ct = c()
for (i in 1:length(rep1)){
	dssP_ct = cbind(dssP_ct, rowMeans(cbind(dssP[,rep1[i]], dssP[,rep2[i]])))
}
# define cell type names
colnames(dssP_ct) = ct_list_human
# umap
dss_umap = umap(dssP_ct[, colnames(dssP_ct)!='NEU_C0011IH2_D'], preserve.seed=T)
# get cCRE label sample
cCRE_label_sample = cCRE_label[used_row,]
# lisi score
cCRE_label_sample_1 = cbind(cCRE_label_sample[,4])
colnames(cCRE_label_sample_1) = 'cCRE_label'
lisi_score = compute_lisi(dss_umap$layout, cCRE_label_sample_1, 'cCRE_label')
lisi_score_mat = rbind(lisi_score_mat, c(mean(lisi_score$cCRE_label), mean(lisi_score$cCRE_label[cCRE_label_sample_1=='Intergenic']), mean(lisi_score$cCRE_label[cCRE_label_sample_1=='Promoter']), mean(lisi_score$cCRE_label[cCRE_label_sample_1=='TES'])))
print(lisi_score_mat)
write.table(lisi_score_mat, 'lisi_score_mat.ATAC.txt', sep='\t', quote=F, row.names=F, col.names=F)
}

write.table(lisi_score_mat, 'lisi_score_mat.ATAC.txt', sep='\t', quote=F, row.names=F, col.names=F)
```
