## Supplementary material for "Interspecies regulatory landscapes and elements revealed by novel joint systematic integration of human and mouse blood cell epigenomes": Scripts for IDEAS modeling of epigenetic states jointly in human and mouse: reviewer1_point5_6_get_cCRE_MKsig.hg38.R

```
args = commandArgs(trailingOnly=TRUE)

Beta_coe_mat_file = args[1]
cCRE_state_coverage_file_start = args[2]
state_max = as.numeric(args[3])
output_filename_start = args[4]
ct_list_human = args[-c(1:4)]

#Beta_coe_mat_file = 'statep_rna_coe_heatmap.human.all.ccre.withcorfilter.AVE.txt'
cCRE_state_coverage_file_start = 'S3V2_IDEAS_hg38_ccre2.cCRE.M.notall0.rmallNEU.withid.S'
state_max = 24
output_filename_start = 'S3V2_IDEAS_hg38_ccre2.cCRE.M.notall0.rmallNEU.withid.ATACsig'
ct_list_human = c('AVE', 'B_B15_50', 'B_NC14_42', 'CD34_E_rep1', 'CD34_E_rep2', 'CLP_100266', 'CLP_100267', 'CMP_100246', 'CMP_100247', 'EOS_S006XEH2', 'EOS_S00BKK', 'ERY_S002R5', 'ERY_S002S3', 'GMP_100256', 'GMP_100257', 'HSC_100258', 'HSC_100259', 'HUDEP1_rep1', 'HUDEP1_rep2', 'HUDEP2_rep1', 'HUDEP2_rep2', 'K562_rep1', 'K562_rep2', 'LMPP_100268', 'LMPP_100269', 'MEP_Donor2596', 'MEP_Donor7256', 'MK_S004BTH2', 'MK_S00VHKH1', 'MONc_C0011IH1', 'MONc_C001UYH2', 'MONp_Prim_mon_C', 'MONp_Prim_mon_F', 'MPP_100272', 'MPP_100273', 'NEU_C0011IH2', 'NEU_C001UYH1', 'NK_S005YG', 'NK_S01E4WH0', 'T_CD4_S008H1', 'T_CD4_S009W4', 'T_CD8_C0066PH1', 'T_CD8_S00C2FH1')


###################
###### human ######
# read table change comment sign
state_para = read.table('/Users/guanjuexiang/Documents/projects/Joint_Human_Mouse_IDEAS_State/VBSJ_052021_outputs_para_pdf/ES/06a_S3V2_IDEAS_hg38_r3_withHg38Mm10prior.para.modified.para', header=T, comment.char = "~")

# state signal mat
state_sig = apply(state_para[,2:9],2,function(x) x/state_para[,1])

# state color
state_color = read.table('/Users/guanjuexiang/Documents/projects/Joint_Human_Mouse_IDEAS_State/VBSJ_052021_outputs_para_pdf/ES/state_color.txt', header=F, comment.char = "~")

# coe
coe_mat = read.table('../coe_analysis/statep_rna_coe_heatmap.HM.all.ccre.withcorfilter.AVE.txt', header=T)
scale_ratio = max(coe_mat[,2])/max(coe_mat[,1])
coe_adj = (coe_mat[,1] * scale_ratio + coe_mat[,2])/2

# plot coe vs state MK average sig
dir.create('MKsig_vs_BetaCoef')
mk_list = colnames(state_sig)
for (i in 1:length(mk_list)){
	mk_i = mk_list[i]
	pdf(paste0('MKsig_vs_BetaCoef/', mk_i, '_vs_BetaCoef.pdf'), width=8, height=8)
	plot(coe_adj, state_sig[,i], cex=2, cex.axis=2, cex.lab=2, xlab='Beta-coef', ylab=mk_i)
	for (j in 1:length(coe_adj)){
		points(coe_adj[j], state_sig[j,i], pch=20, cex=2.1, col=state_color[j,3])
	}
	# add text rownames of coe_mat
	text(coe_adj, state_sig[,i], labels=rownames(coe_mat), pos=3, cex=1.5)
	abline(v=0, lty=2)
	dev.off()
	# mk only
	pdf(paste0('MKsig_vs_BetaCoef/', mk_i, '_only.pdf'), width=3, height=8)
	plot(rep(0,dim(state_sig)[1]), state_sig[,i], cex=2, cex.axis=2, cex.lab=2, xlab='', ylab=mk_i)
	for (j in 1:length(coe_adj)){
		points(0, state_sig[j,i], pch=20, cex=2.1, col=state_color[j,3])
	}
	# add text rownames of coe_mat
	text(rep(0,dim(state_sig)[1]), state_sig[,i], labels=rownames(coe_mat), pos=2, cex=3)
	abline(v=0, lty=2)
	dev.off()
}

pdf('BetaCoef_only.pdf', width=8, height=8)
plot(coe_adj, rep(0,dim(state_sig)[1]), cex=2, cex.axis=2, cex.lab=2, xlab='Beta-coef', ylab='ATAC')
for (i in 1:length(coe_adj)){
	points(coe_adj[i], 0, pch=20, cex=2.1, col=state_color[i,3])
}
# add text rownames of coe_mat
text(coe_adj, rep(0,dim(state_sig)[1]), labels=rownames(coe_mat), pos=2, cex=1.5)
abline(v=0, lty=2)
dev.off()

pdf('BetaCoef_only.vertical.pdf', width=3, height=8)
plot(rep(0,dim(state_sig)[1]), coe_adj, cex=2, cex.axis=2, cex.lab=2, xlab='', ylab='Beta-coef')
for (i in 1:length(coe_adj)){
	points(0, coe_adj[i], pch=20, cex=2.1, col=state_color[i,3])
}
# add text rownames of coe_mat
text(rep(0,dim(state_sig)[1]), coe_adj, labels=rownames(coe_mat), pos=2, cex=1.5)
abline(v=0, lty=2)
dev.off()


print('read state count')
Hs = read.table(paste0(cCRE_state_coverage_file_start, '0.mat.txt'), header=F)

print('normalize dist')
HsP0 = as.matrix(Hs[,-c(1:4)])

print('state count * ATACsig')
HsP = HsP0 * state_sig[1,1]

for (i in 1:state_max){
	print(i)
	Hs_i = read.table(paste0(cCRE_state_coverage_file_start, i, '.mat.txt'), header=F)
	HsP0 = as.matrix(Hs_i[,-c(1:4)])
	HsP = HsP + HsP0 * state_sig[i+1,1]
}

set.seed(2019)
# add noise
HsP = HsP + matrix(rnorm(dim(HsP)[1]*dim(HsP)[2], sd=0.2), nrow=dim(HsP)[1], ncol=dim(HsP)[2])
HsP[HsP<0] = 0

# round
HsP = round(HsP, 3)

print('add colnames')
print(ct_list_human)
colnames(HsP) = ct_list_human

print('output mat')
HsPD_mat = cbind(Hs[,1:4], (HsP))
colnames(HsPD_mat)[1:4] = c('chr','start','end','id')

write.table(HsPD_mat, 'S3V2_IDEAS_hg38_ccre2.cCRE.M.notall0.withid.ATACsig.withid.txt', quote=F, col.names=T, row.names=F, sep='\t')

HsPD_mat = HsPD_mat[,!is.element(colnames(HsPD_mat), c('NEU_C0011IH2', 'NEU_C001UYH1'))]

write.table(HsPD_mat, 'S3V2_IDEAS_hg38_ccre2.cCRE.M.notall0.rmallNEU.withid.ATACsig.withid.txt', quote=F, col.names=T, row.names=F, sep='\t')


# H3K27ac

print('read state count')
Hs = read.table(paste0(cCRE_state_coverage_file_start, '0.mat.txt'), header=F)

print('normalize dist')
HsP0 = as.matrix(Hs[,-c(1:4)])

print('state count * ATACsig')
HsP = HsP0 * state_sig[1,3]

for (i in 1:state_max){
	print(i)
	Hs_i = read.table(paste0(cCRE_state_coverage_file_start, i, '.mat.txt'), header=F)
	HsP0 = as.matrix(Hs_i[,-c(1:4)])
	HsP = HsP + HsP0 * state_sig[i+1,3]
}

set.seed(2019)
# add noise
HsP = HsP + matrix(rnorm(dim(HsP)[1]*dim(HsP)[2], sd=0.2), nrow=dim(HsP)[1], ncol=dim(HsP)[2])
HsP[HsP<0] = 0

# round
HsP = round(HsP, 3)

print('add colnames')
print(ct_list_human)
colnames(HsP) = ct_list_human

print('output mat')
HsPD_mat = cbind(Hs[,1:4], (HsP))
colnames(HsPD_mat)[1:4] = c('chr','start','end','id')

write.table(HsPD_mat, 'S3V2_IDEAS_hg38_ccre2.cCRE.M.notall0.withid.H3K27acsig.withid.txt', quote=F, col.names=T, row.names=F, sep='\t')

HsPD_mat = HsPD_mat[,!is.element(colnames(HsPD_mat), c('NEU_C0011IH2', 'NEU_C001UYH1'))]

write.table(HsPD_mat, 'S3V2_IDEAS_hg38_ccre2.cCRE.M.notall0.rmallNEU.withid.H3K27acsig.withid.txt', quote=F, col.names=T, row.names=F, sep='\t')
```
