## Supplementary material for "Interspecies regulatory landscapes and elements revealed by novel joint systematic integration of human and mouse blood cell epigenomes": Scripts for IDEAS modeling of epigenetic states jointly in human and mouse: reviewer1_point5_6_get_cCRE_MKsig.mm10.R

```
###################
###### mouse ######
ct_list_mouse = c('AVE', 'B_r1', 'B_r2', 'CFUE_r1', 'CFUMK_r1', 'CLP_r1', 'CMP_r1', 'ER4_r1', 'ER4_r2', 'ERY_ad_r1', 'ERY_ad_r2', 'ERY_fl_r1', 'ERY_fl_r2', 'G1E_r1', 'G1E_r2', 'GMP_r1', 'HPC7_r1', 'LSK_r1', 'MEL_r1', 'MEL_r2', 'MEP_r1', 'MK_fl_r1', 'MK_fl_r2', 'MON_r1', 'NEU_r1', 'NK_r1', 'T_CD4_r1', 'T_CD8_r1', 'iMEL_r1', 'iMEL_r2', 'iMK_r1', 'iMK_r2')

# read table change comment sign
state_para = read.table('/Users/guanjuexiang/Documents/projects/Joint_Human_Mouse_IDEAS_State/VBSJ_052021_outputs_para_pdf/ES/06a_S3V2_IDEAS_hg38_r3_withHg38Mm10prior.para.modified.para', header=T, comment.char = "~")

# state signal mat
state_sig = apply(state_para[,2:9],2,function(x) x/state_para[,1])

# ATAC
### read state count
Hs = read.table('../coe_analysis_mouse/S3V2_IDEAS_mm10_ccre2.cCRE.M.notall0.withid.S0.mat.txt', header=F)

### normalize dist
HsP0 = as.matrix(Hs[,-c(1:4)])

### state count * coe
HsP = HsP0 * state_sig[1,1]

for (i in 1:24){
	print(i)
Hs_i = read.table(paste('../coe_analysis_mouse/S3V2_IDEAS_mm10_ccre2.cCRE.M.notall0.withid.S',i,'.mat.txt', sep=''), header=F)
HsP0 = as.matrix(Hs_i[,-c(1:4)])
HsP = HsP + HsP0 * state_sig[i+1,1]
}

### add colnames
colnames(HsP) = paste(ct_list_mouse, '', sep='')

### output mat
HsPD_mat = cbind(Hs[,1:4], (HsP))
colnames(HsPD_mat)[1:4] = c('chr','start','end','id')

write.table(HsPD_mat, '../coe_analysis_mouse/S3V2_IDEAS_mm10_ccre2.cCRE.M.notall0.withid.ATACsig.withid.txt', quote=F, col.names=T, row.names=F, sep='\t')


# H3K27ac
### read state count
Hs = read.table('../coe_analysis_mouse/S3V2_IDEAS_mm10_ccre2.cCRE.M.notall0.withid.S0.mat.txt', header=F)

### normalize dist
HsP0 = as.matrix(Hs[,-c(1:4)])

### state count * coe
HsP = HsP0 * state_sig[1,3]

for (i in 1:24){
	print(i)
Hs_i = read.table(paste('../coe_analysis_mouse/S3V2_IDEAS_mm10_ccre2.cCRE.M.notall0.withid.S',i,'.mat.txt', sep=''), header=F)
HsP0 = as.matrix(Hs_i[,-c(1:4)])
HsP = HsP + HsP0 * state_sig[i+1,3]
}

### add colnames
colnames(HsP) = paste(ct_list_mouse, '', sep='')

### output mat
HsPD_mat = cbind(Hs[,1:4], (HsP))
colnames(HsPD_mat)[1:4] = c('chr','start','end','id')

write.table(HsPD_mat, '../coe_analysis_mouse/S3V2_IDEAS_mm10_ccre2.cCRE.M.notall0.withid.H3K27acsig.withid.txt', quote=F, col.names=T, row.names=F, sep='\t')
```
