## Supplementary material for "Interspecies regulatory landscapes and elements revealed by novel joint systematic integration of human and mouse blood cell epigenomes": Scripts for IDEAS modeling of epigenetic states jointly in human and mouse: reviewer1_point5_6_get_gencode_annotation.R

```
cd /Users/guanjuexiang/Documents/projects/analysis/04_05_2022_coe/coe_analysis

# download data
wget https://ftp.ebi.ac.uk/pub/databases/gencode/Gencode_human/release_35/gencode.v35.annotation.gtf.gz
gunzip gencode.v35.annotation.gtf.gz

# get cCRE bed
cat S3V2_IDEAS_hg38_ccre2.cCRE.M.notall0.withid.ATACsig.withid.txt | tail -n+2 | cut -f1,2,3,4 > S3V2_IDEAS_hg38_ccre2.cCRE.M.notall0.withid.bed

# only get gene rows
cat gencode.v35.annotation.gtf | awk -F '\t' -v OFS='\t' '{if ($3=="gene") print $0}' > gencode.v35.annotation.gene.gtf

# get gene coordinates and strand and type
cat gencode.v35.annotation.gene.gtf | awk -F '\t' -v OFS='\t' '{print $1,$4,$5,$7}' > gencode.v35.annotation.gene.coordinates.bed
cat gencode.v35.annotation.gene.gtf | awk -F '\t' -v OFS='\t' '{print $9}' | awk -F '"' -v OFS='\t' '{print $2,$4,$6}' > gencode.v35.annotation.gene.info.txt
paste gencode.v35.annotation.gene.coordinates.bed gencode.v35.annotation.gene.info.txt > gencode.v35.annotation.gene.coordinates.info.bed
rm gencode.v35.annotation.gene.coordinates.bed gencode.v35.annotation.gene.info.txt

# get TSS +/- 2kb
cat gencode.v35.annotation.gene.coordinates.info.bed | awk -F '\t' -v OFS='\t' '{if ($6=="protein_coding") print $0}' | awk -F '\t' -v OFS='\t' '{if ($4=="+") print $1,$2-2000,$2,$4,$5,$6,$7; else print $1,$3,$3+2000,$4,$5,$6,$7}' | awk -F '\t' -v OFS='\t' '{if ($2<0) $2=0; if ($3<0) $3=0; print $0}' > gencode.v35.annotation.gene.TSS.updown2kb.bed
# get gene body
cat gencode.v35.annotation.gene.coordinates.info.bed | awk -F '\t' -v OFS='\t' '{if ($6=="protein_coding") print $0}' | awk -F '\t' -v OFS='\t' '{print $0}' > gencode.v35.annotation.gene.body.bed
# get TES +/- 2kb
cat gencode.v35.annotation.gene.coordinates.info.bed | awk -F '\t' -v OFS='\t' '{if ($6=="protein_coding") print $0}' | awk -F '\t' -v OFS='\t' '{if ($4=="+") print $1,$3,$3+2000,$4,$5,$6,$7; else print $1,$2-2000,$2,$4,$5,$6,$7}' | awk -F '\t' -v OFS='\t' '{if ($2<0) $2=0; if ($3<0) $3=0; print $0}' > gencode.v35.annotation.gene.TES.updown2kb.bed
# 

# bedtools intersect
bedtools intersect -a S3V2_IDEAS_hg38_ccre2.cCRE.M.notall0.withid.bed -b gencode.v35.annotation.gene.TSS.updown2kb.bed -c > S3V2_IDEAS_hg38_ccre2.cCRE.M.notall0.withid.TSS.updown2kb.bed
bedtools intersect -a S3V2_IDEAS_hg38_ccre2.cCRE.M.notall0.withid.bed -b annotation_orthogonal/human_ccre_hg38_ctcf6ct.bed -c > S3V2_IDEAS_hg38_ccre2.cCRE.M.notall0.withid.gene_body.bed
bedtools intersect -a S3V2_IDEAS_hg38_ccre2.cCRE.M.notall0.withid.bed -b gencode.v35.annotation.gene.TES.updown2kb.bed -c > S3V2_IDEAS_hg38_ccre2.cCRE.M.notall0.withid.TES.updown2kb.bed

# define label
paste S3V2_IDEAS_hg38_ccre2.cCRE.M.notall0.withid.TSS.updown2kb.bed \
S3V2_IDEAS_hg38_ccre2.cCRE.M.notall0.withid.TES.updown2kb.bed \
S3V2_IDEAS_hg38_ccre2.cCRE.M.notall0.withid.gene_body.bed | \
awk -F '\t' -v OFS='\t' '{if ($5!=0) print $1,$2,$3,"Promoter"; else if ($10!=0) print $1,$2,$3,"TES"; else print $1,$2,$3,"Intergenic"}' > S3V2_IDEAS_hg38_ccre2.cCRE.M.notall0.withid.label.bed

#awk -F '\t' -v OFS='\t' '{if ($5!=0) print $1,$2,$3,"Promoter"; else if ($10!=0) print $1,$2,$3,"TES"; else if ($15!=0) print $1,$2,$3,"TADbnd"; else print $1,$2,$3,"Intergenic"}' > S3V2_IDEAS_hg38_ccre2.cCRE.M.notall0.withid.label.bed
```
