## Supplementary material for "Interspecies regulatory landscapes and elements revealed by novel joint systematic integration of human and mouse blood cell epigenomes": Scripts for IDEAS modeling of epigenetic states jointly in human and mouse: reviewer3_point18_plot_state_dif.R

```
library(pheatmap)

# hg38 IDEAS state round 0
dh1 = read.table('01a_S3V2_IDEAS_hg38_r1.para', header=T, comment.char = "~")
dm1 = read.table('01b_S3V2_IDEAS_mm10_r1.para', header=T, comment.char = "~")

# get state average signal
dh1s = apply(dh1[,2:9],2,function(x) x/dh1[,1])
dm1s = apply(dm1[,2:9],2,function(x) x/dm1[,1])

# add noise
set.seed(2019)
dh1s = dh1s + rnorm(dim(dh1s)[1]*dim(dh1s)[2], mean=0, sd=0.1)
dm1s = dm1s + rnorm(dim(dm1s)[1]*dim(dm1s)[2], mean=0, sd=0.1)

# add rownames
rownames(dh1s) = paste0('h',1:dim(dh1s)[1])
rownames(dm1s) = paste0('m',1:dim(dm1s)[1])

# plot heatmap
pdf('heatmap_hg38_mm10.r1.pdf', width=8, height=16)
set.seed(2019)
pheatmap(rbind(dh1s,dm1s), cluster_rows = T, cluster_cols = T, show_rownames = T, show_colnames = T, border_color = NA)
dev.off()

# get the pairwise Euclidean distance between dh1s and dm1s
dh1s_dm1s_dist = matrix(0, nrow = dim(dh1s)[1], ncol = dim(dm1s)[1])
for (i in 1:dim(dh1s)[1]){
    for (j in 1:dim(dm1s)[1]){
        dh1s_dm1s_dist[i,j] = dist(rbind(dh1s[i,],dm1s[j,]), method = "euclidean")
    }
}

# get row/col min binary matrix
dh1s_dm1s_dist_rowmin = t(apply(dh1s_dm1s_dist, 1, function(x) (x == min(x))*1))
dh1s_dm1s_dist_colmin = apply(dh1s_dm1s_dist, 2, function(x) (x == min(x))*1)

# Mutual nearest neighbor
dh1s_dm1s_dist_rowmin_colmin = dh1s_dm1s_dist_rowmin*dh1s_dm1s_dist_colmin

# get the distance between the Mutual nearest neighbor of the two matrices
mnn_mse_vec_r1 = c()
for (i in 1:dim(dh1s_dm1s_dist_rowmin_colmin)[1]){
    dh1s_dm1s_dist_rowmin_colmin_row_i = dh1s_dm1s_dist_rowmin_colmin[i,]
    if (sum(dh1s_dm1s_dist_rowmin_colmin_row_i) > 0){
        mnn_mse_vec_r1 = c(mnn_mse_vec_r1, mean((dh1s[i,]-dm1s[which(dh1s_dm1s_dist_rowmin_colmin_row_i==1),])^2))
    }
}


# hg38 IDEAS state round 1
dp2 = read.table('02_hg38_mm10_joint_states_8mk.52.para', header=T, comment.char = "~")
dh2 = read.table('03_S3V2_IDEAS_hg38_r2_with_prior_J.para', header=T, comment.char = "~")
dm2 = read.table('04_S3V2_IDEAS_mm10_r2_withhg38prior.para', header=T, comment.char = "~")

# get state average signal
dp2s = apply(dp2[,2:9],2,function(x) x/dp2[,1])
dh2s = apply(dh2[,2:9],2,function(x) x/dh2[,1])
dm2s = apply(dm2[,2:9],2,function(x) x/dm2[,1])

# add rownames
rownames(dp2s) = paste0('p',1:dim(dp2s)[1])
rownames(dh2s) = paste0('h',1:dim(dh2s)[1])
rownames(dm2s) = paste0('m',1:dim(dm2s)[1])

# plot heatmap
pdf('heatmap_prior_hg38.r2.pdf', width=8, height=16)
set.seed(2019)
pheatmap(rbind(dp2s,dh2s), cluster_rows = T, cluster_cols = T, show_rownames = T, show_colnames = T, border_color = NA)
dev.off()

pdf('heatmap_hg38_mm10.r2.pdf', width=8, height=16)
set.seed(2019)
pheatmap(rbind(dh2s,dm2s), cluster_rows = T, cluster_cols = T, show_rownames = T, show_colnames = T, border_color = NA)
dev.off()

# get the pairwise Euclidean distance between dp2s and dh2s
dp2s_dh2s_dist = matrix(0, nrow = dim(dp2s)[1], ncol = dim(dh2s)[1])
for (i in 1:dim(dp2s)[1]){
    for (j in 1:dim(dh2s)[1]){
        dp2s_dh2s_dist[i,j] = dist(rbind(dp2s[i,],dh2s[j,]), method = "euclidean")
    }
}

# get row/col min binary matrix
dp2s_dh2s_dist_rowmin = t(apply(dp2s_dh2s_dist, 1, function(x) (x == min(x))*1))
dp2s_dh2s_dist_colmin = apply(dp2s_dh2s_dist, 2, function(x) (x == min(x))*1)

# Mutual nearest neighbor
dp2s_dh2s_dist_rowmin_colmin = dp2s_dh2s_dist_rowmin*dp2s_dh2s_dist_colmin

# get the distance between the Mutual nearest neighbor of the two matrices
mnn_mse_vec_r2_prior_hg38 = c()
for (i in 1:dim(dp2s_dh2s_dist_rowmin_colmin)[1]){
    dp2s_dh2s_dist_rowmin_colmin_row_i = dp2s_dh2s_dist_rowmin_colmin[i,]
    if (sum(dp2s_dh2s_dist_rowmin_colmin_row_i) > 0){
        mnn_mse_vec_r2_prior_hg38 = c(mnn_mse_vec_r2_prior_hg38, mean((dp2s[i,]-dh2s[which(dp2s_dh2s_dist_rowmin_colmin_row_i==1),])^2))
    }
}


# get the pairwise Euclidean distance between dh2s and dm2s
dh2s_dm2s_dist = matrix(0, nrow = dim(dh2s)[1], ncol = dim(dm2s)[1])
for (i in 1:dim(dh2s)[1]){
    for (j in 1:dim(dm2s)[1]){
        dh2s_dm2s_dist[i,j] = dist(rbind(dh2s[i,],dm2s[j,]), method = "euclidean")
    }
}

# get row/col min binary matrix
dh2s_dm2s_dist_rowmin = t(apply(dh2s_dm2s_dist, 1, function(x) (x == min(x))*1))
dh2s_dm2s_dist_colmin = apply(dh2s_dm2s_dist, 2, function(x) (x == min(x))*1)

# Mutual nearest neighbor
dh2s_dm2s_dist_rowmin_colmin = dh2s_dm2s_dist_rowmin*dh2s_dm2s_dist_colmin

# get the distance between the Mutual nearest neighbor of the two matrices
mnn_mse_vec_r2_hg38_mm10 = c()
for (i in 1:dim(dh2s_dm2s_dist_rowmin_colmin)[1]){
    dh2s_dm2s_dist_rowmin_colmin_row_i = dh2s_dm2s_dist_rowmin_colmin[i,]
    if (sum(dh2s_dm2s_dist_rowmin_colmin_row_i) > 0){
        mnn_mse_vec_r2_hg38_mm10 = c(mnn_mse_vec_r2_hg38_mm10, mean((dh2s[i,]-dm2s[which(dh2s_dm2s_dist_rowmin_colmin_row_i==1),])^2))
    }
}


# hg38 IDEAS state round 2
dp3 = read.table('05_S3V2_IDEAS_mm10_r2_withhg38prior.rmh.10.para', header=T, comment.char = "~")
dh3 = read.table('06a_S3V2_IDEAS_hg38_r3_withHg38Mm10prior.para.modified.para', header=T, comment.char = "~")
dm3 = read.table('06b_S3V2_IDEAS_mm10_r3_withHg38Mm10prior.para.modified.para', header=T, comment.char = "~")

# get state average signal
dp3s = apply(dp3[,2:9],2,function(x) x/dp3[,1])
dh3s = apply(dh3[,2:9],2,function(x) x/dh3[,1])
dm3s = apply(dm3[,2:9],2,function(x) x/dm3[,1])

# add rownames
rownames(dp3s) = paste0('p',1:dim(dp3s)[1])
rownames(dh3s) = paste0('h',1:dim(dh3s)[1])
rownames(dm3s) = paste0('m',1:dim(dm3s)[1])

# plot heatmap
pdf('heatmap_prior_hg38.r3.pdf', width=8, height=16)
set.seed(2019)
pheatmap(rbind(dp3s,dh3s), cluster_rows = T, cluster_cols = T, show_rownames = T, show_colnames = T, border_color = NA)
dev.off()

pdf('heatmap_hg38_mm10.r3.pdf', width=8, height=16)
set.seed(2019)
pheatmap(rbind(dh3s,dm3s), cluster_rows = T, cluster_cols = T, show_rownames = T, show_colnames = T, border_color = NA)
dev.off()

# get the pairwise Euclidean distance between dp3s and dh3s
dp3s_dh3s_dist = matrix(0, nrow = dim(dp3s)[1], ncol = dim(dh3s)[1])
for (i in 1:dim(dp3s)[1]){
    for (j in 1:dim(dh3s)[1]){
        dp3s_dh3s_dist[i,j] = dist(rbind(dp3s[i,],dh3s[j,]), method = "euclidean")
    }
}

# get row/col min binary matrix
dp3s_dh3s_dist_rowmin = t(apply(dp3s_dh3s_dist, 1, function(x) (x == min(x))*1))
dp3s_dh3s_dist_colmin = apply(dp3s_dh3s_dist, 2, function(x) (x == min(x))*1)

# Mutual nearest neighbor
dp3s_dh3s_dist_rowmin_colmin = dp3s_dh3s_dist_rowmin*dp3s_dh3s_dist_colmin

# get the distance between the Mutual nearest neighbor of the two matrices
mnn_mse_vec_r3_prior_hg38 = c()
for (i in 1:dim(dp3s_dh3s_dist_rowmin_colmin)[1]){
    dp3s_dh3s_dist_rowmin_colmin_row_i = dp3s_dh3s_dist_rowmin_colmin[i,]
    if (sum(dp3s_dh3s_dist_rowmin_colmin_row_i) > 0){
        mnn_mse_vec_r3_prior_hg38 = c(mnn_mse_vec_r3_prior_hg38, mean((dp3s[i,]-dh3s[which(dp3s_dh3s_dist_rowmin_colmin_row_i==1),])^2))
    }
}


# get the pairwise Euclidean distance between dh3s and dm3s
dh3s_dm3s_dist = matrix(0, nrow = dim(dh3s)[1], ncol = dim(dm3s)[1])
for (i in 1:dim(dh3s)[1]){
    for (j in 1:dim(dm3s)[1]){
        dh3s_dm3s_dist[i,j] = dist(rbind(dh3s[i,],dm3s[j,]), method = "euclidean")
        #dh3s_dm3s_dist[i,j] = 1-cor(dh3s[i,],dm3s[j,])
    }
}

# get row/col min binary matrix
dh3s_dm3s_dist_rowmin = t(apply(dh3s_dm3s_dist, 1, function(x) (x == min(x))*1))
dh3s_dm3s_dist_colmin = apply(dh3s_dm3s_dist, 2, function(x) (x == min(x))*1)

# Mutual nearest neighbor
dh3s_dm3s_dist_rowmin_colmin = dh3s_dm3s_dist_rowmin*dh3s_dm3s_dist_colmin

# get the distance between the Mutual nearest neighbor of the two matrices
mnn_mse_vec_r3_hg38_mm10 = c()
for (i in 1:dim(dh3s_dm3s_dist_rowmin_colmin)[1]){
    dh3s_dm3s_dist_rowmin_colmin_row_i = dh3s_dm3s_dist_rowmin_colmin[i,]
    if (sum(dh3s_dm3s_dist_rowmin_colmin_row_i) > 0){
        mnn_mse_vec_r3_hg38_mm10 = c(mnn_mse_vec_r3_hg38_mm10, mean((dh3s[i,]-dm3s[which(dh3s_dm3s_dist_rowmin_colmin_row_i==1),])^2))
    }
}


mean(mnn_mse_vec_r1)
mean(mnn_mse_vec_r2_prior_hg38)
mean(mnn_mse_vec_r2_hg38_mm10)
mean(mnn_mse_vec_r3_prior_hg38)
mean(mnn_mse_vec_r3_hg38_mm10)

# plot mse
pdf('mse_prior_hg38_mm10.pdf', width=5, height=5)
plot(1:3, c(mean(mnn_mse_vec_r2_hg38_mm10), mean(mnn_mse_vec_r3_prior_hg38), mean(mnn_mse_vec_r3_hg38_mm10)), xlim=c(0.5,3.5), xlab='', ylab='MNN MSE', cex=1.5, pch=19, col='dodgerblue', xaxt='n')
# lines
labels = c('Hr1_Mr1', 'Mr1_Hr2', 'Hr2_Mr2')
axis(1, at=1:3, labels=FALSE)
text(x=1:3+0.3, par("usr")[3], labels=labels, srt=45, adj=1.5, xpd=TRUE)
lines(1:3, c(mean(mnn_mse_vec_r2_hg38_mm10), mean(mnn_mse_vec_r3_prior_hg38), mean(mnn_mse_vec_r3_hg38_mm10)), type='b', col='dodgerblue')
dev.off()
```
