## Supplementary material for "Interspecies regulatory landscapes and elements revealed by novel joint systematic integration of human and mouse blood cell epigenomes": Scripts for IDEAS modeling of epigenetic states jointly in human and mouse: get_state_heatmap_for_workflow_fig.R

```
state_human = c()
for (i in 1:100){
	state_human_i =  read.table(paste('01aa_hg38_r1_tmp_para/S3V2_IDEAS_hg38_r1.tmp.', i,'.para', sep=''), comment="!",header=T)
	m=as.matrix(state_human_i[,1+1:8]/state_human_i[,1])
	m = m[,order(colnames(m))]
	state_human = rbind(state_human, m)
}


state_mouse = c()
for (i in 1:100){
	state_human_i =  read.table(paste('01ba_mm10_r1_tmp_para/S3V2_IDEAS_mm10_r1.tmp.', i,'.para', sep=''), comment="!",header=T)
	m=as.matrix(state_human_i[,1+1:8]/state_human_i[,1])
	m = m[,order(colnames(m))]
	state_mouse = rbind(state_mouse, m)
}

rownames(state_human) = paste('H:', 1:dim(state_human)[1], sep='')
rownames(state_mouse) = paste('M:', 1:dim(state_mouse)[1], sep='')

library(pheatmap)

state_human_mouse = rbind(state_human, state_mouse)

state_dist_mat = as.matrix(dist(state_human_mouse))
state_hclust = hclust(dist(state_human_mouse), method = 'ward.D')

state_hclust_cutree = cutree(state_hclust, k = 100)

pdf('dist_mat.pdf', height=20, , width=20)
used_rows = sample(dim(state_dist_mat)[1], 500)
pheatmap(1-state_dist_mat[used_rows,used_rows],  clustering_method = "average", cluster_cols=T, cluster_rows=T)
dev.off()


full_dist = read.table('dist.mat.full.txt', header=F)

library(pheatmap)

set.seed(2019)
full_dist_sample = sample(dim(full_dist)[1], 50)
pdf('state_dist.heatmap.pdf')
pheatmap(-full_dist[full_dist_sample,full_dist_sample])
dev.off()


my_colorbar=colorRampPalette(c('white', 'blue4'))(n = 128)
pheatmap(state_human_mouse[sample(dim(state_human_mouse)[1], 100),], clustering_method = "average", cluster_cols=F, color=my_colorbar)


pheatmap(state_human_mouse[state_hclust_cutree==36,], clustering_method = "average", cluster_cols=F, color=my_colorbar)
pheatmap(state_human_mouse[state_hclust_cutree==36 | state_hclust_cutree==100,], clustering_method = "average", cluster_cols=F, color=my_colorbar)


state_human_mouse_mean = c()
state_human_mouse_mean_n = c()
for (i in 1:max(state_hclust_cutree)){
	state_human_mouse_mean = rbind(state_human_mouse_mean, colMeans(state_human_mouse[state_hclust_cutree==i,]))
	state_human_mouse_mean_n = c(state_human_mouse_mean_n, sum(state_hclust_cutree==i))
}


rownames(state_human_mouse_mean) = apply(cbind(1:max(state_hclust_cutree), state_human_mouse_mean_n), 1, function(x) paste(x[1],x[2], sep=': '))

pdf('tmp.pdf', height=20)
pheatmap(state_human_mouse_mean,  clustering_method = "average", cluster_cols=F, color=my_colorbar)
dev.off()


pdf('tmp1.pdf', height=20)
pheatmap(rbind(state_human_mouse[state_hclust_cutree==56,], state_human_mouse[state_hclust_cutree==78,]),  clustering_method = "average", cluster_cols=F, cluster_rows=F, color=my_colorbar)
dev.off()

pdf('tmp2.pdf', height=20)
pheatmap(state_human_mouse[state_hclust_cutree==56,],  clustering_method = "average", cluster_cols=F, color=my_colorbar)
dev.off()
```
