## Supplementary material for "Interspecies regulatory landscapes and elements revealed by novel joint systematic integration of human and mouse blood cell epigenomes": Scripts for IDEAS modeling of epigenetic states jointly in human and mouse: get_chr5_NR3C1_fimoscore.sh

```
bedtools getfasta -fi chr5.fa -bed bins.200bp.chr5.bed > bins.200bp.chr5.bed.fa

/storage/home/gzx103/work/software_group/meme_4.12.0/src/fimo --thresh 0.1 --text NR3C1.MA0113.3.meme bins.200bp.chr5.bed.fa > bins.200bp.chr5.bed.fa.NR3C1.MA0113.3.fimo.txt

cut -f3,8 bins.200bp.chr5.bed.fa.NR3C1.MA0113.3.fimo.txt > bins.200bp.chr5.bed.fa.NR3C1.MA0113.3.fimo.info.txt


time python find_best_match_fimo.py -i bins.200bp.chr5.bed.fa.NR3C1.MA0113.3.fimo.txt -o check.txt

cat check.txt | awk -F '\t' -v OFS='\t' '{print $3,$8}' | awk -F ':' -v OFS='\t' '{print $1,$2}' | awk -F '-' -v OFS='\t' '{print $1,$2}' | awk -F '\t' -v OFS='\t' '{if ((-(log($4)/log(10)))>3) print $1,$2,$3,-(log($4)/log(10))}'> check.bed

~/work/software_group/ucsc/bedGraphToBigWig check.bed ~/group/genome/mm10/mm10.chrom.sizes NR3C1.chr5.fimo.bw

R


d = read.table('bins.200bp.chr5.bed.fa.NR3C1.MA0113.3.fimo.info.txt', header=T, sep='\t')
dp = -log10(d[,2])

dpn = d[,1]
d_unique = unique(dpn)


dm = apply(cbind(d_unique[1000]), 1, function(x) max(dpn[dpn==x]))
```
