## Supplementary material for "Interspecies regulatory landscapes and elements revealed by novel joint systematic integration of human and mouse blood cell epigenomes": Scripts for IDEAS modeling of epigenetic states jointly in human and mouse: get_GC_matched_rand_pk.include_othersets.sh

```
cd ~/scratch/gc_percentage/

GC percentage:
wget http://hgdownload.cse.ucsc.edu/gbdb/hg38/bbi/gc5BaseBw/gc5Base.bw

Mappability:
wget https://hgdownload.soe.ucsc.edu/gbdb/hg38/hoffmanMappability/k50.Umap.MultiTrackMappability.bw


target_pk='bp24b_V2_2.nonALL0.cCRE.withid.bed'

### get target pk GC
time ~/group/software/ucsc/bigWigAverageOverBed gc5Base.bw $target_pk tmp.GC.txt
cut -f1,6 tmp.GC.txt | awk -F '_' -v OFS='\t' '{print $1,$2,$3}' | sort -k4,4n > tmp.GC.bed


### get random peaks
for seed in {1..30}
do
	echo $seed
	echo get rand pk
	bedtools random -seed $seed -l 305 -n 2235290 -g /storage/home/gzx103/group/genome/hg38/hg38.chrom.1_22_X_Y.sizes > 'r.bin.'$seed'.bed'
	cut -f1,2,3 'r.bin.'$seed'.bed' | awk -F '\t' -v OFS='\t' '{print $1,$2,$3, $1"_"$2"_"$3}' | sort -u > 'r.bin.'$seed'.withid.bed'
	echo get rand pk GC
	random_pk='r.bin.'$seed'.withid.bed'
	time ~/group/software/ucsc/bigWigAverageOverBed gc5Base.bw $random_pk tmp.r.GC.txt
	cut -f1,6 tmp.r.GC.txt | awk -F '_' -v OFS='\t' '{print $1,$2,$3}' | sort -k4,4n > tmp.r.GC.bed
	echo get adj rand pk
	time Rscript get_GC_matched_rand_pk.R tmp.GC.bed tmp.r.GC.bed 'r.s'$seed'.GCadj.bed' 200 100
	echo filter intersect peaks
	bedtools intersect -a 'r.s'$seed'.GCadj.bed' -b bp24b_V2_2.nonALL0.cCRE.withid.bed -v > 'r.s'$seed'.nocCRE.GCadj.bed'
	wc -l 'r.s'$i'.nocCRE.GCadj.bed'
	time Rscript sampe_pk_n.R bp24b_V2_2.nonALL0.cCRE.withid.bed 'r.s'$seed'.nocCRE.GCadj.bed'
	mv 'r.s'$seed'.nocCRE.GCadj.beds.bed' 'r.s'$seed'.nocCRE.GCadj.s.bed'
done


######## for DHS cCRE

cd ~/scratch/gc_percentage/DHS_rand

GC percentage:
wget http://hgdownload.cse.ucsc.edu/gbdb/hg38/bbi/gc5BaseBw/gc5Base.bw

Mappability:
wget https://hgdownload.soe.ucsc.edu/gbdb/hg38/hoffmanMappability/k50.Umap.MultiTrackMappability.bw


cat DHS.ME.bed | awk -F '\t' -v OFS='\t' '{print $1,$2,$3,$1"_"$2"_"$3}' > DHS.ME.withid.bed
target_pk='DHS.ME.withid.bed'

### get target pk GC
time ~/group/software/ucsc/bigWigAverageOverBed ~/scratch/gc_percentage/gc5Base.bw $target_pk tmp.GC.txt
cut -f1,6 tmp.GC.txt | awk -F '_' -v OFS='\t' '{print $1,$2,$3}' | sort -k4,4n > tmp.GC.bed


### get random peaks
for seed in {1..30}
do
	echo $seed
	echo get rand pk
	bedtools random -seed $seed -l 192 -n 1866160 -g /storage/home/gzx103/group/genome/hg38/hg38.chrom.1_22_X_Y.sizes > 'r.bin.'$seed'.bed'
	cut -f1,2,3 'r.bin.'$seed'.bed' | awk -F '\t' -v OFS='\t' '{print $1,$2,$3, $1"_"$2"_"$3}' | sort -u > 'r.bin.'$seed'.withid.bed'
	echo get rand pk GC
	random_pk='r.bin.'$seed'.withid.bed'
	time ~/group/software/ucsc/bigWigAverageOverBed ~/scratch/gc_percentage/gc5Base.bw $random_pk tmp.r.GC.txt
	cut -f1,6 tmp.r.GC.txt | awk -F '_' -v OFS='\t' '{print $1,$2,$3}' | sort -k4,4n > tmp.r.GC.bed
	echo get adj rand pk
	time Rscript ~/scratch/gc_percentage/get_GC_matched_rand_pk.R tmp.GC.bed tmp.r.GC.bed 'r.s'$seed'.GCadj.bed' 20 100
	echo filter intersect peaks
	bedtools intersect -a 'r.s'$seed'.GCadj.bed' -b $target_pk -v > 'r.s'$seed'.nocCRE.GCadj.bed'
	wc -l 'r.s'$seed'.nocCRE.GCadj.bed'
	time Rscript ~/scratch/gc_percentage/sampe_pk_n.R $target_pk 'r.s'$seed'.nocCRE.GCadj.bed'
	mv 'r.s'$seed'.nocCRE.GCadj.beds.bed' 'r.s'$seed'.nocCRE.GCadj.s.bed'
done


######## for DHSall cCRE

cd ~/scratch/gc_percentage/DHSall_rand


cat DHS.M.bed | awk -F '\t' -v OFS='\t' '{print $1,$2,$3,$1"_"$2"_"$3}' > DHS.M.withid.bed

time Rscript ~/scratch/gc_percentage/sampe_pk_n.R /storage/home/gzx103/scratch/gc_percentage/bp24b_V2_2.nonALL0.cCRE.withid.bed DHS.M.withid.bed

target_pk='DHS.M.withid.beds.bed'
target_pk0='DHS.M.withid.bed'

### get target pk GC
time ~/group/software/ucsc/bigWigAverageOverBed ~/scratch/gc_percentage/gc5Base.bw $target_pk tmp.GC.txt
cut -f1,6 tmp.GC.txt | awk -F '_' -v OFS='\t' '{print $1,$2,$3}' | sort -k4,4n > tmp.GC.bed


### get random peaks
for seed in {1..30}
do
	echo $seed
	echo get rand pk
	bedtools random -seed $seed -l 274 -n 4035290 -g /storage/home/gzx103/group/genome/hg38/hg38.chrom.1_22_X_Y.sizes > 'r.bin.'$seed'.bed'
	cut -f1,2,3 'r.bin.'$seed'.bed' | awk -F '\t' -v OFS='\t' '{print $1,$2,$3, $1"_"$2"_"$3}' | sort -u > 'r.bin.'$seed'.withid.bed'
	echo get rand pk GC
	random_pk='r.bin.'$seed'.withid.bed'
	time ~/group/software/ucsc/bigWigAverageOverBed ~/scratch/gc_percentage/gc5Base.bw $random_pk tmp.r.GC.txt
	cut -f1,6 tmp.r.GC.txt | awk -F '_' -v OFS='\t' '{print $1,$2,$3}' | sort -k4,4n > tmp.r.GC.bed
	echo get adj rand pk
	time Rscript ~/scratch/gc_percentage/get_GC_matched_rand_pk.R tmp.GC.bed tmp.r.GC.bed 'r.s'$seed'.GCadj.bed' 20 100
	echo filter intersect peaks
	bedtools intersect -a 'r.s'$seed'.GCadj.bed' -b $target_pk0 -v > 'r.s'$seed'.nocCRE.GCadj.bed'
	wc -l 'r.s'$seed'.nocCRE.GCadj.bed'
	time Rscript ~/scratch/gc_percentage/sampe_pk_n.R $target_pk 'r.s'$seed'.nocCRE.GCadj.bed'
	mv 'r.s'$seed'.nocCRE.GCadj.beds.bed' 'r.s'$seed'.nocCRE.GCadj.s.bed'
done

time Rscript ~/scratch/gc_percentage/sampe_pk_num.R 136186 $target_pk

for seed in {1..30}
do
	echo $seed
	time Rscript ~/scratch/gc_percentage/sampe_pk_n.R $target_pk's.bed' 'r.s'$seed'.nocCRE.GCadj.bed'
	mv 'r.s'$seed'.nocCRE.GCadj.beds.bed' 'r.s'$seed'.nocCRE.GCadj.s.bed'
done


######## for SCREEN cCRE

cd ~/scratch/gc_percentage/SCREEN_rand


cat SCREEN.M.bed | awk -F '\t' -v OFS='\t' '{print $1,$2,$3,$1"_"$2"_"$3}' > SCREEN.M.withid.bed

time Rscript ~/scratch/gc_percentage/sampe_pk_n.R /storage/home/gzx103/scratch/gc_percentage/bp24b_V2_2.nonALL0.cCRE.withid.bed SCREEN.M.withid.bed

target_pk='SCREEN.M.withid.beds.bed'
target_pk0='SCREEN.M.withid.bed'

### get target pk GC
time ~/group/software/ucsc/bigWigAverageOverBed ~/scratch/gc_percentage/gc5Base.bw $target_pk tmp.GC.txt
cut -f1,6 tmp.GC.txt | awk -F '_' -v OFS='\t' '{print $1,$2,$3}' | sort -k4,4n > tmp.GC.bed


### get random peaks
for seed in {1..30}
do
	echo $seed
	echo get rand pk
	bedtools random -seed $seed -l 274 -n 4035290 -g /storage/home/gzx103/group/genome/hg38/hg38.chrom.1_22_X_Y.sizes > 'r.bin.'$seed'.bed'
	cut -f1,2,3 'r.bin.'$seed'.bed' | awk -F '\t' -v OFS='\t' '{print $1,$2,$3, $1"_"$2"_"$3}' | sort -u > 'r.bin.'$seed'.withid.bed'
	echo get rand pk GC
	random_pk='r.bin.'$seed'.withid.bed'
	time ~/group/software/ucsc/bigWigAverageOverBed ~/scratch/gc_percentage/gc5Base.bw $random_pk tmp.r.GC.txt
	cut -f1,6 tmp.r.GC.txt | awk -F '_' -v OFS='\t' '{print $1,$2,$3}' | sort -k4,4n > tmp.r.GC.bed
	echo get adj rand pk
	time Rscript ~/scratch/gc_percentage/get_GC_matched_rand_pk.R tmp.GC.bed tmp.r.GC.bed 'r.s'$seed'.GCadj.bed' 20 100
	echo filter intersect peaks
	bedtools intersect -a 'r.s'$seed'.GCadj.bed' -b $target_pk0 -v > 'r.s'$seed'.nocCRE.GCadj.bed'
	wc -l 'r.s'$seed'.nocCRE.GCadj.bed'
	time Rscript ~/scratch/gc_percentage/sampe_pk_n.R $target_pk 'r.s'$seed'.nocCRE.GCadj.bed'
	mv 'r.s'$seed'.nocCRE.GCadj.beds.bed' 'r.s'$seed'.nocCRE.GCadj.s.bed'
done

time Rscript ~/scratch/gc_percentage/sampe_pk_num.R 194490 $target_pk

for seed in {1..30}
do
	echo $seed
	time Rscript ~/scratch/gc_percentage/sampe_pk_n.R $target_pk 'r.s'$seed'.nocCRE.GCadj.bed'
	mv 'r.s'$seed'.nocCRE.GCadj.beds.bed' 'r.s'$seed'.nocCRE.GCadj.s.bed'
done


######### get intersect counts

cd ~/scratch/gc_percentage/interrsect_counts

bedtools intersect -a /storage/home/gzx103/group/projects/vision_human/independent_cRE_peaklist/ERY_ep300.fdr02.bed -b ~/scratch/gc_percentage/DHS_rand/DHS.ME.withid.bed -wa -u > test.txt
wc -l test.txt > DHS.intersect.counts.txt
for i in {1..30}
do
	echo $i
	bedtools intersect -a /storage/home/gzx103/group/projects/vision_human/independent_cRE_peaklist/ERY_ep300.fdr02.bed -b ~/scratch/gc_percentage/DHS_rand/r.s$i'.nocCRE.GCadj.s.bed' -wa -u > test.txt
	wc -l test.txt >> DHS.intersect.counts.txt
done


bedtools intersect -a /storage/home/gzx103/group/projects/vision_human/independent_cRE_peaklist/ERY_ep300.fdr02.bed -b ~/scratch/gc_percentage/bp24b_V2_2.nonALL0.cCRE.withid.bed -wa -u > test.txt
wc -l test.txt > VISION.intersect.counts.txt
for i in {1..30}
do
	echo $i
	bedtools intersect -a /storage/home/gzx103/group/projects/vision_human/independent_cRE_peaklist/ERY_ep300.fdr02.bed -b ~/scratch/gc_percentage/r.s$i'.nocCRE.GCadj.s.bed' -wa -u > test.txt
	wc -l test.txt >> VISION.intersect.counts.txt
done


bedtools intersect -a /storage/home/gzx103/group/projects/vision_human/independent_cRE_peaklist/ERY_ep300.fdr02.bed -b ~/scratch/gc_percentage/SCREEN_rand/SCREEN.M.withid.beds.bed -wa -u > test.txt
wc -l test.txt > SCREEN.intersect.counts.txt
for i in {1..30}
do
	echo $i
	bedtools intersect -a /storage/home/gzx103/group/projects/vision_human/independent_cRE_peaklist/ERY_ep300.fdr02.bed -b ~/scratch/gc_percentage/SCREEN_rand/r.s$i'.nocCRE.GCadj.s.bed' -wa -u > test.txt
	wc -l test.txt >> SCREEN.intersect.counts.txt
done


R

V = read.table('VISION.intersect.counts.txt', header=F)
D = read.table('DHS.intersect.counts.txt', header=F)
S = read.table('SCREEN.intersect.counts.txt', header=F)

pnorm(V[1,1],mean(V[-1,1]),sd(V[-1,1]), lower.tail=F)

pnorm(D[1,1],mean(D[-1,1]),sd(D[-1,1]), lower.tail=F)

pnorm(S[1,1],mean(S[-1,1]),sd(S[-1,1]), lower.tail=F)
```
