## Supplementary material for "Interspecies regulatory landscapes and elements revealed by novel joint systematic integration of human and mouse blood cell epigenomes": Scripts for IDEAS modeling of epigenetic states jointly in human and mouse: get_random_ccre_set.mm10.sh

```
#!/bin/bash

#SBATCH --mem=200G
#SBATCH --cpus-per-task=12

source ~/.bashrc

cd /homes1/gxiang/projects/impute_cistrome/get_neg_set_mm10

j=$1
bash -c "RANDOM=$j"

### split orignial cCRE list into 12 peaks
head -10000 S3V2_IDEAS_mm10_ccre2.cCRE.M.notall0.bed > S3V2_IDEAS_mm10_ccre2.cCRE.M.notall0.1.bed
head -20000 S3V2_IDEAS_mm10_ccre2.cCRE.M.notall0.bed | tail -n+10001 > S3V2_IDEAS_mm10_ccre2.cCRE.M.notall0.2.bed
head -30000 S3V2_IDEAS_mm10_ccre2.cCRE.M.notall0.bed | tail -n+20001 > S3V2_IDEAS_mm10_ccre2.cCRE.M.notall0.3.bed
head -40000 S3V2_IDEAS_mm10_ccre2.cCRE.M.notall0.bed | tail -n+30001 > S3V2_IDEAS_mm10_ccre2.cCRE.M.notall0.4.bed
head -50000 S3V2_IDEAS_mm10_ccre2.cCRE.M.notall0.bed | tail -n+40001 > S3V2_IDEAS_mm10_ccre2.cCRE.M.notall0.5.bed
head -60000 S3V2_IDEAS_mm10_ccre2.cCRE.M.notall0.bed | tail -n+50001 > S3V2_IDEAS_mm10_ccre2.cCRE.M.notall0.6.bed
head -70000 S3V2_IDEAS_mm10_ccre2.cCRE.M.notall0.bed | tail -n+60001 > S3V2_IDEAS_mm10_ccre2.cCRE.M.notall0.7.bed
head -80000 S3V2_IDEAS_mm10_ccre2.cCRE.M.notall0.bed | tail -n+70001 > S3V2_IDEAS_mm10_ccre2.cCRE.M.notall0.8.bed
head -90000 S3V2_IDEAS_mm10_ccre2.cCRE.M.notall0.bed | tail -n+80001 > S3V2_IDEAS_mm10_ccre2.cCRE.M.notall0.9.bed
head -100000 S3V2_IDEAS_mm10_ccre2.cCRE.M.notall0.bed | tail -n+90001 > S3V2_IDEAS_mm10_ccre2.cCRE.M.notall0.10.bed


#echo $j
### get random peaks
for i in {1..10}
do
	echo $i
	time bash Negetive_sequence_matched_length_GC.sh S3V2_IDEAS_mm10_ccre2.cCRE.M.notall0.$i.bed mm10.chrom.1_19X.sizes mm10.fa 5 &
done


### pool random peaks
#rm S3V2_IDEAS_mm10_ccre2.cCRE.M.notall0.N.bed_matched.$j.bed
#for i in {1..12}
#do
#	cat S3V2_IDEAS_mm10_ccre2.cCRE.M.notall0.$i.bed_matched.bed >> S3V2_IDEAS_mm10_ccre2.cCRE.M.notall0.N.bed_matched.$j.bed
#done
```
