## Supplementary material for "Interspecies regulatory landscapes and elements revealed by novel joint systematic integration of human and mouse blood cell epigenomes": Scripts for IDEAS modeling of epigenetic states jointly in human and mouse: Negetive_sequence_matched_length_GC.mm10.sh

```
#!/bin/bash
##########################################################################
#This script is for generating length and GC matched background sequence
#Created by Houyu Zhang on 2020-07-07.
#Issue report on
#Copyright (c) 2020 __YenLab@SKLEH__. All rights reserved.
##########################################################################
if [ -n "$1" ]; then
    echo "--->>> You provided the arguments: $@, Let's do it..."
else
    echo "This script is for generating length and GC content matched background sequence, heavily using bedtools" 1>&2
    echo "  $(basename $0) [input bed file] [genome size] [genome fasta] [iteration times]"  1>&2
    echo "  [input bed file]: A bed file as template, just first 3 cols are used" 1>&2
    echo "  [genome size]: Genome size for bedtools" 1>&2
    echo "  [genome fatsa]: Genome Fatsa file for bedtools" 1>&2
    echo "  [iteration times]: Sequence pools for comparison, 100 is pretty good" 1>&2
    exit 1
fi

bed_file=$1
GENOME_SIZE=$2
FASTA_FILE=$3
Iter=$4

#1. Process input bed file to obtain GC content
cut -f 1-3 ${bed_file} | bedtools nuc -fi ${FASTA_FILE} -bed - | sed '1d' | awk -v FS="\t" -v OFS="\t" '{print "ref_"$1,$2,$3,$5}' > ${bed_file}.fabed

#2. Generate sequence pools for comparison
for i in `seq 1 ${Iter}`
do
    echo "Generating ${i} dataset..."
    cut -f 1-3 ${bed_file} | bedtools shuffle -g ${GENOME_SIZE} -noOverlapping -i - \
    | bedtools nuc -fi ${FASTA_FILE} -bed - | sed '1d' | awk -v FS="\t" -v OFS="\t" '{print $1,$2,$3,$5}' > ${bed_file}_shuf_${i}.fabed
done

#3. Compare GC content between ref bed and pools, line by line
line_num=$(cat $bed_file | wc -l)
echo $line_num

rm ${bed_file}_matched.bed

for i in `seq 1 ${line_num}`
do
    #echo $i
    #A tmp file for processing current line
    Proc_file=${bed_file}_Processing.txt

    #Output all ith line from ref bed and pools  
    awk -v FS="\t" -v OFS="\t" -v line=${i} 'FNR==line {print FILENAME, $0}' ${bed_file}*fabed | sort -k5n > ${Proc_file}

    #The rank of ref line will be captured, and the content be kept for output
    ref_line=`grep -n --color="never" "ref" ${Proc_file} | sed "s/:.*//g"`
    awk -v FS="\t" -v OFS="\t" -v line=${ref_line} 'FNR==line {print $0}' ${Proc_file}| cut -f 2-5 | sed "s/ref_//g"> ${bed_file}_tmp

    #just 1 line ahead will be selected with similar GC content. So, larger pool get more similar GC sequence, but slower :)
    if [ ${ref_line} -eq 1 ]
    then
        similar_line=`expr $ref_line + 1`
    else
        similar_line=`expr $ref_line - 1`
    fi
    #Output selected sequence with template sequence together
    awk -v FS="\t" -v OFS="\t" -v line=${similar_line} 'FNR==line {print $0}' ${Proc_file} | cut -f 2-5 | paste - ${bed_file}_tmp >> ${bed_file}_matched.bed
done    

#4. Delete tmp files
rm -f ${bed_file}_Processing.txt ${bed_file}*fabed ${bed_file}_tmp

### Parallel run this scipt
#If you want to accelerate the calculation, you can split original file into "Thread" number files and run simultaneously.
#Thread=10
#for file in *bed
#do
#    split -n l/${Thread} ${file} ${file}_split_
#
#    for j in ${file}_split_*
#    do
#        nohup bash Negetive_sequence_matched_length_GC.sh ${j} ${GENOME_SIZE} ${FASTA_FILE} 500 &
#    done
#done
```
