## Supplementary material for "Interspecies regulatory landscapes and elements revealed by novel joint systematic integration of human and mouse blood cell epigenomes": Scripts for IDEAS modeling of epigenetic states jointly in human and mouse: readme.txt

script for GC matched bed file is downloaded from the following link:
https://github.com/YenLab/Tn5InsertPrefer/blob/main/StandaloneScripts/Negetive_sequence_matched_length

##########################################################################
#This script is for generating length and GC matched background sequence
#Created by Houyu Zhang on 2020-07-07.
#Issue report on
#Copyright (c) 2020 __YenLab@SKLEH__. All rights reserved.
##########################################################################
......
