## Supplementary material for "Interspecies regulatory landscapes and elements revealed by novel joint systematic integration of human and mouse blood cell epigenomes": Scripts for IDEAS modeling of epigenetic states jointly in human and mouse: get_intersect_pk.ERY_EP300.12_2021.sh

```
cd /storage/home/gzx103/scratch/gc_percentage/interrsect_counts_ERYEP300_2021

### get SCREEN peaks
target_pk='SCREEN.M.withid.bed'
target_r_pk='r.s1.GCadj.bed'
data_folder='/storage/home/gzx103/scratch/gc_percentage/SCREEN_rand/'
true_file='ERY_ep300.fdr02.bed'
cp $data_folder$target_pk ./
cp $data_folder$target_r_pk ./
rm target.txt
rm target.rand.txt
for seed in {1..30}
do
	echo $seed
	#time Rscript ~/scratch/gc_percentage/sampe_pk_num.withseed.R 100000 $target_pk $seed
	seed_add1=$((seed+1))
	cat '/storage/home/gzx103/scratch/gc_percentage/interrsect_counts_ERYEP300_2021/SCREEN_202112/SCREEN.M.withid.bed.'$seed'.sample.bed' '/storage/home/gzx103/scratch/gc_percentage/interrsect_counts_ERYEP300_2021/SCREEN_202112/SCREEN.M.withid.bed.'$seed_add1'.sample.bed' > '/storage/home/gzx103/scratch/gc_percentage/interrsect_counts_ERYEP300_2021/SCREEN_202112/SCREEN.M.withid.bed.'$seed'.'$seed_add1'.sample.bed'
	bedtools intersect -a $true_file -b '/storage/home/gzx103/scratch/gc_percentage/interrsect_counts_ERYEP300_2021/SCREEN_202112/SCREEN.M.withid.bed.'$seed'.'$seed_add1'.sample.bed' -wa -u > tmp.txt
	wc -l tmp.txt >> target.txt
	#time Rscript ~/scratch/gc_percentage/sampe_pk_num.withseed.R 100000 $target_r_pk $seed
	cat '/storage/home/gzx103/scratch/gc_percentage/interrsect_counts_ERYEP300_2021/SCREEN_202112/SCREEN.M.withid.bed.'$seed'.sample.bed_matched.bed' '/storage/home/gzx103/scratch/gc_percentage/interrsect_counts_ERYEP300_2021/SCREEN_202112/SCREEN.M.withid.bed.'$seed_add1'.sample.bed_matched.bed' > '/storage/home/gzx103/scratch/gc_percentage/interrsect_counts_ERYEP300_2021/SCREEN_202112/SCREEN.M.withid.bed.'$seed'.'$seed_add1'.sample.bed_matched.bed'
	bedtools intersect -a $true_file -b '/storage/home/gzx103/scratch/gc_percentage/interrsect_counts_ERYEP300_2021/SCREEN_202112/SCREEN.M.withid.bed.'$seed'.'$seed_add1'.sample.bed_matched.bed' -wa -u > tmp.txt
	wc -l tmp.txt >> target.rand.txt
done
paste target.txt target.rand.txt | awk -F ' ' -v OFS='\t' '{print $1,$3}' > SCREEN.counts.txt

### get DHSall peaks
target_pk='DHS.M.withid.bed'
target_r_pk='r.s1.GCadj.bed'
data_folder='/storage/home/gzx103/scratch/gc_percentage/DHSall_rand/'
true_file='ERY_ep300.fdr02.bed'
cp $data_folder$target_pk ./
cp $data_folder$target_r_pk ./
rm target.txt
rm target.rand.txt
for seed in {1..30}
do
	echo $seed
	seed_add1=$((seed+1))
	#time Rscript ~/scratch/gc_percentage/sampe_pk_num.withseed.R 200000 $target_pk $seed
	cat '/storage/home/gzx103/scratch/gc_percentage/interrsect_counts_ERYEP300_2021/DHS_202112/DHS.M.withid.bed.'$seed'.sample.bed' '/storage/home/gzx103/scratch/gc_percentage/interrsect_counts_ERYEP300_2021/DHS_202112/DHS.M.withid.bed.'$seed_add1'.sample.bed' > '/storage/home/gzx103/scratch/gc_percentage/interrsect_counts_ERYEP300_2021/DHS_202112/DHS.M.withid.bed.'$seed'.'$seed_add1'.sample.bed'
	bedtools intersect -a $true_file -b '/storage/home/gzx103/scratch/gc_percentage/interrsect_counts_ERYEP300_2021/DHS_202112/DHS.M.withid.bed.'$seed'.'$seed_add1'.sample.bed' -wa -u > tmp.txt
	wc -l tmp.txt >> target.txt
	#time Rscript ~/scratch/gc_percentage/sampe_pk_num.withseed.R 100000 $target_r_pk $seed
	cat '/storage/home/gzx103/scratch/gc_percentage/interrsect_counts_ERYEP300_2021/DHS_202112/DHS.M.withid.bed.'$seed'.sample.bed_matched.bed' '/storage/home/gzx103/scratch/gc_percentage/interrsect_counts_ERYEP300_2021/DHS_202112/DHS.M.withid.bed.'$seed_add1'.sample.bed_matched.bed' > '/storage/home/gzx103/scratch/gc_percentage/interrsect_counts_ERYEP300_2021/DHS_202112/DHS.M.withid.bed.'$seed'.'$seed_add1'.sample.bed_matched.bed'
	bedtools intersect -a $true_file -b '/storage/home/gzx103/scratch/gc_percentage/interrsect_counts_ERYEP300_2021/DHS_202112/DHS.M.withid.bed.'$seed'.'$seed_add1'.sample.bed_matched.bed' -wa -u > tmp.txt
	wc -l tmp.txt >> target.rand.txt
done
paste target.txt target.rand.txt | awk -F ' ' -v OFS='\t' '{print $1,$3}' > DHSall.counts.txt

### get VISION peaks
target_pk='S3V2_IDEAS_hg38_ccre2.cCRE.M.notall0.rmallNEU.bed'
target_r_pk='VISION_202112/S3V2_IDEAS_hg38_ccre2.cCRE.M.notall0.N.bed_matched.2.bed'
data_folder='/storage/home/gzx103/scratch/gc_percentage/'
true_file='ERY_ep300.fdr02.bed'
#cp $data_folder$target_pk ./
#cp $data_folder$target_r_pk ./

wget https://raw.githubusercontent.com/guanjue/Joint_Human_House_IDEAS_State/main/VBSJ_052021_outputs_para_pdf/CCREs/S3V2_IDEAS_hg38_ccre2.cCRE.M.notall0.rmallNEU.bed
cut -f1,2,3 EP300_ERY_FDR_0.2.interval.txt > ERY_ep300.fdr02.bed
rm target.txt
rm target.rand.txt
for seed in {1..30}
do
	echo $seed
	seed_add1=$((seed+1))
	#time Rscript ~/scratch/gc_percentage/sampe_pk_num.withseed.R 50000 $target_pk $seed
	cat '/storage/home/gzx103/scratch/gc_percentage/interrsect_counts_ERYEP300_2021/VISION_202112/S3V2_IDEAS_hg38_ccre2.cCRE.M.notall0.rmallNEU.withid.bed.'$seed'.sample.bed' '/storage/home/gzx103/scratch/gc_percentage/interrsect_counts_ERYEP300_2021/VISION_202112/S3V2_IDEAS_hg38_ccre2.cCRE.M.notall0.rmallNEU.withid.bed.'$seed_add1'.sample.bed' > '/storage/home/gzx103/scratch/gc_percentage/interrsect_counts_ERYEP300_2021/VISION_202112/S3V2_IDEAS_hg38_ccre2.cCRE.M.notall0.rmallNEU.withid.bed.'$seed'.'$seed_add1'.sample.bed'
	bedtools intersect -a $true_file -b '/storage/home/gzx103/scratch/gc_percentage/interrsect_counts_ERYEP300_2021/VISION_202112/S3V2_IDEAS_hg38_ccre2.cCRE.M.notall0.rmallNEU.withid.bed.'$seed'.'$seed_add1'.sample.bed' -wa -u > tmp.txt
	wc -l tmp.txt >> target.txt
	#time Rscript ~/scratch/gc_percentage/sampe_pk_num.withseed.R 50000 $target_r_pk $seed
	cat '/storage/home/gzx103/scratch/gc_percentage/interrsect_counts_ERYEP300_2021/VISION_202112/S3V2_IDEAS_hg38_ccre2.cCRE.M.notall0.rmallNEU.withid.bed.'$seed'.sample.bed_matched.bed' '/storage/home/gzx103/scratch/gc_percentage/interrsect_counts_ERYEP300_2021/VISION_202112/S3V2_IDEAS_hg38_ccre2.cCRE.M.notall0.rmallNEU.withid.bed.'$seed_add1'.sample.bed_matched.bed' > '/storage/home/gzx103/scratch/gc_percentage/interrsect_counts_ERYEP300_2021/VISION_202112/S3V2_IDEAS_hg38_ccre2.cCRE.M.notall0.rmallNEU.withid.bed.'$seed'.'$seed_add1'.sample.bed_matched.bed'
	bedtools intersect -a $true_file -b '/storage/home/gzx103/scratch/gc_percentage/interrsect_counts_ERYEP300_2021/VISION_202112/S3V2_IDEAS_hg38_ccre2.cCRE.M.notall0.rmallNEU.withid.bed.'$seed'.'$seed_add1'.sample.bed_matched.bed' -wa -u > tmp.txt
	wc -l tmp.txt >> target.rand.txt
done
paste target.txt target.rand.txt | awk -F ' ' -v OFS='\t' '{print $1,$3}' > VISION.counts.txt


R

library(ggplot2)

V = read.table('VISION.counts.txt', header=F)
D = read.table('DHSall.counts.txt', header=F)
#D = D[17:30,]
S = read.table('SCREEN.counts.txt', header=F)

sm_num = 0
wilcox.test((V[,1]+sm_num)/(V[,2]+sm_num), (S[,1]+sm_num)/(S[,2]+sm_num), alternative='greater')
wilcox.test((V[,1]+sm_num)/(V[,2]+sm_num), (D[,1]+sm_num)/(D[,2]+sm_num), alternative='greater')

enrichment = data.frame()
Edf = cbind(as.data.frame(c((D[,1]+sm_num)/((D[,2])+sm_num), (S[,1]+sm_num)/((S[,2])+sm_num), (V[,1]+sm_num)/((V[,2])+sm_num))), c(rep('DHS', dim(D)[1]), rep('SCR', dim(S)[1]), rep('VISION', dim(V)[1]))  )
colnames(Edf) = c('Enrichment', 'Methods')

pdf('enrichment.ERYEP300.12_2021.pdf', width=3, height=2)
p = ggplot(data = Edf, aes(x=Methods, y=Enrichment))
p = p + geom_hline(yintercept=1, linetype="dashed", color = "gray")
p = p + geom_boxplot(aes(fill = Methods), outlier.size = 0.2)
p = p + scale_fill_manual(values=rep(c('gray', 'orange1', 'dodgerblue1'),each = 1))
p = p + theme(panel.background = element_blank(), panel.border = element_rect(colour = "black", fill=NA, size=1))
p = p + ylim(0.5, max(Edf[,1])+1)
plot(p)
dev.off()
```
