## Supplementary material for "Interspecies regulatory landscapes and elements revealed by novel joint systematic integration of human and mouse blood cell epigenomes": Scripts for IDEAS modeling of epigenetic states jointly in human and mouse: get_fdr_p.hg38.R

```
d=read.table('cCRE.ATACseq.NBPsig.mat.txt', header=T)
ds = d[,-c(1:4)]
dsp = 10^(-ds)
dspfdr = p.adjust(as.vector(t(dsp)), 'fdr')

dspfdrm = (t(matrix(dspfdr, nrow=dim(dsp)[2])))

cor(dspfdrm[,1],dsp[,1])
cor(dspfdrm[,2],dsp[,2])
cor(dspfdrm[,3],dsp[,3])
cor(dspfdrm[,1],dsp[,3])
cor(dspfdrm[,1],dsp[,2])
cor(dspfdrm[,3],dsp[,2])

ddspfdrm = cbind(d[,1:4], -log10(dspfdrm))
colnames(dspfdrm) = colnames(d)[-c(1:4)]
write.table(ddspfdrm, 'cCRE.ATACseq.NBPsig.fdr.mat.txt', quote=F, col.names=T, row.names=F, sep='\t')
```
