## Supplementary material for "Interspecies regulatory landscapes and elements revealed by novel joint systematic integration of human and mouse blood cell epigenomes": Scripts for IDEAS modeling of epigenetic states jointly in human and mouse: get_signal_mat.sh

```
#PBS -l nodes=1:ppn=1
#PBS -l walltime=20:00:00
#PBS -j oe
##PBS -A yzz2_e_g_sc_default
#PBS -A rch8_b_g_bc_default
#PBS -l pmem=20gb

cd /storage/home/gzx103/scratch/joint_human_mouse/hg38/S3V2_IDEAS_outputs_hg38/S3V2_IDEAS_hg38_r3_withHg38Mm10prior_IDEAS_snapshot/output

sort -k4,4 snapshot.sort.bed > snapshot.sort.mat.txt

input_folder='/storage/home/gzx103/scratch/joint_human_mouse/hg38/S3V2_IDEAS_outputs_hg38/S3V2_IDEAS_hg38_r1_bws_NBP/'

for file in $(cat /storage/home/gzx103/scratch/joint_human_mouse/hg38/S3V2_IDEAS_outputs_hg38/NBP.bw.list.txt)
do
	echo $file
	~/work/software_group/S3V2_IDEAS_ESMP/bin/bigWigAverageOverBed $input_folder/$file snapshot.sort.bed tmp1.txt
	sort -k1,1 tmp1.txt | cut -f6 > tmp2.txt
	paste snapshot.sort.mat.txt tmp2.txt > snapshot.sort.mat.txt.tmp && mv snapshot.sort.mat.txt.tmp snapshot.sort.mat.txt
done
```
