## Supplementary material for "Interspecies regulatory landscapes and elements revealed by novel joint systematic integration of human and mouse blood cell epigenomes": Scripts for IDEAS modeling of epigenetic states jointly in human and mouse: get_state_barplot.hg38.R

```
### get parameters
args = commandArgs(trailingOnly=TRUE)
index_matrix_ideas_state_inputfile = args[1]
index_matrix_index_inputfile = args[2]
signal_input_list = args[3]
ideas_state_color = args[4]
cREs_IDEASpro_outfile = args[5]

### read index set matrix
read_color = function(x){
	rgb_color_int = as.numeric(unlist(strsplit(x, ',')))
	rgb_color = rgb(rgb_color_int[1],rgb_color_int[2],rgb_color_int[3],max=255)
	return(rgb_color)
}

setwd('/gpfs/scratch/gzx103/joint_human_mouse/hg38/S3V2_IDEAS_outputs_hg38/S3V2_IDEAS_hg38_r3_withHg38Mm10prior_IDEAS_snapshot/output')
index_matrix_index_inputfile = 'snapshot.index.matrix.txt'
index_matrix_ideas_state_inputfile = 'snapshot.IDEAS.matrix.txt'
ideas_state_color = 'state_color.txt'
cREs_IDEASpro_outfile = 'snapshot.hg38.barplot.pdf'

#################################################### 
############ read input files
####################################################
### read signal matrix file
print('read signal matrix file')
index_matrix_od = as.matrix(read.table(index_matrix_index_inputfile, header=TRUE))
print('read ideas state matrix file')
state_matrix_od = as.matrix(read.table(index_matrix_ideas_state_inputfile, header=TRUE))
index_matrix = index_matrix_od[ , -c(1:4)]
index_matrix = apply(index_matrix, 2, as.numeric)
state_matrix = state_matrix_od[,is.element(colnames(state_matrix_od), colnames(index_matrix_od))][,-c(1:4)]
print(dim(index_matrix))
print(dim(state_matrix))
### convert to numeric matrix
class(state_matrix) = 'numeric'

### index set
print('get uniq index set id')
index_set_id = signal_matrix_od[,4]
index_set_id_uniq = unique(index_set_id)
### sort index
index_set_id_uniq_sort = sort(index_set_id_uniq)

### set heatmap colors
print('set heatmap colors')
rgb_col_num0 = read.table(ideas_state_color,header=F)
rgb_col_num = rgb_col_num0[,2]
print(rgb_col_num)
rgb_col=apply((rgb_col_num0),1,function(x) read_color(x[2]))
my_colorbar=colorRampPalette(rgb_col)(n = length(rgb_col_num)[1])

ideas_state_matrix_uniq_sort = 0:(length(rgb_col_num)-1)

### order cell types
ct_list = apply(cbind(colnames(state_matrix)),1,function(x) unlist(strsplit(x,'_'))[1] )
ct_list[34:35] = 'T_CD4'
ct_list[36:37] = 'T_CD8'

###### extract counts matrix
counts_matrix = c()
counts_index_matrix = c()
for (i in c(1: dim(state_matrix)[2]) ){
	### extract ith cell type data
	ct_i = ct_list[i]
	ideas_state_matrix_table_tmp = as.matrix(as.numeric(state_matrix[,i]))
	if (sum(ct_list==ct_i)>1){
		index_state_matrix_table_tmp = as.matrix(rowMeans(index_matrix[,ct_list==ct_i]))
	}else{
		index_state_matrix_table_tmp = as.matrix(cbind(index_matrix[,ct_list==ct_i]))
	}
	#ideas_state_matrix_table_tmp = ideas_state_matrix_table_tmp[index_state_matrix_table_tmp!=0]
	ideas_state_matrix_table_tmp = ideas_state_matrix_table_tmp[ideas_state_matrix_table_tmp!=0]
	table_tmp = c()
	for (j in c( 1: length(ideas_state_matrix_uniq_sort)) ){
		### count the number of cREs have jth IDEAS state
		table_tmp[j] = sum(ideas_state_matrix_table_tmp==(j-1))
	}

	### vector to matrix
	counts_matrix = rbind(counts_matrix, table_tmp)

	### extract ith cell type data
	print('atac-pk start')
	ideas_state_matrix_table_tmp = as.matrix(as.numeric(state_matrix[,i]))
	ideas_state_matrix_table_tmp = ideas_state_matrix_table_tmp[index_state_matrix_table_tmp!=0]
	ideas_state_matrix_table_tmp = ideas_state_matrix_table_tmp[ideas_state_matrix_table_tmp!=0]
	table_tmp = c()
	for (j in c( 1: length(ideas_state_matrix_uniq_sort)) ){
		### count the number of cREs have jth IDEAS state
		table_tmp[j] = sum(ideas_state_matrix_table_tmp==(j-1))
	}

	### vector to matrix
	counts_index_matrix = rbind(counts_index_matrix, table_tmp)

}

### transpose matrix
counts_matrix_t = t( counts_matrix )
### add colnames
colnames(counts_matrix_t) = colnames(state_matrix)


### transpose matrix
counts_index_matrix_t = t( counts_index_matrix )
### add colnames
colnames(counts_index_matrix_t) = colnames(state_matrix)
index_matrix_num = apply(index_matrix,2,as.numeric)


celltype_count = c()
for (ct in ct_list){
#celltype_count = c(celltype_count, sum(index_matrix_num[,ct_list==ct]))
celltype_count = c(celltype_count, max(colSums(cbind(index_matrix_num[,ct_list==ct]))))
}

### get cell type mean
counts_index_matrix_ct = c()
counts_matrix_ct = c()
celltype_count_uniq = c()
for (ct in unique(ct_list)){
	#celltype_count_uniq = c(celltype_count_uniq, mean(celltype_count[ct_list==ct]))
	#counts_index_matrix_ct = cbind(counts_index_matrix_ct, colMeans(rbind(counts_index_matrix[ct_list==ct,])))
	#counts_matrix_ct = cbind(counts_matrix_ct, colMeans(rbind(counts_matrix[ct_list==ct,])))
	celltype_count_uniq = c(celltype_count_uniq, mean(celltype_count[ct_list==ct]))
	counts_index_matrix_ct = cbind(counts_index_matrix_ct, apply(rbind(counts_index_matrix[ct_list==ct,]),2,max))
	counts_matrix_ct = cbind(counts_matrix_ct, apply(rbind(counts_matrix[ct_list==ct,]),2,max))
}
colnames(counts_index_matrix_ct) = unique(ct_list)
colnames(counts_matrix_ct) = unique(ct_list)

barplot_lim = 110000
print(counts_index_matrix_t)
### save figure
pdf(paste(cREs_IDEASpro_outfile, '.ideas.celltype.needpk.max.pdf', sep=''), width=7, height=7)
par(mar=c(6.1,4.1,4.1,2.1))
#barplot(counts_index_matrix_t[,order(-colSums(index_matrix_num))], col=my_colorbar, ylim=c(0,barplot_lim),las=2)
barplot(counts_index_matrix_ct[,order(-(celltype_count_uniq))], col=my_colorbar, ylim=c(0,barplot_lim),las=2)
box()
dev.off()

barplot_lim = 200000
pdf(paste(cREs_IDEASpro_outfile, '.ideas.celltype.noneedpk.max.pdf', sep=''), width=14, height=14)
par(mar=c(10.1,4.1,4.1,2.1))
barplot(counts_matrix_ct[,order(-colSums(counts_matrix_ct))], col=my_colorbar, ylim=c(0,barplot_lim),las=2)
box()
dev.off()


state_order = c(2,1,4,3,6,10,11,5,9,13,8,19,12,25,14,23,22,20,21,17,18,7,16,15,24)


barplot_lim = 120000
print(counts_index_matrix_t)
### save figure
pdf(paste(cREs_IDEASpro_outfile, '.ideas.needpk.pdf', sep=''), width=10, height=7)
par(mar=c(10.1,4.1,4.1,2.1))
#barplot(counts_index_matrix_t[,order(-colSums(index_matrix_num))], col=my_colorbar, ylim=c(0,barplot_lim),las=2)
barplot(counts_index_matrix_t[,order(-(celltype_count))], col=my_colorbar, ylim=c(0,barplot_lim),las=2)
box()
dev.off()


### save figure

non0state = colSums(counts_matrix_t)
celltype_count_non0state = c()
for (ct in ct_list){
#celltype_count = c(celltype_count, sum(index_matrix_num[,ct_list==ct]))
celltype_count_non0state = c(celltype_count_non0state, max(non0state[ct_list==ct]))
}


barplot_lim = 200000
pdf(paste(cREs_IDEASpro_outfile, '.ideas.noneedpk.pdf', sep=''), width=10, height=7)
par(mar=c(10.1,4.1,4.1,2.1))
barplot(counts_matrix_t[,order(-celltype_count_non0state)], col=my_colorbar, ylim=c(0,barplot_lim),las=2)
box()
dev.off()
```
