## Supplementary material for "Interspecies regulatory landscapes and elements revealed by novel joint systematic integration of human and mouse blood cell epigenomes": Scripts for IDEAS modeling of epigenetic states jointly in human and mouse: runall_commandline.sh

```
#PBS -l nodes=1:ppn=1
#PBS -l walltime=20:00:00
#PBS -j oe
##PBS -A yzz2_e_g_sc_default
#PBS -A rch8_b_g_bc_default
#PBS -l pmem=20gb

##################################
script_folder='/storage/home/gzx103/scratch/joint_human_mouse/hg38/S3V2_IDEAS_outputs_hg38/S3V2_IDEAS_hg38_r3_withHg38Mm10prior_IDEAS_snapshot/snapshot/bin/'
input_folder='/storage/home/gzx103/scratch/joint_human_mouse/hg38/S3V2_IDEAS_outputs_hg38/S3V2_IDEAS_hg38_r3_withHg38Mm10prior_IDEAS_snapshot/'
output_folder='/storage/home/gzx103/scratch/joint_human_mouse/hg38/S3V2_IDEAS_outputs_hg38/S3V2_IDEAS_hg38_r3_withHg38Mm10prior_IDEAS_snapshot/output'


### run snapshot (CORE!!!)
echo 'run snapshot :o'
cd $input_folder

cp ../../S3V2_IDEAS_outputs_hg38_CCRE/S3V2_IDEAS_hg38_ccre2_IDEAS_output/S3V2_IDEAS_hg38_ccre2.cCRE.M.notall0.bed all_pk.bed

#cp ../../S3V2_IDEAS_outputs_hg38_CCRE/S3V2_IDEAS_hg38_ccre2_IDEAS_output/S3V2_IDEAS_hg38_ccre2.*.cCRE.bed atac_pk/

coln=4
for ctr in $(cat celltype_rep_list.txt)
do
	echo $ctr
	coln=$((coln+1))
	echo $coln
	### get function label
#	cut --delimiter=' ' -f2,3,4,$coln ../S3V2_IDEAS_hg38_r3_withHg38Mm10prior_IDEAS_output/S3V2_IDEAS_hg38_r3_withHg38Mm10prior.state | tail -n+2 | awk -F ' ' -v OFS='\t' '{print $1,$2,$3,".",$4}' > function_label/$ctr.sig.bed
done

#ls atac_pk/* | awk -F '.' -v OFS='\t' '{print $2}' > atac_ctr.list.txt
for ctr in $(cat atac_ctr.list.txt)
do
	echo $ctr
	### get signal
#	paste ../windowsNoBlack.withid.bed ../S3V2_IDEAS_hg38_r1_IDEAS_input_NB/$ctr.ATAC.S3V2.bedgraph.NBP.txt > atac_sig/$ctr.ATAC.S3V2.bedgraph.NBP.bed
done
#paste ../windowsNoBlack.withid.bed ../S3V2_IDEAS_hg38_r1_IDEAS_input_NB/ATAC.average_sig.NBP.txt > atac_sig/AVE.ATAC.S3V2.bedgraph.NBP.bed


### signal list
#ls atac_pk/* | awk -F '.' -v OFS='\t' '{print $2}' > atac_ctr.list.txt
#ls atac_sig/* > signal_list.txt
#paste signal_list.txt atac_ctr.list.txt > signal_list.txt1 && mv signal_list.txt1 signal_list.txt

### function label list
#ls function_label/* > function_list.txt
#paste function_list.txt celltype_rep_list.txt > function_list.txt1 && mv function_list.txt1 function_list.txt

### pk list
#ls atac_pk/* | awk -F '.' -v OFS='\t' '{print $0, $2}' > peak_list.txt


time python $script_folder'snapshot_v.0.4.py' -p peak_list.txt -n snapshot -t 1 -s signal_list.txt -l F -z F -x 0.01 -f function_list.txt -m mostfreq -c function_color_list.txt -e cd_tree.txt -i $input_folder -o $output_folder -b $script_folder -q 1
echo 'complete :)'
```
