## Supplementary material for "Interspecies regulatory landscapes and elements revealed by novel joint systematic integration of human and mouse blood cell epigenomes": Scripts for IDEAS modeling of epigenetic states jointly in human and mouse: get_fdr_p.mm10.R

```
### read signal matrix; row for cCRE; col for feature
d=read.table('cCRE.ATACseq.NBPsig.mat.mm10.txt', header=T)

### exclude bed file part
ds = d[,-c(1:4)]

### convert -log10-p-value to p-value
dsp = 10^(-ds)

### FDR adjustment for all signals
dspfdr = p.adjust(as.vector(t(dsp)), 'fdr')

### convert back to matrix
dspfdrm = (t(matrix(dspfdr, nrow=dim(dsp)[2])))

### add bed file part
ddspfdrm = cbind(d[,1:4], -log10(dspfdrm))

### write output
colnames(ddspfdrm) = colnames(d)
write.table(ddspfdrm, 'cCRE.ATACseq.NBPsig.fdr.mat.mm10.txt', quote=F, col.names=T, row.names=F, sep='\t')
```
