## Supplementary material for "Interspecies regulatory landscapes and elements revealed by novel joint systematic integration of human and mouse blood cell epigenomes": Scripts for IDEAS modeling of epigenetic states jointly in human and mouse: get_snapshot.mm10.error2.sh

```
#PBS -l nodes=1:ppn=1
#PBS -l walltime=20:00:00
#PBS -j oe
##PBS -A yzz2_e_g_sc_default
#PBS -A rch8_b_g_bc_default
#PBS -l pmem=20gb

##################################
script_folder='/storage/home/gzx103/scratch/joint_human_mouse/hg38/S3V2_IDEAS_outputs_hg38/S3V2_IDEAS_hg38_r3_withHg38Mm10prior_IDEAS_snapshot/snapshot/bin/'

working_folder='/storage/home/gzx103/scratch/joint_human_mouse/mm10/S3V2_IDEAS_outputs_mm10/S3V2_IDEAS_mm10_r3_withHg38Mm10prior_IDEAS_snapshot'
### the working folder should be inside the S3V2-IDEAS output directory
### e.g S3V2_IDEAS_outputs_mm10

output_folder=$working_folder'/output1'
S3V2_IDEAS_cCRE_folder='/storage/home/gzx103/scratch/joint_human_mouse/mm10/S3V2_IDEAS_outputs_mm10_CCRE/S3V2_IDEAS_mm10_ccre2_IDEAS_output'
NBP_txt_folder='/storage/home/gzx103/scratch/joint_human_mouse/mm10/S3V2_IDEAS_outputs_mm10/S3V2_IDEAS_mm10_r1_IDEAS_input_NB/'


S3V2_IDEAS_IS_cCRE_file='/storage/home/gzx103/scratch/joint_human_mouse/mm10/S3V2_IDEAS_outputs_mm10_CCRE/S3V2_IDEAS_mm10_ccre2_IDEAS_output/S3V2_IDEAS_mm10_ccre2.cCRE.M.notall0.bed'
S3V2_IDEAS_ES_state_file='/storage/home/gzx103/scratch/joint_human_mouse/mm10/S3V2_IDEAS_outputs_mm10/S3V2_IDEAS_mm10_r3_withHg38Mm10prior_IDEAS_output/S3V2_IDEAS_mm10_r3_withHg38Mm10prior.state'
bin_file_withid='/storage/home/gzx103/scratch/joint_human_mouse/mm10/S3V2_IDEAS_outputs_mm10_CCRE/windowsNoBlack.withid.bed'

IDEAS_cell_type_rep_list=$working_folder'/celltype_rep_list_ideas.txt'

### remove line 179 - 184 in snapshot/bin/snapshot_v.0.4.py
        #if os.path.isfile('all_pk.bed'):
        #       call('rm all_pk.bed', shell=True)
        ### read filenames in peak list
        #for file_info in open(peak_list, 'r'):
        #       filename = file_info.split('\t')[0]
        #       call('cat ' + filename + ' >> all_pk.bed', shell=True)

### run snapshot (CORE!!!)
echo 'run snapshot :o'
mkdir -p $working_folder
cd $working_folder

### get cCRE list
cp $S3V2_IDEAS_IS_cCRE_file all_pk.bed

### copy all cell type peaks
mkdir -p atac_pk
cp $S3V2_IDEAS_cCRE_folder'/'*.*.cCRE.bed atac_pk/

### get ideas state bed files
mkdir -p function_label
coln=4
for ctr in $(cat $IDEAS_cell_type_rep_list)
do
        echo $ctr
        coln=$((coln+1))
        echo $coln
        ### get function label
       cut --delimiter=' ' -f2,3,4,$coln $S3V2_IDEAS_ES_state_file | tail -n+2 | awk -F ' ' -v OFS='\t' '{print $1,$2,$3,".",$4}' > function_label/$ctr.sig.bed
done

### get signal files
mkdir -p atac_sig
ls atac_pk/* | awk -F '.' -v OFS='\t' '{print $2}' > atac_ctr.list.txt
for ctr in $(cat atac_ctr.list.txt)
do
        echo $ctr
        ### get signal
        paste $bin_file_withid $NBP_txt_folder'/'$ctr.ATAC.S3V2.bedgraph.NBP.txt > atac_sig/$ctr.ATAC.S3V2.bedgraph.NBP.bed
done
paste $bin_file_withid $NBP_txt_folder'/'ATAC.average_sig.NBP.txt > atac_sig/AVE.ATAC.S3V2.bedgraph.NBP.bed

###### get snapshot input lists
### signal list
echo 'AVE' > atac_ctr.list.txt
ls atac_pk/*.cCRE.bed | awk -F '.' -v OFS='\t' '{print $2}' >> atac_ctr.list.txt
ls atac_sig/*.S3V2.bedgraph.NBP.bed > signal_list.txt
paste signal_list.txt atac_ctr.list.txt > signal_list.txt1 && mv signal_list.txt1 signal_list.txt

### function label list
ls function_label/*.sig.bed > function_list.txt
sort $IDEAS_cell_type_rep_list > $IDEAS_cell_type_rep_list'.tmp'
paste function_list.txt $IDEAS_cell_type_rep_list'.tmp' > function_list.txt1 && mv function_list.txt1 function_list.txt

### pk list
ls atac_pk/*.cCRE.bed | awk -F '.' -v OFS='\t' '{print $0, $2}' > peak_list.txt

mkdir -p $output_folder
### run snapshot to get IDEAS matrix
time python $script_folder'snapshot_v.0.4.py' -p peak_list.txt -n snapshot -t 1 -s signal_list.txt -l F -z F -x 0.01 -f function_list.txt -m mostfreq -c function_color_list.txt -e cd_tree.txt -i $input_folder -o $output_folder -b $script_folder -q 1
#echo 'complete :)'
```
