## Supplementary material for "Interspecies regulatory landscapes and elements revealed by novel joint systematic integration of human and mouse blood cell epigenomes": Scripts for IDEAS modeling of epigenetic states jointly in human and mouse: get_cross_species.clusters.sh

```
cd /Users/guanjuexiang/Documents/projects/analysis/04_05_2022_coe

R

### get Human data
dh = read.table('coe_analysis/S3V2_IDEAS_hg38_ccre2.cCRE.M.notall0.rmallNEU.withid.coe_mat.PDmerged.clusterID.txt', header=T, sep='\t')
dhs = dh[,-c(1:6)]
rep1h = c(1, 2, 4, 6, 8, 10, 12, 14, 16, 18, 20, 22, 24, 26, 28, 30, 32, 34, 36, 38, 40)
rep2h = c(1, 3, 5, 7, 9, 11, 13, 15, 17, 19, 21, 23, 25, 27, 29, 31, 33, 35, 37, 39, 41)
ct_list_h = c('AVE','B','CD34','CLP','CMP','EOS','ERY','GMP','HSC','HUDEP1','HUDEP2','K562','LMPP','MEP','MK','MON','MONp','MPP','NK','TCD4','TCD8')
### get rep average mat
dhs_ave = c()
for (i in 1:length(rep1h)){
	dhs_ave = cbind(dhs_ave, rowMeans(cbind(dhs[,rep1h[i]], dhs[,rep2h[i]])) )
}
colnames(dhs_ave) = ct_list_h

### get Mouse data
dm = read.table('coe_analysis_mouse/S3V2_IDEAS_mm10_ccre2.cCRE.M.notall0.withid.coe_mat.PDmerged.clusterID.txt', header=T, sep='\t')
dms = dm[,-c(1:6)]
rep1m = c(1, 2, 4, 5, 6, 7, 8, 10, 12, 14, 16, 17, 18, 19, 21, 22, 24, 25, 26, 27, 28, 29, 31)
rep2m = c(1, 3, 4, 5, 6, 7, 9, 11, 13, 15, 16, 17, 18, 20, 21, 23, 24, 25, 26, 27, 28, 30, 32)
ct_list_m = c('AVE','B','CFUE','CFUMK','CLP','CMP','ER4','ERY','ERYfl','G1E','GMP','HPC7','HSC','MEL','MEP','MK','MON','NEU','NK','TCD4','TCD8','iMEL','iMK')
### get rep average mat
dms_ave = c()
for (i in 1:length(rep1m)){
	dms_ave = cbind(dms_ave, rowMeans(cbind(dms[,rep1m[i]], dms[,rep2m[i]])) )
}
colnames(dms_ave) = ct_list_m


### share mat
dhs_ave_shared = dhs_ave[,is.element(ct_list_h, ct_list_m)]
dhs_ave_shared = dhs_ave_shared[,order(colnames(dhs_ave_shared))]
dms_ave_shared = dms_ave[,is.element(ct_list_m, ct_list_h)]
dms_ave_shared = dms_ave_shared[,order(colnames(dms_ave_shared))]

### ids 
id_h = dh[,5]
id_m = dm[,5]

### mean mat
mean_mat_h = matrix(0, nrow=length(unique(id_h)), ncol=dim(dhs_ave_shared)[2])
for (i in unique(id_h)){
	mean_mat_h[i,] = colMeans(dhs_ave_shared[id_h==i,])
}
colnames(mean_mat_h) = colnames(dhs_ave_shared)
rownames(mean_mat_h) = 1:length(unique(id_h))

mean_mat_m = matrix(0, nrow=length(unique(id_m)), ncol=dim(dms_ave_shared)[2])
for (i in unique(id_m)){
	mean_mat_m[i,] = colMeans(dms_ave_shared[id_m==i,])
}
colnames(mean_mat_m) = colnames(dms_ave_shared)
rownames(mean_mat_m) = 1:length(unique(id_m))

### dist mat
cluster_dist = matrix(0, nrow=dim(mean_mat_h)[1], ncol=dim(mean_mat_m)[1])
cluster_cor = matrix(0, nrow=dim(mean_mat_h)[1], ncol=dim(mean_mat_m)[1])
cluster_cosine = matrix(0, nrow=dim(mean_mat_h)[1], ncol=dim(mean_mat_m)[1])
library(lsa)
for (i in 1:dim(mean_mat_h)[1]){
for (j in 1:dim(mean_mat_m)[1]){
cluster_dist[i,j] = sqrt(mean((mean_mat_h[i,]-mean_mat_m[j,])^2))
cluster_cor[i,j] = cor(mean_mat_h[i,], mean_mat_m[j,])
cluster_cosine[i,j] = cosine(mean_mat_h[i,], mean_mat_m[j,])
}}


library(pheatmap)
rownames(cluster_dist) = paste0('H:', 1:dim(cluster_dist)[1])
colnames(cluster_dist) = paste0('M:', 1:dim(cluster_dist)[2])
png('cross_species.dist.heatmap.png')
pheatmap(max(cluster_dist)-cluster_dist, cluster_row=F, cluster_col=F)
dev.off()

rownames(cluster_cor) = paste0('H:', 1:dim(cluster_dist)[1])
colnames(cluster_cor) = paste0('M:', 1:dim(cluster_dist)[2])
png('cross_species.cor.heatmap.png')
pheatmap(cluster_cor, cluster_row=F, cluster_col=F)
dev.off()

my_colorbar=colorRampPalette(c('white', 'red'))(n = 128)
rownames(cluster_cosine) = paste0('H:', 1:dim(cluster_dist)[1])
colnames(cluster_cosine) = paste0('M:', 1:dim(cluster_dist)[2])
png('cross_species.cosine.heatmap.png')
cluster_cosine[cluster_cosine<0] = 0
pheatmap(cluster_cosine, cluster_row=F, cluster_col=F)
dev.off()

my_colorbar=colorRampPalette(c('white', 'red'))(n = 128)
png('human.mean.heatmap.png')
pheatmap(mean_mat_h, cluster_row=F, cluster_col=F, color=my_colorbar)
dev.off()

png('mouse.mean.heatmap.png')
pheatmap(mean_mat_m, cluster_row=F, cluster_col=F, color=my_colorbar)
dev.off()


png('Human_Mouse.mean.heatmap.png', height=2000, width=1200)
rownames(mean_mat_h) = paste0('H:',1:dim(mean_mat_h)[1])
rownames(mean_mat_m) = paste0('M:',1:dim(mean_mat_m)[1])
pheatmap(rbind(mean_mat_h, mean_mat_m), cluster_row=T, cluster_col=T, color=my_colorbar)
dev.off()


set.seed(2019)
HM_dif = c()
k_vec = 5:150
for (k in k_vec){
print(k)
HM_mean_mat_km = kmeans(HM_mean_mat, k)
HM_count_i = c()
for (j in 1:k){
	id = which(HM_mean_mat_km$cluster==j)
	HM_count_i = rbind(HM_count_i, c(sum(id<=100), sum(id>100)))
}
#HM_dif = c(HM_dif, cor(HM_count_i[,1],HM_count_i[,2]))
HM_dif = c(HM_dif, mean( abs(HM_count_i[,1]-HM_count_i[,2]) / (HM_count_i[,1]+HM_count_i[,2])  ) )
}


lmfit = lm(HM_dif[-c(1:100)]~k_vec[-c(1:100)])
pdf('K_vs_cor.pdf')
plot(k_vec, HM_dif-(k_vec*lmfit$coefficients[2]+lmfit$coefficients[1]))
lines(k_vec, HM_dif-(k_vec*lmfit$coefficients[2]+lmfit$coefficients[1]))
abline(h=0, col='red')
abline(v=36, col='blue')
dev.off()

used_k =36

set.seed(2019)
png('Human_Mouse.mean.heatmap.km.png', height=2000, width=1200)
rownames(mean_mat_h) = paste0('H:',1:dim(mean_mat_h)[1])
rownames(mean_mat_m) = paste0('M:',1:dim(mean_mat_m)[1])
HM_mean_mat = rbind(mean_mat_h, mean_mat_m)
HM_mean_mat_km = kmeans(HM_mean_mat, used_k)
hclust_ct = hclust(dist(t(HM_mean_mat)))
pheatmap(HM_mean_mat[order(HM_mean_mat_km$cluster),hclust_ct$order], cluster_row=F, cluster_col=F, color=my_colorbar)
dev.off()


png('Human_Mouse.mean.heatmap.km_id.png', height=2000, width=1200)
km_id_mat = cbind(HM_mean_mat_km$cluster, HM_mean_mat_km$cluster)
colnames(km_id_mat) = c('TCD4', 'TCD4')
library(RColorBrewer)
n = 200#length(unique(new_id))
qual_col_pals = brewer.pal.info[brewer.pal.info$category == 'qual',]
col_vector = c(unlist(mapply(brewer.pal, qual_col_pals$maxcolors, rownames(qual_col_pals))) )
pheatmap(km_id_mat[order(HM_mean_mat_km$cluster),], cluster_row=F, cluster_col=F, color=col_vector)
dev.off()
```
