## Supplementary material for "Interspecies regulatory landscapes and elements revealed by novel joint systematic integration of human and mouse blood cell epigenomes": Scripts for IDEAS modeling of epigenetic states jointly in human and mouse: get_ave.R

```
#input_folder_start = 'coe_score_no'
#target_file_name = 'statep_rna_coe_heatmap.human.all.ccre.withcorfilter.txt'
#output_name = 'statep_rna_coe_heatmap.human.all.ccre.withcorfilter.AVE'
#cts = c("B", "CLP", "CMP", "EOS", "ERY", "GMP", "LSK", "MK", "MONc", "MONp", "MPP", "NEU", "NK", "CD4", "CD8")
args = commandArgs(trailingOnly=TRUE)
input_folder_start = args[1]
target_file_name = args[2]
output_name = args[3]
cts = args[-c(1:3)]
print(cts)

### PCA lm
sig0 = read.table(paste(input_folder_start,cts[1],'/', target_file_name, sep=''), header=T)

for (i in 2:length(cts)){
	sig0_i = read.table(paste(input_folder_start,cts[i],'/', target_file_name, sep=''), header=T)
	sig0 = sig0+sig0_i
}

sig0 = sig0/length(cts)

library(pheatmap)

plot_lim_P = max(abs(sig0[,1]))
plot_lim_D = max(abs(sig0[,2]))
plot_lim_PD = max(abs(sig0))
pdf(paste0(output_name, '.pdf'), width=3)
rank = c(2,1,4,3,6,10,11,5,9,13,8,19,12,25,14,23,22,20,21,17,18,7,16,15,24)
breaksList = seq(-plot_lim_PD, plot_lim_PD, by = 0.001)
my_colorbar=colorRampPalette(c('blue', 'white', 'red'))(n = length(breaksList))
col_breaks = c(seq(0, 2000,length=33))
pheatmap(sig0[rank,], color=my_colorbar, breaks = breaksList, cluster_cols = FALSE,cluster_rows=F, clustering_method = 'average',annotation_names_row=TRUE,annotation_names_col=TRUE,show_rownames=TRUE,show_colnames=TRUE)
dev.off()


write.table(sig0, paste0(output_name, '.txt'), quote=F, col.names=T, row.names=T, sep='\t')
```
