## Supplementary material for "Interspecies regulatory landscapes and elements revealed by novel joint systematic integration of human and mouse blood cell epigenomes": Scripts for IDEAS modeling of epigenetic states jointly in human and mouse: get_ave_cross_species.R

```
cts = c("B", "CLP", "CMP", "EOS", "ERY", "GMP", "LSK", "MK", "MONc", "MONp", "MPP", "NEU", "NK", "CD4", "CD8")
cts_mouse = c("CFUE", "CFUMK", "CMP", "ERYfl", "GMP", "LSK", "MEP", "MONO", "NEU", "iMK", "G1E", "ER4")


### human
sig0 = read.table(paste('coe_score_no',cts[1],'/statep_rna_coe_heatmap.human.all.ccre.withcorfilter.txt', sep=''), header=T)

for (i in 2:length(cts)){
	sig0_i = read.table(paste('coe_score_no',cts[i],'/statep_rna_coe_heatmap.human.all.ccre.withcorfilter.txt', sep=''), header=T)
	sig0 = sig0+sig0_i
}

sig0 = sig0/length(cts)


### mouse
sig0_mouse = read.table(paste('../coe_analysis_mouse/coe_score_no',cts_mouse[1],'/statep_rna_coe_heatmap.mouse.all.ccre.withcorfilter.txt', sep=''), header=T)

for (i in 2:length(cts_mouse)){
	sig0_i = read.table(paste('../coe_analysis_mouse/coe_score_no',cts_mouse[i],'/statep_rna_coe_heatmap.mouse.all.ccre.withcorfilter.txt', sep=''), header=T)
	sig0_mouse = sig0_mouse+sig0_i
}

sig0_mouse = sig0_mouse/length(cts_mouse)

sig0_mouse_min_non0 = min(sig0_mouse[sig0_mouse>0])
sig0_mouse = sig0_mouse-sig0_mouse_min_non0
sig0_mouse = sig0_mouse /max(sig0_mouse) * (max(sig0)-sig0_mouse_min_non0)
sig0_mouse = sig0_mouse+sig0_mouse_min_non0


library(pheatmap)


sig0_ave = (sig0+sig0_mouse)/2
#sig0_ave_direction = ((sig0+sig0_mouse)>=0)*1
#print(sig0_ave_direction)
#sig0_ave_direction[sig0_ave_direction==0] = -1
#print(sig0_ave_direction)
#sig0_ave = sqrt(abs(sig0*sig0_mouse)) * sig0_ave_direction

plot_lim_P = max(abs(sig0_ave[,1]))
plot_lim_D = max(abs(sig0_ave[,2]))
plot_lim_PD = max(abs(sig0_ave))
pdf(paste('statep_rna_coe_heatmap.HM.all.ccre.withcorfilter.AVE.pdf', sep=''), width=3)
rank = c(2,1,4,3,6,10,11,5,9,13,8,19,12,25,14,23,22,20,21,17,18,7,16,15,24)
breaksList = seq(-plot_lim_PD, plot_lim_PD, by = 0.001)
my_colorbar=colorRampPalette(c('blue', 'white', 'red'))(n = length(breaksList))
col_breaks = c(seq(0, 2000,length=33))
pheatmap(sig0_ave[rank,], color=my_colorbar, breaks = breaksList, cluster_cols = FALSE,cluster_rows=F, clustering_method = 'average',annotation_names_row=TRUE,annotation_names_col=TRUE,show_rownames=TRUE,show_colnames=TRUE)
dev.off()

pdf(paste('statep_rna_coe_heatmap.HM.P.all.ccre.withcorfilter.AVE.pdf', sep=''), width=1)
rank = c(2,1,4,3,6,10,11,5,9,13,8,19,12,25,14,23,22,20,21,17,18,7,16,15,24)
breaksList = seq(-plot_lim_P, plot_lim_P, by = 0.001)
my_colorbar=colorRampPalette(c('blue', 'white', 'red'))(n = length(breaksList))
col_breaks = c(seq(0, 2000,length=33))
pheatmap(cbind(sig0_ave[rank,1]), color=my_colorbar, breaks = breaksList, cluster_cols = FALSE,cluster_rows=F, clustering_method = 'average',annotation_names_row=TRUE,annotation_names_col=TRUE,show_rownames=TRUE,show_colnames=TRUE)
dev.off()

pdf(paste('statep_rna_coe_heatmap.HM.D.all.ccre.withcorfilter.AVE.pdf', sep=''), width=1)
rank = c(2,1,4,3,6,10,11,5,9,13,8,19,12,25,14,23,22,20,21,17,18,7,16,15,24)
breaksList = seq(-plot_lim_D, plot_lim_D, by = 0.001)
my_colorbar=colorRampPalette(c('blue', 'white', 'red'))(n = length(breaksList))
col_breaks = c(seq(0, 2000,length=33))
pheatmap(cbind(sig0_ave[rank,2]), color=my_colorbar, breaks = breaksList, cluster_cols = FALSE,cluster_rows=F, clustering_method = 'average',annotation_names_row=TRUE,annotation_names_col=TRUE,show_rownames=TRUE,show_colnames=TRUE)
dev.off()

pdf(paste('statep_rna_coe_heatmap.HM.PDsum.all.ccre.withcorfilter.SUM.pdf', sep=''), width=1)
rank = c(2,1,4,3,6,10,11,5,9,13,8,19,12,25,14,23,22,20,21,17,18,7,16,15,24)
plot_lim_PD_sum = max(abs(sig0_ave[rank,1]/max(sig0_ave[rank,1])*max(sig0_ave[rank,2])+sig0_ave[rank,2]))
breaksList = seq(-plot_lim_PD_sum, plot_lim_PD_sum, by = 0.001)
my_colorbar=colorRampPalette(c('blue', 'white', 'red'))(n = length(breaksList))
col_breaks = c(seq(0, 2000,length=33))
pheatmap(cbind(sig0_ave[rank,1]/max(sig0_ave[rank,1])*max(sig0_ave[rank,2])+sig0_ave[rank,2]), color=my_colorbar, breaks = breaksList, cluster_cols = FALSE,cluster_rows=F, clustering_method = 'average',annotation_names_row=TRUE,annotation_names_col=TRUE,show_rownames=TRUE,show_colnames=TRUE)
dev.off()

sig0_ave[1,] = 0
write.table(sig0_ave, 'statep_rna_coe_heatmap.HM.all.ccre.withcorfilter.AVE.txt', quote=F, col.names=T, row.names=T, sep='\t')

output = cbind(sig0_ave[,1]/max(sig0_ave[rank,1])*max(sig0_ave[rank,2])+sig0_ave[,2])
output[1,1] = 0
rownames(output) = 0:(dim(sig0_ave)[1]-1)
write.table(output, 'statep_rna_coe_heatmap.HM.all.ccre.withcorfilter.SUM.txt', quote=F, col.names=T, row.names=T, sep='\t')


pdf('coe_mouse_vs_human.pdf', width=3.6, height=4)
plot_lim_max = max(c(max(abs(sig0)), max(abs(sig0_mouse))) )
plot_lim_min = min(c(min((sig0)), min((sig0_mouse))) )
plot(as.numeric(as.matrix(sig0)), as.numeric(as.matrix(sig0_mouse)), xlim=c(plot_lim_min,plot_lim_max), ylim=c(plot_lim_min,plot_lim_max), xlab='Human', ylab='Mouse', 
	main = paste('Pearson.Cor = ', round(cor(as.numeric(as.matrix(sig0)), as.numeric(as.matrix(sig0_mouse))), 3), sep=''))
abline(0,1, col='red')
dev.off()

### get coe difference
coe_SUM = output
coe_SUM_dist_mat = c()

for (i in 1:length(coe_SUM)){
	coe_SUM_dist_row = c()
for (j in 1:length(coe_SUM)){
	coe_SUM_dist_row = c(coe_SUM_dist_row, coe_SUM[i] - coe_SUM[j])
}
coe_SUM_dist_mat = rbind(coe_SUM_dist_mat, coe_SUM_dist_row)
}
colnames(coe_SUM_dist_mat) = 0:24
rownames(coe_SUM_dist_mat) = 0:24

pdf(paste('statep_rna_coe_heatmap.HM.PDsum.all.ccre.withcorfilter.SUM.difference.pdf', sep=''))
rank = c(2,1,4,3,6,10,11,5,9,13,8,19,12,25,14,23,22,20,21,17,18,7,16,15,24)
plot_lim_PD_sum = max(abs(coe_SUM_dist_mat))
breaksList = seq(-plot_lim_PD_sum, plot_lim_PD_sum, by = 0.001)
my_colorbar=colorRampPalette(c('blue', 'white', 'red'))(n = length(breaksList))
col_breaks = c(seq(0, 2000,length=33))
pheatmap(coe_SUM_dist_mat[rank,rank], color=my_colorbar, breaks = breaksList, cluster_cols = F,cluster_rows=F, clustering_method = 'average',annotation_names_row=TRUE,annotation_names_col=TRUE,show_rownames=TRUE,show_colnames=TRUE)
dev.off()
```
