## Supplementary material for "Interspecies regulatory landscapes and elements revealed by novel joint systematic integration of human and mouse blood cell epigenomes": Scripts for IDEAS modeling of epigenetic states jointly in human and mouse: get_coe_pcareg.human.withccre_withcorfilter.local.iter.25JES_PD.R

```
#R
args = commandArgs(trailingOnly=TRUE)
library(data.table)
library(LSD)
library(pheatmap)
library(pastecs)
library(lmvar)
library(mclust)

R2 = function(obs, pred){
	r2 = 1-mean((obs-pred)^2)/mean((obs-mean(obs))^2)
	return(r2)
}

R2_0 = function(obs, pred){
	r2 = 1-mean((obs-pred)^2)/mean((obs-0)^2)
	return(r2)
}

quantile_norm = function(x){
	xm = rowMeans(x)
	refsig_sort = xm[order(xm)]
	for (i in 1:dim(x)[2]){
		sigtmp = x[,i]
		x[,i] = refsig_sort[rank(sigtmp)]
	}
	return(x)
}

setwd('/Users/guanjuexiang/Downloads/coe_analysis/')

#sp_list = c('B', 'B', 'CD34', 'CD34', 'CD4', 'CD4', 'CD8', 'CD8', 'CLP', 'CLP', 'CMP', 'CMP', 'EOS', 'ERY', 'ERY', 'GMP', 'GMP', 'H1', 'H1', 'H2', 'H2', 'LMPP', 'LMPP', 'LSK', 'LSK', 'MK', 'MK', 'MONc', 'MONc', 'MONp', 'MONp', 'MPP', 'MPP', 'NEU', 'NEU', 'NK', 'NK', 'PBMC', 'PBMC')
rna_list = c('LSK', 'LSK', 'ERY', 'ERY', 'CD4', 'CD4', 'CD8', 'CD8', 'B', 'B', 'CMP', 'CMP', 'MONp', 'MONp', 'NEU', 'NEU', 'MONc', 'MONc', 'GMP', 'GMP', 'CFUE', 'NK', 'NK', 'MK', 'MK', 'CLP', 'MPP', 'MPP', 'EOS', 'EOS', 'MEP', 'MEP', 'MK', 'MK', 'CLP', 'ERY', 'ERY', 'ERY', 'HUDEP1', 'HUDEP1', 'HUDEP2', 'HUDEP2', 'CD34', 'CD34')
sp_list = c('AVE', 'B', 'B', 'CD34', 'CD34', 'CLP', 'CLP', 'CMP', 'CMP', 'EOS', 'EOS', 'ERY', 'ERY', 'GMP', 'GMP', 'HSC', 'HSC', 'HUDEP1', 'HUDEP1', 'HUDEP2', 'HUDEP2', 'K562', 'K562', 'LMPP', 'LMPP', 'MEP', 'MEP', 'MK', 'MK', 'MONc', 'MONc', 'MONp', 'MONp', 'MPP', 'MPP', 'NEU', 'NEU', 'NK', 'NK', 'CD4', 'CD4', 'CD8', 'CD8')

#sp_list = c('B_1', 'B2_2', 'CD34_1', 'CD34_2', 'CD4_1', 'CD4_2', 'CD8_1', 'CD8_2', 'CLP_1', 'CLP_2', 'CMP_1', 'CMP_2', 'EOS_1', 'ERY_1', 'ERY_2', 'GMP_1', 'GMP_2', 'H1_1', 'H1_2', 'H2_1', 'H2_2', 'LMPP_1', 'LMPP_2', 'LSK_1', 'LSK_2', 'MK_1', 'MK_2', 'MONc_1', 'MONc_2', 'MONp_1', 'MONp_2', 'MPP_1', 'MPP_2', 'NEU_1', 'NEU_2', 'NK_1', 'NK_2', 'PBMC_1', 'PBMC_2')
#rna_list = c('LSK_1', 'LSK_2', 'ERY_1', 'ERY_2', 'CD4_1', 'CD4_2', 'CD8_1', 'CD8_2', 'B_1', 'B_2', 'CMP_1', 'CMP_2', 'MONp_1', 'MONp_2', 'NEU_1', 'NEU_2', 'MONc_1', 'MONc_2', 'GMP_1', 'GMP_2', 'CFUE_1', 'NK_1', 'NK_2', 'MK_1', 'MK_2', 'CLP_1', 'CLP_2', 'MPP_1', 'MPP_2', 'EOS_1', 'EOS_2')

common_ct = intersect(sp_list, rna_list)
no_used_ct = c('HUDEP1','HUDEP2','CD34')
common_ct = common_ct[!is.element(common_ct, no_used_ct)]
#common_ct = common_ct[common_ct!='NEU']

leave_out_ct=args[1]
common_ct = common_ct[common_ct!=leave_out_ct]
smallnum = 1e-1
smallnum_sp =1
cor_thresh = 0.2
pcv_var_used = 0.95
state_n = 25
plot_lim_all = c(-2.5,5)
set.seed(2019)

output_folder=paste('coe_score_no', leave_out_ct, sep='')
dir.create(output_folder)
### get RNA
#d = read.table('/gpfs/group/yzz2/default/legacy/group/projects/vision_joint_human_mouse/coefficients_human_with_cCRE_with_corfilter/HumanVISION_RNAseq_hg38_genes_tpm.idsort.protein_coding.txt', header=F)
d = read.table('HumanVISION_RNAseq_hg38_genes_tpm.idsort.protein_coding.txt', header=F)

ds = d[,-c(1:6)]

### QTnorm
ds_qt = quantile_norm(ds)

### split gene groups
dms = rowMeans(ds_qt)
dms_log = log(dms+smallnum)
mod = densityMclust(dms_log,G=2)
gmm_thresh = max(dms_log[mod$classification==1])
rna_l = mod$classification==1
rna_h = mod$classification==2
pdf(paste(output_folder, '/dms_log_rna.hist.pdf',sep=''), width=5, height=5)
plot(mod, what = "density", data = dms_log, breaks = 30)
abline(v=gmm_thresh, col='red', lwd=2, lty=2)
dev.off()

### get gene mat
ds_qt_all = c()

for (ct in common_ct){
ct_pos_rna = which(rna_list==ct)
if (length(ct_pos_rna)>1){
rna_mat_tmp = rowMeans(ds_qt[,ct_pos_rna])
}else{rna_mat_tmp = ds_qt[,ct_pos_rna]}
ds_qt_all = c(ds_qt_all, rna_mat_tmp)
}

ds_qt_all_log = log(ds_qt_all+smallnum)

### get state coverage
sp_all = c()
for (i in c(0:(state_n-1))){
ss = read.table(paste('HumanVISION_RNAseq_hg38_gene.idsort.protein_coding.Nkbupdownexp.S',i,'.bed', sep=''), header=F)
sp = (ss[,-c(1:3)])
sp_i = c()
for (ct in common_ct){
ct_pos_sp = which(sp_list==ct)
### average replicates
if (length(ct_pos_sp)>1){
sp_mat_tmp = rowMeans(sp[,ct_pos_sp])
}else{
sp_mat_tmp = sp[,ct_pos_sp]
}
sp_i = c(sp_i, sp_mat_tmp)
}
print(i)
sp_all = cbind(sp_all, sp_i)
}


sp_all_dist = c()
for (i in c(0:(state_n-1))){
ss = read.table(paste('HumanVISION_RNAseq_hg38_gene.idsort.protein_coding.NHkbupdownexp.S',i,'.bed', sep=''), header=F)
sp = (ss[,-c(1:3)])
sp_i = c()
for (ct in common_ct){
ct_pos_sp = which(sp_list==ct)
### average replicates
if (length(ct_pos_sp)>1){
sp_mat_tmp = rowMeans(sp[,ct_pos_sp])
}else{
sp_mat_tmp = sp[,ct_pos_sp]
}
sp_i = c(sp_i, sp_mat_tmp)
}
print(i)
sp_all_dist = cbind(sp_all_dist, sp_i)
}


colnames(sp_all) = c(0:(state_n-1))
sp_all_log = log((sp_all+smallnum_sp))

colnames(sp_all_dist) = c(0:(state_n-1))
sp_all_dist_log = log(sp_all_dist+smallnum_sp)

### prepare pca
adj_mean_od = mean(sp_all_dist_log)
adj_mean_tar = mean(sp_all_log)
adj_sd_od = sd(sp_all_dist_log)
adj_sd_tar = sd(sp_all_log)
global_scale = 1/mean(sp_all_dist_log)*mean(sp_all_log)
#sp_all_PD_log = cbind(sp_all_log, sp_all_dist_log*global_scale)
sp_all_PD_log = cbind(sp_all_log, (sp_all_dist_log - mean(adj_mean_od)) / adj_sd_od * adj_sd_tar + adj_mean_tar )

### match high low gene means
dmslog_noMean = ds_qt_all_log# - mean(ds_qt_all_log)


### correlation heatmap
sp_mat_for_corheatmap = sp_all_PD_log[,-c(1,(state_n+1))]
colnames(sp_mat_for_corheatmap) = c(paste('P:', 1:(state_n-1)), paste('D:', 1:(state_n-1)))
cor_mat_for_corheatmap = cor(sp_mat_for_corheatmap)

pdf(paste(output_folder, '/cor_mat_for_corheatmap.pdf', sep=''), width=7.5,height=7)
breaksList = seq(-1, 1, by = 0.001)
my_colorbar=colorRampPalette(c('blue', 'white', 'red'))(n = length(breaksList))
col_breaks = c(seq(0, 2000,length=33))
pheatmap(cor_mat_for_corheatmap, color=my_colorbar, breaks = breaksList, cluster_cols = T,cluster_rows=T, clustering_method = 'average',annotation_names_row=TRUE,annotation_names_col=TRUE,show_rownames=TRUE,show_colnames=TRUE)
dev.off()

### raw linear model
used_idtry = dmslog_noMean>(-1000000)
lm_raw_all = lm(dmslog_noMean[used_idtry]~sp_all_PD_log[,-c(1,(state_n+1))])
Bpca_raw_all = lm_raw_all$coefficients[-1]
eRP_mat_human_raw = cbind(Bpca_raw_all[1:(state_n-1)],Bpca_raw_all[(state_n):((state_n-1)*2)])
eRP_mat_human_raw = rbind(c(0,0), eRP_mat_human_raw)
colnames(eRP_mat_human_raw) = c('P','D')
rownames(eRP_mat_human_raw) = 0:(state_n-1)

plot_lim_P = max(abs(eRP_mat_human_raw[,1]))
plot_lim_D = max(abs(eRP_mat_human_raw[,2]))
plot_lim_PD = max(abs(eRP_mat_human_raw))
pdf(paste(output_folder, '/statep_rna_coe_heatmap.human.all.PD.withccre.rawlinear.pdf', sep=''), width=3)
rank = c(2,1,4,3,6,10,11,5,9,13,8,19,12,25,14,23,22,20,21,17,18,7,16,15,24)
breaksList = seq(-plot_lim_PD, plot_lim_PD, by = 0.001)
my_colorbar=colorRampPalette(c('blue', 'white', 'red'))(n = length(breaksList))
col_breaks = c(seq(0, 2000,length=33))
pheatmap(eRP_mat_human_raw[rank,], color=my_colorbar, breaks = breaksList, cluster_cols = FALSE,cluster_rows=F, clustering_method = 'average',annotation_names_row=TRUE,annotation_names_col=TRUE,show_rownames=TRUE,show_colnames=TRUE)
dev.off()
pdf(paste(output_folder, '/statep_rna_coe_heatmap.human.all.P.withccre.rawlinear.pdf', sep=''), width=3)
breaksList = seq(-plot_lim_P, plot_lim_P, by = 0.001)
my_colorbar=colorRampPalette(c('blue', 'white', 'red'))(n = length(breaksList))
col_breaks = c(seq(0, 2000,length=33))
pheatmap(cbind(eRP_mat_human_raw[rank,1],eRP_mat_human_raw[rank,1]), color=my_colorbar, breaks = breaksList, cluster_cols = FALSE,cluster_rows=F, clustering_method = 'average',annotation_names_row=TRUE,annotation_names_col=TRUE,show_rownames=TRUE,show_colnames=TRUE)
dev.off()
pdf(paste(output_folder, '/statep_rna_coe_heatmap.human.all.D.withccre.rawlinear.pdf', sep=''), width=3)
breaksList = seq(-plot_lim_D, plot_lim_D, by = 0.001)
my_colorbar=colorRampPalette(c('blue', 'white', 'red'))(n = length(breaksList))
col_breaks = c(seq(0, 2000,length=33))
pheatmap(cbind(eRP_mat_human_raw[rank,2],eRP_mat_human_raw[rank,2]), color=my_colorbar, breaks = breaksList, cluster_cols = FALSE,cluster_rows=F, clustering_method = 'average',annotation_names_row=TRUE,annotation_names_col=TRUE,show_rownames=TRUE,show_colnames=TRUE)
dev.off()


### dimension reducation
pca_all <- prcomp(sp_all_PD_log[,-c(1,(state_n+1))], center = F,scale. = F)
pca0_rotation = pca_all$rotation 
exp_var = summary(pca_all)$importance[2,]
exp_var_sum = 0
k = 0
for (varexp_tmp in exp_var){
k = k+1
exp_var_sum = exp_var_sum+varexp_tmp
if (exp_var_sum>pcv_var_used){
pcn = k
break
}
}
### PCA lm
used_idtry = dmslog_noMean>(-1000000)


lm_all = lm(dmslog_noMean[used_idtry]~pca_all$x[used_idtry,c(1:pcn)])
Bpca_all = pca_all$rotation[,c(1:pcn)] %*% lm_all$coefficients[-1]

alpha = lm_all$coefficients[1]

pred = lm_all$fitted.value #+ mean(c(ds_qt_all_log_l, ds_qt_all_log_h))
#pca_all$x[,1:pcn] %*% lm_all$coefficients + mean(c(ds_qt_all_log_l, ds_qt_all_log_h))
R2(dmslog_noMean[used_idtry], pred)
0.5628416

eRP_mat_human = cbind(Bpca_all[1:(state_n-1)],Bpca_all[(state_n):((state_n-1)*2)])
eRP_mat_human = rbind(c(0,0), eRP_mat_human)
colnames(eRP_mat_human) = c('P','D')
rownames(eRP_mat_human) = 0:(state_n-1)
write.table(eRP_mat_human, paste(output_folder, '/statep_rna_coe_heatmap.human.all.txt', sep=''), quote=F, col.names=T, row.names=T, sep='\t')

set.seed(2019)
used_id = sample(length(pred), 50000)
plot_lim = plot_lim_all#c(min(c(dmslog_noMean+mean(ds_qt_all_log), pred)), max(c(dmslog_noMean+mean(ds_qt_all_log), pred)))

png(paste(output_folder, '/pred_obs.tpm1.png', sep=''))
#heatscatter(as.vector(pred)[used_id]+mean(ds_qt_all_log), as.vector(dmslog_noMean)[used_id]+mean(ds_qt_all_log), xlim=plot_lim, ylim=plot_lim)
heatscatter(as.vector(pred)[used_id], as.vector(dmslog_noMean)[used_id], xlim=plot_lim, ylim=plot_lim, main=R2(dmslog_noMean[used_idtry], pred))
abline(0,1, col='red', lwd=2)
abline(h=0, col='black', lwd=2)
abline(v=0, col='black', lwd=2)
dev.off()

plot_lim_P = max(abs(eRP_mat_human[,1]))
plot_lim_D = max(abs(eRP_mat_human[,2]))
plot_lim_PD = max(abs(eRP_mat_human))
pdf(paste(output_folder, '/statep_rna_coe_heatmap.human.all.PD.withccre.pdf', sep=''), width=3)
plot_lim_PD_max = max(c(abs(plot_lim_P), abs(plot_lim_D)))
breaksList = seq(-plot_lim_PD, plot_lim_PD, by = 0.001)
my_colorbar=colorRampPalette(c('blue', 'white', 'red'))(n = length(breaksList))
col_breaks = c(seq(0, 2000,length=33))
pheatmap((eRP_mat_human[rank,]), color=my_colorbar, breaks = breaksList, cluster_cols = FALSE,cluster_rows=F, clustering_method = 'average',annotation_names_row=TRUE,annotation_names_col=TRUE,show_rownames=TRUE,show_colnames=TRUE)
dev.off()
pdf(paste(output_folder, '/statep_rna_coe_heatmap.human.all.P.withccre.pdf', sep=''), width=3)
plot_lim_PD_max = max(c(abs(plot_lim_P), abs(plot_lim_D)))
breaksList = seq(-plot_lim_P, plot_lim_P, by = 0.001)
my_colorbar=colorRampPalette(c('blue', 'white', 'red'))(n = length(breaksList))
col_breaks = c(seq(0, 2000,length=33))
pheatmap(cbind(eRP_mat_human[rank,1],eRP_mat_human[rank,1]), color=my_colorbar, breaks = breaksList, cluster_cols = FALSE,cluster_rows=F, clustering_method = 'average',annotation_names_row=TRUE,annotation_names_col=TRUE,show_rownames=TRUE,show_colnames=TRUE)
dev.off()
pdf(paste(output_folder, '/statep_rna_coe_heatmap.human.all.D.withccre.pdf', sep=''), width=3)
breaksList = seq(-plot_lim_D, plot_lim_D, by = 0.001)
my_colorbar=colorRampPalette(c('blue', 'white', 'red'))(n = length(breaksList))
col_breaks = c(seq(0, 2000,length=33))
pheatmap(cbind(eRP_mat_human[rank,2],eRP_mat_human[rank,2]), color=my_colorbar, breaks = breaksList, cluster_cols = FALSE,cluster_rows=F, clustering_method = 'average',annotation_names_row=TRUE,annotation_names_col=TRUE,show_rownames=TRUE,show_colnames=TRUE)
dev.off()


### filter by cor
### get state coe predict mat

#for (iter_i in 1:1){
iter_i = 1
### get cCRE eRP score across celltypes
ss00 = read.table(paste('S3V2_IDEAS_hg38_ccre2.cCRE.M.withid.S0.mat.txt', sep=''), header=F)
sp_sig_pred_ct = matrix(0, nrow=dim(ss00)[1], ncol=dim(ss00)[2])
for (i in c(0:(state_n-1))){
print(i)
ss = read.table(paste('S3V2_IDEAS_hg38_ccre2.cCRE.M.withid.S',i,'.mat.txt', sep=''), header=F)
sp = (ss[,-c(1:4)])
#sp_sig = sp*eRP_mat_human[i+1,2]
sp_sig = log(sp+smallnum_sp)*eRP_mat_human[i+1,2]
sp_sig_pred_ct = sp_sig_pred_ct + sp_sig
}

### average the replicates of each celltype
sp_sig_pred_ct_common_ct = c()
for (ct in common_ct){
ct_pos_sp = which(sp_list==ct)
### average replicates
if (length(ct_pos_sp)>1){
sp_mat_tmp = rowMeans(sp_sig_pred_ct[,ct_pos_sp])
}else{
sp_mat_tmp = sp_sig_pred_ct[,ct_pos_sp]
}
sp_sig_pred_ct_common_ct = cbind(sp_sig_pred_ct_common_ct, sp_mat_tmp)
}

### get gene mat
ds_qt_all_mat = c()

for (ct in common_ct){
ct_pos_rna = which(rna_list==ct)
if (length(ct_pos_rna)>1){
rna_mat_tmp = rowMeans(ds_qt[,ct_pos_rna])
}else{rna_mat_tmp = ds_qt[,ct_pos_rna]}
ds_qt_all_mat = cbind(ds_qt_all_mat, rna_mat_tmp)
}

ds_qt_all_mat_log = log(ds_qt_all_mat+smallnum)


### read gene with used id
gene_withccre_id = read.table('HumanVISION_RNAseq_hg38_gene.idsort.protein_coding.NHkbupdownexp.withccreid.bed', header=F)[,5]
gene_withccre_id_str = as.character(gene_withccre_id)

cor_vec_all_select = matrix('.', nrow=length(gene_withccre_id), ncol=1)
cor_vec = c()

for (i in 1:length(gene_withccre_id)){
#for (i in 1:6){
if (i%%1000==0){
	print(i)
}
ds_qt_all_mat_log_i = ds_qt_all_mat_log[i,]
state_id = as.numeric(unlist(strsplit(gene_withccre_id_str[i], ',')))
sp_sig_pred_ct_common_ct_i = sp_sig_pred_ct_common_ct[state_id,]
if (length(state_id)>1){
cor_vec_i = apply(sp_sig_pred_ct_common_ct_i, 1, function(x) cor(x, ds_qt_all_mat_log_i))
} else if (length(state_id)==1){
cor_vec_i = cor(sp_sig_pred_ct_common_ct_i, ds_qt_all_mat_log_i)
}
cor_vec = c(cor_vec, cor_vec_i)
cor_vec_i_select = state_id[(abs(cor_vec_i)>=cor_thresh) & (!is.na(cor_vec_i))]
cor_vec_i_select_str = paste(cor_vec_i_select, collapse=',')
cor_vec_all_select[i,1] = cor_vec_i_select_str
}

print('ccre gene correlation:')
cor_vec = as.matrix(cor_vec)
cor_vec = cor_vec[!is.na(cor_vec)]
print('mean:')
print(mean(cor_vec))
print('sd:')
print(sd(cor_vec))


pdf(paste(output_folder, '/ccre_cor_hist.',iter_i,'.pdf', sep=''))
hist(cor_vec, breaks=100, xlim=c(-1,1))
zcor = (cor_vec-mean(cor_vec))/sd(cor_vec)
zcorp = pnorm(zcor, mean=0, sd=1, lower.tail=F)
lim1 = min(cor_vec[zcorp<0.025])
lim2 = max(cor_vec[zcorp>0.975])
#abline(v=lim1, lwd=2)
#abline(v=lim2, lwd=2)
abline(v=0, lwd=2)
abline(v=mean(cor_vec), lwd=2, lty=2, col='red')
box()
dev.off()

### redo sp coverage after cor filtering
ss = read.table(paste('S3V2_IDEAS_hg38_ccre2.cCRE.M.withid.S0.mat.txt', sep=''), header=F)
ss_all = (ss[,-c(1:4)])
for (i in c(1:(state_n-1))){
print(i)
ss = read.table(paste('S3V2_IDEAS_hg38_ccre2.cCRE.M.withid.S',i,'.mat.txt', sep=''), header=F)
sp = (ss[,-c(1:4)])
ss_all = cbind(ss_all, sp)
}


sp_corfiltered = matrix(0, nrow=dim(cor_vec_all_select)[1], ncol=dim(ss_all)[2])
for (i in 1:dim(cor_vec_all_select)[1]){
if (i%%100==0){
	print(i)
}
used_row_i = as.numeric(unlist(strsplit(cor_vec_all_select[i,1], ',')))
if (length(used_row_i)>0){
sp_corfiltered[i,] = colSums(ss_all[used_row_i,])
}else{
sp_corfiltered[i,] = rep(0, dim(sp_corfiltered)[2])
}
}


sp_corfiltered_all = matrix(0, nrow=dim(sp_corfiltered)[1]*length(common_ct), ncol=(state_n))
k=0
for (ct in common_ct){
print(ct)
ct_pos_sp = which(sp_list==ct)
sp_mat_tmp = matrix(0, nrow=dim(sp_corfiltered)[1], ncol=state_n )
for (i in ct_pos_sp){
#sp_mat_tmp = sp_mat_tmp+sp_corfiltered[,seq(1,39*24, by=39)+i-1]
sp_mat_tmp = sp_mat_tmp+sp_corfiltered[,seq(1,dim(sp_corfiltered)[2], by=dim(sp_corfiltered)[2]/state_n)+i-1]
}
sp_mat_tmp = sp_mat_tmp/length(ct_pos_sp)
sp_corfiltered_all[seq(1,dim(sp_corfiltered)[1])+(k)*dim(sp_corfiltered)[1],] = sp_mat_tmp
k = k+1
}

colnames(sp_corfiltered_all) = c(0:(state_n-1))
sp_corfiltered_all_D_log = log((sp_corfiltered_all+smallnum_sp))
#sp_all_PD_corfilter_log = cbind(sp_all_log, sp_corfiltered_all_D_log*global_scale)
sp_all_PD_corfilter_log = cbind(sp_all_log, (sp_corfiltered_all_D_log - mean(adj_mean_od)) / adj_sd_od * adj_sd_tar + adj_mean_tar )


### dimension reducation rm 0 state info -c(1,25)
### PCA lm
used_idtry = dmslog_noMean>(-1000000)
sp_all_PD_corfilter_log_new = sp_all_PD_corfilter_log[,-c(1,state_n)] %*% pca0_rotation

lm_all = lm(dmslog_noMean[used_idtry]~sp_all_PD_corfilter_log_new[used_idtry,c(1:pcn)])
Bpca_all = pca0_rotation[,c(1:pcn)] %*% lm_all$coefficients[-1]

pred = lm_all$fitted.value #+ mean(c(ds_qt_all_log_l, ds_qt_all_log_h))
#pca_all$x[,1:pcn] %*% lm_all$coefficients + mean(c(ds_qt_all_log_l, ds_qt_all_log_h))
R2(dmslog_noMean[used_idtry], pred)
0.564377


set.seed(2019)
used_id = sample(length(pred), 50000)
plot_lim = plot_lim_all#c(min(c(dmslog_noMean+mean(ds_qt_all_log), pred)), max(c(dmslog_noMean+mean(ds_qt_all_log), pred)))

png(paste(output_folder, '/pred_obs.tpm.',iter_i,'.png', sep=''))
#heatscatter(as.vector(pred)[used_id]+mean(ds_qt_all_log), as.vector(dmslog_noMean)[used_id]+mean(ds_qt_all_log), xlim=plot_lim, ylim=plot_lim)
heatscatter(as.vector(pred)[used_id], as.vector(dmslog_noMean)[used_id], xlim=plot_lim, ylim=plot_lim, main=R2(dmslog_noMean[used_idtry], pred))
abline(0,1, col='red', lwd=2)
abline(h=0, col='black', lwd=2)
abline(v=0, col='black', lwd=2)
dev.off()


eRP_mat_human = cbind(Bpca_all[1:(state_n-1)],Bpca_all[(state_n):((state_n-1)*2)])
eRP_mat_human = rbind(c(0,0), eRP_mat_human)
colnames(eRP_mat_human) = c('P','D')
rownames(eRP_mat_human) = 0:(state_n-1)
write.table(eRP_mat_human, paste(output_folder, '/statep_rna_coe_heatmap.human.all.ccre.withcorfilter.',iter_i,'.txt', sep=''), quote=F, col.names=T, row.names=T, sep='\t')


plot_lim_P = max(abs(eRP_mat_human[,1]))
plot_lim_D = max(abs(eRP_mat_human[,2]))
plot_lim_PD = max(abs(eRP_mat_human))

pdf(paste(output_folder, '/statep_rna_coe_heatmap.human.all.PD.ccre.withcorfilter.',iter_i,'.pdf', sep=''), width=3)
#rank = c(9,2,3,1,7,4,15,14,13,5,11,22,6,18,8,24,12,10,20,16,23,17,19,21)
breaksList = seq(-plot_lim_PD, plot_lim_PD, by = 0.001)
my_colorbar=colorRampPalette(c('blue', 'white', 'red'))(n = length(breaksList))
col_breaks = c(seq(0, 2000,length=33))
pheatmap((eRP_mat_human[rank,]), color=my_colorbar, breaks = breaksList, cluster_cols = FALSE,cluster_rows=F, clustering_method = 'average',annotation_names_row=TRUE,annotation_names_col=TRUE,show_rownames=TRUE,show_colnames=TRUE)
dev.off()
pdf(paste(output_folder, '/statep_rna_coe_heatmap.human.all.P.ccre.withcorfilter.',iter_i,'.pdf', sep=''), width=3)
#rank = c(9,2,3,1,7,4,15,14,13,5,11,22,6,18,8,24,12,10,20,16,23,17,19,21)
breaksList = seq(-plot_lim_P, plot_lim_P, by = 0.001)
my_colorbar=colorRampPalette(c('blue', 'white', 'red'))(n = length(breaksList))
col_breaks = c(seq(0, 2000,length=33))
pheatmap(cbind(eRP_mat_human[rank,1],eRP_mat_human[rank,1]), color=my_colorbar, breaks = breaksList, cluster_cols = FALSE,cluster_rows=F, clustering_method = 'average',annotation_names_row=TRUE,annotation_names_col=TRUE,show_rownames=TRUE,show_colnames=TRUE)
dev.off()
pdf(paste(output_folder, '/statep_rna_coe_heatmap.human.all.D.ccre.withcorfilter.',iter_i,'.pdf', sep=''), width=3)
#rank = c(9,2,3,1,7,4,15,14,13,5,11,22,6,18,8,24,12,10,20,16,23,17,19,21)
breaksList = seq(-plot_lim_D, plot_lim_D, by = 0.001)
my_colorbar=colorRampPalette(c('blue', 'white', 'red'))(n = length(breaksList))
col_breaks = c(seq(0, 2000,length=33))
pheatmap(cbind(eRP_mat_human[rank,2],eRP_mat_human[rank,2]), color=my_colorbar, breaks = breaksList, cluster_cols = FALSE,cluster_rows=F, clustering_method = 'average',annotation_names_row=TRUE,annotation_names_col=TRUE,show_rownames=TRUE,show_colnames=TRUE)
dev.off()

#}


#############################
#############################
############################# Round2
### filter by cor
### get state coe predict mat
sp_sig_pred_ct = matrix(0, nrow=dim(ss00)[1], ncol=dim(ss00)[2])
for (i in c(0:(state_n-1))){
print(i)
ss = read.table(paste('S3V2_IDEAS_hg38_ccre2.cCRE.M.withid.S',i,'.mat.txt', sep=''), header=F)
sp = (ss[,-c(1:4)])
#sp_sig = sp*eRP_mat_human[i+1,2]
sp_sig = log(sp+smallnum_sp)*eRP_mat_human[i+1,2]
sp_sig_pred_ct = sp_sig_pred_ct + sp_sig
}

sp_sig_pred_ct_common_ct = c()
for (ct in common_ct){
ct_pos_sp = which(sp_list==ct)
### average replicates
if (length(ct_pos_sp)>1){
sp_mat_tmp = rowMeans(sp_sig_pred_ct[,ct_pos_sp])
}else{
sp_mat_tmp = sp_sig_pred_ct[,ct_pos_sp]
}
sp_sig_pred_ct_common_ct = cbind(sp_sig_pred_ct_common_ct, sp_mat_tmp)
}

### get gene mat
ds_qt_all_mat = c()
for (ct in common_ct){
ct_pos_rna = which(rna_list==ct)
if (length(ct_pos_rna)>1){
rna_mat_tmp = rowMeans(ds_qt[,ct_pos_rna])
}else{rna_mat_tmp = ds_qt[,ct_pos_rna]}
ds_qt_all_mat = cbind(ds_qt_all_mat, rna_mat_tmp)
}

ds_qt_all_mat_log = log(ds_qt_all_mat+smallnum)


### read gene with used id
gene_withccre_id = read.table('HumanVISION_RNAseq_hg38_gene.idsort.protein_coding.NHkbupdownexp.withccreid.bed', header=F)[,5]
gene_withccre_id_str = as.character(gene_withccre_id)

cor_vec_all_select = matrix('.', nrow=length(gene_withccre_id), ncol=1)

for (i in 1:length(gene_withccre_id)){
#for (i in 1:6){
if (i%%1000==0){
	print(i)
}
ds_qt_all_mat_log_i = ds_qt_all_mat_log[i,]
state_id = as.numeric(unlist(strsplit(gene_withccre_id_str[i], ',')))
sp_sig_pred_ct_common_ct_i = sp_sig_pred_ct_common_ct[state_id,]
if (length(state_id)>1){
cor_vec_i = apply(sp_sig_pred_ct_common_ct_i, 1, function(x) cor(x, ds_qt_all_mat_log_i))
} else if (length(state_id)==1){
cor_vec_i = cor(sp_sig_pred_ct_common_ct_i, ds_qt_all_mat_log_i)
}
cor_vec_i_select = state_id[(abs(cor_vec_i)>=cor_thresh) & (!is.na(cor_vec_i))]
cor_vec_i_select_str = paste(cor_vec_i_select, collapse=',')
cor_vec_all_select[i,1] = cor_vec_i_select_str
}


###### cCRE per gene after selection: abs(cor)>=0.2
#[1] "summary(all_ccre_num):"
#   Min. 1st Qu.  Median    Mean 3rd Qu.    Max. 
#   1.00    8.00   13.00   14.87   21.00   64.00 
#[1] "summary(select_ccre_num):"
#   Min. 1st Qu.  Median    Mean 3rd Qu.    Max. 
#  0.000   3.000   7.000   8.101  11.000  43.000
#[1] 936
# 4.7%

###### cCRE per gene after selection: (cor)>=0.2
#[1] "summary(all_ccre_num):"
#   Min. 1st Qu.  Median    Mean 3rd Qu.    Max. 
#   1.00    8.00   13.00   14.87   21.00   64.00 
#[1] "summary(select_ccre_num):"
#   Min. 1st Qu.  Median    Mean 3rd Qu.    Max. 
#  0.000   2.000   4.000   4.714   7.000  34.000 
#[1] 2194
# 11.1%


### redo sp coverage after cor filtering
ss = read.table(paste('S3V2_IDEAS_hg38_ccre2.cCRE.M.withid.S0.mat.txt', sep=''), header=F)
ss_all = (ss[,-c(1:4)])
for (i in c(1:(state_n-1))){
print(i)
ss = read.table(paste('S3V2_IDEAS_hg38_ccre2.cCRE.M.withid.S',i,'.mat.txt', sep=''), header=F)
sp = (ss[,-c(1:4)])
ss_all = cbind(ss_all, sp)
}

sp_corfiltered = matrix(0, nrow=dim(cor_vec_all_select)[1], ncol=dim(ss_all)[2])

for (i in 1:dim(cor_vec_all_select)[1]){
if (i%%100==0){
	print(i)
}
used_row_i = as.numeric(unlist(strsplit(cor_vec_all_select[i,1], ',')))
if (length(used_row_i)>0){
sp_corfiltered[i,] = colSums(ss_all[used_row_i,])
}else{
sp_corfiltered[i,] = rep(0, dim(sp_corfiltered)[2])
}
}


sp_corfiltered_all = matrix(0, nrow=dim(sp_corfiltered)[1]*length(common_ct), ncol=state_n)
k=0
for (ct in common_ct){
print(ct)
ct_pos_sp = which(sp_list==ct)
sp_mat_tmp = matrix(0, nrow=dim(sp_corfiltered)[1], ncol=state_n )
for (i in ct_pos_sp){
sp_mat_tmp = sp_mat_tmp+sp_corfiltered[,seq(1,dim(sp_corfiltered)[2], by=dim(sp_corfiltered)[2]/state_n)+i-1]
}
sp_mat_tmp = sp_mat_tmp/length(ct_pos_sp)
sp_corfiltered_all[seq(1,dim(sp_corfiltered)[1])+(k)*dim(sp_corfiltered)[1],] = sp_mat_tmp
k = k+1
}

colnames(sp_corfiltered_all) = c(0:(state_n-1))
#sp_corfiltered_all_D_log = sp_corfiltered_all
#sp_corfiltered_all_D_log = log(sp_corfiltered_all+smallnum_sp)
sp_corfiltered_all_D_log = log((sp_corfiltered_all+smallnum_sp))
#sp_corfiltered_all_D_log = sp_corfiltered_all

#sp_all_PD_corfilter_log = cbind(sp_all_log, sp_corfiltered_all_D_log*global_scale)
sp_all_PD_corfilter_log = cbind(sp_all_log, (sp_corfiltered_all_D_log - mean(adj_mean_od)) / adj_sd_od * adj_sd_tar + adj_mean_tar )


### dimension reducation rm 0 state info -c(1,25)
### PCA lm

used_idtry = dmslog_noMean>(-1000000)
sp_all_PD_corfilter_log_new = sp_all_PD_corfilter_log[,-c(1,(state_n+1))] %*% pca0_rotation

lm_all = lm(dmslog_noMean[used_idtry]~sp_all_PD_corfilter_log_new[used_idtry,c(1:pcn)])
Bpca_all = pca0_rotation[,c(1:pcn)] %*% lm_all$coefficients[-1]

pred = lm_all$fitted.value #+ mean(c(ds_qt_all_log_l, ds_qt_all_log_h))
#pca_all$x[,1:pcn] %*% lm_all$coefficients + mean(c(ds_qt_all_log_l, ds_qt_all_log_h))
R2(dmslog_noMean[used_idtry], pred)
0.5644413

###
set.seed(2019)
used_id = sample(length(pred), 50000)
plot_lim = plot_lim_all#c(min(c(dmslog_noMean+mean(ds_qt_all_log), pred)), max(c(dmslog_noMean+mean(ds_qt_all_log), pred)))

png(paste(output_folder, '/pred_obs.tpm3.png', sep=''))
#heatscatter(as.vector(pred)[used_id]+mean(ds_qt_all_log), as.vector(dmslog_noMean)[used_id]+mean(ds_qt_all_log), xlim=plot_lim, ylim=plot_lim)
heatscatter(as.vector(pred)[used_id], as.vector(dmslog_noMean)[used_id], xlim=plot_lim, ylim=plot_lim, main=R2(dmslog_noMean[used_idtry], pred))
abline(0,1, col='red', lwd=2)
abline(h=0, col='black', lwd=2)
abline(v=0, col='black', lwd=2)
dev.off()


eRP_mat_human = cbind(Bpca_all[1:(state_n-1)],Bpca_all[(state_n):((state_n-1)*2)])
eRP_mat_human = rbind(c(0,0), eRP_mat_human)
colnames(eRP_mat_human) = c('P','D')
rownames(eRP_mat_human) = 0:(state_n-1)
write.table(eRP_mat_human, paste(output_folder, '/statep_rna_coe_heatmap.human.all.ccre.withcorfilter.txt', sep=''), quote=F, col.names=T, row.names=T, sep='\t')

plot_lim_P = max(abs(eRP_mat_human[,1]))
plot_lim_D = max(abs(eRP_mat_human[,2]))
plot_lim_PD = max(abs(eRP_mat_human))

pdf(paste(output_folder, '/statep_rna_coe_heatmap.human.all.ccre.withcorfilter.pdf', sep=''), width=3)
#rank = c(9,2,3,1,7,4,15,14,13,5,11,22,6,18,8,24,12,10,20,16,23,17,19,21)
breaksList = seq(-plot_lim_PD, plot_lim_PD, by = 0.001)
my_colorbar=colorRampPalette(c('blue', 'white', 'red'))(n = length(breaksList))
col_breaks = c(seq(0, 2000,length=33))
pheatmap(eRP_mat_human[rank,], color=my_colorbar, breaks = breaksList, cluster_cols = FALSE,cluster_rows=F, clustering_method = 'average',annotation_names_row=TRUE,annotation_names_col=TRUE,show_rownames=TRUE,show_colnames=TRUE)
dev.off()

pdf(paste(output_folder, '/statep_rna_coe_heatmap.human.all.P.ccre.withcorfilter.pdf', sep=''), width=3)
#rank = c(9,2,3,1,7,4,15,14,13,5,11,22,6,18,8,24,12,10,20,16,23,17,19,21)
breaksList = seq(-plot_lim_PD, plot_lim_PD, by = 0.001)
my_colorbar=colorRampPalette(c('blue', 'white', 'red'))(n = length(breaksList))
col_breaks = c(seq(0, 2000,length=33))
pheatmap(cbind(eRP_mat_human[rank,1],eRP_mat_human[rank,1]), color=my_colorbar, breaks = breaksList, cluster_cols = FALSE,cluster_rows=F, clustering_method = 'average',annotation_names_row=TRUE,annotation_names_col=TRUE,show_rownames=TRUE,show_colnames=TRUE)
dev.off()

pdf(paste(output_folder, '/statep_rna_coe_heatmap.human.all.D.ccre.withcorfilter.pdf', sep=''), width=3)
#rank = c(9,2,3,1,7,4,15,14,13,5,11,22,6,18,8,24,12,10,20,16,23,17,19,21)
breaksList = seq(-plot_lim_PD, plot_lim_PD, by = 0.001)
my_colorbar=colorRampPalette(c('blue', 'white', 'red'))(n = length(breaksList))
col_breaks = c(seq(0, 2000,length=33))
pheatmap(cbind(eRP_mat_human[rank,2],eRP_mat_human[rank,2]), color=my_colorbar, breaks = breaksList, cluster_cols = FALSE,cluster_rows=F, clustering_method = 'average',annotation_names_row=TRUE,annotation_names_col=TRUE,show_rownames=TRUE,show_colnames=TRUE)
dev.off()

write.table(sp_all_PD_corfilter_log, paste(output_folder, '/sp_all_PD_corfilter_log.human.txt', sep=''), quote=F, col.names=F, row.names=F, sep='\t')
write.table(dmslog_noMean, paste(output_folder, '/dmslog_noMean.human.txt', sep=''), quote=F, col.names=F, row.names=F, sep='\t')
```
