## Supplementary material for "Interspecies regulatory landscapes and elements revealed by novel joint systematic integration of human and mouse blood cell epigenomes": Scripts for IDEAS modeling of epigenetic states jointly in human and mouse: README.md

###
```
# get_coe_score.05292022.JustHuman.sh: is used for Only Human data analysis. (From Downloading data to the final outputs)

# get_coe_score.04292022.human.sh: is used for joint Human & Mouse data analysis

# get_cCRE_coe_mat.human.R: is used for getting esRP (Human only) for all cCREs across all cell-types' replicates

# get_state_Beta_coefficients.human.R: is the core function to calculate Beta coefficients for all IDEAS states

# get_ave.R: is used for get the average Beta coefficients for all leave-one-out runs

```
