## Supplementary material for "Interspecies regulatory landscapes and elements revealed by novel joint systematic integration of human and mouse blood cell epigenomes": Scripts for IDEAS modeling of epigenetic states jointly in human and mouse: add_id.R

```
library(data.table)
args = commandArgs(trailingOnly=TRUE)

input_bed = args[1]
output_bed = args[2]

bed = as.data.frame(fread(input_bed))
bed_id = 1:dim(bed)[1]
bed_out = cbind(bed, bed_id)

write.table(bed_out, output_bed, quote=F, col.names=F, row.names=F, sep='\t')
```
