## Supplementary material for "Interspecies regulatory landscapes and elements revealed by novel joint systematic integration of human and mouse blood cell epigenomes": Scripts for IDEAS modeling of epigenetic states jointly in human and mouse: get_coe_score.04292022.mouse.sh

```
cd /Users/guanjuexiang/Downloads/coe_analysis_mouse

### download data
wget https://usevision.org/data/mm10/rnaTPMall_withcoordinates.0.txt
Rscript get_new_RNA_mat.mouse.R

cut -f1,2,3,4 rnaTPM_withcoordinates.txt | tail -n+2 | sort -k1,1 -k2,2n > RNA_gene.bed
cat rnaTPM_withcoordinates.txt | head -1 > MouseVISION_RNAseq_mm10_genes_tpm.idsort.header.txt
tail -n+2 rnaTPM_withcoordinates.txt | sort -k4,4 > MouseVISION_RNAseq_mm10_genes_tpm.idsort.txt

wget https://usevision.org/data/mm10/ideasJointMay2021/S3V2_IDEAS_mm10_ccre2.cCRE.M.notall0.bed
wget https://usevision.org/data/mm10/ideasJointMay2021/S3V2_IDEAS_mm10_r3_withHg38Mm10prior.state


cat MouseVISION_RNAseq_mm10_genes_tpm.idsort.txt  | awk -F '\t' -v OFS='\t' '{if ($5=="protein_coding") print $1,$2,$3,$5,$6,$4, $7,$8,$9,$10,$11,$12,$13,$14,$15,$16,$17,$18}' > MouseVISION_RNAseq_mm10_genes_tpm.idsort.protein_coding.txt
cat MouseVISION_RNAseq_mm10_genes_tpm.idsort.txt  \
| awk -F '\t' -v OFS='\t' '{print $1,$2,$3,$5,$6,$4}' | cut -f1,2,3,5\
| awk -F '\t' -v OFS='\t' -v exp_win=1000 '{if (($2-exp_win>0) && ($4=="+")) print $1,$2-exp_win, $2+exp_win; if (($3-exp_win>0) && ($4=="-")) print $1,$3-exp_win, $3+exp_win; if (($2-exp_win<0) && ($4=="+")) print $1,0, $2+exp_win; if (($3-exp_win<0) && ($4=="-")) print $1,0, $3+exp_win}' \
> MouseVISION_RNAseq_mm10_genes_tpm.idsort.all.Nkbupdownexp.bed

cut -f1,2,3,5 MouseVISION_RNAseq_mm10_genes_tpm.idsort.protein_coding.txt > MouseVISION_RNAseq_mm10_genes_tpm.idsort.protein_coding.bed

### get protein coding gene proximal regions +-1kb 
cat MouseVISION_RNAseq_mm10_genes_tpm.idsort.protein_coding.bed \
| awk -F '\t' -v OFS='\t' -v exp_win=1000 '{if (($2-exp_win>0) && ($4=="+")) print $1,$2-exp_win, $2+exp_win; if (($3-exp_win>0) && ($4=="-")) print $1,$3-exp_win, $3+exp_win; if (($2-exp_win<0) && ($4=="+")) print $1,0, $2+exp_win; if (($3-exp_win<0) && ($4=="-")) print $1,0, $3+exp_win}' \
> MouseVISION_RNAseq_mm10_genes_tpm.idsort.protein_coding.Nkbupdownexp.bed

### get protein coding gene distal regions +-50kb 
cat MouseVISION_RNAseq_mm10_genes_tpm.idsort.protein_coding.bed \
| awk -F '\t' -v OFS='\t' -v exp_win=50000 '{if (($2-exp_win>0) && ($4=="+")) print $1,$2-exp_win, $2+exp_win; if (($3-exp_win>0) && ($4=="-")) print $1,$3-exp_win, $3+exp_win; if (($2-exp_win<0) && ($4=="+")) print $1,0, $2+exp_win; if (($3-exp_win<0) && ($4=="-")) print $1,0, $3+exp_win}' \
> MouseVISION_RNAseq_mm10_genes_tpm.idsort.protein_coding.NHkbupdownexp.bed

### add cCRE ids
bedtools intersect -a S3V2_IDEAS_mm10_ccre2.cCRE.M.notall0.bed -b MouseVISION_RNAseq_mm10_genes_tpm.idsort.protein_coding.Nkbupdownexp.bed -v > S3V2_IDEAS_mm10_ccre2.cCRE.M.notall0.noProximal.bed
time Rscript add_id.R S3V2_IDEAS_mm10_ccre2.cCRE.M.notall0.bed S3V2_IDEAS_mm10_ccre2.cCRE.M.withid.bed 
bedtools intersect -a S3V2_IDEAS_mm10_ccre2.cCRE.M.withid.bed -b MouseVISION_RNAseq_mm10_genes_tpm.idsort.all.Nkbupdownexp.bed -c > S3V2_IDEAS_mm10_ccre2.cCRE.M.withid.atProximal.bed


### convert .state file to .bed file
tail -n+2 S3V2_IDEAS_mm10_r3_withHg38Mm10prior.state \
| awk -F ' ' -v OFS='\t' '{print $2,$3,$4, $0}' > J_IDEAS.state.bed

### only save the state bin intersect with cCRE
bedtools intersect -a J_IDEAS.state.bed -b S3V2_IDEAS_mm10_ccre2.cCRE.M.notall0.bed -wa -u \
> J_IDEAS.state.ccre.bed
cut -f4 J_IDEAS.state.ccre.bed > J_IDEAS.state.inccre

### only save the state bin intersect with genes promoter +-1kb
bedtools intersect -a J_IDEAS.state.bed -b MouseVISION_RNAseq_mm10_genes_tpm.idsort.protein_coding.Nkbupdownexp.bed -wa -u \
> J_IDEAS.state.gene.bed
cut -f4 J_IDEAS.state.gene.bed > J_IDEAS.state.ingene

### get TSS state
for i in {0..24}
do
echo $i
cp MouseVISION_RNAseq_mm10_genes_tpm.idsort.protein_coding.Nkbupdownexp.bed MouseVISION_RNAseq_mm10_genes_tpm.idsort.protein_coding.Nkbupdownexp.S$i.bed
for j in {5..36}
do
echo $j
cat J_IDEAS.state.ingene \
| awk -F ' ' -v OFS='\t' -v c=$j -v s=$i '{if ($c==s) print $2,$3,$4}' > ct$j.s$i.bed
echo 'NO filter cCRE'
bedtools intersect -a MouseVISION_RNAseq_mm10_genes_tpm.idsort.protein_coding.Nkbupdownexp.bed -b ct$j.s$i.bed -c > ct$j.s$i.bed.count.bed
cut -f4 ct$j.s$i.bed.count.bed > ct$j.s$i.bed.count.txt
paste MouseVISION_RNAseq_mm10_genes_tpm.idsort.protein_coding.Nkbupdownexp.S$i.bed ct$j.s$i.bed.count.txt > MouseVISION_RNAseq_mm10_genes_tpm.idsort.protein_coding.Nkbupdownexp.S$i.bed.tmp \
&& mv MouseVISION_RNAseq_mm10_genes_tpm.idsort.protein_coding.Nkbupdownexp.S$i.bed.tmp MouseVISION_RNAseq_mm10_genes_tpm.idsort.protein_coding.Nkbupdownexp.S$i.bed
rm ct$j.s$i.bed.count.txt ct$j.s$i.bed.count.bed ct$j.s$i.bed
done
done


### get distal states
for i in {0..24}
do
echo $i
cp MouseVISION_RNAseq_mm10_genes_tpm.idsort.protein_coding.NHkbupdownexp.bed MouseVISION_RNAseq_mm10_genes_tpm.idsort.protein_coding.NHkbupdownexp.S$i.bed
for j in {5..36}
do
echo $j
cat J_IDEAS.state.inccre \
| awk -F ' ' -v OFS='\t' -v c=$j -v s=$i '{if ($c==s) print $2,$3,$4}' > ct$j.s$i.ccre.bed
echo 'filter cCRE'
bedtools intersect -a MouseVISION_RNAseq_mm10_genes_tpm.idsort.protein_coding.NHkbupdownexp.bed -b ct$j.s$i.ccre.bed -c > ct$j.s$i.bed.count.bed
cut -f4 ct$j.s$i.bed.count.bed > ct$j.s$i.bed.count.txt
paste MouseVISION_RNAseq_mm10_genes_tpm.idsort.protein_coding.NHkbupdownexp.S$i.bed ct$j.s$i.bed.count.txt > MouseVISION_RNAseq_mm10_genes_tpm.idsort.protein_coding.NHkbupdownexp.S$i.bed.tmp \
&& mv MouseVISION_RNAseq_mm10_genes_tpm.idsort.protein_coding.NHkbupdownexp.S$i.bed.tmp MouseVISION_RNAseq_mm10_genes_tpm.idsort.protein_coding.NHkbupdownexp.S$i.bed
rm ct$j.s$i.bed.count.txt ct$j.s$i.bed.count.bed ct$j.s$i.ccre.bed
done
done

time Rscript add_id.R MouseVISION_RNAseq_mm10_genes_tpm.idsort.protein_coding.NHkbupdownexp.bed MouseVISION_RNAseq_mm10_genes_tpm.idsort.protein_coding.NHkbupdownexp.withid.bed 
sort -k1,1 -k2,2n MouseVISION_RNAseq_mm10_genes_tpm.idsort.protein_coding.NHkbupdownexp.withid.bed > MouseVISION_RNAseq_mm10_genes_tpm.idsort.protein_coding.NHkbupdownexp.withid.sort.bed
bedtools map -a MouseVISION_RNAseq_mm10_genes_tpm.idsort.protein_coding.NHkbupdownexp.withid.sort.bed -b S3V2_IDEAS_mm10_ccre2.cCRE.M.withid.bed -c 4 -o distinct > MouseVISION_RNAseq_mm10_genes_tpm.idsort.protein_coding.NHkbupdownexp.withccreid.bed
sort -k4,4n MouseVISION_RNAseq_mm10_genes_tpm.idsort.protein_coding.NHkbupdownexp.withccreid.bed > MouseVISION_RNAseq_mm10_genes_tpm.idsort.protein_coding.NHkbupdownexp.withccreid.bed.tmp
mv MouseVISION_RNAseq_mm10_genes_tpm.idsort.protein_coding.NHkbupdownexp.withccreid.bed.tmp MouseVISION_RNAseq_mm10_genes_tpm.idsort.protein_coding.NHkbupdownexp.withccreid.bed


### get cCREs
for i in {0..24}
do
echo $i
cp S3V2_IDEAS_mm10_ccre2.cCRE.M.withid.bed S3V2_IDEAS_mm10_ccre2.cCRE.M.withid.S$i.mat.txt
for j in {5..36}
do
echo $j
cat J_IDEAS.state.inccre \
| awk -F ' ' -v OFS='\t' -v c=$j -v s=$i '{if ($c==s) print $2,$3,$4}' > ct$j.s$i.bed
### NO filter cCRE
bedtools intersect -a S3V2_IDEAS_mm10_ccre2.cCRE.M.withid.bed -b ct$j.s$i.bed -c > ct$j.s$i.bed.count.bed
cut -f5 ct$j.s$i.bed.count.bed > ct$j.s$i.bed.count.txt
paste S3V2_IDEAS_mm10_ccre2.cCRE.M.withid.S$i.mat.txt ct$j.s$i.bed.count.txt > S3V2_IDEAS_mm10_ccre2.cCRE.M.withid.S$i.mat.txt.tmp \
&& mv S3V2_IDEAS_mm10_ccre2.cCRE.M.withid.S$i.mat.txt.tmp S3V2_IDEAS_mm10_ccre2.cCRE.M.withid.S$i.mat.txt
rm ct$j.s$i.bed.count.txt ct$j.s$i.bed.count.bed ct$j.s$i.bed
done
done

### get cCREs All
bedtools intersect -a J_IDEAS.state.bed -b S3V2_IDEAS_mm10_ccre2.cCRE.M.notall0.bed -wa -u \
> J_IDEAS.state.ccre.all.bed
cut -f4 J_IDEAS.state.ccre.all.bed > J_IDEAS.state.inccre.all
###
time Rscript add_id.R S3V2_IDEAS_mm10_ccre2.cCRE.M.notall0.bed S3V2_IDEAS_mm10_ccre2.cCRE.M.notall0.withid.bed
###
for i in {0..24}
do
echo $i
cp S3V2_IDEAS_mm10_ccre2.cCRE.M.notall0.withid.bed S3V2_IDEAS_mm10_ccre2.cCRE.M.notall0.withid.S$i.mat.txt
for j in {5..36}
do
echo $j
cat J_IDEAS.state.inccre.all \
| awk -F ' ' -v OFS='\t' -v c=$j -v s=$i '{if ($c==s) print $2,$3,$4}' > ct$j.s$i.bed
### NO filter cCRE
bedtools intersect -a S3V2_IDEAS_mm10_ccre2.cCRE.M.notall0.withid.bed -b ct$j.s$i.bed -c > ct$j.s$i.bed.count.bed
cut -f5 ct$j.s$i.bed.count.bed > ct$j.s$i.bed.count.txt
paste S3V2_IDEAS_mm10_ccre2.cCRE.M.notall0.withid.S$i.mat.txt ct$j.s$i.bed.count.txt > S3V2_IDEAS_mm10_ccre2.cCRE.M.notall0.withid.S$i.mat.txt.tmp \
&& mv S3V2_IDEAS_mm10_ccre2.cCRE.M.notall0.withid.S$i.mat.txt.tmp S3V2_IDEAS_mm10_ccre2.cCRE.M.notall0.withid.S$i.mat.txt
rm ct$j.s$i.bed.count.txt ct$j.s$i.bed.count.bed ct$j.s$i.bed
done
done


### get cCRE expand states
#cat S3V2_IDEAS_hg38_ccre2.cCRE.M.withid.bed | awk -F '\t' -v OFS='\t' -v exp_win=50000 '{if (($2-exp_win)>0) print $1,$2-exp_win, $3+exp_win, $4; else print $1,0, $3+exp_win, $4}' > S3V2_IDEAS_hg38_ccre2.cCRE.M.withid.expand.bed
#bedtools intersect -a J_IDEAS.state.bed -b S3V2_IDEAS_hg38_ccre2.cCRE.M.withid.expand.bed -wa -u \
#> J_IDEAS.state.ccre_exp.bed
#cut -f4 J_IDEAS.state.ccre_exp.bed > J_IDEAS.state.inccre_exp

#for i in {0..24}
#do
#echo $i
#cp S3V2_IDEAS_hg38_ccre2.cCRE.M.withid.bed S3V2_IDEAS_hg38_ccre2.cCRE.expand.M.withid.S$i.mat.txt
#for j in {5..47}
#do
#echo $j
#cat J_IDEAS.state.inccre_exp \
#| awk -F ' ' -v OFS='\t' -v c=$j -v s=$i '{if ($c==s) print $2,$3,$4}' > ct$j.s$i.bed
#### NO filter cCRE
#bedtools intersect -a S3V2_IDEAS_hg38_ccre2.cCRE.M.withid.expand.bed -b ct$j.s$i.bed -c > ct$j.s$i.bed.count.bed
#cut -f5 ct$j.s$i.bed.count.bed > ct$j.s$i.bed.count.txt
#paste S3V2_IDEAS_hg38_ccre2.cCRE.expand.M.withid.S$i.mat.txt ct$j.s$i.bed.count.txt > S3V2_IDEAS_hg38_ccre2.cCRE.expand.M.withid.S$i.mat.txt.tmp \
#&& mv S3V2_IDEAS_hg38_ccre2.cCRE.expand.M.withid.S$i.mat.txt.tmp S3V2_IDEAS_hg38_ccre2.cCRE.expand.M.withid.S$i.mat.txt
#rm ct$j.s$i.bed.count.txt ct$j.s$i.bed.count.bed ct$j.s$i.bed
#done
#done

declare -a common_ct=("CFUE" "CFUMK" "CMP" "ERYfl" "GMP" "LSK" "MEP" "MONO" "NEU" "iMK" "G1E" "ER4")
## now loop through the above array
for ct in "${common_ct[@]}"
do
   echo "$ct"
   time Rscript /Users/guanjuexiang/Documents/projects/Joint_Human_House_IDEAS_State/other_scripts/state_coe_score/coe_mouse/get_coe_pcareg.mouse.withccre_withcorfilter.local.iter.25JES_PD.R $ct
done

Rscript get_ave.R
```
