## Supplementary material for "Interspecies regulatory landscapes and elements revealed by novel joint systematic integration of human and mouse blood cell epigenomes": Scripts for IDEAS modeling of epigenetic states jointly in human and mouse: get_new_RNA_mat.mouse.R

```
d = read.table('rnaTPMall_withcoordinates.0.txt', header=T)

ds = 2^(d[,-c(1:6)])
ds = ds-1e-3
ds[ds<0] = 0

dnew = cbind(d[,1:6],ds)
write.table(dnew, 'rnaTPM_withcoordinates.txt', quote=F, col.names=T, row.names=F, sep='\t')
```
