## Supplementary material for "Interspecies regulatory landscapes and elements revealed by novel joint systematic integration of human and mouse blood cell epigenomes": Scripts for IDEAS modeling of epigenetic states jointly in human and mouse: get_new_RNA_mat.R

```
d1 = read.table('tpmFeb21_v3.tab', sep='\t', header=T)
d2 = read.table('HumanVISION_RNAseq_hg38_genes_tpm.old.txt', sep='\t', header=T)

d2_id = apply(d2,1, function(x) unlist(strsplit(x[4], '[.]'))[1])

shared_d2_id = is.element(d2_id, d1[,1])

d2a = d2[shared_d2_id,]

d1a = cbind(d2a[,1:4],d1[,-1])

write.table(d1a, 'HumanVISION_RNAseq_hg38_genes_tpm.txt', quote=F, sep='\t', col.names=T, row.names=F)
```
