## Supplementary material for "Interspecies regulatory landscapes and elements revealed by novel joint systematic integration of human and mouse blood cell epigenomes": Scripts for IDEAS modeling of epigenetic states jointly in human and mouse: get_cor_heatmap_km_qda.human.withmodification.R

```
library(dynamicTreeCut)
library(pheatmap)

#d = read.table('S3V2_IDEAS_hg38_ccre2.cCRE.M.notall0.rmallNEU.withid.coe_mat.txt', header=T)
HsP_cCRE = read.table('../coe_analysis/S3V2_IDEAS_hg38_ccre2.cCRE.M.withid.atProximal.bed', header=F)
d = read.table('../coe_analysis/S3V2_IDEAS_hg38_ccre2.cCRE.M.notall0.rmallNEU.withid.coe_mat.PDmerged.txt', header=T)
dPDsep = read.table('../coe_analysis/S3V2_IDEAS_hg38_ccre2.cCRE.M.notall0.rmallNEU.withid.coe_mat.txt', header=T)

ds = d[,-c(1:4)]


set.seed(2019)
#used_row = sample(dim(ds)[1], dim(ds)[1]
dss = ds
dss[ds>quantile(as.numeric(as.matrix(ds)),0.99)] = quantile(as.numeric(as.matrix(ds)),0.99)


### ct dist cor
dPDsep = dPDsep[,!is.element(colnames(dPDsep), c('chr', 'start', 'end', 'id', 'NEU_C0011IH2_P','NEU_C001UYH1_P', 'NEU_C0011IH2_D','NEU_C001UYH1_D'))]
ds_forcor = dPDsep[,(dim(dPDsep)[2]/2+1):dim(dPDsep)[2]]
#ds_forcor[ds_forcor>quantile(as.numeric(as.matrix(ds)),1)] = quantile(as.numeric(as.matrix(ds)),1)
ds_cor_D = cor(ds_forcor)
#ds_cor_D = cor(ds)
hclust_cor = hclust(as.dist(1-ds_cor_D))
hclust_cor_order = hclust_cor$order

png('cCRE_coe.human.heatmap.D.cttree.png', height=600, width=1200)
plot(hclust_cor, hang = -1, cex = 2, lwd=2)
dev.off()

rep1 = c(1, 2, 4, 6, 8, 10, 12, 14, 16, 18, 20, 22, 24, 26, 28, 30, 32, 34, 36, 38, 40)
rep2 = c(1, 3, 5, 7, 9, 11, 13, 15, 17, 19, 21, 23, 25, 27, 29, 31, 33, 35, 37, 39, 41)

dss_ct = c()
for (i in 1:length(rep1)){
	dss_ct = cbind(dss_ct, rowMeans(cbind(dss[,rep1[i]], dss[,rep2[i]])))
}

within_c_dist = function(x, c_vec){
	within_dist = 0
	for (i in min(c_vec):max(c_vec)){
		x_i = x[c_vec==i,]
		if (sum(c_vec==i)>1){
			x_i_colm = colMeans(x_i)
			x_i_dist = sum(apply(x_i, 1, function(x) sum((x-x_i_colm)^2)))
		} else{
			x_i_colm = as.numeric(x_i)
			x_i_dist = 0
		}
		within_dist = within_dist+x_i_dist
	}
	return(within_dist)
}

set.seed(2019)
dist_ratio = c()
for (n in seq(10,200,10)){
	print(n)
	dss_km_n = kmeans(dss_ct, n)
	dist_km = within_c_dist(dss_ct, dss_km_n$cluster)
	dss_km_ave_n = kmeans(cbind(rowMeans(dss_ct)), n)
	dist_ave = within_c_dist(dss_ct, dss_km_ave_n$cluster)
	dist_ratio = rbind(dist_ratio, c(n, dist_km, dist_ave, dss_km_n$totss))	
}

pdf('km_k_dist.pdf')
plot(c(1,seq(10,200,10)), c(dss_km_n$totss, dist_ratio[,2]))
lines(c(1,seq(10,200,10)), c(dss_km_n$totss, dist_ratio[,2]))
dev.off()

library(MASS)
### get 100 round of km k=50
set.seed(2019)
km_k = 100
km_centers = c()
iter_n = 2
for (r in 1:iter_n){
	print(r)
	dss_km_i = kmeans(dss_ct, km_k)
	for (k_i in 1:km_k){
	km_centers = rbind(km_centers, colMeans(dss_ct[dss_km_i$cluster==k_i,]))
	}
}

### get reproducible clusters
set.seed(2019)
kmkm = kmeans(km_centers, km_k)

pdf('kmkm_n.hist.pdf')
hist(table(kmkm$cluster), breaks=20)
dev.off()

kmkm_n_z = ( table(kmkm$cluster) - mean(table(kmkm$cluster)) ) /sd(table(kmkm$cluster))
#used_kmkm_centers = which(pnorm(kmkm_n_z, lower.tail=TRUE)>=0.05)
#used_kmkm_centers = which(table(kmkm$cluster)>=(iter_n/2))
used_kmkm_centers = which(table(kmkm$cluster)>=0)
### get new km centers
km_new_centers = c()
for (i in 1:length(used_kmkm_centers)){
	if (sum(kmkm$cluster==i)>1){
		km_new_centers = rbind(km_new_centers, colMeans(as.matrix(km_centers)[kmkm$cluster==i,]))
	}else{
		km_new_centers = rbind(km_new_centers, as.matrix(km_centers)[kmkm$cluster==i,])
	}
}

### reassign cCREs into reproducible clusters
x_dist_to_newcenter = c()
for (i in 1:dim(km_new_centers)[1]){
	print(i)
x_i_dist = apply(dss_ct, 1, function(x) sum((x-km_new_centers[i,])^2))
x_dist_to_newcenter = cbind(x_dist_to_newcenter, x_i_dist)
}

x_dist_to_newcenter_reassign = apply(x_dist_to_newcenter,1,which.min)

png('cCRE_coe.human.heatmap.D.png', width=1200, height=1800)
plot_lim_PD = quantile(as.numeric(as.matrix(dss)),0.99)
breaksList = seq(-plot_lim_PD, plot_lim_PD, by = 0.001)
my_colorbar=colorRampPalette(c('blue', 'white', 'red'))(n = length(breaksList))
col_breaks = c(seq(0, 2000,length=33))
dss_plot = round(dss,3)
dss_plot[dss>plot_lim_PD]=plot_lim_PD
pheatmap(dss_plot[order(x_dist_to_newcenter_reassign),hclust_cor_order], color=my_colorbar, breaks = breaksList, cluster_cols = F,cluster_rows=F, clustering_method = 'complete',annotation_names_row=F,annotation_names_col=TRUE,show_rownames=F,show_colnames=TRUE)
dev.off()


### QDA rescue
set.seed(2019)
#pre_cluster = x_dist_to_newcenter_reassign
#for (i in 1:1){
#qda_model = qda(round(dss_ct,3), pre_cluster)
#fit_cluster_reorder_vec_after_rescue = predict(qda_model, dss_ct)$class
##fit_cluster_reorder_vec[fit_cluster_reorder_vec==0] = fit_cluster_reorder_vec_after_rescue
#print(sum(pre_cluster!=fit_cluster_reorder_vec_after_rescue))
#if (sum(pre_cluster!=fit_cluster_reorder_vec_after_rescue)==0){break}
#pre_cluster = fit_cluster_reorder_vec_after_rescue
#print(cbind(table(pre_cluster)))
#}
#fit_cluster_reorder_vec_after_rescue = pre_cluster
fit_cluster_reorder_vec_after_rescue = x_dist_to_newcenter_reassign

### reorder clustering results
### get mean vec
dss_mean_vec = c()
for (i in c(1:length(unique(fit_cluster_reorder_vec_after_rescue)))){
	dss_mean_vec = rbind(dss_mean_vec, colMeans(dss[fit_cluster_reorder_vec_after_rescue==i,]))
}


### kmean meta
set.seed(2019)
dist_ratio = c()
for (n in seq(2,30,1)){
	print(n)
	dss_km_n = kmeans(dss_mean_vec, n)
	#print(table(dss_km_n$cluster))
	dist_km = within_c_dist(dss_mean_vec, dss_km_n$cluster)
	#print(dist_km)
	dist_ratio = rbind(dist_ratio, c(n, dist_km, dss_km_n$totss))	
}

pdf('km_k_dist.ave.pdf')
plot(c(1,seq(2,30,1)), c(dss_km_n$totss, dist_ratio[,2]))
lines(c(1,seq(2,30,1)), c(dss_km_n$totss, dist_ratio[,2]))
dev.off()


### kmean meta
#set.seed(2019)
#kmeta = 20
#kmeans_meta = kmeans(dss_mean_vec, centers = kmeta)
#old_km_ids = 1:length(unique(fit_cluster_reorder_vec_after_rescue))
#cluster_id_order = c(old_km_ids)[order(kmeans_meta$cluster)]

#### get kmeans meta clusterID
#new_id_meta = rep(-1, length(fit_cluster_reorder_vec_after_rescue))
#for (i in 1:kmeta){
#meta_i_vec = old_km_ids[kmeans_meta$cluster==i]
#new_id_meta[is.element(fit_cluster_reorder_vec_after_rescue, meta_i_vec)] = i
#}

#### reorder clusters
#new_id = fit_cluster_reorder_vec_after_rescue
#k = 0
#for (i in 1:length(cluster_id_order)){
#new_id[fit_cluster_reorder_vec_after_rescue==cluster_id_order[i]] = i
#}


### kmean meta
set.seed(2019)
rownames(dss_mean_vec) = 1:dim(dss_mean_vec)[1]
### add noise to split high vs low sig
dss_mean_vec_add_noise = dss_mean_vec #+ matrix(runif(dim(dss_mean_vec)[1]*dim(dss_mean_vec)[2], -0.05, 0.05), dim(dss_mean_vec)[1], dim(dss_mean_vec)[2])
dss_mean_vec_add_noise_merge_rep = c()
for (i in 1:length(rep1)){
	dss_mean_vec_add_noise_merge_rep = cbind(dss_mean_vec_add_noise_merge_rep, rowMeans(cbind(dss_mean_vec_add_noise[,rep1[i]], dss_mean_vec_add_noise[,rep2[i]])))
}
colnames(dss_mean_vec_add_noise_merge_rep) = colnames(ds_forcor)[rep1]


### 5 ct group
#ct_group_HSC = dss_mean_vec_add_noise_merge_rep[,c(4,5,8,9,13,14,18)]
#ct_group_ERY = dss_mean_vec_add_noise_merge_rep[,c(3,7,10,11,12,15)]
#ct_group_MON = dss_mean_vec_add_noise_merge_rep[,c(6,16,17)]
#ct_group_LYM = dss_mean_vec_add_noise_merge_rep[,c(2,19,20,21)]

#ct_5group_mat = cbind(rowMeans(ct_group_HSC), rowMeans(ct_group_ERY), rowMeans(ct_group_MON), rowMeans(ct_group_LYM))
#ct_5group_mat = cbind(rowMeans(ct_group_HSC), rowMeans(ct_group_ERY), rowMeans(ct_group_MON), rowMeans(ct_group_LYM))
#colnames(ct_5group_mat) = c(colnames(dss_mean_vec_add_noise_merge_rep),'HSC','ERY','MONO','LYMP')
allct_ct_5group_mat = cbind(dss_mean_vec_add_noise_merge_rep)

library(lsa)
#dss_mean_vec_cor_dist = as.dist(1 - (cosine(t(allct_ct_5group_mat))) )
dss_mean_vec_cor_dist = as.dist(1 - cor(t(dss_mean_vec_add_noise)))
#dss_mean_vec_cor_dist_ct5group = as.dist(1 - cosine(t(ct_5group_mat)))
#dss_mean_vec_cor_dist = as.dist(1 - (cosine(t(allct_ct_5group_mat))*cor(t(allct_ct_5group_mat))) )
#dss_mean_vec_cor_dist = as.dist(1 - (cor(t(allct_ct_5group_mat))^2 * (cor(t(allct_ct_5group_mat))/abs(cor(t(allct_ct_5group_mat))))) )
#dss_mean_vec_cor_dist = as.dist(1 - cosine(t(pcs$x[,1:12])))
#dss_mean_vec_cor_dist = as.dist(1 - cor(t(dss_mean_vec_add_noise_merge_rep)))
#dss_mean_vec_cor_dist = dist(allct_ct_5group_mat)
#kmeans_meta_ct5group = hclust(dss_mean_vec_cor_dist_ct5group, method = 'complete')
kmeans_meta = hclust(dss_mean_vec_cor_dist, method = 'complete')

pdf('meta.tree.pdf', width=30)
plot(kmeans_meta)
dev.off()

### reorder clusters
k = 0
new_id = fit_cluster_reorder_vec_after_rescue
for (i in kmeans_meta$order){
k = k+1
new_id[fit_cluster_reorder_vec_after_rescue==i] = k
}

### cut tree
#kmeans_meta_dynamicTC = cutreeDynamic(kmeans_meta, distM=as.matrix(dss_mean_vec_cor_dist), minClusterSize=1, deepSplit=2, method='hybrid')
#kmeans_meta_dynamicTC_ct5group = cutreeDynamic(kmeans_meta_ct5group, minClusterSize=2, deepSplit=4, method='tree')
#kmeans_meta_dynamicTC_ct = cutreeDynamic(kmeans_meta, minClusterSize=2, deepSplit=4, method='tree')
#kmeans_meta_dynamicTC = paste(kmeans_meta_dynamicTC_ct5group, kmeans_meta_dynamicTC)
kmeans_meta_dynamicTC = cutreeDynamic(kmeans_meta, minClusterSize=2, deepSplit=4, method='tree')
if (min(kmeans_meta_dynamicTC)==0){kmeans_meta_dynamicTC = kmeans_meta_dynamicTC+1}
old_km_ids = 1:km_k

table_pre = table(kmeans_meta_dynamicTC)
for (k in 1:10){
### get new centers
allct_ct_5group_mat_new_center = matrix(0, nrow=length(unique(kmeans_meta_dynamicTC)), ncol=dim(allct_ct_5group_mat)[2])
for (ki in 1:length(unique(kmeans_meta_dynamicTC))){
if (sum(kmeans_meta_dynamicTC==ki)>1){
allct_ct_5group_mat_new_center[ki,] = colMeans(allct_ct_5group_mat[kmeans_meta_dynamicTC==ki,])
} else if (sum(kmeans_meta_dynamicTC==ki)==1){
allct_ct_5group_mat_new_center[ki,] = allct_ct_5group_mat[kmeans_meta_dynamicTC==ki,]
}
}

### new meta cluster
new_cosine_mat = matrix(0, nrow=dim(allct_ct_5group_mat)[1], ncol=dim(allct_ct_5group_mat_new_center)[1])
for (i in 1:dim(allct_ct_5group_mat)[1]){
for (j in 1:dim(allct_ct_5group_mat_new_center)[1]){
	new_cosine_mat[i,j] = cosine(allct_ct_5group_mat[i,],allct_ct_5group_mat_new_center[j,])
}
}
###
kmeans_meta_dynamicTC = apply(new_cosine_mat,1,which.max)
print(table(kmeans_meta_dynamicTC))
if (length(table_pre)==length(table(kmeans_meta_dynamicTC))){
if (sum(table_pre == table(kmeans_meta_dynamicTC))==length(table_pre)){break}
}
table_pre = table(kmeans_meta_dynamicTC)
}

### get kmeans meta clusterID
new_id_meta = rep(-1, length(fit_cluster_reorder_vec_after_rescue))
for (i in old_km_ids){
meta_i_vec = i #old_km_ids[kmeans_meta_dynamicTC==i]
new_id_meta[fit_cluster_reorder_vec_after_rescue == i] = kmeans_meta_dynamicTC[i]
}
#new_id_meta = as.numeric(as.factor(new_id_meta))

### adjust meta-clusters 
new_id_meta_1 = new_id_meta
table(new_id[new_id_meta==6])
new_id_meta_1[is.element(new_id, c(11,12))]  = max(new_id_meta_1)+1
new_id_meta_1[is.element(new_id, c(37,38))]  = max(new_id_meta_1)+1
table(new_id[new_id_meta==14])
table(new_id[new_id_meta==20])
new_id_meta_1[is.element(new_id, c(89,90))]  = max(new_id_meta_1)+1
table(new_id[new_id_meta==16])
table(new_id[new_id_meta==17])
new_id_meta_1[is.element(new_id, c(24,32,33,34,15,16,20))]  = max(new_id_meta_1)+1
table(new_id[new_id_meta==20])
new_id_meta_1[is.element(new_id, c(48,49,91,92))]  = max(new_id_meta_1)+1
table(new_id[new_id_meta==20])
table(new_id[new_id_meta==21])
new_id_meta_1[is.element(new_id, c(50,85,93,77,78))]  = max(new_id_meta_1)+1

#table(new_id[new_id_meta==1])
#table(new_id[new_id_meta==3])
#new_id_meta_1[is.element(new_id, c(1,27,90))]  = max(new_id_meta_1)+1
#table(new_id[new_id_meta==2])
#new_id_meta_1[is.element(new_id, c(6,7))]  = max(new_id_meta_1)+1
#new_id_meta_1[is.element(new_id, c(11:13))]  = max(new_id_meta_1)+1
#table(new_id[new_id_meta==4])
#new_id_meta_1[is.element(new_id, c(75:78))]  = max(new_id_meta_1)+1
#table(new_id[new_id_meta==6])
#new_id_meta_1[is.element(new_id, c(4))]  = max(new_id_meta_1)+1
#table(new_id[new_id_meta==8])
#table(new_id[new_id_meta==13])
#new_id_meta_1[is.element(new_id, c(25,26,68:71))]  = max(new_id_meta_1)+1


### plot heatmap
png('cCRE_coe.human.heatmap.D.qda.png', width=1200, height=1800)
plot_lim_PD = quantile(as.numeric(as.matrix(dss)),1)
breaksList = seq(-plot_lim_PD, plot_lim_PD, by = 0.001)
my_colorbar=colorRampPalette(c('blue', 'white', 'red'))(n = length(breaksList))
col_breaks = c(seq(0, 2000,length=33))
dss_plot = dss
dss_plot[dss>plot_lim_PD]=plot_lim_PD
pheatmap(dss_plot[order(new_id),hclust_cor_order][order(new_id_meta[order(new_id)]),], color=my_colorbar, breaks = breaksList, cluster_cols = F,cluster_rows=F, clustering_method = 'complete',annotation_names_row=F,annotation_names_col=TRUE,show_rownames=F,show_colnames=TRUE)
#pheatmap(dss_plot[,hclust_cor_order][order(new_id_meta),], color=my_colorbar, breaks = breaksList, cluster_cols = F,cluster_rows=F, clustering_method = 'complete',annotation_names_row=F,annotation_names_col=TRUE,show_rownames=F,show_colnames=TRUE)
dev.off()


png('cCRE_coe.human.heatmap.D.qda.clusterID.png', width=200, height=1800)
library(RColorBrewer)
n = 200#length(unique(new_id))
qual_col_pals = brewer.pal.info[brewer.pal.info$category == 'qual',]
col_vector = c(unlist(mapply(brewer.pal, qual_col_pals$maxcolors, rownames(qual_col_pals))), 
unlist(mapply(brewer.pal, qual_col_pals$maxcolors, rownames(qual_col_pals))), 
unlist(mapply(brewer.pal, qual_col_pals$maxcolors, rownames(qual_col_pals))), 
unlist(mapply(brewer.pal, qual_col_pals$maxcolors, rownames(qual_col_pals))) )
dss_cluster_id = cbind(new_id, new_id_meta)
longest_colname = max(nchar(colnames(dss)))
colnames(dss_cluster_id) = c(colnames(dss)[nchar(colnames(dss))==longest_colname][1], paste(c(rep('b', longest_colname-2), 'ID'), collapse=''))
pheatmap(dss_cluster_id[order(new_id),][order(new_id_meta[order(new_id)]),], color=col_vector, cluster_cols = F,cluster_rows=F, clustering_method = 'complete',annotation_names_row=F,annotation_names_col=TRUE,show_rownames=F,show_colnames=TRUE)
#pheatmap(dss_cluster_id[,][order(new_id_meta),], color=col_vector, cluster_cols = F,cluster_rows=F, clustering_method = 'complete',annotation_names_row=F,annotation_names_col=TRUE,show_rownames=F,show_colnames=TRUE)
dev.off()


### plot heatmap
png('cCRE_coe.human.heatmap.D.qda.adj.png', width=1200, height=1800)
plot_lim_PD = quantile(as.numeric(as.matrix(dss)),1)
breaksList = seq(-plot_lim_PD, plot_lim_PD, by = 0.001)
my_colorbar=colorRampPalette(c('blue', 'white', 'red'))(n = length(breaksList))
col_breaks = c(seq(0, 2000,length=33))
dss_plot = dss
dss_plot[dss>plot_lim_PD]=plot_lim_PD
pheatmap(dss_plot[order(new_id),hclust_cor_order][order(new_id_meta_1[order(new_id)]),], color=my_colorbar, breaks = breaksList, cluster_cols = F,cluster_rows=F, clustering_method = 'complete',annotation_names_row=F,annotation_names_col=TRUE,show_rownames=F,show_colnames=TRUE)
#pheatmap(dss_plot[,hclust_cor_order][order(new_id_meta),], color=my_colorbar, breaks = breaksList, cluster_cols = F,cluster_rows=F, clustering_method = 'complete',annotation_names_row=F,annotation_names_col=TRUE,show_rownames=F,show_colnames=TRUE)
dev.off()


png('cCRE_coe.human.heatmap.D.qda.clusterID.adj.png', width=200, height=1800)
library(RColorBrewer)
n = 200#length(unique(new_id))
qual_col_pals = brewer.pal.info[brewer.pal.info$category == 'qual',]
col_vector = c(unlist(mapply(brewer.pal, qual_col_pals$maxcolors, rownames(qual_col_pals))), 
unlist(mapply(brewer.pal, qual_col_pals$maxcolors, rownames(qual_col_pals))), 
unlist(mapply(brewer.pal, qual_col_pals$maxcolors, rownames(qual_col_pals))), 
unlist(mapply(brewer.pal, qual_col_pals$maxcolors, rownames(qual_col_pals))) )
dss_cluster_id = cbind(new_id, new_id_meta_1)
longest_colname = max(nchar(colnames(dss)))
colnames(dss_cluster_id) = c(colnames(dss)[nchar(colnames(dss))==longest_colname][1], paste(c(rep('b', longest_colname-2), 'ID'), collapse=''))
pheatmap(dss_cluster_id[order(new_id),][order(new_id_meta_1[order(new_id)]),], color=col_vector, cluster_cols = F,cluster_rows=F, clustering_method = 'complete',annotation_names_row=F,annotation_names_col=TRUE,show_rownames=F,show_colnames=TRUE)
#pheatmap(dss_cluster_id[,][order(new_id_meta),], color=col_vector, cluster_cols = F,cluster_rows=F, clustering_method = 'complete',annotation_names_row=F,annotation_names_col=TRUE,show_rownames=F,show_colnames=TRUE)
dev.off()


png('cCRE_coe.human.heatmap.D.qda.shufflekm.png', width=1200, height=1800)
plot_lim_PD = quantile(as.numeric(as.matrix(dss)),1)
breaksList = seq(-plot_lim_PD, plot_lim_PD, by = 0.001)
my_colorbar=colorRampPalette(c('blue', 'white', 'red'))(n = length(breaksList))
col_breaks = c(seq(0, 2000,length=33))
dss_plot = dss
dss_plot[dss>plot_lim_PD]=plot_lim_PD
pheatmap(dss_plot[,hclust_cor_order][order(new_id_meta_1),], color=my_colorbar, breaks = breaksList, cluster_cols = F,cluster_rows=F, clustering_method = 'complete',annotation_names_row=F,annotation_names_col=TRUE,show_rownames=F,show_colnames=TRUE)
#pheatmap(dss_plot[,hclust_cor_order][order(new_id_meta),], color=my_colorbar, breaks = breaksList, cluster_cols = F,cluster_rows=F, clustering_method = 'complete',annotation_names_row=F,annotation_names_col=TRUE,show_rownames=F,show_colnames=TRUE)
dev.off()


###
ddd = read.table('S3V2_IDEAS_hg38_ccre2.cCRE.M.notall0.rmallNEU.withid.coe_mat.PDmerged.clusterID.100.txt', header=T, sep='\t')


png('cCRE_coe.human.heatmap.D.qda.meanvec.png', width=1200, height=1800)
plot_lim_PD = quantile(as.numeric(as.matrix(dss)),1)
breaksList = seq(-plot_lim_PD, plot_lim_PD, by = 0.001)
my_colorbar=colorRampPalette(c('blue', 'white', 'red'))(n = length(breaksList))
col_breaks = c(seq(0, 2000,length=33))
dss_plot = dss
dss_plot[dss>plot_lim_PD]=plot_lim_PD
pheatmap(dss_mean_vec[kmeans_meta$order,hclust_cor_order], color=my_colorbar, breaks = breaksList, cluster_cols = F,cluster_rows=F, clustering_method = 'complete',annotation_names_row=F,annotation_names_col=TRUE,show_rownames=F,show_colnames=TRUE)
dev.off()
unique(kmeans_meta_dynamicTC[kmeans_meta$order])

new_id_meta_uniq = as.numeric(names(table(new_id_meta_1)))
new_id_meta_rename = rep(-1, length(new_id_meta_1))
for (i in 1:length(new_id_meta_uniq)){
	print(c(new_id_meta_uniq[i], sum(new_id_meta_1==new_id_meta_uniq[i])))
	### rename based on heatmap order
	new_id_meta_rename[new_id_meta_1==new_id_meta_uniq[i]] = i
}

dss_out = cbind(d[,c(1:4)], new_id, new_id_meta_rename, ds)
colnames(dss_out)[5:6] = c('clusterID', 'meta_clusterID')

write.table(dss_out, 'S3V2_IDEAS_hg38_ccre2.cCRE.M.notall0.rmallNEU.withid.coe_mat.PDmerged.clusterID.txt', quote=F, row.names=F, col.names=T, sep='\t')
#write.table(cbind(table(kmeans_meta_dynamicTC[kmeans_meta$order]))[unique(kmeans_meta_dynamicTC[kmeans_meta$order]),], 'S3V2_IDEAS_hg38_ccre2.cCRE.M.notall0.rmallNEU.withid.coe_mat.PDmerged.metaCluster_Heatmap_order.txt', quote=F, row.names=T, col.names=T, sep='\t')

### plot mean matrix
library(pheatmap)
coe = read.table('S3V2_IDEAS_hg38_ccre2.cCRE.M.notall0.rmallNEU.withid.coe_mat.PDmerged.clusterID.txt', header=T, sep='\t')

Meta_cluster_id = unique(coe[,6])[order(unique(coe[,6]))]
meta_cluster_mean_mat = c()
for (meta_i in Meta_cluster_id){
	meta_cluster_mean_mat = rbind(meta_cluster_mean_mat, colMeans(coe[coe[,6]==meta_i,-c(1:6)]))
}
rownames(meta_cluster_mean_mat) = Meta_cluster_id
remove_str_end = function(x){
	x_split = unlist(strsplit(x, '_'))
	x_split = x_split[-length(x_split)]
	return(paste(x_split, collapse='_'))
}

colnames(meta_cluster_mean_mat) = apply(cbind(colnames(meta_cluster_mean_mat)), 1, remove_str_end )
png('VISION.hg38.cCRE.meta_cluster.meansig.png', width=1000, height=1000)
plot_lim_PD = quantile(as.numeric(as.matrix(meta_cluster_mean_mat)),0.99)
breaksList = seq(-plot_lim_PD, plot_lim_PD, by = 0.001)
my_colorbar=colorRampPalette(c('blue', 'white', 'red'))(n = length(breaksList))
pheatmap(meta_cluster_mean_mat[,hclust_cor_order], cluster_cols=F, cluster_rows=F, color=my_colorbar, breaks = breaksList, cex=1.5)
dev.off()


KM_cluster_id = unique(coe[,5])[order(unique(coe[,5]))]
km_cluster_mean_mat = c()
for (km_i in KM_cluster_id){
	km_cluster_mean_mat = rbind(km_cluster_mean_mat, colMeans(coe[coe[,5]==km_i,-c(1:6)]))
}
rownames(km_cluster_mean_mat) = KM_cluster_id
colnames(km_cluster_mean_mat) = apply(cbind(colnames(km_cluster_mean_mat)), 1, remove_str_end )

png('VISION.hg38.cCRE.KM100_cluster.meansig.png', width=1000, height=2000)
plot_lim_PD = quantile(as.numeric(as.matrix(km_cluster_mean_mat)),0.99)
breaksList = seq(-plot_lim_PD, plot_lim_PD, by = 0.001)
my_colorbar=colorRampPalette(c('blue', 'white', 'red'))(n = length(breaksList))
pheatmap(km_cluster_mean_mat[,hclust_cor_order], cluster_cols=F, cluster_rows=F, color=my_colorbar, breaks = breaksList, cex=1.5)
dev.off()


#Great analysis:
#Proximal: 5.0kb upstream, 1.0kb downstream, 
#Distal: up to 100kb

#cat S3V2_IDEAS_hg38_ccre2.cCRE.M.notall0.rmallNEU.withid.coe_mat.clusterID.txt | awk -F '\t' -v OFS='\t' '{if ($5!=1000 && $6==19) print $1,$2,$3}' > S3V2_IDEAS_hg38_ccre2.cCRE.M.notall0.rmallNEU.withid.coe_mat.clusterID.metaERY.txt
#cat S3V2_IDEAS_hg38_ccre2.cCRE.M.notall0.rmallNEU.withid.coe_mat.clusterID.txt | awk -F '\t' -v OFS='\t' '{if ($5!=1000 && $6==9) print $1,$2,$3}' > S3V2_IDEAS_hg38_ccre2.cCRE.M.notall0.rmallNEU.withid.coe_mat.clusterID.metaMONc.txt
#cat S3V2_IDEAS_hg38_ccre2.cCRE.M.notall0.rmallNEU.withid.coe_mat.clusterID.txt | awk -F '\t' -v OFS='\t' '{if (($5!=1000 && $6==9) || ($5!=1000 && $6==19)) print $1,$2,$3}' > S3V2_IDEAS_hg38_ccre2.cCRE.M.notall0.rmallNEU.withid.coe_mat.clusterID.metaERY_MONc.txt
#cat S3V2_IDEAS_hg38_ccre2.cCRE.M.notall0.rmallNEU.withid.coe_mat.clusterID.txt | awk -F '\t' -v OFS='\t' '{if ($5!=1000 && $6==13) print $1,$2,$3}' > S3V2_IDEAS_hg38_ccre2.cCRE.M.notall0.rmallNEU.withid.coe_mat.clusterID.metaERY.13.txt
#cat S3V2_IDEAS_hg38_ccre2.cCRE.M.notall0.rmallNEU.withid.coe_mat.clusterID.txt | awk -F '\t' -v OFS='\t' '{if ($5!=1000 && $6==4) print $1,$2,$3}' > S3V2_IDEAS_hg38_ccre2.cCRE.M.notall0.rmallNEU.withid.coe_mat.clusterID.metaMONc.4.txt
```
