## Supplementary material for "Interspecies regulatory landscapes and elements revealed by novel joint systematic integration of human and mouse blood cell epigenomes": Scripts for IDEAS modeling of epigenetic states jointly in human and mouse: get_cor_heatmap.human_vs_mouse.addmultipleIDs.0829.R

```
#conda activate som
library(dynamicTreeCut)
library(pheatmap)
library(lsa)
library(LSD)
#################################################
dh = read.table('S3V2_IDEAS_hg38_ccre2.cCRE.M.notall0.rmallNEU.withid.coe_mat.PDmerged.clusterID.txt', header=T, sep='\t')
dm = read.table('S3V2_IDEAS_mm10_ccre2.cCRE.M.notall0.withid.coe_mat.PDmerged.clusterID.txt', header=T, sep='\t')

dhs = dh[,-c(1:6)]
rep1_h = c(1, 2, 4, 6, 8, 10, 12, 14, 16, 18, 20, 22, 24, 26, 28, 30, 32, 34, 36, 38, 40)
rep2_h = c(1, 3, 5, 7, 9, 11, 13, 15, 17, 19, 21, 23, 25, 27, 29, 31, 33, 35, 37, 39, 41)
ct_h = c('AVE', 'B', 'CD34', 'CLP', 'CMP', 'EOS', 'ERY', 'GMP', 'HSC', 'HUDEP1', 'HUDEP2', 'K562', 'LMPP', 'MEP', 'MK', 'MON', 'MONp', 'MPP', 'NK', 'TCD4', 'TCD8')
dhs_ctmerge = matrix(0, nrow=dim(dhs)[1], ncol=length(ct_h))
colnames(dhs_ctmerge) = ct_h
for (j in 1:length(ct_h)){
	dhs_ctmerge[,j] = (dhs[,rep1_h[j]] + dhs[,rep2_h[j]])/2
}

dms = dm[,-c(1:6)]
rep1_m = c(1, 2, 4, 5, 6, 7, 8, 10, 12, 14, 16, 17, 18, 19, 21, 22, 24, 25, 26, 27, 28, 29, 31)
rep2_m = c(1, 3, 4, 5, 6, 7, 9, 11, 13, 15, 16, 17, 18, 20, 21, 23, 24, 25, 26, 27, 28, 30, 32)
ct_m = c('AVE', 'B', 'CFUE', 'CFUMK', 'CLP', 'CMP', 'ER4', 'ERY', 'ERYfl', 'G1E', 'GMP', 'HPC7', 'HSC', 'MEL', 'MEP', 'MK', 'MON', 'NEU', 'NK', 'TCD4', 'TCD8', 'iMEL', 'iMK')
dms_ctmerge = matrix(0, nrow=dim(dms)[1], ncol=length(ct_m))
colnames(dms_ctmerge) = ct_m
for (j in 1:length(ct_m)){
	dms_ctmerge[,j] = (dms[,rep1_m[j]] + dms[,rep2_m[j]])/2
}

### shared cts
ct_shared = ct_h[is.element(ct_h, ct_m)]
### human
dhs_ctmerge_shared = dhs_ctmerge[,is.element(colnames(dhs_ctmerge), ct_shared)]
dhs_ctmerge_shared = dhs_ctmerge_shared[,order(colnames(dhs_ctmerge_shared))]
### mouse
dms_ctmerge_shared = dms_ctmerge[,is.element(colnames(dms_ctmerge), ct_shared)]
dms_ctmerge_shared = dms_ctmerge_shared[,order(colnames(dms_ctmerge_shared))]

### ct reorder
ct_shared_reorder = c('AVE', 'HSC', 'CMP', 'CLP', 'MEP', 'ERY', 'MK', 'GMP', 'MON', 'NK', 'B', 'TCD4', 'TCD8')
dhs_ctmerge_shared_reorder = c()
dms_ctmerge_shared_reorder = c()
for (i in 1:length(ct_shared_reorder)){
	dhs_ctmerge_shared_reorder = cbind(dhs_ctmerge_shared_reorder, dhs_ctmerge_shared[,colnames(dhs_ctmerge_shared)==ct_shared_reorder[i]])
	dms_ctmerge_shared_reorder = cbind(dms_ctmerge_shared_reorder, dms_ctmerge_shared[,colnames(dms_ctmerge_shared)==ct_shared_reorder[i]])
}
colnames(dhs_ctmerge_shared_reorder) = ct_shared_reorder
colnames(dms_ctmerge_shared_reorder) = ct_shared_reorder

### recalcualte average
dhs_ctmerge_shared_reorder[,1] = rowMeans(dhs_ctmerge_shared_reorder[,-1])
dms_ctmerge_shared_reorder[,1] = rowMeans(dms_ctmerge_shared_reorder[,-1])
#################################################


#################################################
### get lm batch effect removal
ct_i = 1
ct_name_i = colnames(dhs_ctmerge_shared_reorder)[ct_i]
km_mean_sig_h = as.numeric(dhs_ctmerge_shared_reorder[,ct_i])
km_mean_sig_h = km_mean_sig_h[order(-km_mean_sig_h)][1:10000]
km_mean_sig_m = as.numeric(dms_ctmerge_shared_reorder[,ct_i])
km_mean_sig_m = km_mean_sig_m[order(-km_mean_sig_m)][1:10000]
plot_lim_kmmean = c(min(c(km_mean_sig_h, km_mean_sig_m)), max(c(km_mean_sig_h, km_mean_sig_m)))
### lm regression model
lm_kmmean = lm(km_mean_sig_h~km_mean_sig_m)
### 
for (ct_i in 1:dim(dhs_ctmerge_shared_reorder)[2]){
	ct_name_i = colnames(dhs_ctmerge_shared_reorder)[ct_i]
km_mean_sig_h = as.numeric(dhs_ctmerge_shared_reorder[,ct_i])
km_mean_sig_h = km_mean_sig_h[order(-km_mean_sig_h)][1:10000]
km_mean_sig_m = as.numeric(dms_ctmerge_shared_reorder[,ct_i])
km_mean_sig_m = km_mean_sig_m[order(-km_mean_sig_m)][1:10000]
plot_lim_kmmean = c(min(c(km_mean_sig_h, km_mean_sig_m)), max(c(km_mean_sig_h, km_mean_sig_m)))
### lm regression model
#lm_kmmean = lm(km_mean_sig_h~km_mean_sig_m)
png(paste0(ct_name_i,'.km_ctave_sig.heatscatter.ave.png'), width=1000)
par(mfrow=c(1,2))
heatscatter(km_mean_sig_h[order(km_mean_sig_h)], km_mean_sig_m[order(km_mean_sig_m)], pch=16, xlim=plot_lim_kmmean, ylim=plot_lim_kmmean)
abline(0,1)
heatscatter(km_mean_sig_h[order(km_mean_sig_h)], km_mean_sig_m[order(km_mean_sig_m)]*lm_kmmean$coefficients[2]+lm_kmmean$coefficients[1], pch=16, xlim=plot_lim_kmmean, ylim=plot_lim_kmmean)
abline(0,1)
dev.off()
}
### batch removal
km_mean_sig_h = as.numeric(dhs_ctmerge_shared_reorder[,1])
km_mean_sig_m = as.numeric(dms_ctmerge_shared_reorder[,1])
dms_ctmerge_shared_reorder_batcheffect_rm = dms_ctmerge_shared_reorder*lm_kmmean$coefficients[2]+lm_kmmean$coefficients[1]
###
png('check_cross_ct.mean.before.png')
plot(1:13, apply(dhs_ctmerge_shared_reorder,2,function(x) x[order(-x)][5000]), ylim=c(0.0, 1.5))
lines(1:13, apply(dhs_ctmerge_shared_reorder,2,function(x) x[order(-x)][5000]), col='blue')
points(1:13, apply(dms_ctmerge_shared_reorder,2,function(x) x[order(-x)][5000]))
lines(1:13, apply(dms_ctmerge_shared_reorder,2,function(x) x[order(-x)][5000]))
###
points(1:13, apply(dhs_ctmerge_shared_reorder,2,function(x) x[order(-x)][1000]))
lines(1:13, apply(dhs_ctmerge_shared_reorder,2,function(x) x[order(-x)][1000]), col='blue')
points(1:13, apply(dms_ctmerge_shared_reorder,2,function(x) x[order(-x)][1000]))
lines(1:13, apply(dms_ctmerge_shared_reorder,2,function(x) x[order(-x)][1000]))
###
points(1:13, apply(dhs_ctmerge_shared_reorder,2,function(x) x[order(-x)][50000]))
lines(1:13, apply(dhs_ctmerge_shared_reorder,2,function(x) x[order(-x)][50000]), col='blue')
points(1:13, apply(dms_ctmerge_shared_reorder,2,function(x) x[order(-x)][50000]))
lines(1:13, apply(dms_ctmerge_shared_reorder,2,function(x) x[order(-x)][50000]))
dev.off()
###
png('check_cross_ct.mean.after.png')
plot(1:13, apply(dhs_ctmerge_shared_reorder,2,function(x) x[order(-x)][5000]), ylim=c(0.0, 1.5))
lines(1:13, apply(dhs_ctmerge_shared_reorder,2,function(x) x[order(-x)][5000]), col='blue')
points(1:13, apply(dms_ctmerge_shared_reorder_batcheffect_rm,2,function(x) x[order(-x)][5000]))
lines(1:13, apply(dms_ctmerge_shared_reorder_batcheffect_rm,2,function(x) x[order(-x)][5000]))
###
points(1:13, apply(dhs_ctmerge_shared_reorder,2,function(x) x[order(-x)][1000]))
lines(1:13, apply(dhs_ctmerge_shared_reorder,2,function(x) x[order(-x)][1000]), col='blue')
points(1:13, apply(dms_ctmerge_shared_reorder_batcheffect_rm,2,function(x) x[order(-x)][1000]))
lines(1:13, apply(dms_ctmerge_shared_reorder_batcheffect_rm,2,function(x) x[order(-x)][1000]))
###
points(1:13, apply(dhs_ctmerge_shared_reorder,2,function(x) x[order(-x)][50000]))
lines(1:13, apply(dhs_ctmerge_shared_reorder,2,function(x) x[order(-x)][50000]), col='blue')
points(1:13, apply(dms_ctmerge_shared_reorder_batcheffect_rm,2,function(x) x[order(-x)][50000]))
lines(1:13, apply(dms_ctmerge_shared_reorder_batcheffect_rm,2,function(x) x[order(-x)][50000]))
dev.off()
#################################################


#################################################
### merge Human and Mouse matrix
dhs_dms_ctmerge_shared_reorder = rbind(dhs_ctmerge_shared_reorder, dms_ctmerge_shared_reorder)
cCRE_id = c(paste0(rep('H', dim(dhs_ctmerge_shared_reorder)[1])), paste0(rep('M', dim(dms_ctmerge_shared_reorder)[1])))
coordinates = rbind(dh[,c(1:6)], dm[,c(1:6)])
### Human & mouse cCRE PCA
dhs_ctmerge_shared_reorder_pca = prcomp(dhs_dms_ctmerge_shared_reorder, center = F, scale. = F)
dhs_ctmerge_shared_reorder_pca_summary = summary(dhs_ctmerge_shared_reorder_pca)
print(dhs_ctmerge_shared_reorder_pca_summary$importance)
pca_used_num = sum(dhs_ctmerge_shared_reorder_pca_summary$importance[3,]<0.95)+1
set.seed(2019)
used_id = sample(dim(dhs_ctmerge_shared_reorder_pca$x)[1], 10000)
dhs_ctmerge_shared_reorder_pca_x_plot = dhs_ctmerge_shared_reorder_pca$x[used_id,]
cCRE_id_plot = cCRE_id[used_id]
png('PCA.plot.raw.png')
plot(dhs_ctmerge_shared_reorder_pca_x_plot[,1], dhs_ctmerge_shared_reorder_pca_x_plot[,2])
points(dhs_ctmerge_shared_reorder_pca_x_plot[cCRE_id_plot=='H',1], dhs_ctmerge_shared_reorder_pca_x_plot[cCRE_id_plot=='H',2], col='red')
points(dhs_ctmerge_shared_reorder_pca_x_plot[cCRE_id_plot=='M',1], dhs_ctmerge_shared_reorder_pca_x_plot[cCRE_id_plot=='M',2], col='blue')
dev.off()
#################################################


#################################################
### cluster cCRE by PCA-KM by subsampling and identify reproducible clusters pca 1:5 cover 95% of variance
### 1 example
set.seed(2019)
KMPCA_meansig_mat_1 = c()
start_km_num = 100
km_dhs_ctmerge_shared1 = kmeans(dhs_ctmerge_shared_reorder_pca$x[,1:pca_used_num], centers=start_km_num)
### check KM cluster cCRE Human/Mouse ratio
for (i in 1:start_km_num){
	dhs_dms_ctmerge_shared_reorder_ji = dhs_dms_ctmerge_shared_reorder[km_dhs_ctmerge_shared1$cluster==i,]
	KMPCA_meansig_mat_1 = rbind(KMPCA_meansig_mat_1, colMeans(dhs_dms_ctmerge_shared_reorder_ji))
}
colnames(KMPCA_meansig_mat_1) = colnames(dhs_ctmerge_shared_reorder)
pdf('KMPCA.Joint.1.cluster.pdf', height=30)
pheatmap(KMPCA_meansig_mat_1, cluster_col=T, cluster_rows=T)
dev.off()
#################################################


#################################################
### Select reproducible State
#################################################
### check initial KM-K
set.seed(2019)
cCRE_esRP_KM_determineK_KM_ratio = c()
start_km_num = 110
for (ini_k in seq(10,200,by=10)){
	print(ini_k)
	used_id_j = sample(dim(dhs_ctmerge_shared_reorder_pca$x)[1], 50000)
	km_dhs_ctmerge_shared_check = kmeans(dhs_ctmerge_shared_reorder_pca$x[used_id_j,1:pca_used_num], centers=ini_k)
	cCRE_esRP_KM_determineK_KM_ratio = c(cCRE_esRP_KM_determineK_KM_ratio, km_dhs_ctmerge_shared_check$tot.withinss/km_dhs_ctmerge_shared_check$betweenss)
}
pdf('cCRE_esRP_KM.determineK.pdf')
plot(seq(10,200,by=10), cCRE_esRP_KM_determineK_KM_ratio)
lines(seq(10,200,by=10), cCRE_esRP_KM_determineK_KM_ratio)
abline(v=start_km_num)
dev.off()

### Initial Kmeans
set.seed(2019)
KMPCA_meansig_mat = c()
KMPCA_iteration = c()
KMPCA_cCRE_num = c()
HM_count = c()
check_reproducible = 100
start_km_num = 110
for (j in 1:check_reproducible){
print(j)
used_id_j = sample(dim(dhs_ctmerge_shared_reorder_pca$x)[1], 50000)
km_dhs_ctmerge_shared1 = kmeans(dhs_ctmerge_shared_reorder_pca$x[used_id_j,1:pca_used_num], centers=start_km_num)
### check KM cluster cCRE Human/Mouse ratio
for (i in 1:start_km_num){
	if (sum(km_dhs_ctmerge_shared1$cluster==i)>100){
	dhs_dms_ctmerge_shared_reorder_ji = dhs_dms_ctmerge_shared_reorder[used_id_j,][km_dhs_ctmerge_shared1$cluster==i,]
	KMPCA_meansig_mat = rbind(KMPCA_meansig_mat, colMeans(dhs_dms_ctmerge_shared_reorder_ji))
	KMPCA_cCRE_num = c(KMPCA_cCRE_num, dim(dhs_dms_ctmerge_shared_reorder_ji)[1])
	cCRE_id_ji = cCRE_id[used_id_j][km_dhs_ctmerge_shared1$cluster==i]
	HM_count = rbind(HM_count, c(sum(cCRE_id_ji=='H'), sum(cCRE_id_ji=='M')))
	}
}
KMPCA_iteration = c(KMPCA_iteration, rep(j, start_km_num))
}
#################################################


#################################################
### identify reproducible state
set.seed(2019)
KMPCA_meansig_mat_pca_x = KMPCA_meansig_mat %*% dhs_ctmerge_shared_reorder_pca$rotation
KMPCA_meansig_mat_pca_joint_KM = kmeans(KMPCA_meansig_mat_pca_x[,1:pca_used_num], centers=start_km_num)

KMPCA_iteration_KM_k = c()
KMPCA_meansig_mat_meansig_mat = c()
HM_count_mean = c()
cCRE_count_KMPCA_iteration_KM_k = c()
cluster_set = min(unique(KMPCA_meansig_mat_pca_joint_KM$cluster)):max(unique(KMPCA_meansig_mat_pca_joint_KM$cluster))
for (KM_k in cluster_set){
	used_KM_k = KMPCA_meansig_mat_pca_joint_KM$cluster==KM_k
	KMPCA_iteration_KM_k = c(KMPCA_iteration_KM_k, length(unique(KMPCA_iteration[used_KM_k])) )
	if (sum(used_KM_k)>1){
		KMPCA_meansig_mat_meansig_mat = rbind(KMPCA_meansig_mat_meansig_mat, colMeans(KMPCA_meansig_mat[used_KM_k,]))
		HM_count_mean = rbind(HM_count_mean, apply(HM_count[used_KM_k,], 2, sum))
		cCRE_count_KMPCA_iteration_KM_k = c(cCRE_count_KMPCA_iteration_KM_k, sum(KMPCA_cCRE_num[used_KM_k]))
	} else{
		KMPCA_meansig_mat_meansig_mat = rbind(KMPCA_meansig_mat_meansig_mat, KMPCA_meansig_mat[used_KM_k,])
		HM_count_mean = rbind(HM_count_mean, HM_count[used_KM_k,])
		cCRE_count_KMPCA_iteration_KM_k = c(cCRE_count_KMPCA_iteration_KM_k, KMPCA_cCRE_num[used_KM_k])
	}
}

HM_count_mean_log2FC = log2((HM_count_mean[,1]+1)/(HM_count_mean[,2]/mean(HM_count_mean[,2])*mean(HM_count_mean[,1])+1))
#select_Kclusters = ((KMPCA_iteration_KM_k>=(check_reproducible*0.9)) * (abs(HM_count_mean_log2FC)<=1000))!=0
pdf('KM.reproducible.p.hist.pdf')
hist(KMPCA_iteration_KM_k/check_reproducible, breaks=20)
abline(v=0.6)
box()
dev.off()

pdf('KM.cross_species_abslog2FC.p.hist.pdf')
hist(abs(HM_count_mean_log2FC), breaks=15)
abline(v=2)
box()
dev.off()

pdf('KM.cross_species_abslog2FC.MA.pdf')
M = log2((HM_count_mean[,1]+1)/(HM_count_mean[,2]/mean(HM_count_mean[,2])*mean(HM_count_mean[,1])+1))
A = log2((HM_count_mean[,1]+1)*(HM_count_mean[,2]/mean(HM_count_mean[,2])*mean(HM_count_mean[,1])+1))
plot(A, M, ylim=c(-max(abs(M)), max(abs(M))))
abline(h=0)
dev.off()


select_Kclusters = ((KMPCA_iteration_KM_k>=(check_reproducible*0.6)) * (abs(HM_count_mean_log2FC)<=1))!=0
KMPCA_meansig_mat_meansig_mat_reproducible = KMPCA_meansig_mat_meansig_mat[select_Kclusters,]
cCRE_count_KMPCA_iteration_KM_k_reproducible = cCRE_count_KMPCA_iteration_KM_k[select_Kclusters]
colnames(KMPCA_meansig_mat_meansig_mat_reproducible) = colnames(dhs_ctmerge_shared_reorder)
dim(KMPCA_meansig_mat_meansig_mat_reproducible)

pdf('KMPCA.Joint.reproducible.cluster.pdf', height=10, width=6)
pheatmap(KMPCA_meansig_mat_meansig_mat_reproducible, cluster_col=T, cluster_rows=T, clustering_distance_rows=dist(1-cor(t(KMPCA_meansig_mat_meansig_mat_reproducible))))
#pheatmap(KMPCA_meansig_mat_meansig_mat_reproducible, cluster_col=T, cluster_rows=T, clustering_distance_rows=dist(1-cosine(t(KMPCA_meansig_mat_meansig_mat_reproducible))))
#pheatmap(KMPCA_meansig_mat_meansig_mat_reproducible, cluster_col=T, cluster_rows=T, clustering_distance_rows=dist(1-cosine(t(KMPCA_meansig_mat_meansig_mat_reproducible))/2 - cor(t(KMPCA_meansig_mat_meansig_mat_reproducible))/2 ) )
dev.off()
#################################################


#################################################
### reproducible cluster ids & prepare Train data for QDA
reproducible_clusters = cluster_set[select_Kclusters]
reproducible_clusters_rows = is.element(KMPCA_meansig_mat_pca_joint_KM$cluster, reproducible_clusters)
train_data_PCs = KMPCA_meansig_mat_pca_x[reproducible_clusters_rows,1:pca_used_num]
train_data_y = KMPCA_meansig_mat_pca_joint_KM$cluster[reproducible_clusters_rows]
QDA_train_data = cbind(train_data_y, train_data_PCs)
colnames(QDA_train_data)[1] = 'Y'
QDA_cluster_train_data_prior = cCRE_count_KMPCA_iteration_KM_k_reproducible / sum(cCRE_count_KMPCA_iteration_KM_k_reproducible)

### Train QDA
library(MASS)
QDA_model = qda(Y~., data=as.data.frame(QDA_train_data), prior=QDA_cluster_train_data_prior)
#QDA_model = lda(Y~., data=as.data.frame(QDA_train_data), prior=QDA_cluster_train_data_prior)

### predict QDA id for each cCRE
all_cCREs_prediction_Y = predict(QDA_model, newdata = as.data.frame(dhs_ctmerge_shared_reorder_pca$x))$class
#################################################


#################################################
### check cCRE counts
cCRE_id_HM_count = c()
dhs_dms_ctmerge_shared_reorder_meansig_H = c()
dhs_dms_ctmerge_shared_reorder_meansig_M = c()
dhs_dms_ctmerge_shared_reorder_meansig =c()
dhs_dms_ctmerge_shared_reorder_meansig_rowN = c()
for (KM_i in unique(all_cCREs_prediction_Y)){
	cCRE_id_KM_i = cCRE_id[all_cCREs_prediction_Y==KM_i]
	cCRE_id_HM_count = rbind(cCRE_id_HM_count, c(sum(cCRE_id_KM_i=='H'), sum(cCRE_id_KM_i=='M')))
	dhs_dms_ctmerge_shared_reorder_meansig = rbind(dhs_dms_ctmerge_shared_reorder_meansig, colMeans(dhs_dms_ctmerge_shared_reorder[all_cCREs_prediction_Y==KM_i,]))
	dhs_dms_ctmerge_shared_reorder_meansig_rowN = c(dhs_dms_ctmerge_shared_reorder_meansig_rowN, sum(all_cCREs_prediction_Y==KM_i))
	###
	H_rows = cCRE_id=='H'
	dhs_dms_ctmerge_shared_reorder_meansig_H = rbind(dhs_dms_ctmerge_shared_reorder_meansig_H, colMeans(dhs_dms_ctmerge_shared_reorder[H_rows,][all_cCREs_prediction_Y[H_rows]==KM_i,]))
	M_rows = cCRE_id=='M'
	dhs_dms_ctmerge_shared_reorder_meansig_M = rbind(dhs_dms_ctmerge_shared_reorder_meansig_M, colMeans(dhs_dms_ctmerge_shared_reorder[M_rows,][all_cCREs_prediction_Y[M_rows]==KM_i,]))
}
rownames(dhs_dms_ctmerge_shared_reorder_meansig) = unique(all_cCREs_prediction_Y)

### add noise
dhs_dms_ctmerge_shared_reorder_meansig_OD = dhs_dms_ctmerge_shared_reorder_meansig
dhs_dms_ctmerge_shared_reorder_meansig = dhs_dms_ctmerge_shared_reorder_meansig_OD
#dhs_dms_ctmerge_shared_reorder_meansig[dhs_dms_ctmerge_shared_reorder_meansig<0.01] = 0
dhs_dms_ctmerge_shared_reorder_meansig = dhs_dms_ctmerge_shared_reorder_meansig + matrix(runif(dim(dhs_dms_ctmerge_shared_reorder_meansig)[1]*dim(dhs_dms_ctmerge_shared_reorder_meansig)[2], -0.01, 0.01), dim(dhs_dms_ctmerge_shared_reorder_meansig)[1], dim(dhs_dms_ctmerge_shared_reorder_meansig)[2])

hclust_cCRE = hclust(dist(1 - cosine(t(dhs_dms_ctmerge_shared_reorder_meansig))))
#hclust_cCRE = hclust(dist(1 - cosine(t(dhs_dms_ctmerge_shared_reorder_meansig))))
#hclust_cCRE = hclust( dist(1-cosine(t(dhs_dms_ctmerge_shared_reorder_meansig))/2 - cor(t(dhs_dms_ctmerge_shared_reorder_meansig))/2 ) )
#hclust_cCRE_col = hclust(dist(t(dhs_dms_ctmerge_shared_reorder_meansig) ))
#hclust_cCRE_col = hclust(dist(1-cosine((dhs_dms_ctmerge_shared_reorder_meansig))/2 - cor((dhs_dms_ctmerge_shared_reorder_meansig))/2 ))
#################################################


#################################################
### meta-cluster
'''
hclust_cCRE_DTC = cutreeDynamic(hclust_cCRE, minClusterSize=1, deepSplit=4, dist= as.matrix(dist(1 - cosine(t(dhs_dms_ctmerge_shared_reorder_meansig)))), method='hybrid')
table(hclust_cCRE_DTC)
### modified
hclust_cCRE_DTC_modified = hclust_cCRE_DTC
#hclust_cCRE_DTC_modified[hclust_cCRE_DTC==1] = 1
#hclust_cCRE_DTC_modified[hclust_cCRE_DTC==3] = 1
#hclust_cCRE_DTC_modified[hclust_cCRE_DTC==4] = 1
#hclust_cCRE_DTC_modified[hclust_cCRE_DTC==7] = 1
#hclust_cCRE_DTC_modified[hclust_cCRE_DTC==12] = 5


set.seed(2019)
dhs_dms_ctmerge_shared_reorder_meansig_pca = dhs_dms_ctmerge_shared_reorder_meansig %*% dhs_ctmerge_shared_reorder_pca$rotation
hclust_cCRE_DTC_modified = kmeans(dhs_dms_ctmerge_shared_reorder_meansig_pca[,1:pca_used_num], centers=11)$cluster
'''
### check initial KM-K
set.seed(2019)
dhs_dms_ctmerge_shared_reorder_meansig_pca = dhs_dms_ctmerge_shared_reorder_meansig %*% dhs_ctmerge_shared_reorder_pca$rotation
KM_Jmet_KM_ratio_mat = c()
for (iter_i in 1:30){
	KM_Jmet_KM_ratio = c()
for (ini_k in seq(2,20)){
	print(ini_k)
KM_Jmet_KM = kmeans(dhs_dms_ctmerge_shared_reorder_meansig_pca[,1:pca_used_num], centers=ini_k)
	KM_Jmet_KM_ratio = c(KM_Jmet_KM_ratio, KM_Jmet_KM$tot.withinss/KM_Jmet_KM$betweenss)
}
KM_Jmet_KM_ratio_mat = cbind(KM_Jmet_KM_ratio_mat, KM_Jmet_KM_ratio)
}
KM_Jmet_KM_ratio_mat = rowMeans(KM_Jmet_KM_ratio_mat)
pdf("KM_Jmet_KM.determineK.pdf")
plot(seq(2,20), KM_Jmet_KM_ratio_mat)
lines(seq(2,20), KM_Jmet_KM_ratio_mat)
abline(v=15)
dev.off()

###
set.seed(2019)
K = 15
iter_n = 30
dhs_dms_ctmerge_shared_reorder_meansig_pca = dhs_dms_ctmerge_shared_reorder_meansig %*% dhs_ctmerge_shared_reorder_pca$rotation
### run Kmeans 30 times 
KM_by_KM_SameClu_count = matrix(0, dim(dhs_dms_ctmerge_shared_reorder_meansig_pca)[1], dim(dhs_dms_ctmerge_shared_reorder_meansig_pca)[1])
for (iter_i in 1:iter_n){
	print(iter_i)
hclust_cCRE_DTC_modified_iter_i = kmeans(dhs_dms_ctmerge_shared_reorder_meansig_pca[,1:pca_used_num], centers=K)$cluster
for (Ki in 1:K){
	used_rows = which(hclust_cCRE_DTC_modified_iter_i==Ki)
	for (i in 1:length(used_rows)){
		for (j in 1:length(used_rows)){
			KM_by_KM_SameClu_count[used_rows[i],used_rows[j]] = KM_by_KM_SameClu_count[used_rows[i],used_rows[j]]+1
		}
	}
}
}
### count the number runs each row-pair are in the same cluster
colnames(KM_by_KM_SameClu_count) = rownames(dhs_dms_ctmerge_shared_reorder_meansig_pca)
rownames(KM_by_KM_SameClu_count) = rownames(dhs_dms_ctmerge_shared_reorder_meansig_pca)
### plot count in same clusters 
pdf('KM_by_KM_SameClu_count.reproducible.pdf', height=10, width=10)
pheatmap(KM_by_KM_SameClu_count, cex = 1.5, cluster_col=T, cluster_rows=T, clustering_distance_rows=dist(KM_by_KM_SameClu_count), clustering_distance_cols=dist(KM_by_KM_SameClu_count))#, clustering_distance_rows=dist(dhs_dms_ctmerge_shared_reorder_meansig %*% dhs_ctmerge_shared_reorder_pca$rotation[,]) )
dev.off()
### Use cutreeDynamic to cluster KMs
hclust_bySameClu_count = hclust(dist(KM_by_KM_SameClu_count), method='complete')
hclust_cCRE_DTC = cutreeDynamic(hclust_bySameClu_count, minClusterSize=1, deepSplit=4, dist= as.matrix(dist(KM_by_KM_SameClu_count)), method='hybrid')
table(hclust_cCRE_DTC)
#################################################


#################################################
### get Joint meta-cluster ID
hclust_cCRE_DTC_modified = hclust_cCRE_DTC
all_cCREs_prediction_Y_Jmeta = all_cCREs_prediction_Y
class(all_cCREs_prediction_Y_Jmeta) = 'numeric'
KM_i = 0
for (id_KM in unique(all_cCREs_prediction_Y)){
	KM_i = KM_i+1
	print(as.numeric(id_KM))
	all_cCREs_prediction_Y_Jmeta[all_cCREs_prediction_Y==unique(all_cCREs_prediction_Y)[KM_i]] = hclust_cCRE_DTC_modified[KM_i]
}

row_names_combined = paste(hclust_cCRE_DTC_modified, ':', unique(all_cCREs_prediction_Y), sep='')
rownames(dhs_dms_ctmerge_shared_reorder_meansig) = row_names_combined
rownames(dhs_dms_ctmerge_shared_reorder_meansig_H) = row_names_combined
rownames(dhs_dms_ctmerge_shared_reorder_meansig_M) = row_names_combined

pdf('KMPCA.Joint.cluster.reproducible.pdf', height=10)
pheatmap(dhs_dms_ctmerge_shared_reorder_meansig[order(hclust_cCRE_DTC_modified),], cex = 1.5, cluster_col=T, cluster_rows=F, clustering_distance_cols = dist(t(dhs_dms_ctmerge_shared_reorder_meansig)), clustering_distance_rows=dist(1 - cosine(t(dhs_dms_ctmerge_shared_reorder_meansig))) )#, clustering_distance_rows=dist(dhs_dms_ctmerge_shared_reorder_meansig %*% dhs_ctmerge_shared_reorder_pca$rotation[,]) )
dev.off()

pdf('KMPCA.Joint.cluster.reproducible_H.pdf', height=10)
pheatmap(dhs_dms_ctmerge_shared_reorder_meansig_H[order(hclust_cCRE_DTC_modified),], cluster_col=T, cluster_rows=F, cutree_rows=20, clustering_distance_cols = dist(t(dhs_dms_ctmerge_shared_reorder_meansig)), clustering_distance_rows=dist(1 - cosine(t(dhs_dms_ctmerge_shared_reorder_meansig))) )#, clustering_distance_cols = dist(1-cosine(dhs_dms_ctmerge_shared_reorder_meansig)/2 - cor(dhs_dms_ctmerge_shared_reorder_meansig)/2), cclustering_distance_rows=dist(1-cosine(t(dhs_dms_ctmerge_shared_reorder_meansig))/2 - cor(t(dhs_dms_ctmerge_shared_reorder_meansig))/2 ) )
dev.off()

pdf('KMPCA.Joint.cluster.reproducible_M.pdf', height=10)
pheatmap(dhs_dms_ctmerge_shared_reorder_meansig_M[order(hclust_cCRE_DTC_modified),], cluster_col=T, cluster_rows=F, cutree_rows=20, clustering_distance_cols = dist(t(dhs_dms_ctmerge_shared_reorder_meansig)), clustering_distance_rows=dist(1 - cosine(t(dhs_dms_ctmerge_shared_reorder_meansig))) )#, clustering_distance_cols = dist(1-cosine(dhs_dms_ctmerge_shared_reorder_meansig)/2 - cor(dhs_dms_ctmerge_shared_reorder_meansig)/2), cclustering_distance_rows=dist(1-cosine(t(dhs_dms_ctmerge_shared_reorder_meansig))/2 - cor(t(dhs_dms_ctmerge_shared_reorder_meansig))/2 ) )
dev.off()

png('cCRE_id_HM_count.png')
plot_lim = c(5, 50000)
plot(cCRE_id_HM_count[,1],cCRE_id_HM_count[,2]/mean(cCRE_id_HM_count[,2])*mean(cCRE_id_HM_count[,1]), log='xy', xlim = plot_lim, ylim=plot_lim, xlab='H', ylab='M')
abline(0,1)
dev.off()

### get Jmet meansignal mat
dhs_dms_ctmerge_shared_reorder_meansig_Jmet_meansig = c()
for (i in unique(hclust_cCRE_DTC_modified)){
if (sum(hclust_cCRE_DTC_modified==i)>1){dhs_dms_ctmerge_shared_reorder_meansig_Jmet_meansig = rbind(dhs_dms_ctmerge_shared_reorder_meansig_Jmet_meansig, colMeans(dhs_dms_ctmerge_shared_reorder_meansig[hclust_cCRE_DTC_modified==i,]))}
if (sum(hclust_cCRE_DTC_modified==i)==1){dhs_dms_ctmerge_shared_reorder_meansig_Jmet_meansig = rbind(dhs_dms_ctmerge_shared_reorder_meansig_Jmet_meansig, dhs_dms_ctmerge_shared_reorder_meansig[hclust_cCRE_DTC_modified==i,])}
}
rownames(dhs_dms_ctmerge_shared_reorder_meansig_Jmet_meansig) = unique(hclust_cCRE_DTC_modified)
###
Jmet_order = as.numeric(rownames(dhs_dms_ctmerge_shared_reorder_meansig_Jmet_meansig)[hclust(dist(dhs_dms_ctmerge_shared_reorder_meansig_Jmet_meansig))$order])
pdf('KMPCA.Joint.cluster.reproducible.Jmet.pdf', height=5)
pheatmap(dhs_dms_ctmerge_shared_reorder_meansig_Jmet_meansig, cluster_col=T, cluster_rows=T, clustering_distance_cols = dist(t(dhs_dms_ctmerge_shared_reorder_meansig)), cex=1.5)#, clustering_distance_rows=dist(dhs_dms_ctmerge_shared_reorder_meansig %*% dhs_ctmerge_shared_reorder_pca$rotation[,]) )
dev.off()
#################################################


#################################################
### write table
dh_with_JointClusterID_mat = cbind(dh[,c(1:6)], all_cCREs_prediction_Y_Jmeta[H_rows], dh[,-c(1:6)])
dm_with_JointClusterID_mat = cbind(dm[,c(1:6)], all_cCREs_prediction_Y_Jmeta[M_rows], dm[,-c(1:6)])
colnames(dh_with_JointClusterID_mat)[7] = 'J_meta'
colnames(dm_with_JointClusterID_mat)[7] = 'J_meta'
write.table(dh_with_JointClusterID_mat, 'S3V2_IDEAS_hg38_ccre2.cCRE.M.notall0.rmallNEU.withid.coe_mat.PDmerged.clusterID.JclusterID.txt', quote=F, sep='\t', col.names=T, row.names=F)
write.table(dm_with_JointClusterID_mat, 'S3V2_IDEAS_mm10_ccre2.cCRE.M.notall0.withid.coe_mat.PDmerged.clusterID.JclusterID.txt', quote=F, sep='\t', col.names=T, row.names=F)
###### ct esRP mat for RNA correlation analysis
dh_with_JointClusterID_mat_ct = cbind(apply(cbind(dh[,1], as.character(dh[,2]), as.character(dh[,3])) ,1, function(x) paste(x, collapse='_')), dh[,c(1:6)], all_cCREs_prediction_Y_Jmeta[H_rows], dhs_ctmerge_shared_reorder)
dm_with_JointClusterID_mat_ct = cbind(apply(cbind(dm[,1], as.character(dm[,2]), as.character(dm[,3])) ,1, function(x) paste(x, collapse='_')), dm[,c(1:6)], all_cCREs_prediction_Y_Jmeta[M_rows], dms_ctmerge_shared_reorder)
###### check JMeta number
### hg38
JMeta_vs_KM_H = t(apply(cbind(rownames(dhs_dms_ctmerge_shared_reorder_meansig_H)), 1, function(x) as.numeric(unlist(strsplit(x, ':'))) ))
JMeta_vs_KM_M = t(apply(cbind(rownames(dhs_dms_ctmerge_shared_reorder_meansig_M)), 1, function(x) as.numeric(unlist(strsplit(x, ':'))) ))
JMeta_count_H = c()
JMeta_count_M = c()
for (JMeta_i in unique(JMeta_vs_KM_H[,1])){
	table_vec_names = as.numeric(names(table(dh_with_JointClusterID_mat$J_meta)))
	JMeta_vs_KM_H_i = JMeta_vs_KM_H[is.element(JMeta_vs_KM_H[,1], JMeta_i),2]
	JMeta_vs_KM_M_i = JMeta_vs_KM_M[is.element(JMeta_vs_KM_M[,1], JMeta_i),2]
	print(c(JMeta_i, JMeta_vs_KM_H_i))
	print(c(JMeta_i, JMeta_vs_KM_M_i))
	JMeta_count_H = c(JMeta_count_H, sum(table(dh_with_JointClusterID_mat$J_meta)[is.element(table_vec_names, JMeta_vs_KM_H_i)]))
	JMeta_count_M = c(JMeta_count_M, sum(table(dm_with_JointClusterID_mat$J_meta)[is.element(table_vec_names, JMeta_vs_KM_M_i)]))
}
JMeta_count_H = cbind(unique(JMeta_vs_KM_H[,1]), JMeta_count_H)
JMeta_count_M = cbind(unique(JMeta_vs_KM_M[,1]), JMeta_count_M)
###
JMeta_count_HM = cbind(JMeta_count_H[,2], JMeta_count_M[,2])
JMeta_count_HM[,2] = JMeta_count_HM[,2]/mean(JMeta_count_HM[,2])*mean(JMeta_count_HM[,1])
rownames(JMeta_count_HM) = JMeta_count_H[,1]
colnames(JMeta_count_HM) = c('Human','Mouse')
pdf('KMPCA.Joint.cluster.reproducible.Jmet.cCRE_count.pdf', height=5,width=3)
pheatmap(log10(JMeta_count_HM), cluster_col=F, cluster_rows=T, cex=1.5, clustering_distance_rows = dist(dhs_dms_ctmerge_shared_reorder_meansig_Jmet_meansig))#, clustering_distance_rows=dist(dhs_dms_ctmerge_shared_reorder_meansig %*% dhs_ctmerge_shared_reorder_pca$rotation[,]) )
dev.off()
#################################################


#################################################
### gene gene locus
dh_with_JointClusterID = cbind(dh[,c(1:3)], all_cCREs_prediction_Y_Jmeta[H_rows])
dm_with_JointClusterID = cbind(dm[,c(1:3)], all_cCREs_prediction_Y_Jmeta[M_rows])
JointCluster_count_HM = cbind(table(all_cCREs_prediction_Y_Jmeta[H_rows]), table(all_cCREs_prediction_Y_Jmeta[M_rows]))
JointCluster_exp_P_H = JointCluster_count_HM[,1]/sum(JointCluster_count_HM[,1])
JointCluster_exp_P_M = JointCluster_count_HM[,2]/sum(JointCluster_count_HM[,2])
All_Jmet = as.numeric(names(table(all_cCREs_prediction_Y_Jmeta)))

###
get_Function_conserve_cCRE = function(chrH, startH, endH, chrM, startM, endM, Jmet_enrichment_scores_all_zpfdr_thresh){
#chrH = 'chrX'
#startH = 48760001
#endH = 48836000
#chrM = 'chrX'
#startM = 7919401
#endM = 8020800
add_sm_num = 1
included_rows = ((dh_with_JointClusterID[,1]==chrH) * (dh_with_JointClusterID[,2]>=startH) * (dh_with_JointClusterID[,3]<=endH)) != 0
dh_with_JointClusterID_Gene = dh_with_JointClusterID[included_rows,]
included_rows = ((dm_with_JointClusterID[,1]==chrM) * (dm_with_JointClusterID[,2]>=startM) * (dm_with_JointClusterID[,3]<=endM)) != 0
dm_with_JointClusterID_Gene = dm_with_JointClusterID[included_rows,]
#print(dh_with_JointClusterID_GATA1)
#print(dm_with_JointClusterID_GATA1)
### get share state
### expect state count
JointCluster_count_H_exp = (dim(dh_with_JointClusterID_Gene)[1])*JointCluster_exp_P_H
JointCluster_count_M_exp = (dim(dm_with_JointClusterID_Gene)[1])*JointCluster_exp_P_M
JointCluster_count_HM_exp = (JointCluster_count_H_exp+add_sm_num)*(JointCluster_count_M_exp+add_sm_num)
#print(JointCluster_count_HM_exp)
#print(JointCluster_count_H_exp)
#print(JointCluster_count_M_exp)
#print(dim(dh_with_JointClusterID_Gene)[1])
#print(dim(dm_with_JointClusterID_Gene)[1])
### obs state count
JointCluster_count_H_obs = c()
JointCluster_count_M_obs = c()
for (i in All_Jmet){
JointCluster_count_H_obs = c(JointCluster_count_H_obs, sum(dh_with_JointClusterID_Gene[,4]==i))
JointCluster_count_M_obs = c(JointCluster_count_M_obs, sum(dm_with_JointClusterID_Gene[,4]==i))
}
### get odd ratio
#gene_exp = cbind((JointCluster_count_H_obs+add_sm_num)/(JointCluster_count_H_exp+add_sm_num), (JointCluster_count_M_obs+add_sm_num)/(JointCluster_count_M_exp+add_sm_num), ((JointCluster_count_H_obs+add_sm_num)*(JointCluster_count_M_obs+add_sm_num)) / ((JointCluster_count_H_exp+add_sm_num)*(JointCluster_count_M_exp+add_sm_num)) )
#gene_exp = ((JointCluster_count_H_obs+add_sm_num)*(JointCluster_count_M_obs+add_sm_num)) / ((JointCluster_count_H_exp+add_sm_num)*(JointCluster_count_M_exp+add_sm_num))
gene_exp = ((JointCluster_count_H_obs+add_sm_num)*(JointCluster_count_M_obs+add_sm_num)) / JointCluster_count_HM_exp
#print(dh_with_JointClusterID_GATA1)
#print(dm_with_JointClusterID_GATA1)
#print(gene_exp)
Jmet = list()
Jmet$Jmet_score = gene_exp
#Jmet$cCRE_H = cbind(dh_with_JointClusterID_GATA1[is.element(dh_with_JointClusterID_GATA1[,4], dm_with_JointClusterID_GATA1[,4]),1:3])
#Jmet$cCRE_M = cbind(dm_with_JointClusterID_GATA1[is.element(dm_with_JointClusterID_GATA1[,4], dh_with_JointClusterID_GATA1[,4]),1:3])
#Jmet$cCRE_H = cbind(dh_with_JointClusterID_Gene[is.element(dh_with_JointClusterID_Gene[,4], as.numeric(names(gene_exp)[gene_exp>=2])),])
#Jmet$cCRE_M = cbind(dm_with_JointClusterID_Gene[is.element(dm_with_JointClusterID_Gene[,4], as.numeric(names(gene_exp)[gene_exp>=2])),])
#Jmet_enrichment_scores_all_zpfdr_thresh = 2.922063
Jmet$cCRE_H = cbind(dh_with_JointClusterID_Gene, is.element(dh_with_JointClusterID_Gene[,4], as.numeric(names(gene_exp)[gene_exp>=Jmet_enrichment_scores_all_zpfdr_thresh]))*1)
Jmet$cCRE_M = cbind(dm_with_JointClusterID_Gene, is.element(dm_with_JointClusterID_Gene[,4], as.numeric(names(gene_exp)[gene_exp>=Jmet_enrichment_scores_all_zpfdr_thresh]))*1)
return(Jmet)
}
#################################################


#################################################
### read gene pairs
hg38_gene = read.table('/Users/guanjuexiang/Documents/projects/analysis/0813_human_mouse_state_compare_heatmap/hg38.gene.bed', header=F, sep='\t')
mm10_gene = read.table('/Users/guanjuexiang/Documents/projects/analysis/0813_human_mouse_state_compare_heatmap/mm10.gene.bed', header=F, sep='\t')
### mm10
### change Gata1 gene locus from
# chrX	7959260	7978071	-	Gata1
# to
# chrX	7959260	7967910	-	Gata1 
mm10_gene[,5] = toupper(mm10_gene[,5])
###
shared_genes = hg38_gene[is.element(hg38_gene[,5], mm10_gene[,5]),5]
### get input mat
hg38_gene_shared = hg38_gene[is.element(hg38_gene[,5], shared_genes),]
hg38_gene_shared = hg38_gene_shared[order(hg38_gene_shared[,5]),]
mm10_gene_shared = mm10_gene[is.element(mm10_gene[,5], shared_genes),]
mm10_gene_shared = mm10_gene_shared[order(mm10_gene_shared[,5]),]
### duplicate genes
duplicated_genes = unique(c(names(table(hg38_gene_shared[,5])[table(hg38_gene_shared[,5])>1]), names(table(mm10_gene_shared[,5])[table(mm10_gene_shared[,5])>1])))
### remove duplicate genes
shared_genes = shared_genes[!is.element(shared_genes, duplicated_genes)]
hg38_gene_shared = hg38_gene_shared[!is.element(hg38_gene_shared[,5], duplicated_genes),]
mm10_gene_shared = mm10_gene_shared[!is.element(mm10_gene_shared[,5], duplicated_genes),]
### order genes
shared_genes = shared_genes[order(shared_genes)]
hg38_gene_shared = hg38_gene_shared[order(hg38_gene_shared[,5]),]
mm10_gene_shared = mm10_gene_shared[order(mm10_gene_shared[,5]),]
### expand 50KB
hg38_gene_shared_exp = hg38_gene_shared
hg38_gene_shared_exp[hg38_gene_shared_exp[,4]=='+',2] = hg38_gene_shared_exp[hg38_gene_shared_exp[,4]=='+',2] - 50000
hg38_gene_shared_exp[hg38_gene_shared_exp[,4]=='+',3] = hg38_gene_shared_exp[hg38_gene_shared_exp[,4]=='+',2] + 100000
hg38_gene_shared_exp[hg38_gene_shared_exp[,4]=='-',2] = hg38_gene_shared_exp[hg38_gene_shared_exp[,4]=='-',3] - 50000
hg38_gene_shared_exp[hg38_gene_shared_exp[,4]=='-',3] = hg38_gene_shared_exp[hg38_gene_shared_exp[,4]=='-',3] + 50000
hg38_gene_shared_exp[hg38_gene_shared_exp[,2]<0,2] = 0
###
mm10_gene_shared_exp = mm10_gene_shared
mm10_gene_shared_exp[mm10_gene_shared_exp[,4]=='+',2] = mm10_gene_shared_exp[mm10_gene_shared_exp[,4]=='+',2] - 50000
mm10_gene_shared_exp[mm10_gene_shared_exp[,4]=='+',3] = mm10_gene_shared_exp[mm10_gene_shared_exp[,4]=='+',2] + 100000
mm10_gene_shared_exp[mm10_gene_shared_exp[,4]=='-',2] = mm10_gene_shared_exp[mm10_gene_shared_exp[,4]=='-',3] - 50000
mm10_gene_shared_exp[mm10_gene_shared_exp[,4]=='-',3] = mm10_gene_shared_exp[mm10_gene_shared_exp[,4]=='-',3] + 50000
mm10_gene_shared_exp[mm10_gene_shared_exp[,2]<0,2] = 0
###
#hg38_mm10_gene_shared_exp = cbind(hg38_gene_shared_exp[,1:3], mm10_gene_shared_exp[,1:3])
#################################################


#################################################
### get Gene-by-Jmeta matrix: Round1: Determine enrichment threshold of each Jmet cluster
GATA1_exp = get_Function_conserve_cCRE('chrX', 48760001, 48836000, 'chrX', 7919401,8020800, 2.922063)
meta_cluster_mat = as.data.frame(matrix(0, nrow=length(shared_genes), ncol=length(GATA1_exp$Jmet_score)))
cCRE_H_Jmeta = c()
cCRE_M_Jmeta = c()
ptm <- proc.time()
for (i in 1:length(shared_genes)){
	if (i%%1000==0){print(i)}
	shared_genes_i = shared_genes[i]
	bed_H = hg38_gene_shared_exp[i,1:3][1,]
	bed_M = mm10_gene_shared_exp[i,1:3][1,]
	JMeta_i = get_Function_conserve_cCRE(bed_H[1,1], bed_H[1,2], bed_H[1,3], bed_M[1,1], bed_M[1,2], bed_M[1,3], 2.922063)
	meta_cluster_mat[i,] = JMeta_i$Jmet_score
	if (dim(JMeta_i$cCRE_H)[1]>0){cCRE_H_Jmeta = rbind(cCRE_H_Jmeta, cbind(JMeta_i$cCRE_H, shared_genes_i))}
	if (dim(JMeta_i$cCRE_M)[1]>0){cCRE_M_Jmeta = rbind(cCRE_M_Jmeta, cbind(JMeta_i$cCRE_M, shared_genes_i))}
}
proc.time() - ptm
###
colnames(meta_cluster_mat) = rownames(JointCluster_count_HM)
rownames(meta_cluster_mat) = shared_genes
### get enrichment threshold for each Jmet
get_thresh = function(x){
	xzp = pnorm((x-mean(x))/sd(x), lower.tail=F)
	thresh0 = 0.05
	thresh1 = 0.01
	xzp = pnorm((x-mean(x[xzp>=thresh1]))/sd(x[xzp>=thresh1]), lower.tail=F)
	return(min(x[xzp<thresh0]))
}
meta_cluster_mat_thresh = apply(meta_cluster_mat,2,get_thresh)
meta_cluster_mat_thresh

for (i in 1:dim(meta_cluster_mat)[2]){
png(paste0('Enrichment.hist.',i,'.png'))
hist(meta_cluster_mat[,i], breaks=50)
dev.off()
}
#################################################


#################################################
### get Gene-by-Jmeta matrix: Round2
GATA1_exp = get_Function_conserve_cCRE('chrX', 48760001, 48836000, 'chrX', 7919401,8020800, meta_cluster_mat_thresh)
meta_cluster_mat = as.data.frame(matrix(0, nrow=length(shared_genes), ncol=length(GATA1_exp$Jmet_score)))
cCRE_H_Jmeta = c()
cCRE_M_Jmeta = c()
### change some genes' TSS
hg38_gene_shared_exp[hg38_gene_shared_exp[,5]=='CSF1R',2:3] = c(150036554, 150136554)
###
ptm <- proc.time()
for (i in 1:length(shared_genes)){
	if (i%%1000==0){print(i)}
	shared_genes_i = shared_genes[i]
	bed_H = hg38_gene_shared_exp[i,1:3][1,]
	bed_M = mm10_gene_shared_exp[i,1:3][1,]
	JMeta_i = get_Function_conserve_cCRE(bed_H[1,1], bed_H[1,2], bed_H[1,3], bed_M[1,1], bed_M[1,2], bed_M[1,3], meta_cluster_mat_thresh)
	meta_cluster_mat[i,] = JMeta_i$Jmet_score
	if (dim(JMeta_i$cCRE_H)[1]>0){cCRE_H_Jmeta = rbind(cCRE_H_Jmeta, cbind(JMeta_i$cCRE_H, shared_genes_i))}
	if (dim(JMeta_i$cCRE_M)[1]>0){cCRE_M_Jmeta = rbind(cCRE_M_Jmeta, cbind(JMeta_i$cCRE_M, shared_genes_i))}
}
proc.time() - ptm
###
colnames(meta_cluster_mat) = rownames(JointCluster_count_HM)
rownames(meta_cluster_mat) = shared_genes
write.table(cbind(shared_genes, round(meta_cluster_mat,5)), 'Human_Mouse_shared_genes.Jmeta.enrich.txt', quote=F, sep='\t', col.names=T, row.names=F)
write.table(cCRE_H_Jmeta, 'cCRE.Gene50KB.hg38.bed', quote=F, sep='\t', col.names=T, row.names=F)
write.table(cCRE_M_Jmeta, 'cCRE.Gene50KB.mm10.bed', quote=F, sep='\t', col.names=T, row.names=F)
#################################################
chrX 48783000 48783200


#################################################
### cluster gene by Jmet Joint-Enrichment
meta_cluster_mat_log2 = log2(meta_cluster_mat+1)
### decide K
set.seed(2019)
meta_cluster_mat_log2_km_ratio = c()
for (k in 2:30){
	meta_cluster_mat_log2_km_test = kmeans(meta_cluster_mat_log2, centers=k)
	meta_cluster_mat_log2_km_ratio = c(meta_cluster_mat_log2_km_ratio, meta_cluster_mat_log2_km_test$tot.withinss/meta_cluster_mat_log2_km_test$betweenss)
}
pdf('KM_gene_Jmet_enrich.determineK.pdf')
plot(2:30, meta_cluster_mat_log2_km_ratio, cex.axis=2)
lines(2:30, meta_cluster_mat_log2_km_ratio)
abline(v=15)
dev.off()
### KM cluster gene
set.seed(2019)
meta_cluster_mat_log2_km = kmeans(meta_cluster_mat_log2, centers=15)
meta_cluster_mat_log2_km_mean = c()
for (i in as.numeric(names(table(meta_cluster_mat_log2_km$cluster)))){
	meta_cluster_mat_log2_km_mean = rbind(meta_cluster_mat_log2_km_mean, colMeans(meta_cluster_mat_log2[meta_cluster_mat_log2_km$cluster==i,])) 
}
rownames(meta_cluster_mat_log2_km_mean) = as.numeric(names(table(meta_cluster_mat_log2_km$cluster)))
### write gene-by-Jmet-enrichment matrix with Gene KM-ID
meta_cluster_mat_log2_df = cbind(as.data.frame(shared_genes), meta_cluster_mat_log2_km$cluster, meta_cluster_mat_log2)
colnames(meta_cluster_mat_log2_df)[2] = 'GeneKM_ID'
pdf('meta_cluster_mat_log2_df.hist.pdf')
Jmet_enrichment_scores_all = c(as.matrix(meta_cluster_mat_log2_df[,-c(1:2)]))
Jmet_enrichment_scores_all_zpfdr = p.adjust(pnorm((Jmet_enrichment_scores_all - mean(Jmet_enrichment_scores_all)) / sd(Jmet_enrichment_scores_all), lower.tail=F), 'fdr')
Jmet_enrichment_scores_all_zpfdr = p.adjust(pnorm((Jmet_enrichment_scores_all - mean(Jmet_enrichment_scores_all[Jmet_enrichment_scores_all_zpfdr>=0.1])) / sd(Jmet_enrichment_scores_all[Jmet_enrichment_scores_all_zpfdr>=0.1]), lower.tail=F), 'fdr')
Jmet_enrichment_scores_all_zpfdr_thresh = min(Jmet_enrichment_scores_all[Jmet_enrichment_scores_all_zpfdr<0.1])
hist(Jmet_enrichment_scores_all, breaks=50, cex.axis=2)
abline(v=Jmet_enrichment_scores_all_zpfdr_thresh)
box()
dev.off()
write.table(meta_cluster_mat_log2_df, 'Human_Mouse_shared_genes.Jmeta.enrich.txt', quote=F, sep='\t', col.names=T, row.names=F)
#################################################


#################################################
### plot heatmap
png('GeneGroup_by_JMeta.Joint.cluster.png', width = 1000, height = 600)
meta_cluster_mat_GeneGroup_Jmet = c()
used_order = Jmet_order
for (coli in used_order){
	meta_cluster_mat_GeneGroup_Jmet = cbind(meta_cluster_mat_GeneGroup_Jmet, meta_cluster_mat_log2_km_mean[,colnames(meta_cluster_mat)==coli])
}
colnames(meta_cluster_mat_GeneGroup_Jmet) = used_order
rownames(meta_cluster_mat_GeneGroup_Jmet) = rownames(meta_cluster_mat_log2_km_mean)
pheatmap((t(meta_cluster_mat_GeneGroup_Jmet)), cex=2, cluster_col=F, cluster_rows=F, show_rownames=T, show_colnames=T, clustering_distance_rows = dist(dhs_dms_ctmerge_shared_reorder_meansig_Jmet_meansig), cex=1.5)
dev.off()
### get binary enrichment based on GeneGroup
meta_cluster_mat_GeneGroup_Jmet_vec = c(meta_cluster_mat_GeneGroup_Jmet)
for (i in 1:10){
	print(length(meta_cluster_mat_GeneGroup_Jmet_vec))
#meta_cluster_mat_GeneGroup_Jmet_vec_zp_fdr = p.adjust(pnorm((meta_cluster_mat_GeneGroup_Jmet_vec - mean(meta_cluster_mat_GeneGroup_Jmet_vec)) / sd(meta_cluster_mat_GeneGroup_Jmet_vec), lower.tail=F), 'fdr')
#meta_cluster_mat_GeneGroup_Jmet_vec = meta_cluster_mat_GeneGroup_Jmet_vec[meta_cluster_mat_GeneGroup_Jmet_vec_zp_fdr>=0.1]
meta_cluster_mat_GeneGroup_Jmet_vec_zp_fdr = pnorm((meta_cluster_mat_GeneGroup_Jmet_vec - mean(meta_cluster_mat_GeneGroup_Jmet_vec)) / sd(meta_cluster_mat_GeneGroup_Jmet_vec), lower.tail=F)
meta_cluster_mat_GeneGroup_Jmet_vec = meta_cluster_mat_GeneGroup_Jmet_vec[meta_cluster_mat_GeneGroup_Jmet_vec_zp_fdr>=0.001]
}
### get Jmet enrichment for each GeneGroup for Downstream analysis
meta_cluster_mat_GeneGroup_Jmet_z = (meta_cluster_mat_GeneGroup_Jmet - mean(meta_cluster_mat_GeneGroup_Jmet_vec)) / sd(meta_cluster_mat_GeneGroup_Jmet_vec)
meta_cluster_mat_GeneGroup_Jmet_zp_01 = t((apply(meta_cluster_mat_GeneGroup_Jmet_z, 2, function(x) pnorm(x, lower.tail=F)) < 0.001) * 1)
meta_cluster_mat_GeneGroup_Jmet_zp_01
### 
png('Gene_by_JMeta.Joint.cluster.png', width = 1000, height = 600)
meta_cluster_mat_plot = c()
used_order = Jmet_order
for (coli in used_order){
	meta_cluster_mat_plot = cbind(meta_cluster_mat_plot, meta_cluster_mat[,colnames(meta_cluster_mat)==coli])
}
colnames(meta_cluster_mat_plot) = used_order
rownames(meta_cluster_mat_plot) = rownames(meta_cluster_mat)
meta_cluster_mat_plot = meta_cluster_mat_plot[order(meta_cluster_mat_log2_km$cluster),]
pheatmap(log2(t(meta_cluster_mat_plot)+1), cex=2, cluster_col=F, cluster_rows=F, show_rownames=T, show_colnames=F, clustering_distance_rows = dist(dhs_dms_ctmerge_shared_reorder_meansig_Jmet_meansig))
dev.off()
#################################################


#################################################
### prepare SFN files
bash1 = 'mkdir /Users/guanjuexiang/Documents/projects/analysis/04_05_2022_coe/coe_analysis/SF_S_N/'
system(bash1)
### get cCRE with H/MIDs
bash2 = 'cat /Users/guanjuexiang/Documents/projects/analysis/04_05_2022_coe/coe_analysis/SF_S_N/mouse_ccre_mm10_sf.bed /Users/guanjuexiang/Documents/projects/analysis/04_05_2022_coe/coe_analysis/SF_S_N/mouse_ccre_mm10_s.bed /Users/guanjuexiang/Documents/projects/analysis/04_05_2022_coe/coe_analysis/SF_S_N/mouse_ccre_mm10_n.bed | sort -k1,1 -k2,2n > /Users/guanjuexiang/Documents/projects/analysis/04_05_2022_coe/coe_analysis/SF_S_N/mouse_ccre_mm10.withMID.bed'
system(bash2)
bash3 = 'cat /Users/guanjuexiang/Documents/projects/analysis/04_05_2022_coe/coe_analysis/SF_S_N/mouse_ccre_hg38_sf.bed /Users/guanjuexiang/Documents/projects/analysis/04_05_2022_coe/coe_analysis/SF_S_N/mouse_ccre_hg38_s.bed | awk -F \'\t\' -v OFS=\'\t\' \'{print $2,$3,$4,$1}\' | sort -k1,1 -k2,2n > /Users/guanjuexiang/Documents/projects/analysis/04_05_2022_coe/coe_analysis/SF_S_N/mouse_ccre_hg38_sf_s.withMID.bed'
system(bash3)
bash4 = 'cat /Users/guanjuexiang/Documents/projects/analysis/04_05_2022_coe/coe_analysis/SF_S_N/human_ccre_hg38_sf.bed /Users/guanjuexiang/Documents/projects/analysis/04_05_2022_coe/coe_analysis/SF_S_N/human_ccre_hg38_s.bed /Users/guanjuexiang/Documents/projects/analysis/04_05_2022_coe/coe_analysis/SF_S_N/human_ccre_hg38_n.bed | sort -k1,1 -k2,2n > /Users/guanjuexiang/Documents/projects/analysis/04_05_2022_coe/coe_analysis/SF_S_N/human_ccre_hg38.withHID.bed'
system(bash4)
bash5 = 'cat /Users/guanjuexiang/Documents/projects/analysis/04_05_2022_coe/coe_analysis/SF_S_N/human_ccre_mm10_sf.bed /Users/guanjuexiang/Documents/projects/analysis/04_05_2022_coe/coe_analysis/SF_S_N/human_ccre_mm10_s.bed | awk -F \'\t\' -v OFS=\'\t\' \'{print $2,$3,$4,$1}\' | sort -k1,1 -k2,2n > /Users/guanjuexiang/Documents/projects/analysis/04_05_2022_coe/coe_analysis/SF_S_N/human_ccre_mm10_sf_s.withHID.bed'
system(bash5)
### add both H & M IDs
bash6 = 'bedtools map -a /Users/guanjuexiang/Documents/projects/analysis/04_05_2022_coe/coe_analysis/SF_S_N/human_ccre_hg38.withHID.bed -b /Users/guanjuexiang/Documents/projects/analysis/04_05_2022_coe/coe_analysis/SF_S_N/mouse_ccre_hg38_sf_s.withMID.bed -c 4 -o concat -null NA > /Users/guanjuexiang/Documents/projects/analysis/04_05_2022_coe/coe_analysis/SF_S_N/human_ccre_hg38.withHID.with_Mouse_S_MID.bed'
system(bash6)
bash7 = 'bedtools map -a /Users/guanjuexiang/Documents/projects/analysis/04_05_2022_coe/coe_analysis/SF_S_N/mouse_ccre_mm10.withMID.bed -b /Users/guanjuexiang/Documents/projects/analysis/04_05_2022_coe/coe_analysis/SF_S_N/human_ccre_mm10_sf_s.withHID.bed -c 4 -o concat -null NA > /Users/guanjuexiang/Documents/projects/analysis/04_05_2022_coe/coe_analysis/SF_S_N/mouse_ccre_mm10.withMID.with_Human_S_HID.bed'
system(bash7)
#################################################


#################################################
######################## hg38
### Intersect with SF_S cCREs hg38
bedtools1 = 'cat cCRE.Gene50KB.hg38.bed | awk -F \'\t\' \'{if ($1!="NA" && $1!="chr") print $0}\' | sort -k1,1 -k2,2n > cCRE.Gene50KB.hg38.sort.bed'
system(bedtools1)
bedtools2a = 'bedtools map -a cCRE.Gene50KB.hg38.sort.bed -b SF_S_N/human_ccre_hg38.withHID.bed -c 4 -o concat -null NA > cCRE.Gene50KB.hg38.SF_S.HID.bed'
system(bedtools2a)
bedtools2b = 'bedtools map -a cCRE.Gene50KB.hg38.sort.bed -b SF_S_N/human_ccre_hg38.withHID.with_Mouse_S_MID.bed -c 5 -o concat -null NA > cCRE.Gene50KB.hg38.SF_S.MID.bed'
system(bedtools2b)
bedtools2c = 'paste cCRE.Gene50KB.hg38.SF_S.HID.bed cCRE.Gene50KB.hg38.SF_S.MID.bed | cut -f1,2,3,4,5,6,7,14 > cCRE.Gene50KB.hg38.SF_S.bed'
system(bedtools2c)
### Intersect with SF_S cCREs mm10
bedtools3 = 'cat cCRE.Gene50KB.mm10.bed | awk -F \'\t\' \'{if ($1!="NA" && $1!="chr") print $0}\' | sort -k1,1 -k2,2n > cCRE.Gene50KB.mm10.sort.bed '
system(bedtools3)
bedtools4 = 'bedtools map -a cCRE.Gene50KB.mm10.sort.bed -b SF_S_N/mouse_ccre_mm10.withMID.with_Human_S_HID.bed -c 4 -o concat -null NA > cCRE.Gene50KB.mm10.withMID.bed'
system(bedtools4)
bedtools5 = 'cat cCRE.Gene50KB.hg38.SF_S.bed | awk \'{if ($6=="GATA1") print $0}\''
bedtools6 = 'cat cCRE.Gene50KB.mm10.withMID.bed | awk \'{if ($6=="GATA1") print $0}\''
system(bedtools5)
system(bedtools6)
######################## mm10
### Intersect with SF_S cCREs mm10
bedtools1 = 'cat cCRE.Gene50KB.mm10.bed | awk -F \'\t\' \'{if ($1!="NA" && $1!="chr") print $0}\' | sort -k1,1 -k2,2n > cCRE.Gene50KB.mm10.sort.bed'
system(bedtools1)
#bedtools2 = 'bedtools map -a cCRE.Gene50KB.mm10.sort.bed -b SF_S_N/mouse_ccre_mm10.withMID.with_Human_S_HID.bed -c 5 -o concat -null NA > cCRE.Gene50KB.mm10.SF_S.bed'
#system(bedtools2)
bedtools2a = 'bedtools map -a cCRE.Gene50KB.mm10.sort.bed -b SF_S_N/mouse_ccre_mm10.withMID.bed -c 4 -o concat -null NA > cCRE.Gene50KB.mm10.SF_S.MID.bed'
system(bedtools2a)
bedtools2b = 'bedtools map -a cCRE.Gene50KB.mm10.sort.bed -b SF_S_N/mouse_ccre_mm10.withMID.with_Human_S_HID.bed -c 5 -o concat -null NA > cCRE.Gene50KB.mm10.SF_S.HID.bed'
system(bedtools2b)
bedtools2c = 'paste cCRE.Gene50KB.mm10.SF_S.MID.bed cCRE.Gene50KB.mm10.SF_S.HID.bed | cut -f1,2,3,4,5,6,7,14 > cCRE.Gene50KB.mm10.SF_S.bed'
system(bedtools2c)
### Intersect with SF_S cCREs hg38
bedtools3 = 'cat cCRE.Gene50KB.hg38.bed | awk -F \'\t\' \'{if ($1!="NA" && $1!="chr") print $0}\' | sort -k1,1 -k2,2n > cCRE.Gene50KB.hg38.sort.bed '
system(bedtools3)
bedtools4 = 'bedtools map -a cCRE.Gene50KB.hg38.sort.bed -b SF_S_N/human_ccre_hg38.withHID.with_Mouse_S_MID.bed -c 4 -o concat -null NA > cCRE.Gene50KB.hg38.withMID.bed'
system(bedtools4)
bedtools5 = 'cat cCRE.Gene50KB.mm10.SF_S.bed | awk \'{if ($6=="GATA1") print $0}\''
bedtools6 = 'cat cCRE.Gene50KB.hg38.withMID.bed | awk \'{if ($6=="GATA1") print $0}\''
system(bedtools5)
system(bedtools6)
#################################################


#################################################
##################
### read cCRE with SF_S MID hg38
cCRE_H_Jmeta_withMSF_hg38_regions = read.table('cCRE.Gene50KB.hg38.SF_S.bed', header=F)
cCRE_M_Jmeta_withMSF_mm10_regions = read.table('cCRE.Gene50KB.mm10.withMID.bed', header=F)
colnames(cCRE_H_Jmeta_withMSF_hg38_regions) = c('chr','start','end','JmetID','F01','GeneName','HID','MID')
cCRE_H_Jmeta_withMSF_hg38_regions_S = rep(0, dim(cCRE_H_Jmeta_withMSF_hg38_regions)[1])
all_genes_hg38 = unique(cCRE_H_Jmeta_withMSF_hg38_regions[,6])
kkk = 0
for (gene_i in all_genes_hg38){
	if (kkk%%1000==0){print(kkk)}
	kkk = kkk+1
	### get mm10 cCRE MID in gene_i
	cCRE_M_Jmeta_withMSF_mm10_regions_gene_i = cCRE_M_Jmeta_withMSF_mm10_regions[cCRE_M_Jmeta_withMSF_mm10_regions[,6]==gene_i,7]
	### get hg38 cCRE MID in gene_i
	cCRE_H_Jmeta_withMSF_hg38_regions_MID_gene_i = cCRE_H_Jmeta_withMSF_hg38_regions[cCRE_H_Jmeta_withMSF_hg38_regions[,6]==gene_i,8]
	### define the S cCRE 
	cCRE_H_Jmeta_withMSF_hg38_regions_S[cCRE_H_Jmeta_withMSF_hg38_regions[,6]==gene_i] = is.element(cCRE_H_Jmeta_withMSF_hg38_regions_MID_gene_i, cCRE_M_Jmeta_withMSF_mm10_regions_gene_i)*1
}
### add GeneS01 binary label
cCRE_H_Jmeta_withMSF_hg38_regions_set1 = cbind(cCRE_H_Jmeta_withMSF_hg38_regions, cCRE_H_Jmeta_withMSF_hg38_regions_S)
colnames(cCRE_H_Jmeta_withMSF_hg38_regions_set1) = c(colnames(cCRE_H_Jmeta_withMSF_hg38_regions), 'GeneS01')
write.table(cCRE_H_Jmeta_withMSF_hg38_regions_set1, 'cCRE.Gene50KB.hg38.JmetID.FS01.geneName.HID.MID.S01.bed', quote=F, sep='\t', col.names=F, row.names=F)
#####################
### read cCRE with SF_S MID mm10
cCRE_M_Jmeta_withHSF_mm10_regions = read.table('cCRE.Gene50KB.mm10.SF_S.bed', header=F)
cCRE_H_Jmeta_withHSF_hg38_regions = read.table('cCRE.Gene50KB.hg38.withMID.bed', header=F)
colnames(cCRE_M_Jmeta_withHSF_mm10_regions) = c('chr','start','end','JmetID','F01','GeneName','MID','HID')
cCRE_M_Jmeta_withHSF_mm10_regions_S = rep(0, dim(cCRE_M_Jmeta_withHSF_mm10_regions)[1])
all_genes_hg38 = unique(cCRE_M_Jmeta_withHSF_mm10_regions[,6])
kkk = 0
for (gene_i in all_genes_hg38){
	if (kkk%%1000==0){print(kkk)}
	kkk = kkk+1
	### get mm10 cCRE MID in gene_i
	cCRE_H_Jmeta_withHSF_hg38_regions_gene_i = cCRE_H_Jmeta_withHSF_hg38_regions[cCRE_H_Jmeta_withHSF_hg38_regions[,6]==gene_i,7]
	### get hg38 cCRE MID in gene_i
	cCRE_M_Jmeta_withHSF_mm10_regions_HID_gene_i = cCRE_M_Jmeta_withHSF_mm10_regions[cCRE_M_Jmeta_withHSF_mm10_regions[,6]==gene_i,8]
	### define the S cCRE 
	cCRE_M_Jmeta_withHSF_mm10_regions_S[cCRE_M_Jmeta_withHSF_mm10_regions[,6]==gene_i] = is.element(cCRE_M_Jmeta_withHSF_mm10_regions_HID_gene_i, cCRE_H_Jmeta_withHSF_hg38_regions_gene_i)*1
}
### add GeneS01 binary label
cCRE_M_Jmeta_withHSF_mm10_regions_set1 = cbind(cCRE_M_Jmeta_withHSF_mm10_regions, cCRE_M_Jmeta_withHSF_mm10_regions_S)
colnames(cCRE_M_Jmeta_withHSF_mm10_regions_set1) = c(colnames(cCRE_M_Jmeta_withHSF_mm10_regions), 'GeneS01')
write.table(cCRE_M_Jmeta_withHSF_mm10_regions_set1, 'cCRE.Gene50KB.mm10.JmetID.FS01.geneName.MID.HID.S01.bed', quote=F, sep='\t', col.names=F, row.names=F)
bedtools5 = 'cat cCRE.Gene50KB.hg38.JmetID.FS01.geneName.HID.MID.S01.bed | awk \'{if ($6=="GATA1") print $0}\''
bedtools6 = 'cat cCRE.Gene50KB.mm10.JmetID.FS01.geneName.MID.HID.S01.bed | awk \'{if ($6=="GATA1") print $0}\''
system(bedtools5)
system(bedtools6)
#################################################


#################################################
### Add protein coding gene TSS label
bedtools1 = 'cat /Users/guanjuexiang/Documents/projects/analysis/0813_human_mouse_state_compare_heatmap/hg38.gene.bed | awk -F \'\t\' -v OFS=\'\t\' -v exp_win=1000 \'{if (($2-exp_win>0) && ($4=="+")) print $1,$2-exp_win, $2+exp_win, $4,$5; if (($3-exp_win>0) && ($4=="-")) print $1,$3-exp_win, $3+exp_win, $4,$5; if (($2-exp_win<0) && ($4=="+")) print $1,0, $2+exp_win, $4,$5; if (($3-exp_win<0) && ($4=="-")) print $1,0, $3+exp_win, $4,$5}\' | sort -k1,1 -k2,2n > /Users/guanjuexiang/Documents/projects/analysis/0813_human_mouse_state_compare_heatmap/hg38.gene.sort.TSS.bed'
bedtools2 = 'bedtools map -a cCRE.Gene50KB.hg38.JmetID.FS01.geneName.HID.MID.S01.bed -b /Users/guanjuexiang/Documents/projects/analysis/0813_human_mouse_state_compare_heatmap/hg38.gene.sort.TSS.bed -c 5 -o concat -null NA > cCRE.Gene50KB.hg38.JmetID.FS01.geneName.HID.MID.S01.TSS.bed'
system(bedtools1)
system(bedtools2)
bedtools1 = 'cat /Users/guanjuexiang/Documents/projects/analysis/0813_human_mouse_state_compare_heatmap/mm10.gene.bed | awk -F \'\t\' -v OFS=\'\t\' -v exp_win=1000 \'{if (($2-exp_win>0) && ($4=="+")) print $1,$2-exp_win, $2+exp_win, $4,$5; if (($3-exp_win>0) && ($4=="-")) print $1,$3-exp_win, $3+exp_win, $4,$5; if (($2-exp_win<0) && ($4=="+")) print $1,0, $2+exp_win, $4,$5; if (($3-exp_win<0) && ($4=="-")) print $1,0, $3+exp_win, $4,$5}\' | sort -k1,1 -k2,2n > /Users/guanjuexiang/Documents/projects/analysis/0813_human_mouse_state_compare_heatmap/mm10.gene.sort.TSS.bed'
bedtools2 = 'bedtools map -a cCRE.Gene50KB.mm10.JmetID.FS01.geneName.MID.HID.S01.bed -b /Users/guanjuexiang/Documents/projects/analysis/0813_human_mouse_state_compare_heatmap/mm10.gene.sort.TSS.bed -c 5 -o concat -null NA > cCRE.Gene50KB.mm10.JmetID.FS01.geneName.MID.HID.S01.TSS.bed'
system(bedtools1)
system(bedtools2)
###
bedtools5 = 'cat cCRE.Gene50KB.hg38.JmetID.FS01.geneName.HID.MID.S01.TSS.bed | awk \'{if ($6=="GATA1") print $0}\''
system(bedtools5)
bedtools5 = 'cat cCRE.Gene50KB.mm10.JmetID.FS01.geneName.MID.HID.S01.TSS.bed | awk \'{if ($6=="GATA1") print $0}\''
system(bedtools5)
#################################################


#################################################
### add Gene Jmet enrichment KM cluster ID
add_Gene_KMID = function(input_mat_file, output_mat_file, meta_cluster_mat_log2_km, cCRE_HM_Jmeta_withMSF_hg38mm10_regions_set1) {
	cCRE_gene_JmetID_FS01_geneName_MID_HID_S01_TSS = read.table(input_mat_file, header=F)
	cCRE_gene_JmetID_FS01_geneName_MID_HID_S01_TSS_GeneKM = rep(0,dim(cCRE_gene_JmetID_FS01_geneName_MID_HID_S01_TSS)[1])
	for (gene_i in unique(cCRE_gene_JmetID_FS01_geneName_MID_HID_S01_TSS[,6])){
	cCRE_gene_JmetID_FS01_geneName_MID_HID_S01_TSS_GeneKM[cCRE_gene_JmetID_FS01_geneName_MID_HID_S01_TSS[,6]==gene_i] = meta_cluster_mat_log2_km$cluster[names(meta_cluster_mat_log2_km$cluster)==gene_i]
	}
	cCRE_gene_JmetID_FS01_geneName_MID_HID_S01_TSS_GeneKM_mat = cbind(cCRE_gene_JmetID_FS01_geneName_MID_HID_S01_TSS, cCRE_gene_JmetID_FS01_geneName_MID_HID_S01_TSS_GeneKM)
	colnames(cCRE_gene_JmetID_FS01_geneName_MID_HID_S01_TSS_GeneKM_mat) = c(colnames(cCRE_HM_Jmeta_withMSF_hg38mm10_regions_set1), 'TSS', 'GeneKMID')
	write.table(cCRE_gene_JmetID_FS01_geneName_MID_HID_S01_TSS_GeneKM_mat, output_mat_file, quote=F, sep='\t', col.names=T, row.names=F)
	return(cCRE_gene_JmetID_FS01_geneName_MID_HID_S01_TSS_GeneKM_mat)
}
cCRE_gene_JmetID_FS01_geneName_MID_HID_S01_TSS_GeneKM_mat_hg38 = add_Gene_KMID('cCRE.Gene50KB.hg38.JmetID.FS01.geneName.HID.MID.S01.TSS.bed', 'cCRE.Gene50KB.hg38.JmetID.FS01.geneName.HID.MID.S01.TSS.GeneKMID.bed', meta_cluster_mat_log2_km, cCRE_H_Jmeta_withMSF_hg38_regions_set1)
cCRE_gene_JmetID_FS01_geneName_MID_HID_S01_TSS_GeneKM_mat_mm10 = add_Gene_KMID('cCRE.Gene50KB.mm10.JmetID.FS01.geneName.MID.HID.S01.TSS.bed', 'cCRE.Gene50KB.mm10.JmetID.FS01.geneName.MID.HID.S01.TSS.GeneKMID.bed', meta_cluster_mat_log2_km, cCRE_M_Jmeta_withHSF_mm10_regions_set1)
### check
cCRE_gene_JmetID_FS01_geneName_MID_HID_S01_TSS_GeneKM_mat_hg38[cCRE_gene_JmetID_FS01_geneName_MID_HID_S01_TSS_GeneKM_mat_hg38[,6]=='GATA1',]
cCRE_gene_JmetID_FS01_geneName_MID_HID_S01_TSS_GeneKM_mat_mm10[cCRE_gene_JmetID_FS01_geneName_MID_HID_S01_TSS_GeneKM_mat_mm10[,6]=='GATA1',]
cCRE_gene_JmetID_FS01_geneName_MID_HID_S01_TSS_GeneKM_mat_hg38[cCRE_gene_JmetID_FS01_geneName_MID_HID_S01_TSS_GeneKM_mat_hg38[,6]=='ALAS2',]
cCRE_gene_JmetID_FS01_geneName_MID_HID_S01_TSS_GeneKM_mat_mm10[cCRE_gene_JmetID_FS01_geneName_MID_HID_S01_TSS_GeneKM_mat_mm10[,6]=='ALAS2',]
cCRE_gene_JmetID_FS01_geneName_MID_HID_S01_TSS_GeneKM_mat_hg38[cCRE_gene_JmetID_FS01_geneName_MID_HID_S01_TSS_GeneKM_mat_hg38[,6]=='CSF1R',]
#################################################


'''
#################################################
### get F01 based on Gene Kmeans clusters
add_F01_KM = function(cCRE_gene_JmetID_FS01_geneName_MID_HID_S01_TSS_GeneKM_mat, meta_cluster_mat_GeneGroup_Jmet_zp_01){
	F01KM = rep(0, dim(cCRE_gene_JmetID_FS01_geneName_MID_HID_S01_TSS_GeneKM_mat)[1])
	for (i in as.numeric(colnames(meta_cluster_mat_GeneGroup_Jmet_zp_01))){
		print(i)
		meta_cluster_mat_GeneGroup_Jmet_zp_01_i = as.numeric(rownames(meta_cluster_mat_GeneGroup_Jmet_zp_01)[meta_cluster_mat_GeneGroup_Jmet_zp_01[,i]==1])
		meta_cluster_mat_GeneGroup_Jmet_zp_01_i_binary = is.element(cCRE_gene_JmetID_FS01_geneName_MID_HID_S01_TSS_GeneKM_mat$JmetID, meta_cluster_mat_GeneGroup_Jmet_zp_01_i) & (cCRE_gene_JmetID_FS01_geneName_MID_HID_S01_TSS_GeneKM_mat$GeneKMID==i)
		F01KM[meta_cluster_mat_GeneGroup_Jmet_zp_01_i_binary] = 1
	}
	return(F01KM)
}
### get F01 based on Gene Kmeans clusters
hg38_F01KM = add_F01_KM(cCRE_gene_JmetID_FS01_geneName_MID_HID_S01_TSS_GeneKM_mat_hg38, meta_cluster_mat_GeneGroup_Jmet_zp_01)
mm10_F01KM = add_F01_KM(cCRE_gene_JmetID_FS01_geneName_MID_HID_S01_TSS_GeneKM_mat_mm10, meta_cluster_mat_GeneGroup_Jmet_zp_01)
### check
cbind(cCRE_gene_JmetID_FS01_geneName_MID_HID_S01_TSS_GeneKM_mat_hg38, hg38_F01KM)[cCRE_gene_JmetID_FS01_geneName_MID_HID_S01_TSS_GeneKM_mat_hg38$GeneName=='GATA1',]
cbind(cCRE_gene_JmetID_FS01_geneName_MID_HID_S01_TSS_GeneKM_mat_hg38, hg38_F01KM)[cCRE_gene_JmetID_FS01_geneName_MID_HID_S01_TSS_GeneKM_mat_hg38$GeneName=='ALAS2',]
### 
hg38_F01_OD = cCRE_gene_JmetID_FS01_geneName_MID_HID_S01_TSS_GeneKM_mat_hg38$F01
mm10_F01_OD = cCRE_gene_JmetID_FS01_geneName_MID_HID_S01_TSS_GeneKM_mat_mm10$F01
### replace original F01
cCRE_gene_JmetID_FS01_geneName_MID_HID_S01_TSS_GeneKM_mat_hg38$F01 = hg38_F01KM
cCRE_gene_JmetID_FS01_geneName_MID_HID_S01_TSS_GeneKM_mat_mm10$F01 = mm10_F01KM
### 
hg38_F01_new = cCRE_gene_JmetID_FS01_geneName_MID_HID_S01_TSS_GeneKM_mat_hg38$F01
mm10_F01_new = cCRE_gene_JmetID_FS01_geneName_MID_HID_S01_TSS_GeneKM_mat_mm10$F01
###
#cCRE_gene_JmetID_FS01_geneName_MID_HID_S01_TSS_GeneKM_mat_hg38$F01 = hg38_F01_OD
#cCRE_gene_JmetID_FS01_geneName_MID_HID_S01_TSS_GeneKM_mat_mm10$F01 = mm10_F01_OD
#################################################
'''


#################################################
### Get Gene-by-Jmet RNA cor mat hg38
RNA_mat = read.table('HumanVISION_RNAseq_hg38_genes_tpm.txt', header=T)
RNA_mat_ct_list = c('HSC', 'HSC', 'ERY', 'ERY', 'TCD4', 'TCD4', 'TCD8', 'TCD8', 'B', 'B', 'CMP', 'CMP', 'MON', 'MON', 'NEU', 'NEU', 'MON', 'MON', 'GMP', 'GMP', 'CFUE', 'NK', 'NK', 'MK', 'MK', 'CLP', 'MPP', 'MPP', 'EOS', 'EOS', 'MEP', 'MEP', 'MK', 'MK', 'CLP', 'ERY', 'ERY', 'ERY', 'HUDEP1', 'HUDEP1', 'HUDEP2', 'HUDEP2', 'CD34', 'CD34')
RNA_mat_sig = RNA_mat[,-c(1:4)]
### ct average
unique_ct = unique(RNA_mat_ct_list)
RNA_mat_ct_ave = matrix(0, nrow=dim(RNA_mat)[1], ncol=length(unique_ct))
k = 0
for (ct in unique_ct){
	k = k+1
	if (sum(RNA_mat_ct_list==ct)>1){
		RNA_mat_ct_ave[,k] = rowMeans(RNA_mat_sig[,RNA_mat_ct_list==ct])
	} else{
		RNA_mat_ct_ave[,k] = RNA_mat_sig[,RNA_mat_ct_list==ct]
	}
}
colnames(RNA_mat_ct_ave) = unique_ct
### shared ct with HM_cts
RNA_mat_ct_ave_shared = c()
for (ct_i in colnames(dhs_dms_ctmerge_shared_reorder_meansig_Jmet_meansig)[-1]){
	RNA_mat_ct_ave_shared = cbind(RNA_mat_ct_ave_shared, RNA_mat_ct_ave[,colnames(RNA_mat_ct_ave)==ct_i])
}
#RNA_mat_ct_ave_shared = RNA_mat_ct_ave[,is.element(colnames(RNA_mat_ct_ave), colnames(dhs_dms_ctmerge_shared_reorder_meansig_Jmet_meansig))]
RNA_mat_ct_ave_shared_ave = rowMeans(RNA_mat_ct_ave_shared)
RNA_mat_ct_ave_shared = cbind(RNA_mat_ct_ave_shared_ave, RNA_mat_ct_ave_shared)
colnames(RNA_mat_ct_ave_shared) = c(colnames(dhs_dms_ctmerge_shared_reorder_meansig_Jmet_meansig))
### log QTnorm
quantile_norm = function(x){
	xm = (x[,1])
	refsig_sort = xm[order(xm)]
	for (i in 1:dim(x)[2]){
		sigtmp = x[,i]
		x[,i] = refsig_sort[rank(sigtmp)]
	}
	return(x)
}
smallnum = 1e-1 
RNA_mat_ct_ave_shared_logqtnorm = quantile_norm(log(RNA_mat_ct_ave_shared+smallnum))
### select used genes
RNA_ENSID2GeneName = read.table('gene.ENS_id2GeneName.hg38.txt', header=F)
colnames(RNA_ENSID2GeneName) = c('gene_id', 'gene_name')
RNA_mat_ENSID_GeneName = merge(RNA_mat[,c(1:4)], RNA_ENSID2GeneName, by='gene_id', all.x=T)
### select used genes
RNA_mat_ENSID_GeneName_used_binary = is.element(RNA_mat_ENSID_GeneName$gene_name, cCRE_gene_JmetID_FS01_geneName_MID_HID_S01_TSS_GeneKM_mat_hg38$GeneName)
RNA_mat_ENSID_GeneName_used = RNA_mat_ENSID_GeneName[RNA_mat_ENSID_GeneName_used_binary,]
RNA_mat_ct_ave_shared_logqtnorm_used = RNA_mat_ct_ave_shared_logqtnorm[RNA_mat_ENSID_GeneName_used_binary,]
rownames(RNA_mat_ct_ave_shared_logqtnorm_used) = RNA_mat_ENSID_GeneName_used[,5]
### gene-by-Jmet RNA-esRP correlation matrix
gene_RNA_Jmet_cor = cor(t(RNA_mat_ct_ave_shared_logqtnorm_used), t(dhs_dms_ctmerge_shared_reorder_meansig_Jmet_meansig))
gene_RNA_Jmet_cor = gene_RNA_Jmet_cor[,order(as.numeric(colnames(gene_RNA_Jmet_cor)))]
png('gene_Jmet_RNA_esRP_cor.png', width = 1000, height = 600)
set.seed(2019)
gene_RNA_Jmet_cor_plot_df = cbind(as.data.frame(rownames(gene_RNA_Jmet_cor)), gene_RNA_Jmet_cor)
colnames(gene_RNA_Jmet_cor_plot_df)[1] = 'shared_genes'

gene_RNA_Jmet_cor_plot_df_merge = merge(gene_RNA_Jmet_cor_plot_df, meta_cluster_mat_log2_df, by='shared_genes', all.x=T)
gene_RNA_Jmet_cor_plot_df_merge_plot = gene_RNA_Jmet_cor_plot_df_merge[order(gene_RNA_Jmet_cor_plot_df_merge$GeneKM_ID),2:(dim(dhs_dms_ctmerge_shared_reorder_meansig_Jmet_meansig)[1]+1)]
#gene_RNA_Jmet_cor_plot_df_merge_plot = gene_RNA_Jmet_cor_plot_df_merge[order(gene_RNA_Jmet_cor_plot_df_merge$GeneKM_ID),11:18]
colnames(gene_RNA_Jmet_cor_plot_df_merge_plot) = 1:dim(dhs_dms_ctmerge_shared_reorder_meansig_Jmet_meansig)[1]

gene_RNA_Jmet_cor_plot_df_merge_plot1 = c()
used_order = Jmet_order
for (coli in used_order){
	gene_RNA_Jmet_cor_plot_df_merge_plot1 = cbind(gene_RNA_Jmet_cor_plot_df_merge_plot1, gene_RNA_Jmet_cor_plot_df_merge_plot[,colnames(gene_RNA_Jmet_cor_plot_df_merge_plot)==coli])
}
colnames(gene_RNA_Jmet_cor_plot_df_merge_plot1) = used_order
png('gene_Jmet_RNA_esRP_cor.png', width = 1000, height = 600)
pheatmap(t(abs(gene_RNA_Jmet_cor_plot_df_merge_plot1)), cluster_col=F, cluster_rows=F, show_colnames=F, cex=2)
dev.off()
###
pdf('gene_Jmet_RNA_esRP_cor.hist.pdf')
hist(as.numeric(gene_RNA_Jmet_cor), breaks=50, cex=1.5)
box()
dev.off()
print(gene_RNA_Jmet_cor[rownames(gene_RNA_Jmet_cor)=='GATA1',])
#################################################


#################################################
### add Jmet-GeneTPM correlation hg38
#Jmet_gene_pairs = unique(cbind(as.data.frame(cCRE_gene_JmetID_FS01_geneName_MID_HID_S01_TSS_GeneKM_mat_hg38$Jmet), cCRE_gene_JmetID_FS01_geneName_MID_HID_S01_TSS_GeneKM_mat_hg38$GeneName))
#hg38_Jmet_RNA_cor = rep(0, dim(cCRE_gene_JmetID_FS01_geneName_MID_HID_S01_TSS_GeneKM_mat_hg38)[1])
#for (i in 1:dim(Jmet_gene_pairs)[1]){
#	if (i%%10000==0){print(i)}
#	if (sum(rownames(gene_RNA_Jmet_cor)==Jmet_gene_pairs[i,2])>0){
#	used_rows = cCRE_gene_JmetID_FS01_geneName_MID_HID_S01_TSS_GeneKM_mat_hg38$Jmet==Jmet_gene_pairs[i,1] & cCRE_gene_JmetID_FS01_geneName_MID_HID_S01_TSS_GeneKM_mat_hg38$GeneName==Jmet_gene_pairs[i,2]
#	hg38_Jmet_RNA_cor[used_rows] = gene_RNA_Jmet_cor[rownames(gene_RNA_Jmet_cor)==Jmet_gene_pairs[i,2],Jmet_gene_pairs[i,1]]
#	}
#}
#cCRE_gene_JmetID_FS01_geneName_MID_HID_S01_TSS_GeneKM_mat_hg38_RNAcor = cbind(cCRE_gene_JmetID_FS01_geneName_MID_HID_S01_TSS_GeneKM_mat_hg38, hg38_Jmet_RNA_cor)
### get esRP RNAlogTPM cor
cCRE_hg38_coordinates = cCRE_gene_JmetID_FS01_geneName_MID_HID_S01_TSS_GeneKM_mat_hg38[,1:3]
cCRE_gene_JmetID_FS01_geneName_MID_HID_S01_TSS_GeneKM_mat_hg38_withPKID = cbind(apply(cbind(cCRE_hg38_coordinates[,1], as.character(cCRE_hg38_coordinates[,2]), as.character(cCRE_hg38_coordinates[,3])), 1, function(x) paste(x, collapse='_')), cCRE_gene_JmetID_FS01_geneName_MID_HID_S01_TSS_GeneKM_mat_hg38)
cCRE_gene_PKID = cCRE_gene_JmetID_FS01_geneName_MID_HID_S01_TSS_GeneKM_mat_hg38_withPKID[,1]
cCRE_gene_genes = cCRE_gene_JmetID_FS01_geneName_MID_HID_S01_TSS_GeneKM_mat_hg38_withPKID[,7]
RNA_mat_genes = rownames(RNA_mat_ct_ave_shared_logqtnorm_used)
cCRE_PKID = dh_with_JointClusterID_mat_ct[,1]
cCRE_esRPmat = dh_with_JointClusterID_mat_ct[,-c(1:8)]
###
ptm <- proc.time()
cCRE_gene_esRP_RNA_cor = rep(-100,dim(cCRE_gene_JmetID_FS01_geneName_MID_HID_S01_TSS_GeneKM_mat_hg38_withPKID)[1])
for (i in 1:dim(cCRE_gene_JmetID_FS01_geneName_MID_HID_S01_TSS_GeneKM_mat_hg38_withPKID)[1]){
	if (i%%10000==0){print(i)}
	gene_i = cCRE_gene_genes[i]
	cCRE_i = cCRE_gene_PKID[i]
	if (is.element(gene_i, RNA_mat_genes)){
	cCRE_esRP_i = cCRE_esRPmat[cCRE_PKID==cCRE_i,]
	RNA_tpm_i = RNA_mat_ct_ave_shared_logqtnorm_used[RNA_mat_genes==gene_i,]
	if (!is.null(dim(RNA_tpm_i))){
			cCRE_gene_esRP_RNA_cor[i] = cor(as.numeric(cCRE_esRP_i), RNA_tpm_i[1,])
		} else{
			cCRE_gene_esRP_RNA_cor[i] = cor(as.numeric(cCRE_esRP_i), RNA_tpm_i)
		}
	}
}
proc.time() - ptm
###
cCRE_gene_JmetID_FS01_geneName_MID_HID_S01_TSS_GeneKM_mat_hg38_RNAcor = cbind(cCRE_gene_JmetID_FS01_geneName_MID_HID_S01_TSS_GeneKM_mat_hg38, cCRE_gene_esRP_RNA_cor)
colnames(cCRE_gene_JmetID_FS01_geneName_MID_HID_S01_TSS_GeneKM_mat_hg38_RNAcor)[dim(cCRE_gene_JmetID_FS01_geneName_MID_HID_S01_TSS_GeneKM_mat_hg38_RNAcor)[2]] = 'RNAcor'
cCRE_gene_JmetID_FS01_geneName_MID_HID_S01_TSS_GeneKM_mat_hg38_RNAcor[cCRE_gene_JmetID_FS01_geneName_MID_HID_S01_TSS_GeneKM_mat_hg38_RNAcor$GeneName=='GATA1',-c(1:3)]
### add SFN label
cCRE_gene_JmetID_FS01_geneName_MID_HID_S01_TSS_GeneKM_mat_hg38_RNAcor_SFN = rep('N', dim(cCRE_gene_JmetID_FS01_geneName_MID_HID_S01_TSS_GeneKM_mat_hg38_RNAcor)[1])
# exclude TSS
exclude_TSS = (cCRE_gene_JmetID_FS01_geneName_MID_HID_S01_TSS_GeneKM_mat_hg38_RNAcor$GeneName!=cCRE_gene_JmetID_FS01_geneName_MID_HID_S01_TSS_GeneKM_mat_hg38_RNAcor$TSS) | is.na(cCRE_gene_JmetID_FS01_geneName_MID_HID_S01_TSS_GeneKM_mat_hg38_RNAcor$TSS)
#exclude_TSS = cCRE_gene_JmetID_FS01_geneName_MID_HID_S01_TSS_GeneKM_mat_hg38_RNAcor$F01==0
#cCRE_gene_JmetID_FS01_geneName_MID_HID_S01_TSS_GeneKM_mat_hg38_RNAcor_SFN[((cCRE_gene_JmetID_FS01_geneName_MID_HID_S01_TSS_GeneKM_mat_hg38_RNAcor$F01==1) | (!exclude_TSS)) & (cCRE_gene_JmetID_FS01_geneName_MID_HID_S01_TSS_GeneKM_mat_hg38_RNAcor$GeneS01==1) ] = 'SF'
#cCRE_gene_JmetID_FS01_geneName_MID_HID_S01_TSS_GeneKM_mat_hg38_RNAcor_SFN[((cCRE_gene_JmetID_FS01_geneName_MID_HID_S01_TSS_GeneKM_mat_hg38_RNAcor$F01==1) | (!exclude_TSS)) & (cCRE_gene_JmetID_FS01_geneName_MID_HID_S01_TSS_GeneKM_mat_hg38_RNAcor$GeneS01==0) ] = 'F'
#cCRE_gene_JmetID_FS01_geneName_MID_HID_S01_TSS_GeneKM_mat_hg38_RNAcor_SFN[((cCRE_gene_JmetID_FS01_geneName_MID_HID_S01_TSS_GeneKM_mat_hg38_RNAcor$F01==0) & (exclude_TSS)) & (cCRE_gene_JmetID_FS01_geneName_MID_HID_S01_TSS_GeneKM_mat_hg38_RNAcor$GeneS01==1) ] = 'S'
cCRE_gene_JmetID_FS01_geneName_MID_HID_S01_TSS_GeneKM_mat_hg38_RNAcor_SFN[(cCRE_gene_JmetID_FS01_geneName_MID_HID_S01_TSS_GeneKM_mat_hg38_RNAcor$F01==1) & (cCRE_gene_JmetID_FS01_geneName_MID_HID_S01_TSS_GeneKM_mat_hg38_RNAcor$GeneS01==1) ] = 'SF'
cCRE_gene_JmetID_FS01_geneName_MID_HID_S01_TSS_GeneKM_mat_hg38_RNAcor_SFN[(cCRE_gene_JmetID_FS01_geneName_MID_HID_S01_TSS_GeneKM_mat_hg38_RNAcor$F01==1) & (cCRE_gene_JmetID_FS01_geneName_MID_HID_S01_TSS_GeneKM_mat_hg38_RNAcor$GeneS01==0) ] = 'F'
cCRE_gene_JmetID_FS01_geneName_MID_HID_S01_TSS_GeneKM_mat_hg38_RNAcor_SFN[(cCRE_gene_JmetID_FS01_geneName_MID_HID_S01_TSS_GeneKM_mat_hg38_RNAcor$F01==0) & (cCRE_gene_JmetID_FS01_geneName_MID_HID_S01_TSS_GeneKM_mat_hg38_RNAcor$GeneS01==1) ] = 'S'
cCRE_gene_JmetID_FS01_geneName_MID_HID_S01_TSS_GeneKM_mat_hg38_RNAcor_SFN[!exclude_TSS] = 'P'
###
cCRE_gene_JmetID_FS01_geneName_MID_HID_S01_TSS_GeneKM_mat_hg38_RNAcor_SFN_mat = cbind(cCRE_gene_JmetID_FS01_geneName_MID_HID_S01_TSS_GeneKM_mat_hg38_RNAcor, cCRE_gene_JmetID_FS01_geneName_MID_HID_S01_TSS_GeneKM_mat_hg38_RNAcor_SFN)
colnames(cCRE_gene_JmetID_FS01_geneName_MID_HID_S01_TSS_GeneKM_mat_hg38_RNAcor_SFN_mat)[dim(cCRE_gene_JmetID_FS01_geneName_MID_HID_S01_TSS_GeneKM_mat_hg38_RNAcor_SFN_mat)[2]] = 'SFNID'
cCRE_gene_JmetID_FS01_geneName_MID_HID_S01_TSS_GeneKM_mat_hg38_RNAcor_SFN_mat[cCRE_gene_JmetID_FS01_geneName_MID_HID_S01_TSS_GeneKM_mat_hg38_RNAcor_SFN_mat$GeneName=='GATA1',]
meta_cluster_mat_log2_df[meta_cluster_mat_log2_df$shared_genes=='GATA1',]
cCRE_gene_JmetID_FS01_geneName_MID_HID_S01_TSS_GeneKM_mat_hg38_RNAcor_SFN_mat[cCRE_gene_JmetID_FS01_geneName_MID_HID_S01_TSS_GeneKM_mat_hg38_RNAcor_SFN_mat$GeneName=='CSF1R',]
meta_cluster_mat_log2_df[meta_cluster_mat_log2_df$shared_genes=='CSF1R',]
write.table(cCRE_gene_JmetID_FS01_geneName_MID_HID_S01_TSS_GeneKM_mat_hg38_RNAcor_SFN_mat, 'cCRE.Gene50KB.hg38.JmetID.FS01.geneName.HID.MID.S01.TSS.GeneKMID.RNAcor.SFNID.bed', quote=F, sep='\t', col.names=T, row.names=F)
### Gene cor of 'SF','F','S','N' of each gene
SFN_ID_list = c('P', 'SF','F','S','N')
score_mat = matrix(0, nrow = length(unique(cCRE_gene_JmetID_FS01_geneName_MID_HID_S01_TSS_GeneKM_mat_hg38_RNAcor_SFN_mat$GeneName)), ncol=length(SFN_ID_list))
k = 0
for (gene_i in unique(cCRE_gene_JmetID_FS01_geneName_MID_HID_S01_TSS_GeneKM_mat_hg38_RNAcor_SFN_mat$GeneName)){
	k = k+1
	if (k%%1000==0){print(k)}
	SFN_mat_gene_i = cCRE_gene_JmetID_FS01_geneName_MID_HID_S01_TSS_GeneKM_mat_hg38_RNAcor_SFN_mat[cCRE_gene_JmetID_FS01_geneName_MID_HID_S01_TSS_GeneKM_mat_hg38_RNAcor_SFN_mat$GeneName==gene_i,]
	SFN_mat_gene_i_dat = unique(SFN_mat_gene_i[,c(4,5,9,12,13,2)])
	score_mat_i = rep(-100, length(SFN_ID_list))
	for (j in 1:length(SFN_ID_list)){
		id = SFN_ID_list[j]
		if (sum(SFN_mat_gene_i_dat$SFNID==id)>0){
			score_mat_i[j] = mean(SFN_mat_gene_i_dat[SFN_mat_gene_i_dat$SFNID==id,4], na.rm=T)
		} 
	}
	score_mat[k,] = score_mat_i
}
### 
colnames(score_mat) = SFN_ID_list
score_mat_abs = abs(score_mat)
used_rows = apply(score_mat, 1, sum)>-10
wilcox.test(score_mat[used_rows,1], score_mat[used_rows,4], alternative='greater', paired=T)
wilcox.test(score_mat[used_rows,2], score_mat[used_rows,4], alternative='greater', paired=T)
wilcox.test(score_mat[used_rows,3], score_mat[used_rows,4], alternative='greater', paired=T)
wilcox.test(score_mat[used_rows,2], score_mat[used_rows,3], alternative='greater', paired=T)
wilcox.test(score_mat[used_rows,3], score_mat[used_rows,5], alternative='greater', paired=T)
### cor boxplot
pdf('corabs_SFN_mat.box.pdf')
#boxplot(score_mat[used_rows,]-score_mat[used_rows,4], cex.axis=2)
boxplot(score_mat[used_rows,], cex.axis=2, ylim=c(-1,1))
points(1:dim(score_mat)[2], colMeans(score_mat[used_rows,], na.rm=T))
lines(1:dim(score_mat)[2], colMeans(score_mat[used_rows,], na.rm=T))
abline(h=0)
dev.off()
pdf('corabs_SFN_mat.box.dif.pdf')
boxplot(score_mat[used_rows,]-score_mat[used_rows,4], cex.axis=2)
#boxplot(score_mat[used_rows,], cex.axis=2, ylim=c(-1,1))
abline(h=0)
dev.off()
########################
### add SFNID to mm10
cCRE_gene_JmetID_FS01_geneName_MID_HID_S01_TSS_GeneKM_mat_mm10_SFN = rep('N', dim(cCRE_gene_JmetID_FS01_geneName_MID_HID_S01_TSS_GeneKM_mat_mm10)[1])
cCRE_gene_JmetID_FS01_geneName_MID_HID_S01_TSS_GeneKM_mat_mm10_SFN[(cCRE_gene_JmetID_FS01_geneName_MID_HID_S01_TSS_GeneKM_mat_mm10$F01==1) & (cCRE_gene_JmetID_FS01_geneName_MID_HID_S01_TSS_GeneKM_mat_mm10$GeneS01==1)] = 'SF'
cCRE_gene_JmetID_FS01_geneName_MID_HID_S01_TSS_GeneKM_mat_mm10_SFN[(cCRE_gene_JmetID_FS01_geneName_MID_HID_S01_TSS_GeneKM_mat_mm10$F01==1) & (cCRE_gene_JmetID_FS01_geneName_MID_HID_S01_TSS_GeneKM_mat_mm10$GeneS01==0)] = 'F'
cCRE_gene_JmetID_FS01_geneName_MID_HID_S01_TSS_GeneKM_mat_mm10_SFN[(cCRE_gene_JmetID_FS01_geneName_MID_HID_S01_TSS_GeneKM_mat_mm10$F01==0) & (cCRE_gene_JmetID_FS01_geneName_MID_HID_S01_TSS_GeneKM_mat_mm10$GeneS01==1)] = 'S'
cCRE_gene_JmetID_FS01_geneName_MID_HID_S01_TSS_GeneKM_mat_mm10_SFN_mat = cbind(cCRE_gene_JmetID_FS01_geneName_MID_HID_S01_TSS_GeneKM_mat_mm10, cCRE_gene_JmetID_FS01_geneName_MID_HID_S01_TSS_GeneKM_mat_mm10_SFN)
colnames(cCRE_gene_JmetID_FS01_geneName_MID_HID_S01_TSS_GeneKM_mat_mm10_SFN_mat)[dim(cCRE_gene_JmetID_FS01_geneName_MID_HID_S01_TSS_GeneKM_mat_mm10_SFN_mat)[2]] = 'SFNID'
### add P to mm10
cCRE_gene_JmetID_FS01_geneName_MID_HID_S01_TSS_GeneKM_mat_mm10_SFN_mat[,10] = toupper(cCRE_gene_JmetID_FS01_geneName_MID_HID_S01_TSS_GeneKM_mat_mm10_SFN_mat[,10])
exclude_TSS_mm10 = (cCRE_gene_JmetID_FS01_geneName_MID_HID_S01_TSS_GeneKM_mat_mm10_SFN_mat$GeneName!=cCRE_gene_JmetID_FS01_geneName_MID_HID_S01_TSS_GeneKM_mat_mm10_SFN_mat$TSS) | is.na(cCRE_gene_JmetID_FS01_geneName_MID_HID_S01_TSS_GeneKM_mat_mm10_SFN_mat$TSS)
cCRE_gene_JmetID_FS01_geneName_MID_HID_S01_TSS_GeneKM_mat_mm10_SFN_mat$SFNID[!exclude_TSS_mm10] = 'P'
###
write.table(cCRE_gene_JmetID_FS01_geneName_MID_HID_S01_TSS_GeneKM_mat_mm10_SFN_mat, 'cCRE.Gene50KB.mm10.JmetID.FS01.geneName.MID.HID.S01.TSS.GeneKMID.SFNID.bed', quote=F, sep='\t', col.names=T, row.names=F)
write.table(cCRE_gene_JmetID_FS01_geneName_MID_HID_S01_TSS_GeneKM_mat_hg38_RNAcor_SFN_mat[,colnames(cCRE_gene_JmetID_FS01_geneName_MID_HID_S01_TSS_GeneKM_mat_hg38_RNAcor_SFN_mat)!='RNAcor'], 'cCRE.Gene50KB.hg38.JmetID.FS01.geneName.HID.MID.S01.TSS.GeneKMID.SFNID.bed', quote=F, sep='\t', col.names=T, row.names=F)
### info
# 118435 unique hg38 cCREs & 66106 unique mm10 cCREs
# 234762 unique hg38 cCREs-protein-coding-gene pairs & 145109 unique mm10 cCREs-protein-coding-gene pairs (TSS +/-50kb)
table(cCRE_gene_JmetID_FS01_geneName_MID_HID_S01_TSS_GeneKM_mat_hg38_RNAcor_SFN_mat$SFNID)
# hg38
#    F      N      P      S     SF 
# 53638 125967  15275  26034  13848
     F      N      P      S     SF 
 53638 125967  15275  26034  13848 
table(cCRE_gene_JmetID_FS01_geneName_MID_HID_S01_TSS_GeneKM_mat_mm10_SFN_mat$SFNID)
# mm10
#    F     N     P     S    SF 
# 26630 64980 12324 26961 14214
    F     N     P     S    SF 
26630 64980 12324 26961 14214 
#################################################


#################################################
### count number 
SFNP_ID_order = c('P','SF','F','S','N')
Jmet_i_SFNP_count = c()
Jmet_i_SFNP_enrich = c()
for (Jmet_i in used_order){
	print(Jmet_i)
	Jmet_i_binary = cCRE_gene_JmetID_FS01_geneName_MID_HID_S01_TSS_GeneKM_mat_mm10_SFN_mat$JmetID==Jmet_i
	SFNP_ID_order_count_i = c()
	SFNP_ID_order_enrich_i = c()
	for (SFNP_j in SFNP_ID_order){
		obs_Jmet_i_SFNP_j = sum((Jmet_i_binary * (cCRE_gene_JmetID_FS01_geneName_MID_HID_S01_TSS_GeneKM_mat_mm10_SFN_mat$SFNID==SFNP_j))!=0)
		exp_Jmet_i_SFNP_j = sum(Jmet_i_binary) / dim(cCRE_gene_JmetID_FS01_geneName_MID_HID_S01_TSS_GeneKM_mat_mm10_SFN_mat)[1] * sum(cCRE_gene_JmetID_FS01_geneName_MID_HID_S01_TSS_GeneKM_mat_mm10_SFN_mat$SFNID==SFNP_j) 
		SFNP_ID_order_count_i = c(SFNP_ID_order_count_i, obs_Jmet_i_SFNP_j)
		SFNP_ID_order_enrich_i = c(SFNP_ID_order_enrich_i, (obs_Jmet_i_SFNP_j+100)/(exp_Jmet_i_SFNP_j+100))
	}
	Jmet_i_SFNP_count = rbind(Jmet_i_SFNP_count, SFNP_ID_order_count_i)
	Jmet_i_SFNP_enrich = rbind(Jmet_i_SFNP_enrich, SFNP_ID_order_enrich_i)
}
###
colnames(Jmet_i_SFNP_count) = SFNP_ID_order
rownames(Jmet_i_SFNP_count) = used_order
colnames(Jmet_i_SFNP_enrich) = SFNP_ID_order
rownames(Jmet_i_SFNP_enrich) = used_order
###
pdf('Jmet_i_SFNP_count.pdf', width=3, height=5)
pheatmap(log10(Jmet_i_SFNP_count), cluster_cols=F, cluster_rows=F, cex=1.5)
dev.off()
pdf('Jmet_i_SFNP_enrich.pdf', width=3, height=5)
pheatmap(log10(Jmet_i_SFNP_enrich), cluster_cols=F, cluster_rows=F, cex=1.5)
dev.off()
#################################################


#################################################
### write Gene-cCRE loop file
hg38_gene_locus = read.table('/Users/guanjuexiang/Documents/projects/analysis/0813_human_mouse_state_compare_heatmap/hg38.gene.bed', header=F)
hg38_gene_locus_TSS = hg38_gene_locus
hg38_gene_locus_TSS[hg38_gene_locus_TSS[,4]=='+',3] = hg38_gene_locus_TSS[hg38_gene_locus_TSS[,4]=='+',2]+1
hg38_gene_locus_TSS[hg38_gene_locus_TSS[,4]=='-',2] = hg38_gene_locus_TSS[hg38_gene_locus_TSS[,4]=='-',3]-1
### add TSS position
hg38_SFN_loops = cCRE_gene_JmetID_FS01_geneName_MID_HID_S01_TSS_GeneKM_mat_hg38_RNAcor_SFN_mat
hg38_SFN_loops_TSS = hg38_SFN_loops[,1:3]
k = 0
for (gene_i in unique(hg38_SFN_loops$GeneName)){
	k = k+1
	if (k%%1000==0){print(k)}
	used_n = sum(hg38_SFN_loops$GeneName==gene_i)
	TSS_gene_i = hg38_gene_locus_TSS[hg38_gene_locus_TSS[,5]==gene_i,1:3]
	hg38_SFN_loops_TSS[hg38_SFN_loops$GeneName==gene_i,] = cbind(as.data.frame(rep(TSS_gene_i[1,1], used_n)), rep(TSS_gene_i[1,2], used_n), rep(TSS_gene_i[1,3], used_n))
}
###
hg38_SFN_loops = cbind(hg38_SFN_loops[,1:5], round(hg38_SFN_loops[12]*1000), hg38_SFN_loops[,13], rep('#DEDEDE', dim(hg38_SFN_loops)[1]), hg38_SFN_loops[,c(1:3,7)], rep('.', dim(hg38_SFN_loops)[1]), hg38_SFN_loops_TSS, hg38_SFN_loops$GeneName, rep('.', dim(hg38_SFN_loops)[1]) )
### remove neg
hg38_SFN_loops[(hg38_SFN_loops[,6]<0) | (is.na(hg38_SFN_loops[,6])),6] = 0
### change color
hg38_SFN_loops[hg38_SFN_loops[,7]=='SF',8] = '#FD0606'
hg38_SFN_loops[hg38_SFN_loops[,7]=='F',8] = '#FFAB00'
hg38_SFN_loops[hg38_SFN_loops[,7]=='S',8] = '#696969'
hg38_SFN_loops[hg38_SFN_loops[,7]=='N',8] = '#DEDEDE'
hg38_SFN_loops[hg38_SFN_loops[,7]=='P',8] = '#FF0000'
hg38_SFN_loops[hg38_SFN_loops[,17]=='GATA1',]
hg38_gene_shared_exp[hg38_gene_shared_exp[,5]=='GATA1',]
hg38_SFN_loops[hg38_SFN_loops[,17]=='CSF1R',]
hg38_gene_shared_exp[hg38_gene_shared_exp[,5]=='CSF1R',]
options("scipen"=100, "digits"=4)
hg38_SFN_loops$HID = apply(cbind(hg38_SFN_loops$JmetID, hg38_SFN_loops$HID), 1, function(x) paste(x[1], x[2], sep=':'))
write.table(hg38_SFN_loops[hg38_SFN_loops[,6]>=0,], 'hg38.SFN.loop.interact', quote=F, sep='\t', col.names=F, row.names=F)
bash1 = 'cat loop.header.txt > hg38.SFN.loop.interact.tmp'
bash2 = 'cat hg38.SFN.loop.interact >> hg38.SFN.loop.interact.tmp && mv hg38.SFN.loop.interact.tmp hg38.SFN.loop.interact'
system(bash1)
system(bash2)
bash1 = 'cat loop.header.txt > hg38.SFN.loop.interact.tmp'
bash2 = 'cat hg38.SFN.loop.interact | awk -F \'\t\' -v OFS=\'\t\' \'{if ($17=="GATA1") print $0}\' >> hg38.SFN.loop.interact.tmp && mv hg38.SFN.loop.interact.tmp hg38.SFN.loop.GATA1.interact'
system(bash1)
system(bash2)
bash1 = 'cat loop.header.txt > hg38.SFN.loop.interact.tmp'
bash2 = 'cat hg38.SFN.loop.interact | awk -F \'\t\' -v OFS=\'\t\' \'{if ($17=="HBA1") print $0}\' >> hg38.SFN.loop.interact.tmp && mv hg38.SFN.loop.interact.tmp hg38.SFN.loop.HBA1.interact'
system(bash1)
system(bash2)
bash1 = 'cat loop.header.txt > hg38.SFN.loop.interact.tmp'
bash2 = 'cat hg38.SFN.loop.interact | awk -F \'\t\' -v OFS=\'\t\' \'{if ($17=="CSF1R") print $0}\' >> hg38.SFN.loop.interact.tmp && mv hg38.SFN.loop.interact.tmp hg38.SFN.loop.CSF1R.interact'
system(bash1)
system(bash2)
bash1 = 'cat loop.header.txt > hg38.SFN.loop.interact.tmp'
bash2 = 'cat hg38.SFN.loop.interact | awk -F \'\t\' -v OFS=\'\t\' \'{if ($17=="IFNG") print $0}\' >> hg38.SFN.loop.interact.tmp && mv hg38.SFN.loop.interact.tmp hg38.SFN.loop.IFNG.interact'
system(bash1)
system(bash2)
bash1 = 'cat loop.header.txt > hg38.SFN.loop.interact.tmp'
bash2 = 'cat hg38.SFN.loop.interact | awk -F \'\t\' -v OFS=\'\t\' \'{if ($17=="H2AC4") print $0}\' >> hg38.SFN.loop.interact.tmp && mv hg38.SFN.loop.interact.tmp hg38.SFN.loop.H2AC4.interact'
system(bash1)
system(bash2)
bash1 = 'cat loop.header.txt > hg38.SFN.loop.interact.tmp'
bash2 = 'cat hg38.SFN.loop.interact | awk -F \'\t\' -v OFS=\'\t\' \'{if ($17=="ALAS2") print $0}\' >> hg38.SFN.loop.interact.tmp && mv hg38.SFN.loop.interact.tmp hg38.SFN.loop.ALAS2.interact'
system(bash1)
system(bash2)
hg38_gene_shared_exp[hg38_gene_shared_exp[,5]=='ALAS2',]
#################################################


#################################################
### write Gene-cCRE loop file mm10
mm10_gene_locus = read.table('/Users/guanjuexiang/Documents/projects/analysis/0813_human_mouse_state_compare_heatmap/mm10.gene.bed', header=F)
mm10_gene_locus_TSS = mm10_gene_locus
mm10_gene_locus_TSS[mm10_gene_locus_TSS[,4]=='+',3] = mm10_gene_locus_TSS[mm10_gene_locus_TSS[,4]=='+',2]+1
mm10_gene_locus_TSS[mm10_gene_locus_TSS[,4]=='-',2] = mm10_gene_locus_TSS[mm10_gene_locus_TSS[,4]=='-',3]-1
mm10_gene_locus_TSS[,5] = toupper(mm10_gene_locus_TSS[,5])
### add TSS position
mm10_SFN_loops = cCRE_gene_JmetID_FS01_geneName_MID_HID_S01_TSS_GeneKM_mat_mm10_SFN_mat
mm10_SFN_loops_TSS = mm10_SFN_loops[,1:3]
k = 0
for (gene_i in unique(mm10_SFN_loops$GeneName)){
	k = k+1
	if (k%%1000==0){print(k)}
	used_n = sum(mm10_SFN_loops$GeneName==gene_i)
	TSS_gene_i = mm10_gene_locus_TSS[mm10_gene_locus_TSS[,5]==gene_i,1:3]
	mm10_SFN_loops_TSS[mm10_SFN_loops$GeneName==gene_i,] = cbind(as.data.frame(rep(TSS_gene_i[1,1], used_n)), rep(TSS_gene_i[1,2], used_n), rep(TSS_gene_i[1,3], used_n))
}
###
mm10_SFN_loops = cbind(mm10_SFN_loops[,1:5], round(rep(1,dim(mm10_SFN_loops)[1])*1000), mm10_SFN_loops[,12], rep('#DEDEDE', dim(mm10_SFN_loops)[1]), mm10_SFN_loops[,c(1:3,7)], rep('.', dim(mm10_SFN_loops)[1]), mm10_SFN_loops_TSS, mm10_SFN_loops$GeneName, rep('.', dim(mm10_SFN_loops)[1]) )
### remove neg
mm10_SFN_loops[(mm10_SFN_loops[,6]<0) | (is.na(mm10_SFN_loops[,6])),6] = 0
### change color
mm10_SFN_loops[mm10_SFN_loops[,7]=='SF',8] = '#FD0606'
mm10_SFN_loops[mm10_SFN_loops[,7]=='F',8] = '#FFAB00'
mm10_SFN_loops[mm10_SFN_loops[,7]=='S',8] = '#696969'
mm10_SFN_loops[mm10_SFN_loops[,7]=='N',8] = '#DEDEDE'
mm10_SFN_loops[mm10_SFN_loops[,7]=='P',8] = '#FF0000'
mm10_SFN_loops[mm10_SFN_loops[,17]=='GATA1',]
mm10_gene_shared_exp[mm10_gene_shared_exp[,5]=='GATA1',]
#mm10_SFN_loops[mm10_SFN_loops[,17]=='CSF1R',]
#mm10_gene_shared_exp[mm10_gene_shared_exp[,5]=='CSF1R',]
options("scipen"=100, "digits"=4)
mm10_SFN_loops$MID = apply(cbind(mm10_SFN_loops$JmetID, mm10_SFN_loops$MID), 1, function(x) paste(x[1], x[2], sep=':'))
write.table(mm10_SFN_loops[mm10_SFN_loops[,6]>=0,], 'mm10.SFN.loop.interact', quote=F, sep='\t', col.names=F, row.names=F)
bash1 = 'cat loop.header.txt > mm10.SFN.loop.interact.tmp'
bash2 = 'cat mm10.SFN.loop.interact >> mm10.SFN.loop.interact.tmp && mv mm10.SFN.loop.interact.tmp mm10.SFN.loop.interact'
system(bash1)
system(bash2)
bash1 = 'cat loop.header.txt > mm10.SFN.loop.interact.tmp'
bash2 = 'cat mm10.SFN.loop.interact | awk -F \'\t\' -v OFS=\'\t\' \'{if ($17=="GATA1") print $0}\' >> mm10.SFN.loop.interact.tmp && mv mm10.SFN.loop.interact.tmp mm10.SFN.loop.GATA1.interact'
system(bash1)
system(bash2)
bash1 = 'cat loop.header.txt > mm10.SFN.loop.interact.tmp'
bash2 = 'cat mm10.SFN.loop.interact | awk -F \'\t\' -v OFS=\'\t\' \'{if ($17=="ALAS2") print $0}\' >> mm10.SFN.loop.interact.tmp && mv mm10.SFN.loop.interact.tmp mm10.SFN.loop.ALAS2.interact'
system(bash1)
system(bash2)
#################################################


#################################################
### write cCRE-cCRE loop file
### add TSS position
hg38_SFN_CC_loops_new = hg38_SFN_loops[hg38_SFN_loops[,1]!='chrXasdasd',]
### enriched F set in each gene
hg38_SFN_set_per_gene = apply(hg38_SFN_loops[hg38_SFN_loops[,1]!='chrXasdasd',c(4,17)], 1, function(x) paste(x[1], x[2], sep='_'))
hg38_SFN_CC_loops_Jmet_gene_new = c()
k = 0
for (Jmet_gene_i in unique(hg38_SFN_set_per_gene)){
	k = k+1
	if (k%%100==0){print(k)}
	hg38_SFN_CC_loops_Jmet_gene_i = hg38_SFN_CC_loops_new[hg38_SFN_set_per_gene==Jmet_gene_i,]
	if (hg38_SFN_CC_loops_Jmet_gene_i[1,5]!=0){
		hg38_SFN_CC_loops_Jmet_gene_i_new = c()
		for (i in 1:dim(hg38_SFN_CC_loops_Jmet_gene_i)[1]){
			for (j in 1:dim(hg38_SFN_CC_loops_Jmet_gene_i)[1]){
				if (i<j){
					hg38_SFN_CC_loops_Jmet_gene_i_new_j = hg38_SFN_CC_loops_Jmet_gene_i[i,]
					hg38_SFN_CC_loops_Jmet_gene_i_new_j[14:18] = hg38_SFN_CC_loops_Jmet_gene_i[j,c(9:13)]
					hg38_SFN_CC_loops_Jmet_gene_i_new = rbind(hg38_SFN_CC_loops_Jmet_gene_i_new, hg38_SFN_CC_loops_Jmet_gene_i_new_j)
				}
			}
		}
		hg38_SFN_CC_loops_Jmet_gene_new = rbind(hg38_SFN_CC_loops_Jmet_gene_new, hg38_SFN_CC_loops_Jmet_gene_i_new)
	}
}

### change colors
hg38_SFN_CC_loops_Jmet_gene_new_unique[hg38_SFN_CC_loops_Jmet_gene_new_unique[,7]=='SF',8] = '#FD0606'
hg38_SFN_CC_loops_Jmet_gene_new_unique[hg38_SFN_CC_loops_Jmet_gene_new_unique[,7]=='F',8] = '#FFAB00'
hg38_SFN_CC_loops_Jmet_gene_new_unique[hg38_SFN_CC_loops_Jmet_gene_new_unique[,7]=='S',8] = '#000000'
hg38_SFN_CC_loops_Jmet_gene_new_unique[hg38_SFN_CC_loops_Jmet_gene_new_unique[,7]=='N',8] = '#DEDEDE'
### 
hg38_SFN_CC_loops_Jmet_gene_new_unique = unique(hg38_SFN_CC_loops_Jmet_gene_new)
options("scipen"=100, "digits"=4)
write.table(hg38_SFN_CC_loops_Jmet_gene_new_unique[hg38_SFN_CC_loops_Jmet_gene_new_unique[,6]>0.2,], 'hg38.SFN.loop.CC.interact', quote=F, sep='\t', col.names=F, row.names=F)
bash1 = 'cat loop.header.txt > hg38.SFN.loop.CC.interact.tmp'
bash2 = 'cat hg38.SFN.loop.CC.interact >> hg38.SFN.loop.CC.interact.tmp && mv hg38.SFN.loop.CC.interact.tmp hg38.SFN.loop.CC.interact'
system(bash1)
system(bash2)


hg38_SFN_loops = cbind(hg38_SFN_loops[,1:5], round(hg38_SFN_loops[12]*1000), hg38_SFN_loops[,13], rep('#DEDEDE', dim(hg38_SFN_loops)[1]), hg38_SFN_loops[,c(1:3,7)], rep('.', dim(hg38_SFN_loops)[1]), hg38_SFN_loops_TSS, hg38_SFN_loops$GeneName, rep('.', dim(hg38_SFN_loops)[1]) )
### remove neg
hg38_SFN_loops[(hg38_SFN_loops[,6]<0) | (is.na(hg38_SFN_loops[,6])),6] = 0
### change color
hg38_SFN_loops[hg38_SFN_loops[,7]=='SF',8] = '#FD0606'
hg38_SFN_loops[hg38_SFN_loops[,7]=='F',8] = '#FFAB00'
hg38_SFN_loops[hg38_SFN_loops[,7]=='S',8] = '#000000'
hg38_SFN_loops[hg38_SFN_loops[,7]=='N',8] = '#DEDEDE'
hg38_SFN_loops[hg38_SFN_loops[,7]=='P',8] = '#FF0000'
hg38_SFN_loops[hg38_SFN_loops[,17]=='GATA1',]
hg38_gene_shared_exp[hg38_gene_shared_exp[,5]=='GATA1',]
hg38_SFN_loops[hg38_SFN_loops[,17]=='CSF1R',]
hg38_gene_shared_exp[hg38_gene_shared_exp[,5]=='CSF1R',]
options("scipen"=100, "digits"=4)
write.table(hg38_SFN_loops[hg38_SFN_loops[,6]>0.2,], 'hg38.SFN.loop.interact', quote=F, sep='\t', col.names=F, row.names=F)
bash1 = 'cat loop.header.txt > hg38.SFN.loop.interact.tmp'
bash2 = 'cat hg38.SFN.loop.interact >> hg38.SFN.loop.interact.tmp && mv hg38.SFN.loop.interact.tmp hg38.SFN.loop.interact'
system(bash1)
system(bash2)
#################################################


### mm10
cCRE_M_Jmeta1 = cbind(cCRE_M_Jmeta, cCRE_M_Jmeta[,4])
for (gene_i in unique(cCRE_M_Jmeta[,5])){
cCRE_M_Jmeta1[cCRE_M_Jmeta[,5]==gene_i,6] = meta_cluster_mat_log2_km$cluster[shared_genes==gene_i]
}
colnames(cCRE_M_Jmeta1)[4] = 'JmetID'
colnames(cCRE_M_Jmeta1)[6] = 'Gene_Jmet_KMID'
write.table(cCRE_M_Jmeta1, 'cCRE.Gene50KB.All_Info.mm10.bed', quote=F, sep='\t', col.names=T, row.names=F)


cCRE_H_Jmeta1[cCRE_H_Jmeta1[,5]=='GATA1',]
cCRE_M_Jmeta1[cCRE_M_Jmeta1[,5]=='GATA1',]

hg38_gene_shared_exp[hg38_gene_shared_exp[,5]=='GATA1',]
mm10_gene_shared_exp[mm10_gene_shared_exp[,5]=='GATA1',]

i = which(shared_genes == 'GATA1')
bed_H = hg38_gene_shared_exp[i,1:3][1,]
bed_M = mm10_gene_shared_exp[i,1:3][1,]
JMeta_i = get_Function_conserve_cCRE(bed_H[1,1], bed_H[1,2], bed_H[1,3], bed_M[1,1], bed_M[1,2], bed_M[1,3])

### check Jmet_GeneKM count in H & M
pdf('Jmet.GeneKM.count.pdf')
plot_lim = c(1,10000)
Human_count = as.numeric(table(apply(cCRE_H_Jmeta1,1, function(x) paste(x[4],x[6]))))
Mouse_count = as.numeric(table(apply(cCRE_M_Jmeta1,1, function(x) paste(x[4],x[6])))) 
Mouse_count_adj = Mouse_count / mean(Mouse_count) * mean(Human_count)
plot(Human_count,  Mouse_count_adj, xlim=plot_lim, ylim=plot_lim, log='xy', cex.axis=2, pch=16)
abline(0,1)
dev.off()
#################################################


#################################################
### check
module_per_gene = apply(cCRE_H_Jmeta, 1, function(x) paste(x[4], x[5], sep=':'))
head(table(module_per_gene)[order(-table(module_per_gene))])

meta_cluster_mat_log2_df[meta_cluster_mat_log2_df[,1]=='FIBCD1',]
meta_cluster_mat_log2_df[meta_cluster_mat_log2_df[,1]=='GATA1',]
meta_cluster_mat_log2_df[meta_cluster_mat_log2_df[,1]=='CD8A',]
meta_cluster_mat_log2_df[meta_cluster_mat_log2_df[,1]=='CSF1R',]

GATA1_exp = get_Function_conserve_cCRE('chrX', 48760001, 48836000, 'chrX', 7919401,8020800)
CD8A_exp = get_Function_conserve_cCRE('chr2', 86758396, 86858396, 'chr6', 71323427,71423427,JointCluster_count_HM)
CSF1R_exp = get_Function_conserve_cCRE('chr5', 150063372, 150163372, 'chr18', 61050598,61150598,JointCluster_count_HM)
IFNG_exp = get_Function_conserve_cCRE('chr12', 68109740, 68209740, 'chr10', 118391046,118491046,JointCluster_count_HM)


length(table(apply(cCRE_H_Jmeta1,1, function(x) paste(x[4],x[6]))))
length(table(apply(cCRE_M_Jmeta1,1, function(x) paste(x[4],x[6]))))


### check EP-pairs num
EP_pair_num = rep(0, dim(hg38_gene_shared_exp)[1])
for (i in 1:dim(hg38_gene_shared_exp)[1]){
	if (i%%1000==0){print(i)}
hg38_gene_shared_exp_i = hg38_gene_shared_exp[i,]
included_rows_i = ((dh_with_JointClusterID[,1]==hg38_gene_shared_exp_i[1,1]) * (dh_with_JointClusterID[,2]>=hg38_gene_shared_exp_i[1,2]) * (dh_with_JointClusterID[,3]<=hg38_gene_shared_exp_i[1,3])) != 0
EP_pair_num[i] = sum(included_rows_i)
}

#> summary(EP_pair_num)
#   Min. 1st Qu.  Median    Mean 3rd Qu.    Max. 
#   0.00    9.00   16.00   17.88   24.00   97.00 
#> sum(EP_pair_num)
#[1] 286027


bedtools intersect -a cCRE.Gene50KB.hg38.bed -b human_ccre_hg38_sf.bed  -c > cCRE.Gene50KB.hg38.SF01.bed
bedtools intersect -a cCRE.Gene50KB.mm10.bed -b mouse_ccre_mm10_sf.bed  -c > cCRE.Gene50KB.mm10.SF01.bed

input_file_hg38 = 'cCRE.Gene50KB.hg38.SF01.bed'
input_file_mm10 = 'cCRE.Gene50KB.mm10.SF01.bed'
output_file = 'Jmet.SF.p.pdf'

dd_hg38 = read.table(input_file_hg38, header=F)
dd_mm10 = read.table(input_file_mm10, header=F)
dd = rbind(dd_hg38, dd_mm10)
### check gene group SF count
Gene_SF_count = c()
for (gene_i in unique(dd[,5])){
Gene_SF_count = rbind(Gene_SF_count, c(sum(dd[,5]==gene_i), sum(dd[dd[,5]==gene_i,7])))
}
Gene_SF_count = cbind(as.data.frame(unique(dd[,5])), Gene_SF_count)
colnames(Gene_SF_count) = c('gene', 'F_cCREn', 'SF_cCREn')

### check Jmet SF proportion
Jmet_SF = c()
for (i in unique(dd[,4])){
Jmet_SF = rbind(Jmet_SF, c(sum(dd[dd[,4]==i,7]!=0), sum(dd[,4]==i)))
}
Jmet_SF = cbind(unique(dd[,4]), Jmet_SF)
Jmet_SF = cbind(Jmet_SF,Jmet_SF[,2]/Jmet_SF[,3])
colnames(Jmet_SF) = c('Jmet','SF_nCREn','F_nCREn','ratio')
Jmet_SF_plot = c()
used_order = c(1,11,5,8,2,6,9,10)
for (Jmeti in used_order){
	Jmet_SF_plot = rbind(Jmet_SF_plot, Jmet_SF[Jmet_SF[,1]==Jmeti,4])
}
rownames(Jmet_SF_plot) = used_order
colnames(Jmet_SF_plot) = NULL

pdf(output_file, width=3)
pheatmap(cbind(Jmet_SF_plot,Jmet_SF_plot), cluster_rows=F, cex=1.5, )
dev.off()

#################################################


chr     start   end     cCRE    AVE     B_B15_50        B_NC14_42 CD34_E_rep1     CD34_E_rep2     CLP_100266      CLP_100267      CMP_100246 CMP_100247      EOS_S006XEH2    EOS_S00BKK      ERY_S002R5      ERY_S002S3
GMP_100256      GMP_100257      HSC_100258      HSC_100259
HUDEP1_rep1     HUDEP1_rep2     HUDEP2_rep1     HUDEP2_rep2     K562_rep1
K562_rep2       LMPP_100268     LMPP_100269     MEP_Donor2596
MEP_Donor7256   MK_S004BTH2     MK_S00VHKH1     MONc_C0011IH1
MONc_C001UYH2   MONp_Prim_mon_C MONp_Prim_mon_F MPP_100272      MPP_100273
NEU_C0011IH2    NEU_C001UYH1    NK_S005YG       NK_S01E4WH0
T_CD4_S008H1    T_CD4_S009W4    T_CD8_C0066PH1  T_CD8_S00C2FH1
chr2    60550400        60552800        chr2_60550400_60552800  12      14
14      14      20      2       2       2       14      18      18      14
14      2       12      2       14      15      15      15      15      2
2       12      12      2       12      12      14      14      14      14
14      12      12      12      18      18      18      18      18      12


bedtools5 = 'cat cCRE.Gene50KB.hg38.SF_S.bed | awk \'{if ($6=="GATA1") print $0}\''
bedtools6 = 'cat cCRE.Gene50KB.mm10.withMID.bed | awk \'{if ($6=="GATA1") print $0}\''
system(bedtools5)
system(bedtools6)
gene_RNA_Jmet_cor[rownames(gene_RNA_Jmet_cor)=='GATA1',]
hg38_gene_shared_exp[hg38_gene_shared_exp[,5]=='GATA1',]
mm10_gene_shared_exp[mm10_gene_shared_exp[,5]=='GATA1',]


bedtools5 = 'cat cCRE.Gene50KB.hg38.SF_S.bed | awk \'{if ($6=="GLOD5") print $0}\''
bedtools6 = 'cat cCRE.Gene50KB.mm10.withMID.bed | awk \'{if ($6=="GLOD5") print $0}\''
system(bedtools5)
system(bedtools6)
gene_RNA_Jmet_cor[rownames(gene_RNA_Jmet_cor)=='GLOD5',]
hg38_gene_shared_exp[hg38_gene_shared_exp[,5]=='GLOD5',]
mm10_gene_shared_exp[mm10_gene_shared_exp[,5]=='GLOD5',]

bedtools5 = 'cat cCRE.Gene50KB.hg38.SF_S.bed | awk \'{if ($6=="CSF1R") print $0}\''
bedtools6 = 'cat cCRE.Gene50KB.mm10.withMID.bed | awk \'{if ($6=="CSF1R") print $0}\''
bedtools5 = 'cat cCRE.Gene50KB.hg38.SF_S.bed | awk \'{if ($6=="CSF1R" && ($5==1 || $7!="NA")) print $0}\''
bedtools6 = 'cat cCRE.Gene50KB.mm10.withMID.bed | awk \'{if ($6=="CSF1R" && ($5==1 || $7!="NA")) print $0}\''
system(bedtools5)
system(bedtools6)
gene_RNA_Jmet_cor[rownames(gene_RNA_Jmet_cor)=='CSF1R',]
hg38_gene_shared_exp[hg38_gene_shared_exp[,5]=='CSF1R',]
mm10_gene_shared_exp[mm10_gene_shared_exp[,5]=='CSF1R',]


###
Gene_Jmet_RNA_shared_genes = rownames(meta_cluster_mat)[is.element(rownames(meta_cluster_mat), rownames(gene_RNA_Jmet_cor))]
meta_cluster_mat_check_shared = meta_cluster_mat[is.element(rownames(meta_cluster_mat), Gene_Jmet_RNA_shared_genes),]
gene_RNA_Jmet_cor_check_shared = gene_RNA_Jmet_cor[is.element(rownames(gene_RNA_Jmet_cor), Gene_Jmet_RNA_shared_genes),]
### remove uniq
gene_RNA_Jmet_cor_check_shared_unique = c()
k = 0
for (gene_i in rownames(meta_cluster_mat_check_shared)){
	k =k+1
	if (k%%10000==0){print(k)}
	if (sum(rownames(gene_RNA_Jmet_cor_check_shared)==gene_i)>1){
		gene_RNA_Jmet_cor_check_shared_unique = rbind(gene_RNA_Jmet_cor_check_shared_unique, colMeans(gene_RNA_Jmet_cor_check_shared[rownames(gene_RNA_Jmet_cor_check_shared)==gene_i,]))
	} else{
		gene_RNA_Jmet_cor_check_shared_unique = rbind(gene_RNA_Jmet_cor_check_shared_unique, gene_RNA_Jmet_cor_check_shared[rownames(gene_RNA_Jmet_cor_check_shared)==gene_i,])

	}
}
rownames(gene_RNA_Jmet_cor_check_shared_unique) = rownames(meta_cluster_mat_check_shared)
colnames(gene_RNA_Jmet_cor_check_shared_unique) = colnames(gene_RNA_Jmet_cor_check_shared)


gene_RNA_Jmet_cor_check_shared_unique = gene_RNA_Jmet_cor_check_shared_unique[order(rownames(gene_RNA_Jmet_cor_check_shared_unique)),order(colnames(gene_RNA_Jmet_cor_check_shared_unique))]
meta_cluster_mat_check_shared = meta_cluster_mat_check_shared[order(rownames(meta_cluster_mat_check_shared)),order(colnames(meta_cluster_mat_check_shared))]

gene_RNA_Jmet_cor_check_shared_unique_vec = as.numeric(as.matrix(gene_RNA_Jmet_cor_check_shared_unique))
meta_cluster_mat_check_shared_vec = as.numeric(as.matrix(meta_cluster_mat_check_shared))


png('gene_RNA_Jmet_cor.png')
used_id = sample(length(gene_RNA_Jmet_cor_check_shared_unique_vec), 10000)
heatscatter(gene_RNA_Jmet_cor_check_shared_unique_vec[used_id], meta_cluster_mat_check_shared_vec[used_id], log='y')
dev.off()
```
