## Supplementary material for "Interspecies regulatory landscapes and elements revealed by novel joint systematic integration of human and mouse blood cell epigenomes": Scripts for IDEAS modeling of epigenetic states jointly in human and mouse: SF_S_N.data.preprocess.sh

```
cd /Users/guanjuexiang/Documents/projects/analysis/04_05_2022_coe/coe_analysis/SF_S_N/

cat mouse_ccre_mm10_sf.bed mouse_ccre_mm10_s.bed mouse_ccre_mm10_n.bed | sort -k1,1 -k2,2n > mouse_ccre_mm10.withMID.bed
cat mouse_ccre_hg38_sf.bed mouse_ccre_hg38_s.bed | awk -F '\t' -v OFS='\t' '{print $2,$3,$4,$1}' | sort -k1,1 -k2,2n > mouse_ccre_hg38_sf_s.withMID.bed

cat human_ccre_hg38_sf.bed human_ccre_hg38_s.bed human_ccre_hg38_n.bed | sort -k1,1 -k2,2n > human_ccre_hg38.withHID.bed
cat human_ccre_mm10_sf.bed human_ccre_mm10_s.bed | awk -F '\t' -v OFS='\t' '{print $2,$3,$4,$1}' | sort -k1,1 -k2,2n > human_ccre_mm10_sf_s.withHID.bed

bedtools map -a human_ccre_hg38.withHID.bed -b mouse_ccre_hg38_sf_s.withMID.bed -c 4 -o concat -null NA > human_ccre_hg38.withHID.with_Mouse_S_MID.bed
bedtools map -a mouse_ccre_mm10.withMID.bed -b human_ccre_mm10_sf_s.withHID.bed -c 4 -o concat -null NA > mouse_ccre_mm10.withMID.with_Human_S_HID.bed


cd /Users/guanjuexiang/Documents/projects/analysis/04_05_2022_coe/coe_analysis/


cat cCRE.Gene50KB.hg38.bed | awk -F '\t' '{if ($1!="NA" && $1!="chr") print $0}' | sort -k1,1 -k2,2n > cCRE.Gene50KB.hg38.sort.bed 
bedtools map -a cCRE.Gene50KB.hg38.sort.bed -b SF_S_N/human_ccre_hg38.withHID.with_Mouse_S_MID.bed -c 5 -o concat -null NA > cCRE.Gene50KB.hg38.SF_S.bed

cat cCRE.Gene50KB.mm10.bed | awk -F '\t' '{if ($1!="NA" && $1!="chr") print $0}' | sort -k1,1 -k2,2n > cCRE.Gene50KB.mm10.sort.bed 
bedtools map -a cCRE.Gene50KB.mm10.sort.bed -b SF_S_N/mouse_ccre_mm10.withMID.with_Human_S_HID.bed -c 4 -o concat -null NA > cCRE.Gene50KB.mm10.withMID.bed


cat cCRE.Gene50KB.hg38.SF_S.bed | awk '{if ($6=="GATA1") print $0}' 

cat cCRE.Gene50KB.mm10.withMID.bed | awk '{if ($6=="GATA1") print $0}' 
```
