## Supplementary material for "Interspecies regulatory landscapes and elements revealed by novel joint systematic integration of human and mouse blood cell epigenomes": Scripts for IDEAS modeling of epigenetic states jointly in human and mouse: prepare_snapshot_input.R

```
#d = read.table('S3V2_IDEAS_hg38_ccre2.cCRE.M.notall0.rmallNEU.withid.coe_mat.txt', header=T)
HsP_cCRE = read.table('../coe_analysis/S3V2_IDEAS_hg38_ccre2.cCRE.M.withid.atProximal.bed', header=F)
d = read.table('../coe_analysis/S3V2_IDEAS_hg38_ccre2.cCRE.M.notall0.rmallNEU.withid.coe_mat.PDmerged.txt', header=T)
dPDsep = read.table('../coe_analysis/S3V2_IDEAS_hg38_ccre2.cCRE.M.notall0.rmallNEU.withid.coe_mat.txt', header=T)

ds = d[,-c(1:4)]

library(pheatmap)

set.seed(2019)
#used_row = sample(dim(ds)[1], dim(ds)[1]
dss = ds

### ct dist cor
dPDsep = dPDsep[,!is.element(colnames(dPDsep), c('chr', 'start', 'end', 'id', 'NEU_C0011IH2_P','NEU_C001UYH1_P', 'NEU_C0011IH2_D','NEU_C001UYH1_D'))]
ds_forcor = dPDsep[,(dim(dPDsep)[2]/2+1):dim(dPDsep)[2]]
#ds_forcor[ds_forcor>quantile(as.numeric(as.matrix(ds)),1)] = quantile(as.numeric(as.matrix(ds)),1)
ds_cor_D = cor(ds_forcor)
#ds_cor_D = cor(ds)
hclust_cor = hclust(as.dist(1-ds_cor_D))
hclust_cor_order = hclust_cor$order

png('cCRE_coe.human.heatmap.D.cttree.png', height=600, width=1200)
plot(hclust_cor, hang = -1, cex = 2, lwd=2)
dev.off()

rep1 = c(1, 2, 4, 6, 8, 10, 12, 14, 16, 18, 20, 22, 24, 26, 28, 30, 32, 34, 36, 38, 40)
rep2 = c(1, 3, 5, 7, 9, 11, 13, 15, 17, 19, 21, 23, 25, 27, 29, 31, 33, 35, 37, 39, 41)

dss_ct = c()
for (i in 1:length(rep1)){
	dss_ct = cbind(dss_ct, rowMeans(cbind(dss[,rep1[i]], dss[,rep2[i]])))
}
ct_list = apply(cbind(colnames(ds_forcor)[rep1]), 1, function(x) unlist(strsplit(x, '_'))[1])
ct_list[length(ct_list)-1] = 'CD4'
ct_list[length(ct_list)] = 'CD8'
colnames(dss_ct) = ct_list

### write bg
for (i in 1:length(ct_list)){
	bg_i = cbind(d[,1:3], dss_ct[,i])
	write.table(bg_i, paste0('Snapshot/input_data/signal_bgbw/', ct_list[i], '.esRP.bedgraph'), sep='\t', quote=F, col.names=F, row.names=F)
}

### get binary
sig_vec0 = as.numeric(dss_ct)
sig_vec = sig_vec0#[sig_vec0!=0]
print(length(sig_vec))
length_pre = length(sig_vec)
for (i in 1:100){
sig_vec_zp = pnorm((sig_vec-mean(sig_vec))/sd(sig_vec), lower.tail=F)
sig_vec = sig_vec[sig_vec_zp>=0.01]
if (length_pre == length(sig_vec)){break}
length_pre = length(sig_vec)
print(length(sig_vec))
}
sig_vec_thresh = max(sig_vec)
print(sig_vec_thresh)

png('hist.coe.png')
hist(sig_vec0, breaks=50)
abline(v=sig_vec_thresh)
box()
dev.off()

###
dss_ct_binary = (dss_ct > sig_vec_thresh)*1
### write peak bed
for (i in 1:length(ct_list)){
	pk_i = cbind(d[dss_ct_binary[,i]!=0,1:3])
	write.table(pk_i, paste0('Snapshot/input_data/peak_bed/', ct_list[i], '.pk.bed'), sep='\t', quote=F, col.names=F, row.names=F)
}
```
