## Supplementary material for "Interspecies regulatory landscapes and elements revealed by novel joint systematic integration of human and mouse blood cell epigenomes": Scripts for IDEAS modeling of epigenetic states jointly in human and mouse: get_cCRE_coe_mat.R

```
coe = read.table('../coe_analysis/statep_rna_coe_heatmap.HM.all.ccre.withcorfilter.AVE.txt', header=T)

###################
###### human ######
ct_list_human = c('AVE', 'B_B15_50', 'B_NC14_42', 'CD34_E_rep1', 'CD34_E_rep2', 'CLP_100266', 'CLP_100267', 'CMP_100246', 'CMP_100247', 'EOS_S006XEH2', 'EOS_S00BKK', 'ERY_S002R5', 'ERY_S002S3', 'GMP_100256', 'GMP_100257', 'HSC_100258', 'HSC_100259', 'HUDEP1_rep1', 'HUDEP1_rep2', 'HUDEP2_rep1', 'HUDEP2_rep2', 'K562_rep1', 'K562_rep2', 'LMPP_100268', 'LMPP_100269', 'MEP_Donor2596', 'MEP_Donor7256', 'MK_S004BTH2', 'MK_S00VHKH1', 'MONc_C0011IH1', 'MONc_C001UYH2', 'MONp_Prim_mon_C', 'MONp_Prim_mon_F', 'MPP_100272', 'MPP_100273', 'NEU_C0011IH2', 'NEU_C001UYH1', 'NK_S005YG', 'NK_S01E4WH0', 'T_CD4_S008H1', 'T_CD4_S009W4', 'T_CD8_C0066PH1', 'T_CD8_S00C2FH1')

### read state count
Hs = read.table('../coe_analysis/S3V2_IDEAS_hg38_ccre2.cCRE.M.notall0.rmallNEU.withid.S0.mat.txt', header=F)

### normalize dist
HsP0 = as.matrix(Hs[,-c(1:4)])
HsD0 = as.matrix(Hs[,-c(1:4)])

### state count * coe
HsP = log(HsP0+1) * coe[1,1]
HsD = log(HsD0+1) * coe[1,2]

for (i in 1:24){
	print(i)
Hs_i = read.table(paste('../coe_analysis/S3V2_IDEAS_hg38_ccre2.cCRE.M.notall0.rmallNEU.withid.S',i,'.mat.txt', sep=''), header=F)
HsP0 = as.matrix(Hs_i[,-c(1:4)])
HsD0 = as.matrix(Hs_i[,-c(1:4)])
HsP = HsP + log(HsP0+1) * coe[i+1,1]
HsD = HsD + log(HsD0+1) * coe[i+1,2]
}

### merge P & D
#HsPD = HsD

### add colnames
colnames(HsP) = paste(ct_list_human, '_P', sep='')
colnames(HsD) = paste(ct_list_human, '_D', sep='')

### output mat
HsPD_mat = cbind(Hs[,1:4], (HsP), (HsD))
colnames(HsPD_mat)[1:4] = c('chr','start','end','id')

write.table(HsPD_mat, '../coe_analysis/S3V2_IDEAS_hg38_ccre2.cCRE.M.notall0.rmallNEU.withid.coe_mat.txt', quote=F, col.names=T, row.names=F, sep='\t')


###################
###### mouse ######
ct_list_mouse = c('AVE', 'B_r1', 'B_r2', 'CFUE_r1', 'CFUMK_r1', 'CLP_r1', 'CMP_r1', 'ER4_r1', 'ER4_r2', 'ERY_ad_r1', 'ERY_ad_r2', 'ERY_fl_r1', 'ERY_fl_r2', 'G1E_r1', 'G1E_r2', 'GMP_r1', 'HPC7_r1', 'LSK_r1', 'MEL_r1', 'MEL_r2', 'MEP_r1', 'MK_fl_r1', 'MK_fl_r2', 'MON_r1', 'NEU_r1', 'NK_r1', 'T_CD4_r1', 'T_CD8_r1', 'iMEL_r1', 'iMEL_r2', 'iMK_r1', 'iMK_r2')

### read state count
Hs = read.table('../coe_analysis_mouse/S3V2_IDEAS_mm10_ccre2.cCRE.M.notall0.withid.S0.mat.txt', header=F)

### normalize dist
HsP0 = as.matrix(Hs[,-c(1:4)])
HsD0 = as.matrix(Hs[,-c(1:4)])

### state count * coe
HsP = log(HsP0+1) * coe[1,1]
HsD = log(HsD0+1) * coe[1,2]

for (i in 1:24){
	print(i)
Hs_i = read.table(paste('../coe_analysis_mouse/S3V2_IDEAS_mm10_ccre2.cCRE.M.notall0.withid.S',i,'.mat.txt', sep=''), header=F)
HsP0 = as.matrix(Hs_i[,-c(1:4)])
HsD0 = as.matrix(Hs_i[,-c(1:4)])
HsP = HsP + log(HsP0+1) * coe[i+1,1]
HsD = HsD + log(HsD0+1) * coe[i+1,2]
}

### merge P & D
#HsPD = HsD

### add colnames
colnames(HsP) = paste(ct_list_mouse, '_P', sep='')
colnames(HsD) = paste(ct_list_mouse, '_D', sep='')

### output mat
HsPD_mat = cbind(Hs[,1:4], (HsP), (HsD))
colnames(HsPD_mat)[1:4] = c('chr','start','end','id')

write.table(HsPD_mat, '../coe_analysis_mouse/S3V2_IDEAS_mm10_ccre2.cCRE.M.notall0.withid.coe_mat.txt', quote=F, col.names=T, row.names=F, sep='\t')
```
