## Supplementary material for "Interspecies regulatory landscapes and elements revealed by novel joint systematic integration of human and mouse blood cell epigenomes": Scripts for IDEAS modeling of epigenetic states jointly in human and mouse: get_cor_heatmap.R

```
d = read.table('S3V2_IDEAS_hg38_ccre2.cCRE.M.notall0.rmallNEU.withid.coe_mat.txt', header=T)

ds = d[,-c(1:4)]

library(pheatmap)

set.seed(2019)
used_row = sample(dim(ds)[1], 50000)
ds_cor_P = cor(ds[used_row,1:(dim(ds)[2]/2)])
ds_cor_D = cor(ds[used_row,(dim(ds)[2]/2+1):dim(ds)[2]])

pdf('state_coe_cor.human.heatmap.P.pdf', width=10, height=10)
breaksList = seq(-1, 1, by = 0.001)
my_colorbar=colorRampPalette(c('blue', 'blue', 'blue', 'white', 'red'))(n = length(breaksList))
col_breaks = c(seq(0, 2000,length=33))
pheatmap(ds_cor_P, cluster_cols = T,cluster_rows=T, clustering_method = 'complete',annotation_names_row=TRUE,annotation_names_col=TRUE,show_rownames=TRUE,show_colnames=TRUE)
dev.off()

pdf('state_coe_cor.human.heatmap.D.pdf', width=10, height=10)
pheatmap(ds_cor_D, cluster_cols = T,cluster_rows=T, clustering_method = 'complete',annotation_names_row=TRUE,annotation_names_col=TRUE,show_rownames=TRUE,show_colnames=TRUE)
dev.off()

pdf('state_coe_cor.human.heatmap.PD.pdf', width=10, height=10)
pheatmap(ds_cor_P/2+ds_cor_D/2, cluster_cols = T,cluster_rows=T, clustering_method = 'complete',annotation_names_row=TRUE,annotation_names_col=TRUE,show_rownames=TRUE,show_colnames=TRUE)
dev.off()


set.seed(2019)
used_row = sample(dim(ds)[1], dim(ds)[1])
dss = ds[used_row,(dim(ds)[2]/2+1):dim(ds)[2]]

ds_cor_P = cor(ds[used_row,1:(dim(ds)[2]/2)])
ds_cor_D = cor(ds[used_row,(dim(ds)[2]/2+1):dim(ds)[2]])

hclust_cor = hclust(as.dist(1-ds_cor_D))
hclust_cor_order = hclust_cor$order

png('cCRE_coe.human.heatmap.D.cttree.png', height=600, width=1200)
plot(hclust_cor, hang = -1, cex = 2, lwd=2)
dev.off()


png('cCRE_coe.human.heatmap.D.png', width=1200, height=1800)
plot_lim_PD = quantile(as.numeric(as.matrix(dss)),0.999)
breaksList = seq(-plot_lim_PD, plot_lim_PD, by = 0.001)
my_colorbar=colorRampPalette(c('blue', 'white', 'red'))(n = length(breaksList))
col_breaks = c(seq(0, 2000,length=33))
dss_plot = dss
set.seed(2019)
dss_km = kmeans(dss, 50)
dss_plot[dss>plot_lim_PD]=plot_lim_PD
pheatmap(dss_plot[order(dss_km$cluster),hclust_cor_order], color=my_colorbar, breaks = breaksList, cluster_cols = F,cluster_rows=F, clustering_method = 'complete',annotation_names_row=F,annotation_names_col=TRUE,show_rownames=F,show_colnames=TRUE)
dev.off()

pdf('cCRE_coe.human.heatmap.D.pdf', width=8, height=12)
plot_lim_PD = max(dss)
breaksList = seq(-plot_lim_PD, plot_lim_PD, by = 0.001)
my_colorbar=colorRampPalette(c('blue', 'white', 'red'))(n = length(breaksList))
col_breaks = c(seq(0, 2000,length=33))
dss_plot = dss
dss_km = kmeans(dss, 20)
dss_plot[dss>plot_lim_PD]=plot_lim_PD
pheatmap(dss_plot[order(dss_km$cluster),], color=my_colorbar, breaks = breaksList, cluster_cols = T,cluster_rows=F, clustering_method = 'complete',annotation_names_row=F,annotation_names_col=TRUE,show_rownames=F,show_colnames=TRUE)
dev.off()


######

d = read.table('S3V2_IDEAS_mm10_ccre2.cCRE.M.notall0.withid.coe_mat.txt', header=T)

ds = d[,-c(1:4)]


set.seed(2019)
used_row = sample(dim(ds)[1], dim(ds)[1])
dss = ds[used_row,(dim(ds)[2]/2+1):dim(ds)[2]]

ds_cor_P = cor(ds[used_row,1:(dim(ds)[2]/2)])
ds_cor_D = cor(ds[used_row,(dim(ds)[2]/2+1):dim(ds)[2]])

hclust_cor = hclust(as.dist(1-ds_cor_D))
hclust_cor_order = hclust_cor$order

png('cCRE_coe.mouse.heatmap.D.cttree.png', height=600, width=1200)
plot(hclust_cor, hang = -1, cex = 2, lwd=2)
dev.off()


png('cCRE_coe.mouse.heatmap.D.png', width=600, height=900)
plot_lim_PD = quantile(as.numeric(as.matrix(dss)),0.999)
breaksList = seq(-plot_lim_PD, plot_lim_PD, by = 0.001)
my_colorbar=colorRampPalette(c('blue', 'white', 'red'))(n = length(breaksList))
col_breaks = c(seq(0, 2000,length=33))
dss_plot = dss
set.seed(2019)
dss_km = kmeans(dss, 50)
dss_plot[dss>plot_lim_PD]=plot_lim_PD
pheatmap(dss_plot[order(dss_km$cluster),hclust_cor_order], color=my_colorbar, breaks = breaksList, cluster_cols = F,cluster_rows=F, clustering_method = 'complete',annotation_names_row=F,annotation_names_col=TRUE,show_rownames=F,show_colnames=TRUE)
dev.off()


set.seed(2019)
used_row = sample(dim(ds)[1], 50000)
ds_cor_P = cor(ds[used_row,1:(dim(ds)[2]/2)])
ds_cor_D = cor(ds[used_row,(dim(ds)[2]/2+1):dim(ds)[2]])

pdf('state_coe_cor.mouse.heatmap.P.pdf', width=10, height=10)
breaksList = seq(-1, 1, by = 0.001)
my_colorbar=colorRampPalette(c('blue', 'blue', 'blue', 'white', 'red'))(n = length(breaksList))
col_breaks = c(seq(0, 2000,length=33))
pheatmap(ds_cor_P, cluster_cols = T,cluster_rows=T, clustering_method = 'complete',annotation_names_row=TRUE,annotation_names_col=TRUE,show_rownames=TRUE,show_colnames=TRUE)
dev.off()

pdf('state_coe_cor.mouse.heatmap.D.pdf', width=10, height=10)
pheatmap(ds_cor_D, cluster_cols = T,cluster_rows=T, clustering_method = 'complete',annotation_names_row=TRUE,annotation_names_col=TRUE,show_rownames=TRUE,show_colnames=TRUE)
dev.off()

pdf('state_coe_cor.mouse.heatmap.PD.pdf', width=10, height=10)
pheatmap(ds_cor_P/2+ds_cor_D/2, cluster_cols = T,cluster_rows=T, clustering_method = 'complete',annotation_names_row=TRUE,annotation_names_col=TRUE,show_rownames=TRUE,show_colnames=TRUE)
dev.off()
```
