## Supplementary material for "Interspecies regulatory landscapes and elements revealed by novel joint systematic integration of human and mouse blood cell epigenomes": Scripts for IDEAS modeling of epigenetic states jointly in human and mouse: get_coe_umap.R

```
d = read.table('S3V2_IDEAS_hg38_ccre2.cCRE.M.notall0.rmallNEU.withid.coe_mat.txt', header=T)

ds = d[,-c(1:4)]

library(pheatmap)
library(umap)
library(LSD)


###### For Distal
set.seed(2019)
used_row = sample(dim(ds)[1], 10000)
dss = ds[used_row,(dim(ds)[2]/2+1):dim(ds)[2]]
### run umap
dss_umap = umap(round(dss,2), preserve.seed=T)

cbind(1:dim(dss)[2], colnames(dss))

dir.create('coe_umap_D')
### plot heatscatter plot
png('coe_umap_D/state_coe.human.umap.D.png')
heatscatter(dss_umap$layout[,1], dss_umap$layout[,2])
dev.off()

for (i in 1:dim(dss)[2]){
	ct_i = colnames(dss)[i]
png(paste('coe_umap_D/', 'state_coe.human.umap.D.', ct_i,'.png', sep=''))
rbPal = colorRampPalette(c('gray90','red'))
color_sig_range = (dss[,i])
color_sig_range[color_sig_range>quantile(as.numeric(as.matrix(dss)),0.99)] = quantile(as.numeric(as.matrix(dss)),0.99)
plot_color = rbPal(50)[as.numeric(cut(color_sig_range,breaks = 50))]
plot(dss_umap$layout[,1], dss_umap$layout[,2], col=plot_color, pch=16)
dev.off()
}


###### For Proximal
set.seed(2019)
used_row = sample(dim(ds)[1], 10000)
dss = ds[used_row,1:(dim(ds)[2]/2)]
### run umap
dss_umap = umap(round(dss, 3), preserve.seed=T)

cbind(1:dim(dss)[2], colnames(dss))

dir.create('coe_umap_P')
### plot heatscatter plot
png('coe_umap_P/state_coe.human.umap.P.png')
heatscatter(dss_umap$layout[,1], dss_umap$layout[,2])
dev.off()

for (i in 1:dim(dss)[2]){
	ct_i = colnames(dss)[i]
png(paste('coe_umap_P/', 'state_coe.human.umap.P.', ct_i,'.png', sep=''))
rbPal = colorRampPalette(c('gray90','red'))
color_sig_range = (dss[,i])
color_sig_range[color_sig_range>quantile(as.numeric(as.matrix(dss)),0.99)] = quantile(as.numeric(as.matrix(dss)),0.99)
plot_color = rbPal(50)[as.numeric(cut(color_sig_range,breaks = 50))]
plot(dss_umap$layout[,1], dss_umap$layout[,2], col=plot_color, pch=16)
dev.off()
}


###### For Proximal & Distal
set.seed(2019)
used_row = sample(dim(ds)[1], 10000)
dssP = as.matrix(ds[used_row,1:(dim(ds)[2]/2)])
dssD = as.matrix(ds[used_row,(dim(ds)[2]/2+1):dim(ds)[2]])
dssDadj = (dssD - mean(dssD))/sd(dssD) * sd(dssP) + mean(dssP)
dss = cbind(dssP, dssDadj)

### run umap
dss_umap = umap(dss, preserve.seed=T)

cbind(1:dim(dss)[2], colnames(dss))

dir.create('coe_umap_PD')
### plot heatscatter plot
png('coe_umap_PD/state_coe.human.umap.PD.png')
heatscatter(dss_umap$layout[,1], dss_umap$layout[,2])
dev.off()

for (i in 1:dim(dss)[2]){
	ct_i = colnames(dss)[i]
png(paste('coe_umap_PD/', 'state_coe.human.umap.PD.', ct_i,'.png', sep=''))
rbPal = colorRampPalette(c('gray90','red'))
color_sig_range = (dss[,i])
color_sig_range[color_sig_range>quantile(as.numeric(as.matrix(dss)),0.99)] = quantile(as.numeric(as.matrix(dss)),0.99)
plot_color = rbPal(50)[as.numeric(cut(color_sig_range,breaks = 50))]
plot(dss_umap$layout[,1], dss_umap$layout[,2], col=plot_color, pch=16)
dev.off()
}
```
