## Supplementary material for "Interspecies regulatory landscapes and elements revealed by novel joint systematic integration of human and mouse blood cell epigenomes": Scripts for IDEAS modeling of epigenetic states jointly in human and mouse: get_cor_heatmap_som_metacluster.R

```
conda create -n som r=4.1 r-FlowSOM

d = read.table('S3V2_IDEAS_hg38_ccre2.cCRE.M.notall0.rmallNEU.withid.coe_mat.txt', header=T)

ds = d[,-c(1:4)]

library(pheatmap)

set.seed(2019)
#used_row = sample(dim(ds)[1], dim(ds)[1])
dss = ds[,(dim(ds)[2]/2+1):dim(ds)[2]]

ds_cor_P = cor(ds[,1:(dim(ds)[2]/2)])
ds_cor_D = cor(ds[,(dim(ds)[2]/2+1):dim(ds)[2]])

hclust_cor = hclust(as.dist(1-ds_cor_D))
hclust_cor_order = hclust_cor$order

png('cCRE_coe.human.heatmap.D.cttree.png', height=600, width=1200)
plot(hclust_cor, hang = -1, cex = 2, lwd=2)
dev.off()

rep1 = c(1, 2, 4, 6, 8, 10, 12, 14, 16, 18, 20, 22, 24, 26, 28, 30, 32, 34, 36, 38, 40, 42)
rep2 = c(1, 3, 5, 7, 9, 11, 13, 15, 17, 19, 21, 23, 25, 27, 29, 31, 33, 35, 37, 39, 41, 43)

dss_ct = c()
for (i in 1:length(rep1)){
	dss_ct = cbind(dss_ct, rowMeans(cbind(dss[,rep1[i]], dss[,rep2[i]])))
}

library(FlowSOM)

#d = read.table('S3V2_IDEAS_hg38_ccre2.cCRE.M.notall0.rmallNEU.withid.coe_mat.txt', header=T)
HsP_cCRE = read.table('../coe_analysis/S3V2_IDEAS_hg38_ccre2.cCRE.M.withid.atProximal.bed', header=F)
d = read.table('../coe_analysis/S3V2_IDEAS_hg38_ccre2.cCRE.M.notall0.rmallNEU.withid.coe_mat.PDmerged.txt', header=T)
dPDsep = read.table('../coe_analysis/S3V2_IDEAS_hg38_ccre2.cCRE.M.notall0.rmallNEU.withid.coe_mat.txt', header=T)


library(data.table)
d = as.data.frame(fread('../coe_analysis/S3V2_IDEAS_hg38_ccre2.cCRE.M.notall0.rmallNEU.withid.coe_mat.PDmerged.txt', header=T))


ds = d[,-c(1:4)]

library(pheatmap)

set.seed(2019)
used_row = sample(dim(ds)[1], 10000)
dss = ds[used_row,]
write.table(dss, 'tmp.txt')


### som
library(kohonen)

som_grid <- somgrid(xdim = 20, ydim=20, topo="hexagonal")

som_model <- som(as.matrix(dss), 
    grid=som_grid, 
    rlen=500, 
    alpha=c(0.05,0.01), 
    keep.data = TRUE )

plot(som_model, type="changes")

plot(som_model, type="count", main="Node Counts")

#plot(som_model, type="codes")

coolBlueHotRed <- function(n, alpha = 1) {rainbow(n, end=4/6, alpha=alpha)[n:1]}
plot(som_model, type = "property", property = getCodes(som_model)[,4], main=colnames(getCodes(som_model))[4], palette.name=coolBlueHotRed)

som.events <- som_model$codes[[1]]
som.events.colors <- rgb(som.events[,1], som.events[,2], som.events[,3], maxColorValue = 255)
som.dist <- as.matrix(dist(som.events))


clusterMeanDist <- function(clusters){
  cluster.means = c()
  
  for(c in unique(clusters)){
    temp.members <- which(clusters == c)
    
    if(length(temp.members) > 1){
      temp.dist <- som.dist[temp.members,]
      temp.dist <- temp.dist[,temp.members]
      cluster.means <- append(cluster.means, mean(temp.dist))
    }else(cluster.means <- 0)
  }
  
  return(mean(cluster.means))
  
}

try.k <- 2:100
cluster.dist.eval <- as.data.frame(matrix(ncol = 3, nrow = (length(try.k))))
colnames(cluster.dist.eval) <- c('k', 'kmeans', 'hclust')

for(i in 1:length(try.k)) {
  cluster.dist.eval[i, 'k'] <- try.k[i]
  cluster.dist.eval[i, 'kmeans'] <- clusterMeanDist(kmeans(som.events, centers = try.k[i], iter.max = 20)$cluster)
  cluster.dist.eval[i, 'hclust'] <- clusterMeanDist(cutree(hclust(vegdist(som.events)), k = try.k[i]))
}

plot(cluster.dist.eval[, 'kmeans'] ~ try.k,
     type = 'l')

lines(cluster.dist.eval[, 'hclust'] ~ try.k,
      col = 'red')

legend('topright',
       legend = c('k-means', 'hierarchical'),
       col = c('black', 'red'),
       lty = c(1, 1))


cluster.tries <- list()

for(k in c(20,30,50)){
  print(k)

  ## k-means clustering
  
  som.cluster.k <- kmeans(som.events, centers = k, iter.max = 100, nstart = 10)$cluster # k-means
  
  ## hierarchical clustering
  
  som.dist <- dist(som.events) # hierarchical, step 1
  som.cluster.h <- cutree(hclust(som.dist), k = k) # hierarchical, step 2
  
  ## capture outputs
  
  cluster.tries[[paste0('som.cluster.k.', k)]] <- som.cluster.k
  cluster.tries[[paste0('som.cluster.h.', k)]] <- som.cluster.h
}

## Take a look at the various clusters.  You're looking for the algorithm that produces the
## least fragmented clusters.

som.events <- som_model$codes[[1]]


plotSOM <- function(clusters){
  plot(som_model,
       type = 'mapping',
       keepMargins = F,
       bg = som_color_vec,
       col = NA)
  add.cluster.boundaries(som_model, clusters)
}


rbPal <- colorRampPalette(c('white','green4'))
som.events.1 = som.events[,38]
som.events.1[som.events.1>quantile(som.events, 0.99)]=quantile(som.events, 0.99)
som_color_vec <- rbPal(10)[as.numeric(cut(som.events.1,breaks = 10))]
plotSOM(cluster.tries$som.cluster.h.50)

rbPal <- colorRampPalette(c('white','red'))
som.events.1 = som.events[,12]
som.events.1[som.events.1>quantile(som.events, 0.99)]=quantile(som.events, 0.99)
som_color_vec <- rbPal(10)[as.numeric(cut(som.events.1,breaks = 10))]
plotSOM(cluster.tries$som.cluster.h.30)

rbPal <- colorRampPalette(c('white','blue'))
som.events.1 = som.events[,30]
som.events.1[som.events.1>quantile(som.events, 0.99)]=quantile(som.events, 0.99)
som_color_vec <- rbPal(10)[as.numeric(cut(som.events.1,breaks = 10))]
plotSOM(cluster.tries$som.cluster.h.30)

rbPal <- colorRampPalette(c('blue','white','red'))
som.events.1 = som.events[,12] - som.events[,30]
som.events.1[som.events.1>quantile(som.events, 0.99)]=quantile(som.events, 0.99)
som.events.1[som.events.1<(-quantile(som.events, 0.99))]=-quantile(som.events, 0.99)
som_color_vec <- rbPal(10)[as.numeric(cut(som.events.1,breaks = 10))]
plotSOM(cluster.tries$som.cluster.h.30)


### ct dist cor
dPDsep = dPDsep[,!is.element(colnames(dPDsep), c('chr', 'start', 'end', 'id', 'NEU_C0011IH2_P','NEU_C001UYH1_P', 'NEU_C0011IH2_D','NEU_C001UYH1_D'))]
ds_forcor = dPDsep[,(dim(dPDsep)[2]/2+1):dim(dPDsep)[2]]


flowSOM.res <- ReadInput('tmp.txt')

flowSOM.res <- BuildSOM(dss)


fileName <- system.file("extdata","lymphocytes.fcs",package="FlowSOM")


   flowSOM.res <- ReadInput(fileName, compensate=TRUE,transform=TRUE,
                            scale=TRUE)

   flowSOM.res <- BuildSOM(flowSOM.res,colsToUse=c(9,12,14:18))

   flowSOM.res <- BuildMST(flowSOM.res)
   
   # Apply metaclustering
   metacl <- MetaClustering(flowSOM.res$map$codes,
                            "metaClustering_consensus",
                            max=10)
   
   # Get metaclustering per cell
   flowSOM.clustering <- metacl[flowSOM.res$map$mapping[,1]]    


within_c_dist = function(x, c_vec){
	within_dist = 0
	for (i in min(c_vec):max(c_vec)){
		x_i = x[c_vec==i,]
		x_i_colm = colMeans(x_i)
		x_i_dist = sum(apply(x_i, 1, function(x) sum((x-x_i_colm)^2)))
		within_dist = within_dist+x_i_dist
	}
	return(within_dist)
}

set.seed(2019)
dist_ratio = c()
for (n in seq(10,200,10)){
	print(n)
	dss_km_n = kmeans(dss_ct, n)
	dist_km = within_c_dist(dss_ct, dss_km_n$cluster)
	dss_km_ave_n = kmeans(cbind(rowMeans(dss_ct)), n)
	dist_ave = within_c_dist(dss_ct, dss_km_ave_n$cluster)
	dist_ratio = rbind(dist_ratio, c(n, dist_km, dist_ave, dss_km_n$totss))	
}

pdf('km_k_dist.pdf')
plot(c(1,seq(10,200,10)), c(dss_km_n$totss, dist_ratio[,2]))
lines(c(1,seq(10,200,10)), c(dss_km_n$totss, dist_ratio[,2]))
dev.off()

library(MASS)

### get 100 round of km k=50
km_k = 50
km_centers = c()
iter_n = 100
for (r in 1:iter_n){
	print(r)
	dss_km_i = kmeans(dss_ct, km_k)
	for (k_i in 1:km_k){
	km_centers = rbind(km_centers, colMeans(dss_ct[dss_km_i$cluster==k_i,]))
	}
}

### get reproducible clusters
set.seed(2019)
kmkm = kmeans(km_centers, km_k)

pdf('kmkm_n.hist.pdf')
hist(table(kmkm$cluster), breaks=20)
dev.off()

kmkm_n_z = ( table(kmkm$cluster) - mean(table(kmkm$cluster)) ) /sd(table(kmkm$cluster))
#used_kmkm_centers = which(pnorm(kmkm_n_z, lower.tail=TRUE)>=0.05)
used_kmkm_centers = which(table(kmkm$cluster)>=(iter_n/2))
### get new km centers
km_new_centers = c()
for (i in 1:length(used_kmkm_centers)){
km_new_centers = rbind(km_new_centers, colMeans(as.matrix(km_centers)[kmkm$cluster==i,]))
}

### reassign cCREs into reproducible clusters
x_dist_to_newcenter = c()
for (i in 1:dim(km_new_centers)[1]){
	print(i)
x_i_dist = apply(dss_ct, 1, function(x) sum((x-km_new_centers[i,])^2))
x_dist_to_newcenter = cbind(x_dist_to_newcenter, x_i_dist)
}

x_dist_to_newcenter_reassign = apply(x_dist_to_newcenter,1,which.min)

png('cCRE_coe.human.heatmap.D.png', width=1200, height=1800)
plot_lim_PD = quantile(as.numeric(as.matrix(dss)),0.99)
breaksList = seq(-plot_lim_PD, plot_lim_PD, by = 0.001)
my_colorbar=colorRampPalette(c('blue', 'white', 'red'))(n = length(breaksList))
col_breaks = c(seq(0, 2000,length=33))
dss_plot = round(dss,3)
dss_plot[dss>plot_lim_PD]=plot_lim_PD
pheatmap(dss_plot[order(x_dist_to_newcenter_reassign),hclust_cor_order], color=my_colorbar, breaks = breaksList, cluster_cols = F,cluster_rows=F, clustering_method = 'complete',annotation_names_row=F,annotation_names_col=TRUE,show_rownames=F,show_colnames=TRUE)
dev.off()


set.seed(2019)
pre_cluster = x_dist_to_newcenter_reassign
for (i in 1:2){
qda_model = qda(round(dss_ct,3), pre_cluster)
fit_cluster_reorder_vec_after_rescue = predict(qda_model, dss_ct)$class
#fit_cluster_reorder_vec[fit_cluster_reorder_vec==0] = fit_cluster_reorder_vec_after_rescue
print(sum(pre_cluster!=fit_cluster_reorder_vec_after_rescue))
if (sum(pre_cluster!=fit_cluster_reorder_vec_after_rescue)==0){break}
pre_cluster = fit_cluster_reorder_vec_after_rescue
print(cbind(table(pre_cluster)))
}

new_id = fit_cluster_reorder_vec_after_rescue
cluster_id_order = c(1:length(unique(fit_cluster_reorder_vec_after_rescue)))[order(-table(fit_cluster_reorder_vec_after_rescue))]
k = 0
for (i in 1:length(cluster_id_order)){
new_id[fit_cluster_reorder_vec_after_rescue==cluster_id_order[i]] = i
}

png('cCRE_coe.human.heatmap.D.qda.png', width=1200, height=1800)
plot_lim_PD = quantile(as.numeric(as.matrix(dss)),0.99)
breaksList = seq(-plot_lim_PD, plot_lim_PD, by = 0.001)
my_colorbar=colorRampPalette(c('blue', 'white', 'red'))(n = length(breaksList))
col_breaks = c(seq(0, 2000,length=33))
dss_plot = dss
dss_plot[dss>plot_lim_PD]=plot_lim_PD
pheatmap(dss_plot[order(new_id),hclust_cor_order], color=my_colorbar, breaks = breaksList, cluster_cols = F,cluster_rows=F, clustering_method = 'complete',annotation_names_row=F,annotation_names_col=TRUE,show_rownames=F,show_colnames=TRUE)
dev.off()


dss_out = cbind(d[,c(1:4)], new_id, ds)
colnames(dss_out)[5] = 'clusterID'

write.table(dss_out, 'S3V2_IDEAS_hg38_ccre2.cCRE.M.notall0.rmallNEU.withid.coe_mat.clusterID.txt', quote=F, row.names=F, col.names=T, sep='\t')
```
