## Supplementary material for "Interspecies regulatory landscapes and elements revealed by novel joint systematic integration of human and mouse blood cell epigenomes": Scripts for IDEAS modeling of epigenetic states jointly in human and mouse: README.md

# Joint_Human_House_IDEAS_State
# Joint_Human_House_IDEAS_State

###### The whole pipeline is in the "RUNME_joint_IDEAS_pipeline_8mk.sh" file
```
##################
### joint human-mouse IDEAS Epigenetic State run
### Human hg38 R1
###### step1: hg38 initial S3V2-IDEAS run (hg38 r1)
cd /storage/home/gzx103/scratch/joint_human_mouse/hg38
qsub run_S3V2_IDEAS_ESMP_hg38_r1.sh

### Mouse mm10 R1
###### step2: mm10 initial S3V2-IDEAS run (mm10 r1)
cd /storage/home/gzx103/scratch/joint_human_mouse/mm10
qsub run_S3V2_IDEAS_ESMP_mm10_r1.sh


### Joint States
###### step3: get joint states
cd /storage/home/gzx103/scratch/joint_human_mouse/joint_states
#     time Rscript get_joint_mouse_human_states.R /storage/home/gzx103/scratch/joint_human_mouse/hg38/S3V2_IDEAS_outputs_hg38/S3V2_IDEAS_hg38_r1_IDEAS_output/S3V2_IDEAS_hg38_r1 /storage/home/gzx103/scratch/joint_human_mouse/mm10/S3V2_IDEAS_outputs_mm10/S3V2_IDEAS_mm10_r1_IDEAS_output/S3V2_IDEAS_mm10_r1 hg38_mm10_joint_states_8mk.55.para 100 50 0.55
time Rscript ~/scratch/joint_human_mouse/S3V2_IDEAS_ESMP/bin/get_joint_mouse_human_states.R /storage/home/gzx103/scratch/joint_human_mouse/hg38/S3V2_IDEAS_outputs_hg38/S3V2_IDEAS_hg38_r1_IDEAS_output1/S3V2_IDEAS_hg38_r1 /storage/home/gzx103/scratch/joint_human_mouse/mm10/S3V2_IDEAS_outputs_mm10/S3V2_IDEAS_mm10_r1_IDEAS_output1/S3V2_IDEAS_mm10_r1 hg38_mm10_joint_states_8mk.52.para 100 50 0.52


### Human hg38 R2
###### step4: rescan hg38 with prior joint states (hg38 r2)
cd /storage/home/gzx103/scratch/joint_human_mouse/hg38/S3V2_IDEAS_outputs_hg38
qsub S3V2_IDEAS_hg38_r2.sh
### step4a: remove heterognenious-state
cd /storage/home/gzx103/scratch/joint_human_mouse/hg38/
time Rscript ~/scratch/joint_human_mouse/S3V2_IDEAS_ESMP/bin/get_state_mean_var.maxnum.R 'S3V2_IDEAS_outputs_hg38/' 'S3V2_IDEAS_hg38_r2' '.m_v.r2.pdf' '_IDEAS_output/'

### Mouse mm10 R2
###### step5: rescan mm10 with prior joint states (mm10 r2)
cd /storage/home/gzx103/scratch/joint_human_mouse/mm10/S3V2_IDEAS_outputs_mm10
qsub S3V2_IDEAS_mm10_r2_withhg38prior.sh
### step5a: remove heterognenious-state
cd /storage/home/gzx103/scratch/joint_human_mouse/mm10/
time Rscript ~/scratch/joint_human_mouse/S3V2_IDEAS_ESMP/bin/get_state_mean_var.maxnum.R 'S3V2_IDEAS_outputs_mm10/' 'S3V2_IDEAS_mm10_r2_withhg38prior' '.m_v.r2.pdf' '_IDEAS_output/'


### final state
# /storage/home/gzx103/scratch/joint_human_mouse/mm10/S3V2_IDEAS_outputs_mm10/S3V2_IDEAS_mm10_r2_withhg38prior_IDEAS_output/S3V2_IDEAS_mm10_r2_withhg38prior.rmh.10.para

### Human hg38 R3
###### step6a: rescan hg38 with prior joint states (hg38 r3)
cd /storage/home/gzx103/scratch/joint_human_mouse/hg38/S3V2_IDEAS_outputs_hg38
###### add feature average signal into the IDEAS input file in the 3rd run
declare -a mk_arr=("ATAC" "CTCF" "H3K27ac" "H3K27me3"  "H3K36me3" "H3K4me1" "H3K4me3" "H3K9me3")
for mk in "${mk_arr[@]}"
do
	cut -f4 $mk'.average_sig.bedgraph.S3V2.ave.bedgraph.NBP.bedgraph' > 'S3V2_IDEAS_hg38_r1_IDEAS_input_NB/'$mk'.average_sig.NBP.txt'
done
###### add average signal into the IDEAS.input file
### e.g.: AVE ATAC /storage/home/gzx103/scratch/joint_human_mouse/hg38/S3V2_IDEAS_outputs_hg38//S3V2_IDEAS_hg38_r1_IDEAS_input_NB/ATAC.average_sig.NBP.txt
qsub S3V2_IDEAS_hg38_r3_withHg38Mm10prior.sh
###### step6a: remeasure the state coverage 
cd /storage/home/gzx103/scratch/joint_human_mouse/hg38/S3V2_IDEAS_outputs_hg38/S3V2_IDEAS_hg38_r3_withHg38Mm10prior_IDEAS_output
time Rscript ../bin/get_modified_para_heatmap.R S3V2_IDEAS_hg38_r3_withHg38Mm10prior.para S3V2_IDEAS_hg38_r3_withHg38Mm10prior.state F ../bin/createGenomeTracks_np.R
### get non all 0 bin bed
time Rscript ~/scratch/joint_human_mouse/S3V2_IDEAS_ESMP/bin/get_notall0_bed.R S3V2_IDEAS_hg38_r3_withHg38Mm10prior.state S3V2_IDEAS_hg38_r3_withHg38Mm10prior.nonall0.bed

### Mouse mm10 R3
###### step6b: rescan mm10-hg38 with prior joint states (mm10 r3)
cd /storage/home/gzx103/scratch/joint_human_mouse/mm10/S3V2_IDEAS_outputs_mm10
###### add feature average signal into the IDEAS input file in the 3rd run
declare -a mk_arr=("ATAC" "CTCF" "H3K27ac" "H3K27me3"  "H3K36me3" "H3K4me1" "H3K4me3" "H3K9me3")
for mk in "${mk_arr[@]}"
do
	cut -f4 $mk'.average_sig.bedgraph.S3V2.ave.bedgraph.NBP.bedgraph' > 'S3V2_IDEAS_hg38_r1_IDEAS_input_NB/'$mk'.average_sig.NBP.txt'
done
###### add average signal into the IDEAS.input file
### e.g.: AVE ATAC /storage/home/gzx103/scratch/joint_human_mouse/hg38/S3V2_IDEAS_outputs_hg38//S3V2_IDEAS_hg38_r1_IDEAS_input_NB/ATAC.average_sig.NBP.txt
qsub S3V2_IDEAS_mm10_r3_withHg38Mm10prior.sh
###### step6b: remeasure the state coverage 
cd /storage/home/gzx103/scratch/joint_human_mouse/hg38/S3V2_IDEAS_outputs_hg38/S3V2_IDEAS_mm10_r3_withHg38Mm10prior_IDEAS_output
time Rscript ../bin/get_modified_para_heatmap.R S3V2_IDEAS_mm10_r3_withHg38Mm10prior.para S3V2_IDEAS_mm10_r3_withHg38Mm10prior.state F ../bin/createGenomeTracks_np.R
### get non all 0 bin bed
time Rscript ~/scratch/joint_human_mouse/S3V2_IDEAS_ESMP/bin/get_notall0_bed.R S3V2_IDEAS_mm10_r3_withHg38Mm10prior.state S3V2_IDEAS_mm10_r3_withHg38Mm10prior.nonall0.bed

###### check R3 mean vs var
cd /storage/home/gzx103/scratch/joint_human_mouse/hg38/
time Rscript ~/scratch/joint_human_mouse/S3V2_IDEAS_ESMP/bin/get_state_mean_var.maxnum.R 'S3V2_IDEAS_outputs_hg38/' 'S3V2_IDEAS_hg38_r3_withHg38Mm10prior' '.m_v.r3.pdf' '_IDEAS_output/'
cd /storage/home/gzx103/scratch/joint_human_mouse/mm10/
time Rscript ~/scratch/joint_human_mouse/S3V2_IDEAS_ESMP/bin/get_state_mean_var.maxnum.R 'S3V2_IDEAS_outputs_mm10/' 'S3V2_IDEAS_mm10_r3_withHg38Mm10prior' '.m_v.r3.pdf' '_IDEAS_output/'


##################
### Human hg38 R0
### step1a: remove heterognenious-state 
cd /storage/home/gzx103/scratch/joint_human_mouse/hg38
time Rscript ~/scratch/joint_human_mouse/S3V2_IDEAS_ESMP/bin/get_state_mean_var.maxnum.R 'S3V2_IDEAS_outputs_hg38/' 'S3V2_IDEAS_hg38_r1' '.m_v.pdf' '_IDEAS_output/'
### step1b: remove heterognenious-state
### step1c: rerun hg38 only S3V2-IDEAS run 
cd /storage/home/gzx103/scratch/joint_human_mouse/hg38/S3V2_IDEAS_outputs_hg38
qsub S3V2_IDEAS_hg38_r1_rmh_rerun.sh
cd /storage/home/gzx103/scratch/joint_human_mouse/hg38/S3V2_IDEAS_outputs_hg38/S3V2_IDEAS_hg38_r1_rmh_rerun_IDEAS_output
time Rscript ../bin/get_modified_para_heatmap.R S3V2_IDEAS_hg38_r1_rmh_rerun.para S3V2_IDEAS_hg38_r1_rmh_rerun.state F ../bin/createGenomeTracks_np.R


##################
### CCRE 
### Use the mm10 ATAC ave signal as the overall reference
### /storage/home/gzx103/scratch/joint_human_mouse/mm10/S3V2_IDEAS_outputs_mm10_CCRE/ATAC.average_sig.bedgraph
# 1st run mm10 
cd /storage/home/gzx103/scratch/joint_human_mouse/mm10
qsub run_S3V2_IDEAS_CCRE_mm10.sh
# 1st run hg38
cd /storage/home/gzx103/scratch/joint_human_mouse/hg38
qsub run_S3V2_IDEAS_CCRE_hg38.sh
# get shared states
cd /storage/home/gzx103/scratch/joint_human_mouse/joint_states
time Rscript get_joint_mouse_human_states.R /storage/home/gzx103/scratch/joint_human_mouse/hg38/S3V2_IDEAS_outputs_hg38_CCRE/S3V2_IDEAS_hg38_ccre_IDEAS_output1/S3V2_IDEAS_hg38_ccre /storage/home/gzx103/scratch/joint_human_mouse/mm10/S3V2_IDEAS_outputs_mm10_CCRE/S3V2_IDEAS_mm10_ccre_IDEAS_output1/S3V2_IDEAS_mm10_ccre hg38_mm10_joint_states_IS.52.1.para 100 50 0.52

# 2nd run mm10 
cd /storage/home/gzx103/scratch/joint_human_mouse/mm10/S3V2_IDEAS_outputs_mm10_CCRE
qsub S3V2_IDEAS_mm10_ccre2.sh
# 2nd run hg38
cd /storage/home/gzx103/scratch/joint_human_mouse/hg38/S3V2_IDEAS_outputs_hg38_CCRE
qsub S3V2_IDEAS_hg38_ccre2.sh

### remove all 0 bins
cd ~/scratch/joint_human_mouse/hg38/S3V2_IDEAS_outputs_hg38_CCRE/S3V2_IDEAS_hg38_ccre2_IDEAS_output
nonall0_bedfile='/storage/home/gzx103/scratch/joint_human_mouse/hg38/S3V2_IDEAS_outputs_hg38/S3V2_IDEAS_hg38_r3_withHg38Mm10prior_IDEAS_output/S3V2_IDEAS_hg38_r3_withHg38Mm10prior.nonall0.bed'
bedtools intersect -a S3V2_IDEAS_hg38_ccre2.cCRE.M.bed -b $nonall0_bedfile -wa -u > S3V2_IDEAS_hg38_ccre2.cCRE.M.notall0.bed
cd ~/scratch/joint_human_mouse/mm10/S3V2_IDEAS_outputs_mm10_CCRE/S3V2_IDEAS_mm10_ccre2_IDEAS_output
nonall0_bedfile='/storage/home/gzx103/scratch/joint_human_mouse/mm10/S3V2_IDEAS_outputs_mm10/S3V2_IDEAS_mm10_r3_withHg38Mm10prior_IDEAS_output/S3V2_IDEAS_mm10_r3_withHg38Mm10prior.nonall0.bed'
bedtools intersect -a S3V2_IDEAS_mm10_ccre2.cCRE.M.bed -b $nonall0_bedfile -wa -u > S3V2_IDEAS_mm10_ccre2.cCRE.M.notall0.bed


##################
###### cp everything to /storage/home/gzx103/scratch/joint_human_mouse/transfer_files/

###### hg38 J25
cd /storage/home/gzx103/scratch/joint_human_mouse/hg38/S3V2_IDEAS_outputs_hg38/S3V2_IDEAS_hg38_r3_withHg38Mm10prior_IDEAS_output
cp -r S3V2_IDEAS_hg38_r3_withHg38Mm10prior_IDEAS_output/S3V2_IDEAS_hg38_r3_withHg38Mm10prior.para.modified.para /storage/home/gzx103/scratch/joint_human_mouse/transfer_files/Epigenetic_State/VBSJ_ES_J25_hg38/
cp -r S3V2_IDEAS_hg38_r3_withHg38Mm10prior_IDEAS_output/S3V2_IDEAS_hg38_r3_withHg38Mm10prior.para.modified.para.pdf /storage/home/gzx103/scratch/joint_human_mouse/transfer_files/Epigenetic_State/VBSJ_ES_J25_hg38/
cp -r S3V2_IDEAS_hg38_r3_withHg38Mm10prior_IDEAS_output/S3V2_IDEAS_hg38_r3_withHg38Mm10prior.state /storage/home/gzx103/scratch/joint_human_mouse/transfer_files/Epigenetic_State/VBSJ_ES_J25_hg38/
cp -r S3V2_IDEAS_hg38_r3_withHg38Mm10prior_IDEAS_output/Tracks /storage/home/gzx103/scratch/joint_human_mouse/transfer_files/Epigenetic_State/VBSJ_ES_J25_hg38/
### bigwig
cp -r S3V2_IDEAS_hg38_r1_bws_RC /storage/home/gzx103/scratch/joint_human_mouse/transfer_files/Epigenetic_State/VBSJ_ES_J25_hg38/
cp -r S3V2_IDEAS_hg38_r1_bws_NBP /storage/home/gzx103/scratch/joint_human_mouse/transfer_files/Epigenetic_State/VBSJ_ES_J25_hg38/
### tmp para
cd /storage/home/gzx103/scratch/joint_human_mouse/hg38/S3V2_IDEAS_outputs_hg38/S3V2_IDEAS_hg38_r1_IDEAS_output1
mkdir 01aa_hg38_r1_tmp_para
cp S3V2_IDEAS_hg38_r1.tmp.*.para 01aa_hg38_r1_tmp_para/
cp -r 01aa_hg38_r1_tmp_para /storage/home/gzx103/scratch/joint_human_mouse/transfer_files/Epigenetic_State/VBSJ_ES_J25_hg38/

### mm10 J25
cd /storage/home/gzx103/scratch/joint_human_mouse/mm10/S3V2_IDEAS_outputs_mm10/
cp -r S3V2_IDEAS_mm10_r3_withHg38Mm10prior_IDEAS_output/S3V2_IDEAS_mm10_r3_withHg38Mm10prior.para.modified.para /storage/home/gzx103/scratch/joint_human_mouse/transfer_files/Epigenetic_State/VBSJ_ES_J25_mm10/
cp -r S3V2_IDEAS_mm10_r3_withHg38Mm10prior_IDEAS_output/S3V2_IDEAS_mm10_r3_withHg38Mm10prior.para.modified.para.pdf /storage/home/gzx103/scratch/joint_human_mouse/transfer_files/Epigenetic_State/VBSJ_ES_J25_mm10/
cp -r S3V2_IDEAS_mm10_r3_withHg38Mm10prior_IDEAS_output/S3V2_IDEAS_mm10_r3_withHg38Mm10prior.state /storage/home/gzx103/scratch/joint_human_mouse/transfer_files/Epigenetic_State/VBSJ_ES_J25_mm10/
cp -r S3V2_IDEAS_mm10_r3_withHg38Mm10prior_IDEAS_output/Tracks /storage/home/gzx103/scratch/joint_human_mouse/transfer_files/Epigenetic_State/VBSJ_ES_J25_mm10/
### bigwig
cp -r S3V2_IDEAS_mm10_r1_bws_RC /storage/home/gzx103/scratch/joint_human_mouse/transfer_files/Epigenetic_State/VBSJ_ES_J25_mm10/
cp -r S3V2_IDEAS_mm10_r1_bws_NBP /storage/home/gzx103/scratch/joint_human_mouse/transfer_files/Epigenetic_State/VBSJ_ES_J25_mm10/
### tmp para
cd /storage/home/gzx103/scratch/joint_human_mouse/mm10/S3V2_IDEAS_outputs_mm10/S3V2_IDEAS_mm10_r1_IDEAS_output1
mkdir 01ba_mm10_r1_tmp_para
cp S3V2_IDEAS_mm10_r1.tmp.*.para 01ba_mm10_r1_tmp_para/
cp -r 01ba_mm10_r1_tmp_para /storage/home/gzx103/scratch/joint_human_mouse/transfer_files/Epigenetic_State/VBSJ_ES_J25_mm10/


### hg38 H29
cd /storage/home/gzx103/scratch/joint_human_mouse/hg38/S3V2_IDEAS_outputs_hg38/S3V2_IDEAS_hg38_r1_rmh_rerun2_IDEAS_output


### hg38 IS & cCRE
cd /storage/home/gzx103/scratch/joint_human_mouse/hg38/S3V2_IDEAS_outputs_hg38_CCRE/S3V2_IDEAS_hg38_ccre2_IDEAS_output
cp S3V2_IDEAS_hg38_ccre2.cCRE.M.notall0.bed /storage/home/gzx103/scratch/joint_human_mouse/transfer_files/cCRE_IS/VBSJ_IS_cCRE_hg38/
cp S3V2_IDEAS_hg38_ccre2.para /storage/home/gzx103/scratch/joint_human_mouse/transfer_files/cCRE_IS/VBSJ_IS_cCRE_hg38/
cp S3V2_IDEAS_hg38_ccre2.pdf /storage/home/gzx103/scratch/joint_human_mouse/transfer_files/cCRE_IS/VBSJ_IS_cCRE_hg38/
cp -r Tracks /storage/home/gzx103/scratch/joint_human_mouse/transfer_files/cCRE_IS/VBSJ_IS_cCRE_hg38/
cp S3V2_IDEAS_hg38_ccre2.*.cCRE.bed /storage/home/gzx103/scratch/joint_human_mouse/transfer_files/cCRE_IS/VBSJ_IS_cCRE_hg38/celltype_pk_bed/

### mm10 IS & cCRE
cd /storage/home/gzx103/scratch/joint_human_mouse/mm10/S3V2_IDEAS_outputs_mm10_CCRE/S3V2_IDEAS_mm10_ccre2_IDEAS_output
cp S3V2_IDEAS_mm10_ccre2.cCRE.M.notall0.bed /storage/home/gzx103/scratch/joint_human_mouse/transfer_files/cCRE_IS/V_IS_cCRE_mm10/
cp S3V2_IDEAS_mm10_ccre2.para /storage/home/gzx103/scratch/joint_human_mouse/transfer_files/cCRE_IS/V_IS_cCRE_mm10/
cp S3V2_IDEAS_mm10_ccre2.pdf /storage/home/gzx103/scratch/joint_human_mouse/transfer_files/cCRE_IS/V_IS_cCRE_mm10/
cp -r Tracks /storage/home/gzx103/scratch/joint_human_mouse/transfer_files/cCRE_IS/V_IS_cCRE_mm10/
cp S3V2_IDEAS_mm10_ccre2.*.cCRE.bed /storage/home/gzx103/scratch/joint_human_mouse/transfer_files/cCRE_IS/V_IS_cCRE_mm10/celltype_pk_bed/

```
