## Supplementary material for "Interspecies regulatory landscapes and elements revealed by novel joint systematic integration of human and mouse blood cell epigenomes": Scripts for IDEAS modeling of epigenetic states jointly in human and mouse: run_S3V2_IDEAS_CCRE_hg38.sh

```
#PBS -l nodes=1:ppn=4
#PBS -l walltime=40:00:00
#PBS -j oe
#PBS -A yzz2_e_g_sc_default
##PBS -A rch8_b_g_bc_default
#PBS -l pmem=60gb

module load bedtools

### required inputs
###### the absolute path to the bin folder
script_dir='/storage/home/gzx103/scratch/joint_human_mouse/S3V2_IDEAS_ESMP/bin/'
###### your output folder
output_dir='/storage/home/gzx103/scratch/joint_human_mouse/hg38/S3V2_IDEAS_outputs_hg38_CCRE/'
###### the absolute path to the your modified "metadata.for_master_peak_calls.txt" file
metadata='/storage/home/gzx103/scratch/joint_human_mouse/hg38/metadata.human.atac.txt'
###### The output name
id_name='S3V2_IDEAS_hg38_ccre'

###### genome
GENOME='hg38'
###### genome size (can be found in the "S3V2_IDEAS_ESMP/genomesize/" folder)
GENOMESIZES='/storage/home/gzx103/scratch/test_S3V2/S3V2_IDEAS_ESMP/genomesize/hg38.chrom.chr1_22XY.sizes'
###### blacklist (can be found in the "S3V2_IDEAS_ESMP/blacklist/" folder)
BLACK='/storage/home/gzx103/scratch/test_S3V2/S3V2_IDEAS_ESMP/blacklist/hg38-blacklist.v2.bed'

###### number of threads in system
###### When the number of threads is too large, the multi-threads in python3 may fail. So it is more stable to keep it below 4. 
threads=4
###### bin size of the signal resolution
bin_size=200
###### email address
email=''

###### other parameters 
get_sigtrack='F'
normalization='T'
get_bw='F'
run_ideas='T'
local_bg_bin=5
cap_sig=16
### User can use the "other_parafile" parameter to incorporate previous epigenetic state model
### We provided two epigenetic state models with 8/7 epigenetic features that can be found in the "prior_ES_models/" folder
other_parafile='F'
IDEAS_track_link='http://your_acess_link_that_can_be_used_for_track_hub_in_genome_browser/'

#[[ -d $output_dir ]] || mkdir $output_dir
cd $output_dir
time python3 $script_dir/S3V2_IDEAS_pipeline.py \
-u $get_sigtrack -v $normalization -y $get_bw -z $run_ideas \
-s $script_dir -o $output_dir -g $GENOME -c $GENOMESIZES -b $BLACK \
-i $metadata -d $id_name -e $email -t $threads -w $IDEAS_track_link -x $other_parafile \
-l $bin_size -n $local_bg_bin -a $cap_sig
```
