## Supplementary material for "Interspecies regulatory landscapes and elements revealed by novel joint systematic integration of human and mouse blood cell epigenomes": Scripts for IDEAS modeling of epigenetic states jointly in human and mouse: readme.hg38.txt

IDEAS OUTPUT FILES:

(A) The IDEAS states generated by joinly using human and mouse data:
(A1) S3V2_IDEAS_hg38_r3_withHg38Mm10prior.state: 
The Epigenetic states and position classes for the human (hg38) hematopoietic system. The Joint epigenetic states were used as prior IDEAS epigenetic state.
	First 4 columns are index, chr, position_st position_ed (position_ed will be the same as position_st if only one position for each window is provided in input) 
	The next N columns are epigenetic states, where N=total number of cell types, including replicates. 
	The last column is the position class label in IDEAS local clustering.

(A2) S3V2_IDEAS_hg38_r3_withHg38Mm10prior.para.modified.para: 
Frequency, sum for each of the joint human and mouse epigenetic states in human (hg38) hematopoietic system.
		The first column in the Frequency of the state
		The next N columns are the sum and variance parameters for each epigenetic states.


(B) The IDEAS states generated by only using human data
(B1) S3V2_IDEAS_hg38_r1_rmh_rerun.state: 
The Epigenetic states and position classes for the human (hg38) hematopoietic system. The Joint epigenetic states were NOT used as prior IDEAS epigenetic state.

(B2) S3V2_IDEAS_hg38_r1_rmh_rerun.para.modified.para: 
Frequency, sum for each of the epigenetic states in human (hg38) hematopoietic system.


(C) The cCRE list in human (hg38) hematopoietic system identified by S3V2-IDEAS IS module. 
S3V2_IDEAS_hg38_ccre.cCRE.M.rmall0.bed


The files for the tracks associated with this data are available from
https://newbadger.bx.psu.edu/downloads/hg38/bbi/ideasJointFeb2021/
