## Supplementary material for "Interspecies regulatory landscapes and elements revealed by novel joint systematic integration of human and mouse blood cell epigenomes": Scripts for IDEAS modeling of epigenetic states jointly in human and mouse: readme.txt

The files for the genome browser:
IDEAS states Tracks
IDEAS states pdf
S3V2 signal (bigwig) Read count
S3V2 signal (bigwig) -log10-pvalue
cCRE list
cell type peaks
IDEAS IS tracks
IDEAS IS states pdf

The files for the usevision.org download page:
IDEAS states file along the genome in all cell types (For RNA prediction)
IDEAS states para
IDEAS states pdf
IDEAS states tmp para (for generating joint states)
cCRE list


A. Epigenetic-State: 
/tmp/May_IDEASruns/Epigenetic_State/Joint-human-mouse-state (25 states)

(A0) Joint IDEAS states:
IDEAS states para file: 
/tmp/May_IDEASruns/Epigenetic_State/VBSJ_ES_J25_hg38/S3V2_IDEAS_hg38_r3_withHg38Mm10prior.para.modified.para.pdf
IDEAS states pdf file: 
/tmp/May_IDEASruns/Epigenetic_State/VBSJ_ES_J25_hg38/S3V2_IDEAS_hg38_r3_withHg38Mm10prior.para.modified.para.pdf

(A1) hg38
IDEAS states Tracks: The IDEAS state state tracks for genome browser in bigbed format
/tmp/May_IDEASruns/Epigenetic_State/VBSJ_ES_J25_hg38/Tracks/
IDEAS states pdf:
/tmp/May_IDEASruns/Epigenetic_State/VBSJ_ES_J25_hg38/S3V2_IDEAS_hg38_r3_withHg38Mm10prior.para.modified.para.pdf
S3V2 signal (bigwig) Read count:
/tmp/May_IDEASruns/Epigenetic_State/VBSJ_ES_J25_hg38/S3V2_IDEAS_hg38_r1_bws_RC/
S3V2 signal (bigwig) -log10-pvalue:
/tmp/May_IDEASruns/Epigenetic_State/VBSJ_ES_J25_hg38/S3V2_IDEAS_hg38_r1_bws_NBP/
IDEAS states file along the genome in all cell types: (For RNA prediction)
/tmp/May_IDEASruns/Epigenetic_State/VBSJ_ES_J25_hg38/S3V2_IDEAS_hg38_r3_withHg38Mm10prior.state
IDEAS states para: 
/tmp/May_IDEASruns/Epigenetic_State/VBSJ_ES_J25_hg38/S3V2_IDEAS_hg38_r3_withHg38Mm10prior.para.modified.para
IDEAS states tmp para: (for generating joint states)
/tmp/May_IDEASruns/Epigenetic_State/VBSJ_ES_J25_hg38/01aa_hg38_r1_tmp_para/

(A2) mm10
IDEAS states Tracks:
/tmp/May_IDEASruns/Epigenetic_State/VBSJ_ES_J25_mm10/Tracks/
IDEAS states pdf:
/tmp/May_IDEASruns/Epigenetic_State/VBSJ_ES_J25_mm10/S3V2_IDEAS_mm10_r3_withHg38Mm10prior.para.modified.para.pdf
S3V2 signal (bigwig) Read count:
/tmp/May_IDEASruns/Epigenetic_State/VBSJ_ES_J25_mm10/S3V2_IDEAS_mm10_r1_bws_RC/
S3V2 signal (bigwig) -log10-pvalue:
/tmp/May_IDEASruns/Epigenetic_State/VBSJ_ES_J25_mm10/S3V2_IDEAS_mm10_r1_bws_NBP/
IDEAS states file along the genome in all cell types: (For RNA prediction)
/tmp/May_IDEASruns/Epigenetic_State/VBSJ_ES_J25_mm10/S3V2_IDEAS_mm10_r3_withHg38Mm10prior.state
IDEAS states para:
/tmp/May_IDEASruns/Epigenetic_State/VBSJ_ES_J25_mm10/S3V2_IDEAS_mm10_r3_withHg38Mm10prior.para.modified.para
IDEAS states tmp para: (for generating joint states)
/tmp/May_IDEASruns/Epigenetic_State/VBSJ_ES_J25_mm10/01aa_mm10_r1_tmp_para/

B. Signal-Intensity-State & cCRE:
/tmp/May_IDEASruns/cCRE_IS/

(B1) hg38
cCRE list:
/tmp/May_IDEASruns/cCRE_IS/VBSJ_IS_cCRE_hg38/S3V2_IDEAS_hg38_ccre2.cCRE.M.notall0.bed
cell type peaks:
/tmp/May_IDEASruns/cCRE_IS/VBSJ_IS_cCRE_hg38/celltype_pk_bed/
IDEAS IS tracks:
/tmp/May_IDEASruns/cCRE_IS/VBSJ_IS_cCRE_hg38/Tracks/
IDEAS IS states pdf:
/tmp/May_IDEASruns/cCRE_IS/VBSJ_IS_cCRE_hg38/S3V2_IDEAS_hg38_ccre2.pdf

(B2) mm10
cCRE list:
/tmp/May_IDEASruns/cCRE_IS/V_IS_cCRE_mm10/S3V2_IDEAS_mm10_ccre2.cCRE.M.notall0.bed
cell type peaks:
/tmp/May_IDEASruns/cCRE_IS/V_IS_cCRE_mm10/celltype_pk_bed/
IDEAS IS tracks:
/tmp/May_IDEASruns/cCRE_IS/V_IS_cCRE_mm10/Tracks/
IDEAS IS states pdf:
/tmp/May_IDEASruns/cCRE_IS/V_IS_cCRE_mm10/S3V2_IDEAS_mm10_ccre2.pdf
